# G-Graph: An interactive genomic graph viewer

**DOI:** 10.1101/803015

**Authors:** Peter A. Andrews, Joan Alexander, Jude Kendall, Michael Wigler

## Abstract

**Motivation:** Effective and efficient exploration of numeric data and annotations as a function of genomic position requires specialized software.

**Results:** We present G-Graph, an interactive genomic scatter plot viewer. G-Graph stacks or tiles multiple data series in one graph using different colors and markers. It displays gene annotation and other metadata, allows easy changes to the appearance of data series, implements stack-based undo functionality, and saves user-selected application views as image and pdf files. G-Graph delivers smooth and rapid scrolling and zooming even for datasets with millions of points and line segments. The primary target user is a researcher examining many copy number profiles to identify potentially deleterious variants. G-Graph runs under Linux, Mac OSX and Windows.

**Availability:** https://github.com/docpaa/mumdex/ or https://mumdex.com/ggraph/

**Contact:** (or)

## 1 Introduction

G-Graph is a free-software (MIT license) desktop application for interactive plotting of numeric data as a function of genomic position. The primary purpose for G-Graph is to enable efficient exploratory analysis of copy number and other datasets by genomic researchers. This requires the ability to quickly scroll and zoom in to and out of regions of interest, to distinguish samples and data types, to change the stacking order of samples, switch between stacked and tiled views, to revert to previous views, to interpret events with an integrated display of gene annotation, and to save selected images.

G-Graph is distributed as part of the MUMdex genome analysis software package (Andrews *et al*., 2016). G-Graph compiles in Linux or Unix with X11, Mac OSX with XQuartz, and Windows with Cygwin. The system requirements for G-Graph installation are a C++11 or later compiler, an X11 development environment, and optionally the ImageMagick convert program to allow image output in png and pdf format.

At the core of G-Graph is a custom-built generic scatterplot graphing module which is designed to be extensible. G-Graph uses this extensibility to incorporate capabilities that are appropriate for genomic analysis. Users with specific requirements can similarly change G-Graph functionality. G-Graph is written in the C++ programming language to permit development at a high-level of abstraction without sacrificing run-time efficiency. G-Graph code wraps low-level details such as bytes, memory locations, fundamental types and library interfaces in higher-level objects representing concepts such as fonts, windows, datasets, graphs, etc.

G-Graph employs the low-level and stable X11 Xlib for visualization without the aid of a widget toolkit such as Tk. This increases G-Graph portability and ease of installation while allowing unique modes of interaction, making its distinctive look and feel invariant under different window managers and operating systems. G-Graph suffers minimal degradation of responsiveness even for remote sessions over slow networks because X11 enables very efficient data display.

## 2 Properties

### 2.1 Launching G-Graph

We illustrate the launching of G-Graph using Figure 1, which shows a megabase portion of the X chromosome from a 500,000 bin copy number analysis of one family from the autism Simons Simplex Collection (Fischbach *et al*., 2010), processed with MUMdex alignment and copy number software. The four horizontal lines in the figure indicate copy number values of 1, 2, 3 and 4. The father (blue markers) and son (yellow) have a copy number value of 1, while the mother (red) and daughter (green) are mostly at a copy number value of 2, but there are also amplifications in the daughter that were inherited from the mother. This figure illustrates the display of both bin ratio data as unconnected points and segmentation profiles as lines for four different samples.

**Figure 1:**
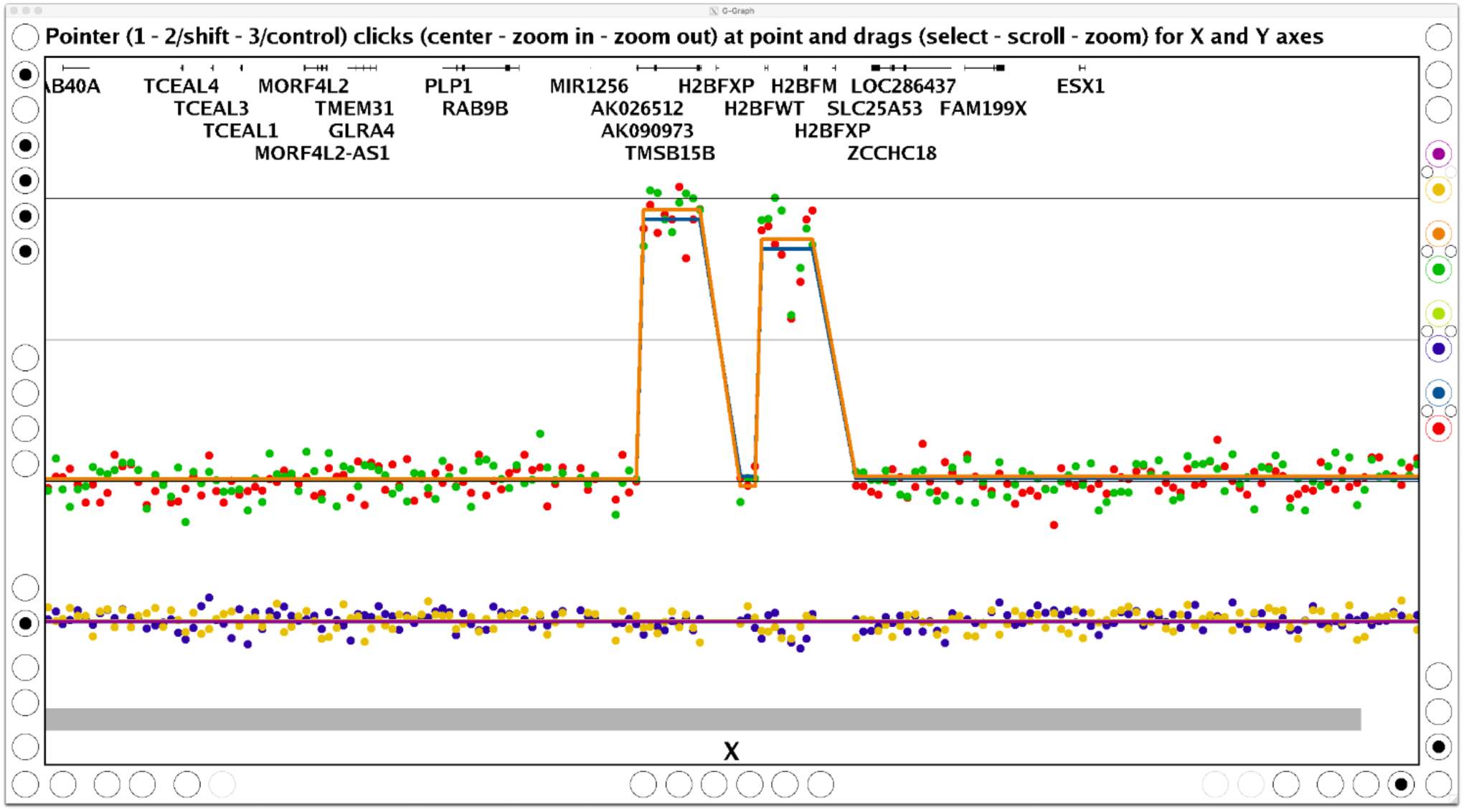
A zoomed in G-Graph application view showing inherited X chromosome amplification in family females (see text for a full description).

The input text data files for Figure 1 are m.txt, f.txt, d.txt and s.txt (corresponding to mother, father, daughter, and son). These files contain, among others, columns with names abspos, ratio, and seg. Input files must be whitespace-separated and tabular in form, with numeric x values in one column and corresponding y values in other columns, in any order. From the directory of the files, the command line for Figure 1 was:

~~~
    **ggraph cn hg19.fa abspos,ratio,seg {m,f,d,s}.txt**
~~~

Here, ggraph is the command name, which is assumed to be in the shell search path. The first argument cn indicates that copy number ratio lines should be drawn and genome information displayed, while hg19.fa indicates the reference genome fasta file used. Next, the x axis absolute genome position and two y axis variables are given as column names. The input files are specified at the end of the command line using shell brace expansion for compactness.

The G-Graph tutorial at https://mumdex.com/ggraph/ explains how to input chromosomal coordinates, how to control the selection of point or line display for each series, how to use keyboard shortcuts for various actions, how to use G-Graph for batch image generation, and other details.

The sample dataset used here, UCSC genome fasta files, plus gene and cytoband definition files for hg19 and hg38 are available for download at https://mumdex.com/ggraph/.

### 2.2 User Interface

The G-Graph user interface (see Figure 1) consists of a central graph region and a number of grouped radio button controls along three borders. At the top border the status line displays a tooltip message for each interface element that the pointer (e.g. mouse, trackpad) hovers over. The controls and status display disappear when the pointer exits the window to provide an uncluttered view of application content. The window can be adjusted to any size possible on the user’s screen.

Clicks and drags of the three pointer buttons are used to center, scroll and zoom the view. Pointer clicks perform immediate centering, zoom ins and zoom outs, while pointer drags perform region selection or smooth scrolling and zooming. In the central graph region these actions affect both axes, while along the borders they affect only the adjacent axis. The shift and control keys are used to emulate the secondary and tertiary pointer button for users without a full-function mouse. The status line shown in Figure 1 describes this pointer action behavior (for the central graph region), which was designed for rapid multiscale exploration.

The radio buttons come in togglable and non-togglable varieties, and are grouped by function. The top right group allows reversion to successively earlier views, saves views in both png and pdf formats, and selects between stacked or tiled views. The colored controls select which data series are displayed and allow color changes, while the adjacent small buttons toggle series group display and advance series stacking order. The bottom right controls modify the appearance of the markers and lines. The bottom left controls perform zoom-outs to the full data range, toggle the display of grid lines and axis labels, and control y axis scale. Scale choices include linear, logarithmic and a new scale smoothly covering all values from 0 to infinity while emphasizing common ploidies (see next section). The top left controls load the G-Graph tutorial and toggle tooltip, coordinate, and genomic metadata display (such as chromosome label, gene, and cytoband). Typing a query into the coordinate radio button also controls genome region displayed by search criteria, such as genome position or gene. The central group of controls along each axis perform discrete jumps and continuous scrolling.

The genic structure and gene names are displayed at the top of the graph region when the view is sufficiently zoomed in. If the user’s pointer hovers over any gene name, the standard description of the gene is displayed in the status line. If a gene name is clicked with the pointer, the user’s web browser is instructed to load the UCSC gene browser (Kent *et al*., 2002) view for the gene. Cytobands are displayed as colored thick horizontal bands at the bottom of the graph region.

### 2.3 Universal Copy Number Scale

The default G-Graph Y axis display scale has been designed to provide a uniform scale smoothly covering copy number values from 0 to infinity, with an emphasis on common copy number values from 0.1 to 30. The utility of this new scale is displayed in Figure 2, which shows 128 copy number profiles, two of which are of breast cancer and the rest are examples of developmental syndromes caused by large chromosomal deletions and duplications.

**Figure 2:**
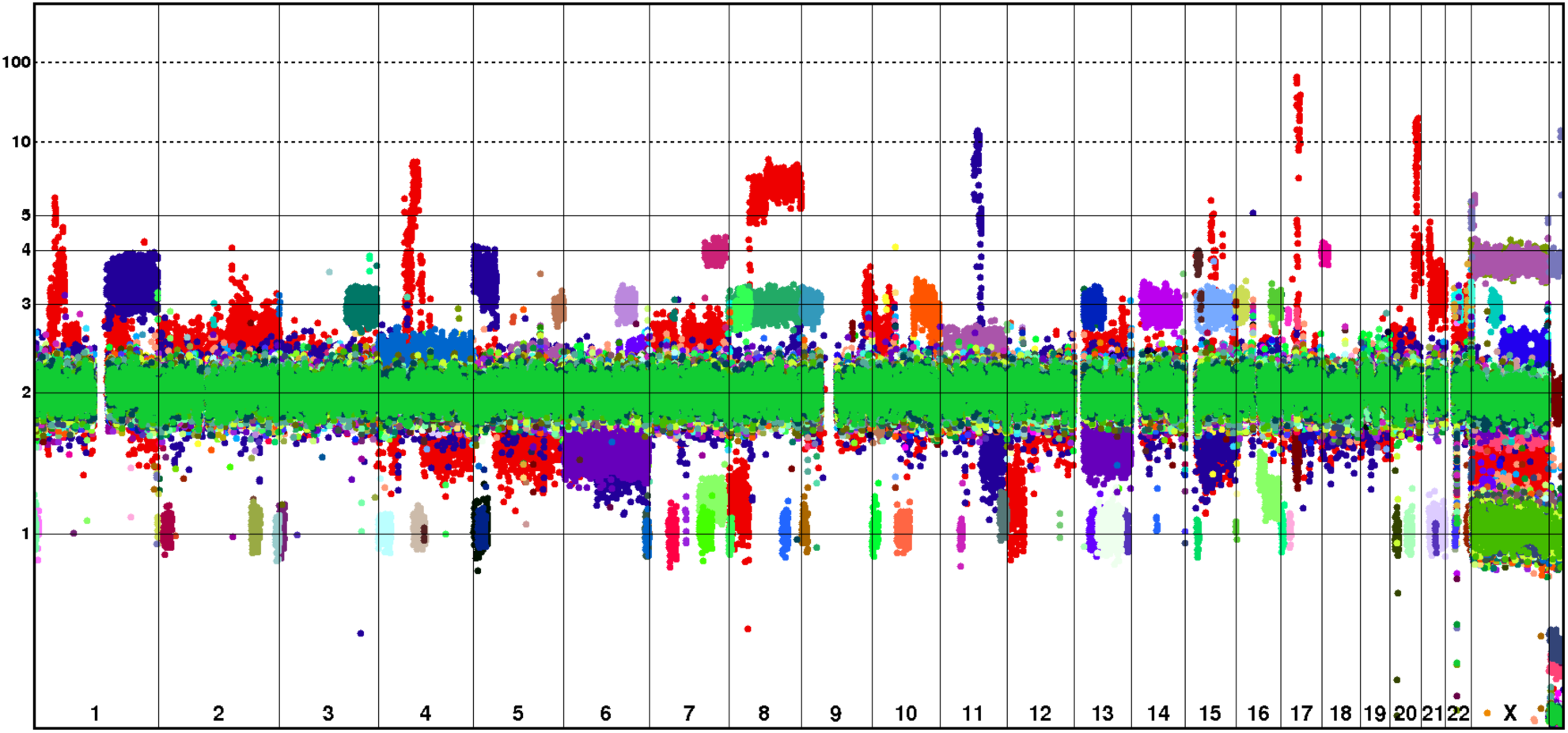
128 samples with large duplications and deletions displayed in one graph, to show the utility of G-Graph’s new universal copy number scale.

The universal copy number scale is defined by a somewhat complicated function, chosen after much experimentation to yield a visually pleasing display scale. The main novelty is to use the arctan function to map all positive values to a finite range. The scale is defined by the following equations:

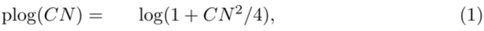

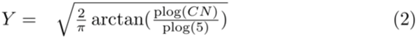

The log, the power of two, the square root and the various constants used are to refine the arctan mapping to produce a useful scale for copy number data that is usually centered at a value of 2, while mapping the range of all nonnegative real values to a domain from 0 to 1. The mapping from copy number to the new scale is shown in Figure 3.

**Figure 3:**
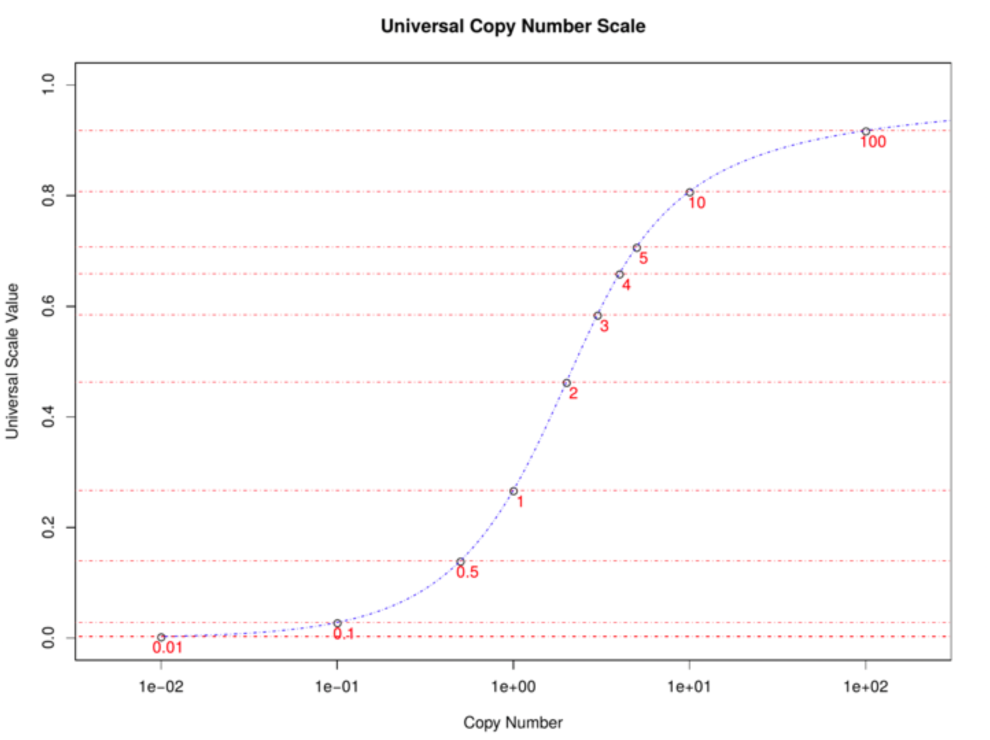
The mapping from copy number on the X-axis to the new universal copy number scale on the Y-axis.

### 2.4 Performance and Functionality Comparisons

On a 2018 mac mini with a 3.2GHz Intel Core i7 processor, it takes 3 seconds for G-Graph to load and display the test dataset of two million points and two million line segments. Once loaded, a zoom action induces a scan over all data plus a redraw of in-frame data, which takes from 3 ms for a megabase-sized region to 500 ms for the entire dataset. This level of performance permits efficient exploratory analysis of large copy number datasets even on midrange desktop computers.

We developed an R interface to G-Graph’s plotting engine so that we could directly compare the speed of native plotting in R (using the plot command with pch=‘.’ to increase plot speed) to G-Graph. For 1 million to 100 million data points already loaded in R memory, display latency from G-Graph launch was 7 to 20 times shorter than for native R plotting.

We also compared G-Graph features and performance with the genomic scatterplot capabilities present in the CNVkit copy number analysis python package (Talevich 2016). The first difference encountered is that CNVkit requires a separate input file to display segmented profiles, while G-Graph reads both ratio and segmented data from the same input file. A second difference is that CNVkit only displays log-ratio data while G-Graph only displays ratio data. A third difference is that CNVkit will only display specific gene names that are listed in input files, and does not display the genic structure or extent of the gene. A fourth difference is that panning and zooming in CNVkit uses standard matplotlib python controls which tend to be more limited and difficult to use than G-Graph’s custom interface. A fifth difference is that CNVkit has the ability to display b allele frequencies, copy number ideograms, and copy number heatmaps, while G-Graph lacks these features. Finally, when viewing a 1 million bin copy number dataset, it takes CNVkit over 16 seconds to output an image from command start, and 14 seconds to update the image after a window resize or zoom when running on a powerful server. For G-Graph, these same functions take less than 4 seconds and 200 ms respectively, on the same server. The conclusion is that G-Graph is superior to CNVkit for interactive copy number profile exploration, especially in light of its many additional features mentioned in earlier sections.

Other copy number visualization packages have similar shortcomings, or do not have comparable features to G-Graph to justify direct comparison. For example, the R copynumber (Nilsen 2012) and DNAcopy (Seshan 2019) packages can generate copy number scatter plots, but the time for redraw after a window resize takes over 8 times the time G-Graph takes, and there is no interactive zoom or scroll function present in these utilities. The IGV (Robinson 2011) copy number display component is only capable of segmented heatmap visualization, and the same is true of several other utilities.

## Supporting information

MUMdex Source Code for G-Graph

Genome Configuration Files (included for completeness - better to use install instructions on website)

Sample Data Files (included for completeness - better to use install instructions on website)

## Funding

This work was supported by a grant from the Simons Foundation (497800, MW).

## Conflict of Interest

none declared.

## Notes

https://mumdex.com/ggraph/

https://github.com/docpaa/mumdex/

