## Supplementary material for "G-Graph: An interactive genomic graph viewer": MUMdex Source Code for G-Graph: cn.h

```
//
// cn.h
//
// Copy Number Utilities
//
// copyright 2016 Peter Andrews
//


#ifndef PAA_CN_H_
#define PAA_CN_H_

#include <algorithm>
#include <fstream>
#include <map>
#include <limits>
#include <memory>
#include <mutex>
#include <numeric>
#include <set>
#include <string>
#include <thread>
#include <utility>
#include <vector>

#include "error.h"
#include "files.h"
#include "genes.h"
#include "lowess.h"
#include "mumdex.h"
#include "psplot.h"
#include "strings.h"
#include "threads.h"

namespace paa {

class PSCNGraph : public PSGraph {
  using PSGraph::PSGraph;

 public:
  PSCNGraph(PSDoc & doc, const Reference & ref_,
            const std::vector<unsigned int> & chromosomes_,
            const std::string & title__ = "") :
      PSGraph{doc, title__ + ";Genome Position (Gb);Copy Number"},
    ref{ref_}, chromosomes{chromosomes_} { }

  PSCNGraph(PSDoc & doc, const Reference & ref_,
            const unsigned int chromosome, const std::string & title__ = "") :
      PSGraph{doc, title__ + ";Chromosome " +
        remove_substring(ref_.name(chromosome), "chr") +
        " Position;Copy Number Ratio"},
    ref{ref_}, chromosomes{chromosome} { }

  virtual void finalize(PSDoc & doc, Bounds bounds) {
    const Bounds saved_range{range_};

    const double scale{pow(bounds.xw() * bounds.yw() /
                           doc.width() / doc.height(), 0.2)};

    // Graph labels, possibly with extra room at top for chromosome labels
    do_labels(doc, bounds, scale,
              chromosomes.size() == 1 ? 0.0 : tick_size() * scale);

    // Chromosome offsets
    const std::vector<unsigned int> offsets{[this]() {
        std::vector<unsigned int> result(chromosomes.size() + 1);
        unsigned int offset{0};
        for (unsigned int ci{0}; ci <= chromosomes.size(); ++ci) {
          result[ci] = offset;
          if (ci == chromosomes.size()) break;
          offset += ref.size(chromosomes[ci]);
        }
        return result;
      }()};

    // Graph range
    this->log_y_ = true;
    const double border{0.02};
    const Bounds min_range{0.0, static_cast<double>(offsets.back()), 0.1, 10.0};
    do_range(border, min_range);

    // Axis label bounds adjustment
    const double tick_size__{tick_size() * scale};
    const double page_size{sqrt(doc.width() * doc.height())};
    const double x_page_scale{pow(bounds.xw() / page_size, 0.5)};
    const double y_page_scale{pow(bounds.yw() / page_size, 0.5)};
    const Axis x_axis{range().xl(), range().xh(), 7 * x_page_scale};
    const Axis y_axis{range().yl(), range().yh(), 2 * y_page_scale,
          this->log_y_};
    const Ticks x_ticks{x_axis.ticks()};
    const Ticks y_ticks{y_axis.ticks()};
    std::ostringstream sample;
    const double high_val{this->log_y_ ?
          pow(10, y_ticks.back().first) : y_ticks.back().first};
    sample << std::setprecision(10) << high_val;
        // y_axis.format(0.99 * range().yh());
    bounds.xl() += 0.75 * sample.str().size() * tick_size__;
    bounds.yl() += 1.3 * tick_size__;

    // Set x y scales
    double scales[2]{bounds.xw() / range().xw(), bounds.yw() / range().yw()};

    // Bounding box
    const double eb{border_width() * scale / 2};
    do_bbox(doc, scale, bounds, eb);

    // Axes and ticks and labels
    const double width1{0.5 * grid_width() * scale};
    const double width2{grid_width() * scale};
    const std::string dash1{minor_dash()};
    const std::string dash2{major_dash()};
    std::ostringstream grid2_out;
    grid2_out << std::setprecision(default_precision)
              << "np " << width2 << " lw " << dash2 << " sd\n";

    std::ostringstream ticks_out;
    ticks_out  << std::setprecision(default_precision)
               << "np bk c " << tick_size__ << " sf "
               << tick_width() << " lw [] 0 sd\n";

    doc << "/glx {" << "0 " << bounds.yw() + 2 * eb << " rl} def "
        << "/gly {" << bounds.xw() + 2 * eb << " 0" << " rl} def "
        << "/tmx {" << "0 " << tick_length() * scale << " rl} def "
        << "/tmy {" << tick_length() * scale << " 0" << " rl} def\n"
        << "np 0.8 0.8 0.8 c " << width1 << " lw " << dash1 << " sd\n";

    unsigned int n_pname{0};
    for (const bool y : {false, true}) {
      for (const std::pair<double, bool> tick : (y ? y_ticks : x_ticks)) {
        std::ostringstream pname_out;
        pname_out << "p" << ++n_pname;
        const std::string pname{pname_out.str()};
        // Saved position command
        doc << "/" << pname << " {"
            << (y ? bounds.xl() - eb :
                (tick.first - range().xl()) * scales[0] + bounds.xl()) << " "
            << (y ? (tick.first - range().yl()) * scales[1] + bounds.yl() :
                bounds.yl() - eb) << " m} def\n";

        // Grid
        if (y || chromosomes.size() == 1) {
          if (tick.second) {
            grid2_out << pname << " gl" << (y ? "y" : "x") << "\n";
          } else {
            doc << pname << " gl" << (y ? "y" : "x") << "\n";
          }
        }

        // Ticks and labels
        bool force_label{false};
        if (y) {
          for (const double force_major_tick :
            {0.02, 0.05, 0.2, 0.5, 2.0, 3.0, 4.0, 5.0, 20.0, 50.0, 200.0}) {
            if (!(pow(10, tick.first) < force_major_tick * 0.999||
                  pow(10, tick.first) > force_major_tick * 1.001)) {
              force_label = true;
            }
          }
        }
       if (tick.second || force_label) {
          ticks_out << pname << " tm" << (y ? "y" : "x") << " "
                    << pname << " (" << std::setprecision(10)
                    << ((y && this->log_y_) ? pow(10, tick.first) :
                        (chromosomes.size() > 1 ?
                         tick.first / 1000000000 : tick.first))
                    << ") " << std::setprecision(default_precision);
          if (y) {
            ticks_out << "jrx " << 0.4 * tick_size__
                      << " sub " << 0.35 * tick_size__;
          } else {
            ticks_out << "jcx " << 1.1 * tick_size__;
          }
          ticks_out << " nrs\n";
        }
      }
    }
    doc << "sp\n"
        << grid2_out.str() << "sp [] 0 sd 0 0 0 c\n";

    // Finalize series
    for (PSSeries * series : children()) {
      series->finalize(doc, bounds, range_, *this);
    }
    doc << "ic\n";
    doc << ticks_out.str() << "sp\n";

    write_text(doc, bounds, scale);

    // Heavy horizontal lines
    doc << "np " << width1 << " lw [] 0 sd ";
    for (const double force_heavy_line : {0.5, 1.0, 2.0, 3.0, 4.0, 5.0}) {
        doc << bounds.xl() - eb << " "
            << (log10(force_heavy_line) - range().yl()) *
            scales[1] + bounds.yl()
            << " m gly sp\n";
    }


    // Vertical chromosome lines and labels at top
    if (chromosomes.size() > 1) {
      doc << tick_size() * 0.8 << " sf\n";
      for (unsigned int ci{0}; ci <= chromosomes.size(); ++ci) {
        doc << (offsets[ci] - range().xl()) * scales[0] + bounds.xl() << " "
            << bounds.yl() - eb << " m glx sp\n";
        if (ci == chromosomes.size()) break;
        const unsigned int c{chromosomes[ci]};
        doc << (offsets[ci] + ref.size(c) / 2 - range().xl()) *
            scales[0] + bounds.xl() << " "
            << bounds.yh() + tick_size() * 0.2 << " m "
            << "(" << remove_substring(ref.name(c), "chr")
            << ") jc s\n";
      }
    }

    // Arbitrary text
    write_text(doc, bounds, scale);

    // Arbitrary ps
    do_ps(doc, scales, bounds, ps_);

    // Restore range
    range_ = saved_range;
  }

 private:
  const Reference & ref;
  std::vector<unsigned int> chromosomes;
};

void add_gaps(PSGraph & graph, const Gaps & gaps, const unsigned int chr) {
  std::ostringstream gap_lines;
  gap_lines << "0 0 1 c [] 0 sd 1 setlinewidth np\n";
  for (const Gap & gap : gaps) {
    if (gap.chr == chr) {
      gap_lines << gap.start << " xc 0 yfc m " << gap.start << " xc 1 yfc l\n";
      gap_lines << gap.stop << " xc 0 yfc m " << gap.stop << " xc 1 yfc l\n";
    }
  }
  gap_lines << "sp";
  graph.ps(gap_lines.str());
}

class CN_abspos {
 public:
  using ChrPos = std::pair<unsigned int, unsigned int>;
  explicit CN_abspos(const Reference & ref,
                     const unsigned int min_size = 40000000) :
      ref_offsets(ref.n_chromosomes() + 1) {
    const unsigned int bad(-1);
    unsigned int offset{0};
    for (unsigned int c{0}; c != ref.n_chromosomes(); ++c) {
      const unsigned int length{ref.size(c)};
      if (length > min_size) {
        chromosomes_.push_back(c);
        ref_offsets[c] = offset;
        cn_offsets.push_back(offset);
        offset += length;
      } else {
        ref_offsets[c] = bad;
      }
    }
    for (unsigned int c{0}; c != ref.n_chromosomes(); ++c) {
      if (ref_offsets[c] == bad) {
        ref_offsets[c] = offset;
      }
    }
    ref_offsets.back() = offset;
    cn_offsets.push_back(offset);
  }
  unsigned int operator()(const unsigned int chromosome,
                          const unsigned int position) const {
    return ref_offsets[chromosome] + position;
  }
  unsigned int ref_offset(const unsigned int chr) const {
    return ref_offsets[chr];
  }
  unsigned int cn_offset(const unsigned int chr) const {
    return cn_offsets[chr];
  }
  unsigned int ref_size(const unsigned int chr) const {
    return ref_offsets[chr + 1] - ref_offsets[chr];
  }
  ChrPos chrpos(const unsigned int abspos) const {
    const unsigned int c(static_cast<unsigned int>(
        upper_bound(cn_offsets.begin(), cn_offsets.end(),
                    abspos) - cn_offsets.begin() - 1));
    if (c >= chromosomes_.size()) {
      const unsigned int chr{chromosomes_.back()};
      return {chr, ref_size(chr)};
    } else {
      return {chromosomes_[c], abspos - cn_offset(c)};
    }
  }
  const std::vector<unsigned int> & chromosomes() const {
    return chromosomes_;
  }
  const std::vector<unsigned int> & offsets() const { return cn_offsets; }

  unsigned int bad_abspos() const { return ref_offsets.back(); }
  unsigned int n_positions() const { return ref_offsets.back(); }

 private:
  std::vector<unsigned int> ref_offsets{};
  std::vector<unsigned int> cn_offsets{};
  std::vector<unsigned int> chromosomes_{};
};

class RefCN : public RefPlus {
 public:
  explicit RefCN(const std::string & fasta_file_name) :
      RefPlus{fasta_file_name}, cn_abspos{*this} { }
  ~RefCN() {}

  const CN_abspos cn_abspos;
};

class Bin {
 public:
  Bin(const unsigned int chromosome__,
      const unsigned int start_position__,
      const unsigned int abspos__,
      const unsigned int length__,
      const double gc__,
      const double map__,
      const double norm__,
      const double corr__,
      const bool bad__) :
      chromosome_{chromosome__},
    start_position_{start_position__},
    abspos_start_{abspos__},
    abspos_{abspos_start_ + length__ / 2},
    length_{length__},
    gc_{gc__},
    map_{map__},
    norm_{norm__},
    corr_{corr__},
    bad_{bad__} { }

  unsigned int chromosome() const { return chromosome_; }
  unsigned int start_position() const { return start_position_; }
  unsigned int stop_position() const { return start_position_ + length_; }
  unsigned int chr_position() const { return start_position_ + length_ / 2; }
  unsigned int abspos() const { return abspos_; }
  unsigned int abspos_start() const { return abspos_start_; }
  unsigned int abspos_stop() const { return abspos_start_ + length_; }
  unsigned int length() const { return length_; }
  double gc() const { return gc_; }
  double map() const { return map_; }
  double norm() const { return norm_; }
  double corr() const { return corr_; }
  bool bad() const { return bad_; }
  void set_bad() {
    bad_ = true;
  }
  void adjust_corr(const double average) {
    corr_ /= average;
  }

  bool operator<(const PosInfo & pos) const {
    if (chromosome_ == pos.chr) {
      return start_position_ < pos.pos;
    } else {
      return chromosome_ < pos.chr;
    }
  }

  bool operator<(const unsigned int pos) const {
    return abspos_start_ < pos;
  }

  void output(std::ostream & out, const Reference & ref,
              const bool set_bad_ = false) const {
    out << ref.name(chromosome_) << " "
        << start_position_ << " "
        << length_ << " "
        << gc_ << " "
        << map_ << " "
        << norm_ << " "
        << (bad_ || set_bad_)
        << "\n";
  }

 private:
  unsigned int chromosome_;
  unsigned int start_position_;
  unsigned int abspos_start_;
  unsigned int abspos_;
  unsigned int length_;
  double gc_;
  double map_;
  double norm_;
  double corr_;
  bool bad_;
};

inline bool operator>(const PosInfo & pos, const Bin & bin) {
  if (bin.chromosome() == pos.chr) {
    return bin.start_position() < pos.pos;
  } else {
    return bin.chromosome() < pos.chr;
  }
}

inline bool operator<(const PosInfo & pos, const Bin & bin) {
  if (bin.chromosome() == pos.chr) {
    return bin.start_position() > pos.pos;
  } else {
    return bin.chromosome() > pos.chr;
  }
}

inline bool operator<(const unsigned int pos, const Bin & bin) {
  return pos < bin.abspos_start();
}

std::vector<Bin> load_bins(const std::string & bins_name,
                           const Reference & ref,
                           const bool cut = false,
                           const bool mask = false) {
  std::ifstream bins_file{bins_name.c_str()};
  if (!bins_file) throw Error("Could not open bins file") << bins_name;
  const CN_abspos cn_abspos{ref};
  const ChromosomeIndexLookup lookup{ref};
  std::vector<Bin> bins_;
  std::string chromosome_name;
  unsigned int start_position;
  unsigned int length;
  double gc;
  double map;
  bool bad{false};
  double corr;
  double average_corr{0.0};
  std::vector<unsigned int> lengths;
  std::vector<double> corrs;
  while (bins_file >> chromosome_name >> start_position
         >> length >> gc >> map >> corr >> bad) {
    const unsigned int norm = corr;
    const unsigned int chromosome{lookup[chromosome_name]};
    const unsigned int abspos{cn_abspos(chromosome, start_position)};
    bins_.emplace_back(chromosome, start_position, abspos, length,
                       gc, map, norm, corr, bad);
    average_corr += corr;
    corrs.push_back(corr);
    lengths.push_back(length);
  }
  if (!bins_.size()) throw Error("No bins loaded");
  sort(lengths.begin(), lengths.end());
  sort(corrs.begin(), corrs.end());
  const unsigned int median_length{lengths[lengths.size() / 2]};
  const double median_corr{corrs[corrs.size() / 2]};
  average_corr /= bins_.size();
  unsigned int n_bad{0};
  for (Bin & bin : bins_) {
    bin.adjust_corr(median_corr);
    if (cut) {
      if (bin.map() > 150) {
        bin.set_bad();
      }
      if (bin.gc() < 0.3) {
        bin.set_bad();
      }
      if (bin.length() < median_length / 2) {
        bin.set_bad();
      }
      if (bin.length() < median_length / 1.3 ||
        bin.length() > median_length * 1.3) {
        // bin.set_bad();
      }
      if (bin.corr() < median_corr / 1.2 || bin.corr() > median_corr * 1.1) {
        // bin.set_bad();
      }
      if (bin.corr() > average_corr * 1.2) {
        // bin.set_bad();
      }
    }
    if (bin.bad()) {
      ++n_bad;
    }
  }

  // Unstable intervals near centromeres, telomeres - HG19 ONLY
  std::map<std::string,
      std::vector<std::pair<unsigned int, unsigned int>>> bad_zones{
    {"1", { {0, 864000}, {120560000, 149810000} }},
    {"2", { {87090000, 98340000}}},
    {"3", { {90300000, 93580000}, {197913000, 198022430}}},
    {"4", { {0, 43800}, {49080000, 52710000}, {190781000, 191154276}}},
    {"5", { {46310000, 49568000}, {68820000, 70810000}}},
    {"6", { {57680000, 61990000}}},
    {"7", { {57500000, 62060000}}},
    {"8", { {6945000, 8111000}, {11859000, 12572000}, {43741000, 46895000}}},
    {"9", { {0, 232000}, {38450000, 71066000}}},
    {"10", { {38410000, 43010000}, {135432700, 135534747}}},
    {"11", { {48360000, 55040000}, {89550000, 89851000}}},
    {"12", { {34856000, 38225000}}},
    {"13", { {0, 19540000}}},
    {"14", { {0, 20540000}}},
    {"15", { {0, 22960000}, {102395000, 102531392}}},
    {"16", { {35120000, 46610000}}},
    {"17", { {22180000, 25350000}, {44370000, 44810000}}},
    {"18", { {14102000, 18566000}}},
    {"19", { {24540000, 28110000}}},
    {"20", { {25666000, 29898000}}},
    {"21", { {0, 15424000}}},
    {"22", { {0, 17103000}, {51174000, 51304566}}},
    {"X", { {58330000, 61932000}}},
    {"Y", { {9739000, 13994000}, {29340000, 59373566}}}};

  const unsigned int x_chr{ref.find_x_chromosome()};
  if (mask && ref.size(x_chr) != 155270560)
    throw Error("Code designed for hg19 only - need to change") << mask;

  const std::vector<unsigned char> masked_bins{
    [&ref, &bins_, &bad_zones, mask]() {
      std::vector<unsigned char> result(bins_.size());
      if (!mask) return result;
      unsigned int b{0};
      unsigned int n{0};
      for (unsigned int c{0}; c != ref.n_chromosomes(); ++c) {
        const std::string name{ref.name(c)};
        const std::string name_no_chr{remove_substring(name, "chr")};
        if (bad_zones.count(name_no_chr)) {
          const std::vector<std::pair<unsigned int, unsigned int>> &
              zones{bad_zones[name_no_chr]};
          for (const std::pair<unsigned int, unsigned int> zone : zones) {
            while (b + 1 != bins_.size() && bins_[b].chromosome() < c) ++b;
            if (bins_[b].chromosome() > c)
              throw Error("Unexpected chromosome ordering");
            while (b + 1 != bins_.size() &&
                   bins_[b].chromosome() == c &&
                   bins_[b].stop_position() <= zone.first) ++b;
            while (bins_[b].chromosome() == c &&
                   bins_[b].start_position() < zone.second) {
              if (0)
                std::cerr << "Marked bad " << ref.name(bins_[b].chromosome())
                          << " " << bins_[b].start_position()
                          << " " << bins_[b].stop_position() << std::endl;
              result[b] = true;
              ++n;
              if (b + 1 == bins_.size()) break;
              ++b;
            }
          }
        }
      }
      if (0)
        std::cerr << n << " bins marked bad or " << 1.0 * n / bins_.size()
                  << std::endl;
      return result;
    }()};

  for (unsigned int b{0}; b != bins_.size(); ++b) {
    if (masked_bins[b]) bins_[b].set_bad();
  }

  if (0)
    std::cout << "Loaded " << bins_.size() << " bins, with "
              << n_bad << " bad" << std::endl;
  return bins_;
}

void plot_bins(const std::vector<Bin> & bins, const std::string & out_name) {
  if (0) {
    std::cout << bins.size() << std::endl;
  }
  PSDoc ps{out_name};

  PSGraph counts_vs_pos_graph{ps, ";Absolute Position;Bin Count"};
  counts_vs_pos_graph.log_y(true);
  const Marker small_red_marker{paa::circle(), 0.1, "1 0 0", 1, true};
  PSXYSeries counts_vs_pos_series{counts_vs_pos_graph, small_red_marker};

  PSGraph length_vs_pos_graph{ps, ";Absolute Position;Bin Length"};
  length_vs_pos_graph.log_y(true);
  PSXYSeries length_vs_pos_series{length_vs_pos_graph, small_red_marker};

  PSGraph counts_vs_length_graph{ps, ";Bin Count;Bin Length"};
  counts_vs_length_graph.log_y(true);
  PSXYSeries counts_vs_length_series{counts_vs_length_graph, small_red_marker};
}

unsigned int default_min_length{20};
unsigned int default_min_excess{20};
unsigned int default_max_mappability{50};
bool default_use_midpoint{false};

// Temporary function used during testing only
inline void set_cn_parameters() {
  default_min_excess = 10;
  default_max_mappability = 100;
  default_use_midpoint = false;
  return;
  const std::string pwd{getenv("PWD")};
  unsigned int parsed{0};
  const size_t ex{pwd.find("-ex")};
  if (ex != std::string::npos) {
    std::istringstream in{pwd.substr(ex + 3).c_str()};
    in >> default_min_excess;
    if (in) ++parsed;
  }
  const size_t mm{pwd.find("-mm")};
  if (mm != std::string::npos) {
    std::istringstream in{pwd.substr(mm + 3).c_str()};
    in >> default_max_mappability;
    if (in) ++parsed;
  }
  const size_t mid{pwd.find("-midpoint")};
  if (mid != std::string::npos) {
    default_use_midpoint = true;
    ++parsed;
  }
  if (parsed < 2) throw Error("Problem parsing PWD") << pwd << parsed;
  std::cout << "CN Paremeters: "
            << "min excess " << default_min_excess << " "
            << "max_mappability " << default_max_mappability << " "
            << "use midpoint " << default_use_midpoint
            << std::endl;
}

template <class MUMDEX, class MAPPABILITY>
inline unsigned int pair_cn_abspos(const MUMDEX & mumdex,
                                   const MAPPABILITY & mappability,
                                   const CN_abspos & cn_abspos,
                                   const uint64_t pair_index) {
  const MUM * cn_mum{mumdex.cn_MUM(pair_index, mappability,
                                   default_min_length,
                                   default_min_excess,
                                   default_max_mappability)};
  if (cn_mum == nullptr) return cn_abspos.bad_abspos();
  const MUM mum{*cn_mum};
  if (default_use_midpoint) {
    return cn_abspos(mum.chromosome(), mum.position0() + mum.length() / 2);
  } else {
    return cn_abspos(mum.chromosome(), mum.position0());
  }
}

template <template <class ...> class VECTOR>
class FinestBinsT {
  using Vector = VECTOR<unsigned int>;

 public:
  explicit FinestBinsT(const Reference & ref_) :
      ref{ref_}, cn_abspos{ref},
    counts{cn_abspos.n_positions() + 3} { }
  FinestBinsT(const Reference & ref_, const unsigned int nsamples,
              const unsigned int nx, unsigned int ny) :
      FinestBinsT{ref_} {
    n_samples(nsamples);
    n_x(nx);
    n_y(ny);
  }
  FinestBinsT(const Reference & ref_,
              const std::string & file_name,
              const bool mapped,
              const bool readahead = false) :
      ref{ref_}, cn_abspos{ref}, counts{file_name, mapped, readahead} {
        if (counts.size() != cn_abspos.n_positions() + 3)
          throw Error("FinestBins counts size mismatch");
      }
  FinestBinsT(const Reference & ref_,
              const std::string & file_name) :
      ref{ref_}, cn_abspos{ref}, counts{file_name} {
        if (counts.size() != cn_abspos.n_positions() + 3)
          throw Error("FinestBins counts size mismatch");
      }
  void add(const MUMdex & mumdex, const Mappability & map,
           const unsigned int nx, unsigned int ny) {
    n_samples(n_samples() + 1);
    n_x(n_x() + nx);
    n_y(n_y() + ny);
    for (uint64_t p{0}; p < mumdex.n_pairs(); ++p) {
      const unsigned int abspos{pair_cn_abspos(mumdex, map, cn_abspos, p)};
      if (abspos >= cn_abspos.bad_abspos()) continue;
      ++counts[abspos];
    }
  }
  template <class MUMDEX>
  void add(const MUMDEX & mumdex, const Mappability & map,
           const unsigned int nx, const unsigned int ny,
           const unsigned int n_threads) {
    n_samples(n_samples() + 1);
    n_x(n_x() + nx);
    n_y(n_y() + ny);

    ThreadPool pool{n_threads};
    ThreadPool::Results<void> results;
    const uint64_t block_size{
      std::max(static_cast<uint64_t>(1000), mumdex.n_pairs() / n_threads / 10)};

    const uint64_t mutex_block{16000};
    std::vector<std::mutex> mutexes(counts.size() / mutex_block + 2);

    for (uint64_t b{0}; b < mumdex.n_pairs(); b += block_size) {
      pool.run(results, [n_threads, &mutexes, &mumdex, &map,
                         this, b, block_size] () {
          for (uint64_t p{b}; p < std::min(b + block_size, mumdex.n_pairs());
               ++p) {
            const unsigned int abspos{pair_cn_abspos(mumdex, map,
                                                     cn_abspos, p)};
            if (abspos >= cn_abspos.bad_abspos()) continue;
            if (n_threads > 1) {
              std::lock_guard<std::mutex> lock{mutexes[abspos / mutex_block]};
              ++counts[abspos];
            } else {
              ++counts[abspos];
            }
          }
        });
    }

    // Progress progress{results.size(), 0.01, "Binning"};
    while (results.size()) {
      results.get();
      // progress();
    }
  }
  void add(const FinestBinsT & other) {
    if (counts.size() != other.counts.size())
      throw Error("FinestBins other counts size mismatch");
    for (unsigned int p{0}; p != counts.size(); ++p) {
      counts[p] += other.counts[p];
    }
  }
  void save(const std::string & file_name) const {
    counts.save(file_name);
  }
  unsigned int n_x() const { return counts[counts.size() - 2]; }
  unsigned int n_y() const { return counts[counts.size() - 1]; }
  unsigned int n_samples() const { return counts[counts.size() - 3]; }
  FinestBinsT & n_x(const unsigned int val) {
    counts[counts.size() - 2] = val;
    return *this;
  }
  FinestBinsT & n_y(const unsigned int val) {
    counts[counts.size() - 1] = val;
    return *this;
  }
  FinestBinsT & n_samples(const unsigned int val) {
    counts[counts.size() - 3] = val;
    return *this;
  }
  unsigned int size() const {
    return static_cast<unsigned int>(counts.size() - 3);
  }
  unsigned int operator[](const unsigned int i) const { return counts[i]; }
  unsigned int & operator[](const unsigned int i) { return counts[i]; }

  using ChrBins = std::pair<unsigned int, std::vector<unsigned int> >;
  using VII = typename Vector::const_iterator;
  ChrBins get_chr_bins(const VII counts_origin,
                       const VII counts_begin,
                       const VII counts_end,
                       const unsigned int n_bins,
                       const unsigned int cpb_) const {
    ChrBins result{cpb_, std::vector<unsigned int>(n_bins)};
    unsigned int & cpb{result.first};
    std::vector<unsigned int> & bin_starts{result.second};
    std::vector<unsigned int> bin_counts(n_bins);
    std::vector<unsigned int> saved_starts;
    std::vector<unsigned int> saved_counts;
    bool excess{false};
    while (true) {
      // std::cout << " " << cpb << std::flush;
      bin_counts.assign(n_bins, 0);
      unsigned int b{0};
      bin_starts[0] = static_cast<unsigned int>(counts_begin - counts_origin);
      for (VII c{counts_begin}; c != counts_end; ++c) {
        bin_counts[b] += *c;
        if (bin_counts[b] >= cpb && b + 1 != n_bins) {
          ++b;
          bin_starts[b] = static_cast<unsigned int>(c + 1 - counts_origin);
        }
      }
      if (excess || bin_counts.back() >= cpb) {
        // have excess now or previously
        if (bin_counts.back() >= cpb) {
          // have excess now
          excess = true;
          saved_starts = bin_starts;
          saved_counts = bin_counts;
          ++cpb;
        } else {
          // reached deficit again
          bin_starts = saved_starts;
          --cpb;
          break;
        }
      } else {
        // deficit
        cpb = std::min(cpb * sqrt(1.0 * b / n_bins), cpb - 5.0);
      }
    }
    auto minmax = std::minmax_element(saved_counts.begin(),
                                      saved_counts.end() - 1);
    std::cout << " min " << *minmax.first
              << " max " << *minmax.second
              << " last " << saved_counts.back();
    return result;
  }

  void bin(const unsigned int n_bins, const std::string & out_name,
           const double xc, const double yc) const {
    // Get chromosome and genome total counts
    std::cout << "Getting chromosome counts" << std::endl;
    const std::vector<unsigned int> & chromosomes{cn_abspos.chromosomes()};
    std::vector<uint64_t> chromosome_totals(chromosomes.size());
    std::vector<uint64_t> corr_chromosome_totals(chromosomes.size());
    double corr_total_count{0.0};
    std::vector<unsigned int> is_x(chromosomes.size());
    std::vector<unsigned int> is_y(chromosomes.size());
    for (unsigned int c{0}; c != chromosomes.size(); ++c) {
      for (unsigned int b{cn_abspos.cn_offset(c)};
           b != cn_abspos.cn_offset(c + 1); ++b) {
        chromosome_totals[c] += counts[b];
        corr_chromosome_totals[c] += counts[b];
      }
      if (ref.name(chromosomes[c]).find('X') != std::string::npos) {
        corr_chromosome_totals[c] *= xc;
        is_x[c] = 1;
      } else if (ref.name(chromosomes[c]).find('Y') != std::string::npos) {
        corr_chromosome_totals[c] *= yc;
        is_y[c] = 1;
      }
      corr_total_count += corr_chromosome_totals[c];
      std::cout << ref.name(chromosomes[c])
                << " " << chromosome_totals[c]
                << " " << corr_chromosome_totals[c]
                << " " << corr_total_count
                << " " << is_x[c]
                << " " << is_y[c]
                << std::endl;
    }

    // Allocate bins to chromosomes
    std::cout << "Allocating bins to chromosomes" << std::endl;
    std::vector<unsigned int> chromosome_n_bins(chromosomes.size());
    unsigned int n_bins_used{0};
    for (unsigned int c{0}; c != chromosomes.size(); ++c) {
      const double this_n_bins{corr_chromosome_totals[c] /
            corr_total_count * n_bins};
      chromosome_n_bins[c] = this_n_bins;
      n_bins_used += this_n_bins;
    }
    if (n_bins_used > n_bins) throw Error("Unexpected too many bins used");
    while (n_bins_used < n_bins) {
      unsigned int best_choice{0};
      double best_per_bin{0.0};
      for (unsigned int c{0}; c != chromosomes.size(); ++c) {
        const double n_per_bin{
          corr_chromosome_totals[c] / (chromosome_n_bins[c] + 0.0)};
        if (n_per_bin > best_per_bin) {
          best_per_bin = n_per_bin;
          best_choice = c;
        }
      }
      ++chromosome_n_bins[best_choice];
      ++n_bins_used;
    }

    // Bin each chromosome
    std::cout << "Setting chromosome bin boundaries" << std::endl;
    std::vector<ChrBins> chr_bins(chromosomes.size());
    for (unsigned int c{0}; c != chromosomes.size(); ++c) {
      std::cout << "Chromosome " << ref.name(chromosomes[c])
                << " bins " << chromosome_n_bins[c]
                << " cpbs" << std::flush;
      chr_bins[c] = get_chr_bins(counts.begin(),
                                 counts.begin() + cn_abspos.cn_offset(c),
                                 counts.begin() + cn_abspos.cn_offset(c + 1),
                                 chromosome_n_bins[c],
                                 static_cast<unsigned int>(
                                     1.0 * chromosome_totals[c] /
                                     chromosome_n_bins[c]));
      std::cout << std::endl;
    }
    std::cout << "Swapping bins between chromosomes" << std::endl;
    std::set<std::vector<unsigned int> > tried_configs;
    // const unsigned int max_swap{5};
    unsigned int n_swaps{0};
    for (const unsigned int to_swap : {16, 8, 4, 2, 1}) {
      // for (unsigned int to_swap{0}; to_swap != 0;) {
      //    for (unsigned int nsr{0}; nsr != max_swap; ++nsr) {
      //  const unsigned int to_swap{max_swap - nsr};
      std::cout << "Swapping " << to_swap << " bins" << std::endl;
      while (true) {
        // Get cpbs under different scenarios of adding and removing bins
        std::vector<double> cpbs;
        std::vector<double> cpbs_m;
        std::vector<double> cpbs_p;
        for (unsigned int c{0}; c != chromosomes.size(); ++c) {
          const double corr{is_x[c] ? xc : (is_y[c] ? yc : 1.0)};
          const unsigned int cpb(chr_bins[c].first * corr);
          const unsigned int chr_n_bins{chromosome_n_bins[c]};
          cpbs.push_back(cpb);
          cpbs_m.push_back(cpb * chr_n_bins / (chr_n_bins - to_swap));
          cpbs_p.push_back(cpb * chr_n_bins / (chr_n_bins + to_swap));
        }

        // Find best switch to raise lowest cpb
        auto min_cpb = min_element(cpbs.begin(), cpbs.end());
        const unsigned int to_remove_bin{static_cast<unsigned int>(
            min_cpb - cpbs.begin())};
        const double to_remove_corr{is_x[to_remove_bin] ?
              xc : (is_y[to_remove_bin] ? yc : 1.0)};
        auto max_cpb_p = max_element(cpbs_p.begin(), cpbs_p.end());
        const unsigned int to_add_bin{static_cast<unsigned int>(
            max_cpb_p - cpbs_p.begin())};
        const double to_add_corr{is_x[to_add_bin] ?
              xc : (is_y[to_add_bin] ? yc : 1.0)};

        // Is best switch good?
        bool swapped{false};
        if (to_remove_bin != to_add_bin &&
            *min_cpb < *max_cpb_p) {
          chromosome_n_bins[to_remove_bin] -= to_swap;
          chromosome_n_bins[to_add_bin] += to_swap;
          // Check for already tried configuration first!
          if (tried_configs.count(chromosome_n_bins)) {
            chromosome_n_bins[to_remove_bin] += to_swap;
            chromosome_n_bins[to_add_bin] -= to_swap;
          } else {
            auto minmax = minmax_element(cpbs.begin(), cpbs.end());
            std::cout << "Swap " << n_swaps << " range "
                      << *minmax.second - *minmax.first
                      << " so swap " << to_swap
                      << " bin" << (to_swap == 1 ? " " : "s ")
                      << ref.name(chromosomes[to_remove_bin])
                      << " --> "
                      << ref.name(chromosomes[to_add_bin]) << std::endl;
            tried_configs.insert(chromosome_n_bins);
            swapped = true;
            chr_bins[to_remove_bin].first = cpbs_m[to_remove_bin] /
                to_remove_corr;
            chr_bins[to_add_bin].first = cpbs_p[to_add_bin] / to_add_corr;
            ++n_swaps;
            for (const unsigned int c : { to_remove_bin, to_add_bin }) {
              std::cout << "Chromosome " << ref.name(chromosomes[c])
                        << " bins " << chromosome_n_bins[c]
                        << " cpbs" << std::flush;
              chr_bins[c] = get_chr_bins(
                  counts.begin(),
                  counts.begin() + cn_abspos.cn_offset(c),
                  counts.begin() + cn_abspos.cn_offset(c + 1),
                  chromosome_n_bins[c],
                  chr_bins[c].first);
              std::cout << std::endl;
            }
          }
        }
        if (!swapped) {
          break;
        }
      }
    }
    std::cout << "Did " << n_swaps << " chromosome bin swaps" << std::endl;

    // Even out and then join bin starts
    std::vector<unsigned int> bin_starts(n_bins + 1);
    std::vector<unsigned int> bin_counts(n_bins);
    std::vector<unsigned int> chr_starts(chromosomes.size() + 1);
    chr_starts[chromosomes.size()] = n_bins;
    unsigned int bs{0};
    uint64_t n_moves{0};
    for (unsigned int c{0}; c != chromosomes.size(); ++c) {
      chr_starts[c] = bs;
      std::cout << "Even out chromosome " << ref.name(chromosomes[c])
                << std::endl;
      std::vector<unsigned int> chr_bin_starts(chr_bins[c].second);
      std::vector<unsigned int> chr_bin_counts(chr_bin_starts.size());
      chr_bin_starts.push_back(c + 1 == chromosomes.size() ?
                               cn_abspos.n_positions() :
                               chr_bins[c + 1].second[0]);

      // Calculate bin counts
      for (unsigned int b{0}; b != chr_bin_counts.size(); ++b) {
        chr_bin_counts[b] = std::accumulate(
            counts.begin() + chr_bin_starts[b],
            counts.begin() + chr_bin_starts[b + 1], 0U);
      }
      // Even out
      for (unsigned int rb{0}; rb != chr_bin_counts.size() - 1; ++rb) {
        const unsigned int b{static_cast<unsigned int>(
            chr_bin_counts.size() - rb - 1)};
        while (true) {
          bool modified = false;
          bool not_modified = false;
          for (unsigned int u{b}; u != 0; --u) {
            const unsigned int l{u - 1};
            const uint64_t uc{chr_bin_counts[u]};
            const uint64_t lc{chr_bin_counts[l]};
            const uint64_t ufc{counts[chr_bin_starts[u]]};
            if (2 * ufc + lc < uc) {
              chr_bin_counts[l] += ufc;
              chr_bin_counts[u] -= ufc;
              ++chr_bin_starts[u];
              ++n_moves;
              modified = true;
            } else {
              not_modified = true;
            }
            if (not_modified) break;
          }
          if (!modified) break;
        }
      }

      // Join chromosome bins to one list
      for (unsigned int b{0}; b != chr_bin_counts.size(); ++b) {
        bin_starts[bs] = chr_bin_starts[b];
        bin_counts[bs] = chr_bin_counts[b];
        ++bs;
      }
    }
    bin_starts[bs] = cn_abspos.n_positions();
    std::cout << "Moved counts " << n_moves << " times" << std::endl;

    // Report stats
    std::cout << "Stats" << std::endl;
    unsigned int min_cpb(-1);
    unsigned int max_cpb{0};
    unsigned int total_bins{0};
    for (unsigned int c{0}; c != chromosomes.size(); ++c) {
      const double corr{is_x[c] ? xc : (is_y[c] ? yc : 1.0)};
      auto minmax = minmax_element(
          bin_counts.begin() + chr_starts[c],
          bin_counts.begin() + chr_starts[c + 1]);
      const unsigned int corr_cpb{
        static_cast<unsigned int>(chr_bins[c].first * corr)};
      std::cout << ref.name(chromosomes[c]) << " "
                << "bins " << chromosome_n_bins[c] << " "
                << "counts " << chr_bins[c].first << " "
                << "min " << *minmax.first << " "
                << "max " << *minmax.second << " "
                << "corr " << corr_cpb
                << std::endl;
      total_bins += chromosome_n_bins[c];
      if (min_cpb > corr_cpb) min_cpb = corr_cpb;
      if (max_cpb < corr_cpb) max_cpb = corr_cpb;
    }
    std::cout << "Total bins " << total_bins << std::endl;
    std::cout << "CPB range " << max_cpb - min_cpb << " "
              << "min " << min_cpb << " "
              << "max " << max_cpb << std::endl;

    // Output bin file
    const Mappability mappability{ref};
    std::cout << "Output bin file" << std::endl;
    std::ofstream bins_out{
      (out_name + "." + std::to_string(n_bins) + ".txt").c_str()};
    unsigned int c{0};
    for (unsigned int b{0}; b != bin_counts.size(); ++b) {
      const unsigned int p{bin_starts[b]};
      if (p >= cn_abspos.cn_offset(c + 1)) {
        ++c;
      }
      const double corr{is_x[c] ? xc : (is_y[c] ? yc : 1.0)};
      const unsigned int chr_bin_start{bin_starts[b] - cn_abspos.cn_offset(c)};
      const unsigned int length{bin_starts[b + 1] - bin_starts[b]};
      bins_out << ref.name(chromosomes[c]) << " "
               << chr_bin_start << " "
               << length << " "
               << get_gc_content(chromosomes[c],
                                 chr_bin_start,
                                 chr_bin_start + length) << " "
               << get_mappability(mappability,
                                  chromosomes[c],
                                  chr_bin_start,
                                  chr_bin_start + length) << " "
               << bin_counts[b] * corr << " 0\n";
    }
  }

  double get_gc_content(const unsigned int chromosome,
                        const unsigned int start,
                        const unsigned int stop) const {
    const unsigned int edge{25};
    const unsigned int low{edge < start ? start - edge : 0};
    const unsigned int high{edge + stop < ref.size(chromosome) ? stop + edge :
          ref.size(chromosome)};
    unsigned int gc{0};
    for (unsigned int b{low}; b != high; ++b) {
      if (ref[chromosome][b] == 'G' || ref[chromosome][b] == 'C') {
        ++gc;
      }
    }
    return 1.0 * gc / (high - low);
  }

  double get_mappability(const Mappability & mappability,
                         const unsigned int chromosome,
                         const unsigned int start,
                         const unsigned int stop) const {
    const unsigned int edge{25};
    const unsigned int low{edge < start ? start - edge : 0};
    const unsigned int high{edge + stop < ref.size(chromosome) ? stop + edge :
          ref.size(chromosome)};
    double map{0};
    for (unsigned int b{low}; b != high; ++b) {
      const unsigned int abspos(ref.abspos(chromosome, b));
      map += mappability.low(abspos) + mappability.high(abspos);
    }
    return 0.5 * map / (high - low);
  }

 private:
  const Reference & ref;
  const CN_abspos cn_abspos;
  Vector counts;
};
using FinestBins = FinestBinsT<FlexVector>;
using FileFinestBins = FinestBinsT<FileVector>;

void copy_number(const Reference & ref,
                 const std::vector<Bin> bins,
                 const std::vector<unsigned int> & raw_counts,
                 const std::string & title,
                 const bool create_pdf = false,
                 const bool minimal = false,
                 const double alpha = 0.05,
                 const double undo = 1.0,
                 const unsigned int minw = 3) {
  std::cout << "Running copy number analysis with "
            << bins.size() << " bins" << std::endl;

  // Normalize counts
  std::cout << "Normalizing Counts" << std::endl;
  const std::vector<double> norm_counts{[&bins, &raw_counts]() {
      std::vector<double> result(raw_counts.size());
      for (unsigned int b{0}; b != bins.size(); ++b) {
        result[b] = raw_counts[b] / bins[b].corr();
      }
      return result;
    }()};

  // List of good bins
  std::cout << "Selecting good bins" << std::endl;
  const std::vector<unsigned int> good_bins{[&bins]() {
      std::vector<unsigned int> result;
      for (unsigned int b{0}; b != bins.size(); ++b) {
        const Bin & bin{bins[b]};
        if (bin.bad()) continue;
        result.push_back(b);
      }
      return result;
    }()};

  // Reference and CN chromosomes
  std::cout << "Getting chromosome lists" << std::endl;
  const std::vector<unsigned int> & chrs{ref.chromosomes()};
  const std::vector<unsigned int> used_chrs{[&bins, &good_bins]() {
      std::vector<unsigned int> result;
      for (const unsigned int b : good_bins) {
        const Bin & bin{bins[b]};
        if (result.empty() || result.back() != bin.chromosome()) {
          result.push_back(bin.chromosome());
        }
      }
      return result;
    }()};

  // Bins need to be sorted by GC
  std::cout << "Ordering bins by gc content" << std::endl;
  const std::vector<unsigned int> gc_indexes{[&bins, &good_bins]() {
      std::vector<unsigned int> result{good_bins};
      sort(result.begin(), result.end(),
           [&bins](const unsigned int lhs, const unsigned int rhs) {
             return bins[lhs].gc() < bins[rhs].gc();
           });
      return result;
    }()};

  // Input data for GC lowess: gc, counts
  std::cout << "Preparing data for GC lowess" << std::endl;
  const std::vector<double> gc_gc{[&gc_indexes, &bins]() {
      std::vector<double> result;
      result.reserve(gc_indexes.size());
      for (const unsigned int b : gc_indexes) {
        result.push_back(bins[b].gc());
      }
      return result;
    }()};
  const std::vector<double> gc_counts{[&gc_indexes, &norm_counts]() {
      std::vector<double> result;
      result.reserve(gc_indexes.size());
      for (const unsigned int b : gc_indexes) {
        result.push_back(norm_counts[b]);
      }
      return result;
    }()};

  std::cout << "Running Lowess for GC correction" << std::endl;
  const std::vector<double> gc_smoothed{[&gc_gc, &gc_counts]() {
      std::vector<double> result(gc_gc.size());
      std::vector<double> gc_resid_weights(gc_gc.size());
      std::vector<double> gc_weights(gc_gc.size());
      CppLowess::TemplatedLowess<std::vector<double>, double> Lowess;
      Lowess.lowess(gc_gc, gc_counts, 0.1, 5, 0.01,
                    result, gc_resid_weights, gc_weights);
      return result;
    }()};

  // Run GC Lowess and return gc corrected ratios
  const std::vector<double> gc_corr_ratios{
    [&gc_indexes, &gc_counts, &gc_smoothed]() {
      // Need inverse gc ordering to get ratios in genome order
      std::cout << "Getting inverse GC ordering" << std::endl;
      const std::vector<unsigned int> gc_inverse_indexes{[&gc_indexes]() {
          std::vector<unsigned int> result;
          for (unsigned int g{0}; g != gc_indexes.size(); ++g) {
            result.push_back(g);
          }
          sort(result.begin(), result.end(),
               [&gc_indexes](const unsigned int lhs, const unsigned int rhs) {
                 return gc_indexes[lhs] < gc_indexes[rhs];
               });
          return result;
        }()};

      // Prepare ratios applying correction factor and un-sorting by GC.
      std::cout << "Calculating GC Lowess-corrected ratios" << std::endl;
      std::vector<double> result;
      result.reserve(gc_counts.size());
      for (const unsigned int g : gc_inverse_indexes) {
        result.push_back(std::max(0.01, gc_counts[g] / gc_smoothed[g]));
      }
      return result;
    }()};

  // Base name for output files
  std::string out_name{title.size() ? title : std::string("copy_number")};
  replace_inplace(out_name, ' ', '_');

  // Output data for segmentation
  std::cout << "Output data for segmentation" << std::endl;
  std::ofstream ratios_out{out_name + ".ratios.txt"};
  ratios_out << "chr\tabspos\tratio\n";
  for (unsigned int b{0}; b != good_bins.size(); ++b) {
    const Bin & bin{bins[good_bins[b]]};
    ratios_out << bin.chromosome() << "\t"
               << bin.abspos() << "\t"
               << gc_corr_ratios[b] << "\n";
  }
  ratios_out.close();

  // Write R code to a local file
  const std::string r_code{R"foo(library("DNAcopy")
cbs.segment <- function(sample, alpha, nperm, undo.SD, min.width) {
	data <- read.table(paste(sample, "ratios.txt", sep="."), header=T) 
	set.seed(25) 
	CNA.object <- CNA(log(data$ratio, base=2), data$chr, data$abspos,
                          data.type="logratio", sampleid=sample) 
	smoothed.CNA.object <- smooth.CNA(CNA.object) 
	segment.smoothed.CNA.object <- segment(smoothed.CNA.object,
          alpha=alpha, nperm=nperm, undo.splits="sdundo", undo.SD=undo.SD,
          min.width=min.width) 
	Short <- segment.smoothed.CNA.object[[2]]
	m <- matrix(data=0, nrow=nrow(data), ncol=1)	
	prevEnd <- 0
	for (i in 1:nrow(Short)) {
		Start <- prevEnd + 1
		End <- prevEnd + Short$num.mark[i]
                Short$seg.mean[i] <- 2^Short$seg.mean[i]
		m[Start:End, 1] <- Short$seg.mean[i]
		prevEnd = End
	}
	data$seg_ratio <- m[, 1]
	Grid <- seq(1.5, 5.5, by=0.05)
	Outer <- data$seg_ratio %o% Grid
	OuterRound <- round(Outer)
	OuterDiff <- (Outer - OuterRound) ^ 2
	OuterColsums <- colSums(OuterDiff, na.rm = FALSE, dims = 1)
	Multiplier <- Grid[which.min(OuterColsums)]
	data$quantal_ratio <- data$ratio * Multiplier
	data$seg_quantal_ratio <- data$seg_ratio * Multiplier
        data$chr <- NULL
        data$abspos <- NULL
        data$ratio <- NULL
	write.table(data, sep="\t", file=paste(sample, "_segmented.txt",
          sep=""), quote=F, row.names=F) 
}
# Parse command line - no spaces in arguments
args=(commandArgs(TRUE))
for (i in 1:length(args)) {
    eval(parse(text=args[i]))
}
if (!exists("sample")) {
   stop("missing command line argument: sample")
}
# alpha 0.05 undo.sd 1.0 good for normal profile
# alpha 0.02 undo.sd 0.5 good for cancer)foo" +
std::string("\ncbs.segment(sample=sample, alpha=") +
    std::to_string(alpha) + ", nperm=1000, undo.SD=" + std::to_string(undo) +
    ", min.width=" + std::to_string(minw) + ")"};

  // Output R code
  const std::string r_name{out_name + "_cbs.r"};
  std::ofstream r_file{r_name.c_str()};
  if (!r_file) throw Error("Problem writing R code in") << r_name;
  r_file << r_code << std::endl;
  r_file.close();

  try {
    // Calculate segmented results in R
    std::cout << "Performing Segmentation" << std::endl;
    if (system(("R CMD BATCH --slave --no-restore --no-save '--args sample=\"" +
                out_name + "\"' " + r_name + " > /dev/null 2>&1").c_str()) ==
        -1) {
      throw Error("Segmentation in R has failed");
    }
    const std::string segmented_name{out_name + "_segmented.txt"};
    std::ifstream seg_in{segmented_name.c_str()};
    seg_in.ignore(10000, '\n');  // skip header
    if (!seg_in) {
      throw Error("Could not open segmented results file") << segmented_name;
    }

    // Create plots
    std::cout << "Creating plots" << std::endl;
    const std::string light{"0 0 0"};
    const std::string dark{"1 0 0"};
    const Marker light_circle_marker{
      paa::circle(), 0.3, light, 0.2, true, light};
    const Marker dark_circle_marker{paa::circle(), 0.3, dark, 0.2, false};
    const Marker small_red_marker{paa::circle(), 0.1, "1 0 0", 1, true};
    const Marker small_blue_marker{paa::circle(), 0.1, "0 0 1", 1, true};

    // Copy number plots
    PSDoc profiles_ps{out_name + "_profiles"};
    profiles_ps.pdf(create_pdf);

    // Regular copy number plots
    PSCNGraph ratio_graph{profiles_ps, ref, used_chrs, title};
    PSXYSeries ratio_series{ratio_graph, light_circle_marker};
    PSXYSeries segmented_ratio_series{ratio_graph, dark_circle_marker};
    segmented_ratio_series.do_lines(true);

    PSCNGraph quantal_graph{profiles_ps, ref, used_chrs, title};
    PSXYSeries quantal_series{quantal_graph, light_circle_marker};
    PSXYSeries segmented_quantal_series{quantal_graph, dark_circle_marker};
    segmented_quantal_series.do_lines(true);

    const bool do_chr_graphs{false};
#define NORM 1
#if NORM
    // Histograms
    PSGraph ratio_hist_graph{profiles_ps, "Ratio results;Relative CN value;N",
          Bounds{0, 2.5}};
    PSHSeries<double, unsigned int> ratio_hist{ratio_hist_graph, 100};
    PSGraph seg_ratio_hist_graph{profiles_ps,
          "Segmented ratio results;Relative CN value;N", Bounds{0.0, 2.0}};
    PSHSeries<double, unsigned int> seg_ratio_hist{
      seg_ratio_hist_graph, 100};

    // Regular copy number plots by chromosome
    std::vector<std::unique_ptr<PSCNGraph>> chr_ratio_graphs(chrs.size());
    std::vector<std::unique_ptr<PSXYSeries>> chr_ratio_series(chrs.size());
    std::vector<std::unique_ptr<PSXYSeries>> chr_segmented_ratio_series(
        chrs.size());
    if (do_chr_graphs) {
      for (const unsigned int chr : used_chrs) {
        chr_ratio_graphs[chr] = std::make_unique<PSCNGraph>(
            profiles_ps, ref, chr, title);
        chr_ratio_series[chr] = std::make_unique<PSXYSeries>(
            *chr_ratio_graphs[chr], light_circle_marker);
        chr_segmented_ratio_series[chr] = std::make_unique<PSXYSeries>(
            *chr_ratio_graphs[chr], dark_circle_marker);
        chr_segmented_ratio_series[chr]->do_lines(true);
      }
    }
#endif
#define QUANTAL 0
#if QUANTAL
    // Histograms
    PSGraph quantal_hist_graph{profiles_ps, "Quantal results;CN value;N",
          Bounds{0, 5}};
    PSHSeries<double, unsigned int> quantal_hist{quantal_hist_graph, 100};
    PSGraph seg_quantal_hist_graph{profiles_ps,
          "Segmented quantal results;CN value;N", Bounds{0.0, 4.0}};
    PSHSeries<double, unsigned int> seg_quantal_hist{
      seg_quantal_hist_graph, 1000};

    // Quantal graphs by chromosome
    std::vector<std::unique_ptr<PSCNGraph>> chr_quantal_graphs(chrs.size());
    std::vector<std::unique_ptr<PSXYSeries>> chr_quantal_series(chrs.size());
    std::vector<std::unique_ptr<PSXYSeries>> chr_segmented_quantal_series(
        chrs.size());
    if (do_chr_graphs) {
      for (const unsigned int chr : used_chrs) {
        chr_quantal_graphs[chr] = std::make_unique<PSCNGraph>(
            profiles_ps, ref, chr, title);
        chr_quantal_series[chr] = std::make_unique<PSXYSeries>(
            *chr_quantal_graphs[chr], light_circle_marker);
        chr_segmented_quantal_series[chr] = std::make_unique<PSXYSeries>(
            *chr_quantal_graphs[chr], dark_circle_marker);
        chr_segmented_quantal_series[chr]->do_lines(true);
      }
    }
#endif

    // PSDoc counts_ps{out_name + "_counts"};
#if 0
    // Counts vs Position
    counts_ps.pdf(create_pdf);
    PSGraph counts_vs_pos_graph{counts_ps,
          ";Absolute Position;Normalized Bin Count"};
    counts_vs_pos_graph.log_y(true);
    PSXYSeries counts_vs_pos_series_bad{counts_vs_pos_graph, small_blue_marker};
    PSXYSeries counts_vs_pos_series{counts_vs_pos_graph, small_red_marker};
#endif

    // Counts vs GC, and Lowess correction factor
    PSGraph counts_vs_gc_graph{profiles_ps,
          ";GC content;Normalized Bin Count"};
    counts_vs_gc_graph.log_y(true);
    PSXYSeries counts_vs_gc_series_bad{counts_vs_gc_graph, small_blue_marker};
    PSXYSeries counts_vs_gc_series{counts_vs_gc_graph, small_red_marker};
    Marker lowess_marker{paa::circle(), 0.3, "0 1 0", 1, true};
    PSXYSeries counts_vs_gc_lowess_series{counts_vs_gc_graph, lowess_marker};

#if 0
    // Counts vs mappability
    PSGraph counts_vs_map_graph{counts_ps,
          ";Average Mappability Length;Normalized Bin Count"};
    counts_vs_map_graph.log_y(true);
    PSXYSeries counts_vs_map_series_bad{counts_vs_map_graph, small_blue_marker};
    PSXYSeries counts_vs_map_series{counts_vs_map_graph, small_red_marker};

    // Counts vs bin length
    PSGraph counts_vs_length_graph{
      counts_ps, ";Bin Length;Normalized Bin Count"};
    counts_vs_length_graph.log_x(true).log_y(true);
    PSXYSeries counts_vs_length_series_bad{
      counts_vs_length_graph, small_blue_marker};
    PSXYSeries counts_vs_length_series{
      counts_vs_length_graph, small_red_marker};
#endif

    // Read in segmented results to plot them and output results
    std::cout << "Read segmented results and fill plots and output"
              << std::endl;
    std::ofstream results{(out_name + "_results.txt").c_str()};
    if (!results) throw Error("Problem opening CN results file");
    if (minimal) {
      results << "chr\tstart\tstop\tchrpos\tabspos\t"
              << "count\tncount\tratio\tseg_ratio\n";
    } else {
      results << "chr\tstart\tstop\tchrpos\tabspos\tlength\t"
              << "norm\tcorr\tgc\tmap\tcount\tncount\t"
              << "ratio\tseg_ratio\tquantal\tseg_quantal\n";
    }
    for (unsigned int gb{0}; gb != good_bins.size(); ++gb) {
      const unsigned int b{good_bins[gb]};
      const Bin & bin{bins[b]};
      const unsigned int chr{bin.chromosome()};
      const unsigned int abspos{bin.abspos()};
      const unsigned int chrpos{bin.chr_position()};
      const double ratio{gc_corr_ratios[gb]};
      double segmented_ratio;
      double quantal_ratio;
      double segmented_quantal_ratio;
      seg_in >> segmented_ratio >> quantal_ratio >> segmented_quantal_ratio;
      if (!seg_in) throw Error("Problem parsing file") << segmented_name;

      // Quantal profiles
      quantal_series.add_point(abspos, quantal_ratio);
      segmented_quantal_series.add_point(abspos, segmented_quantal_ratio);
#if QUANTAL
      quantal_hist.add_point(quantal_ratio);
      seg_quantal_hist.add_point(segmented_quantal_ratio);
      if (do_chr_graphs) {
        chr_quantal_series[chr]->add_point(chrpos, quantal_ratio);
        chr_segmented_quantal_series[chr]->add_point(
            chrpos, segmented_quantal_ratio);
      }
#endif
#if NORM
      // Regular profiles
      ratio_series.add_point(abspos, ratio);
      segmented_ratio_series.add_point(abspos, segmented_ratio);
      ratio_hist.add_point(ratio);
      seg_ratio_hist.add_point(segmented_ratio);

      if (do_chr_graphs) {
        chr_ratio_series[chr]->add_point(chrpos, ratio);
        chr_segmented_ratio_series[chr]->add_point(chrpos, segmented_ratio);
      }
#endif

      // Counts, gc, mappability, length
      const double count{norm_counts[b]};
      const double clipped_count{std::max(0.5, count)};
      // counts_vs_pos_series.add_point(abspos, clipped_count);
      counts_vs_gc_series.add_point(bin.gc(), clipped_count);
      counts_vs_gc_lowess_series.add_point(gc_gc[gb], gc_smoothed[gb]);
      // counts_vs_map_series.add_point(bin.map(), clipped_count);
      // counts_vs_length_series.add_point(bin.length(), clipped_count);

      // Output results
      if (minimal) {
        results << ref.name(chr) << "\t"
                << bin.start_position() << "\t"
                << bin.stop_position() << "\t"
                << chrpos << "\t"
                << abspos << "\t"
                << raw_counts[b] << "\t"
                << count << "\t"
                << ratio << "\t"
                << segmented_ratio << "\n";
      } else {
        results << ref.name(chr) << "\t"
                << bin.start_position() << "\t"
                << bin.stop_position() << "\t"
                << chrpos << "\t"
                << abspos << "\t"
                << bin.length() << "\t"
                << bin.norm() << "\t"
                << bin.corr() << "\t"
                << bin.gc() << "\t"
                << bin.map() << "\t"
                << raw_counts[b] << "\t"
                << count << "\t"
                << ratio << "\t"
                << segmented_ratio << "\t"
                << quantal_ratio << "\t"
                << segmented_quantal_ratio << "\n";
      }
    }

    // List of bad bins
    std::cout << "Selecting bad bins" << std::endl;
    const std::vector<unsigned int> bad_bins{[&bins]() {
        std::vector<unsigned int> result;
        for (unsigned int b{0}; b != bins.size(); ++b) {
          const Bin & bin{bins[b]};
          if (!bin.bad()) continue;
          result.push_back(b);
        }
        return result;
      }()};

    // Plot data for bad bins
    std::cout << "Plot bad bin data" << std::endl;
    if (bad_bins.size()) {
      for (const unsigned int b : bad_bins) {
        const Bin & bin{bins[b]};
        // const unsigned int abspos{bin.abspos()};
        const double count{norm_counts[b]};
        const double clipped_count{std::max(0.5, count)};
        // counts_vs_pos_series_bad.add_point(abspos, clipped_count);
        counts_vs_gc_series_bad.add_point(bin.gc(), clipped_count);
        // counts_vs_map_series_bad.add_point(bin.map(), clipped_count);
        // counts_vs_length_series_bad.add_point(bin.length(), clipped_count);
      }
    }

    // Remove R temporary files
    if (system(("rm " + r_name).c_str()) == -1) {
      std::cerr << "Problem removing cbs.r file" << std::endl;
    }
    if (system((std::string("rm ") + out_name + ".ratios.txt").c_str()) == -1) {
      std::cerr << "Problem removing ratios.txt file" << std::endl;
    }
    if (system((std::string("rm ") + segmented_name).c_str()) == -1) {
      std::cerr << "Problem removing segmented file" << std::endl;
    }

    std::cout << "Creating ps and pdf plot output files" << std::endl;
  } catch (Error & e) {
    std::cerr << "paa::Error:" << std::endl;
    std::cerr << e.what() << std::endl;
    std::cerr << "Skipping this binning due to segmentation failure"
              << std::endl;
  }
}

class CN_Bin {
 public:
  CN_Bin(const unsigned int count__,
         const double norm_count__,
         const double ratio__,
         const double seg_ratio__,
         const double quantal__,
         const double seg_quantal__) :
      count_{count__}, norm_count_{norm_count__},
    ratio_{ratio__}, seg_ratio_{seg_ratio__},
    quantal_{quantal__}, seg_quantal_{seg_quantal__} { }

  unsigned int count() const { return count_; }
  double norm_count() const { return norm_count_; }
  double ratio() const { return ratio_; }
  double seg_ratio() const { return seg_ratio_; }
  double quantal() const { return quantal_; }
  double seg_quantal() const { return seg_quantal_; }

 private:
  unsigned int count_;
  double norm_count_;
  double ratio_;
  double seg_ratio_;
  double quantal_;
  double seg_quantal_;
};

class CN_Bins {
 public:
  CN_Bins(const CN_Bins &) = delete;
  CN_Bins(CN_Bins &&) = default;
  CN_Bins & operator=(const CN_Bins &) = delete;
  CN_Bins & operator=(CN_Bins &&) = default;
  explicit CN_Bins(const std::string & file_name) {
    // Open file
    std::ifstream input{file_name.c_str()};
    if (!input) throw Error("Problem opening bin results file") << file_name;

    // Determine column structure
    std::string line;
    getline(input, line);
    std::istringstream header_stream{line.c_str()};
    const unsigned int n_columns{[&header_stream]() {
        unsigned int result{0};
        std::string col;
        while (header_stream >> col) ++result;
        return result;
      }()};

    // Read file
    unsigned int count;
    double norm_count;
    double ratio;
    double seg_ratio;
    double quantal{0};
    double seg_quantal{0};
    std::string dummy;
    while (input >> dummy >> dummy >> dummy >> dummy >> dummy
           >> count >> norm_count >> ratio >> seg_ratio) {
      if (n_columns == 11) input >> quantal >> seg_quantal;
      if (!input) throw Error("Parse error for bin results file") << file_name;
      bins.emplace_back(count, norm_count, ratio, seg_ratio,
                        quantal, seg_quantal);
    }
  }

  uint64_t size() const { return bins.size(); }
  const CN_Bin & operator[](const unsigned int b) const { return bins[b]; }
  std::vector<CN_Bin>::const_iterator begin() const { return bins.begin(); }
  std::vector<CN_Bin>::const_iterator end() const { return bins.end(); }

 private:
  std::vector<CN_Bin> bins{};
};

struct Segment {
  unsigned int chr{0};
  unsigned int startpos{0};
  unsigned int stoppos{0};
  unsigned int start{0};
  unsigned int stop{0};
  unsigned int n_bins{0};
  double count{0};
  double cn{0.0};
  unsigned int expected{0};
  unsigned int int_cn{0};
  std::string type{""};
  double score{0.0};
  static std::vector<Segment> load(const std::string & file_name,
                                   const ChromosomeIndexLookup & chr_lookup) {
    std::ifstream file{file_name.c_str()};
    if (!file) throw Error("Could not open file") << file_name;
    Segment segment;
    std::vector<Segment> segs;
    std::string chr;
    file.ignore(1000000, '\n');
    while (file >> chr >> segment.startpos >> segment.stoppos >> segment.n_bins
           >> segment.start >> segment.stop
           >> segment.count >> segment.cn >> segment.expected
           >> segment.int_cn >> segment.type
           >> segment.score) {
      if (!file) throw Error("File input error on") << file_name;
      file.ignore(1000000, '\n');
      segment.chr = chr_lookup(chr);
      segs.push_back(segment);
    }
    if (0) std::cerr << "Loaded " << segs.size()
                     << " segments from " << file_name << std::endl;
    return segs;
  }
  bool pass_score() const {
    return score > 7;  // && stoppos - startpos > 30000;
  }
  bool called_event() const {
    return type != "norm" && pass_score();
  }
};

bool overlap_good(const unsigned int seg_size,
                  const unsigned int overlap_size) {
  return seg_size >= 5 ? (4 * seg_size >= 5 * overlap_size) :
      (seg_size - overlap_size <= 1);
}

class Segments {
 public:
  Segments(const std::string & file_name,
           const std::string & name__,
           const Reference & ref,
           const ChromosomeIndexLookup & chr_lookup) :
      file_{file_name}, name_{name__},
    segs(Segment::load(file_name, chr_lookup)),
    starts_(ref.n_chromosomes() + 1),
    profile{replace_substring(file_name, "segments", "results")}
  {
      for (const Segment & seg : segs) {
        total_count_ += seg.count;
        n_bins_ += seg.n_bins;
        if (seg.pass_score()) {
          ++n_segs_good_;
          n_bins_good_ += seg.n_bins;
          if (seg.called_event()) {
            ++n_segs_good_event_;
            n_bins_good_event_ += seg.n_bins;
          }
        }
      }
      unsigned int s{0};
      unsigned int c{0};
      while (s != segs.size() || c != ref.n_chromosomes()) {
        if (s != segs.size() && c > segs[s].chr) {
          ++s;
        } else {
          starts_[c++] = s;
        }
      }
      starts_.back() = static_cast<unsigned int>(segs.size());
    }
  const std::string & file() const { return file_; }
  const std::string & name() const { return name_; }
  std::vector<Segment>::const_iterator begin() const { return segs.begin(); }
  std::vector<Segment>::const_iterator end() const { return segs.end(); }
  const Segment & operator[](const uint64_t s) const { return segs[s]; }
  uint64_t size() const { return segs.size(); }
  double total_count() const { return total_count_; }

  unsigned int n_segs() const { return static_cast<unsigned int>(segs.size()); }
  unsigned int n_bins() const { return n_bins_; }
  unsigned int n_segs_good() const { return n_segs_good_; }
  unsigned int n_bins_good() const { return n_bins_good_; }
  unsigned int n_segs_good_event() const { return n_segs_good_event_; }
  unsigned int n_bins_good_event() const { return n_bins_good_event_; }
  const std::vector<unsigned int> & starts() const { return starts_; }

 private:
  std::string file_;
  std::string name_{};
  std::vector<Segment> segs;
  std::vector<unsigned int> starts_;

  double total_count_{0};
  unsigned int n_bins_{0};
  unsigned int n_segs_good_{0};
  unsigned int n_bins_good_{0};
  unsigned int n_segs_good_event_{0};
  unsigned int n_bins_good_event_{0};

 public:
  CN_Bins profile;
};

std::ostream & operator<<(std::ostream & out, const Segments & seg) {
  return out << seg.file()
             << " " << seg.total_count()
             << " " << seg.n_segs()
             << " " << seg.n_bins()
             << " " << seg.n_segs_good()
             << " " << 1.0 * seg.n_segs_good() / seg.n_segs()
             << " " << seg.n_bins_good()
             << " " << 1.0 * seg.n_bins_good() / seg.n_bins()
             << " " << seg.n_segs_good_event()
             << " " << 1.0 * seg.n_segs_good_event() / seg.n_segs_good()
             << " " << seg.n_bins_good_event()
             << " " << 1.0 * seg.n_bins_good_event() / seg.n_bins_good();
}


}  // namespace paa

#endif  // PAA_CN_H_
```
