## Supplementary material for "G-Graph: An interactive genomic graph viewer": MUMdex Source Code for G-Graph: cngsl.h

```
//
// cngsl.h
//
// Copy Number Utilities that use gsl
//
// copyright 2017 Peter Andrews
//

#ifndef PAA_CNGSL_H_
#define PAA_CNGSL_H_

#include <gsl/gsl_cdf.h>

#include <algorithm>
#include <limits>
#include <string>
#include <vector>

#include "cn.h"

namespace paa {

class CNStateCall {
 public:
  CNStateCall(const std::vector<double> & state_probs_,
              const unsigned int expected_state_,
              const double segment_count_) :
      state_probs{state_probs_},
    not_state_probs(state_probs.size()),
    expected_state{expected_state_},
    segment_count{segment_count_} {
    double loss_prob{0.0};
    double not_loss_prob{0.0};
    double expected_prob{0.0};
    double not_expected_prob{0.0};
    double gain_prob{0.0};
    double not_gain_prob{0.0};
    for (unsigned int s{0}; s != state_probs.size(); ++s) {
      if (state_probs[s] > state_probs[best_call]) {
        best_call = s;
      }
      if (s < expected_state) {
        loss_prob += state_probs[s];
        not_gain_prob += state_probs[s];
        not_expected_prob += state_probs[s];
      } else if (s > expected_state) {
        gain_prob += state_probs[s];
        not_loss_prob += state_probs[s];
        not_expected_prob += state_probs[s];
      } else {
        expected_prob += state_probs[s];
        not_loss_prob += state_probs[s];
        not_gain_prob += state_probs[s];
      }
    }
    for (unsigned int n{0}; n != state_probs.size(); ++n) {
      for (unsigned int s{0}; s != state_probs.size(); ++s) {
        if (n == s) continue;
        not_state_probs[n] += state_probs[s];
      }
    }

    type = best_call < expected_state ? "loss" :
        (best_call > expected_state ? "gain" : "norm");
    const double min_prob{std::numeric_limits<double>::min()};
    score = -log10(max(best_call < expected_state ? not_loss_prob :
                       (best_call > expected_state ? not_gain_prob :
                        not_expected_prob),
                       min_prob));
    prob_not_expected = max(not_expected_prob, min_prob);
    prob_not_loss = max(not_loss_prob, min_prob);
    prob_not_gain = max(not_gain_prob, min_prob);
    }

  std::vector<double> state_probs;
  std::vector<double> not_state_probs;
  unsigned int expected_state;
  unsigned int best_call{0};
  double segment_count{0.0};
  std::string type{""};
  double score{0.0};
  double prob_not_expected{0.0};
  double prob_not_loss{0.0};
  double prob_not_gain{0.0};
};

class CNStateCaller {
 public:
  CNStateCaller(const Reference & ref_,
                const std::vector<Bin> & bins_,
                const CN_Bins & profile_) :
      ref{ref_}, bins{bins_}, profile{profile_},
    x_chr{ref.find_x_chromosome()},
    y_chr{ref.find_y_chromosome()},
    is_male{is_profile_male()},
    expected_states{get_expected_states()},
    average_autosome_count{get_average_autosome_count()},
    zero_state_count{average_autosome_count / 10} { }

  double get_average_autosome_count() const {
    double count{0};
    unsigned int n_count{0};
    for (unsigned int b{0}; b != bins.size(); ++b) {
      const unsigned int chr{bins[b].chromosome()};
      if (chr == x_chr || chr == y_chr) continue;
      count += profile[b].norm_count();
      ++n_count;
    }
    return count / n_count;
  }

  std::vector<unsigned char> get_expected_states() const {
    std::vector<unsigned char> result;
    for (unsigned int c{0}; c != ref.n_chromosomes(); ++c) {
      const bool is_x{c == x_chr};
      const bool is_y{c == y_chr};
      if (is_male && (is_x || is_y)) {
        result.push_back(1);
      } else if (!is_male && is_y) {
        result.push_back(0);
      } else {
        result.push_back(2);
      }
    }
    return result;
  }

  bool is_profile_male() const {
    double ratio{0};
    unsigned int n_ratio{0};
    for (unsigned int b{0}; b != bins.size(); ++b) {
      const unsigned int chr{bins[b].chromosome()};
      if (chr == y_chr) {
        ratio += profile[b].ratio();
        ++n_ratio;
      }
    }
    return ratio / n_ratio > 0.3333;
  }

  CNStateCall call(const unsigned int start,
                   const unsigned int stop) const {
    const double par_fraction{fraction_par(start, stop)};
    const unsigned int expected_par_copy{2};
    const unsigned int expected_state{static_cast<unsigned int>(
        0.5 + par_fraction * expected_par_copy +
        (1 - par_fraction) * expected_states[bins[start].chromosome()])};
    const double segment_count{[start, stop, this]() {
        double result{0};
        for (unsigned int b{start}; b != stop; ++b) {
          result += profile[b].norm_count();
        }
        return result;
      }()};

    std::vector<double> state_chi_squareds(n_states);
    std::vector<double> state_probs(n_states);
    double total_prob{0.0};
    double diff_power{2.0};
    while (total_prob <= 0) {
      for (unsigned int s{0}; s != n_states; ++s) {
        const double expected_count{(stop - start) *
              (s ? s * average_autosome_count / 2 : zero_state_count)};
        const double diff{expected_count > segment_count ?
              expected_count - segment_count : segment_count - expected_count};
        const double chi_squared{pow(diff, diff_power) / expected_count};
        state_chi_squareds[s] = chi_squared;
        const double prob{gsl_cdf_chisq_Q(chi_squared, 1)};
        total_prob += state_probs[s] = prob;
      }
      diff_power -= 0.1;
    }
    diff_power += 0.1;

    for (unsigned int s{0}; s != n_states; ++s) {
      state_probs[s] /= total_prob;
    }

    return CNStateCall{move(state_probs), expected_state, segment_count};
  }

  double fraction_par(const unsigned int start,
                      const unsigned int stop) const {
    if (bins[start].chromosome() != x_chr) {
      return 0.0;
    }
    const unsigned int par_coords[2][2]{{60000, 2699520},
      {154931043, 155260560}};
    if (bins[stop - 1].stop_position() <= par_coords[0][0] ||
        bins[start].start_position() >= par_coords[1][1] ||
        (bins[start].start_position() >= par_coords[0][1] &&
         bins[stop - 1].stop_position() <= par_coords[1][0])) {
      return 0.0;
    }
    double result{0.0};
    for (unsigned int b{start}; b != stop; ++b) {
      double frac_pos{0.0};
      for (const unsigned int p : {0, 1}) {
        const unsigned int par_len{par_coords[p][1] - par_coords[p][0]};
        if (bins[b].start_position() <= par_coords[p][0]) {
          if (bins[b].stop_position() >= par_coords[p][1]) {
            frac_pos += 1.0 * par_len / bins[b].length();
          } else if (bins[b].stop_position() >= par_coords[p][0]) {
            frac_pos += 1.0 * (bins[b].stop_position() - par_coords[p][0]) /
                bins[b].length();
          }
        } else if (bins[b].start_position() <= par_coords[p][1]) {
          if (bins[b].stop_position() <= par_coords[p][1]) {
            frac_pos += 1.0;
          } else {
            frac_pos += 1.0 * (par_coords[p][1] - bins[b].start_position()) /
                bins[b].length();
          }
        }
      }
      result += frac_pos;
    }
    return result / (stop - start);
  }

  const Reference & ref;
  const std::vector<Bin> & bins;
  const CN_Bins & profile;
  const unsigned int x_chr;
  const unsigned int y_chr;
  const bool is_male;
  const std::vector<unsigned char> expected_states;
  const double average_autosome_count;
  const double zero_state_count;
  const unsigned int n_states{20};
};


}  // namespace paa

#endif  // PAA_CNGSL_H_
```
