## Supplementary material for "G-Graph: An interactive genomic graph viewer": MUMdex Source Code for G-Graph: download_and_setup_ggraph.sh

```
#! /bin/bash

# Genome selection
genomes=$@
if [ "$genomes" != "hg19" ] &&
    [ "$genomes" != "hg38" ] &&
    [ "$genomes" != "hg19 hg38" ] ; then
    echo Required command line arguments must be exactly \
        hg19, hg38, or hg19 hg38 1>&2
    exit 1
fi

echo This program will download, install, and setup G-Graph for you.
echo First it will try to install all prerequisites on Mac OSX and Windows
echo It will also download the "genome(s)" for \
    $genomes, annotations, and sample data.
echo Please contact Peter Andrews at if you encounter difficulties.
echo This has been tested on CentOS Linux, Mac OSX, and Windows under Cygwin.
echo

# Special Cygwin setup on Windows
STARTX=""
if [ "$OSTYPE" = cygwin ] ; then
    STARTX=startxwin

    if [ ! -e /bin/git.exe ] ; then
        echo You need git to be installed in Cygwin before rerunning this script 1>&2
        exit 1
    fi

    if [ ! -e /bin/wget.exe ] ; then
        echo You need wget to be installed in Cygwin before rerunning this script 1>&2
        exit 1
    fi

    if [ ! -e /usr/local/bin/apt-cyg ] ; then
        git clone https://github.com/transcode-open/apt-cyg/
        mv apt-cyg/apt-cyg /usr/local/bin
        rm -Rf apt-cyg
    fi

    apt-cyg install perl-libwww-perl unzip make gcc-g++ libX11-devel xinit ImageMagick xorg-x11-fonts-Type1 libgsl-devel python

fi

if [ "$(uname)" = Darwin ] ; then
    xcode-select --install 2>&1 | grep -v 'already installed'

    if [ ! -e /usr/local/Homebrew ] ; then
        echo Installing Homebrew
        ruby -e "$(curl -fsSL https://raw.githubusercontent.com/Homebrew/install/master/install)"
    fi

    if [ ! -e /usr/local/include/gsl ] ||
        [ ! -e /usr/local/bin/convert ] ; then
        echo Installing ImageMagick and gsl
        brew install imagemagick gsl
    fi

    if [ ! -e /opt/X11/bin/xterm ] ; then
        echo Installing XQuartz
        lwp-request -m GET http://dl.bintray.com/xquartz/downloads/XQuartz-2.7.11.dmg > XQuartz-2.7.11.dmg
        hdiutil attach XQuartz-2.7.11.dmg
        open /Volumes/XQuartz-2.7.11/XQuartz.pkg
        while [ ! -e /opt/X11/bin/xterm ] ; do
            echo waiting for XQuartz install to almost finish
            sleep 30
        done
    fi
fi

# Get and compile ggraph
if [ -e mumdex/ggraph ] ; then
    echo mumdex directory and ggraph program already exists - skipping 1>&2
else
    if [ -e mumdex/ ] ; then
        echo mumdex directory already exists - skipping 1>&2
    else
        url=http://mumdex.com/mumdex.zip
        echo Downloading $url
        git clone https://github.com/docpaa/mumdex/
    fi

    echo Compiling mumdex/ggraph and mumdex/x11plot
    cd mumdex
    which make 1>&2 > /dev/null
    if [ "$?" = 1 ] ; then
        echo You need make to be installed before re-running this script - quitting 1>&2
        exit 1
    fi
    make -j 4 ggraph x11plot > /dev/null
    if [ $? != 0 ] ; then
        (
            echo Problem compiling ggraph and x11plot
            echo Please try to copile manually before continuing
            echo Look at above messages for a hint as to why it failed
            echo You may need a more recent compiler "(gcc 4.9 or later)"
            echo Quitting
        ) 1>&2
        exit 1
    fi
    cd ../
fi

# Get sample data
data="{m,f,d,s}.txt"
if [ -e s.txt ] ; then
    echo sample data already exists - skipping 1>&2
else
    for file in $(eval echo $data) ; do
        url=http://mumdex.com/ggraph/data/$file
        echo Downloading $url
        lwp-request -m GET $url > $file
    done
fi

# Get genomes
for genome in $genomes ; do
    gzip=$genome.fa.gz
    fasta=$genome.fa
    bin=$genome.fa.bin
    if [ -e $bin ] ; then
        echo Genome files for $genome already exist - skipping 1>&2
    else
        config="{knownGene,knownIsoforms,kgXref,cytoBand}.txt"
        for file in $(eval echo $config $gzip) ; do
            url=http://mumdex.com/ggraph/config/$genome/$file
            echo Downloading $url
            if [ $file = $gzip ] ; then
                echo Please be patient for this one
            fi
            lwp-request -m GET $url > $file
        done
        echo Unzipping $gzip
        gunzip $gzip
        mkdir $bin
        eval mv $config $bin
    fi

    if [ ! -e $bin/ref.seq.bin ] ; then
        ./mumdex/ggraph --setup $genome.fa
    fi
done

# make permissions on xwin directory good for all users
mkdir -p /var/log/xwin
chmod 777 /var/log/xwin

# Give instructions to user
echo
echo Setup all done!
for genome in $genomes ; do
    echo
    echo You can now run G-Graph with $genome using the following command:
    echo
    echo $STARTX ./mumdex/ggraph cn $genome.fa abspos,ratio,seg "$data"
    if [ $genome = hg38 ] ; then
        echo 
        echo Note the sample data is processed for hg19 \
            but it will still display "(slightly incorrectly)" for hg38
    fi
done
```
