## Supplementary material for "G-Graph: An interactive genomic graph viewer": MUMdex Source Code for G-Graph: empirical_bins.cpp

```
//
// empirical bins
//
// load up finebin files and generate empirical binning of them
//
// Copyright 2016 Peter Andrews @ CSHL
//

#include <algorithm>
#include <exception>
#include <functional>
#include <iostream>
#include <random>
#include <sstream>
#include <string>
#include <utility>
#include <vector>

#include "cn.h"
#include "error.h"
#include "files.h"
#include "mumdex.h"
#include "psplot.h"
#include "threads.h"
#include "utility.h"

using std::cerr;
using std::cout;
using std::endl;
using std::exception;
using std::function;
using std::istringstream;
using std::min;
using std::move;
using std::mt19937_64;
using std::ostringstream;
using std::random_device;
using std::string;
using std::uniform_real_distribution;
using std::vector;

using paa::Bounds;
using paa::CN_abspos;
using paa::Error;
using paa::FileFinestBins;
using paa::FinestBins;
using paa::FlexVector;
using paa::Mappability;
using paa::Marker;
using paa::MUMdex;
using paa::Progress;
using paa::PSDoc;
using paa::PSGraph;
using paa::PSHSeries;
using paa::PSXYSeries;
using paa::Reference;
using paa::ThreadPool;

using AllFinestBins = FinestBins;

unsigned int total_position_count(const vector<AllFinestBins> & bin_files,
                                  const unsigned int p) {
  unsigned int total_count{0};
  for (unsigned int s{0}; s != bin_files.size(); ++s) {
    total_count += bin_files[s][p];
  }
  return total_count;
}

int main(int argc, char* argv[])  try {
  // Check usage
  if (--argc < 5) throw Error(
          "usage: empirical_bins ref n_bins out_name n_threads bin_file ...");

  // Process command line arguments
  const Reference ref{argv[1]};
  const CN_abspos cn_abspos{ref};
  const string bins_string{argv[2]};
  const string out_name{argv[3]};
  const unsigned int n_threads(atoi(argv[4]));

  argc -= 4;
  argv += 4;

  // Bin file names
  vector<string> bin_names{[argc, argv]() {
      vector<string> result;
      result.reserve(argc);
      for (int s{0}; s != argc; ++s) {
        result.emplace_back(argv[s + 1]);
      }
      return result;
    }()};

  // Open position count files
  const vector<vector<AllFinestBins>> count_files{[&bin_names, &ref]() {
      vector<vector<AllFinestBins>> result(2);
      for (const string & name : bin_names) {
        AllFinestBins bins{ref, name};
        const bool is_female{bins.n_y() == 0};
        if (is_female) {
          if ((bins.n_x() % 2) != 0) throw Error("Bad females");
        } else {
          if (bins.n_y() != bins.n_x()) throw Error("Bad males");
        }
        cout << "Loaded " << name
             << " " << bins.n_samples()
             << " " << bins.n_x()
             << " " << bins.n_y()
             << " " << is_female
             << endl;
        result[is_female].push_back(move(bins));
      }
      return result;
    }()};

  const vector<AllFinestBins> & male_files{count_files[0]};
  const vector<AllFinestBins> & female_files{count_files[1]};

  const unsigned int x{ref.find_x_chromosome()};
  const unsigned int y{ref.find_y_chromosome()};
  const unsigned int x_start{cn_abspos(x, 0)};

  const double total_cutoff_factor{20.0};

  // Sample total counts across genome to get median and make plots
  cout << "Sampling counts to get approximate median total count" << endl;
  const unsigned int median_total_count{
    [&male_files, &female_files,
     total_cutoff_factor, x_start, &out_name]() {
      const unsigned int n_checks{100};
      const unsigned int n_region{100000};
      const unsigned int n_skip{x_start / n_checks};
      vector<unsigned int> total_counts(n_checks * n_region);
      total_counts.clear();
      Progress total_progress{n_checks * n_region, 0.1, "Total count"};
      for (unsigned int c{0}; c != n_checks; ++c) {
        const unsigned int start{c * n_skip};
        const unsigned int stop{min(start + n_region, x_start)};
        for (unsigned int p{start}; p != stop; ++p) {
          const unsigned int total_count{
            total_position_count(male_files, p) +
            total_position_count(female_files, p)};
          total_progress();
          if (total_count) total_counts.push_back(total_count);
        }
      }
      sort(total_counts.begin(), total_counts.end());
      const unsigned int result{total_counts[total_counts.size() / 2]};
      cout << "Sampled median total count is " << result
           << " (excluding counts of 0)" << endl;

      const double total_cutoff{total_cutoff_factor * result};

      // Graphs
      PSDoc ps{out_name + ".finebin"};
      ps.pdf(false);
      const Marker small_red_marker{paa::circle(), 0.1, "1 0 0", 1, true};

      // Total counts hist
      const unsigned int hist_count{static_cast<unsigned int>(
          total_cutoff * 1.1)};
      PSGraph total_counts_hist{ps, ";Total Count;N", Bounds(0, hist_count)};
      PSHSeries<unsigned int, unsigned int> total_counts_series{
        total_counts_hist, hist_count, "1 0 0", false};

      // Counts vs position
      PSGraph counts_vs_pos_graph{ps, ";Position;Total Count"};
      counts_vs_pos_graph.log_y(true);
      PSXYSeries counts_vs_pos_series{counts_vs_pos_graph, small_red_marker};

      // Random scatter
      random_device rd;
      auto mersenne = mt19937_64(rd());
      uniform_real_distribution<double> dist{-0.5, 0.5};
      function<double()> gen{bind(dist, mersenne)};

      const double min_total{total_cutoff};
      Progress plot_progress{n_checks * n_region, 0.1, "Plots"};
      unsigned int loci_seen{0};
      for (unsigned int c{0}; c != n_checks; ++c) {
        const unsigned int start{c * n_skip};
        const unsigned int stop{min(start + n_region, x_start)};
        for (unsigned int p{start}; p != stop; ++p) {
          ++loci_seen;
          const unsigned int total_count{
            total_position_count(male_files, p) +
                total_position_count(female_files, p)};
          plot_progress();
          if (total_count < hist_count) {
            total_counts_series.add_point(total_count);
          }
          if (total_count) {
            if (total_count >= min_total) {
              counts_vs_pos_series.add_point(loci_seen, total_count);
            }
          }
        }
      }
      ostringstream cutoff_line;
      cutoff_line << "2 lw 0 0 1 c np "
                  << total_cutoff << " xc 0 yfc m "
                  << total_cutoff << " xc 1 yfc l sp";
      total_counts_hist.ps(cutoff_line.str());

      return result;
    }()};

  const double total_cutoff{total_cutoff_factor * median_total_count};

  cout << "Total cutoff is " << total_cutoff << endl;

#if 0
  // Population info
  const unsigned int n_samples{[&count_files]() {
      unsigned int result{0};
      for (unsigned int s{0}; s != count_files.size(); ++s) {
        result += count_files[s].n_samples();
      }
      return result;
    }()};
  // Assumes males and females are not mixed together in finebin files
  const unsigned int n_x{[&count_files]() {
      unsigned int result{0};
      for (unsigned int s{0}; s != count_files.size(); ++s) {
        if (count_files[s].n_y() == 0)  // Now only count female Xs
          result += count_files[s].n_x();
      }
      return result;
    }()};
  const unsigned int n_y{[&count_files]() {
      unsigned int result{0};
      for (unsigned int s{0}; s != count_files.size(); ++s) {
        result += count_files[s].n_y();
      }
      return result;
    }()};
  cout << "Samples " << n_samples << " X " << n_x << " Y " << n_y << endl;

  // BAD X and Y chromosome correction factors
  const double f_x_corr{2.0 * n_samples / n_x};
  const double f_y_corr{2.0 * n_samples / n_y};
  cout << "Borrection X " << f_x_corr << " Y " << f_y_corr << endl;
#endif

  struct locus_counts {
    uint64_t n_cut_total{0};
    uint64_t n_cut{0};
    uint64_t loci_seen{0};
    uint64_t male_count{0};
    uint64_t female_count{0};
    locus_counts & operator+=(const locus_counts & rhs) {
      n_cut_total += rhs.n_cut_total;
      n_cut += rhs.n_cut;
      loci_seen += rhs.loci_seen;
      male_count += rhs.male_count;
      female_count += rhs.female_count;
      return *this;
    }
  };

  FlexVector<unsigned char> bad_positions(cn_abspos.n_positions());

  // Sum of all bin counts
  FinestBins combined_counts{ref};
  /*
  combined_counts.n_samples(n_samples);
  combined_counts.n_x(n_x);
  combined_counts.n_y(n_y);
  */
  auto block_function = [&combined_counts, &bad_positions,
                         total_cutoff, &male_files, &female_files]
      (const unsigned int start_pos,
       const unsigned int stop_pos,
       const double corr,
       const bool use_males,
       const bool use_females) {
    // Loop over loci
    locus_counts result;
    for (unsigned int p{start_pos}; p != stop_pos; ++p) {
      ++result.loci_seen;

      const unsigned int male_count{use_males ?
            total_position_count(male_files, p) : 0U};
      const unsigned int female_count{use_females ?
            total_position_count(female_files, p) : 0U};

      const unsigned int total_count{male_count + female_count};
      const double corr_total_count{corr * total_count};
      if (corr_total_count > total_cutoff) {
        ++result.n_cut_total;
        ++result.n_cut;
        bad_positions[p] = 1;
      } else {
        result.male_count += male_count;
        result.female_count += female_count;
        combined_counts[p] = total_count;
      }
    }
    return result;
  };

  // Examine loci one by one in blocks in parallel
  cout << "Looping over loci to total counts and exclude loci" << endl;
  const unsigned int block_size{16 * 10000};
  ThreadPool pool{n_threads};
  ThreadPool::Results<locus_counts> results;
  locus_counts totals;
  double x_corr{1.0};
  double y_corr{1.0};
  for (const unsigned int c : cn_abspos.chromosomes()) {
    const bool use_males{c != x};
    const bool use_females{c != y};
    if (c == x) {
      // Calculate male and female corr from autosome
      Progress progress{results.size(), 0.1, "Autosome Positions"};
      std::cout << "Waiting for " << results.size() << " results" << endl;
      while (results.size()) {
        const locus_counts result{results.get()};
        progress();
        totals += result;
      }
      const uint64_t total_count{totals.male_count + totals.female_count};
      y_corr = 2.0 * total_count / totals.male_count;
      x_corr = 1.0 * total_count / totals.female_count;
      cout << "Correction X " << x_corr << " Y " << y_corr << endl;
    }
    const double corr{c == y ? y_corr : (c == x ? x_corr : 1.0)};
    for (unsigned int b{0}; b < ref.size(c); b += block_size) {
      const unsigned int start{cn_abspos(c, b)};
      const unsigned int chr_stop{cn_abspos.ref_offset(c) + ref.size(c)};
      const unsigned int stop{start + block_size > chr_stop ?
            chr_stop : start + block_size};
      pool.run(results, block_function, start, stop, corr,
               use_males, use_females);
    }
  }

  // Gather X and Y results
  // locus_counts xy_totals;
  Progress progress{results.size(), 0.1, "XY Positions"};
  std::cout << "Waiting for " << results.size() << " results" << endl;
  while (results.size()) {
    const locus_counts result{results.get()};
    progress();
    totals += result;
  }

  bad_positions.save(out_name + ".bad.bin");

  cout << "Total cut: Cut " << totals.n_cut_total << " of "
       << totals.loci_seen << " loci, or "
       << 1.0 * totals.n_cut_total / totals.loci_seen << " of loci" << endl;
  cout << "All cut: Cut " << totals.n_cut << " of " << totals.loci_seen
       << " loci, or "
       << 1.0 * totals.n_cut / totals.loci_seen << " of loci" << endl;

  if (0) {
    cout << "Saving combined bins" << endl;
    combined_counts.save(out_name + ".combined.bin");
  }

  cout << "Finding bin boundaries" << endl;
  istringstream bins_stream{bins_string.c_str()};
  unsigned int n_bins;
  while (bins_stream >> n_bins) {
    cout << "N bins = " << n_bins << endl;
    combined_counts.bin(n_bins, out_name, x_corr, y_corr);
    bins_stream.get();
  }

  return 0;
} catch (Error & e) {
  cerr << "paa::Error:" << endl;
  cerr << e.what() << endl;
  return 1;
} catch (exception & e) {
  cerr << "std::exception" << endl;
  cerr << e.what() << endl;
  return 1;
} catch (...) {
  cerr << "unknown exception was caught" << endl;
  return 1;
}
```
