## Supplementary material for "G-Graph: An interactive genomic graph viewer": MUMdex Source Code for G-Graph: extract_cn_segments.cpp

```
//
// extract_cn_segments
//
// Create segment files from bin results
//
// Copyright 2017 Peter Andrews @ CSHL
//

#include <exception>
#include <iostream>
#include <set>
#include <string>
#include <vector>

#include "cn.h"
#include "cngsl.h"
#include "error.h"
#include "genes.h"
#include "mumdex.h"
#include "strings.h"

using std::cerr;
using std::cout;
using std::endl;
using std::exception;
using std::set;
using std::string;
using std::vector;

using paa::replace;
using paa::Bin;
using paa::ChromosomeIndexLookup;
using paa::CN_Bin;
using paa::CN_Bins;
using paa::CNStateCall;
using paa::CNStateCaller;
using paa::Error;
using paa::GeneHitFinder;
using paa::GeneInfoInterval;
using paa::GeneXrefs;
using paa::HitType;
using paa::KnownGene;
using paa::KnownGenes;
using paa::Reference;

char hit_char(const HitType info) {
  switch (info) {
    case HitType::none:
      return 'n';
      break;
    case HitType::gene:
      return 'g';
      break;
    case HitType::exon:
      return 'e';
      break;
    case HitType::intron:
      return 'i';
      break;
    default:
      throw Error("unexpected HitType");
  }
}

class Segment {
 public:
  Segment(const unsigned int start__, const unsigned int stop__) :
      start_{start__}, stop_{stop__} {}

  unsigned int start() const { return start_; }
  unsigned int stop() const { return stop_; }
  unsigned int n_bins() const { return stop_ - start_; }

  unsigned int start_{0};
  unsigned int stop_{0};
};

int main(int argc, char* argv[])  try {
  // Check usage
  if (--argc != 3) {
    throw Error("usage: extract_cn_segments ref bins cn_output_file");
  }

  const string reference_file{argv[1]};
  const Reference ref{reference_file};

  const ChromosomeIndexLookup chr_lookup{ref};
  const string genes_name{reference_file + ".bin/knownGene.txt"};
  const string isoforms_name{reference_file + ".bin/knownIsoforms.txt"};
  const string kgXrefs_name{reference_file + ".bin/kgXref.txt"};
  const KnownGenes genes{genes_name, isoforms_name, chr_lookup, ref};
  const GeneXrefs xref{kgXrefs_name};
  const GeneHitFinder gene_finder{genes};

  const string bins_name{argv[2]};
  const vector<Bin> all_bins{load_bins(bins_name, ref, false, true)};

  // List of previously determined good bins
  const vector<Bin> bins{[&all_bins]() {
      vector<Bin> result;
      for (const Bin & bin : all_bins)
        if (!bin.bad()) result.push_back(bin);
      return result;
    }()};

  const CN_Bins profile{argv[3]};

  // Bin length sanity check
  if (profile.size() != bins.size()) {
    throw Error("Bin size mismatch") << profile.size() << bins.size();
  }

  // Extract segment data
  const vector<Segment> segments{[&bins, &profile]() {
      unsigned int segment_bin_start{0};
      vector<Segment> result;
      for (unsigned int b{0}; b <= bins.size(); ++b) {
        if (b == bins.size() ||
            bins[b].chromosome() != bins[segment_bin_start].chromosome() ||
            profile[b].seg_ratio() < profile[segment_bin_start].seg_ratio() ||
            profile[b].seg_ratio() > profile[segment_bin_start].seg_ratio()) {
          result.emplace_back(segment_bin_start, b);
          if (b == bins.size()) break;
          segment_bin_start = b;
        }
      }
      return result;
    }()};

  // Prepare to call states
  const CNStateCaller caller{ref, bins, profile};

  cout << "chr\tchrpos_start\tchrpos_stop\tn_bins\tbin_start\tbin_stop\t"
       << "norm_count\tseg_cn\tnorm_cn\tcn_call\tcall\t"
       << "prob_event\tprob_not_loss\tprob_not_gain\tcall_score\tgenes" << endl;
  for (const Segment & segment : segments) {
    const unsigned int chr{bins[segment.start()].chromosome()};

    // Make copy number calls
    const CNStateCall call{caller.call(segment.start(), segment.stop())};

    // Output segment info
    cout << ref.name(bins[segment.start()].chromosome()) << "\t"
         << bins[segment.start()].start_position() << "\t"
         << bins[segment.stop() - 1].stop_position() << "\t"
         << segment.stop() - segment.start() << "\t"
         << segment.start() << "\t"
         << segment.stop() << "\t"
         << call.segment_count << "\t"
         << 2 * profile[segment.start()].seg_ratio() << "\t"
         << call.expected_state << "\t"
         << call.best_call << "\t"
         << call.type << "\t"
         << call.prob_not_expected << "\t"
         << call.prob_not_loss << "\t"
         << call.prob_not_gain << "\t"
         << call.score << "\t";

    // Annotate calls with genes hit
    set<string> gene_hits;
    set<string> exon_hits;
    if (call.best_call != call.expected_state) {
      auto intervals = gene_finder.lookup(
          chr, bins[segment.start()].start_position(),
          bins[segment.stop() - 1].stop_position());
      for (const GeneInfoInterval * i : intervals) {
        const GeneInfoInterval & interval{*i};
        for (const auto & item : interval.info) {
          const unsigned int gene{item.first};
          const string name{xref[genes[gene].name].geneSymbol};
          const char hitchar{hit_char(item.second.hit)};
          if (hitchar == 'e') {
            exon_hits.insert(name);
          }
          gene_hits.insert(name);
        }
      }
      if (exon_hits.empty() && gene_hits.empty()) {
        cout << "-";
      } else {
        bool first{true};
        for (const string & name : exon_hits) {
          if (first) {
            first = false;
          } else {
            cout << " ";
          }
          cout << name;
          gene_hits.erase(name);
        }
        if (gene_hits.size()) {
          if (exon_hits.size()) cout << " ";
          cout << "not_exon_hits: ";
          first = true;
          for (const string & name : gene_hits) {
            if (first) {
              first = false;
            } else {
              cout << " ";
            }
            cout << replace(name, ' ', '_');
          }
        }
      }
    } else {
      cout << "-";
    }

    cout << endl;
  }

  cerr << "done" << endl;

  return 0;
} catch (Error & e) {
  cerr << "paa::Error:" << endl;
  cerr << e.what() << endl;
  return 1;
} catch (exception & e) {
  cerr << "std::exception" << endl;
  cerr << e.what() << endl;
  return 1;
} catch (...) {
  cerr << "unknown exception was caught" << endl;
  return 1;
}
```
