## Supplementary material for "G-Graph: An interactive genomic graph viewer": MUMdex Source Code for G-Graph: find_diploid_exons.cpp

```
//
// find_diploid_exons.cpp
//
// Find exons on each chromosome that are always diploid
//
// Copyright 2019 Peter Andrews @ CSHL
//

#include <algorithm>
#include <exception>
#include <fstream>
#include <future>
#include <iostream>
#include <map>
#include <sstream>
#include <string>
#include <utility>
#include <vector>

#include "cn.h"
#include "error.h"
#include "mumdex.h"
#include "psplot.h"
#include "stats.h"
#include "strings.h"
#include "threads.h"
#include "utility.h"

using std::cin;
using std::cout;
using std::cerr;
using std::endl;
using std::exception;
using std::future;
using std::ifstream;
using std::map;
using std::ofstream;
using std::ostringstream;
using std::pair;
using std::string;
using std::vector;

using paa::remove_substring;
using paa::Bin;
using paa::Bounds;
using paa::ChromosomeIndexLookup;
using paa::CN_abspos;
using paa::CN_Bin;
using paa::CN_Bins;
using paa::Error;
using paa::GeneHitFinder;
using paa::GeneInfoInterval;
using paa::GeneXrefs;
using paa::HitType;
using paa::KnownGene;
using paa::KnownGenes;
using paa::MAD;
using paa::Marker;
using paa::Progress;
using paa::PSDoc;
using paa::PSGraph;
using paa::PSHSeries;
using paa::PSXYSeries;
using paa::Reference;
using paa::ThreadPool;

using GeneInfo = pair<string, bool>;
using GeneInfos = vector<GeneInfo>;
using Strings = vector<string>;
using OutResult = pair<Strings, GeneInfos>;

int main(int argc, char* argv[]) try {
  if (--argc < 3)
    throw Error("usage: find_diploid_exons ref bin_file results_file ...");

  // Process arguments
  const string reference_file{argv[1]};
  const Reference ref{reference_file};
  const ChromosomeIndexLookup lookup{ref};
  const CN_abspos cn_abspos{ref};

  const string bins_name{argv[2]};
  const vector<Bin> all_bins{load_bins(bins_name, ref)};
  // List of previously determined good bins
  const vector<Bin> bins{[&all_bins]() {
      vector<Bin> result;
      for (const Bin & bin : all_bins) {
        if (!bin.bad()) {
          result.push_back(bin);
        }
      }
      return result;
    }()};

  const double resolution{1.0 * ref.size() / all_bins.size()};
  cerr << "Resolution is " << resolution << endl;

  vector<unsigned char> bad_bins(bins.size());

  // Graphs
  const string out_base{"out"};
  PSDoc ps{out_base};
  ps.pdf(false);
  const Marker small_red_marker{paa::circle(), 0.1, "1 0 0", 1, true};

  // Bin Length cut
  PSHSeries<unsigned int, unsigned int> length_hist{
    ps, "Bin lengths;Length;N", 100, Bounds{resolution / 2, resolution * 1.5}};
  vector<double> lengths;
  for (unsigned int b{0}; b != bins.size(); ++b) {
    length_hist.add_point(bins[b].length());
    lengths.push_back(bins[b].length());
  }
  sort(lengths.begin(), lengths.end());
  const MAD length_mad{lengths};
  const double length_cutoff{1.0};
  const double length_dist{length_cutoff * length_mad.mad()};
  unsigned int n_bad_length{0};
  for (unsigned int b{0}; b != bins.size(); ++b) {
    if (bins[b].length() < length_mad.median() - length_dist ||
        bins[b].length() > length_mad.median() + length_dist) {
      bad_bins[b] = 1;
      ++n_bad_length;
    }
  }
  cerr << "Length median " << length_mad.median()
       << " +/- " << length_mad.mad() * length_cutoff
       << " with " << 100.0 * n_bad_length / bins.size()
       << "% cut" << endl;

  // Bin Gc cut
  PSHSeries<double, unsigned int> gc_hist{
    ps, "Bin GC content;GC proportion;N", 100, Bounds{0.0, 1.0}};
  vector<double> gcs;
  for (unsigned int b{0}; b != bins.size(); ++b) {
    gc_hist.add_point(bins[b].gc());
    gcs.push_back(bins[b].gc());
  }
  sort(gcs.begin(), gcs.end());
  const MAD gc_mad{gcs};
  const double gc_cutoff{0.5};
  const double gc_dist{gc_cutoff * gc_mad.mad()};
  unsigned int n_bad_gc{0};
  for (unsigned int b{0}; b != bins.size(); ++b) {
    if (bins[b].gc() < gc_mad.median() - gc_dist ||
        bins[b].gc() > gc_mad.median() + gc_dist) {
      bad_bins[b] = 1;
      ++n_bad_gc;
    }
  }
  cerr << "GC median " << gc_mad.median()
       << " +/- " << gc_mad.mad() * gc_cutoff
       << " with " << 100.0 * n_bad_gc / bins.size()
       << "% cut" << endl;

  // Bin Mappability cut
  PSHSeries<unsigned int, unsigned int> mappability_hist{
    ps, "Bin mappabilities;Mappability;N", 50,
        Bounds{0, 50}};
  vector<double> mappabilitys;
  for (unsigned int b{0}; b != bins.size(); ++b) {
    mappability_hist.add_point(bins[b].map());
    mappabilitys.push_back(bins[b].map());
  }
  sort(mappabilitys.begin(), mappabilitys.end());
  const MAD mappability_mad{mappabilitys};
  const double mappability_cutoff{1.0};
  const double mappability_dist{mappability_cutoff * mappability_mad.mad()};
  unsigned int n_bad_mappability{0};
  for (unsigned int b{0}; b != bins.size(); ++b) {
    if (bins[b].map() < mappability_mad.median() - mappability_dist ||
        bins[b].map() > mappability_mad.median() + mappability_dist) {
      bad_bins[b] = 1;
      ++n_bad_mappability;
    }
  }
  cerr << "Mappability median " << mappability_mad.median()
       << " +/- " << mappability_mad.mad() * mappability_cutoff
       << " with " << 100.0 * n_bad_mappability / bins.size()
       << "% cut" << endl;

  // X limits
  const unsigned int x_chr{ref.find_x_chromosome()};
  const unsigned int y_chr{ref.find_y_chromosome()};
  unsigned int x_start{0};
  unsigned int x_stop{0};
  for (unsigned int b{0}; b != bins.size(); ++b) {
    if (!x_start && bins[b].chromosome() == x_chr) {
      x_start = b;
    }
    if (!x_stop && bins[b].chromosome() == y_chr) {
      x_stop = b;
    }
  }
  if (false) cerr << "X start " << x_start << " stop " << x_stop
      << " of " << bins.size() << endl;


  argc -= 2;
  argv += 3;
  const vector<CN_Bins> results{[argv, argc, &bins] () {
      ThreadPool pool{6};
      vector<future<CN_Bins>> futures;
      for (int a{0}; a != argc; ++a)
        futures.push_back(pool.run([](const string in_file) {
              return CN_Bins(in_file);
            }, argv[a]));
      vector<CN_Bins> result;
      result.reserve(futures.size());
      Progress progress{futures.size(), 0.001, "get sample info"};
      for (future<CN_Bins> & fut : futures) {
        result.push_back(fut.get());
        progress();
        if (result.back().size() != bins.size())
          throw Error("Bin size mismatch");
      }
      return result;
    }()};

  // Sex determination
  PSHSeries<double, unsigned int> sex_hist{
    ps, "Sex Determination;X ratio median;N", 100, Bounds{0.25, 1.25}};
  // Get sex by looking at X ratio median by sample
  vector<double> x_ratios;
  vector<unsigned char> sample_is_male;
  for (const CN_Bins & sample_data : results) {
    double ratio{0.0};
    for (unsigned int b{x_start}; b != x_stop; ++b)
      ratio += sample_data[b].ratio();
    ratio /= x_stop - x_start;
    sex_hist.add_point(ratio);
    sample_is_male.push_back(ratio < 0.75);
  }

  // bin ratio spreads (max - min) by sex and region
  vector<vector<vector<double>>> spreads_by_sex_region(
      2, vector<vector<double>>(3));
  vector<vector<double>> spreads_by_sex(2);
  for (unsigned int b{0}; b != bins.size(); ++b) {
    const Bin & bin{bins[b]};
    const unsigned int region{bin.chromosome() == x_chr ? 1U :
          (bin.chromosome() == y_chr ? 2U : 0U)};
    vector<vector<double>> ratios_by_sex(2);
    for (unsigned int s{0}; s != results.size(); ++s) {
      const CN_Bins & sample_data{results[s]};
      const bool is_male{!!sample_is_male[s]};
      ratios_by_sex[is_male].push_back(sample_data[b].ratio());
    }
    for (const bool is_male : {false, true}) {
      vector<double> & ratios{ratios_by_sex[is_male]};
      sort(ratios.begin(), ratios.end());
      const double spread{ratios.back() - ratios.front()};
      spreads_by_sex_region[is_male][region].push_back(spread);
      spreads_by_sex[is_male].push_back(spread);
    }
  }
  const string region_names[3]{"autosome", "X", "Y"};
  const string sex_names[3]{"female", "male"};
  const double spread_rank_cutoff{0.5};
  vector<vector<double>> spread_cutoffs(2, vector<double>(3));
  for (const unsigned int region : {0, 1, 2}) {
    for (const bool is_male : {false, true}) {
      vector<double> & spreads{spreads_by_sex_region[is_male][region]};
      sort(spreads.begin(), spreads.end());
      auto hist = PSHSeries<double, unsigned int>::create(
          ps, "Ratio spreads for " + sex_names[is_male] + " " +
          region_names[region] + ";spread;N", 100u,
          Bounds{0.0, 1.0});
      for (const double & spread : spreads) hist->add_point(spread);
      spread_cutoffs[is_male][region] =
          spreads[spread_rank_cutoff * spreads.size()];
    }
  }
  vector<vector<unsigned int>> n_bad_spread(2, vector<unsigned int>(3));
  vector<double> average_spread(bins.size());
  for (unsigned int b{0}; b != bins.size(); ++b) {
    const Bin & bin{bins[b]};
    const unsigned int region{bin.chromosome() == x_chr ? 1U :
          (bin.chromosome() == y_chr ? 2U : 0U)};
    bool is_bad{false};
    for (const bool is_male : {false, true}) {
      average_spread[b] += spreads_by_sex[is_male][b];
      if ((is_male || region != 2) &&
          spreads_by_sex[is_male][b] > spread_cutoffs[is_male][region]) {
        is_bad = true;
        ++n_bad_spread[is_male][region];
      }
    }
    if (region < 2) average_spread[b] /= 2;
    if (is_bad) bad_bins[b] = true;
  }
  const uint64_t region_size[3]{
    x_start, x_stop - x_start, bins.size() - x_stop};
  for (const bool is_male : {false, true})
    for (const unsigned int region : {0, 1, 2})
      if (is_male || region != 2)
          cerr << "Spread cutoff for " << sex_names[is_male]
          << " " << region_names[region]
          << " is " << spread_cutoffs[is_male][region]
          << " with "
          << 100.0 * n_bad_spread[is_male][region] / region_size[region]
          << "% cut" << endl;

  // Load gene info
  const string genes_name{reference_file + ".bin/knownGene.txt"};
  const string isoforms_name{reference_file + ".bin/knownIsoforms.txt"};
  const string kgXrefs_name{reference_file + ".bin/kgXref.txt"};
  const KnownGenes genes{genes_name, isoforms_name, lookup, ref};
  const GeneXrefs xref{kgXrefs_name};
  const GeneHitFinder gene_finder{genes};

  // Determine gene overlaps for each bin
  vector<string> bin_gene_annotations;
  vector<unsigned int> bin_gene_n_exons;
  vector<unsigned int> bin_gene_n_exon_genes;
  Progress gene_progress{bins.size(), 0.01, "get gene overlaps"};
  for (const Bin & bin : bins) {
    // get gene info for segment
    ostringstream gene_info;
    const KnownGenes::GeneOverlaps gene_overlaps{
      genes.find_genes(bin.chromosome(),
                       bin.start_position(), bin.stop_position())};
    map<string, unsigned int> named_overlaps;
    for (const KnownGenes::GeneOverlap & overlap : gene_overlaps) {
      const string symbol{xref[genes[overlap.first].name].geneSymbol};
      const unsigned int old_value{named_overlaps[symbol]};
      if (old_value < overlap.second) {
        named_overlaps[symbol] = overlap.second;
      }
    }
    using Overlap = pair<string, unsigned int>;
    vector<Overlap> overlaps(named_overlaps.begin(), named_overlaps.end());
    sort(overlaps.begin(), overlaps.end(),
         [](const Overlap & lhs, const Overlap & rhs) {
           if (lhs.second == rhs.second) {
             return lhs.first < rhs.first;
           } else {
             return lhs.second > rhs.second;
           }
         });
    unsigned int genes_with_exons{0};
    unsigned int total_exons{0};
    for (unsigned int o{0}; o != overlaps.size(); ++o) {
      genes_with_exons += static_cast<bool>(overlaps[o].second);
      total_exons += overlaps[o].second;
    }
    GeneInfos gene_infos;
    for (unsigned int o{0}; o != overlaps.size(); ++o) {
      const Overlap & overlap{overlaps[o]};
      if (o) gene_info << ",";
      if (overlap.second)
        gene_info << overlap.second << "-exon"
                  << (overlap.second > 1 ? "s" : "") << "-";
      gene_info << overlap.first;
      gene_infos.emplace_back(overlap.first, static_cast<bool>(overlap.second));
    }
    if (overlaps.empty()) {
      gene_info << "intergenic";
    }
    const string gene_string{gene_info.str()};
    if (false && gene_string != "intergenic")
      cout << ref.name(bin.chromosome())
           << '\t' << bin.start_position()
           << '\t' << bin.stop_position()
           << '\t' << bin.length()
           << '\t' << bin.map()
           << '\t' << total_exons
           << '\t' << genes_with_exons
           << '\t' << gene_string
           << endl;
    bin_gene_annotations.push_back(gene_string);
    bin_gene_n_exons.push_back(total_exons);
    bin_gene_n_exon_genes.push_back(genes_with_exons);
    gene_progress();
  }

  // pick best bins (lowest spread) to use by chromosome.
  map<unsigned int, vector<unsigned int>> best_bins;
  const uint64_t max_returned{5};
  for (unsigned int b{0}; b != bins.size(); ++b)
    if (!bad_bins[b] && bin_gene_n_exons[b])
      best_bins[bins[b].chromosome()].push_back(b);
  cout << "chr\tabspos\tstart\tstop\tlength\tmap_length\tgc\tratio_spread"
       << "\tn_exons\tn_exonic\tgene_description" << endl;
  for (const unsigned int chr : cn_abspos.chromosomes()) {
    try {
      auto & chosen = best_bins.at(chr);
      sort(chosen.begin(), chosen.end(),
           [&average_spread](const unsigned int b1, const unsigned int b2) {
             return average_spread[b1] < average_spread[b2];
           });
      if (max_returned > chosen.size()) {
        cerr << "Fewer good bins than desired found for "
             << ref.name(chr) << " (" << chosen.size() << ")" << endl;
      } else {
        chosen.resize(max_returned);
      }
      for (const unsigned int b : chosen) {
        const Bin & bin{bins[b]};
        cout << ref.name(bin.chromosome())
             << '\t' << bin.abspos()
             << '\t' << bin.start_position()
             << '\t' << bin.stop_position()
             << '\t' << bin.length()
             << '\t' << bin.map()
             << '\t' << bin.gc()
             << '\t' << average_spread[b]
             << '\t' << bin_gene_n_exons[b]
             << '\t' << bin_gene_n_exon_genes[b]
             << '\t' << bin_gene_annotations[b]
             << endl;
      }
    } catch (...) {
      cerr << "No good bins found for " << ref.name(chr) << endl;
      continue;
    }
  }

  return 0;
} catch (Error & e) {
  cerr << "paa::Error:" << endl;
  cerr << e.what() << endl;
  return 1;
} catch (exception & e) {
  cerr << "std::exception" << endl;
  cerr << e.what() << endl;
  return 1;
} catch (...) {
  cerr << "unknown exception was caught" << endl;
  return 1;
}
```
