## Supplementary material for "G-Graph: An interactive genomic graph viewer": MUMdex Source Code for G-Graph: finebin_cn.cpp

```
//
// finebin_cn.cpp
//
// Copy number from a finebin file
//
// Copyright 2017 Peter Andrews @ CSHL
//

#include <algorithm>
#include <exception>
#include <iostream>
#include <sstream>
#include <string>
#include <vector>

#include "cn.h"
#include "error.h"
#include "files.h"
#include "mumdex.h"
#include "threads.h"
#include "utility.h"

using std::cout;
using std::cerr;
using std::endl;
using std::exception;
using std::istringstream;
using std::max;
using std::min;
using std::string;
using std::to_string;
using std::vector;

using paa::Bin;
using paa::CN_abspos;
using paa::Error;
using paa::FinestBins;
using paa::Mappability;
using paa::MappedVector;
using paa::MUMdex;
using paa::PosInfo;
using paa::Progress;
using paa::Reference;
using paa::ThreadPool;

int main(int argc, char * argv[]) try {
  paa::exit_on_pipe_close();

  if (--argc != 5)
    throw Error("usage: finebin_cn ref bins bad title finebin");

  paa::set_cn_parameters();

  const bool create_pdf{false};

  cerr << "Loading reference and mappability" << endl;
  const Reference ref{argv[1]};
  const CN_abspos cn_abspos{ref};
  const Mappability mappability{ref};

  cerr << "Loading bins" << endl;
  const string bins_string{argv[2]};
  using AllBins = vector<vector<Bin> >;
  const AllBins bins{[&bins_string, &ref]() {
      vector<string> bins_names;
      istringstream bins_stream{bins_string.c_str()};
      string bins_name;
      while (getline(bins_stream, bins_name, ',')) {
        bins_names.push_back(bins_name);
      }
      AllBins result(bins_names.size());
      for (unsigned int n{0}; n != bins_names.size(); ++n) {
        result[n] = load_bins(bins_names[n], ref);
      }
      return result;
    }()};
  const MappedVector<unsigned char> bad{argv[3]};

  const string title{argv[4]};

  cerr << "Loading finebins" << endl;
  const FinestBins finebins{ref, argv[5]};

  cerr << "Assigning mappings to bins" << endl;

  vector<unsigned int> b(bins.size());
  vector<vector<unsigned int>> counts(bins.size());
  for (unsigned int i{0}; i != bins.size(); ++i) {
    counts[i].resize(bins[i].size());
  }
  for (unsigned int p{0}; p != finebins.size(); ++p) {
    if (bad[p]) continue;
    const unsigned int count{finebins[p]};
    if (!count) continue;
    for (unsigned int i{0}; i != bins.size(); ++i) {
      while (p >= bins[i][b[i]].abspos_stop()) {
        ++b[i];
      }
      if (p >= bins[i][b[i]].abspos_start()) {
        counts[i][b[i]] += count;
      }
    }
  }

  for (unsigned int i{0}; i != bins.size(); ++i) {
    copy_number(ref, bins[i], counts[i],
                title + " " + to_string(bins[i].size()) + " bins",
                create_pdf);
  }

  cerr << "Done" << endl;

  return 0;
} catch (Error & e) {
  cerr << "paa::Error:" << endl;
  cerr << e.what() << endl;
  return 1;
} catch (exception & e) {
  cerr << "std::exception" << endl;
  cerr << e.what() << endl;
  return 1;
} catch (...) {
  cerr << "unknown exception was caught" << endl;
  return 1;
}
```
