## Supplementary material for "G-Graph: An interactive genomic graph viewer": MUMdex Source Code for G-Graph: ggraph.cpp

```
//
// ggraph.cpp
//
// CN view gui
//
// Copyright 2016-2019 Peter Andrews @ CSHL
//
//
// example command line:
// ggraph cn hg19.fa chr,pos,ratio,seg {m,f,d,s}.txt
//

#include <algorithm>
#include <exception>
#include <iostream>
#include <fstream>
#include <functional>
#include <future>
#include <map>
#include <memory>
#include <mutex>
#include <random>
#include <sstream>
#include <string>
#include <thread>
#include <utility>
#include <vector>

#include "cn.h"
#include "error.h"
#include "genes.h"
#include "mumdex.h"
#include "strings.h"
#include "threads.h"
#include "x11plot.h"

using std::cout;
using std::cerr;
using std::endl;
using std::exception;
using std::ifstream;
using std::future;
using std::istringstream;
using std::make_unique;
using std::map;
using std::max;
using std::min;
using std::multimap;
using std::mutex;
using std::ostringstream;
using std::pair;
using std::placeholders::_1;
using std::placeholders::_2;
using std::string;
using std::unique_lock;
using std::unique_ptr;
using std::vector;

using paa::dne;
using paa::inv_atanlog;
using paa::rect;
using paa::remove_substring;
using paa::remove_substring_inplace;
using paa::remove_including_final;
using paa::iBounds;
using paa::CN_abspos;
using paa::Error;
using paa::Event;
using paa::GeneXrefs;
using paa::Geometry;
using paa::KnownGene;
using paa::KnownGenes;
using paa::Point;
using paa::Radio;
using paa::Reference;
using paa::RefCN;
using paa::ThreadPool;
using paa::UsageError;
using paa::X11App;
using paa::X11Font;
using paa::X11Graph;

#ifdef __CYGWIN__
unsigned int n_threads{1};
#else
unsigned int n_threads{std::max(std::thread::hardware_concurrency(), 1U)};
#endif

constexpr unsigned int max_gene_mb{100};
constexpr unsigned int max_name_mb{10};

class GeneInfo {
 public:
  using Rect = vector<vector<int>>;
  GeneInfo(const string & name_, const string & description_,
           const Rect & bounds_, const int low_, const int high_) :
      name{name_}, description{description_}, bounds{bounds_},
    low{low_}, high{high_} { }

  const string name;
  const string description;
  Rect bounds;
  int low;
  int high;
};

struct BestVariant {
  BestVariant(const string & description_,
              const double start_, const double stop_) :
      description{description_}, start{start_}, stop{stop_},
    low{start}, high{stop} { }
  string description;
  double start;
  double stop;
  double middle() const { return (start + stop) / 2; }
  bool overlaps(const BestVariant & rhs) const {
    return start <= rhs.stop && rhs.start <= stop;
  }
  bool merge(const BestVariant & rhs) {
    if (overlaps(rhs)) {
      start = min(start, rhs.start);
      stop = min(stop, rhs.stop);
      if (width() < rhs.width()) {
        low = rhs.low;
        high = rhs.high;
        description = rhs.description;
      }
      return true;
    } else {
      return false;
    }
  }

 private:
  // Longest isoform
  double low;
  double high;
  double width() const { return high - low; }
};

bool add_genes(const RefCN & ref,
               X11Graph & graph, const Event & event = Event()) {
  static bool first_call{true};
  if (first_call && graph.range[0][2] > max_gene_mb * 1000000.0) return false;
  static mutex genes_mutex;
  unique_lock<mutex> lock{genes_mutex};
  static const KnownGenes genes{[&ref]() {
      try {
        const string & reference_file{ref.fasta_file()};
        const string genes_name{reference_file + ".bin/knownGene.txt"};
        const string isoforms_name{reference_file + ".bin/knownIsoforms.txt"};
        return KnownGenes{genes_name, isoforms_name, ref.chr_lookup, ref};
      } catch (Error & err) {
        cerr << err.what() << endl;
        return KnownGenes{ref};
      }
    }()};
  static bool missing_message_shown = false;
  if (genes.size() == 0) {
    if (!missing_message_shown) {
      cerr << "******************************************************" << endl;
      cerr << "No gene information was loaded" << endl;
      cerr << "Please download the files" << endl;
      cerr << "knownGenes.txt, knownIsoforms.txt and kgXref.txt" << endl;
      cerr << "from the UCSC website, from mumdex.com or from the" << endl;
      cerr << "G-Graph paper supplementary materials" << endl;
      cerr << "to match your reference, and place in the directory" << endl;
      cerr << ref.fasta_file() + ".bin/" << endl;
      cerr << "Only hg19 and hg38 versions of these files were tested" << endl;
      cerr << "Note chromosome names must match your reference" << endl;
      cerr << "G-Graph will continue to run without this gene info" << endl;
      cerr << "******************************************************" << endl;
      missing_message_shown = true;
    }
    return false;
  }
  static const string kgXrefs_name{ref.fasta_file() + ".bin/kgXref.txt"};
  static const GeneXrefs xref{kgXrefs_name};
  static const paa::GeneLookup gene_lookup{genes, xref};
  lock.unlock();
  if (first_call) {
    first_call = false;  // to enable pre-loading
    return false;
  }

  static map<X11Graph *, vector<GeneInfo>> all_gene_info;
  vector<GeneInfo> & gene_info{all_gene_info[&graph]};

  if (graph.log_radios[0]) return false;

  if (event.type == Event::X) {
    static map<X11Graph *, bool> all_inside_coord;
    bool & inside_coord{all_inside_coord[&graph]};
    switch (event.x->type) {
#if 1
      case KeyPress:
        if (inside_coord) {
          const std::pair<char, KeySym> char_key{graph.get_char_and_keysym(
              event.x->xkey)};
          const KeySym & sym{char_key.second};
          const char c{char_key.first};
          if (0) std::cerr << "keypress " << static_cast<int>(c) << std::endl;
          static map<X11Graph *, std::string> all_entered_search;
          std::string & entered_search{all_entered_search[&graph]};
          if (c == 13 || sym == XK_Return) {
            if (0) std::cerr << "Return" << std::endl;
            const size_t chr_end{entered_search.find_first_of(" :")};
            int abspos_set{0};
            double abspos = ref.cn_abspos.n_positions();
            double abspos_stop = abspos;
            const double winsize{max_name_mb * 1000000.0 / 2};
            const std::string first{entered_search.substr(0, chr_end)};
            for (const bool with_chr : {false, true}) {
              const std::string s_chr_name{
                remove_substring(first, "chr")};
              const std::string chr_name{std::string(with_chr ? "chr" : "") +
                    s_chr_name};
              if (!ref.chr_lookup.exists(chr_name)) continue;
              const unsigned int chr{ref.chr_lookup[chr_name]};
              if (ref.cn_abspos.ref_offset(chr) == ref.cn_abspos.bad_abspos())
                continue;
              if (chr_end == std::string::npos) {
                abspos = ref.cn_abspos(chr, 0);
                abspos_stop = abspos + ref.size(chr);
                abspos_set = 1;
              } else {
                const size_t pos_end{
                  entered_search.find_first_of("- ", chr_end + 1)};
                const std::string chr_pos(entered_search.substr(
                    chr_end + 1, pos_end - chr_end - 1));
                const uint32_t cp{static_cast<uint32_t>(atol(chr_pos.c_str()))};
                abspos = ref.cn_abspos(chr, cp);
                if (pos_end != std::string::npos) {
                  const std::string end_pos(entered_search.substr(pos_end + 1));
                  const uint32_t end_cp{
                    static_cast<uint32_t>(atol(end_pos.c_str()))};
                  if (end_cp > cp) {
                    if (0)
                    std::cerr << "Set " << chr << " "
                              << chr_pos << " " << abspos << " "
                              << end_pos << " " << end_cp << std::endl;
                    abspos_stop = ref.cn_abspos(chr, end_cp);
                    abspos_set = 2;
                  }
                }
                if (!abspos_set) {
                  abspos_stop = abspos + winsize / 2;
                  abspos -= winsize / 2;
                  abspos_set = 3;
                }
              }
              if (abspos_set) break;
            }
            std::string gene_found{""};
            if (!abspos_set) {
              for (const bool uc : {false, true}) {
                const std::string gene_name{[&first, uc] () {
                    std::string result;
                    for (const char g : first) result += uc ? toupper(g) : g;
                    return result;
                  }()};
                const paa::GeneLookup::ER lookup{
                  gene_lookup.isoforms(gene_name)};
                if (lookup.first != lookup.second) {
                  const KnownGene * longest{lookup.first->second};
                  for (paa::GeneLookup::Iter i{lookup.first};
                       i != lookup.second; ++i) {
                    const KnownGene * g{i->second};
                    if (g->length() > longest->length()) longest = g;
                  }
                  abspos_set = uc ? 5 : 4;
                  abspos = ref.cn_abspos(
                      longest->chr, (longest->t_start + longest->t_stop) / 2);
                  abspos_stop = abspos + winsize / 2;
                  abspos -= winsize / 2;
                  gene_found = gene_name;
                  break;
                }
              }
            }
            if (0) std::cerr << "abspos " << abspos << std::endl;
            if (abspos_set) {
              graph.set_range(0, abspos, abspos_stop);
              graph.prepare_draw();
              graph.status =  "Zoomed to ";
              if (gene_found.size()) graph.status += gene_found + " ";
              const CN_abspos::ChrPos start_pos{ref.cn_abspos.chrpos(abspos)};
              CN_abspos::ChrPos stop_pos{
                ref.cn_abspos.chrpos(abspos_stop)};
              if (stop_pos.first > start_pos.first &&
                  stop_pos.second == 0) {
                stop_pos.first = start_pos.first;
                stop_pos.second = ref.size(stop_pos.first);
              }
              graph.status += ref.name(start_pos.first) + ":" +
                  std::to_string(start_pos.second) + "-" +
                  (start_pos.first != stop_pos.first ?
                   ref.name(stop_pos.first) + ":" : std::string("")) +
                  std::to_string(stop_pos.second);
              graph.draw_status();
              entered_search.clear();
              return true;
            }
            graph.status = "Bad query: ";
            graph.status += entered_search;
            graph.draw_status();
            entered_search.clear();;
            return false;
          } else {
            if (c >= XK_space && c < XK_asciitilde) {
              entered_search += c;
            } else if (sym == XK_Delete || sym == XK_BackSpace) {
              if (entered_search.size()) entered_search.pop_back();
            }
            graph.status = std::string("search: ") + entered_search;
          }
          if (0) std::cerr << graph.status << std::endl;
          graph.draw_status(true);
          return true;
        }
        return false;
#endif
      case MotionNotify:
        {
          const XMotionEvent & xmotion{event.x->xmotion};
          // For coordinate text entry
          // if (graph.call_back_radios[1].contains(xmotion)) {
          if (graph.coord_radio.contains(xmotion)) {
            inside_coord = true;
          } else {
            inside_coord = false;
          }

          // Show gene description on hover
          for (const GeneInfo & gene : gene_info) {
            if (xmotion.x >= max(graph.bounds[0][0], gene.bounds[0][0]) &&
                xmotion.x <= min(graph.bounds[0][1], gene.bounds[0][1]) &&
                xmotion.y >= max(graph.bounds[1][0], gene.bounds[1][0]) &&
                xmotion.y <= min(graph.bounds[1][1], gene.bounds[1][1])) {
              graph.status = gene.description;
              remove_substring_inplace(&graph.status, "Homo sapiens ");
              remove_substring_inplace(&graph.status,
                               std::string(" (") + gene.name + ")");
              remove_substring_inplace(
                  &graph.status, std::string(" ") + gene.name);
              graph.draw_status(true);
              graph.status = "";
              return true;
            }
          }
          // Show chromosomal position
          if (graph.coord_radio) {
            if (xmotion.x >= graph.bounds[0][0] &&
                xmotion.x <= graph.bounds[0][1] &&
                xmotion.y >= graph.bounds[1][0] &&
                xmotion.y <= graph.bounds[1][1]) {
              const double abspos{graph.icoord(0, xmotion.x)};
              if (abspos <= 0 || abspos >= ref.cn_abspos.n_positions()) {
                graph.status = "";
                graph.draw_status();
                return true;
              }
              const pair<unsigned int, unsigned int> chrpos{
                ref.cn_abspos.chrpos(abspos)};
              std::ostringstream coordinates;
              coordinates << std::setprecision(12) << "("
                          << ref.name(chrpos.first);
              const double vals[2]{static_cast<double>(chrpos.second),
                    graph.icoord(1, xmotion.y)};
              for (const bool y : {false, true}) {
                const double val{vals[y]};
                const double res{graph.range[y][2] / graph.bounds[y][2]};
                const double pres{pow(10, floor(log10(res)))};
                const double rval{round(val / pres) * pres};
                const double nval{(y ? graph.y_cn_scale : 1) *
                      (graph.log_radios[y].state() >= 2 ? inv_atanlog(rval) :
                       (graph.log_radios[y] ? pow(10, rval) : rval))};
                if (y && graph.tiled_radio) {
                  coordinates << " , Y coordinate not available in tiled mode";
                } else {
                  coordinates << (y ? " , " : " ") << nval;
                }
              }
              coordinates << " )";
              graph.status = coordinates.str();
              graph.draw_status();
              return true;
            }
          }
        }
        return false;
      case ButtonRelease:
        {
          // Open genome browser
          for (const GeneInfo & gene : gene_info) {
            if (graph.click.x >= gene.bounds[0][0] &&
                graph.click.x <= gene.bounds[0][1] &&
                graph.click.y >= gene.bounds[1][0] &&
                graph.click.y <= gene.bounds[1][1]) {
              const pair<unsigned int, unsigned int> chrpos_start{
                ref.cn_abspos.chrpos(graph.icoord(0, gene.low))};
              const pair<unsigned int, unsigned int> chrpos_stop{
                ref.cn_abspos.chrpos(graph.icoord(0, gene.high))};
              const unsigned int chr{chrpos_start.first};
              const unsigned int range{
                chrpos_stop.second - chrpos_start.second};
              const unsigned int display_start(
                  chrpos_start.second > 0.1 * range + 1 ?
                  chrpos_start.second - 0.1 * range + 1 : 1);
              const unsigned int display_stop(
                  chrpos_stop.second + 0.1 * range < ref.size(chr) ?
                  chrpos_stop.second + 0.1 * range : ref.size(chr));
              ostringstream gene_url;
              gene_url << "'http://genome-mirror.cshl.edu/"
                       << "cgi-bin/hgTracks?db=" << ref.name() << "&"
                       << "position=" << ref.name(chr) << ":"
                       << display_start << "-" << display_stop << "'";
              graph.open_url(gene_url.str());
              return true;
            }
          }
        }
        return false;;

      default:
        break;
    }
  }

  if (event.type != Event::Draw) return false;

  gene_info.clear();
  if (graph.range[0][2] > max_gene_mb * 1000000.0) return false;

  // Get gene limits from graph range
  using ChromPos = pair<unsigned int, unsigned int>;
  std::vector<KnownGene>::const_iterator gene_limits[2];
  for (const bool high : {false, true}) {
    const unsigned int abspos{
      min(ref.cn_abspos.n_positions() - 1,
          static_cast<unsigned int>(max(1.0, graph.range[0][high])))};
    const ChromPos chrpos_bound{ref.cn_abspos.chrpos(abspos)};
    gene_limits[high] = upper_bound(
        genes.begin(), genes.end(), chrpos_bound,
        [high](const ChromPos cp, const KnownGene & gene) {
          if (gene.chr == cp.first) {
            const double e{1000000};
            if (high) {
              return gene.t_stop >= cp.second + e;
            } else {
              return gene.t_start + e >= cp.second;
            }
          }
          return gene.chr >= cp.first;
        });
  }

  if (gene_limits[0] > gene_limits[1]) gene_limits[0] = gene_limits[1];

  static GC gc{graph.create_gc(graph.app.black, graph.app.white, 2)};

  // Set clip rectangle
  static iBounds last_bounds;
  if (graph.bounds != last_bounds) {
    XRectangle clip_rectangle(rect(graph.bounds));
    XSetClipRectangles(graph.display(), gc, 0, 0, &clip_rectangle, 1, YXBanded);
  }

  // Draw all exons and genes in range
  const int exon_height{10};  // Should scale with window?
  const unsigned int gene_y(graph.bounds[1][0] * 1.3);
  const unsigned int exon_y{gene_y - exon_height / 2};
  using Item = pair<string, BestVariant>;
  multimap<string, BestVariant> names;
  for (std::vector<KnownGene>::const_iterator g{gene_limits[0]};
       g != gene_limits[1]; ++g) {
    // Gene line
    const KnownGene & gene{*g};
    const unsigned int min_gene{ref.cn_abspos(gene.chr, gene.t_start)};
    const unsigned int max_gene{ref.cn_abspos(gene.chr, gene.t_stop)};
    if (max_gene <= graph.range[0][0] - 0.2 * graph.range[0][2] ||
        min_gene >= graph.range[0][1] + 0.2 * graph.range[0][2])
      continue;
    const double gene_start{graph.dcoord(0, min_gene)};
    const double gene_stop{max(graph.dcoord(0, max_gene), gene_start + 1)};
    if (graph.range[0][2] <= max_gene_mb * 1000000.0) {
      const Item to_add{
        xref[gene.name].geneSymbol,
        {xref[gene.name].description, gene_start, gene_stop}};
      auto bounds = names.equal_range(to_add.first);
      bool merged{false};
      for (auto i = bounds.first; i != bounds.second; ++i) {
        if (i->second.merge(to_add.second)) {
          merged = true;
          for (auto j = next(i); j != bounds.second; ++j) {
            if (i->second.merge(j->second)) {
              --j;
              names.erase(next(j));
            }
          }
        }
      }
      if (!merged) names.insert(bounds.second, to_add);
    }
    XDrawLine(graph.display(), graph.pixmap, gc,
              max(graph.bounds[0][0], static_cast<int>(gene_start)), gene_y,
              min(graph.bounds[0][1], static_cast<int>(gene_stop)), gene_y);

    // Exon boxes
    for (unsigned int e{0}; e != gene.exon_starts.size(); ++e) {
      const unsigned int exon_start{
        ref.cn_abspos(gene.chr, gene.exon_starts[e])};
      const unsigned int exon_stop{ref.cn_abspos(gene.chr, gene.exon_stops[e])};
      if (exon_start >= graph.range[0][1] || exon_stop <= graph.range[0][0])
        continue;
      const double mod_start{max(graph.range[0][0], 1.0 * exon_start)};
      const double mod_stop{min(graph.range[0][1], 1.0 * exon_stop)};
      const double frac_pixel{(mod_stop - mod_start) * graph.scale[0]};
      if (frac_pixel < 0.5) continue;
      const int box_start{graph.coord(0, mod_start)};
      const int box_stop{frac_pixel < 1.5 ? box_start + 1 :
            graph.coord(0, mod_stop)};
      if (1) {
        XFillRectangle(graph.display(), graph.pixmap, graph.gc,
                       box_start, exon_y, box_stop - box_start, exon_height);
      }
    }
  }

  if (graph.range[0][2] > max_name_mb * 1000000.0) return false;

  // Get good font for gene names
  const double avail_height{(graph.bounds[1][0] - graph.border_width) * 0.6};
  const X11Font * fits{graph.app.good_font("A", 1000, avail_height, gc)};

  // Draw gene names nicely
  std::vector<Item> snames(names.begin(), names.end());
  sort(snames.begin(), snames.end(), [](const Item & lhs, const Item & rhs) {
      return lhs.second.middle() < rhs.second.middle();
      const double l{lhs.second.middle()};
      const double r{rhs.second.middle()};
      if (!dne(l, r)) return lhs.first < rhs.first;
      return l < r;
    });
  static vector<double> last_right(100);
  last_right.clear();
  last_right.resize(100, -1000000.0);
  for (const Item & entry : snames) {
    const string & name{entry.first};
    const double tpos{entry.second.middle()};
    const int width(fits->string_width(name + "  "));
    const double left(tpos - width / 2.0);
    const double xpos{fits->d_centered_x(name, tpos)};
    for (unsigned int i{0}; i != last_right.size(); ++i) {
      if (left > last_right[i]) {
        const int ypos(exon_y + 2 * graph.border_width +
                       exon_height + i * fits->height());
        XDrawString(graph.display(), graph.pixmap, gc, xpos,
                    fits->below_y(ypos), const_cast<char *>(name.c_str()),
                    static_cast<unsigned int>(name.size()));
        const double right{tpos + width / 2};
        last_right[i] = right;
        const GeneInfo::Rect gene_rect{
          {static_cast<int>(left), static_cast<int>(right)},
          {ypos, ypos + fits->height()}};
        gene_info.emplace_back(name, entry.second.description, gene_rect,
                               entry.second.start, entry.second.stop);
        break;
      }
    }
  }

  // Show sequence!
  const string test_bases{"GGGGGGGGGG"};
  const double base_width{graph.bounds[0][2] / graph.range[0][2]};
  const X11Font * bfont{graph.app.good_font(
      test_bases, test_bases.size() * base_width, 1000)};
  const int actual_width(bfont->string_width(test_bases) / test_bases.size());
  if (actual_width <= base_width) {
    unsigned int n_bases{0};
    const int y_pos{graph.bounds[1][1] - 20 * graph.border_width};
    for (unsigned int b = max(0.0, graph.range[0][0]);
         b < graph.range[0][1]; ++b) {
      const pair<unsigned int, unsigned int> chrpos{ref.cn_abspos.chrpos(b)};
      const string base{ref[chrpos.first][chrpos.second]};
      const int x_coord{graph.coord(0, b)};
      const int x_pos{bfont->centered_x(base, x_coord)};
      if (x_coord - actual_width / 2 > graph.bounds[0][0] &&
          x_coord + actual_width / 2 < graph.bounds[0][1]) {
        XDrawString(graph.display(), graph.pixmap, gc,
                    x_pos, y_pos, base.c_str(), 1);
        ++n_bases;
      }
    }
    // Stupid easter egg "it's turtles all the way down"
    if (n_bases <= 1) {
      const string turtle{"turtle"};
      fits = graph.app.good_font(turtle, graph.bounds[0][2] / 10, 1000, gc);
      const paa::Axis axis{graph.range[0][0], graph.range[0][1], 8, false};
      for (const std::pair<double, bool> tick : axis.ticks()) {
        if (!tick.second) continue;
        const int x_coord{graph.coord(0, tick.first)};
        const int x_pos{bfont->centered_x(turtle, x_coord)};
        XDrawString(graph.display(), graph.pixmap, gc,
                    x_pos, y_pos - 1.2 * bfont->height(),
                    turtle.c_str(), static_cast<unsigned int>(turtle.size()));
      }
    }
  }
  return false;
}

bool add_chromosomes(const RefCN & ref,
                     X11Graph & graph, const Event & event = Event()) {
  if (event.type != Event::Draw) return false;

  const vector<unsigned int> & chromosomes{ref.cn_abspos.chromosomes()};
  int min_w{100000000};
  vector<unsigned int> chr;
  vector<int> pos;
  vector<int> widths;
  for (unsigned int c{0}; c != chromosomes.size(); ++c) {
    double cl(ref.cn_abspos.cn_offset(c));
    double ch(ref.cn_abspos.cn_offset(c + 1));
    if (graph.log_radios[0]) cl = log10(cl);
    if (graph.log_radios[0]) ch = log10(ch);
    const double gl{graph.range[0][0]};
    const double gh{graph.range[0][1]};
    if (cl < gh && ch > gl) {
      const double l{max(cl, gl)};
      const double h{min(ch, gh)};
      const int tl{graph.coord(0, l)};
      const int th{graph.coord(0, h)};
      const int m{(th + tl) / 2};
      const int w{th - tl};
      if (cl > gl && ch < gh) min_w = min(min_w, w);
      widths.push_back(w);
      chr.push_back(chromosomes[c]);
      pos.push_back(m + 1);
    }
  }

  // Chromosome name
  const double avail_height{0.8 * (graph.bounds[1][0] - graph.border_width)};
  const std::string longest{"22"};  // In practice, does "22" fit into chr 21?
  static GC gc{graph.create_gc(graph.app.black, graph.app.white)};
  const X11Font * fits{graph.app.good_font(longest, min_w, avail_height, gc)};
  static iBounds last_bounds;
  if (graph.bounds != last_bounds) {
    XRectangle clip_rectangle(rect(graph.bounds));
    XSetClipRectangles(graph.display(), gc, 0, 0, &clip_rectangle, 1, YXBanded);
  }
  for (unsigned int c{0}; c != chr.size(); ++c) {
    const std::string name{remove_substring(ref.name(chr[c]), "chr")};
    if (fits->string_width(name) < widths[c])
      XDrawString(graph.display(), graph.pixmap, gc, fits->centered_x(
          name, pos[c]), fits->centered_y(
              graph.bounds[1][1] - graph.border_width - avail_height * 3 / 8),
                  const_cast<char *>(name.c_str()),
                  static_cast<unsigned int>(name.size()));
  }

  // Chromosome boundary lines
  for (unsigned int c{0}; c <= chromosomes.size(); ++c) {
    double chr_start(ref.cn_abspos.cn_offset(c));
    if (graph.log_radios[0]) chr_start = log10(chr_start);
    if (chr_start > graph.range[0][0] && chr_start < graph.range[0][1]) {
      const unsigned int x_pos(graph.coord(0, chr_start));
      XDrawLine(graph.display(), graph.pixmap, graph.gc,
                x_pos, graph.bounds[1][0], x_pos, graph.bounds[1][1]);
    }
  }
  return false;
}

bool add_ratio_lines(const vector<double> cn_lines,  // Not a reference
                     X11Graph & graph, const Event & event = Event()) {
  if (event.type != Event::Draw) return false;
  // if (graph.tiled_radio) return false;

  // Ratio lines
  for (double y : cn_lines) {
    GC gc{(y >= 10) ? graph.major_gc :
          (fabs(y - floor(y)) < 0.01 ? graph.gc : graph.minor_gc)};
    y /= graph.y_cn_scale;
    if (graph.log_radios[1]) {
      if (y <= 0) continue;
      y = graph.log_radios[1].state() >= 2 ? paa::atanlog(y) : log10(y);
    }
    if (y > graph.range[1][0] && y < graph.range[1][1]) {
      const unsigned int y_pos(graph.coord(1, y));
      if (graph.tiled_radio) {
        for (unsigned int s{0}; s != graph.n_files(); ++s) {
          const int y_tiled{graph.y_tile(s, y_pos)};
          XDrawLine(graph.display(), graph.pixmap, gc,
                    graph.bounds[0][0], y_tiled, graph.bounds[0][1], y_tiled);
        }
      } else {
        XDrawLine(graph.display(), graph.pixmap, gc,
                  graph.bounds[0][0], y_pos, graph.bounds[0][1], y_pos);
      }
    }
  }
  return false;
}

struct CytobandInfo {
  CytobandInfo(const string & chr,
               const unsigned int start, const unsigned int stop,
               const string & name_, const GC gc_) :
      abspos_start{start}, abspos_stop{stop},
    name{chr + " " + name_}, gc{gc_} { }
  CytobandInfo(const int start, const int stop,
               const string & name_) :
      abspos_start{static_cast<unsigned int>(start)},
    abspos_stop{static_cast<unsigned int>(stop)}, name{name_} { }
  CytobandInfo(const CytobandInfo &) = default;
  CytobandInfo & operator=(const CytobandInfo &) = default;
  unsigned int abspos_start;
  unsigned int abspos_stop;
  string name;
  GC gc{nullptr};
  bool operator<(const CytobandInfo & rhs) const {
    return abspos_start < rhs.abspos_start;
  }
};

bool add_cytobands(const RefCN & ref,
                   X11Graph & graph, const Event & event = Event()) {
  static vector<CytobandInfo> cytobands{
    [&ref, &graph]() {
      // GCs for different stain colors
      const map<string, string> stain_colors{
        {"gneg", "grey100"},
        {"gpos25", "grey90"},
        {"gpos50", "grey70"},

        {"gpos75", "grey40"},
        {"gpos100", "grey0"},
        {"gvar", "grey100"},
        {"stalk", "brown3"},
        {"acen", "brown4"}};
      map<string, GC> stain_gcs;
      for (const auto & sc : stain_colors) {
        const string stain_name{sc.first};
        const string color_name{sc.second};
        auto inserted = stain_gcs.emplace(stain_name, nullptr);
        if (inserted.second) {
          GC & gc{inserted.first->second};
          XColor color;
          if (!XAllocNamedColor(graph.display(), graph.app.colormap,
                                stain_colors.at(stain_name).c_str(),
                                &color, &color))
            throw Error("Could not get color") << color_name;
          gc = graph.create_gc(color.pixel, graph.app.white, 1,
                               LineSolid, CapButt, JoinMiter);
        }
      }

      // Read bands file
      const string bands_name{ref.fasta_file() + ".bin/cytoBand.txt"};
      ifstream bands{bands_name.c_str()};
      if (!bands) {
        cerr << "Could not load cytobands file " << bands_name << endl;
        return vector<CytobandInfo>{};
      }
      // bands.ignore(1000, '\n');
      string chr_name;
      unsigned int start;
      unsigned int stop;
      string name;
      string stain;
      vector<CytobandInfo> result;
      while (bands >> chr_name >> start >> stop) {
        bands.get();
        getline(bands, name, '\t');
        bands >> stain;
        if (!bands) throw Error("Problem parsing cytoband file") << bands_name;
        if (chr_name.find("_") != string::npos ||
            chr_name.find("M") != string::npos) {
          continue;
        }
        const unsigned int chr{ref.chr_lookup[chr_name]};
        result.emplace_back(chr_name,
                            ref.cn_abspos(chr, start), ref.cn_abspos(chr, stop),
                            name, stain_gcs[stain]);
      }
      sort(result.begin(), result.end());
      return result;
    }()};

  static bool shown_missing_message{false};
  if (cytobands.empty()) {
    if (!shown_missing_message) {
      cerr << "******************************************************" << endl;
      cerr << "No cytoband information was loaded" << endl;
      cerr << "Please download the file cytoBand.txt" << endl;
      cerr << "from the UCSC website, from mumdex.com or from the" << endl;
      cerr << "G-Graph paper supplementary materials" << endl;
      cerr << "to match your reference, and place in the directory" << endl;
      cerr << ref.fasta_file() + ".bin/" << endl;
      cerr << "Only hg19 and hg38 versions of these files were tested" << endl;
      cerr << "Note chromosome names must match your reference" << endl;
      cerr << "G-Graph will continue to run without this info" << endl;
      cerr << "******************************************************" << endl;
      shown_missing_message = true;
    }
    return false;
  }

  static int band_low{0};
  static int band_high{0};
  static map<X11Graph *, vector<CytobandInfo>> all_band_info;
  vector<CytobandInfo> & band_info{all_band_info[&graph]};

  if (event.type == Event::X && event.x->type == MotionNotify) {
    // Show gene description on hover
    const XMotionEvent & xmotion{event.x->xmotion};
    if (xmotion.y < band_low || xmotion.y > band_high) return false;
    if (xmotion.x < graph.bounds[0][0] ||
        xmotion.x > graph.bounds[0][1]) return false;
    auto found = upper_bound(band_info.begin(), band_info.end(), xmotion.x,
                             [](const unsigned int lhs,
                                const CytobandInfo & rhs) {
                               return lhs < rhs.abspos_start;
                             });
    if (found == band_info.begin()) return false;
    --found;
    if (xmotion.x > static_cast<int>(found->abspos_stop)) return false;
    graph.status = found->name;
    graph.draw_status(true);
    graph.status = "";
    return true;
  }
  if (event.type != Event::PreDraw) return false;

  // Draw stains
  band_info.clear();
  for (const CytobandInfo & band : cytobands) {
    if (band.abspos_start > graph.range[0][1] ||
        band.abspos_stop < graph.range[0][0]) continue;
    const double start{max(1.0 * band.abspos_start, graph.range[0][0])};
    const double stop{min(1.0 * band.abspos_stop, graph.range[0][1])};
    const double avail_height{(graph.bounds[1][0] - graph.border_width) * 0.6};
    const int x_start{graph.coord(0, start)};
    const int x_stop{graph.coord(0, stop)};
    band_low = graph.bounds[1][1] - 2.5 * avail_height;
    band_high = band_low + avail_height;
    XFillRectangle(graph.display(), graph.pixmap, band.gc, x_start, band_low,
                   x_stop - x_start, avail_height);
    band_info.emplace_back(x_start, x_stop, band.name);
  }

  return false;
}

using String = std::string;
using Strings = std::vector<String>;

// Read Specific columns from disk
// and convert chr chrpos to abspos in process if requested
class AbsposColumns {
 public:
  using String = std::string;
  using Strings = std::vector<String>;
  using Column = std::vector<double>;
  using Data = std::vector<Column>;
  using iStream = std::istringstream;
  AbsposColumns(const String & file_name,
                const String & columns,
                const RefCN * ref,
                const uint64_t size_hint = 500000,
                const bool add_jitter = false,
                const double percent = 100.0) {
    // Get list of desired column names or numbers
    iStream column_input{columns.c_str()};
    String column_spec;
    String chr_col_name;
    bool header{false};
    bool implicit_index{false};
    bool has_chr{false};
    while (getline(column_input, column_spec, ',')) {
      // column spec is either name or name:type
      iStream spec_stream{column_spec.c_str()};
      String column_name{};
      String column_type{};
      getline(spec_stream, column_name, ':');
      if (spec_stream) spec_stream >> column_type;
      if (column_name.find_first_not_of("0123456789") != string::npos)
        header = true;
      if (column_names.empty() && column_name == "implicit") {
        implicit_index = true;
        continue;
      }
      // chr column must be first and marked or named appropriately
      // and then if chr column exists the next column must be a chrpos column
      if (ref && column_names.empty() &&
          (column_type == "c" ||
           (column_name[0] == 'c' &&
            (column_name == "chr" || column_name == "chrom" ||
             column_name == "chromosome")) ||
           (column_name[0] == 'C' &&
            (column_name == "Chr" || column_name == "Chrom" ||
             column_name == "Chromosome")))) {
        has_chr = true;
      }
      column_names.push_back(column_name);
      column_types.push_back(column_type);
    }

    if (column_names.size() + implicit_index < 2)
      throw UsageError("Column loader expects to load at least two columns "
                       "from column string") << columns;
    if (ref && implicit_index)
      throw UsageError("Plot types genome and cn cannot be used "
                       "with an implicit index");

    // Input file name
    std::ifstream input{file_name.c_str()};
    if (!input) throw UsageError("Could not open input file") << file_name;

    // Process header, or just use column numbers passed
    using ColumnInfo = std::pair<unsigned int, unsigned int>;
    using ColumnLookup = std::vector<ColumnInfo>;
    const ColumnLookup column_numbers{
      [this, header, columns, &input] () {
        ColumnLookup result;
        if (header) {
          // Column names are strings
          String header_line;
          getline(input, header_line);
          String column_name;
          iStream header_stream{header_line.c_str()};
          unsigned int column{0};
          while (header_stream >> column_name) {
            const Strings::const_iterator found{
              find(column_names.begin(), column_names.end(), column_name)};
            if (found != column_names.end())
              result.emplace_back(column, found - column_names.begin());
            ++column;
          }
        } else {
          // Column names are numbers, starting with 1
          for (unsigned int c{0}; c != column_names.size(); ++c) {
            result.emplace_back(static_cast<unsigned int>(
                stoul(column_names[c]) - 1), c);
            column_names[c] = String(header ? "" : "column ") +
                column_names[c];
          }
          sort(result.begin(), result.end(),
               [] (const ColumnInfo & lhs, const ColumnInfo & rhs) {
                 return lhs.first < rhs.first;
               });
        }
        if (column_names.size() != result.size())
          throw UsageError("Could not find all columns specified in")
              << columns;
        return result;
      }()};

    // Reserve space for data
    data.resize(column_names.size());
    for (unsigned int c{0}; c != data.size(); ++c)
      if (!has_chr || c) data.reserve(size_hint);

    std::random_device rd{};
    std::mt19937_64 mersenne{rd()};
    auto randGen = std::bind(
        std::uniform_real_distribution<double>(0.0, 100.0), mersenne);

    // Read in data line by line
    String line;
    String text;
    String chr_name;
    double value{0};
    while (getline(input, line)) {
      if (percent < 100.0 && randGen() > percent) continue;
      iStream stream{line.c_str()};
      ColumnLookup::const_iterator nc{column_numbers.begin()};
      unsigned int c{0};
      while (stream) {
        if (nc != column_numbers.end() && nc->first == c) {
          if (has_chr && nc->second == 0) {
            stream >> chr_name;
          } else {
            if (stream >> value) {
              data[nc->second].push_back(value);
            } else {
              throw Error("Could not parse data line:\n")
                  << line << "\nfrom data file" << file_name;
            }
          }
          ++nc;
        } else {
          stream >> text;  // should ignore?  space vs tab is why not
        }
        ++c;
        if (nc == column_numbers.end()) break;
      }

      // Process chrpos column into abspos column
      if (has_chr && data[1].size()) {
        const unsigned int chr{ref->chr_lookup[chr_name]};
        data[1].back() = ref->cn_abspos(chr, data[1].back());
      }
    }
    if (has_chr) {
      column_names.erase(column_names.begin());
      column_types.erase(column_types.begin());
      data.erase(data.begin());
    }

    // Check load state
    const uint64_t constant_size{data.front().size()};
    for (unsigned int col{0}; col != data.size(); ++col)
      if (data[col].size() != constant_size)
        throw Error("Inconsistent column data sizes");
    if (constant_size == 0) throw Error("Data load size was zero");

    if (implicit_index) {
      column_names.insert(column_names.begin(), "index");
      column_types.insert(column_types.begin(), "");
      data.emplace(data.begin(), data[0].size());
      for (uint64_t i{0}; i != data[0].size(); ++i) data[0][i] = i + 1;
    }

    // Add X jitter if requested
    if (!add_jitter) return;
    vector<double> & x_vals{data[0]};
    if (x_vals.size() < 2) return;
    double last_value{2 * x_vals[0] - x_vals[1]};
    for (uint64_t i{0}; i != x_vals.size(); ++i) {
      const double low{(x_vals[i] + last_value) / 2};
      const double next_value{i + 1 == x_vals.size() ?
            2 * x_vals[i] - x_vals[i - 1] : x_vals[i + 1]};
      const double high{(x_vals[i] + next_value) / 2};
      last_value = x_vals[i];
      x_vals[i] = std::uniform_real_distribution<double>(low, high)(mersenne);
    }
  }

  using OneColInfo = std::pair<String, String>;
  using XColInfo = std::vector<OneColInfo>;
  const XColInfo info() const {
    if (column_names.empty())
      throw Error("Unexpected empty columns in AbsposColumns::info");
    XColInfo result;
    for (unsigned int c{1}; c != column_names.size(); ++c)
      result.emplace_back(column_names[c], column_types[c]);
    return result;
  }

  const String & name(unsigned int c) const { return column_names[c]; }
  const String & type(unsigned int c) const { return column_types[c]; }
  uint64_t n_rows() const { return n_cols() ? data[0].size() : 0; }
  uint64_t n_cols() const { return data.size(); }
  const Column & operator[](const unsigned int c) const { return data[c]; }

 private:
  Strings column_names{};
  Strings column_types{};
  Data data{};
};

const std::string show_help{"Showing extra help"};

int main(int argc, char * argv[]) try {
  // Process optional command line arguments
  bool set_geometry{false};
  Geometry geometry{X11Graph::default_geometry()};
  char ** initial{nullptr};
  bool fullscreen{false};
  uint64_t n_rows{471000};  // size of trial data, to minimize extra memory
  bool x_jitter{false};
  double percent{100.0};
  bool output{false};
  double scale{2};
  unsigned char log_y{2};
  vector<double> ratio_lines{};
  std::string display_name{X11Graph::default_title};
  --argc;
  while (argc) {
    if (argv[1][0] == '-') {
      const string option{argv[1]};
      auto matches = [&option] (const std::string & full_option) {
        if (option.size() == 2 && option[1] == full_option[2]) return true;
        if (option == full_option) return true;
        return false;
      };

      if (matches("--help")) {
        throw UsageError(show_help);
      } else if (matches("--geometry")) {
        const string geometry_string{argv[2]};
        istringstream geometry_stream{geometry_string.c_str()};
        char dummy;
        int width, height, x_off, y_off;
        geometry_stream >> width >> dummy >> height >> x_off >> y_off;
        if (!geometry_stream)
          throw UsageError("Problem parsing geometry") << geometry_string;
        geometry = {{width, height}, {x_off, y_off}};
        set_geometry = true;
        argv += 2;
        argc -= 2;
      } else if (matches("--initial")) {
        initial = argv + 2;
        argc -= 5;
        argv += 5;
      } else if (matches("--threads")) {
#ifndef __CYGWIN__
        n_threads = atoi(argv[2]);
#else
        cerr << "Ignoring --threads under Cygwin OS" << endl;
#endif
        argc -= 2;
        argv += 2;
      } else if (matches("--fullscreen")) {
        fullscreen = true;
        argc -= 1;
        argv += 1;
      } else if (matches("--rows")) {
        n_rows = atoi(argv[2]);
        argc -= 2;
        argv += 2;
      } else if (matches("--jitter")) {
        x_jitter = true;
        --argc;
        ++argv;
      } else if (matches("--percent")) {
        percent = atof(argv[2]);
        if (percent <= 0 || percent > 100)
          throw UsageError("Percent value must be between 0 and 100");
        argc -= 2;
        argv += 2;
      } else if (matches("--name")) {
        display_name = argv[2];
        argc -= 2;
        argv += 2;
      } else if (matches("--setup")) {
        if (argc != 2) throw UsageError("Only two arguments allowed "
                                        "if first is --setup");
        const Reference ref{argv[2]};
        return 0;
      } else if (matches("--output")) {
        output = true;
        --argc;
        ++argv;
      } else if (matches("--scale")) {
        scale = atof(argv[2]);
        argc -= 2;
        argv += 2;
      } else if (matches("--atan")) {
        log_y = 2;
        --argc;
        ++argv;
      } else if (matches("--atanS")) {
        log_y = 3;
        --argc;
        ++argv;
      } else if (matches("--log")) {
        log_y = 1;
        --argc;
        ++argv;
      } else if (matches("--linear")) {
        log_y = 0;
        --argc;
        ++argv;
      } else if (matches("--lines")) {
        istringstream lines{argv[2]};
        double value;
        while (lines && lines >> value) {
          ratio_lines.push_back(value);
          lines.get();
        }
        argc -= 2;
        argv += 2;
      } else {
        throw UsageError("Unrecognized command line option") << option;
      }
    } else {
      break;
    }
  }
  if (!argc) throw UsageError("No default arguments were used");
  if (argc < 2) throw UsageError("At least two arguments must be used");

  const std::string first_argument{argv[1]};
  const bool do_cn{first_argument == "cn"};

  const bool do_genome{do_cn || first_argument == "genome"};
  unique_ptr<const RefCN> ref_ptr{nullptr};
  if (do_genome) {
    if (argc < 4) throw UsageError("Not enough arguments");
    ref_ptr = make_unique<const RefCN>(argv[2]);
    argc -= 2; argv += 2;
  }
  if (fullscreen && set_geometry)
    cerr << "Note that --fullscreen option overrides --geometry" << endl;

  // Columns to show
  const std::string columns{argv[1]};
  if (columns.find_first_of(',') == string::npos)
    throw UsageError("Did not find mandatory comma in column specification ")
        << columns;

  // Names of input files
  argc -= 1; argv += 1;
  const vector<string> names{[argc, argv] () {
      vector<string> result;
      result.reserve(argc);
      for (int a{0}; a != argc; ++a) result.emplace_back(argv[a + 1]);
      return result;
    }()};
  const Strings short_names{[&names] () {
      Strings result;
      for (const String & name : names)
        result.push_back(remove_including_final(name, '/'));
      return result;
    }()};

  ThreadPool pool{n_threads};

  // Read in columns from multiple data files in parallel
  using InputData = vector<AbsposColumns>;
  const InputData input_data{
    [&names, &columns, &ref_ptr, &pool, n_rows, x_jitter, percent] () {
      using Future = future<AbsposColumns>;
      vector<Future> futures;
      for (const string & name : names)
        futures.push_back(pool.run(
            [&columns, &ref_ptr, n_rows, x_jitter, percent]
            (const string & file_name) {
              return AbsposColumns{
                file_name, columns, ref_ptr.get(), n_rows, x_jitter, percent};
            }, name));
      InputData result;
      result.reserve(names.size());
      for (Future & fut : futures) result.push_back(fut.get());
      return result;
    }()};

  // App to display multiple windows
  X11App app{&pool};
  if (fullscreen)
    geometry = {{app.display_size[0], app.display_size[1]}, {0, 0}};

  // Rearrange data in specific X11Graph format for all individuals
  const int n_sets{static_cast<int>(input_data.size())};
  const int n_y{static_cast<int>(input_data.front().n_cols() - 1)};
  using XYSeries = X11Graph::XYSeries;
  using Data = X11Graph::Data;
  Data data(n_sets * n_y, XYSeries(2));
  for (int r{0}; r != n_sets; ++r)
    for (int y{0}; y != n_y; ++y) {
      data[n_sets * y + r][0] = input_data[r][0];
      data[n_sets * y + r][1] = input_data[r][y + 1];
    }
  using Info = std::pair<Strings, AbsposColumns::XColInfo>;
  using DataInfo = std::pair<Data, Info>;
  const DataInfo info{std::move(data),
        Info{short_names, input_data.front().info()}};

  auto add_special_features =
      [do_genome, do_cn, &ref_ptr, &ratio_lines, scale]
      (X11Graph & graph) {
    if (do_genome) {
      // Chromosomes and ratio lines
      graph.add_call_back("Toggle chromosome display",
                          X11Graph::CallBack{std::bind(
                              &add_chromosomes, std::cref(*ref_ptr), _1, _2)});

      // Gene display
      graph.add_call_back(
          "Toggle chrpos and gene display (only shown for X axis range below " +
          std::to_string(max_gene_mb) + "MB for genes and " +
          std::to_string(max_name_mb) + " MB for names)",
          X11Graph::CallBack{std::bind(
              &add_genes, std::cref(*ref_ptr), _1, _2)});

      // Cytobands
      graph.add_call_back("Toggle cytobands and names",
                          X11Graph::CallBack{std::bind(
                              &add_cytobands, std::cref(*ref_ptr), _1, _2)},
                          true, false);

      // Other genome-specific tweaks
      graph.grid_radios[0][0].toggled(false);
      graph.grid_radios[1][0].toggled(false);
      graph.log_radios[0].actions.visible = [] () { return false; };
      graph.coord_radio.description +=
      " or search (enter text query while pointer is in this radio control)";
    }

    if (do_cn) {
      graph.y_cn_scale = scale;
      const vector<double> cn_lines{[&ratio_lines]() {
          if (ratio_lines.size()) {
            return ratio_lines;
          } else {
            vector<double> result{
              0.0, 1.0, 2.0, 3.0, 4.0, 5.0, 10, 100};
            // for (double & line : result) line /= graph.y_cn_scale;
            return result;
          }
        }()};

      graph.add_call_back("Toggle ratio lines", X11Graph::CallBack{std::bind(
          &add_ratio_lines, cn_lines, _1, _2)}, false, true,
        []() { return true; /* !graph.tiled_radio; */ });

      graph.grid_radios[1][1].toggled(false);
      graph.grid_radios[0][1].toggled(false);
      graph.tick_radios[1].toggled(true);
    }
  };

  // All individuals graph
  X11Graph & graph{app.create<X11Graph>(info, geometry, display_name,
                                        add_special_features)};

  // Preload gene info to avoid wait time after zoom
  future<bool> gene_future;
  if (do_genome) gene_future = pool.run(
          add_genes, std::cref(*ref_ptr), std::ref(graph), Event());

  // Process initial view command line arguments
  if (log_y) {
    graph.log_radios[1].state(log_y);
    graph.prepare_log();
  }
  if (initial) {
    graph.get_range();
    graph.set_range(0, atof(initial[0]), atof(initial[1]));
    if (string(initial[2]) != "X")
      graph.set_range(1, atof(initial[2]), atof(initial[3]));
  }
  if (initial || log_y) graph.prepare();

  // Just print the initial view and quit
  graph.output(output);

  // Run the app
  app.run();

  if (gene_future.valid()) gene_future.get();

  return 0;
} catch (UsageError & e) {
  const std::string err{e.what()};
  if (err != show_help) {
    std::cerr << "Usage Error: ";
    cerr << e.what() << "\n\n";
  }

  std::string usage{std::string() + R"xxx(Usage options:

    ggraph [genome|cn ref_fasta] xn,yn1,yn2... data_file ...

Optional leading arguments (CAPS for numeric):
  -g | --geometry WIDTHxHEIGHT+XOFF+YOFF
  -i | --initial XLOW XHIGH YLOW YHIGH
  -t | --threads NTHREADS
  -n | --name graph_title_word
  -r | --rows NROWS
  -f | --fullscreen
  -s | --setup ref_fasta
  -j | --jitter
  -p | --percent NN
  -a | --atan
  -l | --log
       --linear
       --lines Y1,Y2,Y3...
  -o | --output
  -h | --help

Use optional leading argument --help to display additional usage information
or visit http://mumdex.com/ggraph/ to view the G-Graph tutorial)xxx"};

  std::cerr << usage << std::endl;

  if (err != show_help) return 1;

  std::cerr << R"xxx(Finer usage details:
  1. 'genome' as the first argument includes genome features in plots
  2. 'cn' as the first argument also includes copy number ratio lines in plots
  3. ref_fasta is the name of (or full path to) the genome reference fasta file
  4. Without 'genome' or 'cn' as the first argument, the graph is a plain plot
     and the command requires the shorter usage form shown above
  5. For the column specification 'xn,yn1,yn2...':
    a. It is a list of columns to display from the data files
    b. Starts with the single x variable, followed by y variables
    c. Can be comma separated column numbers if the data file has no header line
    d. In cn mode, by default the second of two y columns is displayed as lines
    e. After any 'yn', ':l' forces line display while ':p' forces point display
    f. If 'xn' is 'implicit' then the x variable is the line number
    g. If 'xn' is one of 'chr', 'chrom' or 'chromosome', or is followed by ':c',
       then that column is assumed to be a chromosome name column and the 'y1'
       column is interpreted as chromosomal position to calculate abspos
    h. An abspos column as 'xn' will work, provided it is calculated correctly.
       G-Graph calculates abspos in the order specified by ref_fasta,
       ignoring chromosomes less than 40 megabases in size.
  6. The data_file arguments are names of (or full paths to) the data files,
     which must be space or tab separated text tables with an optional header.
     Each text file may be a different length but all must share the column
     names (possibly in different order) specified in the column specification.
     Referenced columns (except for the optional chr column) must be numeric.
     Columns after the referenced columns may be ragged.
  7. For the optional argument '--geometry', a '+' before 'XOFF' or 'YOFF'
     attempts window placement relative to the upper left of screen,
     while a '-' attempts placement relative to the bottom right screen corner
  8. The --initial option specifies the initial view to show
  9. The --threads option uses up to that number for file load and display
  10. The --name option specifies a plot title for viewing and saving
  11. The --rows option reserves space for input files of that dimension
  12. The --fullscreen option only tries to make a maximum window size 
  13. The --setup option allows binary reference cache pre-generation, but
      that would happen anyway on first loading. This option was added only to
      reduce first load wait time when using the G-Graph install script
  14. The --jitter option adds a little randomness to loaded X coordinates
  15. The --percent option only loads a fraction of the data, randomly
  16. The --log option sets log mode initially, overriding any config file
  17. The --linear option sets log mode initially, overriding any config file
  18. The --lines option specifies alternative ratio lines for cn mode only
  19. The --output option saves the initial view and then exits
  20. The --help option displays this text and then exits
)xxx"
            << std::endl;

  return 0;
} catch (Error & e) {
  cerr << "paa::Error:" << endl;
  cerr << e.what() << endl;
  return 1;
} catch (exception & e) {
  cerr << "std::exception:" << endl;
  cerr << e.what() << endl;
  return 1;
} catch (...) {
  cerr << "unknown exception was caught" << endl;
  return 1;
}
```
