## Supplementary material for "G-Graph: An interactive genomic graph viewer": MUMdex Source Code for G-Graph: ggraph_paper_figure.sh

```
#! /bin/bash

#
# ggraph_paper_figure.sh
#
# Allows a user to reproduce the specific figure used in the ggraph paper
# 
# Assumes that mumdex code is compiled and available in ./mumdex,
# the hg19 genome is located at ./hg19.fa,
# that gene and cytoband definition files are in ./hg19.fa.bin/ 
# and that the sample files {m,f,d,s}.txt are in the current directory.
# The gene definition, cytoband definition and sample files are available at 
# http://mumdex.com/ggraph/
#

n_threads=2
uname=$(uname)
if [ $uname = "Linux" ] ; then
    n_threads=$(grep processor /proc/cpuinfo | wc -l)
fi
if [ $uname = "Darwin" ] ; then
    n_threads=$(sysctl -n hw.logicalcpu)
fi

abspos_left=2983775000
geometry=2400x1300+0+0

(cd ./mumdex ; make -j $n_threads ggraph > /dev/null) &&
(
echo 'Note: to exactly reproduce the figure from the paper:'
echo 'you need to run on a mac, to get the exact fonts and gui used in the paper'
echo "you need a big enough screen to contain the image (${geometry%%+*} pixels min usable)"
echo 'you need to manually turn on cytoband display (near top left)'
echo 'you need to manually increase point display size by three levels (near bottom right)'
echo 'you need to manually increase line display size by two levels (near bottom right)'
echo 'have the mouse focus remain in the window, in the central graphing region'
echo 'take a window-screenshot: command-shift-4 space, then click on window'
) &&
./mumdex/ggraph --linear --threads $n_threads --geometry $geometry --initial $abspos_left $((abspos_left+1000000)) 0 2.5 cn ./hg19.fa abspos,ratio,seg {m,f,d,s}.txt
```
