## Supplementary material for "G-Graph: An interactive genomic graph viewer": MUMdex Source Code for G-Graph: mumdex_cn.cpp

```
//
// mumdex_cn.cpp
//
// Copy number from a mumdex file
//
// Copyright 2016 Peter Andrews @ CSHL
//

#include <algorithm>
#include <exception>
#include <iostream>
#include <sstream>
#include <string>
#include <vector>

#include "cn.h"
#include "error.h"
#include "files.h"
#include "mumdex.h"
#include "threads.h"
#include "utility.h"

using std::cout;
using std::cerr;
using std::cref;
using std::endl;
using std::exception;
using std::istringstream;
using std::max;
using std::min;
using std::string;
using std::to_string;
using std::vector;

using paa::copy_number;
using paa::Bin;
using paa::CN_abspos;
using paa::Error;
using paa::MappedVector;
using paa::PosInfo;
using paa::Progress;
using paa::Reference;
using paa::ThreadPool;

// using paa::MUMdex;
using MUMdex = paa::MemoryMUMdex;
// using Reference = paa::MemoryReference;  // MUMdex::Reference;
using Mappability = paa::MemoryMappability;

int main(int argc, char * argv[]) try {
  if (--argc != 6)
    throw Error("usage: mumdex_cn mumdex bins bad sample n_threads title");

  paa::set_cn_parameters();

  const bool create_pdf{false};
  cerr << "Loading mumdex" << endl;
  const MUMdex mumdex{argv[1]};
  const string bins_string{argv[2]};
  const MappedVector<unsigned char> bad{argv[3]};
  const unsigned int sample{static_cast<unsigned int>(atoi(argv[4]))};
  if (!sample) throw Error("Sample must be a nonzero integer");
  const unsigned int n_threads{static_cast<unsigned int>(atoi(argv[5]))};
  const string title{argv[6]};

  cerr << "Loading reference and mappability" << endl;
  const Reference & ref{mumdex.reference()};
  const CN_abspos cn_abspos{ref};
  const Mappability mappability{ref};

  cerr << "Loading bins" << endl;
  using AllBins = vector<vector<Bin> >;
  const AllBins bins{[&bins_string, &ref]() {
      vector<string> bins_names;
      istringstream bins_stream{bins_string.c_str()};
      string bins_name;
      while (getline(bins_stream, bins_name, ','))
        bins_names.push_back(bins_name);
      AllBins result(bins_names.size());
      for (unsigned int n{0}; n != bins_names.size(); ++n)
        result[n] = load_bins(bins_names[n], ref);
      return result;
    }()};

  cerr << "Filtering mappings from " << mumdex.n_pairs()
       << " pairs and assigning to bins" << endl;

  ThreadPool pool{n_threads};
  ThreadPool::Results<vector<vector<unsigned int>>> results;
  const uint64_t block_size{max(static_cast<uint64_t>(1000),
                                mumdex.n_pairs() / n_threads / 10)};

  auto block_fun = [&mumdex, &mappability, &cn_abspos, &bad, &bins,
                    block_size, sample] (const uint64_t b) {
    try {
      vector<vector<unsigned int>> counts(bins.size());
      for (unsigned int i{0}; i != bins.size(); ++i)
        counts[i].resize(bins[i].size());
      for (uint64_t p{b}; p < min(b + block_size, mumdex.n_pairs());
           p += sample) {
        const unsigned int abspos{pair_cn_abspos(mumdex, mappability,
                                                 cn_abspos, p)};
        if (abspos >= cn_abspos.bad_abspos()) continue;
        if (bad[abspos]) continue;
        for (unsigned int i{0}; i != bins.size(); ++i) {
          vector<Bin>::const_iterator found{
            upper_bound(bins[i].begin(), bins[i].end(), abspos)};
          if (found-- != bins[i].begin()) {
            if (abspos >= found->abspos_start() &&
                abspos < found->abspos_start() + found->length())
              ++counts[i][found - bins[i].begin()];
          }
        }
      }
      return counts;
    } catch (Error & e) {
      cerr << "paa::Error:" << endl;
      cerr << e.what() << endl;
      throw;
    } catch (exception & e) {
      cerr << "std::exception" << endl;
      cerr << e.what() << endl;
      throw;
    } catch (...) {
      cerr << "unknown exception was caught" << endl;
      throw;
    }
  };

  for (uint64_t b_{0}; b_ < mumdex.n_pairs(); b_ += block_size)
    pool.run(results, block_fun, b_);

  Progress progress{results.size(), 0.01, "Binning"};
  vector<vector<unsigned int>> counts(bins.size());
  for (unsigned int i{0}; i != bins.size(); ++i)
    counts[i].resize(bins[i].size());
  while (results.size()) {
    const vector<vector<unsigned int>> block_counts{results.get()};
    for (unsigned int i{0}; i != bins.size(); ++i)
      for (unsigned int b{0}; b != counts[i].size(); ++b)
        counts[i][b] += block_counts[i][b];
    progress();
  }

  const bool minimal{true};
  ThreadPool::Results<void> cn_results;
  for (unsigned int i{0}; i != bins.size(); ++i)
    pool.run(cn_results, copy_number, cref(ref), cref(bins[i]), cref(counts[i]),
             title + " " + to_string(bins[i].size()) + " bins", create_pdf,
             minimal, 0.05, 1.0, 3);
  while (cn_results.size()) cn_results.get();

  cerr << "Done" << endl;

  return 0;
} catch (Error & e) {
  cerr << "paa::Error:" << endl;
  cerr << e.what() << endl;
  return 1;
} catch (exception & e) {
  cerr << "std::exception" << endl;
  cerr << e.what() << endl;
  return 1;
} catch (...) {
  cerr << "unknown exception was caught" << endl;
  return 1;
}
```
