## Supplementary material for "G-Graph: An interactive genomic graph viewer": MUMdex Source Code for G-Graph: mumdex_finebin.cpp

```
//
// mumdex_finebin
//
// finely bin mappings from one or more mumdex files
//
// Copyright 2016 Peter Andrews @ CSHL
//

#include <exception>
#include <iostream>
#include <string>

#include "cn.h"
#include "error.h"
#include "mumdex.h"
#include "population.h"

using std::cerr;
using std::cout;
using std::endl;
using std::exception;
using std::string;

using paa::saved_ref_name;
using paa::Error;
using paa::Family;
using paa::FinestBins;
using paa::Mappability;
using paa::Population;
using paa::Reference;
using paa::Sample;

using MUMdex = paa::MemoryMUMdex;

int main(int argc, char* argv[])  try {
  // Check usage
  if (--argc < 4) {
    throw Error("usage: mumdex_finebin samples_dir pop_file out_name "
                "[sample|family] ...");
  }

  paa::set_cn_parameters();

  // Process command line arguments
  const string samples_dir{argv[1]};
  const Population pop{argv[2]};
  const string out_name{argv[3]};
  argc -= 3;
  argv += 3;

  // Load reference information
  const string first_sample_name{pop.sample(pop.samples(argv[1]).front())};
  const Reference ref{saved_ref_name(
      samples_dir + "/" + first_sample_name + "/mumdex")};
  const Mappability mappability{ref};
  FinestBins bins{ref};

  while (argc--) {
    // Get samples for sample or family name
    const string sample_or_family{argv++[1]};
    for (const Sample sample : pop.samples(sample_or_family)) {
      // Get sample info
      const Family family{pop.family(sample)};
      const string sample_name{pop.sample(sample)};
      const string family_name{pop.family(family)};

      cerr << "Processing " << sample_name << " " << pop.member(sample) << endl;

      // Load the mumdex for the sample
      const MUMdex mumdex{samples_dir + "/" + sample_name + "/mumdex", &ref};
      bins.add(mumdex, mappability, pop.nX(sample), pop.nY(sample), 12);
    }
  }
  cerr << "Saving output to " << out_name << endl;
  bins.save(out_name);

  cerr << "Done" << endl;

  return 0;
} catch (Error & e) {
  cerr << "paa::Error:" << endl;
  cerr << e.what() << endl;
  return 1;
} catch (exception & e) {
  cerr << "std::exception" << endl;
  cerr << e.what() << endl;
  return 1;
} catch (...) {
  cerr << "unknown exception was caught" << endl;
  return 1;
}
```
