## Supplementary material for "G-Graph: An interactive genomic graph viewer": MUMdex Source Code for G-Graph: pop_cn.cpp

```
//
// pop_cn.cpp
//
// Examine cn for a population, and call denovos
//
// Copyright 2018 Peter Andrews @ CSHL
//

#include <gsl/gsl_cdf.h>

#include <algorithm>
#include <exception>
#include <fstream>
#include <iostream>
#include <limits>
#include <map>
#include <sstream>
#include <string>
#include <thread>
#include <utility>
#include <vector>

#include "cn.h"
#include "error.h"
#include "files.h"
#include "genes.h"
#include "mumdex.h"
#include "population.h"
#include "psplot.h"
#include "stats.h"
#include "strings.h"
#include "threads.h"
#include "utility.h"

using std::cin;
using std::cout;
using std::cerr;
using std::cref;
using std::endl;
using std::exception;
using std::flush;
using std::ifstream;
using std::lock_guard;
using std::map;
using std::move;
using std::mutex;
using std::numeric_limits;
using std::ofstream;
using std::ostream;
using std::ostringstream;
using std::pair;
using std::string;
using std::to_string;
using std::vector;

using paa::commas;
using paa::doc_defaults;
using paa::mkdir;
using paa::nunset;
using paa::unset;
using paa::Bin;
using paa::Bounds;
using paa::ChromosomeIndexLookup;
using paa::CN_abspos;
using paa::CN_Bin;
using paa::CN_Bins;
using paa::Error;
using paa::GeneXrefs;
using paa::KnownGenes;
using paa::MAD;
using paa::Marker;
using paa::Population;
using paa::Progress;
using paa::PSDoc;
using paa::PSGraph;
using paa::PSHSeries;
using paa::PSPage;
using paa::PSXYSeries;
using paa::Reference;
using paa::Sample;
using paa::ThreadPool;

// xxxxxxxxxxxxxxxxxxxxxxxxxxxxxxxxxxxxxxxxxxxxxxxxxxxxxxxxxxxxxxxxxxxxxxxxxxxxx
ostream & separator(ostream & stream) {
  return stream
      << "\n"
      << "-\t-\t-\t-\t-\t-\t-\t-\t-\t-\t-\t-\t-\t-\t-\t-\t-\t-\t-\t-\t-"
      << "\n\n";
}

bool is_par(const Bin & bin, const unsigned int chrX) {
  const unsigned int par_coords[2][2]{{60000, 2699520},
                                      {154931043, 155260560}};
  if (bin.chromosome() != chrX) return false;
  if (bin.stop_position() <= par_coords[0][0] ||
      bin.start_position() >= par_coords[1][1] ||
      (bin.start_position() >= par_coords[0][1] &&
       bin.stop_position() <= par_coords[1][0])) {
    return false;
  } else {
    return true;
  }
}

double get_autosome_average(const vector<Bin> & bins,
                            const CN_Bins & profile,
                            const unsigned int chrX) {
  uint64_t autosome_n_bins{0};
  double autosome_total_count{0};
  for (unsigned int b{0}; b != profile.size(); ++b) {
    const Bin & bin{bins[b]};
    const CN_Bin & bin_counts{profile[b]};
    if (bin.chromosome() >= chrX) break;
    ++autosome_n_bins;
    autosome_total_count += bin_counts.norm_count();
  }
  return autosome_total_count / autosome_n_bins / 2;
}

unsigned char get_expected_copy(
    const unsigned int chr, const bool is_male,
    const unsigned int chrX, const unsigned int chrY) {
  const bool is_x{chr == chrX};
  const bool is_y{chr == chrY};
  if (is_male && (is_x || is_y)) {
    return 1;
  } else if (!is_male && is_y) {
    return 0;
  } else {
    return 2;
  }
}

std::vector<unsigned char> get_expected_copy(
    const Reference & ref, const bool is_male,
    const unsigned int chrX, const unsigned int chrY) {
  std::vector<unsigned char> result;
  for (unsigned int c{0}; c != ref.n_chromosomes(); ++c)
    result.push_back(get_expected_copy(c, is_male, chrX, chrY));
  return result;
}

double score(const double prob) {
  return static_cast<uint64_t>(
      1000 * fabs(log10(std::max(prob, numeric_limits<double>::min())))) /
      1000.0;
}

const char * const call_names[]{
  "norm", "loss", "gain",
      "trans_loss", "trans_gain", "denovo_loss", "denovo_gain"
      };

class Segment {
 public:
  enum Call { Norm, Loss, Gain };
  Segment(const vector<Bin> &bins,
          const CN_Bins & profile,
          const bool is_male,
          const unsigned int chrX,
          const unsigned int chrY,
          const unsigned int start__,
          const unsigned int stop__,
          const Call null_call = Norm) :
      start_{start__},
    stop_{stop__},
    copy_1_count_{get_autosome_average(bins, profile, chrX)},
    expected_fcopy_{get_expected_fcopy(bins, is_male, chrX, chrY)},
    count_{0},
    ratio_{profile[start_].seg_ratio()} {
      if (bins[start_].chromosome() != bins[stop_ - 1].chromosome())
        throw Error("Cross-chromosome bin");

      for (unsigned int b{start_}; b != stop_; ++b)
        count_ += profile[b].norm_count();

      call_ = fcopy() > expected_fcopy_ + 0.5 ? Gain :
          (fcopy() < expected_fcopy_ - 0.5 ? Loss : Norm);

      const bool gain_side{null_call == Norm ?
            fcopy() > expected_fcopy_ : null_call == Gain};
      constexpr uint64_t one{1};
      if (gain_side) {
        const uint64_t null_count{static_cast<uint64_t>(
            expected_count(expected_fcopy_ + 0.5) + 0.5)};
        p_null_ = gsl_cdf_poisson_Q(count_, std::max(one, null_count - 1));
        p_alt_ = gsl_cdf_poisson_P(count_, std::max(one, null_count));
      } else {
        const uint64_t null_count{static_cast<uint64_t>(
            expected_count(expected_fcopy_ - 0.5) + 0.5)};
        p_null_ = gsl_cdf_poisson_P(count_, std::max(one, null_count));
        p_alt_ = gsl_cdf_poisson_Q(count_, std::max(one, null_count - 1));
      }
    }

  double get_expected_fcopy(
      const vector<Bin> & bins, const bool is_male,
      const unsigned int chrX, const unsigned int chrY) const {
    double result{0.0};
    for (unsigned int b{start_}; b != stop_; ++b)
      result += is_par(bins[b], chrX) ? 2 :
          get_expected_copy(bins[b].chromosome(), is_male, chrX, chrY);
    result /= n_bins();
    return result;
  }

  unsigned int start() const { return start_; }
  unsigned int stop() const { return stop_; }
  double copy_1_count() const { return copy_1_count_; }
  double expected_fcopy() const { return expected_fcopy_; }
  unsigned int count() const { return count_; }
  double ratio() const { return ratio_; }
  Call call() const { return call_; }
  double p_null() const { return p_null_; }
  double p_alt() const { return p_alt_; }

  unsigned int n_bins() const { return stop_ - start_; }
  double min_count() const { return n_bins() * copy_1_count_ / 100; }
  double expected_count(double fcopy__ = -1) const {
    if (fcopy__ < 0) fcopy__ = expected_fcopy_;
    const double expected{n_bins() * fcopy__ * copy_1_count_};
    return std::max(expected, 1 + min_count());
  }
  unsigned int expected_copy() const {
    return static_cast<unsigned int>(expected_fcopy() + 0.5);
  }
  double fcopy() const { return count_ / (copy_1_count_ * n_bins()); }
  unsigned int copy() const {
    return static_cast<unsigned int>(fcopy() + 0.5);
  }
  bool is_event() const { return !(call_ == Norm); }
  bool is_gain() const { return call_ == Gain; }
  bool is_loss() const { return call_ == Loss; }
  bool is_good(const double p_val = 0.05) const {
    return p_null_ < p_val;
  }
  string all_string() const { return call_names[call()]; }
  double score() const { return ::score(p_null_); }

  unsigned int start_;
  unsigned int stop_;
  double copy_1_count_;
  double expected_fcopy_;
  double count_;
  double ratio_;
  Call call_{Norm};
  double p_null_{0};
  double p_alt_{0};
};

bool operator<(const Segment & lhs, const Segment & rhs) {
  if (lhs.is_event() == rhs.is_event()) {
    return rhs.score() < lhs.score();
    if (lhs.call() == rhs.call()) {
      return rhs.score() < lhs.score();
    } else {
      return lhs.call() < rhs.call();
    }
  } else {
    return rhs.is_event() < lhs.is_event();
  }
}

class Segments {
 public:
  Segments() = default;
  Segments(Segments && other) = default;
  Segments(Segments & other) = delete;
  Segments & operator=(const Segments &) = delete;
  Segments & operator=(Segments &&) = default;
  Segments(const Reference & ref,
           const vector<Bin> & bins,
           const CN_Bins & profile,
           const bool is_male) {
    const unsigned int chrX{ref.find_x_chromosome()};
    const unsigned int chrY{ref.find_y_chromosome()};
    const std::vector<unsigned char> expected_copy{
      get_expected_copy(ref, is_male, chrX, chrY)};

    unsigned int segment_start{0};
    for (unsigned int b{0}; b <= bins.size(); ++b) {
      if (b == bins.size() ||
          bins[b].chromosome() != bins[segment_start].chromosome() ||
          profile[b].seg_ratio() < profile[segment_start].seg_ratio() ||
          profile[b].seg_ratio() > profile[segment_start].seg_ratio()) {
        segments_.emplace_back(
            bins, profile, is_male, chrX, chrY, segment_start, b);
        segment_start = b;
        if (b == bins.size()) break;
      }
    }
  }

  vector<Segment> & segments() { return segments_; }
  const Segment & back() const { return segments_.back(); }


  const Segment & operator[](const uint64_t index) const {
    return segments_[index];
  }

  uint64_t size() const { return segments_.size(); }
  vector<Segment>::const_iterator begin() const { return segments_.begin(); }
  vector<Segment>::const_iterator end() const { return segments_.end(); }

 private:
  vector<Segment> segments_{};
};

struct SampleInfo {
  SampleInfo(const Population & pop_, const Reference & ref_,
             const vector<Bin> & bins_, const Sample sample_) :
      pop{&pop_}, ref{&ref_}, bins{&bins_}, sample{sample_} {
        if (profile.size() != bins->size())
          throw Error("Bin size mismatch")
              << input_name("results")
              << bins->size()
              << profile.size();
      }

  SampleInfo(const SampleInfo &) = delete;
  SampleInfo(SampleInfo &&) = default;
  SampleInfo & operator=(const SampleInfo &) = delete;
  SampleInfo & operator=(SampleInfo &&) = default;
  const Population * pop;
  const Reference * ref;
  unsigned int chrX{ref->find_x_chromosome()};
  unsigned int chrY{ref->find_y_chromosome()};
  const vector<Bin> * bins;
  Sample sample;
  CN_Bins profile{input_name("results")};
  bool is_male{pop->sex(sample) == "XY"};
  Segments segments{*ref, *bins, profile, is_male};

  bool operator<(const SampleInfo & rhs) const {
    return sample < rhs.sample;
  }
  string input_name(const string type) const {
    ostringstream result;
    const string sample_name{pop->sample(sample)};
    result << sample_name << "/" << sample_name
           << "_" << bins->size() << "_bins_" + type + ".txt";
    return result.str();
  }
  static SampleInfo make_sample_info(
      const Population & pop_,
      const Reference & ref_,
      const vector<Bin> & bins_,
      const Sample sample_) {
    return SampleInfo{pop_, ref_, bins_, sample_};
  }
};
using AllSampleInfo = vector<SampleInfo>;

using GeneInfo = pair<string, bool>;
using GeneInfos = vector<GeneInfo>;
using Strings = vector<string>;
using OutResult = pair<Strings, GeneInfos>;

class PopSegment {
 public:
  enum Call {
    Norm = Segment::Norm,
    Loss = Segment::Loss,
    Gain = Segment::Gain,
    TransLoss,
    TransGain,
    DenovoLoss,
    DenovoGain
  };
  PopSegment(const Sample sample_,
             const AllSampleInfo & all_info,
             const Segment & segment_) :
      sample{sample_}, segment{segment_} {
    const Population & pop{*all_info.front().pop};
    for (unsigned int p{0}; p != pop.n_samples(); ++p) {
      const Sample pop_sample{p};
      const SampleInfo & info{all_info[p]};
      segments.emplace_back(
          *info.bins, info.profile, info.is_male, info.chrX, info.chrY,
          segment.start(), segment.stop(), segment.call());
      const Segment & pop_segment{segments.back()};
      if (pop.family(pop_sample) == pop.family(sample)) {
        if (pop.is_parent(pop_sample)) {
          (pop.is_mother(pop_sample) ? mother_call : father_call) =
              pop_segment.call();
          (pop.is_mother(pop_sample) ? mother_null_prob : father_null_prob) =
              pop_segment.p_null();
          (pop.is_mother(pop_sample) ? mother_alt_prob : father_alt_prob) =
              pop_segment.p_alt();
          (pop.is_mother(pop_sample) ? mother_copy : father_copy) =
              pop_segment.fcopy();
        }
      } else {
        const bool is_sex_chr{
          (*info.bins)[segment.start()].chromosome() == info.chrX ||
              (*info.bins)[segment.start()].chromosome() == info.chrY};
        ++pop_size;
        if (pop_segment.call() == segment.call()) ++pop_n;
        if (!is_sex_chr || info.is_male == all_info[sample].is_male) {
          ++pop_sex_size;
          fcopies.push_back(pop_segment.fcopy());
          if (segment.fcopy() >= pop_segment.fcopy()) ++pop_rank;
        }
      }
    }
    sort(fcopies.begin(), fcopies.end());
    const MAD mad{fcopies};
    pop_mad = mad.mad();
    pop_prob = segment.call() == Segment::Gain ?
        gsl_cdf_gaussian_Q(segment.fcopy() - mad.median(), mad.stdev()) :
        gsl_cdf_gaussian_P(segment.fcopy() - mad.median(), mad.stdev());
    fcopies.push_back(mother_copy);
    fcopies.push_back(father_copy);
    fcopies.push_back(segment.fcopy());
    sort(fcopies.begin(), fcopies.end());
  }

  string call_string() const { return call_names[call()]; }
  Call call() const {
    return static_cast<Call>(static_cast<int>(
        is_trans() ? TransLoss :
        (is_denovo() ? DenovoLoss :
         (segment.is_event() ? Loss :
          Norm))) + static_cast<int>(segment.is_gain()));
  }

  bool is_denovo() const {
    return segment.is_event() && segment.is_good() &&
        not_trans_prob() < 0.05 && pop_prob < 0.2;
  }
  bool is_trans() const {
    return segment.is_event() &&
        (mother_call == segment.call() ||
         father_call == segment.call() ||
         fabs(segment.fcopy() - mother_copy) < 0.25 ||
         fabs(segment.fcopy() - father_copy) < 0.25 ||
         trans_prob() < 0.75);
  }

  unsigned int call_class() const {
    return call() == Norm ? 0 :
        (call() >= DenovoLoss ? 3 :
         (call() >= TransLoss ? 1 : 2));
  }

  bool operator<(const PopSegment & rhs) const {
    const unsigned int lhs_class{call_class()};
    const unsigned int rhs_class{rhs.call_class()};
    if (lhs_class == rhs_class) {
      return segment.p_null() < rhs.segment.p_null();
    } else {
      return rhs_class < lhs_class;
    }
  }
  OutResult out(const SampleInfo & info,
                const KnownGenes & genes, const GeneXrefs & xref) const {
    const Population & pop{*info.pop};
    const Reference & ref{*info.ref};
    const vector<Bin> & bins{*info.bins};
    const unsigned int chr{bins[segment.start()].chromosome()};
    const unsigned int start{bins[segment.start()].start_position()};
    const unsigned int stop{bins[segment.stop() - 1].stop_position()};

    // get gene info for segment
    ostringstream gene_info;
    const KnownGenes::GeneOverlaps gene_overlaps{
      genes.find_genes(chr, start, stop)};
    map<string, unsigned int> named_overlaps;
    for (const KnownGenes::GeneOverlap & overlap : gene_overlaps) {
      const string symbol{xref[genes[overlap.first].name].geneSymbol};
      const unsigned int old_value{named_overlaps[symbol]};
      if (old_value < overlap.second) {
        named_overlaps[symbol] = overlap.second;
      }
    }
    using Overlap = pair<string, unsigned int>;
    vector<Overlap> overlaps(named_overlaps.begin(), named_overlaps.end());
    sort(overlaps.begin(), overlaps.end(),
         [](const Overlap & lhs, const Overlap & rhs) {
           if (lhs.second == rhs.second) {
             return lhs.first < rhs.first;
           } else {
             return lhs.second > rhs.second;
           }
         });
    unsigned int genes_with_exons{0};
    unsigned int total_exons{0};
    for (unsigned int o{0}; o != overlaps.size(); ++o) {
      genes_with_exons += static_cast<bool>(overlaps[o].second);
      total_exons += overlaps[o].second;
    }
    GeneInfos gene_infos;
    for (unsigned int o{0}; o != overlaps.size(); ++o) {
      const Overlap & overlap{overlaps[o]};
      if (o) gene_info << ",";
      if (overlap.second)
        gene_info << overlap.second << "-exon"
                  << (overlap.second > 1 ? "s" : "") << "-";
      gene_info << overlap.first;
      gene_infos.emplace_back(overlap.first, static_cast<bool>(overlap.second));
    }
    if (overlaps.empty()) {
      gene_info << "intergenic";
    }

    vector<string> output{
      // who
      "sample", pop.sample(sample),
          "member", pop.member(sample),
          "sex", pop.sex(sample),
          "family", pop.family(pop.family(sample)),
          // where
          "chr", ref.name(chr),
          "start", to_string(start),
          "stop", to_string(stop),
          "size", commas(stop - start),
          "n_bins", to_string(segment.stop() - segment.start()),
          // genes
          "n_genes", to_string(overlaps.size()),
          "n_exon_genes", to_string(genes_with_exons),
          "n_exons", to_string(total_exons),

          // call
          "call", call_string(),
          "mom_call", call_names[mother_call],
          "dad_call", call_names[father_call],
          // scores
          "denovo_score", to_string(denovo_score()),
          "cn_score", to_string(segment.score()),
          "trans_score", to_string(trans_score()),
          "not_trans_score", to_string(not_trans_score()),
          "pop_score", to_string(pop_score()),
          // pop
          "pop_n", to_string(pop_n),
          "pop_rank", to_string(1.0 * pop_rank / pop_sex_size),
          "pop_mad", to_string(pop_mad),
          "pop_p", to_string(pop_prob),

          // values
          "copy", to_string(segment.fcopy()),
          "exp_copy", to_string(segment.expected_fcopy()),
          "mom_copy", to_string(mother_copy),
          "dad_copy", to_string(father_copy),
          "count", to_string(static_cast<uint64_t>(segment.count())),
          "seg_copy", to_string(2 * segment.ratio()),
          // probs
          "denovo_p", to_string(denovo_prob()),
          "cn_p", to_string(segment.p_null()),
          "mom_p", to_string(mother_null_prob),
          "dad_p", to_string(father_null_prob),
          "mom_alt_p", to_string(mother_alt_prob),
          "dad_alt_p", to_string(father_alt_prob),

          // gene list
          "genes", gene_info.str()};

    return OutResult{output, gene_infos};
  }

  uint64_t pop_size{0};
  uint64_t pop_sex_size{0};
  Sample mother{0};
  Sample father{0};
  Segment::Call mother_call{Segment::Norm};
  Segment::Call father_call{Segment::Norm};
  double mother_copy{0};
  double father_copy{0};
  double mother_null_prob{0};
  double father_null_prob{0};
  double mother_alt_prob{0};
  double father_alt_prob{0};
  double trans_prob() const {
    return std::min(mother_null_prob, father_null_prob);
  }
  double not_trans_prob() const {
    return std::max(mother_alt_prob, father_alt_prob);
  }
  double trans_score() const { return score(trans_prob()); }
  double not_trans_score() const { return score(not_trans_prob()); }

  double pop_prob{0};
  double pop_mad{0};
  uint64_t pop_n{0};
  uint64_t pop_rank{0};
  double pop_score() const { return score(pop_prob); }

  double denovo_prob() const {
    return std::max(segment.p_null(), not_trans_prob());
  }
  double denovo_score() const { return score(denovo_prob()); }

  Sample sample;
  Segment segment;
  vector<double> fcopies{};

 private:
  vector<Segment> segments{};
};
using PopSegments = vector<PopSegment>;

const char * const colors[3]{"1 0 0", "0 0 1", "0 1 0"};
// const char * const members[3]{"M", "F", "P"};

int main(int argc, char* argv[]) try {
  if (--argc != 5)
    throw Error("usage: pop_cn ref family_file bin_file out_dir n_threads");

  // Process arguments
  const Reference ref{argv[1]};
  const ChromosomeIndexLookup lookup{ref};
  const CN_abspos cn_abspos{ref};
  const Population pop{argv[2]};
  const string bins_name{argv[3]};
  const string out_dir{argv[4]};
  const unsigned int n_threads{static_cast<unsigned int>(atoi(argv[5]))};
  ThreadPool pool{n_threads};
  const vector<Bin> bins{load_bins(bins_name, ref)};

  // Gene info
  const string reference_file{ref.fasta_file()};
  const string genes_name{reference_file + ".bin/knownGene.txt"};
  const string isoforms_name{reference_file + ".bin/knownIsoforms.txt"};
  const string kgXrefs_name{reference_file + ".bin/kgXref.txt"};
  const KnownGenes genes{genes_name, isoforms_name, lookup, ref};
  const GeneXrefs xref{kgXrefs_name};

  // load sample information in parallel
  const AllSampleInfo all_sample_info{
    [&ref, &pop, &bins, &pool]() {
      // run
      ThreadPool::Results<SampleInfo> results;
      for (unsigned int s{0}; s != pop.n_samples(); ++s)
        pool.run(results,
                 SampleInfo::make_sample_info,
                 cref(pop), cref(ref), cref(bins), Sample(s));
      // get
      AllSampleInfo result;
      result.reserve(pop.n_samples());
      Progress progress{results.size(), 0.001, "get sample info"};
      while (results.size()) {
        result.push_back(results.get());
        progress();
      }
      sort(result.begin(), result.end());
      return result;
    }()};

  // population stats function
  using SamplePopSegments = pair<Sample, PopSegments>;
  auto pop_fun = [&all_sample_info](const Sample sample) {
    const SampleInfo & info{all_sample_info[sample]};
    // const CN_Bins & profile{info.profile};
    const Segments & segments{info.segments};

    PopSegments displayed;
    for (const Segment & segment : segments)
      if (segment.is_event())
        displayed.emplace_back(sample, all_sample_info, segment);
    sort(displayed.begin(), displayed.end());
    return SamplePopSegments{sample, displayed};
  };

  // Get population stats in parallel
  using AllPopSegments = vector<SamplePopSegments>;
  const AllPopSegments all_pop_segments{
    [&pool, &pop, pop_fun]() {
      ThreadPool::Results<SamplePopSegments> results;
      for (unsigned int s{0}; s != pop.n_samples(); ++s) {
        const Sample sample{s};
        if (pop.is_parent(sample)) continue;
        pool.run(results, pop_fun, sample);
      }
      AllPopSegments result;
      result.reserve(results.size());
      Progress progress{results.size(), 0.001, "get population info"};
      while (results.size()) {
        result.push_back(results.get());
        progress();
      }
      return result;
    }()};

  // Output
  const string chd_dir{out_dir + "/chd"};
  mkdir(chd_dir);
  const string bins_dir{chd_dir + "/" + to_string(bins.size())};
  mkdir(bins_dir);

  ofstream samples_out_file{bins_dir + "/samples.txt"};
  if (!samples_out_file) throw Error("Problem opening samples output file");
  doc_defaults.png(true);
  PSPage cand_pos_graph{bins_dir + "/cand_pos", "", "1 1"};
  PSHSeries<uint64_t, uint64_t> cand_pos_hist{cand_pos_graph,
        "Candidate genome distribution;Genome Position;N",
        Bounds{0, cn_abspos.n_positions() + 0.0}, 200};
  PSPage cand_cn_graph{bins_dir + "/cand_cn", "", "1 1"};
  PSHSeries<double, uint64_t> cand_cn_hist{cand_cn_graph,
        "Candidate copy number distribution;CN;N", Bounds{0, 10.0}, 200};
  PSPage cand_size_graph{bins_dir + "/cand_size", "", "1 1"};
  PSHSeries<double, uint64_t> cand_size_hist{cand_size_graph,
        "Candidate size distribution;log10(size);N", Bounds{0, 10.0}, 200};
  PSPage cand_mad_graph{bins_dir + "/cand_mad", "", "1 1"};
  PSHSeries<double, uint64_t> cand_mad_hist{cand_mad_graph,
        "Candidate mad distribution;MAD;N", Bounds{0, 1}, 200};
  PSPage cand_score_graph{bins_dir + "/cand_score", "", "1 1"};
  PSHSeries<double, uint64_t> cand_score_hist{cand_score_graph,
        "Candidate CN score distribution;CN score;N", Bounds{0, 310}, 310};
  PSPage cand_dscore_graph{bins_dir + "/cand_dscore", "", "1 1"};
  PSHSeries<double, uint64_t> cand_dscore_hist{cand_dscore_graph,
        "Candidate denovo score distribution;denovo score;N",
        Bounds{0, 310}, 310};
  PSPage cand_nscore_graph{bins_dir + "/cand_nscore", "", "1 1"};
  PSHSeries<double, uint64_t> cand_nscore_hist{cand_nscore_graph,
        "Candidate non-trans score distribution;non-trans score;N",
        Bounds{0, 310}, 310};
  PSPage cand_pscore_graph{bins_dir + "/cand_pscore", "", "1 1"};
  PSHSeries<double, uint64_t> cand_pscore_hist{cand_pscore_graph,
        "Candidate population score distribution;population score;N",
        Bounds{0, 310}, 310};
  doc_defaults.png(false);

  auto html = [&bins]
      (const string & dir, const string & html_name,
       const string & title, const string html_text,
       const string sample) {
    ofstream html_file{(dir + "/" + html_name).c_str()};
    if (!html_file) throw Error("Problem opening html file") << html_name;

    html_file << R"xxx(
<html>
<head>
<style>
body { padding: 10px; margin:10px; }
div.header { position:fixed; top:0; left:0; min-height:2em; width:100%;
             border-bottom:3px solid #a00; background-color:#ccc;
             text-align:center; }
div.header p { margin-top:0px; margin-bottom:0px; padding:0.3em; }
div.rthumb { float:right; margin-bottom:0.5em;}
div.thumb { float:left; margin-bottom:0.5em; min-width:300px; }
div.thumb p { float:left; margin-bottom:0.5em;}
div.thumb img { display:block; width:100%; }
b.chosen { color:#BB0000; }
dl { float:left; }
dt { font-weight: bold;}
tr:nth-child(even) { background-color:#f2f2f2; }
tr:hover { background-color:#ddd; }
td, th { font-size:75%; padding:3px; }
th { text-align:left; background-color:#4c50af; color:white; }
h1 { margin-top:1.5em; text-align:center; }
h1, h2, h3, p { clear:left; }
</style>
)xxx";
    html_file << "<title>" << title << "</title>\n";
    html_file << "</head>\n</body>\n";
    vector<unsigned int> all_n_bins{
      20000, 50000, 100000, 200000, 300000, 400000, 500000, 1000000};
    html_file << "<div class=\"header\"><p>";
    if (false) html_file << "<a href=\"/chd/" << bins.size()
                         << "/\"><b>defaults</b></a>&nbsp- ";
    html_file << "<b>n_bins</b>:&nbsp;";
    for (const unsigned int n_bins : all_n_bins) {
      if (n_bins != all_n_bins.front()) html_file << "&nbsp;|&nbsp;";
      if (n_bins == bins.size()) {
        html_file << "<b class=\"chosen\">" <<  n_bins << "</b>";
      } else {
        html_file << "<a href=\"/chd/" << n_bins << "/\">"
                  << n_bins << "</a>";
      }
    }
    vector<string> choices{
      "stats", "samples", "denovo", "transmitted", "unsure"};
    if (sample.size()) choices.push_back(sample);
    html_file << "&nbsp;- <b>views</b>:&nbsp;";
    for (const string & choice : choices) {
      const string file_name{
        choice == sample ? sample + "/" : choice + ".html"};
      if (choice != choices.front()) html_file << "&nbsp;|&nbsp;";
      if (file_name == html_name) {
        html_file << "<b class=\"chosen\">" << choice << "</b>";
      } else {
        html_file << "<a href=\"/chd/" << bins.size() << "/"
                  << file_name << "\">" << choice << "</a>";
      }
    }
    html_file << " - <b>by</b>:&nbsp;"
    << "<a href=\"http://drpa.us\">Peter&nbsp;Andrews</a>&nbsp;- ";

    html_file << "<b>" << title << "</b></p></div>\n";
    html_file << "<br /><h1>" << title << "</h1>\n";
    html_file << html_text;
    html_file << "<p><br /><br /><br /><br /><br /></p></body></html>\n";
  };

  // redirect html
  ofstream redirect{chd_dir + "/index.html"};
  if (!redirect) throw Error("Problem opening redirect html file");
  redirect << "<html>\n<head>\n"
           << "<meta http-equiv=\"Refresh\" "
           << "content=\"0; url=http://mumdex.com/chd/500000/\" />\n"
           << "</head>\n<body>\n"
           << "<h1><a href=\"http://mumdex.com/chd/500000/\">"
           << "Redirect to 500,000 bins page</a></h1>\n</body>\n</html>\n";

  // main html file
  ostringstream main_html;
  vector<string> hist_names{
    "pos", "cn", "size", "mad", "score", "dscore", "nscore", "pscore"};
  for (const string & hist_name : hist_names)
    main_html << "<div class=\"thumb\" style=\"width:33.333%;\">"
              << "<a href=\"cand_" << hist_name << ".png\">"
              << "<img src=\"cand_" << hist_name << ".png\" "
              << "title=\"" << hist_name << "\" "
              << "alt=\"" << hist_name << "\" />"
              << "</a></div>\n";
  html(bins_dir, "stats.html", "CHD&nbsp;statistics", main_html.str(),
       "");
  if (system(("ln -sf ./stats.html " + bins_dir + "/index.html").c_str()) != 0)
    throw Error("Could not link stats.html");

  using EventInfo = pair<double, string>;
  vector<EventInfo> denovo_events;
  vector<EventInfo> transmitted_events;
  vector<EventInfo> unsure_events;

  // samples html
  ostringstream samples_html;
  vector<EventInfo> samples_samples;

  mutex list_mutex;
  // mutex html_mutex;
  mutex hist_mutex;
  mutex file_mutex;
  bool first_out{true};
  auto output_fun =
      [&bins, &pop, &ref, &cn_abspos,
       &genes, &xref, &all_sample_info, &bins_dir, &out_dir,
       &denovo_events, &transmitted_events, &unsure_events,
       &html, &samples_html, &samples_samples, &samples_out_file,
       &list_mutex, &hist_mutex, &file_mutex, &first_out,
       &cand_pos_hist, &cand_cn_hist, &cand_size_hist, &cand_mad_hist,
       &cand_score_hist, &cand_dscore_hist,
       &cand_nscore_hist, &cand_pscore_hist]
      (const SamplePopSegments & sample_segments) {
    const Sample sample{sample_segments.first};
    const PopSegments segments{sample_segments.second};
    uint64_t n_denovo_gain{0};
    uint64_t n_denovo_loss{0};
    uint64_t n_trans_gain{0};
    uint64_t n_trans_loss{0};
    uint64_t n_unsure_gain{0};
    uint64_t n_unsure_loss{0};
    uint64_t n_strong_gain{0};
    uint64_t n_strong_loss{0};
    uint64_t n_weak_gain{0};
    uint64_t n_weak_loss{0};
    for (const PopSegment & pop_segment : segments) {
      const Segment & segment{pop_segment.segment};
      if (segment.call() == Segment::Gain) {
        if (segment.is_good()) {
          ++n_strong_gain;
          if (pop_segment.is_trans()) {
            ++n_trans_gain;
          } else if (pop_segment.is_denovo()) {
            ++n_denovo_gain;
          } else {
            ++n_unsure_gain;
          }
        } else {
          ++n_weak_gain;
        }
      } else if (segment.call() == Segment::Loss) {
        if (segment.is_good()) {
          ++n_strong_loss;
          if (pop_segment.is_trans()) {
            ++n_trans_loss;
          } else if (pop_segment.is_denovo()) {
            ++n_denovo_loss;
          } else {
              ++n_unsure_loss;
          }
        } else {
          ++n_weak_loss;
        }
      }
    }
    const SampleInfo & info{all_sample_info[sample]};
    vector<string> values{
      "family", pop.family(pop.family(sample)),
          "sample", pop.sample(sample),
          "member", pop.member(sample),
          "sex", pop.sex(sample),
          "devent", to_string(n_denovo_gain + n_denovo_loss),
          "dgain", to_string(n_denovo_gain),
          "dloss", to_string(n_denovo_loss),
          "tevent", to_string(n_trans_gain + n_trans_loss),
          "tgain", to_string(n_trans_gain),
          "tloss", to_string(n_trans_loss),
          "uevent", to_string(n_unsure_gain + n_unsure_loss),
          "ugain", to_string(n_unsure_gain),
          "uloss", to_string(n_unsure_loss),
          "wevent", to_string(n_weak_gain + n_weak_loss),
          "wgain", to_string(n_weak_gain),
          "wloss", to_string(n_weak_loss)};

    ostringstream out;

    // sample info
    ostringstream sample_base;
    sample_base << bins_dir << "/" << pop.sample(sample);
    mkdir(sample_base.str());
    ofstream sample_out_file{(sample_base.str() + "/stats.txt").c_str()};
    if (!sample_out_file)
      throw Error("Problem opening sample file for") << pop.sample(sample);

    ostringstream sample_html;
    ostringstream sample_images;
    ostringstream header_out;
    for (unsigned int n{0}; n != values.size(); n += 2)
      header_out << (n ? "\t" : "") << values[n];
    header_out << "\n";
    ostringstream sample_out;
    for (unsigned int n{0}; n != values.size(); n += 2)
      sample_out << (n ? "\t" : "") << values[n + 1];
    sample_out << "\n";
    sample_out_file << header_out.str() << sample_out.str();
    out << header_out.str() << sample_out.str() << "\n";

    ostringstream samples_samples_html;
    samples_samples_html << "<tr>";
    for (unsigned int n{0}; n != values.size(); n += 2) {
      samples_samples_html << "<td>";
      if (values[n] == "sample")
        samples_samples_html << "<a href=\"/chd/" << bins.size()
                             << "/" << pop.sample(sample) << "/\">";
      samples_samples_html << values[n + 1];
      if (values[n] == "sample") samples_samples_html << "</a>";
      samples_samples_html << "</td>";
    }
    samples_samples_html << "</tr>\n";

    {
      lock_guard<mutex> file_lock{file_mutex};
      if (first_out) {
        samples_html << "<table><tr>";
        // samples_html << "<th>view</th>";
        for (unsigned int n{0}; n != values.size(); n += 2)
          samples_html << "<th>" << values[n] << "</th>";
        samples_html << "</tr>\n";
        samples_out_file << header_out.str();
        first_out = false;
      }
      // -1.0 * (n_denovo_gain + n_denovo_loss), when ordered by double
      samples_samples.emplace_back(1.0 * sample,
                                   samples_samples_html.str());
      samples_out_file << sample_out.str();
    }

    ofstream cand_out_file{(sample_base.str() + "/cand.txt").c_str()};
    if (!cand_out_file)
      throw Error("Problem opening candidate file for") << pop.sample(sample);

    ofstream cand_ggraph_file{(sample_base.str() + "/ggraph.txt").c_str()};
    if (!cand_ggraph_file)
      throw Error("Problem opening ggraph file for") << pop.sample(sample);

    ostringstream cand_out;
    string last_type{""};
    for (unsigned int s{0}; s != segments.size(); ++s) {
      const PopSegment & psegment{segments[s]};
      const Segment & segment{psegment.segment};
      const unsigned int chr{bins[segment.start()].chromosome()};
      const unsigned int start{bins[segment.start()].start_position()};
      const unsigned int stop{bins[segment.stop() - 1].stop_position()};
      ostringstream cand_id;
      cand_id << psegment.call_string()
              << "." << ref.name(chr) << "." << start << "-" << stop;
      ostringstream cand_base;
      cand_base << sample_base.str() << "/" << cand_id.str();
      mkdir(cand_base.str());

      string cand_type{""};
      {
        lock_guard<mutex> list_lock{list_mutex};
        if (psegment.call() >= PopSegment::DenovoLoss) {
          cand_type = "denovo";
          denovo_events.emplace_back(
              segment.p_null(), pop.sample(sample) + "/" + cand_id.str());
        } else if (psegment.call() >= PopSegment::TransLoss) {
          cand_type = "transmitted";
          transmitted_events.emplace_back(
              segment.p_null(), pop.sample(sample) + "/" + cand_id.str());
        } else {
          cand_type = "unsure";
          unsure_events.emplace_back(
              segment.p_null(), pop.sample(sample) + "/" + cand_id.str());
        }
      }
      const vector<double> fcopies{psegment.fcopies};
      PSGraph cand_pop_graph{cand_base.str() + "/pop", ";Copy Number;N",
            Bounds{fcopies.front() - 0.5, fcopies.back() + 0.5}};
      PSHSeries<double, uint64_t> cand_pop_hist{cand_pop_graph, 200,
            "0 0 0", false};
      for (const double copy : fcopies) cand_pop_hist.add_point(copy);
      const vector<double> fcvalues{
        psegment.mother_copy, psegment.father_copy, segment.fcopy()};
      ostringstream cand_ps;
      cand_ps << "6 lw\n";
      for (unsigned int v{0}; v != fcvalues.size(); ++v)
        cand_ps << "np "
                << colors[v] << " c "
                << fcvalues[v] << " xc " << 1 << " yfc m "
                << fcvalues[v] << " xc " << 0.8 << " yfc l sp\n";
      cand_pop_graph.ps(cand_ps.str());

      if (psegment.call() >= PopSegment::DenovoLoss) {
        lock_guard<mutex> hist_lock{hist_mutex};
        cand_pos_hist.add_point(cn_abspos(chr, (start + stop) / 2));
        cand_cn_hist.add_point(segment.fcopy());
        cand_size_hist.add_point(log10(stop - start));
        cand_mad_hist.add_point(psegment.pop_mad);
        cand_score_hist.add_point(segment.score());
        cand_dscore_hist.add_point(psegment.denovo_score());
        cand_nscore_hist.add_point(psegment.not_trans_score());
        cand_pscore_hist.add_point(psegment.pop_score());
      }
      const OutResult out_result{psegment.out(info, genes, xref)};
      const Strings & seg_info{out_result.first};
      // const GeneInfos & gene_infos{out_result.second};
      if (!s) {
        for (unsigned int n{0}; n != seg_info.size(); n += 2)
          cand_out << (n ? "\t" : "") << seg_info[n];
        cand_out << "\n";

        sample_html << "<table><tr><th>view</th>";
        for (unsigned int n{0}; n + 2 != seg_info.size(); n += 2)
          sample_html << "<th>" << seg_info[n] << "</th>";
        sample_html << "</tr>\n";
      }
      for (unsigned int n{0}; n != seg_info.size(); n += 2)
        cand_out << (n ? "\t" : "") << seg_info[n + 1];
      cand_out << "\n";

      if (last_type != cand_type) {
        sample_images << "<h3>" << cand_type << " events</h3>";
        sample_html << "<tr><th style=\"background-color:#aaaaef;\">"
                    << cand_type << "</th></tr>\n";
        last_type = cand_type;
      }
      sample_images << "<div class=\"thumb\" style=\"width:33.3%;\">"
                    << "<a href=\"./" << cand_id.str() << "/\">"
                    << "<img src=\"./" << cand_id.str() << "/ggraph.png\" "
                    << "title=\"" << cand_id.str() << "\" "
                    << "alt=\"" << cand_id.str() << "\" /></a>"
                    << "</div>\n";
      sample_html << "<td><a href=\"./"
                  << cand_id.str() << "/\">view</a></td>";
      for (unsigned int n{0}; n + 2 != seg_info.size(); n += 2)
        sample_html << "<td>" << seg_info[n + 1] << "</td>";
      sample_html << "</tr>\n";

      ostringstream ggraph;
      const unsigned int eastart{cn_abspos(chr, start)};
      const unsigned int eastop{cn_abspos(chr, stop)};
      const unsigned int edge{(stop - start) / 1};
      const unsigned int astart{eastart > edge ? eastart - edge : 0};
      const unsigned int astop{eastop + edge < cn_abspos.n_positions() ?
            eastop + edge : cn_abspos.n_positions()};
      const double high_cn{std::max(2.0, segment.fcopy() + 1)};

      ggraph << " --initial "
             << astart << " " << astop << " 0 " << high_cn
             << " cn /data/safe/paa/analysis/mums/hg19/chrAll.fa "
             << "chr,start,ratio,seg_ratio "
             << out_dir << "/" << pop.family(pop.family(sample)) << "{P,M,F}/*"
             << bins.size() << "_bins_results.txt";
      cand_ggraph_file << "ggraph " << ggraph.str() << "\n";
      ofstream command{(cand_base.str() + "/ggraph.sh").c_str()};
      if (!command) throw Error("Problem opening ggraph command file");
      command << "cd " << cand_base.str() << "\n"
              << "rm -f *.{xpm,png,pdf}\n"
              << "(ggraph --output " << ggraph.str() << " 2>&1) |\n"
              << "grep -v -e ^Saved -e ^Converted\n"
              << "mv cn.1.png ggraph.png\n"
              << "[ -e pop.ps ] && convert pop.ps pop.png && rm pop.ps\n"
              << "rm -f *.{pdf,xpm,new}\n";

      ostringstream cand_html;
      cand_html << "<div class=\"thumb\" style=\"width:57%;\">"
                << "<img src=\"./ggraph.png\" "
                << "title=\"candidate profile\" "
                << "alt=\"candidate profile\" />"
                << "</div>\n";
      cand_html << "<div class=\"thumb\" style=\"width:43%;\">"
                << "<img src=\"./pop.png\" "
                << "title=\"candidate population copy\" "
                << "alt=\"candidate population copy\" />"
                << "</div>\n";
      cand_html << "<h2>candidate properties</h2>\n";
      const vector<string> break_points{"n_exons", "pop_p", "dummy"};
      uint64_t d{0};
      uint64_t b{0};
      while (d + 2 != seg_info.size()) {
        cand_html << "<dl style=\"width:33%\">";
        for (; d + 2 != seg_info.size(); d += 2) {
          cand_html << "<dt>" << seg_info[d] << "</dt>"
                    << "<dd>" << seg_info[d + 1] << "</dd>";
          if (seg_info[d] == break_points[b]) {
            ++b;
            d += 2;
            break;
          }
        }
        cand_html << "</dl>\n";
      }

      cand_html << "<dl style=\"clear:left;\"><dt>genes</dt><dd>"
                << seg_info.back() << "</dd></dl>\n";
      cand_html << "<h2>view command</h2><p>ggraph " << ggraph.str()
                << "</p>\n";
      html(cand_base.str(), "/index.html",
           "CHD sample " + pop.sample(sample) + " event " +
           cand_id.str(), cand_html.str(), pop.sample(sample));
    }
    cand_out_file << cand_out.str();
    out << cand_out.str();
    separator(out);
    sample_html << "</table>\n" << sample_images.str();
    sample_html << "<h2>text output files</h2>";
    sample_html << "<p><a href=\"./cand.txt\">candidate data</a></p>\n";
    sample_html << "<p><a href=\"./ggraph.txt\">ggraph view commands</a></p>\n";
    html(sample_base.str(), "/index.html",
         "CHD sample " + pop.sample(sample), sample_html.str(), "");

    return out.str();
  };

  // Output for all probands
  ThreadPool::Results<string> results;
  for (unsigned int p{0}; p != all_pop_segments.size(); ++p) {
    pool.run(results, output_fun, cref(all_pop_segments[p]));
  }
  Progress progress{results.size(), 0.001, "output results"};
  while (results.size()) {
    cout << results.get();
    progress();
  }

  cerr << "finishing up" << endl;

  sort(samples_samples.begin(), samples_samples.end());
  for (const EventInfo & info : samples_samples) samples_html << info.second;

  samples_html << "</table>\n";
  samples_html << "<h2>text output files</h2>"
               << "<p><a href=\"./samples.txt\">sample data text</a></p>\n";

  html(bins_dir, "samples.html", "CHD samples", samples_html.str(), "");

  const vector<std::string> cand_list_names{"denovo", "transmitted", "unsure"};
  const vector<vector<EventInfo> *> event_lists{
    &denovo_events, &transmitted_events, &unsure_events};
  for (unsigned int l{0}; l != event_lists.size(); ++l) {
    const string & name{cand_list_names[l]};
    vector<EventInfo> & events{*event_lists[l]};
    sort(events.begin(), events.end());
    ostringstream event_html;
    for (const EventInfo & info : events) {
      const string dir{info.second};
      event_html << "<div class=\"thumb\" style=\"width:25%;\">"
                 << "<a href=\"./" << dir << "\">"
                 << "<img src=\"./" << dir << "/ggraph.png\" "
                 << "title=\"" << dir << " profile\" "
                 << "alt=\"" << dir << " profile\" />"
                 << "</a></div>\n";
    }
    html(bins_dir, name + ".html", "CHD " + name + " events",
         event_html.str(), "");
  }

  return 0;
} catch (Error & e) {
  cerr << "paa::Error:" << endl;
  cerr << e.what() << endl;
  return 1;
} catch (exception & e) {
  cerr << "std::exception" << endl;
  cerr << e.what() << endl;
  return 1;
} catch (...) {
  cerr << "unknown exception was caught" << endl;
  return 1;
}
```
