## Supplementary material for "G-Graph: An interactive genomic graph viewer": MUMdex Source Code for G-Graph: recurrent_cn.cpp

```
//
// recurrent_cn.cpp
//
// Find repeated events and add gene annotations
//
// Copyright 2019 Peter Andrews @ CSHL
//

#include <algorithm>
#include <exception>
#include <fstream>
#include <future>
#include <iostream>
#include <map>
#include <sstream>
#include <string>
#include <utility>
#include <vector>

#include "cn.h"
#include "error.h"
#include "mumdex.h"
#include "psplot.h"
#include "stats.h"
#include "strings.h"
#include "threads.h"
#include "utility.h"

using std::cin;
using std::cout;
using std::cerr;
using std::endl;
using std::exception;
using std::future;
using std::ifstream;
using std::map;
using std::ofstream;
using std::ostringstream;
using std::pair;
using std::string;
using std::vector;

using paa::remove_substring;
using paa::Bin;
using paa::Bounds;
using paa::ChromosomeIndexLookup;
using paa::CN_abspos;
using paa::CN_Bin;
using paa::CN_Bins;
using paa::Error;
using paa::GeneHitFinder;
using paa::GeneInfoInterval;
using paa::GeneXrefs;
using paa::HitType;
using paa::KnownGene;
using paa::KnownGenes;
using paa::MAD;
using paa::Marker;
using paa::Progress;
using paa::PSDoc;
using paa::PSGraph;
using paa::PSHSeries;
using paa::PSXYSeries;
using paa::Reference;
using paa::ThreadPool;

using GeneInfo = pair<string, bool>;
using GeneInfos = vector<GeneInfo>;
using Strings = vector<string>;
using OutResult = pair<Strings, GeneInfos>;

int main(int argc, char* argv[]) try {
  paa::exit_on_pipe_close();
  std::ios_base::sync_with_stdio(false);
  cin.tie(nullptr);

  if (--argc != 2)
    throw Error("usage: cat events.txt | recurrent_events ref n_binnings");

  // Settings
  const uint64_t bin_size{1000};
  const unsigned int max_event_size{1000000};
  const double recurrence_cutoff{5};

  // Process arguments
  const string reference_file{argv[1]};
  const Reference ref{reference_file};
  const ChromosomeIndexLookup lookup{ref};
  const CN_abspos cn_abspos{ref};
  const unsigned int n_binnings{static_cast<unsigned int>(atoi(argv[2]))};

  // Graphs
  const string out_base{"recurrent"};
  PSDoc ps{out_base};
  ps.pdf(true);
  const Marker marker{paa::circle(), 0.1, "1 0 0", 1, true};
  PSXYSeries mountain{ps, "Recurrent CN Events;Abspos;Recurrence", marker};

  cerr << "Read events" << endl;
  string sample;
  string chr_name;
  unsigned int start;
  unsigned int stop;
  string call;
  vector<unsigned int> hits(cn_abspos.n_positions());
  uint64_t n_lines{0};
  while (cin >> sample >> chr_name >> start >> stop >> call) {
    const unsigned int chr{lookup[chr_name]};
    const unsigned int abspos_start{cn_abspos(chr, start)};
    const unsigned int abspos_stop{abspos_start + stop - start};
    if (stop - start < max_event_size) {
      for (unsigned int abspos{abspos_start}; abspos != abspos_stop; ++abspos) {
        ++hits[abspos];
      }
    }
    ++n_lines;
  }
  cerr << "Read " << n_lines << " events" << endl;

  vector<double> hits_hist(hits.size() / bin_size + 1);
  uint64_t bin_total{0};
  double largest_value{0};
  ofstream recurrence_out{"recurrence.txt"};
  for (unsigned int abspos{0}; abspos != hits.size(); ++abspos) {
    if ((abspos && (abspos % bin_size == 0)) || abspos + 1 == hits.size()) {
      if (bin_total) {
        const uint64_t bin{abspos + 1 == hits.size() ?
              hits_hist.size() - 1 : abspos / bin_size};
        const double value{1.0 * bin_total / bin_size / n_binnings};
        largest_value = std::max(largest_value, value);
        hits_hist[bin] = value;
        // cout << abspos << " " << value << endl;
        recurrence_out << abspos << " " << value << endl;
        mountain.add_point(abspos - bin_size / 2, value);
        // mountain[chr].add_point(abspos - bin_size / 2, value);
        // hist.add_point(value);
      }
      bin_total = 0;
    }
    bin_total += hits[abspos];
  }
  cerr << "Made mountain.  Largest value is " << largest_value << endl;

  largest_value += 1;
  const unsigned int n_hist_bins{300};
  vector<unsigned int> recurrence_hist(n_hist_bins);
  for (const double value : hits_hist) {
    const unsigned int bin{static_cast<unsigned int>(
        n_hist_bins * value / largest_value)};
    ++recurrence_hist[bin];
  }
  PSGraph hist_graph{ps, "Recurrent CN histograms;Recurrence;N"};
  hist_graph.log_y(true);
  const Marker bigger_marker{paa::circle(), 0.5, "1 0 0", 1, true, "1 0 0"};
  PSXYSeries hist{hist_graph, bigger_marker};
  uint64_t total{0};
  PSGraph cutoff_graph{ps, "Recurrence Cutoff;Recurrence;% Genome Cut"};
  PSXYSeries cutoff_series{cutoff_graph, bigger_marker};
  for (unsigned int b{0}; b != recurrence_hist.size(); ++b) {
    const double value{(b+1) * largest_value / n_hist_bins};
    const unsigned int count{recurrence_hist[b]};
    total += count;
    cutoff_series.add_point(
        value, 100.0 * (hits_hist.size() - total) / hits_hist.size());
    hist.add_point(value, count);
  }

  // Look at individual regions to judge cuts
  bool above_cutoff{false};
  unsigned int peak_start{0};
  ofstream cut_file{"cuts.txt"};
  for (unsigned int b{0}; b != hits_hist.size(); ++b) {
    const double value{hits_hist[b]};
    if (above_cutoff) {
      if (value < recurrence_cutoff) {
        const unsigned int peak_stop{b};
        const uint64_t region_center{bin_size * (peak_start + peak_stop) / 2};
        const uint64_t region_edge{bin_size * (peak_stop - peak_start) + 2};
        const uint64_t region_start{
          bin_size * peak_start > region_edge ?
              bin_size * peak_start - region_edge : 0};
        const uint64_t region_stop{
            bin_size * peak_stop + region_edge > hits.size() ?
            hits.size() : bin_size * peak_stop + region_edge};
        ostringstream region_file_name;
        region_file_name << "regions/data." << region_start << ".txt";
        ofstream region_data{region_file_name.str().c_str()};
        ostringstream region_limits_name;
        region_limits_name << "regions/data." << region_start << ".limits.txt";
        ofstream limits_data{region_limits_name.str().c_str()};
        region_data << "abspos\trecurrence\n";
        limits_data << "abspos\trecurrence\n";
        const CN_abspos::ChrPos chrpos{
          cn_abspos.chrpos(static_cast<unsigned int>(region_center))};
        cut_file << region_file_name.str()
                 << " " << region_limits_name.str()
                 << " " << region_start
                 << " " << region_stop
                 << " " << largest_value
                 << " " << ref.name(chrpos.first)
                 << " " << chrpos.second
                 << " " << (peak_stop - peak_start) * bin_size / 1000.0 << "kb"
                 << endl;
        for (double l{0}; l < largest_value + 5; l += 0.1) {
          limits_data << peak_start * bin_size << "\t" << l << "\n";
          limits_data << peak_stop * bin_size << "\t" << l << "\n";
        }
        for (uint64_t r{region_start}; r != region_stop; ++r)
          region_data << r << "\t" << hits[r] / n_binnings << '\n';
        above_cutoff = false;
      }
    } else {
      if (value > recurrence_cutoff) {
        peak_start = b;
        above_cutoff = true;
      }
    }
  }

  return 0;
  // Load gene info
  const string genes_name{reference_file + ".bin/knownGene.txt"};
  const string isoforms_name{reference_file + ".bin/knownIsoforms.txt"};
  const string kgXrefs_name{reference_file + ".bin/kgXref.txt"};
  const KnownGenes genes{genes_name, isoforms_name, lookup, ref};
  const GeneXrefs xref{kgXrefs_name};
  const GeneHitFinder gene_finder{genes};

#if 0

  PSGraph hist_graph{ps, "Recurrent CN Events;Recurrence;N",
        Bounds{0.0, 100.0}};
  hist_graph.log_y(true);


  // Determine gene overlaps for each bin
  vector<string> bin_gene_annotations;
  vector<unsigned int> bin_gene_n_exons;
  vector<unsigned int> bin_gene_n_exon_genes;
  Progress gene_progress{bins.size(), 0.01, "get gene overlaps"};
  for (const Bin & bin : bins) {
    // get gene info for segment
    ostringstream gene_info;
    const KnownGenes::GeneOverlaps gene_overlaps{
      genes.find_genes(bin.chromosome(),
                       bin.start_position(), bin.stop_position())};
    map<string, unsigned int> named_overlaps;
    for (const KnownGenes::GeneOverlap & overlap : gene_overlaps) {
      const string symbol{xref[genes[overlap.first].name].geneSymbol};
      const unsigned int old_value{named_overlaps[symbol]};
      if (old_value < overlap.second) {
        named_overlaps[symbol] = overlap.second;
      }
    }
    using Overlap = pair<string, unsigned int>;
    vector<Overlap> overlaps(named_overlaps.begin(), named_overlaps.end());
    sort(overlaps.begin(), overlaps.end(),
         [](const Overlap & lhs, const Overlap & rhs) {
           if (lhs.second == rhs.second) {
             return lhs.first < rhs.first;
           } else {
             return lhs.second > rhs.second;
           }
         });
    unsigned int genes_with_exons{0};
    unsigned int total_exons{0};
    for (unsigned int o{0}; o != overlaps.size(); ++o) {
      genes_with_exons += static_cast<bool>(overlaps[o].second);
      total_exons += overlaps[o].second;
    }
    GeneInfos gene_infos;
    for (unsigned int o{0}; o != overlaps.size(); ++o) {
      const Overlap & overlap{overlaps[o]};
      if (o) gene_info << ",";
      if (overlap.second)
        gene_info << overlap.second << "-exon"
                  << (overlap.second > 1 ? "s" : "") << "-";
      gene_info << overlap.first;
      gene_infos.emplace_back(overlap.first, static_cast<bool>(overlap.second));
    }
    if (overlaps.empty()) {
      gene_info << "intergenic";
    }
    const string gene_string{gene_info.str()};
    if (false && gene_string != "intergenic")
      cout << ref.name(bin.chromosome())
           << '\t' << bin.start_position()
           << '\t' << bin.stop_position()
           << '\t' << bin.length()
           << '\t' << bin.map()
           << '\t' << total_exons
           << '\t' << genes_with_exons
           << '\t' << gene_string
           << endl;
    bin_gene_annotations.push_back(gene_string);
    bin_gene_n_exons.push_back(total_exons);
    bin_gene_n_exon_genes.push_back(genes_with_exons);
    gene_progress();
  }

  // pick best bins (lowest spread) to use by chromosome.
  map<unsigned int, vector<unsigned int>> best_bins;
  const uint64_t max_returned{5};
  for (unsigned int b{0}; b != bins.size(); ++b)
    if (!bad_bins[b] && bin_gene_n_exons[b])
      best_bins[bins[b].chromosome()].push_back(b);
  cout << "chr\tabspos\tstart\tstop\tlength\tmap_length\tgc\tratio_spread"
       << "\tn_exons\tn_exonic\tgene_description" << endl;
  for (const unsigned int chr : cn_abspos.chromosomes()) {
    try {
      auto & chosen = best_bins.at(chr);
      sort(chosen.begin(), chosen.end(),
           [&average_spread](const unsigned int b1, const unsigned int b2) {
             return average_spread[b1] < average_spread[b2];
           });
      if (max_returned > chosen.size()) {
        cerr << "Fewer good bins than desired found for "
             << ref.name(chr) << " (" << chosen.size() << ")" << endl;
      } else {
        chosen.resize(max_returned);
      }
      for (const unsigned int b : chosen) {
        const Bin & bin{bins[b]};
        cout << ref.name(bin.chromosome())
             << '\t' << bin.abspos()
             << '\t' << bin.start_position()
             << '\t' << bin.stop_position()
             << '\t' << bin.length()
             << '\t' << bin.map()
             << '\t' << bin.gc()
             << '\t' << average_spread[b]
             << '\t' << bin_gene_n_exons[b]
             << '\t' << bin_gene_n_exon_genes[b]
             << '\t' << bin_gene_annotations[b]
             << endl;
      }
    } catch (...) {
      cerr << "No good bins found for " << ref.name(chr) << endl;
      continue;
    }
  }
#endif

  return 0;
} catch (Error & e) {
  cerr << "paa::Error:" << endl;
  cerr << e.what() << endl;
  return 1;
} catch (exception & e) {
  cerr << "std::exception" << endl;
  cerr << e.what() << endl;
  return 1;
} catch (...) {
  cerr << "unknown exception was caught" << endl;
  return 1;
}
```
