## Supplementary material for "G-Graph: An interactive genomic graph viewer": MUMdex Source Code for G-Graph: smash.cpp

```
//
// smash.cpp
//
// Process smash data
//
// Not for clinical use
//
// Copyright 2016 Peter Andrews @ CSHL
//

#include <climits>

#include <algorithm>
#include <array>
#include <exception>
#include <fstream>
#include <future>
#include <iostream>
#include <memory>
#include <mutex>
#include <random>
#include <set>
#include <sstream>
#include <string>
#include <utility>
#include <vector>

#include "./cn.h"
#include "./error.h"
#include "./files.h"
#include "./longSA.h"
#include "./lowess.h"
#include "./mumdex.h"
#include "./psplot.h"
#include "./utility.h"

using std::array;
using std::async;
using std::binomial_distribution;
using std::cout;
using std::cerr;
using std::endl;
using std::exception;
using std::future;
using std::ifstream;
using std::istream;
using std::istringstream;
using std::lock_guard;
using std::make_unique;
using std::max;
using std::min;
using std::mt19937_64;
using std::mutex;
using std::ofstream;
using std::ostringstream;
using std::random_device;
using std::set;
using std::swap;
using std::string;
using std::to_string;
using std::unique_ptr;
using std::vector;

using paa::Bin;
using paa::Bounds;
using paa::ChromosomeIndexLookup;
using paa::CN_abspos;
using paa::Error;
using paa::FinestBins;
using paa::PosInfo;
using paa::PSCNGraph;
using paa::PSDoc;
using paa::PSGraph;
using paa::PSHSeries;
using paa::PSXYSeries;
using paa::Reference;
using paa::Sequence;
using paa::match_t;
using paa::nunset;
using paa::reverse_complement;
using paa::serr;
using paa::sout;
using paa::tout;
using paa::unset;

#define SHORT 1
#if SHORT
const bool use_reverse_complement{false};  // false for lower memory usage
using SA = paa::shortSA;
#else
const bool use_reverse_complement{true};  // true for high memory usage
using SA = paa::longSA;
#endif

using Mappability = paa::PreMappedMappability;
using paa::PreMappedVector;

const unsigned int joined_offset{10000};
const unsigned int discarded_offset{20000};

class MappingInfo {
 public:
  MappingInfo(const Sequence & ref, match_t match, const bool read_2__,
              const unsigned int read_length__) :
      position_{0}, chromosome_{0}, offset_{match.offset}, length_{0},
    read_2_{read_2__}, flipped_{false},
    // filtered_length_{false},
    filtered_excess_{false},
    // joined_{false},
    discarded_{false}, seen_before_{false}, pair_end_{false},
    read_length_{read_length__} {
    if (match.len >= discarded_offset) {
      match.len -= discarded_offset;
      discarded_ = true;
    } else {
      discarded_ = false;
    }
    if (match.len >= joined_offset && match.len < discarded_offset) {
      match.len -= joined_offset;
      // joined_ = true;
    } else {
      // joined_ = false;
    }
    length_ = match.len;

    flipped_ = false;
    uint64_t rpos{match.ref};
    if (rpos >= ref.N) {  // for flipped non rcref
      rpos -= ref.N;
      flipped_ = true;
    }
    std::vector<uint64_t>::const_iterator it =
        upper_bound(ref.startpos.begin(), ref.startpos.end(), rpos);

    int64_t chr{it - ref.startpos.begin() - 1};
    position_ = static_cast<unsigned int>(rpos - *--it);
    if (ref.rcref) {
      if ((chr % 2) == 1) {
        position_ = ref.sizes[chr] - position_ - length_;
        flipped_ = true;
      }
      chr /= 2;
    }
    chromosome_ = chr;
  }

  unsigned int chromosome() const { return chromosome_; }
  unsigned int position() const { return position_; }
  unsigned int offset() const { return offset_; }
  unsigned int length() const { return length_; }
  unsigned int read_length() const { return read_length_; }

  bool read_2() const { return read_2_; }
  bool flipped() const { return flipped_; }
  /*
  bool filtered_length() const { return filtered_length_; }
  void filtered_length(const bool filtered_length__) {
    filtered_length_ = filtered_length__;
  }
  */
  bool filtered_excess() const { return filtered_excess_; }
  void filtered_excess(const bool filtered_excess__) {
    filtered_excess_ = filtered_excess__;
  }
  /*
  bool joined() const { return joined_; }
  */
  bool discarded() const { return discarded_; }
  bool seen_before() const { return seen_before_; }
  void seen_before(const bool seen_before__) {
    seen_before_ = seen_before__;
  }
  bool pair_end() const { return pair_end_; }
  void pair_end(const bool pair_end__) {
    pair_end_ = pair_end__;
  }

 private:
  uint64_t position_ : 28;          // 28 x
  uint64_t chromosome_ : 7;         // 35 x
  uint64_t offset_ : 8;             // 43 x
  uint64_t length_ : 8;             // 51 x
  uint64_t read_2_ : 1;             // 52 x
  uint64_t flipped_ : 1;            // 53 x

  // uint64_t filtered_length_ : 1;    // 54 x
  uint64_t filtered_excess_ : 1;    // 54 x
  // uint64_t joined_ : 1;             // 55 x
  uint64_t discarded_ : 1;          // 56 x
  uint64_t seen_before_ : 1;        // 57 x
  uint64_t pair_end_ : 1;           // 58 x
  uint64_t read_length_ : 8;
};

uint64_t rotr(const uint64_t value, unsigned int count) {
  constexpr unsigned int mask = CHAR_BIT * sizeof(value) - 1;
  count &= mask;
  return (value >> count) | (value << ((-count) & mask));
}

class PairHash {
 public:
  explicit PairHash(const vector<MappingInfo> & maps) {
    for (const MappingInfo map : maps) {
      const uint64_t value{map.position() - map.offset() +
            (static_cast<uint64_t>(map.chromosome()) << 32)};
      hash_ = hash_ ^ rotr(value, static_cast<unsigned int>(value));
    }
  }
  uint64_t hash() const {
    return hash_;
  }
  operator uint64_t() const {
    return hash_;
  }

 private:
  uint64_t hash_{0};
};

mutex cout_mutex;

vector<uint64_t> read_length_hist;
uint64_t n_pairs{0};
vector<MappingInfo> smash_map(
    const SA & sa, istream & fq1, istream & fq2, mutex & fq_mutex,
    const unsigned int min_length) {
  array<string, 2> sequences;

  vector<MappingInfo> mappings;
  vector<match_t> MUMs;
  while (true) {
    {
      lock_guard<mutex> fq_lock(fq_mutex);
      fq1.ignore(10000, '\n');
      getline(fq1, sequences[0]);
      if (!fq1) return mappings;
      fq1.ignore(10000, '\n');
      fq1.ignore(10000, '\n');
      fq2.ignore(10000, '\n');
      getline(fq2, sequences[1]);
      fq2.ignore(10000, '\n');
      fq2.ignore(10000, '\n');
      // Update stats
      for (const bool read_2 : { false, true }) {
        const uint64_t read_length{sequences[read_2].size()};
        if (read_length_hist.size() <= read_length) {
          read_length_hist.resize(read_length + 1);
        }
        ++read_length_hist[read_length];
      }
      ++n_pairs;
    }

    for (const bool read_2 : {false, true}) {
      // Map sequence
      MUMs.clear();
      sa.find_mams(MUMs, sequences[read_2], min_length);

      // Merge zero-invariant MUMs, discard MUMs between mergings
      // PROBLEM with flipped ?  YES
      for (unsigned int m1{0}; m1 != MUMs.size(); ++m1) {
        match_t & mum1{MUMs[m1]};
        if (!mum1.len) throw Error("Zero length MUM");
        for (unsigned int m2{m1 + 1}; m2 != MUMs.size(); ++m2) {
          match_t & mum2{MUMs[m2]};
          if (mum2.len >= discarded_offset) continue;
          if (mum1.ref + mum2.offset == mum2.ref + mum1.offset) {
            mum1.len = mum2.offset - mum1.offset + mum2.len + joined_offset;
            while (m1 != m2) {
              ++m1;
              MUMs[m1].len += discarded_offset;
            }
          }
        }
      }

      // Return mappings, after making some cuts
      for (const match_t mum : MUMs) {
        // if (mum.len < min_length) continue;
        const uint64_t read_length{sequences[read_2].size()};
        mappings.emplace_back(sa.ref, mum, read_2, read_length);
        /*
        MappingInfo & map{mappings.back()};
        if (map.length() < min_length) {
          map.filtered_length(true);
        }
        */

#if 0
        // Testing crap
        string read_mum_seq{
          sequences[read_2].substr(map.offset(), map.length())};
        if (map.flipped()) {
          reverse_complement(&read_mum_seq);
        }
        string genome_mum_seq{ref.subseq(map.chromosome(),
                                         map.position(),
                                         map.position() + map.length())};
        if (read_mum_seq != genome_mum_seq && !map.joined()) {
          lock_guard<mutex> cout_lock(cout_mutex);
          cout << "mapping "
               << ref.name(map.chromosome()) << " " << map.position()
               << map.length() << " " << map.flipped() << " "
               << (read_mum_seq != genome_mum_seq) << "\n"
               << read_mum_seq << "\n"
               << genome_mum_seq << "\n";
        }
#endif
      }
    }
    if (mappings.size()) mappings.back().pair_end(true);
  }
}


int main(int argc, char * argv[]) try {
  paa::exit_on_pipe_close();
  const bool do_memory_map{true};
  paa::memory_mapped = do_memory_map;
  paa::read_ahead = true;

  const bool create_pdf{false};

  --argc;

  // process optional arguments first
  double alpha{0.05};
  double undo{1.0};
  unsigned int minw{3};
  while (argc) {
    bool acted{false};
    if (argc > 1 && argv[1] == string("-a")) {
      alpha = atof(argv[2]);
      argc -= 2;
      argv += 2;
      acted = true;
    }
    if (argc > 1 && argv[1] == string("-u")) {
      undo = atof(argv[2]);
      argc -= 2;
      argv += 2;
      acted = true;
    }
    if (argc > 1 && argv[1] == string("-m")) {
      minw = atoi(argv[2]);
      argc -= 2;
      argv += 2;
      acted = true;
    }
    if (!acted) break;
  }

  if (argc != 10 && argc != 11)
    throw Error("normal usage: smash ref fq1 fq2 minLength minExcess "
                "excludeWindow n_threads out_prefix bin_file,... bad_pos\n"
                "or\n"
                "finebin usage: smash ref fq1 fq2 minLength minExcess "
                "excludeWindow n_threads out_prefix n_samples n_x n_y\n");

  const string ref_name{argv[1]};

  const string fq1_name{argv[2]};
  const string fq2_name{argv[3]};
  ifstream fq1_file{fq1_name.c_str()};
  if (!fq1_file) throw Error("Could not open fastq file 1") << fq1_name;
  ifstream fq2_file{fq2_name.c_str()};
  if (!fq2_file) throw Error("Could not open fastq file 2") << fq2_name;

  const unsigned int min_length{static_cast<unsigned int>(atoi(argv[4]))};
  const unsigned int min_excess{static_cast<unsigned int>(atoi(argv[5]))};
  const unsigned int exclude_window{static_cast<unsigned int>(atoi(argv[6]))};
  const unsigned int n_threads{static_cast<unsigned int>(atoi(argv[7]))};
  const string output_prefix{argv[8]};
  const bool normal_output{argc == 10};

  // Map reads
  vector<vector<MappingInfo> > mappings(n_threads);
  uint64_t n_mappings{0};
  {
    cout << "Loading suffix array structures" << endl;
    const SA sa{ref_name, use_reverse_complement, do_memory_map, true};
    cout << "Mapping reads using " << n_threads << " threads" << endl;
    vector<future<vector<MappingInfo> > > futures(n_threads);
    mutex fq_mutex;
    for (unsigned int t{0}; t != n_threads; ++t) {
      futures[t] = async(std::launch::async,
                         smash_map, std::ref(sa),
                         std::ref(fq1_file), std::ref(fq2_file),
                         std::ref(fq_mutex), min_length);
    }
    for (unsigned int t{0}; t != n_threads; ++t) {
      future<vector<MappingInfo> > & result{futures[t]};
      mappings[t] = result.get();
      n_mappings += mappings[t].size();
    }
  }

  cout << "Loading reference and mappability" << endl;
  paa::memory_mapped = false;
  const Reference ref{ref_name};
  const Mappability mappability{ref_name, true};
  const CN_abspos cn_abspos{ref};

  cout << "Filtering mappings and assigning to bins" << endl;
  uint64_t n_used_pairs{0};
  uint64_t n_filtered{0};
  // uint64_t n_filtered_length{0};
  uint64_t n_filtered_excess{0};
  uint64_t n_discarded{0};
  // uint64_t n_joined{0};
  uint64_t n_seen_before{0};
  uint64_t n_dupe_pairs{0};
  uint64_t n_dupe_mappings{0};
  uint64_t n_good_mappings{0};
  uint64_t n_plotted_mappings{0};
  vector<MappingInfo> seen_before;
  vector<MappingInfo> good_mappings;
  vector<uint64_t> map_counts;
  vector<uint64_t> map_offsets;
  vector<uint64_t> map_lengths;
  vector<uint64_t> map_orientations;
  vector<uint64_t> map_mates;
  vector<uint64_t> map_coverage_1;
  vector<uint64_t> map_coverage_2;
  vector<uint64_t> map_unmapped_lengths;
  unsigned int max_template_length{1000};
  vector<uint64_t> template_lengths;
  set<uint64_t> dupes;

  // Bins, only loaded if used according to command line arguments
  using AllBins = vector<vector<Bin> >;
  const unique_ptr<AllBins> bins{[normal_output, argv, &ref]() {
      if (normal_output) {
        const string bins_string{argv[9]};
        vector<string> bins_names;
        istringstream bins_stream{bins_string.c_str()};
        string bins_name;
        while (getline(bins_stream, bins_name, ',')) {
          bins_names.push_back(bins_name);
        }
        unique_ptr<AllBins> result{make_unique<AllBins>(bins_names.size())};
        for (unsigned int n{0}; n != bins_names.size(); ++n) {
          // changed behavior - no masking!
          (*result)[n] = load_bins(bins_names[n], ref, false, false);
        }
        return result;
      } else {
        return unique_ptr<vector<vector<Bin> > >{nullptr};
      }
    }()};

  vector<vector<unsigned int>> counts{[normal_output, &bins]() {
      if (normal_output) {
        vector<vector<unsigned int>> result(bins->size());
        for (unsigned int b{0}; b != result.size(); ++b) {
          result[b].resize((*bins)[b].size());
        }
        return result;
      } else {
        return vector<vector<unsigned int>>();
      }
    }()};

  const unique_ptr<PreMappedVector<unsigned char>> bad{
    [normal_output, argv] () {
      if (normal_output) {
        return make_unique<PreMappedVector<unsigned char>>(argv[10]);
      } else {
        return unique_ptr<PreMappedVector<unsigned char>>{nullptr};
      }
    }()};

  // finebin, only loaded if used according to command line arguments
  unique_ptr<FinestBins> finebins{[normal_output, argv, &ref]() {
      if (normal_output) {
        return unique_ptr<FinestBins>{nullptr};
      } else {
        return make_unique<FinestBins>(
            ref,
            static_cast<unsigned int>(atoi(argv[9])),
            static_cast<unsigned int>(atoi(argv[10])),
            static_cast<unsigned int>(atoi(argv[11])));
      }
    }()};

  for (unsigned int t{0}; t != n_threads; ++t) {
    for (unsigned int m{0}; m != mappings[t].size(); ++m) {
      MappingInfo & map{mappings[t][m]};
      const unsigned int abspos{ref.abspos(map.chromosome(), map.position())};
#if 0
      const unsigned int min_map{mappability.min(abspos, map.length())};
      const unsigned int excess_map{map.length() - min_map};
#else
      const unsigned int max_map{
        max(mappability.low(abspos),
            mappability.high(abspos + map.length() - 1))};
      if (max_map > map.length()) {
        throw Error("Bad excess mappability")
            << ref.name(map.chromosome()) << map.position()
            << max_map << map.length()
            << mappability.low(abspos)
            << mappability.high(abspos + map.length() - 1);
      }
      const unsigned int excess_map{map.length() - max_map};
#endif
      bool filtered{false};
      if (excess_map < min_excess) {  // use < ?
        map.filtered_excess(true);
        ++n_filtered_excess;
        filtered = true;
      }
      /*
      if (map.filtered_length()) {
        ++n_filtered_length;
        filtered = true;
      }
      */
      if (map.discarded()) {
        ++n_discarded;
        filtered = true;
      }
      /*
      if (map.joined()) {
        ++n_joined;
      }
      */
      const bool pair_end{map.pair_end()};
      for (MappingInfo & seen : seen_before) {
        if (map.chromosome() == seen.chromosome() &&
            map.position() <= seen.position() + exclude_window &&
            map.position() + exclude_window >= seen.position()) {
          ++n_seen_before;
          filtered = true;
          if (map.read_2() != seen.read_2() &&
              map.flipped() != seen.flipped()) {
            const unsigned int read_pos_1{map.flipped() ?
                  map.position() + map.read_length()
                  - map.offset() - map.length() :
                  map.position() - map.offset()};
            const unsigned int read_pos_2{seen.flipped() ?
                  seen.position() + seen.read_length()
                  - seen.offset() - seen.length() :
                  seen.position() - seen.offset()};
            const unsigned int pos_diff{read_pos_1 > read_pos_2 ?
                  read_pos_1 - read_pos_2 : read_pos_2 - read_pos_1};
            const unsigned int template_length{pos_diff +
                  max(map.read_length(), seen.read_length())};
            if (template_length < max_template_length) {
              template_lengths.push_back(template_length);
            }
          }
          if (map.length() < seen.length()) {
            swap(seen, map);
          }
        }
      }
      if (filtered) {
        ++n_filtered;
      } else {
        seen_before.push_back(map);
      }
      if (pair_end) {
        // Look for dupes
        const PairHash hash{seen_before};
        if (dupes.count(hash)) {
          ++n_dupe_pairs;
          n_dupe_mappings += seen_before.size();
          n_filtered += seen_before.size();
        } else {
          dupes.insert(hash);
          ++n_used_pairs;
          if (normal_output) {
            // Output mappings
            if (map_counts.size() < seen_before.size() + 1)
              map_counts.resize(seen_before.size() + 1);
            ++map_counts[seen_before.size()];
            // cout << seen_before.size() << endl;
            for (const MappingInfo & seen : seen_before) {
              good_mappings.push_back(seen);
              const PosInfo pos{seen.chromosome(), seen.position()};
              ++n_good_mappings;
              const unsigned int cnabspos{cn_abspos(seen.chromosome(),
                                                    seen.position())};
              if (cnabspos < cn_abspos.bad_abspos() &&
                  !(*bad)[cnabspos]) {
                for (unsigned int b{0}; b != bins->size(); ++b) {
                  vector<Bin>::const_iterator found{
                    lower_bound((*bins)[b].begin(), (*bins)[b].end(), pos)};
                  if (found-- != (*bins)[b].begin()) {
                    if (seen.chromosome() == found->chromosome() &&
                        seen.position() >= found->start_position() &&
                        seen.position() < found->stop_position()) {
                      if (!b) ++n_plotted_mappings;
                      ++counts[b][found - (*bins)[b].begin()];
                    }
                  }
                }
              }
            }
          } else {
            for (const MappingInfo & seen : seen_before) {
              const unsigned int cnabspos{cn_abspos(seen.chromosome(),
                                                    seen.position())};
              if (cnabspos < cn_abspos.bad_abspos()) {
                ++(*finebins)[cnabspos];
              }
            }
          }
        }
        seen_before.clear();
      }
    }
    // mappings[t].swap(good_mappings);
    // good_mappings.clear();
  }

#if 0
  // Only set to 1 for testing !!!!!!!!!
  random_device rd;
  auto mersenne = mt19937_64(rd());
  const unsigned int desired_maps{12000000};
  if (n_plotted_mappings > desired_maps) {
    const double desired_downsample{1.0 * desired_maps / n_plotted_mappings};
    for (unsigned int B{0}; B != bins->size(); ++B) {
      for (unsigned int b{0}; b != (*bins)[B].size(); ++b) {
        unsigned int & count{counts[B][b]};
        count = binomial_distribution<unsigned int>{
          count, desired_downsample}(mersenne);
      }
    }
  }
#endif

  if (!normal_output) {
    cerr << "Saving finebins at " << output_prefix << endl;
    finebins->save(output_prefix + ".finebins.bin");
    return 0;
  }

  cout << "Determine copy number" << endl;
  for (unsigned int b{0}; b != bins->size(); ++b) {
    copy_number(ref, (*bins)[b], counts[b],
                output_prefix + " " + to_string((*bins)[b].size()) + " bins",
                create_pdf, true, alpha, undo, minw);
  }

  cout << "Make mapping statistics plots" << endl;
  // Randomization for some plots
  std::function<float()> gen{bind(
      std::uniform_real_distribution<float>{-0.5, 0.5},
      std::mt19937_64(std::random_device()()))};

  PSDoc ps{output_prefix + "_stats"};
  ps.pdf(create_pdf);

  // Map counts histogram
  PSGraph map_counts_graph{ps, ";Mappings Per Read Pair;N",
        Bounds(0, map_counts.size())};
  PSHSeries<int, uint64_t> map_counts_series{
    map_counts_graph, static_cast<unsigned int>(map_counts.size()),
        "1 0 0", false};
  map_counts[0] = n_pairs - n_used_pairs;
  for (unsigned int i{0}; i != map_counts.size(); ++i) {
    // sout << "map counts" << i << map_counts[i] << endl;
    map_counts_series.add_value(i, map_counts[i]);
  }
  ostringstream mean_string;
  mean_string << "Mean = " << 1.0 * n_good_mappings / n_pairs;
  map_counts_graph.add_text(0.75, 0.9, mean_string.str());

  // Hit length distribution
  const unsigned int max_read_length{
    static_cast<unsigned int>(read_length_hist.size())};
  PSGraph hit_length_graph{ps, ";Hit Length;N", Bounds(0, max_read_length)};
  PSHSeries<int, uint64_t> hit_length_series{hit_length_graph,
        max_read_length, "1 0 0", false};
  for (const MappingInfo & map : good_mappings) {
    hit_length_series.add_point(map.length());
    map_lengths.push_back(map.length());
  }

  // Template length histogram
  PSGraph template_length_graph{ps, ";Template Length;N",
        Bounds(0, max_template_length)};
  PSHSeries<int, uint64_t> template_length_series{
    template_length_graph, max_template_length / 10, "1 0 0", false};
  for (unsigned int i{0}; i != template_lengths.size(); ++i) {
    template_length_series.add_point(static_cast<int>(template_lengths[i]));
  }

  // Read length histogram
  PSGraph read_length_graph{ps, ";Read Length;N",
        Bounds(0, max_read_length)};
  PSHSeries<int, uint64_t> read_length_series{
    read_length_graph, max_read_length, "1 0 0", false};
  for (unsigned int i{0}; i != read_length_hist.size(); ++i) {
    read_length_series.add_value(i, read_length_hist[i]);
  }

#if 0
  // Hit Length vs Hit offset
  PSGraph hit_length_vs_offset_graph{ps, ";Hit Length; Hit Offset"};
  PSXYSeries hit_length_vs_offset_series{
    hit_length_vs_offset_graph, small_red_marker};
  for (const MappingInfo & map : good_mappings) {
    hit_length_vs_offset_series.add_point(
        map.length() + gen(), map.offset() + gen());
  }
#endif

  // Hit offset distribution
  PSGraph hit_offset_graph{ps, ";Hit Offset;N", Bounds(0, max_read_length)};
  PSHSeries<int, uint64_t> hit_offset_series{hit_offset_graph, max_read_length,
        "1 0 0", false};
  for (const MappingInfo & map : good_mappings) {
    hit_offset_series.add_point(map.offset());
  }

  // Read 2 distribution
  PSGraph read_2_graph{ps, ";Read;N", Bounds(0, 2, 0, nunset())};
  PSHSeries<int, uint64_t> read_2_series{read_2_graph, 2, "1 0 0", false};
  for (const MappingInfo & map : good_mappings) {
    read_2_series.add_point(map.read_2());
  }

  // Flipped distribution
  PSGraph flipped_graph{ps, ";Flipped;N", Bounds(0, 2, 0, nunset())};
  PSHSeries<int, uint64_t> flipped_series{flipped_graph, 2, "1 0 0", false};
  for (const MappingInfo & map : good_mappings) {
    flipped_series.add_point(map.flipped());
  }

  cout << "Summary Info:" << endl;
  sout << "Processed" << endl
       << n_pairs << "pairs with" << endl
       << n_dupe_pairs << "dupe pairs and" << endl
       << n_used_pairs << "used pairs and found" << endl
       << n_mappings << "mappings, of which" << endl
       << n_good_mappings << "were good," << endl
       << n_plotted_mappings << "were plotted," << endl
      // << n_joined << "were joined and" << endl
       << n_filtered << "were filtered:" << endl
       << n_discarded << "due to between joined," << endl
      // << n_filtered_length << "due to length and" << endl
       << n_filtered_excess << "due to excess and" << endl
       << n_dupe_mappings << "due to dupe pairs and" << endl
       << n_seen_before << "due to seen in pair" << endl;

  cout << "Done" << endl;

  ofstream stats_file{(output_prefix + "_stats.txt").c_str()};

  stats_file << "sample_id n_pairs n_dupe_pairs n_used_pairs n_maps "
      "n_good_maps n_used_maps n_overlap_maps percent_pair_covered "
      "mean_insert_length median_insert_length mean_map_length "
      "median_map_length maps_per_pair\n";

  const double percent_pair_covered{0.0};

  const double mean_insert_length{1.0 * accumulate(
      template_lengths.begin(), template_lengths.end(), 0UL) /
        template_lengths.size()};
  sort(template_lengths.begin(), template_lengths.end());
  const unsigned int median_insert_length{static_cast<unsigned int>(
      template_lengths[template_lengths.size() / 2])};

  const double mean_map_length{1.0 * accumulate(
      map_lengths.begin(), map_lengths.end(), 0UL) /
        map_lengths.size()};
  sort(map_lengths.begin(), map_lengths.end());
  const unsigned int median_map_length{static_cast<unsigned int>(
      map_lengths[map_lengths.size() / 2])};

  stats_file
      << output_prefix << " "
      << n_pairs << " "
      << n_dupe_pairs << " "
      << n_used_pairs << " "
      << n_mappings << " "
      << n_good_mappings << " "
      << n_plotted_mappings << " "
      << n_seen_before << " "
      << percent_pair_covered << " "
      << mean_insert_length << " "
      << median_insert_length << " "
      << mean_map_length << " "
      << median_map_length << " "
      << (n_pairs ? 1.0 * n_good_mappings / n_used_pairs : 0.0)
      << endl;

  return 0;
} catch (Error & e) {
  cout << "paa::Error:" << endl;
  cout << e.what() << endl;
  return 1;
}
catch (exception & e) {
  cout << "std::exception" << endl;
  cout << e.what() << endl;
  return 1;
} catch (...) {
  cout << "unknown exception was caught" << endl;
  return 1;
}
```
