## Supplementary material for "G-Graph: An interactive genomic graph viewer": MUMdex Source Code for G-Graph: smash.sh

```
#! /bin/bash

if [ $# != 7 ] && [ $# != 8 ] ; then
    echo usage: smash.sh mumdex_dir ref_file bins_dir n_bins,... sample_name fastq1.gz fastq2.gz [n_threads] 1>&2
    exit 1
fi

mumdex_dir="$1"
ref="$2"
bin_dir="$3"
n_bins="$4"
name="$5"
fastq1="$6"
fastq2="$7"
n_threads="${8:-12}"

if [ "$name" = "" ] ; then
    echo sample name is empty 1>&2
    exit 1
fi

if false && [ -e "$name" ] ; then
    echo sample name directory "(or file)" $name already exists 1>&2
    exit 1
fi

mkdir -p "$name"
if [ ! -d "$name" ] ; then
    echo could not create output directory $name 1>&2
    exit 1
fi
cd "$name"

if [ ! -d "$mumdex_dir" ] ; then
    echo mumdex dir $mumdex_dir does not exist 1>&2
    exit 1
fi

if [ ! -x "$mumdex_dir/smash" ] ; then
    echo smash executable $mumdex_dir/smash does not exist 1>&2
    exit 1
fi

if [ ! -f "$ref" ] ; then
    echo reference $ref is not a file 1>&2
    exit 1
fi

if [ ! -d "$ref.bin" ] ; then
    echo binary reference and suffix array $ref.bin does not exist 1>&2
    exit 1
fi

if [ ! -d "$bin_dir" ] ; then
    echo bin_dir directory does not exist 1>&2
    exit 1
fi

bad_pos="$bin_dir/bins.bad.bin"
if [ ! -f "$bad_pos" ] ; then
    echo bad positions file $bad_pos not found 1>&2
    exit 1
fi

if [ "$n_bins" = "" ] ; then
    echo n_bins command line argument is empty 1>&2
    exit 1
fi

bin_files=$(
echo $n_bins | perl -pe 's/[\s,]/\n/g' |
while read bin ; do
    bins_file="$bin_dir"/bins.$bin.txt
    if [ ! -e "$bins_file" ] ; then
        echo bins file $bins_file does not exist 1>&2
        exit 1
    fi
    echo "$bins_file"
done | perl -pe 's/\n/,/g')
bin_files=${bin_files%,}

if [ ! -e "$fastq1" ] ; then
    echo first fastq command line argument $fastq1 does not exist 1>&2
    exit 1
fi

if [ ! -e "$fastq2" ] ; then
    echo second fastq command line argument $fastq2 does not exist 1>&2
    exit 1
fi

echo running smash on sample $name in fastq files $fastq1, $fastq2 using reference $ref and bin files $bin_files

echo $mumdex_dir/smash "$ref" '<('zcat "$fastq1"')' '<('zcat "$fastq2"')' 20 4 1000 $n_threads "$name" "$bin_files" "$bad_pos" '>' "$name".out.txt '2>' "$name".err.txt

if $mumdex_dir/smash "$ref" <(zcat "$fastq1") <(zcat "$fastq2") 20 4 1000 $n_threads "$name" "$bin_files" "$bad_pos" > "$name".out.txt 2> "$name".err.txt ; then
    echo done with smash for sample $name
else
    echo smash failed
    exit 1
fi

# Do segmentation
for bin in $(echo $n_bins | perl -pe 's/,/ /g') ; do for results in $(find $PWD -name '*'_${bin}_bins_results.txt) ; do segments=${results/_results./_segments.} ; if [ ! -e $segments ] ; then $mumdex_dir/extract_cn_segments $ref $bin_dir/bins.$bin.txt $results > $segments 2> ${segments%.txt}.err.txt & fi ; done ; done
wait 

# Generate pdf from postscript
find $PWD -name '*.ps' | while read file ; do pdf=${file%.ps}.pdf ; if [ ! -e $pdf ] ; then echo generate $pdf ; $mumdex_dir/*/ps2pdf.sh $file ; fi ; done

exit 0


exit 0
```
