## Supplementary material for "G-Graph: An interactive genomic graph viewer": MUMdex Source Code for G-Graph: smash_qc.cpp

```
//
// smash_qc.cpp
//
// Copyright 2017 Peter Andrews @ CSHL
//

#include <algorithm>
#include <iostream>
#include <fstream>
#include <set>
#include <string>
#include <utility>

#include "./error.h"
#include "./files.h"

using std::cout;
using std::cerr;
using std::endl;
using std::ifstream;
using std::pair;
using std::set;
using std::string;
using std::to_string;

using paa::file_size;
using paa::Error;

int main(int argc, char ** argv)  try {
  // Check arguments
  --argc;
  if (argc != 1) throw Error("usage: smash_qc fastqs_list");
  ifstream fastqs{argv[1]};
  if (!fastqs) throw Error("Problem opening fastqs file") << argv[1];

  string sample;
  string fastq1;
  string fastq2;
  while (fastqs >> sample >> fastq1 >> fastq2) {
    cout << sample;
    // Can get info from fastq name?
    const string info{[&fastq1]() {
        auto sp = fastq1.find("WGS.");
        if (sp != string ::npos) {
          auto sp2 = fastq1.find("/", sp);
          if (sp2 != string::npos) {
            return fastq1.substr(sp + 4, sp2 - sp - 4);
          }
        }
        return string("-");
      }()};
    cout << " " << info;
    cout << " " << file_size(fastq1)
         << " " << file_size(fastq2);
    const string qc_name{sample + "_stats.txt"};
    ifstream qc{qc_name.c_str()};
    set<pair<string, string>> problems;
    if (qc) {
      qc.ignore(10000, '\n');
      string sample2;
      unsigned int n_pairs;
      unsigned int n_dupe_pairs;
      unsigned int n_used_pairs;
      unsigned int n_maps;
      unsigned int n_good_maps;
      unsigned int n_used_maps;
      unsigned int n_overlap_maps;
      double percent_covered;
      double mean_insert_length;
      unsigned int median_insert_length;
      double mean_map_length;
      unsigned int median_map_length;
      double maps_per_pair;
      qc >> sample2 >> n_pairs >> n_dupe_pairs >> n_used_pairs
         >> n_maps >> n_good_maps >> n_used_maps >> n_overlap_maps
         >> percent_covered >> mean_insert_length >> median_insert_length
         >> mean_map_length >> median_map_length >> maps_per_pair;
      if (!qc) throw Error("Problem parsing qc file") << qc_name;
      if (n_pairs * 2 < 2500000)
        problems.emplace("low_pairs", to_string(n_pairs));
      if (n_pairs * 0.05 < n_dupe_pairs)
        problems.emplace("many_dupes", to_string(1.0 * n_dupe_pairs / n_pairs));
      if (n_maps < 25000000)
        problems.emplace("few_maps", to_string(n_maps));
      // if (n_maps * 0.33 > n_good_maps)
      //   problems.emplace("few_good_maps", to_string());
      if (n_used_maps < 10000000)
        problems.emplace("few_used_maps", to_string(n_used_maps));
      if (n_good_maps * 0.3 < n_overlap_maps)
        problems.emplace("overlaps", to_string(1.0 * n_overlap_maps /
                                               n_good_maps));
      if (maps_per_pair < 3.25)
        problems.emplace("maps_per_pair", to_string(maps_per_pair));
      if (mean_insert_length <= 260)
        problems.emplace("mean_insert", to_string(mean_insert_length));
      if (median_map_length < 31)
        problems.emplace("short_maps", to_string(median_map_length));
      if (median_map_length > 42)
        problems.emplace("long_maps", to_string(median_map_length));
    } else {
      problems.emplace("smash_failed", "");
    }
    for (const pair<string, string> & problem : problems) {
      cout << " " << problem.first;
      if (problem.second.size()) {
        cout << "=" << problem.second;
      }
    }
    if (problems.empty()) cout << " passed";
    cout << endl;
  }

  return 0;
} catch (Error & e) {
  cerr << "paa::Error:" << endl;
  cerr << e.what() << endl;
  return 1;
} catch(std::exception & e) {
  cerr << e.what() << '\n';
  return 1;
} catch(...) {
  cerr << "Some exception was caught.\n";
  return 1;
}
```
