## Supplementary material for "G-Graph: An interactive genomic graph viewer": MUMdex Source Code for G-Graph: theoretical_bins.cpp

```
//
// theoretical_bins.cpp
//
// Determine theoretical bins and compare with empirical bin boundaries
//
// Copyright 2019 Peter Andrews @ CSHL
//

#include <algorithm>
#include <exception>
#include <fstream>
#include <iostream>
#include <map>
#include <sstream>
#include <string>
#include <utility>
#include <vector>

#include "cn.h"
#include "error.h"
#include "files.h"
#include "mumdex.h"
#include "psplot.h"
#include "stats.h"
#include "strings.h"

using std::cin;
using std::cout;
using std::cerr;
using std::endl;
using std::exception;
using std::ifstream;
using std::map;
using std::ofstream;
using std::ostringstream;
using std::pair;
using std::string;
using std::to_string;
using std::vector;

using paa::readable;
using paa::remove_substring;
using paa::Bin;
using paa::Bounds;
using paa::ChromosomeIndexLookup;
using paa::CN_abspos;
using paa::CN_Bin;
using paa::CN_Bins;
using paa::Error;
using paa::FinestBins;
using paa::MAD;
using paa::Mappability;
using paa::Marker;
using paa::PSDoc;
using paa::PSGraph;
using paa::PSHSeries;
using paa::PSXYSeries;
using paa::Reference;

int main(int argc, char* argv[]) try {
  --argc;
  if (argc != 2 && argc != 3)
    throw Error("usage: theoretical_bins ref n_bins [existing_bins]");

  // Process arguments
  const string ref_name{argv[1]};
  const Reference ref{ref_name};
  const ChromosomeIndexLookup lookup{ref};
  const Mappability mappability{ref};
  const CN_abspos cn_abspos{ref};
  const unsigned int X{ref.find_x_chromosome()};
  const unsigned int Y{ref.find_y_chromosome()};

  const unsigned int n_bins{static_cast<unsigned int>(atoi(argv[2]))};

  const string new_bins_prefix{"bins"};
  const string new_bins_name{
    new_bins_prefix + "." + to_string(n_bins) + ".txt"};
  if (readable(new_bins_name)) {
    cerr << "New bins file " << new_bins_name << " already exists" << endl;
  } else {
    const uint64_t min_map_length{15};
    const uint64_t max_map_length{40};
    string finebin_filenames;
    for (const bool male : {false, true}) {
      FinestBins bins{ref};
      bins.n_samples(1);
      bins.n_x(male ? 1 : 2);
      bins.n_y(male ? 1 : 0);
      for (const unsigned int chr : cn_abspos.chromosomes()) {
        cerr << male << " " << ref.name(chr) << endl;
        const unsigned int increment{
          (male && (chr == X || chr == Y)) ? 1u :
              ((!male && chr == Y) ? 0u : 2u)};
        for (unsigned int pos{0}; pos != ref.size(chr); ++pos) {
          const unsigned int abspos{cn_abspos(chr, pos)};
          if (mappability.low(abspos) >= min_map_length &&
              mappability.low(abspos) <= max_map_length) {
            bins[abspos] += increment;
          }
        }
      }
      const string bins_name{string("finebins.") + (male ? "male" : "female")};
      finebin_filenames += " ";
      finebin_filenames += bins_name;
      bins.save(bins_name);
    }

    // run empirical bins
    ostringstream command;
    command << "~/mumdex/empirical_bins " << ref_name << " " << n_bins
            << " " << new_bins_prefix << " " << 24 << finebin_filenames;
    cout << command.str() << endl;
    if (system(command.str().c_str()))
      throw Error("empirical bins failed");
  }

  if (argc == 3) {
    // List of previously determined good bins
    const string bins_name{argv[3]};
    const vector<Bin> existing_bins{load_bins(bins_name, ref)};
    if (n_bins != existing_bins.size())
      throw Error("Bin count mismatch for existing")
          << n_bins << existing_bins.size();
    const vector<Bin> new_bins{load_bins(new_bins_name, ref)};
    if (n_bins != new_bins.size())
      throw Error("Bin count mismatch for new") << n_bins << new_bins.size();
    uint64_t diff{0};
    for (unsigned int b{0}; b != n_bins; ++b) {
      const unsigned int abspos1{existing_bins[b].abspos()};
      const unsigned int abspos2{new_bins[b].abspos()};
      diff += (abspos1 > abspos2 ? abspos1 - abspos2 : abspos2 - abspos1);
    }
    const double aadiff{1.0 * diff / n_bins};
    cerr << "Average absolute bin position difference is "
         << aadiff << " or " << aadiff / cn_abspos.n_positions() << endl;
  }

  return 0;
} catch (Error & e) {
  cerr << "paa::Error:" << endl;
  cerr << e.what() << endl;
  return 1;
} catch (exception & e) {
  cerr << "std::exception" << endl;
  cerr << e.what() << endl;
  return 1;
} catch (...) {
  cerr << "unknown exception was caught" << endl;
  return 1;
}
```
