## Supplementary material for "G-Graph: An interactive genomic graph viewer": MUMdex Source Code for G-Graph: anchors.h

```
//
// anchors.h
//
// anchor and reference counts
//
// Copyright 2015 Peter Andrews @ CSHL
//

#ifndef PAA_ANCHORS_H
#define PAA_ANCHORS_H

#include <algorithm>
#include <iostream>
#include <string>
#include <vector>

#include "bed.h"
#include "error.h"
#include "files.h"
#include "mumdex.h"
#include "utility.h"

namespace paa {

#define USE_ZERO 0
#define USE_SUPPORT 0

class AnchorCountsCreator {
 public:
  AnchorCountsCreator(const MUMdex & mumdex,
                      const BedFile & bed,
                      const std::string & counts_dir) {
    const Reference & ref = mumdex.reference();
    const ChromosomeIndexLookup lookup{ref};
    read_ahead = true;
    const Mappability map{ref.fasta_file(), true};
    read_ahead = false;

    constexpr unsigned int low = 0;
    constexpr unsigned int high = 1;

    std::vector<PositionInfo> positions(ref.size());
    std::vector<unsigned int> coverage(ref.size());

    std::vector<unsigned int> seen_coverage;
    std::vector<unsigned int> seen_anchor[2];
    std::vector<unsigned int> seen_reference[2];
#if USE_ZERO
    std::vector<unsigned int> seen_zero[2];
    std::vector<unsigned int> seen_edge[2];
#endif
    std::cerr << "looping over pairs" << std::endl;
    Progress progress(mumdex.n_pairs(), 0.01, "Pair Loop");
    for (uint64_t pair_index = 0; pair_index != mumdex.n_pairs();
         ++pair_index) {
      progress();
      const auto pair = mumdex.pair(pair_index);
      if (pair.dupe()) continue;
      if (!pair.has_mums()) continue;
      seen_coverage.clear();
      for (const bool lowhigh : {false, true}) {
        seen_anchor[lowhigh].clear();
        seen_reference[lowhigh].clear();
#if USE_ZERO
        seen_zero[lowhigh].clear();
        seen_edge[lowhigh].clear();
#endif
      }
      bool read_1 = true;
      const auto sequences = mumdex.sequences(pair_index);
      const auto mums_begin = mumdex.mums_begin(pair_index);
      const auto mums_end = mumdex.mums_end(pair_index);
      for (auto mum_iter = mums_begin; mum_iter != mums_end; ++mum_iter) {
        const auto mum = *mum_iter;
        if (pair.bad(mum.read_2())) continue;

        // Is mum long enough?
        const auto offset = ref.offset(mum.chromosome());
        if (mum.length() < map.min(offset + mum.position0(), mum.length()) + 5)
          continue;
        // if (mum.length() < 25) continue;

        if (read_1 && mum.read_2()) {
          sort(seen_coverage.begin(), seen_coverage.end());
          for (const bool lowhigh : {false, true}) {
            sort(seen_anchor[lowhigh].begin(), seen_anchor[lowhigh].end());
            sort(seen_reference[lowhigh].begin(),
                 seen_reference[lowhigh].end());
#if USE_ZERO
            sort(seen_zero[lowhigh].begin(), seen_zero[lowhigh].end());
            sort(seen_edge[lowhigh].begin(), seen_edge[lowhigh].end());
#endif
          }
          read_1 = false;
        }

#if USE_ZERO
        // Determine if a zero or edge anchor
        bool zero_anchor[2] = { false, false };
        bool edge_anchor[2] = { true, true };
        for (auto other_mum_iter = mums_begin; other_mum_iter != mums_end;
             ++other_mum_iter) {
          if (mum_iter == other_mum_iter) continue;
          const auto other_mum = *other_mum_iter;
          if (mum.read_2() == other_mum.read_2()) {
            const bool is_high = (other_mum_iter > mum_iter) ^ mum.flipped();
            edge_anchor[is_high] = false;
            if (mum.flipped() == other_mum.flipped() &&
                mum.read_position0(pair.length(mum.read_2())) ==
                other_mum.read_position0(pair.length(other_mum.read_2()))) {
              zero_anchor[is_high] = true;
              if (other_mum_iter > mum_iter) break;
            }
          } else if (!mum.read_2() && other_mum.read_2()) {
            break;
          }
        }
#endif

        for (unsigned int mb = 0; mb != mum.length(); ++mb) {
          const unsigned int base_position = mum.position0() + mb;
          const unsigned int abspos = offset + base_position;
          const auto rb = mum.flipped() ?
              mum.offset() + mum.length() - mb - 1:
              mb + mum.offset();

          // Record basic coverage
          if (!mum.read_2() ||
              !binary_search(seen_coverage.begin(),
                             seen_coverage.end(), abspos)) {
            seen_coverage.push_back(abspos);
            ++coverage[abspos];
          }

          if (mb == 0) {  // potential in anchor
            // Does mum base touch the end of the read?
            if (mum.flipped()) {
              if (mum.touches_end()) {
                continue;
              }
            } else {
              if (mum.offset() == 0) {
                continue;
              }
            }

            // Is anchor due to an N?
            const auto ab = rb + (mum.flipped() ? 1 : -1);
            if (sequences[mum.read_2()][ab] == 'N') continue;

#if USE_SUPPORT
            // Record maximum support
            positions[abspos].max_support[low] =
                std::max(positions[abspos].max_support[low],
                         static_cast<uint8_t>(mum.length()));
#endif

#if USE_ZERO
            // Record if a zero anchor
            if (zero_anchor[low]) {
              if (mum.read_2()) {
                if (!binary_search(seen_zero[low].begin(),
                                   seen_zero[low].end(), abspos)) {
                  ++positions[abspos].counts[low].zero;
                }
              } else {
                ++positions[abspos].counts[low].zero;
                seen_zero[low].push_back(abspos);
              }
            }

            // Record or correct if an edge anchor
            const bool old_edge = mum.read_2() &&
                binary_search(seen_edge[low].begin(), seen_edge[low].end(),
                              abspos);
            if (edge_anchor[low]) {
              if (!old_edge) {
                ++positions[abspos].counts[low].edge;
                if (!mum.read_2()) seen_edge[low].push_back(abspos);
              }
            } else {
              if (old_edge) {
                --positions[abspos].counts[low].edge;
              }
            }
#endif
            // Record anchor, if not seen already in read 1
            if (mum.read_2()) {
              if (binary_search(seen_anchor[low].begin(),
                                seen_anchor[low].end(), abspos)) {
                continue;
              }
            } else {
              seen_anchor[low].push_back(abspos);
            }
            ++positions[abspos].counts[low].anchor;
          } else if (mb + 1 == mum.length()) {
            // Does mum base touch the end of the read?
            if (mum.flipped()) {
              if (mum.offset() == 0) {
                continue;
              }
            } else {
              if (mum.touches_end()) {
                continue;
              }
            }

            // Is anchor due to an N?
            const auto ab = rb + (mum.flipped() ? -1 : 1);
            if (sequences[mum.read_2()][ab] == 'N') continue;

#if USE_SUPPORT
            // Record maximum support
            positions[abspos].max_support[high] =
                std::max(positions[abspos].max_support[high],
                    static_cast<uint8_t>(mum.length()));
#endif
#if USE_ZERO
            // Record if a zero anchor
            if (zero_anchor[high]) {
              if (mum.read_2()) {
                if (!binary_search(seen_zero[high].begin(),
                                   seen_zero[high].end(), abspos)) {
                  ++positions[abspos].counts[high].zero;
                }
              } else {
                ++positions[abspos].counts[high].zero;
                seen_zero[high].push_back(abspos);
              }
            }

            // Record or correct if an edge anchor
            const bool old_edge = mum.read_2() &&
                binary_search(seen_edge[high].begin(),
                              seen_edge[high].end(), abspos);
            if (edge_anchor[high]) {
              if (!old_edge) {
                ++positions[abspos].counts[high].edge;
                if (!mum.read_2()) seen_edge[high].push_back(abspos);
              }
            } else {
              if (old_edge) {
                --positions[abspos].counts[high].edge;
              }
            }
#endif

            // Record anchor, if not seen already in read 1
            if (mum.read_2()) {
              if (binary_search(seen_anchor[high].begin(),
                                seen_anchor[high].end(),
                                abspos)) {
                continue;
              }
            } else {
              seen_anchor[high].push_back(abspos);
            }
            ++positions[abspos].counts[high].anchor;
          } else {
            if (mum.length() - mb >= map.low(abspos)) {
              if (mum.read_2()) {
                if (!binary_search(seen_reference[low].begin(),
                                   seen_reference[low].end(), abspos)) {
                  ++positions[abspos].counts[low].reference;
                }
              } else {
                ++positions[abspos].counts[low].reference;
                seen_reference[low].push_back(abspos);
              }
            }
            if (mb + 1 >= map.high(abspos)) {
              if (mum.read_2()) {
                if (!binary_search(seen_reference[high].begin(),
                                   seen_reference[high].end(), abspos)) {
                  ++positions[abspos].counts[high].reference;
                }
              } else {
                ++positions[abspos].counts[high].reference;
                seen_reference[high].push_back(abspos);
              }
            }
          }
        }
      }
    }

    std::cerr << "saving compressed counts" << std::endl;
    const auto n_bed_positions = [&bed]() {
      unsigned int n_positions = 0;
      for (const auto & interval : bed) {
        n_positions += interval.stop_pos - interval.start_pos;
      }
      return n_positions;
    }();
    CompressedInts<uint16_t, uint8_t> compressed
    {4 * n_bed_positions, bed.size()};
#if USE_SUPPORT
    OldMappedVector<uint8_t> max_support{2 * n_bed_positions};
#endif
    Progress progress2(bed.n_bases(), 0.01, "Save Loop");
    for (const auto & interval : bed) {
      const unsigned int chromosome = lookup[interval.chromosome];
      const unsigned int offset = ref.offset(chromosome);
      compressed.add_lookup_entry();
      for (unsigned int abspos = offset + interval.start_pos;
           abspos != offset + interval.stop_pos; ++abspos) {
        progress2();
        const auto & position = positions[abspos];
        compressed.add_int(coverage[abspos]);
        for (const bool lowhigh : {false, true}) {
#if USE_SUPPORT
          max_support.push_back(position.max_support[lowhigh]);
#endif
          compressed.add_int(position.counts[lowhigh].reference);
          compressed.add_int(position.counts[lowhigh].anchor);
#if USE_ZERO
          compressed.add_int(position.counts[lowhigh].zero);
          compressed.add_int(position.counts[lowhigh].edge);
#endif
        }
      }
    }
    compressed.save(counts_dir);
#if USE_SUPPORT
    max_support.save(counts_dir + "/max_support.bin");
#endif
  }

 private:
  class LowhighInfo {
   public:
    LowhighInfo() noexcept {}
    uint16_noo_t reference{0};
    uint16_noo_t anchor{0};
#if USE_ZERO
    uint16_noo_t zero{0};
    uint16_noo_t edge{0};
#endif
  };

  class PositionInfo {
   public:
    LowhighInfo counts[2]{ LowhighInfo(), LowhighInfo() };
#if USE_SUPPORT
    uint8_t max_support[2]{ 0, 0 };
#endif
  };
};

typedef struct {
  uint16_t reference[2]{static_cast<uint16_t>(0), static_cast<uint16_t>(0)};
  uint16_t anchor[2]{static_cast<uint16_t>(0), static_cast<uint16_t>(0)};
#if USE_ZERO
  uint16_t zero[2]{static_cast<uint16_t>(0), static_cast<uint16_t>(0)};
  uint16_t edge[2]{static_cast<uint16_t>(0), static_cast<uint16_t>(0)};
#endif
#if USE_SUPPORT
  uint8_t max_support[2]{static_cast<uint16_t>(0), static_cast<uint16_t>(0)};
#endif
} Pos_counts;

#if USE_ZERO
#define N_CT 5U
#else
#define N_CT 9U
#endif

class AnchorCounts {
 public:
  AnchorCounts(const std::string & bed_file,
               const std::string & counts_dir) :
    bed_{bed_file}, counts{counts_dir},
    n_counts_per_locus{N_CT},
#if USE_SUPPORT
    max_support{counts_dir + "/max_support.bin"},
    size_ {max_support.size() / 2}
#else
    size_ {static_cast<unsigned int>(counts.size() / N_CT)}
#endif
  {
      if (size_ * n_counts_per_locus != counts.size())
        throw Error("Counts size mismatch");
      if (!more()) throw Error("No counts data to load");
      load_bed_line(0);
      if (0) throw Error("dummy") << dummy;  // avoid unused warning
    }
  void load_bed_line(const unsigned int bed_line_) {
    current_bed = bed_line_;
    current_position = bed_[bed_line_].start_pos;
    n_loci_in = bed_.n_positions_to_line(bed_line_);
    counts.relocate(n_counts_per_locus * n_loci_in, bed_line_);
    load();
  }
  void load_position(const std::string & chr, const unsigned int pos0) {
    const unsigned int bed_line_ = bed_.find_bed_line(chr, pos0);
    if (bed_line_ == bed_.size())
      throw Error("Could not find position in bed:") << chr << pos0;
    load_bed_line(bed_line_);
    advance_n(bed_.n_positions_in_line(bed_line_, pos0));
  }
  void advance_n(const unsigned int n_to_advance) {
    for (unsigned int n = 0; n != n_to_advance; ++n) {
      load_next();
    }
  }
  const Pos_counts & current() const {
    return pos_counts;
  }
  void load_next() {
    ++n_loci_in;
    if (++current_position == bed_[current_bed].stop_pos) {
      ++current_bed;
      current_position = bed_[current_bed].start_pos;
    }
    load();
  }
  void load() {
    for (const bool high : {false, true}) {
      pos_counts.reference[high] = counts.next_int();
      pos_counts.anchor[high] = counts.next_int();
#if USE_ZERO
      pos_counts.zero[high] = counts.next_int();
      pos_counts.edge[high] = counts.next_int();
#endif
#if USE_SUPPORT
      pos_counts.max_support[high] = max_support[2 * n_loci_in + high];
#endif
    }
  }
  bool more() const {
    return n_loci_in + 1 < size();
  }
  bool good() const {
    return n_loci_in < size();
  }
  unsigned int bed_line() const { return current_bed; }
  const std::string & chromosome() const {
    return bed_[current_bed].chromosome;
  }
  unsigned int position0() const {
    return current_position;
  }
  unsigned int position1() const {
    return current_position + 1;
  }
  unsigned int size() const {
    return size_;
  }
  const BedFile & bed() const { return bed_; }
  uint8_t clipped_reference(const unsigned int position,
                            const bool high) const {
    return counts.clipped_result(position * 8 + high * 4);
  }
  uint8_t clipped_anchor(const unsigned int position, const bool high) const {
    return counts.clipped_result(position * 8 + high * 4 + 1);
  }
#if USE_ZERO
  uint8_t clipped_zero(const unsigned int position, const bool high) const {
    return counts.clipped_result(position * 8 + high * 4 + 2);
  }
  uint8_t clipped_edge(const unsigned int position, const bool high) const {
    return counts.clipped_result(position * 8 + high * 4 + 3);
  }
#endif

 private:
  BedFile bed_;
  CompressedInts<uint16_t, uint8_t> counts;
  unsigned int n_counts_per_locus{8};
#if USE_SUPPORT
  OldMappedVector<uint8_t> max_support;
#endif
  unsigned int size_{0};
  unsigned int n_loci_in{0};
  unsigned int current_bed{0};
  unsigned int current_position{0};
  Pos_counts pos_counts{};
  unsigned int dummy{0};
};

}  // namespace paa

#endif  // PAA_ANCHORS_H
```
