## Supplementary material for "G-Graph: An interactive genomic graph viewer": MUMdex Source Code for G-Graph: bed.h

```
//
// bed.h
//
// using bed files
//
// Copyright 2014 Peter Andrews @ CSHL
//

#ifndef PAA_BED_H
#define PAA_BED_H

#include <algorithm>
#include <iostream>
#include <fstream>
#include <sstream>
#include <vector>
#include <cstdint>
#include <map>
#include <string>

#include "error.h"
#include "utility.h"

namespace paa {

class GenomicInterval {
 public:
  GenomicInterval() { }
  GenomicInterval(const GenomicInterval & input,
                  const std::map<std::string, uint32_t> & chromosomes) {
    const unsigned int offset = chromosomes.at(input.chromosome);
    start_pos = offset + input.start_pos;
    stop_pos = offset + input.stop_pos;
  }
  std::istream & in(std::istream & stream) {
    stream >> chromosome >> start_pos >> stop_pos;
    if (stream && start_pos >= stop_pos)
      throw Error("Interval ordering error")
          << chromosome << start_pos << stop_pos;
    return stream;
  }
  std::ostream & out(std::ostream & stream) const {
    if (chromosome.size()) stream << chromosome << "\t";
    return stream << start_pos << "\t" << stop_pos;
  }
  bool operator!=(const GenomicInterval right) const {
    if (start_pos != right.start_pos || stop_pos != right.stop_pos ||
        chromosome != right.chromosome) {
      return true;
    } else {
      return false;
    }
  }
  bool operator<(const GenomicInterval right) const {
    if (chromosome == right.chromosome) {
      return start_pos < right.start_pos;
    } else {
      return chromosome < right.chromosome;
    }
  }
  GenomicInterval & operator=(const GenomicInterval &) = default;
  unsigned int size() const { return stop_pos - start_pos; }

  // private:
  std::string chromosome = "";
  unsigned int start_pos = 0;
  unsigned int stop_pos = 0;
};

class BedFile {
 public:
  explicit BedFile(const std::string & file_name, const bool do_sort = false) {
    std::ifstream input(file_name.c_str());
    GenomicInterval interval;
    if (!input) {
      std::istringstream sinput(file_name.c_str());
      // sinput >> interval;
      interval.in(sinput);
      if (sinput) {
        intervals.push_back(interval);
      } else {
        throw Error("Problem opening bed file or interpreting interval")
            << file_name;
      }
    }
    construct_bed(input, do_sort);
  }
  explicit BedFile(std::istream & input, const bool do_sort = false) {
    construct_bed(input, do_sort);
  }
  void construct_bed(std::istream & input, const bool do_sort = false) {
    GenomicInterval interval;
    while (input) {
      if (interval.in(input)) {
        intervals.push_back(interval);
      }
    }
    if (do_sort) {
      std::sort(intervals.begin(), intervals.end());
    }
    const bool show_bed = false;
    if (show_bed) {
      int n = 0;
      for (const GenomicInterval interval2 : intervals) {
        interval2.out(std::cerr);
        std::cerr << std::endl;
        ++n;
        if (n == 100) break;
      }
    }
    if (0)
      std::cerr << "Read " << intervals.size() << " genomic intervals from bed"
                << std::endl;
  }
  const GenomicInterval & operator[](const unsigned int index) const {
    return intervals[index];
  }
  std::vector<GenomicInterval>::const_iterator begin() const {
    return intervals.cbegin();
  }
  std::vector<GenomicInterval>::const_iterator end() const {
    return intervals.cend();
  }
  unsigned int size() const {
    return static_cast<unsigned int>(intervals.size());
  }
  unsigned int find_bed_line(const std::string & chr,
                             const uint64_t pos) const {
    unsigned int current_line = 0;
    while (current_line != intervals.size()) {
      const auto & interval = intervals[current_line];
      if (chr == interval.chromosome &&
          pos >= interval.start_pos &&
          pos < interval.stop_pos) {
        return current_line;
      }
      ++current_line;
    }
    return current_line;
  }
  unsigned int n_positions_to_line(const unsigned int bed_line) const {
    unsigned int current_line = 0;
    unsigned int n_positions_in = 0;
    while (current_line != bed_line) {
      n_positions_in += intervals[current_line].stop_pos -
          intervals[current_line].start_pos;
      ++current_line;
    }
    return n_positions_in;
  }
  unsigned int n_positions_in_line(const unsigned int bed_line,
                                   const unsigned int pos) const {
    return pos - intervals[bed_line].start_pos;
  }
  unsigned int positions_to_next_chromosome(const unsigned int bed_line,
                                            const unsigned int pos) const {
    unsigned int n_positions_in = intervals[bed_line].stop_pos - pos;
    const auto start_interval = intervals[bed_line];
    for (unsigned int current_bed = bed_line + 1;
         current_bed != size(); ++current_bed) {
      const auto & interval = intervals[current_bed];
      if (interval.chromosome != start_interval.chromosome) break;
      n_positions_in += interval.size();
    }

    return n_positions_in;
  }
  unsigned int positions_to_genome_end(const unsigned int bed_line,
                                       const unsigned int pos) const {
    unsigned int n_positions_in = intervals[bed_line].stop_pos - pos;
    const auto start_interval = intervals[bed_line];
    for (unsigned int current_bed = bed_line + 1;
         current_bed != size(); ++current_bed) {
      const auto & interval = intervals[current_bed];
      n_positions_in += interval.size();
    }

    return n_positions_in;
  }
  unsigned int n_bases() const {
    return positions_to_genome_end(0, 0);
  }

 private:
  std::vector<GenomicInterval> intervals{};
};

}  // namespace paa

#endif  // PAA_BED_H
```
