## Supplementary material for "G-Graph: An interactive genomic graph viewer": MUMdex Source Code for G-Graph: bridge_figure.cpp

```
//
// bridge_figure
//
// look for a specific bridge in one or more samples, make pretty pictures
//
// Copyright 2016 Peter Andrews @ CSHL
//

#include <algorithm>
#include <exception>
#include <iostream>
#include <fstream>
#include <sstream>
#include <string>
#include <utility>
#include <vector>

#include "bridges.h"
#include "error.h"
#include "fasta.h"
#include "mumdex.h"
#include "population.h"
#include "sequence.h"
#include "strings.h"
#include "utility.h"

using std::cerr;
using std::cout;
using std::endl;
using std::exception;
using std::max;
using std::min;
using std::ofstream;
using std::ostream;
using std::ostringstream;
using std::pair;
using std::string;
using std::to_string;
using std::vector;

using paa::complement;
using paa::replace_substring;
using paa::reverse_complement;
using paa::sout;
using paa::Base;
using paa::ChromosomeIndexLookup;
using paa::ConsensusSequence;
using paa::Error;
using paa::Family;
using paa::Mappability;
using paa::MUM;
using paa::MUMindex;
using paa::Pair;
using paa::Population;
using paa::Reference;
using paa::Sample;

#if 1
using MUMdex = paa::FileMUMdex;
using Bridges = paa::FileBridges;
#else
using MUMdex = paa::MUMdex;
using Bridges = paa::Bridges;
#endif

using Anchors = MUMdex::Anchors;
using Bridge = Bridges::Bridge;

string base_color(const char base) {
  switch (base) {
    case 'a': case 'A':
      return "acolor";
    case 'c': case 'C':
      return "ccolor";
    case 'g': case 'G':
      return "gcolor";
    case 't': case 'T':
      return "tcolor";
    default:
      return "bcolor";
  }
}

unsigned int base_value(const char base) {
  switch (base) {
    case 'a': case 'A':
      return 0;
    case 'c': case 'C':
      return 1;
    case 'g': case 'G':
      return 2;
    case 't': case 'T':
      return 3;
    default:
      return 4;
  }
}

const unsigned int page_width{792};
const unsigned int page_height{612};
const unsigned int header_font_size{22};
const unsigned int section_font_size{26};
const double sequence_font_size{18.2};
const double sequence_line_height{sequence_font_size * 1.7};
const double sequence_base_spacing{sequence_font_size * 0.8};
const unsigned int padding{20};
const double usable_width{page_width - 2.0 * padding};

// All reads on reference view
double convert(const int64_t pos,
               const int64_t ref_start, const int64_t ref_stop,
               const bool flip) {
  if (flip) {
    return padding + usable_width -
        usable_width * (pos - ref_start) / (ref_stop - ref_start);
  } else {
    return padding +
        usable_width * (pos - ref_start) / (ref_stop - ref_start);
  }
}

void show_section(ostream & ps, const double y_pos, const string & section) {
  ps << "/Helvetica findfont " << section_font_size
     << " scalefont setfont bcolor "
     << padding << " " << y_pos + 1.0 * sequence_line_height << " moveto "
     << "(" << section << ":) show" << endl;
}

void interlaced_counts(
    ostream & ps, const string & section, const string & sequence,
    const unsigned int mum1_start, const unsigned int mum1_stop,
    const unsigned int mum2_start, const unsigned int mum2_stop,
    const string & ref1, const string & ref2,
    const vector<vector<vector<unsigned int> > > & base_counts,
    const vector<vector<unsigned int> > & total_counts) {
  ps << "% interlaced_counts" << endl;
  const double top_pos{370};
  show_section(ps, top_pos, section);

  ps << "/Helvetica findfont " << sequence_font_size
     << " scalefont setfont 1 setlinewidth" << endl;
  unsigned int line{0};
  unsigned int start_b{0};
  for (unsigned int b{0}; b != sequence.size(); ++b) {
    const double x_pos{padding + (b - start_b + 0.5) * sequence_base_spacing};
    const double y_pos{top_pos - (4 * line + 1) * sequence_line_height};
    ps << x_pos << " " << y_pos << " moveto" << endl;

    for (const bool anchor2 : {false, true}) {
      const unsigned int max_count{*max_element(total_counts[anchor2].begin(),
                                                total_counts[anchor2].end())};
      ps << "  gsave 0 " << sequence_line_height * (anchor2 ? -1.0 : 0.8)
         << " rmoveto" << endl;
      const char * bases{"ACGTN"};
      const string & ref{anchor2 ? ref2 : ref1};
      double cumulative{0};
      for (unsigned int type{0}; type != 5; ++type) {
        const double base_frac{
          1.0 * base_counts[anchor2][b][type] / total_counts[anchor2][b]};
        const double height{base_frac * sequence_font_size};
        ps << "    gsave currentpoint newpath moveto ";
        const double width{0.9 * sequence_base_spacing *
              total_counts[anchor2][b] / max_count};
        ps << "-" << width / 2 << " " << cumulative - 3 << " rmoveto "
           << width << " 0 rlineto 0 " << height << " rlineto -" << width
           << " 0 rlineto closepath "
           << (ref[b] == bases[type] ? "gray" : base_color(bases[type]))
           << " fill stroke grestore" << endl;
        cumulative += height;
      }
      const char max_base{bases[max_element(base_counts[anchor2][b].begin(),
                                            base_counts[anchor2][b].end()) -
                                base_counts[anchor2][b].begin()]};
      ps << "  grestore" << endl << "  gsave " << "/Helvetica findfont " << 12
         << " scalefont setfont " << base_color(ref[b]) << " 0 "
         << (anchor2 ? -45 : 42) << " rmoveto " << "("
         << (max_base == ref[b] ? ' ' : ref[b])
         << ") dup stringwidth pop 2 div neg 0 rmoveto show grestore " << endl;
    }
    ps << "  gsave " << base_color(sequence[b]) << " (" << sequence[b]
       << ") dup stringwidth pop 2 div neg 0 rmoveto show ";
    if (b + 2 < sequence.size() &&
        x_pos + sequence_base_spacing > page_width - padding * 2) {
      start_b = b + 1;
      ++line;
      ps << "bcolor ( ...) show ";
    }
    ps << "grestore ";

    // Draw MUM underline
    if (b >= mum1_start && b < mum1_stop) {
      ps << "gsave currentpoint newpath moveto color1 "
         << "-" << sequence_base_spacing / 2 << " -6 rmoveto "
         << sequence_base_spacing << " 0 rlineto stroke grestore ";
    }
    if (b >= mum2_start && b < mum2_stop) {
      ps << "gsave currentpoint newpath moveto color2 "
         << "-" << sequence_base_spacing / 2 << " -4 rmoveto "
         << sequence_base_spacing << " 0 rlineto stroke grestore";
    }
    ps << endl;
  }
}

void show_read(ostream & ps, const double y_pos,
               const string & section, const string & sequence,
               const string & ref1, const string & ref2,
               const unsigned int mum1_start, const unsigned int mum1_stop,
               const unsigned int mum2_start, const unsigned int mum2_stop) {
  ps << "% show_read" << endl;
  const bool mum1_at_left{mum1_start < mum2_start};
  const unsigned int mum1_color_start{mum1_at_left ?
        0U : max(mum1_start, mum2_stop)};
  const unsigned int mum1_color_stop{mum1_at_left ? min(mum1_stop, mum2_start) :
        static_cast<unsigned int>(sequence.size())};
  const unsigned int mum2_color_start{mum1_at_left ?
        max(mum1_stop, mum2_start) : 0U};
  const unsigned int mum2_color_stop{mum1_at_left ?
        static_cast<unsigned int>(sequence.size()) :
        min(mum1_start, mum2_stop)};

  show_section(ps, y_pos, section);
  ps << "/Helvetica findfont " << sequence_font_size << " scalefont setfont"
     << endl << "1 setlinewidth" << endl;
  unsigned int line{0};
  unsigned int start_b{0};
  for (unsigned int b{0}; b != sequence.size(); ++b) {
    const double x_pos{
      padding + (b - start_b + 0.5) * sequence_base_spacing};
    ps << x_pos << " " << y_pos - line * sequence_line_height
       << " moveto gsave ";

    if (b >= mum1_color_start && b < mum1_color_stop) {
      ps << "color1 (" << (sequence[b] == ref1[b] ? ref1[b] :
                                static_cast<char>(tolower(sequence[b])));
    } else if (b >= mum2_color_start && b < mum2_color_stop) {
      ps << "color2 (" << (sequence[b] == ref2[b] ? ref2[b] :
                           static_cast<char>(tolower(sequence[b])));
    } else {
      const bool mums_overlap{mum1_at_left ?
            mum1_stop > mum2_start : mum2_stop > mum1_start};
      if (mums_overlap || (sequence[b] == ref1[b] && sequence[b] == ref2[b])) {
        ps << "color4 (" << sequence[b];
      } else {
        if (sequence[b] == ref1[b]) {
          ps << "color1 (" << ref1[b];
        } else if (sequence[b] == ref2[b]) {
          ps << "color2 (" << ref2[b];
        } else {
          ps << "color4 (" << static_cast<char>(tolower(sequence[b]));
        }
      }
    }

    ps << ") dup stringwidth pop 2 div neg 0 rmoveto show ";
    if (b + 2 < sequence.size() &&
        x_pos + sequence_base_spacing > page_width - padding * 2) {
      start_b = b + 1;
      ++line;
      ps << "bcolor ( ...) show ";
    }
    ps << "grestore ";
    if (b >= mum1_start && b < mum1_stop) {
      ps << "gsave currentpoint newpath moveto color1 "
         << "-" << sequence_base_spacing / 2 << " -6 rmoveto "
         << sequence_base_spacing << " 0 rlineto stroke grestore ";
    }
    if (b >= mum2_start && b < mum2_stop) {
      ps << "gsave currentpoint newpath moveto color2 "
         << "-" << sequence_base_spacing / 2 << " -4 rmoveto "
         << sequence_base_spacing << " 0 rlineto stroke grestore";
    }
    ps << endl;
  }
}

void show_reference(ostream & ps, const double y_pos,
                    const string & pre_command, const string & section,
                    const string & ref, const string & other,
                    const int mum1_start, const int mum1_stop,
                    const int mum2_start, const int mum2_stop) {
  ps << "% show_reference" << endl;
  show_section(ps, y_pos, section);
  ps << pre_command << endl;

  ps << "/Helvetica findfont " << sequence_font_size
     << " scalefont setfont 1 setlinewidth" << endl;
  unsigned int line{0};
  unsigned int start_b{0};
  for (unsigned int b{0}; b != ref.size(); ++b) {
    const double x_pos{padding + (b - start_b + 0.5) * sequence_base_spacing};
    ps << x_pos << " " << y_pos - line * sequence_line_height << " moveto "
       << "gsave ("
       << (ref[b] == other[b] ? ref[b] : static_cast<char>(tolower(ref[b])))
       << ") dup stringwidth pop 2 div neg 0 rmoveto show ";

    if (b + 2 < ref.size() &&
        x_pos + sequence_base_spacing > page_width - padding * 2) {
      start_b = b + 1;
      ++line;
      ps << "bcolor ( ...) show ";
    }
    ps << "grestore ";

    if (static_cast<int>(b) >= mum1_start && static_cast<int>(b) < mum1_stop) {
      ps << "gsave currentpoint newpath moveto color1 "
         << "-" << sequence_base_spacing / 2 << " -6 rmoveto "
         << sequence_base_spacing << " 0 rlineto stroke grestore ";
    }
    if (static_cast<int>(b) >= mum2_start && static_cast<int>(b) < mum2_stop) {
      ps << "gsave currentpoint newpath moveto color2 "
         << "-" << sequence_base_spacing / 2 << " -4 rmoveto "
         << sequence_base_spacing << " 0 rlineto stroke grestore";
    }
    ps << endl;
  }
}

vector<string> mum_colors{"0 0 0", "0 1 0", "0.8 0.8 0.8", "0 0.7 0",
      "0.5 0 0.5", "0.5 0.5 0.5", "0.5 0.5 0", "0 0.5 0.5"};

void read_pair_view(ostream & page, const string & header,
                    const MUMdex & mumdex, const Bridge & bridge,
                    const unsigned int chr[2], const string chrs[2],
                    const unsigned int pos[2], const MUM mum[2],
                    const std::array<std::string, 2> & sequences,
                    const uint64_t pair_number) {
  if (0) cout << pair_number << endl;
  const Pair pair{bridge.pair()};
  const Reference & ref{mumdex.reference()};

  //
  // Read-pair view page
  //
  page << "% read-pair view" << endl << header;

  // Determine reference boundaries, pair separation, orientation
  const int max_half_insert{2000};
  bool ref_flipped[2]{mum[0].flipped() != mum[0].read_2(),
        mum[1].flipped() != mum[1].read_2()};
  vector<MUM> mums_on_mate[2];
  for (uint64_t mum_index{mumdex.mums_start(bridge.pair_index())};
       mum_index != mumdex.mums_stop(bridge.pair_index()); ++mum_index) {
    const MUM amum{mumdex.mum(mum_index)};
    for (const bool anchor2 : {false, true}) {
      if (chr[anchor2] != amum.chromosome()) continue;
      const int read_pos{amum.read_position0(pair.length(amum.read_2()))};
      const int read_stop_pos{read_pos +
            static_cast<int>(pair.length(amum.read_2()))};
      // Is mum close enough to anchor?
      if (abs(read_stop_pos - static_cast<int>(pos[anchor2])) <
          max_half_insert ||
          abs(read_pos - static_cast<int>(pos[anchor2])) < max_half_insert) {
        if (amum.read_2() != mum[0].read_2()) {
          mums_on_mate[anchor2].push_back(amum);
        }
      }
    }
  }
  for (const bool anchor2 : {false, true}) {
    sort(mums_on_mate[anchor2].begin(), mums_on_mate[anchor2].end(),
         [](const MUM lhs, const MUM rhs) {
           if (lhs.offset() == rhs.offset()) {
             if (lhs.length() == rhs.length()) {
               return lhs < rhs;
             } else {
               return lhs.length() > rhs.length();
             }
           } else {
             return lhs.offset() > rhs.offset();
           }
         });
  }

  // Get insert size
  const double y_center{235};
  const double y_half_height_available{215};

  bool insert_set{false};
  int mate_distance{0};
  unsigned int default_spacing{23};
  bool best{mum[0].offset() < mum[1].offset()};
  if (mums_on_mate[best].size()) {
    const MUM m_mum{mums_on_mate[best].front()};
    if (m_mum.chromosome() == mum[best].chromosome() &&
        m_mum.flipped() != mum[best].flipped() &&
        abs(static_cast<int>(m_mum.position0()) -
            static_cast<int>(mum[best].position0())) < max_half_insert) {
      mate_distance = mum[best].read_position0(
          pair.length(mum[best].read_2())) -
          m_mum.read_position0(pair.length(m_mum.read_2()));
      insert_set = true;
    }
  }
  if (!insert_set && mums_on_mate[!best].size()) {
    const MUM m_mum{mums_on_mate[!best].front()};
    if (m_mum.chromosome() == mum[!best].chromosome() &&
        m_mum.flipped() != mum[!best].flipped() &&
        abs(static_cast<int>(m_mum.position0()) -
            static_cast<int>(mum[!best].position0())) <
        max_half_insert) {
      mate_distance = mum[!best].read_position0(
          pair.length(mum[!best].read_2())) -
          m_mum.read_position0(pair.length(m_mum.read_2()));
      insert_set = true;
    }
  }

  if (mate_distance < 0) mate_distance = -mate_distance;
  if (!insert_set) {
    page << "bcolor " << padding + usable_width / 2 << " " << y_center
         << " moveto (--?--) dup stringwidth pop 2 div neg -8 rmoveto show"
         << endl;
    mate_distance = pair.length(0) + default_spacing;
  }
  const unsigned int pair_length = max(
      pair.length(0), mate_distance + pair.length(1));

  unsigned int low_ref[2];
  unsigned int high_ref[2];

  for (const bool anchor2 : {false, true}) {
    const int read_length{static_cast<int>(pair.length(mum[anchor2].read_2()))};
    const int read_start_pos{mum[anchor2].read_position0(read_length)};

    if (mum[anchor2].flipped()) {
      high_ref[anchor2] = read_start_pos + read_length;
      if (read_start_pos - mate_distance > 0) {
        low_ref[anchor2] = read_start_pos - mate_distance;
      } else {
        low_ref[anchor2] = 0;
      }
    } else {
      if (read_start_pos > 0) {
        low_ref[anchor2] = read_start_pos;
      } else {
        low_ref[anchor2] = 0;
      }
      high_ref[anchor2] = read_start_pos + pair_length;
    }
  }
  if (chr[0] == chr[1]) {
    const unsigned int low_ref_offset{low_ref[0] < low_ref[1] ?
          low_ref[1] - low_ref[0] : low_ref[0] - low_ref[1]};
    const unsigned int high_ref_offset{high_ref[0] < high_ref[1] ?
          high_ref[1] - high_ref[0] : high_ref[0] - high_ref[1]};
    const unsigned int ref_offset{high_ref_offset > low_ref_offset ?
          high_ref_offset : low_ref_offset};
    if (ref_offset < 200) {
      high_ref[0] += ref_offset;
      high_ref[1] += ref_offset;
      if (low_ref[0] > ref_offset) {
        low_ref[0] -= ref_offset;
      } else {
        low_ref[0] = 0;
      }
      if (low_ref[1] > ref_offset) {
        low_ref[1] -= ref_offset;
      } else {
        low_ref[1] = 0;
      }
    }
  }

  const unsigned int ref1_length{high_ref[0] - low_ref[0]};
  const unsigned int ref2_length{high_ref[1] - low_ref[1]};
  const unsigned int extra_bases{5};
  const unsigned int show_bases{2 * extra_bases +
        max(pair_length, max(ref1_length, ref2_length))};
  if (ref1_length < show_bases) {
    const unsigned int half_deficit{(show_bases - ref1_length) / 2};
    if (low_ref[0] > half_deficit) {
      low_ref[0] -= half_deficit;
    } else {
      low_ref[0] = 0;
    }
    high_ref[0] = low_ref[0] + show_bases;
  }
  if (ref2_length < show_bases) {
    const unsigned int half_deficit{(show_bases - ref2_length) / 2};
    if (low_ref[1] > half_deficit) {
      low_ref[1] -= half_deficit;
    } else {
      low_ref[1] = 0;
    }
    high_ref[1] = low_ref[1] + show_bases;
  }

  // Draw references
  double ref_height{20};
  double ref_centers[2]{0.0, 0.0};
  for (const bool anchor2 : {false, true}) {
    page << "% reference " << anchor2 << endl;
    const double y_low{y_center +
          (anchor2 ? -1 : 1) * (y_half_height_available - 20) +
          (anchor2 ? 0 : -ref_height)};
    const double y_high{y_low + ref_height};
    ref_centers[anchor2] = (y_low + y_high) / 2;
    const double y_label{anchor2 ? y_low - 22 : y_high + 8};
    const double x_label{page_width / 2};
    const bool flip{ref_flipped[anchor2]};
    page << (anchor2 ? "color2" : "color1")
         << " (" << (flip ? "flipped " : "") << chrs[anchor2] << ") "
         << "dup stringwidth pop 2 div " << x_label << " exch sub "
         << " " << y_label << " moveto show" << endl;
    const unsigned int ref_start{low_ref[anchor2]};
    const unsigned int ref_stop{high_ref[anchor2]};
    for (unsigned int rpos{ref_start}; rpos != ref_stop; ++rpos) {
      const double x_low{convert(rpos, ref_start, ref_stop, flip)};
      const double x_high{convert(rpos + 1, ref_start, ref_stop, flip)};
      const char base{ref[chr[anchor2]][rpos]};
      const char fbase{flip ? complement(base) : base};
      page << "newpath "
           << x_low << " " << y_low << " moveto "
           << x_high << " " << y_low << " lineto "
           << x_high << " " << y_high << " lineto "
           << x_low << " " << y_high << " lineto "
           << "closepath " << base_color(fbase) << " fill stroke " << endl;
    }
  }

  const double mum_height{4};

  const double base_spacing{usable_width / show_bases};
  // const double pair_width{pair_length * base_spacing};
  // Pair alignment
  const int a_mum_read_stop{mum[best].read_start_position() +
        (mum[best].flipped() ? -1 : 1) *
        static_cast<int>(pair.length(mum[best].read_2())) +
        (ref_flipped[best] == mum[best].read_2() ? 0 : 1)};

  const double a_mum_read_stop_x{
    convert(a_mum_read_stop, low_ref[best], high_ref[best], ref_flipped[best])};
  const double pair_x_start_trial{a_mum_read_stop_x -
        (mum[best].read_2() ? mate_distance : pair.length(0)) * base_spacing };
  const double pair_x_start{
    (pair_x_start_trial < padding ||
     pair_x_start_trial + pair_length * base_spacing >
     padding + usable_width) ?
        (padding + extra_bases * base_spacing) : pair_x_start_trial};
  const double pair_height{20};
  unsigned int color_index{0};
  ostringstream reads;
  vector<vector<std::pair<unsigned int, unsigned int> > > levels1;
  vector<vector<std::pair<unsigned int, unsigned int> > > levels2;
  for (const bool read2 : {false, true}) {
    page << "% read " << read2 << endl;

    const double y_offset{static_cast<unsigned int>(
        mate_distance) > pair.length(0) + 2 ? 0 : read2 ? -20.0 : 20.0};
    const double mate_center{y_center + y_offset};
    const double y_low{mate_center - pair_height / 2};
    const double y_high{mate_center + pair_height / 2};
    const double mate_x_start{pair_x_start +
          (read2 ? mate_distance * base_spacing : 0)};

    // MUM lines
    vector<vector<std::pair<unsigned int, unsigned int> > > levels;
    for (uint64_t mum_index{mumdex.mums_start(bridge.pair_index())};
         mum_index != mumdex.mums_stop(bridge.pair_index()); ++mum_index) {
      const MUM amum{mumdex.mum(mum_index)};
      if (amum.read_2() != read2) continue;
      const bool on_1{amum.chromosome() == chr[0] &&
            amum.position0() >= low_ref[0] &&
            amum.position0() + amum.length() <= high_ref[0]};
      const bool on_2{amum.chromosome() == chr[1] &&
            amum.position0() >= low_ref[1] &&
            amum.position0() + amum.length() <= high_ref[1]};
      if (!on_1 && !on_2) continue;

      string mum_color;
      if (amum == mum[0]) {
        mum_color = "color1";
      } else if (amum == mum[1]) {
        mum_color = "color2";
      } else {
        mum_color = mum_colors[color_index++ % mum_colors.size()] +
            " setrgbcolor";
      }
      page << mum_color << endl;
      const unsigned int b_low{read2 ?
            (pair.length(read2) - amum.offset() - amum.length()) :
            amum.offset()};
      const double x_low_f{mate_x_start + b_low * base_spacing};
      const double x_high_f{x_low_f + amum.length() * base_spacing};
      const double x_low{(amum.flipped() == read2) ? x_low_f : x_high_f};
      const double x_high{(amum.flipped() == read2) ? x_high_f : x_low_f};

      unsigned int unoccupied_level{0};
      const unsigned int m_stop{amum.offset() + amum.length()};
      for (unsigned int l{0}; l != levels.size(); ++l) {
        bool occupied{false};
        for (unsigned int m{0}; m != levels[l].size(); ++m) {
          const unsigned int occ_start{levels[l][m].first};
          const unsigned int occ_stop{levels[l][m].second};
          if ((amum.offset() >= occ_start && amum.offset() < occ_stop) ||
              (m_stop > occ_start && m_stop <= occ_stop) ||
              (occ_start >= amum.offset() && occ_start < m_stop) ||
              (occ_stop > amum.offset() && occ_stop <= m_stop)) {
            occupied = true;
          }
        }
        if (occupied) {
          ++unoccupied_level;
        } else {
          break;
        }
      }
      if (unoccupied_level == levels.size()) {
        levels.resize(levels.size() + 1);
      }
      levels[unoccupied_level].emplace_back(amum.offset(), m_stop);

      const double y_level{mate_center + (read2 ? -1 : 1) *
            (pair_height / 2 + 2 * (unoccupied_level + 1) * mum_height)};
      page << "[] 0 setdash " << mum_height << " setlinewidth ";
      page << "newpath " << x_low << " " << y_level << " moveto "
           << x_high << " " << y_level << " lineto stroke" << endl;

      if (on_1) {
        unsigned int unoccupied_level1{0};
        const unsigned int am_stop{amum.position0() + amum.length()};
        for (unsigned int l{0}; l != levels1.size(); ++l) {
          bool occupied{false};
          for (unsigned int m{0}; m != levels1[l].size(); ++m) {
            const unsigned int occ_start{levels1[l][m].first};
            const unsigned int occ_stop{levels1[l][m].second};
            if ((amum.position0() >= occ_start &&
                 amum.position0() < occ_stop) ||
                (am_stop > occ_start && am_stop <= occ_stop) ||
                (occ_start >= amum.position0() && occ_start < am_stop) ||
                (occ_stop > amum.position0() && occ_stop <= am_stop)) {
              occupied = true;
            }
          }
          if (occupied) {
            ++unoccupied_level1;
          } else {
            break;
          }
        }
        if (unoccupied_level1 == levels1.size()) {
          levels1.resize(levels1.size() + 1);
        }
        levels1[unoccupied_level1].emplace_back(amum.position0(), am_stop);

        const double y_level1{ref_centers[0] -
              (ref_height + 2 * (unoccupied_level1 + 0.5) * mum_height)};
        const double x_start{
          convert(amum.position0(), low_ref[0], high_ref[0], ref_flipped[0])};
        const double x_stop{
          convert(amum.position0() + amum.length(),
                  low_ref[0], high_ref[0], ref_flipped[0])};
        page << "[] 0 setdash " << mum_height << " setlinewidth "
             << "newpath " << x_start << " " << y_level1 << " moveto "
             << x_stop << " " << y_level1 << " lineto stroke" << endl;
        page << "[2 2] 0 setdash 1 setlinewidth "
             << "newpath " << x_start << " " << y_level1 << " moveto "
             << x_low << " " << y_level << " lineto stroke" << endl;
        page << "[2 2] 0 setdash 1 setlinewidth "
             << "newpath " << x_stop << " " << y_level1 << " moveto "
             << x_high << " " << y_level << " lineto stroke" << endl;
      }

      if (on_2) {
        unsigned int unoccupied_level2{0};
        const unsigned int am_stop{amum.position0() + amum.length()};
        for (unsigned int l{0}; l != levels2.size(); ++l) {
          bool occupied{false};
          for (unsigned int m{0}; m != levels2[l].size(); ++m) {
            const unsigned int occ_start{levels2[l][m].first};
            const unsigned int occ_stop{levels2[l][m].second};
            if ((amum.position0() >= occ_start &&
                 amum.position0() < occ_stop) ||
                (am_stop > occ_start && am_stop <= occ_stop) ||
                (occ_start >= amum.position0() && occ_start < am_stop) ||
                (occ_stop > amum.position0() && occ_stop <= am_stop)) {
              occupied = true;
            }
          }
          if (occupied) {
            ++unoccupied_level2;
          } else {
            break;
          }
        }
        if (unoccupied_level2 == levels2.size()) {
          levels2.resize(levels2.size() + 1);
        }
        levels2[unoccupied_level2].emplace_back(amum.position0(), am_stop);

        const double y_level2{ref_centers[1] +
              (ref_height + 2 * (unoccupied_level2 + 0.5) * mum_height)};

        const double x_start{
          convert(amum.position0(), low_ref[1], high_ref[1], ref_flipped[1])};
        const double x_stop{
          convert(amum.position0() + amum.length(),
                  low_ref[1], high_ref[1], ref_flipped[1])};
        page << "[] 0 setdash " << mum_height << " setlinewidth "
             << "newpath " << x_start << " " << y_level2 << " moveto "
             << x_stop << " " << y_level2 << " lineto stroke" << endl;
        page << "[2 2] 0 setdash 1 setlinewidth "
             << "newpath " << x_start << " " << y_level2 << " moveto "
             << x_low << " " << y_level << " lineto stroke" << endl;
        page << "[2 2] 0 setdash 1 setlinewidth "
             << "newpath " << x_stop << " " << y_level2 << " moveto "
             << x_high << " " << y_level << " lineto stroke" << endl;
      }
    }

    // Read sequence
    for (unsigned int b{0}; b != pair.length(read2); ++b) {
      const double x_low{mate_x_start + b * base_spacing};
      const double x_high{x_low + base_spacing};
      const unsigned int bb{read2 ? pair.length(read2) - b - 1 : b};
      const char base{sequences[read2][bb]};
      const char fbase{read2 ? complement(base) : base};
      reads << "newpath "
            << x_low << " " << y_low << " moveto "
            << x_high << " " << y_low << " lineto "
            << x_high << " " << y_high << " lineto "
            << x_low << " " << y_high << " lineto "
            << "closepath " << base_color(fbase) << " fill stroke " << endl;
    }
    // border
    reads << "[] 0 setdash 1 setlinewidth bcolor newpath "
          << mate_x_start << " " << y_low << " moveto "
          << mate_x_start + pair.length(read2) * base_spacing
          << " " << y_low << " lineto "
          << mate_x_start + pair.length(read2) * base_spacing
          << " " << y_high << " lineto " << mate_x_start << " " << y_high
          << " lineto " << "closepath stroke" << endl;
  }
  page << reads.str();
}

int main(int argc, char* argv[])  try {
  // Check usage
  if (--argc != 13) {
    throw Error("usage: bridge_figure samples_dir pop_file "
                "chr1 pos1 high1 chr2 pos2 high2 "
                "inv offset type family member");
  }

  // Process command line arguments and setup
  const string family_name{argv[12]};
  const string samples_dir{argv[1]};
  const Population pop{argv[2]};
  const string first_sample_name{pop.sample(pop.samples(family_name).front())};
  const string mumdex_name{samples_dir + "/" + first_sample_name + "/mumdex"};
  const MUMdex first_mumdex{mumdex_name};
  const Reference & ref{first_mumdex.reference()};
  const Mappability mappability(mumdex_name);
  const ChromosomeIndexLookup chr_lookup{ref};
  const string chrs[2]{argv[3], argv[6]};
  const unsigned int chr[2]{chr_lookup[chrs[0]], chr_lookup[chrs[1]]};
  const unsigned int pos[2]{static_cast<unsigned int>(atoi(argv[4])),
        static_cast<unsigned int>(atoi(argv[7]))};
  const bool high[2]{static_cast<bool>(atoi(argv[5])),
        static_cast<bool>(atoi(argv[8]))};
  const int64_t invariant{atol(argv[9])};
  const string offset{argv[10]};
  const string extra{argv[11]};
  const vector<Sample> family_samples{pop.samples(family_name)};
  const Family family{pop.family(family_samples.front())};
  const string member{argv[13]};
  const vector<Sample> & event_samples{
    member == "proband" ? pop.proband(family) :
        (member == "sibling" ? pop.sibling(family) : pop.child(family))};

  // Pages
  vector<pair<float, vector<string>>> sample_pages;
  ostringstream page;
  ostringstream main_page;

  // Get information on all bridges for main page plot boundaries, etc.
  const unsigned int max_ref_half_size{700};
  unsigned int n_bridges{0};
  unsigned int max_len_1{0};
  unsigned int max_len_2{0};
  unsigned int max_dist[2]{0, 0};
  unsigned int main_low_ref[2]{static_cast<unsigned int>(-1),
        static_cast<unsigned int>(-1)};
  unsigned int main_high_ref[2]{0, 0};
  // Loop over samples
  for (const Sample sample : event_samples) {
    // Load the mumdex for the sample
    const string sample_name{pop.sample(sample)};
    const MUMdex mumdex{samples_dir + "/" + sample_name + "/mumdex"};
    // Loop over bridges found
    for (const Bridge & bridge : Bridges{chr[0], pos[0], high[0],
            chr[1], pos[1], high[1], invariant, mumdex}) {
      max_len_1 = max(max_len_1, bridge.mum1().length());
      max_len_2 = max(max_len_2, bridge.mum2().length());
      ++n_bridges;

      // Determine main figure X range
      for (const bool anchor2 : {false, true}) {
        for (uint64_t mum_index{mumdex.mums_start(bridge.pair_index())};
             mum_index != mumdex.mums_stop(bridge.pair_index());
             ++mum_index) {
          const MUM amum{mumdex.mum(mum_index)};
          if (amum.chromosome() != chr[anchor2]) continue;
          if (amum.position0() >= pos[anchor2] + max_ref_half_size) continue;
          const unsigned int m_stop{amum.position0() + amum.length()};
          if (m_stop + max_ref_half_size <= pos[anchor2]) continue;
          if (main_low_ref[anchor2] > amum.position0()) {
            main_low_ref[anchor2] = amum.position0();
          }
          if (main_high_ref[anchor2] < m_stop) {
            main_high_ref[anchor2] = m_stop;
          }
        }
        if (main_high_ref[anchor2] != 0) {
          max_dist[anchor2] = max(pos[anchor2] - main_low_ref[anchor2],
                                  main_high_ref[anchor2] - pos[anchor2]);
        }
      }
    }
  }
  const unsigned int ref_half_size{max(max_dist[0], max_dist[1]) + 10};

  // Loop over samples
  for (const Sample sample : event_samples) {
    // Load the mumdex for the sample
    const string sample_name{pop.sample(sample)};
    const MUMdex mumdex{samples_dir + "/" + sample_name + "/mumdex"};

    // Loop over bridges found
    for (const Bridge & bridge : Bridges{chr[0], pos[0], high[0],
          chr[1], pos[1], high[1], invariant, mumdex}) {
      // Bridge Pair
      const Pair pair{bridge.pair()};
      if (pair.dupe()) continue;
      const uint64_t pair_number{sample_pages.size() + 1};

      // Anchor mums for bridge
      const MUM mum[2]{bridge.mum1(), bridge.mum2()};

      // Read Pair sequences
      const std::array<std::string, 2> sequences(
          mumdex.sequences(bridge.pair_index()));

      // Closure to make header
      auto make_header = [&mappability, &ref, &family_name, &extra, n_bridges,
                          invariant, offset, chr, chrs, pos, high]
          (const string & sname, const string & member_,
           const unsigned int l1, const unsigned int l2,
           const uint64_t pn) {
        ostringstream header;
        header << "gsave" << endl;  // restored when page is finalized
        const double hpadding{10};
        const double header_line_spacing{1.4 * header_font_size};
        const double header_height{2 * hpadding + 3.6 * header_line_spacing};
        const double first_line_height{
          page_height - padding - header_font_size};
        const double text_left{padding + hpadding};
        // Box
        header << "0.9 0.9 0.9 setrgbcolor newpath "
               << padding << " " << page_height - padding << " moveto "
               << "0 " << -header_height << " rlineto "
               << page_width - 2 * padding << " 0 rlineto "
               << "0 " << header_height << " rlineto "
               << "closepath gsave fill grestore bcolor stroke" << endl;
        // Text Size
        header << "/Helvetica findfont " << header_font_size
               << " scalefont setfont" << endl;
        // Line 1 sample / bridge type
        header << text_left << " " << first_line_height << " moveto ("
               << family_name << " " << sname << " "
               << member_ << " " << extra << ") show" << endl;
        // Line 2 anchor 1
        header << text_left << " " << first_line_height - header_line_spacing
               << " moveto color1 (" << chrs[0] << " " << pos[0] << " "
               << (high[0] ? "high" : "low") << " "
               << mappability.low_high(high[0], ref.abspos(chr[0], pos[0]))
               << " / " << l1 << ") show" << endl;
        // Line 3 anchor 2
        header << text_left << " "
               << first_line_height - 2 * header_line_spacing
               << " moveto color2 (" << chrs[1] << " " << pos[1] << " "
               << (high[1] ? "high" : "low") << " "
               << mappability.low_high(high[1], ref.abspos(chr[1], pos[1]))
               << " / " << l2 << ") show" << endl;
        // Line 4 other info
        header << text_left << " "
               << first_line_height - 3 * header_line_spacing
               << " moveto bcolor (count " << n_bridges << " invariant "
               << invariant << " offset " << offset << " ) show" << endl;
        // Pair number
        if (0) {
          header << page_width - padding
                 << " " << first_line_height << " moveto ("
                 << pn
                 << " ) dup stringwidth pop neg 0 rmoveto show" << endl;
        }
        return header.str();
      };
      const string header{make_header(sample_name, pop.member(sample),
                                      mum[0].length(), mum[1].length(),
                                      pair_number)};

      // Read sequences
      string sequence{sequences[mum[0].read_2()]};
      const bool flip{mum[0].offset() > mum[1].offset()};
      if (flip) reverse_complement(&sequence);

      // MUM positions
      const unsigned int mum1_start{static_cast<unsigned int>(
          flip ? sequence.size() - mum[0].offset() - mum[0].length() :
          mum[0].offset())};
      const unsigned int mum1_stop{static_cast<unsigned int>(
          flip ? sequence.size() - mum[0].offset() :
          mum[0].offset() + mum[0].length())};
      const unsigned int mum2_start{static_cast<unsigned int>(
          flip ? sequence.size() - mum[1].offset() - mum[1].length() :
          mum[1].offset())};
      const unsigned int mum2_stop{static_cast<unsigned int>(
          flip ? sequence.size() - mum[1].offset() :
          mum[1].offset() + mum[1].length())};

      // Holds pages
      sample_pages.emplace_back(static_cast<int>(mum1_stop + mum2_start) -
                                static_cast<int>(sequence.length()) + 0.1,
                                vector<string>());

      //
      // Read-Pair view page
      //
      read_pair_view(page, header, mumdex, bridge, chr, chrs, pos, mum,
                     sequences, pair_number);
      sample_pages.back().second.push_back(page.str());
      page.str("");

      //
      // Read view page
      //
      const double center_height{page_height / 2.0 - 10};
      page << "% read view page" << endl;
      page << header;

      // Reference 1 and positions
      const int ref1_pos{mum[0].read_position0(sequence.size())};
      string ref1{ref.subseq(mum[0].chromosome(), ref1_pos,
                             static_cast<unsigned int>(
                                 ref1_pos + sequence.length()))};
      int ref1_mum2_start{static_cast<int>(mum[1].position0()) - ref1_pos};
      int ref1_mum2_stop{ref1_mum2_start + static_cast<int>(mum[1].length())};
      const bool flip1{flip ? !mum[0].flipped() : mum[0].flipped()};
      if (flip1) {
        const int temp = ref1_mum2_stop;
        ref1_mum2_stop = static_cast<int>(sequence.size()) - ref1_mum2_start;
        ref1_mum2_start = static_cast<int>(sequence.size()) - temp;
        reverse_complement(&ref1);
      }
      if (mum[0].chromosome() != mum[1].chromosome()) {
        ref1_mum2_start = static_cast<unsigned int>(sequence.size());
        ref1_mum2_stop = static_cast<unsigned int>(sequence.size());
      }
      show_reference(page, center_height + 3 * section_font_size, "color1",
                     string(flip1 ? "Flipped " : "") + chrs[0] +
                     " " + to_string(ref1_pos), ref1, sequence,
                     mum1_start, mum1_stop, ref1_mum2_start, ref1_mum2_stop);

      // Reference 2 and positions
      const int ref2_pos{mum[1].read_position0(sequence.size())};
      string ref2{ref.subseq(mum[1].chromosome(), ref2_pos,
                             static_cast<unsigned int>(
                                 ref2_pos + sequence.length()))};
      int ref2_mum1_start{static_cast<int>(mum[0].position0()) - ref2_pos};
      int ref2_mum1_stop{ref2_mum1_start + static_cast<int>(mum[0].length())};
      const bool flip2{flip ? !mum[1].flipped() : mum[1].flipped()};
      if (flip2) {
        const int temp = ref2_mum1_stop;
        ref2_mum1_stop = static_cast<int>(sequence.size()) - ref2_mum1_start;
        ref2_mum1_start = static_cast<int>(sequence.size()) - temp;
        reverse_complement(&ref2);
      }
      if (mum[0].chromosome() != mum[1].chromosome()) {
        ref2_mum1_start = static_cast<unsigned int>(sequence.size());
        ref2_mum1_stop = static_cast<unsigned int>(sequence.size());
      }
      show_reference(page, center_height - 8 * section_font_size, "color2",
                     string(flip2 ? "Flipped " : "") + chrs[1] +
                     " " + to_string(ref2_pos), ref2, sequence,
                     ref2_mum1_start, ref2_mum1_stop, mum2_start, mum2_stop);

      // The read in the center
      show_read(page, center_height - 2.5 * section_font_size,
                flip ? "flipped read" : "read", sequence, ref1, ref2,
                mum1_start, mum1_stop, mum2_start, mum2_stop);

      sample_pages.back().second.push_back(page.str());
      page.str("");

      //
      // Main page
      //
      const double y_center{230};
      const double y_half_height_available{220};
      const double y_half_height{y_half_height_available *
            (n_bridges < 6 ? 0.5 : 1.0)};
      const double ref_half_spacing{15};
      const double pair_spacing{y_half_height / (2 + n_bridges)};
      const double mum_width{pair_spacing / 8};
      const double ref_height{10};

      string sample_names{sample_name};
      const double ref_font_size{16};
      if (pair_number == 1) {
        if (event_samples.size() > 1) {
          sample_names = "";
          for (const Sample esample : event_samples) {
            if (sample_names.size()) sample_names += " ";
            sample_names += pop.sample(esample);
          }
        }
        main_page << "% main page" << endl;
        main_page << make_header(sample_names, member, max_len_1, max_len_2, 0);
        main_page << mum_width << " setlinewidth" << endl;
        main_page << "/Helvetica findfont " << ref_font_size
                  << " scalefont setfont newpath "
                  << padding << " " << y_center - y_half_height
                  << " moveto "
                  << page_width - padding << " " << y_center - y_half_height
                  << " lineto "
                  << page_width - padding << " " << y_center + y_half_height
                  << " lineto "
                  << padding << " " << y_center + y_half_height
                  << " lineto closepath clip" << endl;
      }

      for (const bool anchor2 : {false, true}) {
        const unsigned int ref_start{pos[anchor2] > ref_half_size ?
              pos[anchor2] - ref_half_size : 0};
        const unsigned int ref_stop{pos[anchor2] + ref_half_size <
              ref.size(chr[anchor2]) ? pos[anchor2] + ref_half_size :
              ref.size(chr[anchor2])};
        if (pair_number == 1) {
          const double y_low{y_center +
                (anchor2 ? -(ref_half_spacing + ref_height): ref_half_spacing)};
          const double y_high{y_low + ref_height};
          const double y_label{y_center + (anchor2 ? -0.8 * ref_font_size: 0)};
          const double x_label{page_width / 2};
          const double x_label_offset{100.0};
          main_page << (anchor2 ? "color2" : "color1")
                    << " (" << chrs[anchor2] << ") "
                    << "dup stringwidth pop "
                    << x_label << " " << x_label_offset << " "
                    << (anchor2 ? "add" : "sub")
                    << " exch " << (anchor2 ? "sub" : "pop")
                    << " " << y_label << " moveto show" << endl;
          for (unsigned int rpos{ref_start}; rpos != ref_stop; ++rpos) {
            const double x_low{convert(rpos, ref_start, ref_stop, false)};
            const double x_high{convert(rpos + 1, ref_start, ref_stop, false)};
            main_page << "newpath "
                      << x_low << " " << y_low << " moveto "
                      << x_high << " " << y_low << " lineto "
                      << x_high << " " << y_high << " lineto "
                      << x_low << " " << y_high << " lineto "
                      << "closepath " << base_color(ref[chr[anchor2]][rpos])
                      << " fill stroke " << endl;
          }
        }
        const double y_offset{ref_half_spacing + ref_height +
              pair_spacing * pair_number};
        vector<vector<std::pair<unsigned int, unsigned int> > > levels;
        for (uint64_t mum_index{mumdex.mums_start(bridge.pair_index())};
             mum_index != mumdex.mums_stop(bridge.pair_index()); ++mum_index) {
          const MUM amum{mumdex.mum(mum_index)};
          if (amum.chromosome() != chr[anchor2]) continue;
          if (amum.position0() >= ref_stop) continue;
          const unsigned int m_stop{amum.position0() + amum.length()};
          if (m_stop <= ref_start) continue;
          const double y_main_level{y_center + (anchor2 ? -1 : 1) * y_offset};
          unsigned int unoccupied_level{0};
          for (unsigned int l{0}; l != levels.size(); ++l) {
            bool occupied{false};
            for (unsigned int m{0}; m != levels[l].size(); ++m) {
              const unsigned int occ_start{levels[l][m].first};
              const unsigned int occ_stop{levels[l][m].second};
              if ((amum.position0() >= occ_start &&
                   amum.position0() < occ_stop) ||
                  (m_stop > occ_start && m_stop <= occ_stop) ||
                  (occ_start >= amum.position0() && occ_start < m_stop) ||
                  (occ_stop > amum.position0() && occ_stop <= m_stop)) {
                occupied = true;
              }
            }
            if (occupied) {
              ++unoccupied_level;
            } else {
              break;
            }
          }
          if (unoccupied_level == levels.size()) {
            levels.resize(levels.size() + 1);
          }
          levels[unoccupied_level].emplace_back(amum.position0(), m_stop);
          const double y_level{y_main_level +
                2.0 * mum_width * (anchor2 ? -1 : 1) * unoccupied_level};
          string color{"bcolor"};
          const bool mum2_on_bad_side{mum[1].offset() > mum[0].offset()};
          const bool mum2_colored{
            (mum2_on_bad_side &&
             ((amum.read_2() == mum[1].read_2() &&
               amum.offset() >= mum[1].offset()) ||
              amum.read_2() != mum[1].read_2())) ||
                (!mum2_on_bad_side && amum.read_2() == mum[1].read_2() &&
                 amum.offset() <= mum[1].offset())};
          const bool mum1_colored{
            (!mum2_on_bad_side &&
             ((amum.read_2() == mum[0].read_2() &&
               amum.offset() >= mum[0].offset()) ||
              amum.read_2() != mum[0].read_2())) ||
                (mum2_on_bad_side && amum.read_2() == mum[0].read_2() &&
                 amum.offset() <= mum[0].offset())};
          if (mum1_colored && mum2_colored) {
            throw Error("dual coloring");
          }

          if (mum1_colored) {
            color = "color1";
          } else if (mum2_colored) {
            color = "color2";
          }

          if (amum.read_2() == mum[anchor2].read_2()) {
            if (amum == mum[0] || amum == mum[1]) {
              main_page << "[]";
            } else {
              main_page << "[" << 6 * mum_width << " " << 2 * mum_width << "]";
            }
          } else {
            main_page << "[" << 2 * mum_width << " " << 2 * mum_width << "]";
          }
          main_page << " 0 setdash " << color << " newpath "
                    << convert(amum.position0(), ref_start, ref_stop, false)
                    << " " << y_level << " moveto "
                    << convert(amum.position0() + amum.length(),
                               ref_start, ref_stop, false) << " "
                    << y_level << " lineto stroke" << endl;
        }
      }

      //
      // Base counts pages
      //

      // Loop over parents (actually all family members) for counts
      for (const Sample parent : pop.samples(family)) {
        const string parent_name{pop.sample(parent)};

        // Load the mumdex for the sample
        const MUMdex parent_mumdex{samples_dir + "/" + parent_name + "/mumdex"};

        // Counts of bases seen over region
        vector<vector<vector<unsigned int> > > base_counts(
            2, vector<vector<unsigned int> >(sequence.size(),
                                             vector<unsigned int>(5)));
        vector<vector<unsigned int>> total_counts(
            2, vector<unsigned int>(sequence.size()));

        // Loop over anchors
        const unsigned int max_read_length{152};
        for (const bool anchor2 : {false, true}) {
          const int aref_pos{anchor2 ? ref2_pos : ref1_pos};
          const bool aflip{anchor2 ? flip2 : flip1};

          // Loop over mums covering anchor reference region
          const MUMdex::MUMRegion mums_in_region{parent_mumdex.region(
              mum[anchor2].chromosome(), aref_pos - max_read_length,
              mum[anchor2].chromosome(),
              static_cast<unsigned int>(aref_pos + sequence.length()))};
          for (const MUMindex index : mums_in_region) {
            const uint64_t pair_index{index.pair_index()};
            const Pair ppair{parent_mumdex.pair(index)};
            const MUM pmum{parent_mumdex.mum(index)};
            if (pmum.length() < 25) continue;
            const std::array<std::string, 2> parent_sequences(
                parent_mumdex.sequences(pair_index));
            string parent_sequence{parent_sequences[pmum.read_2()]};
            bool flip_parent{pmum.flipped()};
            if (flip_parent) {
              reverse_complement(&parent_sequence);
            }
            const int read_start{
              pmum.read_position0(ppair.length(pmum.read_2()))};
            for (int b{0}; b != static_cast<int>(parent_sequence.size()); ++b) {
              const int ref_pos{read_start + b};
              const int ref_index{ref_pos - aref_pos};
              if (ref_index >= 0 &&
                  ref_index < static_cast<int>(parent_sequence.size())) {
                ++total_counts[anchor2][ref_index];
                unsigned int base_index{base_value(parent_sequence[b])};
                if (aflip) {
                  switch (base_index) {
                    case 0:
                      base_index = 3;
                      break;
                    case 1:
                      base_index = 2;
                      break;
                    case 2:
                      base_index = 1;
                      break;
                    case 3:
                      base_index = 0;
                      break;
                    default:
                      base_index = 4;
                      break;
                  }
                }
                ++base_counts[anchor2][ref_index][base_index];
              }
            }
          }
          if (aflip) {
            reverse(base_counts[anchor2].begin(), base_counts[anchor2].end());
            reverse(total_counts[anchor2].begin(), total_counts[anchor2].end());
          }
        }

        // Interlaced view (base count page)
        page << "% counts page" << endl;
        page << header;
        interlaced_counts(
            page, pop.member(parent) + " " + chrs[0] + " " + to_string(
                *max_element(total_counts[0].begin(), total_counts[0].end())) +
            " / " + chrs[1] + " " + to_string(
                *max_element(total_counts[1].begin(), total_counts[1].end())),
            sequence, mum1_start, mum1_stop, mum2_start, mum2_stop,
            ref1, ref2, base_counts, total_counts);
        sample_pages.back().second.push_back(page.str());
        page.str("");
      }
    }
  }

  // Output postscript
  ostringstream out_name;
  out_name << family_name << '.' << chrs[0] << '.' << pos[0] << '.'
           << (high[0] ? "high" : "low") << '.'
           << chrs[1] << '.' << pos[1] << '.'
           << (high[1] ? "high" : "low") << '.'
           << invariant << "inv." << offset << "off." << extra << ".ps";
  ofstream ps{out_name.str().c_str()};

  // Postscript header
  ps << R"foo(%!PS-Adobe-3.0
%%BoundingBox: 0 0 )foo" << page_width << " " << page_height
     << R"foo(
%%Pages: (atend)
%%Title: )foo"
     << family_name << " " << member << " "
     << chrs[0] << " " << pos[0] << " " << (high[0] ? "high" : "low") << " "
     << chrs[1] << " " << pos[1] << " " << (high[1] ? "high" : "low") << " "
     << invariant << " " << offset << " " << extra
     << R"foo(
%%Creator: Peter Andrews
%%CreationDate: Today
%%EndComments
%%BeginProlog
/gray{ 0.8 0.8 0.8 setrgbcolor } def
/color1{ 0.7 0 0 setrgbcolor } def
/color2{ 0 0 0.7 setrgbcolor } def
/color3{ 0 0 0 setrgbcolor } def
/color4{ 0 0.7 0 setrgbcolor } def
/bcolor{ 0 0 0 setrgbcolor } def
/acolor{ 0.39 0.91 0.25 setrgbcolor } def
/ccolor{ 1 0.70 0.25 setrgbcolor } def
/gcolor{ 0.92 0.25 0.235 setrgbcolor } def
/tcolor{ 0.235 0.53 0.93 setrgbcolor } def
%%EndProlog
)foo";

  sample_pages.emplace_back(0.0, vector<string>{main_page.str()});

  // Output pages in order
  // most centered page first, then left to right
  sort(sample_pages.begin(), sample_pages.end(),
       [](const pair<double, vector<string>> & lhs,
          const pair<double, vector<string>> & rhs) {
         return fabs(lhs.first) < fabs(rhs.first);
       });
  if (sample_pages.size() > 2) {
    sort(sample_pages.begin() + 2, sample_pages.end(),
         [](const pair<double, vector<string>> & lhs,
            const pair<double, vector<string>> & rhs) {
           return lhs.first < rhs.first;
         });
  }

  const bool best_page_only{false};
  const bool best_bridge_only{true};
  const bool main_pages_only{false};

  unsigned int n_pages_pre{0};
  unsigned int n_pages{0};
  for (const pair<double, vector<string> > & sample : sample_pages) {
    for (const string & spage : sample.second) {
      ++n_pages_pre;
      if (((n_pages_pre % 3) != 1) && main_pages_only) continue;
      ++n_pages;
      ps << endl << "%%Page: " << n_pages << " " << n_pages << endl
         << spage << "showpage" << endl << "grestore" << endl
         << "%%EndPage: " << n_pages << endl;
      if (best_page_only) break;
    }
    if (best_page_only) break;
    if (best_bridge_only && n_pages > 1) break;
  }

  ps << "%%Trailer" << endl;
  ps << "%%Pages: " << n_pages << endl;
  ps << "%%EOF" << endl;
  ps.close();

  // Convert to pdf
  ostringstream ps2pdf;
  string pdf_name{out_name.str()};
  replace_substring(pdf_name, ".ps", ".pdf");
  ps2pdf << "ps2pdf -dDEVICEWIDTHPOINTS=" << page_width
         << " -dDEVICEHEIGHTPOINTS=" << page_height
         << " " << out_name.str() << " "
         << pdf_name;
  if (system(ps2pdf.str().c_str()) == -1) {
    cerr << "Problem creating pdf file" << endl;
  }

  return 0;
} catch (Error & e) {
  cerr << "paa::Error:" << endl;
  cerr << e.what() << endl;
  return 1;
} catch (exception & e) {
  cerr << "std::exception" << endl;
  cerr << e.what() << endl;
  return 1;
} catch (...) {
  cerr << "unknown exception was caught" << endl;
  return 1;
}
```
