## Supplementary material for "G-Graph: An interactive genomic graph viewer": MUMdex Source Code for G-Graph: bridges.cpp

```
//
// bridges
//
// extract bridge information from a mumdex
//
// Copyright 2016 Peter Andrews @ CSHL
//

#include "bridges.h"

#include <algorithm>
#include <exception>
#include <iostream>
#include <future>
#include <mutex>
#include <queue>
#include <sstream>
#include <string>
#include <thread>
#include <utility>
#include <vector>

#include "error.h"
#include "files.h"
#include "mumdex.h"
#include "threads.h"

using std::async;
using std::cerr;
using std::cout;
using std::endl;
using std::exception;
using std::future;
using std::lower_bound;
using std::make_unique;
using std::move;
using std::ostringstream;
using std::priority_queue;
using std::sort;
using std::string;
using std::unique_ptr;
using std::upper_bound;
using std::vector;

using paa::bwritec;
using paa::ref_ptr;
using paa::unlink;
using paa::BridgeInfo;
using paa::Error;
using paa::MappedVector;
using paa::MUMdex;
using paa::OneBridgeInfo;
using paa::Progress;
using paa::Reference;
using paa::ThreadPool;

const Reference * paa::ref_ptr;

std::mutex cerr_mutex;
std::mutex thread_mutex;
uint64_t n_threads_running{0};
bool do_progress{false};

vector<string> bridges(const MUMdex & mumdex,
                       const uint64_t pair_start,
                       const uint64_t pair_stop,
                       const unsigned int thread_id,
                       const uint64_t max_output,
                       Progress & progress) {
  {
    std::lock_guard<std::mutex> thread_lock(thread_mutex);
    ++n_threads_running;
  }
  vector<string> file_names;
  vector<OneBridgeInfo> output;
  output.reserve(max_output + 10000);
  unsigned int output_index{0};
  uint64_t last_start{pair_start};
  for (uint64_t pair_index{pair_start}; pair_index != pair_stop;
       ++pair_index) {
    pair_bridges(mumdex, pair_index, output);
    if (output.size() &&
        (output.size() > max_output || pair_index + 1 == pair_stop)) {
      sort(output.begin(), output.end());
      ostringstream file_name;
      file_name << "bridges." << thread_id << "." << output_index++ << ".bin";
      const string name{file_name.str()};
      bwritec(file_name.str(), &output[0], "OneBridgeInfo vector",
              sizeof(OneBridgeInfo) * output.size());
      file_names.push_back(name);
      std::lock_guard<std::mutex> cerr_lock(cerr_mutex);
      ostringstream message;
      if (pair_index + 1 == pair_stop) {
        std::lock_guard<std::mutex> thread_lock(thread_mutex);
        --n_threads_running;
      }
      if (n_threads_running) {
        message << n_threads_running << " threads";
      }
      if (do_progress) progress(pair_index - last_start, message.str());
      last_start = pair_index;
      output.clear();
    }
  }
  return file_names;
}

class BridgeComparer {
 public:
  bool operator()(const unsigned int chromosome, const OneBridgeInfo & info) {
    return chromosome < info.chr1();
  }
  bool operator()(const OneBridgeInfo & info, const unsigned int chromosome) {
    return info.chr1() < chromosome;
  }
};

std::mutex cout_mutex;
class MergeHelper {
 public:
  explicit MergeHelper(const MappedVector<OneBridgeInfo> & file_,
                       const unsigned int chromosome) :
      file{&file_} {
    const MappedVector<OneBridgeInfo>::const_iterator lower{
      lower_bound(file->begin(), file->end(), chromosome, BridgeComparer())};
    const MappedVector<OneBridgeInfo>::const_iterator upper{
      upper_bound(lower, file->end(), chromosome, BridgeComparer())};
    n = lower - file->begin();
    stop = upper - file->begin();
    if (0) {
      std::lock_guard<std::mutex> cout_lock(cout_mutex);
      cout << chromosome << " " << n << " " << stop << endl;
    }
  }
  bool advance() {
    if (++n == stop) {
      return false;
    } else {
      return true;
    }
  }
  bool operator<(const MergeHelper & other) const {
    return other.current() < current();
  }
  const OneBridgeInfo current() const {
    return (*file)[n];
  }
  uint64_t size() const {
    return stop - n;
  }

 private:
  uint64_t n{0};
  uint64_t stop{0};
  const MappedVector<OneBridgeInfo> * file{nullptr};
};

int main(int argc, char* argv[])  try {
  if (0) {
    cerr << paa::max_support_save_length
         << " " << sizeof(OneBridgeInfo)
         << " " << sizeof(BridgeInfo)
         << " " << NEW_BRIDGE_FORMAT << endl;
  }

  if (argc != 3 && argc != 4) {
    throw Error("usage: bridges mumdex n_threads [progress]");
  }

  if (argc == 4) {
    do_progress = true;
  }

  const uint64_t billion{1000000000UL};
  const uint64_t max_bytes{10 * billion};
  const unsigned int n_threads{static_cast<unsigned int>(atoi(argv[2]))};

  vector<string> filenames;
  vector<string> chromosomes;
  string fasta_name;
  {
    const MUMdex mumdex{argv[1]};
    fasta_name = mumdex.reference().fasta_file();
    ref_ptr = &mumdex.reference();
    for (unsigned int chromosome{0}; chromosome != ref_ptr->n_chromosomes();
         ++chromosome) {
      chromosomes.push_back(ref_ptr->name(chromosome));
    }
    const uint64_t max_output{max_bytes / sizeof(OneBridgeInfo) / n_threads};
    const uint64_t n_pairs{mumdex.n_pairs()};
    const uint64_t pairs_per_thread{n_pairs / n_threads};

    cerr << "Finding bridges in " << n_pairs << " pairs and "
         << mumdex.n_mums() << " mums" << endl;

    // launch bridge info threads
    vector<future<vector<string>>> bridge_futures;
    Progress progress(n_pairs, 0.01, "Bridge Loop");
    for (unsigned int thread{0}; thread != n_threads; ++thread) {
      const uint64_t start_pair{pairs_per_thread * thread};
      const uint64_t stop_pair{thread + 1 == n_threads ?
            n_pairs : pairs_per_thread * (thread + 1)};
      bridge_futures.push_back(async(
          std::launch::async,
          bridges, std::ref(mumdex),
          start_pair, stop_pair, thread, max_output, std::ref(progress)));
    }

    // collect bridge info filenames from threads
    for (future<vector<string>> & future : bridge_futures) {
      const vector<string> thread_filenames{future.get()};
      filenames.insert(filenames.end(),
                       thread_filenames.begin(), thread_filenames.end());
    }
  }
  const Reference ref{fasta_name};
  ref_ptr = &ref;

  const uint64_t max_merged{max_bytes / sizeof(BridgeInfo) /
        chromosomes.size() / 2};
  vector<MappedVector<OneBridgeInfo>> input_files;
  input_files.reserve(filenames.size());
  uint64_t n_to_merge{0};
  for (const string & name : filenames) {
    input_files.emplace_back(name);
    n_to_merge += input_files.back().size();
  }

  cerr << "merging " << n_to_merge << " bridges from "
       << filenames.size() << " files" << endl;

  vector<future<uint64_t>> mergers;
  mergers.reserve(chromosomes.size());
  ThreadPool pool{n_threads};
  for (unsigned int chromosome{0}; chromosome != chromosomes.size();
       ++chromosome) {
    mergers.emplace_back(pool.run(
        [&input_files, &ref, chromosome, max_merged]() {
          vector<BridgeInfo> merged;
          vector<BridgeInfo> to_save;
          if (1) {
            merged.reserve(max_merged);
            to_save.reserve(max_merged);
          }
          future<void> saver;
          priority_queue<MergeHelper> queue;
          uint64_t n_to_merge_chr{0};
          for (unsigned int file{0}; file != input_files.size(); ++file) {
            MergeHelper helper{input_files[file], chromosome};
            if (helper.size()) {
              queue.push(move(helper));
              n_to_merge_chr += helper.size();
            }
          }

          // merge bridge info
          const string bridge_file_name{get_bridge_file_name(ref, chromosome)};
          FILE * output_file{fopen(bridge_file_name.c_str(), "wb")};
          if (output_file == nullptr) {
            throw Error("Could not open output file")
                << bridge_file_name << paa::bridges_bad_message();
          }
          if (queue.empty()) {
            fclose(output_file);
            return static_cast<uint64_t>(0);
          }
          uint64_t n_saved{0};

          Progress progress(n_to_merge_chr, 0.01, "Merge Loop");
          while (true) {
            if (do_progress && chromosome == 0) progress();
            MergeHelper lowest{queue.top()};
            const OneBridgeInfo current{lowest.current()};
            queue.pop();
            if (lowest.advance()) {
              queue.push(lowest);
            }
            const bool empty{queue.empty()};

            if (merged.empty() || merged.back() < current) {
              merged.emplace_back(current);
            } else {
              merged.back().combine(current);
            }
            if (merged.size() == max_merged || empty) {
              if (saver.valid()) {
                saver.get();
              }
              const BridgeInfo last_entry{merged.back()};
              if (!empty) merged.pop_back();
              n_saved += merged.size();
              to_save.swap(merged);
              merged.clear();
              if (!empty) merged.push_back(last_entry);
              saver = async(
                  std::launch::async,
                  [output_file] (const vector<BridgeInfo> & tosave) {
                    bwritec(output_file, &tosave[0], "bridges",
                            tosave.size() * sizeof(BridgeInfo));
                  }, std::ref(to_save));
              if (empty) {
                break;
              }
            }
          }
          if (saver.valid()) {
            saver.get();
          }
          fclose(output_file);
          return n_saved;
        }));
  }

  uint64_t n_saved{0};
  for (unsigned int chromosome{0}; chromosome != chromosomes.size();
       ++chromosome) {
    n_saved += mergers[chromosome].get();
  }
  cerr << "saved " << n_saved << " bridges" << endl;
  for (const string & file : filenames) {
    unlink(file);
  }

  return 0;
} catch (Error & e) {
  cerr << "paa::Error:" << endl;
  cerr << e.what() << endl;
  return 1;
} catch (exception & e) {
  cerr << "std::exception" << endl;
  cerr << e.what() << endl;
  return 1;
} catch (...) {
  cerr << "unknown exception was caught" << endl;
  return 1;
}
```
