## Supplementary material for "G-Graph: An interactive genomic graph viewer": MUMdex Source Code for G-Graph: bridges.h

```
//
// bridges.h
//
// bridge information format and extraction
//
// Copyright 2015 Peter Andrews @ CSHL
//

#ifndef PAA_BRIDGES_H
#define PAA_BRIDGES_H

#include <algorithm>
#include <limits>
#include <map>
#include <mutex>
#include <numeric>
#include <set>
#include <string>
#include <utility>
#include <vector>

#include "error.h"
#include "mumdex.h"
#include "population.h"
#include "utility.h"

namespace paa {

#ifndef NEW_BRIDGE_FORMAT
#define NEW_BRIDGE_FORMAT 1
#endif

std::string bridges_bad_message() {
  return R"xxx(
Perhaps this error was caused by a bridges file version incomatibility

To select old version of bridges structure for processing old chrbridges files:

make clean && make SPECIAL=-DNEW_BRIDGE_FORMAT=0

The new version is the default and works for up to 256 chromosomes
   and the file output names start with newbridges

An error is also thrown if you use the old version with too many chromosomes

All programs are written to never confuse the two versions
  since the output file names are different

If you encounter an error using old files, swich to old version using
  the modified make command above
)xxx";
}

std::string get_bridge_file_name(const Reference & ref,
                                 const unsigned int chromosome) {
  std::ostringstream result;
  if (NEW_BRIDGE_FORMAT) {
    result << "newbridges." << ref.name(chromosome) << ".bin";
  } else {
    if (ref.n_chromosomes() > 128)
      throw Error("Bad reference size for bridges") << bridges_bad_message();
    result << "chrbridges." << chromosome << ".bin";
  }
  return result.str();
}

// run parameters
constexpr bool exclude_snps{false};
constexpr unsigned int min_support_length{20};

// fit in more data for shorter reads and fewer chromosomes than mumdex allows
constexpr unsigned int fewer_chromosome_bits{7};
constexpr unsigned int fewer_position_bits{31};
constexpr unsigned int short_read_bits{9};
constexpr unsigned int max_support_save_length{(1 << short_read_bits) - 1};
constexpr bool longer_reads_than_fit_in_short_read_bits{false};

constexpr bool test_invariant{false};

extern const Reference * ref_ptr;

class OneBridgeInfo {
 public:
  OneBridgeInfo() :
      pos1_{0}, pos2_{0},
    high1_{0}, high2_{0},
    chr1_{0}, chr2_{0},
    padding_{0}, offset_{0},
    anchor1_length_{0}, anchor2_length_{0},
    mate_anchor1_length_{0}, mate_anchor2_length_{0} { }

  OneBridgeInfo(const unsigned int chr1__,
                const unsigned int pos1__,
                const unsigned int high1__,
                const unsigned int chr2__,
                const unsigned int pos2__,
                const unsigned int high2__,
                const int offset__) :
      pos1_{pos1__}, pos2_{pos2__},
    high1_{high1__}, high2_{high2__},
    chr1_{chr1__}, chr2_{chr2__},
    padding_{0}, offset_{offset__},
    anchor1_length_{0}, anchor2_length_{0},
    mate_anchor1_length_{0}, mate_anchor2_length_{0} { }

  OneBridgeInfo(const MUM mum1,
                const bool mum1_is_high,
                const MUM mum2,
                const bool mum2_is_high,
                unsigned int mum1_support = 0,
                unsigned int mum2_support = 0,
                unsigned int mate_mum1_support = 0,
                unsigned int mate_mum2_support = 0) :
      OneBridgeInfo{} {
    const unsigned int mum1_pos{mum1.anchor_position(mum1_is_high)};
    const unsigned int mum2_pos{mum2.anchor_position(mum2_is_high)};

    const bool keep_order{(mum1.chromosome() == mum2.chromosome() ?
                           (mum1_pos == mum2_pos ?
                            mum1_is_high > mum2_is_high :
                            mum1_pos < mum2_pos) :
                            mum1.chromosome() > mum2.chromosome())};

    chr1_ = keep_order ? mum1.chromosome() : mum2.chromosome();
    chr2_ = keep_order ? mum2.chromosome() : mum1.chromosome();

    pos1_ = keep_order ? mum1_pos : mum2_pos;
    pos2_ = keep_order ? mum2_pos : mum1_pos;

    high1_ = keep_order ? mum1_is_high : mum2_is_high;
    high2_ = keep_order ? mum2_is_high : mum1_is_high;

    if (longer_reads_than_fit_in_short_read_bits) {
      if (mum1_support > max_support_save_length) {
        mum1_support = max_support_save_length;
      }
      if (mum2_support > max_support_save_length) {
        mum2_support = max_support_save_length;
      }
      if (mate_mum1_support > max_support_save_length) {
        mate_mum1_support = max_support_save_length;
      }
      if (mate_mum2_support > max_support_save_length) {
        mate_mum2_support = max_support_save_length;
      }
    }

    anchor1_length_ = keep_order ? mum1_support : mum2_support;
    anchor2_length_ = keep_order ? mum2_support : mum1_support;

    mate_anchor1_length_ = keep_order ? mate_mum1_support : mate_mum2_support;
    mate_anchor2_length_ = keep_order ? mate_mum2_support : mate_mum1_support;

    const unsigned int mum1_base{mum1.offset() +
          (mum1_is_high != mum1.flipped() ? mum1.length() - 1 : 0)};
    const unsigned int mum2_base{mum2.offset() +
          (mum2_is_high != mum2.flipped() ? mum2.length() - 1 : 0)};
    offset_ = static_cast<int>(mum2_base) - static_cast<int>(mum1_base);
    padding_ = 0;

    if (test_invariant) {
      const int normal_invariant{
        ((mum1.flipped() ? -1 : 1) * mum1.read_start_position() -
         (mum2.flipped() ? -1 : 1) * mum2.read_start_position())};
      if (invariant() != normal_invariant) {
        // throw Error("Bad invariant calculation");
        sout << high1() << high2() << mum1_base << mum2_base
             << invariant() - normal_invariant << offset()
             << invariant() << std::endl;
      }
    }
  }

  unsigned int chr1() const { return chr1_; }
  unsigned int chr2() const { return chr2_; }
  unsigned int pos1() const { return pos1_; }
  unsigned int pos2() const { return pos2_; }
  bool high1() const { return high1_; }
  bool high2() const { return high2_; }

  int16_t offset() const { return offset_; }
  int64_t invariant() const {
    return (high1_ ? 1 : -1) * static_cast<int64_t>(pos1_) +
        (high2_ ? 1 : -1) * static_cast<int64_t>(pos2_) + offset_;
  }
  char orientation_char() const {
    if (high1_ != high2_) {
      return '=';
    } else {
      if (high1_) {
        return 'i';
      } else {
        return 'o';
      }
    }
  }

  unsigned int anchor1_length() const { return anchor1_length_; }
  unsigned int anchor2_length() const { return anchor2_length_; }
  unsigned int mate_anchor1_length() const { return mate_anchor1_length_; }
  unsigned int mate_anchor2_length() const { return mate_anchor2_length_; }

  void combine(const OneBridgeInfo & other) {
    if (anchor1_length_ < other.anchor1_length_) {
      anchor1_length_ = other.anchor1_length_;
    }
    if (anchor2_length_ < other.anchor2_length_) {
      anchor2_length_ = other.anchor2_length_;
    }
    if (mate_anchor1_length_ < other.mate_anchor1_length_) {
      mate_anchor1_length_ = other.mate_anchor1_length_;
    }
    if (mate_anchor2_length_ < other.mate_anchor2_length_) {
      mate_anchor2_length_ = other.mate_anchor2_length_;
    }
  }

  template <class BRIDGE>
  bool operator<(const BRIDGE & other) const {
    if (chr1_ == other.chr1()) {
      if (pos1_ == other.pos1()) {
        if (high1_ == other.high1()) {
          if (chr2_ == other.chr2()) {
            if (pos2_ == other.pos2()) {
              if (high2_ == other.high2()) {
                return offset_ < other.offset();
              } else {
                return high2_ < other.high2();
              }
            } else {
              return pos2_ < other.pos2();
            }
          } else {
            return chr2_ < other.chr2();
          }
        } else {
          return high1_ < other.high1();
        }
      } else {
        return pos1_ < other.pos1();
      }
    } else {
      return chr1_ < other.chr1();
    }
  }

  template <class STREAM>
  void output(STREAM & stream) const {
    stream << ref_ptr->name(chr1())
           << pos1()
           << high1()
           << ref_ptr->name(chr2())
           << pos2()
           << high2()
           << orientation_char()
           << invariant()
           << offset()
           << anchor1_length()
           << anchor2_length()
           << mate_anchor1_length()
           << mate_anchor2_length();
  }

  std::string description() const {
    const unsigned int hmega{100000000};
    const unsigned int tmega{10000000};
    const unsigned int mega{1000000};
    const unsigned int huge{100000};
    const unsigned int large{10000};
    const unsigned int kilo{1000};
    const unsigned int small{100};
    const unsigned int tiny{10};

    if (chr2_ != chr1_) {
      return "translocation";
    }

    const int64_t sinvariant{invariant()};
    const uint64_t uinvariant{int64_abs(sinvariant)};
    const int64_t sposdiff{(static_cast<int32_t>(pos1_) -
                            static_cast<int32_t>(pos2_))};
    const uint64_t uposdiff{int64_abs(sposdiff)};
    const uint64_t size{((uinvariant == 0 || high1_ == high2_) ?
                         uposdiff : uinvariant)};

    std::string distance;
    if (size <= 1) {
      distance = "single";
    } else if (size <= tiny) {
      distance = "1s";
    } else if (size <= small) {
      distance = "10s";
    } else if (size <= kilo) {
      distance = "100s";
    } else if (size <= large) {
      distance = "1Ks";
    } else if (size <= huge) {
      distance = "10Ks";
    } else if (size <= mega) {
      distance = "100Ks";
    } else if (size <= tmega) {
      distance = "1Ms";
    } else if (size <= hmega) {
      distance = "10Ms";
    } else {
      distance = "100Ms";
    }

    std::string bridge_type;
    if (high1() != high2()) {
      if (sinvariant == 0) {
        if (size <= 2) {
          distance = "single";
        }
        bridge_type = "substitution";
      } else if (sinvariant > 0) {
        bridge_type = "insertion";
      } else {
        bridge_type = "deletion";
      }
    } else if (high1()) {
      bridge_type = "inversion";
    } else {
      bridge_type = "outversion";
    }
    return distance + "_" + bridge_type;
  }

 private:
#if NEW_BRIDGE_FORMAT
  uint64_t pos1_: fewer_position_bits;             // 31 31
  uint64_t pos2_: fewer_position_bits;             // 31 62
  uint64_t high1_: 1;                              // 1  63
  uint64_t high2_: 1;                              // 1  64
  uint64_t chr1_: chromosome_bits;                 // 8  72
  uint64_t chr2_: chromosome_bits;                 // 8  80
  uint64_t padding_: 1;                            // 1  81
  int64_t offset_: read_bits + 1;                  // 11 92
#else
  uint32_t pos1_: position_bits;                   // 32 32
  uint32_t pos2_: position_bits;                   // 32 64
  uint64_t high1_: 1;                              // 1  65
  uint64_t high2_: 1;                              // 1  66
  uint64_t chr1_: fewer_chromosome_bits;           // 7  73
  uint64_t chr2_: fewer_chromosome_bits;           // 7  80
  uint64_t padding_: 1;                            // 1  81
  int64_t offset_: read_bits + 1;                  // 11 92
#endif

 protected:
  uint64_t anchor1_length_: short_read_bits;       // 9  101
  uint64_t anchor2_length_: short_read_bits;       // 9  110
  uint64_t mate_anchor1_length_: short_read_bits;  // 9  119
  uint64_t mate_anchor2_length_: short_read_bits;  // 9  128
};

template <class STREAM>
STREAM & operator<<(STREAM & out, const OneBridgeInfo & bridge) {
  bridge.output(out);
  return out;
}

class BridgeInfo : public OneBridgeInfo {
 public:
  BridgeInfo() : OneBridgeInfo{} { }
  explicit BridgeInfo(const OneBridgeInfo info) :
      OneBridgeInfo{info} {
    if (mate_anchor1_length_) ++mate_anchor1_count_;
    if (mate_anchor2_length_) ++mate_anchor2_count_;
    if (false) padding_ = 0;  // Avoid unused variable warning
  }

  uint16_t bridge_count() const { return bridge_count_; }
  uint16_t mate_anchor1_count() const { return mate_anchor1_count_; }
  uint16_t mate_anchor2_count() const { return mate_anchor2_count_; }

  void combine(const OneBridgeInfo & info) {
    ++bridge_count_;
    if (anchor1_length() < info.anchor1_length()) {
      anchor1_length_ = info.anchor1_length();
    }
    if (anchor2_length() < info.anchor2_length()) {
      anchor2_length_ = info.anchor2_length();
    }
    if (info.mate_anchor1_length()) {
      ++mate_anchor1_count_;
      if (mate_anchor1_length() < info.mate_anchor1_length()) {
        mate_anchor1_length_ = info.mate_anchor1_length();
      }
    }
    if (info.mate_anchor2_length()) {
      ++mate_anchor2_count_;
      if (mate_anchor2_length() < info.mate_anchor2_length()) {
        mate_anchor2_length_ = info.mate_anchor2_length();
      }
    }
  }
  void combine(const BridgeInfo & info) {
    bridge_count_ += info.bridge_count_;
    if (anchor1_length() < info.anchor1_length()) {
      anchor1_length_ = info.anchor1_length();
    }
    if (anchor2_length() < info.anchor2_length()) {
      anchor2_length_ = info.anchor2_length();
    }
    if (info.mate_anchor1_length()) {
      ++mate_anchor1_count_;
      if (mate_anchor1_length() < info.mate_anchor1_length()) {
        mate_anchor1_length_ = info.mate_anchor1_length();
      }
    }
    if (info.mate_anchor2_length()) {
      ++mate_anchor2_count_;
      if (mate_anchor2_length() < info.mate_anchor2_length()) {
        mate_anchor2_length_ = info.mate_anchor2_length();
      }
    }
  }
  void multiply(const uint64_t count) {
    bridge_count_ *= count;
  }

  template <class STREAM>
  void output(STREAM & stream) const {
    OneBridgeInfo::output(stream);
    stream << bridge_count()
           << mate_anchor1_count()
           << mate_anchor2_count();
  }

 private:
  uint16_noo_t bridge_count_{1};
  uint16_noo_t mate_anchor1_count_{0};
  uint16_noo_t mate_anchor2_count_{0};
  uint16_t padding_{0};
};

template <class STREAM>
STREAM & operator<<(STREAM & out, const BridgeInfo & bridge) {
  bridge.output(out);
  return out;
}

class MergeHelper {
 public:
  explicit MergeHelper(const std::string & file_name_,
                       const Sample sample_arg,
                       const unsigned int start_,
                       const unsigned int stop_,
                       const bool initial_advance = true) :
      file_name{file_name_}, sample_{sample_arg},
    start{start_}, stop{stop_} {
      if ((file = fopen(file_name.c_str(), "rb")) == nullptr) {
        sleep(2);
        if ((file = fopen(file_name.c_str(), "rb")) == nullptr) {
          sleep(5);
          if ((file = fopen(file_name.c_str(), "rb")) == nullptr) {
            throw Error("Could not open bridges file")
                << file_name << bridges_bad_message();
          }
        }
      }

      // find starting position
      if (start != 0) {
        const FileVector<BridgeInfo> mapped{file_name};
        const FileVector<BridgeInfo>::const_iterator found{
          std::lower_bound(mapped.begin(), mapped.end(), start,
                           [](const BridgeInfo & bridge,
                              const unsigned int val) {
                             return bridge.pos1() < val;
                           })};
        if (found == mapped.end()) {
          fclose(file);
          file = nullptr;
          return;
        }
        if (fseek(file, (found - mapped.begin()) * sizeof(BridgeInfo),
                  SEEK_SET)) {
          throw Error("Problem seeking in file") << file_name;
        }
      }
      if (!initial_advance || advance()) {
        valid_start_ = true;
      }
    }

  MergeHelper(const MergeHelper &) = delete;
  MergeHelper & operator=(const MergeHelper &) = delete;
  MergeHelper(MergeHelper && rhs) :
      file_name{rhs.file_name}, file{rhs.file},
    sample_{rhs.sample_}, start{rhs.start}, stop{rhs.stop},
    valid_start_{rhs.valid_start_}, current_{rhs.current_} {
      rhs.file = nullptr;
    }

  ~MergeHelper() {
    if (file) {
      fclose(file);
      file = nullptr;
    }
  }

  uint64_t read_block(const uint64_t n_desired, BridgeInfo * data) {
    if (!file || !n_desired) return 0;
    uint64_t n_read{fread(data, sizeof(BridgeInfo), n_desired, file)};
    while (n_read && (data + n_read - 1)->pos1() >= stop) --n_read;
    if (n_read < n_desired) {
      fclose(file);
      file = nullptr;
    }
    return n_read;
  }

  bool advance() {
    if (fread(&current_, sizeof(BridgeInfo), 1, file) == 1) {
      if (current_.pos1() >= stop) return false;
      return true;
    } else {
      fclose(file);
      file = nullptr;
      return false;
    }
  }
  // lower bridges first in priority queue
  // if equal bridges, lower sample ids first
  bool operator<(const MergeHelper & other) const {
    if (other.current() < current()) {
      return true;
    } else if (current() < other.current()) {
      return false;
    } else {
      return other.sample() < sample();
    }
  }
  const BridgeInfo current() const {
    return current_;
  }
  Sample sample() const {
    return sample_;
  }
  bool valid_start() const {
    return valid_start_;
  }

 private:
  std::string file_name;
  FILE * file{nullptr};
  Sample sample_;
  unsigned int start;
  unsigned int stop;
  bool valid_start_{false};
  BridgeInfo current_{};
};

struct MergeHelperCompare {
  bool operator()(const MergeHelper * lhs, const MergeHelper * rhs) {
    return *lhs < *rhs;
  }
};

template <class MUMDEX>
class MUMsupport {
 public:
  MUMsupport(const MUMDEX & mumdex, const Pair pair,
             const uint64_t main_mum_index,
             const uint64_t mums_start,
             const uint64_t read_2_start,
             const uint64_t mums_stop) {
    const MUM main_mum{mumdex.mum(main_mum_index)};
    const unsigned int read_length{pair.length(main_mum.read_2())};
    const int main_mum_read_pos{main_mum.read_position0(read_length)};
    unsigned int min_seen_pos{main_mum.position0()};
    unsigned int max_seen_pos{main_mum.position0() + main_mum.length() - 1};

    // get support on read
    for (const bool high : {false, true}) {
      n_bases_read[high] += main_mum.length();
    }
    const uint64_t read_start{main_mum.read_2() ? read_2_start : mums_start};
    const uint64_t read_stop{main_mum.read_2() ? mums_stop : read_2_start};
    for (uint64_t mum_index{read_start}; mum_index != read_stop; ++mum_index) {
      if (mum_index == main_mum_index) continue;
      const MUM mum{mumdex.mum(mum_index)};
      if (main_mum.chromosome() != mum.chromosome()) continue;
      if (main_mum.flipped() != mum.flipped()) continue;
      if (main_mum_read_pos != mum.read_position0(read_length)) continue;
      const bool high_side{(mum_index > main_mum_index) != main_mum.flipped()};
      n_bases_read[high_side] += mum.length();
      min_seen_pos = std::min(min_seen_pos, mum.position0());
      max_seen_pos = std::max(max_seen_pos, mum.position0() + mum.length() - 1);
    }

    // get support on mate
    const uint64_t mate_start{main_mum.read_2() ? mums_start : read_2_start};
    const uint64_t mate_stop{main_mum.read_2() ? read_2_start : mums_stop};
    const unsigned int max_mate_distance{1000};
    const bool high_side{!main_mum.flipped()};
    for (uint64_t mum_i_1{mate_stop}; mum_i_1 != mate_start; --mum_i_1) {
      const uint64_t mum_index{mum_i_1 - 1};
      const MUM mum{mumdex.mum(mum_index)};
      if (main_mum.chromosome() != mum.chromosome()) continue;
      if (main_mum.flipped() == mum.flipped()) continue;
      const unsigned int max_mate_mum_pos{mum.position0() + mum.length() - 1};
      if (high_side) {
        if (max_mate_mum_pos <= max_seen_pos ||
            mum.position0() > main_mum.position0() + max_mate_distance) {
          continue;
        }
        n_bases_mate[high_side] += max_mate_mum_pos -
            std::max(max_seen_pos + 1, mum.position0()) + 1;
      } else {
        if (mum.position0() >= min_seen_pos ||
            main_mum.position0() > mum.position0() + max_mate_distance) {
          continue;
        }
        n_bases_mate[high_side] +=
            std::min(min_seen_pos, max_mate_mum_pos + 1) - mum.position0();
      }
      min_seen_pos = std::min(min_seen_pos, mum.position0());
      max_seen_pos = std::max(max_seen_pos, max_mate_mum_pos);
    }
  }

  unsigned int n_bases_read[2]{0, 0};
  unsigned int n_bases_mate[2]{0, 0};
};

template <class MUMDEX>
void pair_bridges(const MUMDEX & mumdex,
                  const uint64_t pair_index,
                  std::vector<OneBridgeInfo> & output,
                  const bool skip_dupes = true) {
  // the pair
  const Pair pair{mumdex.pair(pair_index)};
  if (skip_dupes && pair.dupe()) return;

  // mum boundaries for pair
  const uint64_t mums_start{mumdex.mums_start(pair_index)};
  const uint64_t mums_stop{mumdex.mums_stop(pair_index)};

  // find the first mum of read 2, if any
  const uint64_t read_2_start{[mums_start, mums_stop, &mumdex] {
      for (uint64_t mum_index{mums_start}; mum_index != mums_stop;
           ++mum_index) {
        if (mumdex.mum(mum_index).read_2()) {
          return mum_index;
        }
      }
      return mums_stop;
    }()};

  // gather support info for each mum in pair
  static thread_local std::vector<MUMsupport<MUMDEX>> support;
  support.clear();
  for (uint64_t mum_index{mums_start}; mum_index != mums_stop; ++mum_index) {
    support.emplace_back(mumdex, pair, mum_index,
                         mums_start, read_2_start, mums_stop);
  }

  // collect bridges for pair
  static thread_local std::vector<OneBridgeInfo> pair_output;
  pair_output.clear();
  for (uint64_t mum1_index{mums_start}; mum1_index != mums_stop;
       ++mum1_index) {
    const MUM mum1{mumdex.mum(mum1_index)};
    if (pair.bad(mum1.read_2())) continue;

    const MUMsupport<MUMDEX> & mum1_support{support[mum1_index - mums_start]};
    if (mum1_support.n_bases_read[0] < min_support_length &&
        mum1_support.n_bases_read[1] < min_support_length) {
      continue;
    }

    // loop over mum2 mums
    for (uint64_t mum2_index{mum1_index + 1}; mum2_index != mums_stop;
         ++mum2_index) {
      const MUM mum2{mumdex.mum(mum2_index)};
      if (mum2.read_2() != mum1.read_2()) continue;

      const MUMsupport<MUMDEX> & mum2_support{support[mum2_index - mums_start]};
      if (mum2_support.n_bases_read[0] < min_support_length &&
          mum2_support.n_bases_read[1] < min_support_length) {
        continue;
      }

      // Bad logic ??? here or later - but works if mum1 < mum2 in read
      const bool mum1_is_high{(mum2_index > mum1_index) != mum1.flipped()};
      const bool mum2_is_high{(mum2_index > mum1_index) == mum2.flipped()};

      const bool mum1_side{!mum1_is_high};
      const bool mum2_side{!mum2_is_high};

      if (mum1_support.n_bases_read[mum1_side] < min_support_length ||
          mum2_support.n_bases_read[mum2_side] < min_support_length) {
        continue;
      }

      const OneBridgeInfo bridge{
        mum1, mum1_is_high,
            mum2, mum2_is_high,
            mum1_support.n_bases_read[mum1_side],
            mum2_support.n_bases_read[mum2_side],
            mum1_support.n_bases_mate[mum1_side],
            mum2_support.n_bases_mate[mum2_side]
            };
      if (!exclude_snps ||
          mum1.chromosome() != mum2.chromosome() ||
          bridge.invariant() != 0) {
        pair_output.push_back(bridge);
      }
    }
  }
  sort(pair_output.begin(), pair_output.end());
  for (unsigned int bridge_index{0}; bridge_index != pair_output.size();
       ++bridge_index) {
    const OneBridgeInfo & bridge{pair_output[bridge_index]};
    if (bridge_index == 0 || output.back() < bridge) {
      output.push_back(bridge);
    } else {
      output.back().combine(bridge);
    }
  }
}

template <class MUMdex>
class TBridges {
 public:
  using Anchors = typename MUMdex::Anchors;
  class Bridge {
   public:
    using const_iterator = typename MUMdex::Anchors::const_iterator;
    Bridge(const TBridges & bridges_, const const_iterator start) :
        bridges{bridges_}, current{start} {
          if (current != bridges.anchorsA.end() && !prepare()) {
            ++(*this);
          }
        }
    bool operator!=(const Bridge & other) const {
      return current != other.current;
    }
    Bridge & operator++() {
      while (++current != bridges.anchorsA.end()) {
        if (prepare()) {
          break;
        }
      }
      return *this;
    }
    const Bridge & operator*() const {
      return *this;
    }
    bool prepare() {
      // Pair and MUM information on anchor A
      const MUMindex pm{*current};
      pair_index_ = pm.pair_index();
      const MUMdex & mumdex_{bridges.mumdex};
      const Pair pair_{mumdex_.pair(pm)};
      const uint64_t mumA_index{pair_.mums_start() + pm.mum_in_pair_index()};
      const MUM mumA{mumdex_.mum(pm)};

      // Look for anchor B in same pair as anchor A
      for (uint64_t mumB_index{pair_.mums_start()};
           mumB_index != mumdex_.mums_stop(pair_index_) ; ++mumB_index) {
        if (mumA_index == mumB_index) continue;
        const MUM mumB{mumdex_.mum(mumB_index)};
        if (mumA.read_2() != mumB.read_2()) continue;

        // Is anchor B at correct position?
        if (mumB.chromosome() != bridges.chrB) continue;
        if (mumB.position0() + (bridges.highB ? mumB.length() - 1 : 0) !=
            bridges.posB) continue;

        // Mum orientation sanity check
        if (false) {
          if ((bridges.highA == mumA.flipped()) == (mumA_index < mumB_index)) {
            std::cerr << "unexpected highA seen" << std::endl;
            continue;
          }
          if ((bridges.highB != mumB.flipped()) == (mumA_index < mumB_index)) {
            std::cerr << "unexpected highB seen" << std::endl;
            continue;
          }
        }

        // Orient MUMs so that mumL is earlier in the read
        const bool switchMUMs{mumB_index < mumA_index};
        const MUM mumL{switchMUMs ? mumB : mumA};
        const bool highL{switchMUMs ? bridges.highB : bridges.highA};
        const MUM mumR{switchMUMs ? mumA : mumB};
        const bool highR{switchMUMs ? bridges.highA : bridges.highB};

        // Create bridge
        bridge = OneBridgeInfo{mumL, highL, mumR, highR,
                               mumL.length(), mumR.length()};

        // Check that invariant is as expected
        if (bridge.invariant() != bridges.invariant) {
          if (false) {
            sout << "found different bridge invariant"
                 << bridge.invariant() << std::endl;
          }
          continue;
        }

        // Orient MUMs to canonical ordering used by bridge
        const bool canonicalSwitch{bridge.high1() != highL ||
              bridge.pos1() !=
              mumL.position0() + (highL ? mumL.length() - 1 : 0)};
        mum1_ = canonicalSwitch ? mumR : mumL;
        mum2_ = canonicalSwitch ? mumL : mumR;

        // Bridge found in the pair with anchor A
        return true;
      }
      // No bridge found in the pair with anchor A
      return false;
    }

    uint64_t pair_index() const { return pair_index_; }
    Pair pair() const { return bridges.mumdex.pair(pair_index_); }
    MUM mum1() const { return mum1_; }
    MUM mum2() const { return mum2_; }
    unsigned int chr1() const { return bridge.chr1(); }
    unsigned int chr2() const { return bridge.chr2(); }
    unsigned int pos1() const { return bridge.pos1(); }
    unsigned int pos2() const { return bridge.pos2(); }
    bool high1() const { return bridge.high1(); }
    bool high2() const { return bridge.high2(); }
    OneBridgeInfo the_bridge() const {
      return bridge;
    }

    std::array<std::string, 2> adjacent_sequences(
        const unsigned int adj_len) const {
      const MUM mums[2]{mum1_, mum2_};
      const bool highs[2]{high1(), high2()};
      std::array<std::string, 2> seqs;
      for (const unsigned int index : {0, 1}) {
        seqs[index] = bridges.mumdex.adjacent_sequence(
            pair_index_, mums[index], highs[index], adj_len);
      }
      return seqs;
    }

   private:
    const TBridges & bridges;
    const_iterator current;

    OneBridgeInfo bridge{};
    uint64_t pair_index_{0};
    MUM mum1_{};
    MUM mum2_{};
  };
  using const_iterator = Bridge;

 public:
  TBridges(const unsigned int chr1, const unsigned int pos1, const bool high1,
          const unsigned int chr2, const unsigned int pos2, const bool high2,
          const int64_t invariant_arg, const MUMdex & mumdex_arg) :
      chrA{high1 ? chr2 : chr1}, chrB{high1 ? chr1 : chr2},
                    posA{high1 ? pos2 : pos1}, posB{high1 ? pos1 : pos2},
                    highA{high1 ? high2 : high1}, highB{high1 ? high1 : high2},
                    invariant{invariant_arg}, mumdex{mumdex_arg},
                    anchorsA{mumdex.anchors(chrA, posA, highA)} { }

  TBridges(const OneBridgeInfo & b, const MUMdex & mumdex_arg) :
      chrA{b.high1() ? b.chr2() : b.chr1()},
                    chrB{b.high1() ? b.chr1() : b.chr2()},
                    posA{b.high1() ? b.pos2() : b.pos1()},
                    posB{b.high1() ? b.pos1() : b.pos2()},
                    highA{b.high1() ? b.high2() : b.high1()},
                    highB{b.high1() ? b.high1() : b.high2()},
                    invariant{b.invariant()}, mumdex{mumdex_arg},
                    anchorsA{mumdex.anchors(chrA, posA, highA)} { }

  Bridge begin() const { return Bridge{*this, anchorsA.begin()}; }
  Bridge end() const { return Bridge{*this, anchorsA.end()}; }
  bool exist() const { return begin() != end(); }

 private:
  const unsigned int chrA;
  const unsigned int chrB;
  const unsigned int posA;
  const unsigned int posB;
  const bool highA;
  const bool highB;
  const int64_t invariant;
  const MUMdex & mumdex;
  const Anchors anchorsA;
};

using MemoryBridges = TBridges<MemoryMUMdex>;
using PreMappedBridges = TBridges<PreMappedMUMdex>;
using UnMappedBridges = TBridges<UnMappedMUMdex>;
using FileBridges = TBridges<FileMUMdex>;
using Bridges = TBridges<MUMdex>;
using Bridge = Bridges::Bridge;

class PopBridgeInfo {
 public:
  class EndOfFile {};

  PopBridgeInfo(std::istream & in, const unsigned int chr1__) :
      pos1_{0}, pos2_{0},
    chr1_{static_cast<uint8_t>(chr1__)}, chr2_{0},
    n_people_{0}, n_bridges_{0},
    median_bridges_{0}, max_bridges_{0}, offset_{0},
    high1_{0}, high2_{0} {
    unsigned int chr2__;
    in >> pos1_ >> high1_
       >> chr2__ >> pos2_ >> high2_
       >> offset_
       >> n_people_ >> n_bridges_
       >> median_bridges_ >> max_bridges_;
    if (!in) throw EndOfFile();
    chr2_ = chr2__;
  }

  PopBridgeInfo(std::istream & in, const ChromosomeIndexLookup & lookup) :
      pos1_{0}, pos2_{0},
    chr1_{0}, chr2_{0},
    n_people_{0}, n_bridges_{0},
    median_bridges_{0}, max_bridges_{0}, offset_{0},
    high1_{0}, high2_{0} {
      std::string chr1__;
      std::string chr2__;
      in >> chr1__ >> pos1_ >> high1_
         >> chr2__ >> pos2_ >> high2_
         >> offset_
         >> n_people_ >> n_bridges_
         >> median_bridges_ >> max_bridges_;
      if (!in) throw EndOfFile();
      chr1_ = lookup[chr1__];
      chr2_ = lookup[chr2__];
    }

  explicit PopBridgeInfo(const BridgeInfo & bridge) :
      pos1_{bridge.pos1()},
    pos2_{bridge.pos2()},
    chr1_{static_cast<uint8_t>(bridge.chr1())},
    chr2_{static_cast<uint8_t>(bridge.chr2())},
    n_people_{1},
    n_bridges_{bridge.bridge_count()},
    median_bridges_{0}, max_bridges_{0},
    offset_{bridge.offset()},
    high1_{bridge.high1()},
    high2_{bridge.high2()} { }

  unsigned int chr1() const { return chr1_; }
  unsigned int chr2() const { return chr2_; }
  unsigned int pos1() const { return pos1_; }
  unsigned int pos2() const { return pos2_; }
  bool high1() const { return high1_; }
  bool high2() const { return high2_; }

  int16_t offset() const { return offset_; }
  int64_t invariant() const {
    return (high1_ ? 1 : -1) * static_cast<int64_t>(pos1_) +
        (high2_ ? 1 : -1) * static_cast<int64_t>(pos2_) + offset_;
  }
  char orientation_char() const {
    if (high1_ != high2_) {
      return '=';
    } else {
      if (high1_) {
        return 'i';
      } else {
        return 'o';
      }
    }
  }

  unsigned int n_people() const { return n_people_; }
  unsigned int n_bridges() const { return n_bridges_; }
  unsigned int median_bridges() const { return median_bridges_; }
  unsigned int max_bridges() const { return max_bridges_; }
  bool overflow() const {
    return max_bridges_ == std::numeric_limits<uint16_t>::max();
  }

  void combine(const BridgeInfo & bridge) {
    n_bridges_ += bridge.bridge_count();
    ++n_people_;
  }

  bool operator<(const BridgeInfo & other) const {
    if (chr1_ == other.chr1()) {
      if (pos1_ == other.pos1()) {
        if (high1_ == other.high1()) {
          if (chr2_ == other.chr2()) {
            if (pos2_ == other.pos2()) {
              if (high2_ == other.high2()) {
                return offset_ < other.offset();
              } else {
                return high2_ < other.high2();
              }
            } else {
              return pos2_ < other.pos2();
            }
          } else {
            return chr2_ < other.chr2();
          }
        } else {
          return high1_ < other.high1();
        }
      } else {
        return pos1_ < other.pos1();
      }
    } else {
      return chr1_ < other.chr1();
    }
  }

  template <class STREAM>
  void output(STREAM & stream) const {
    stream << ref_ptr->name(chr1())
           << pos1()
           << high1()
           << ref_ptr->name(chr2())
           << pos2()
           << high2()
           << orientation_char()
           << invariant()
           << offset()
           << n_people()
           << n_bridges()
           << median_bridges()
           << max_bridges();
  }

  template <class STREAM>
  void compact_output(STREAM & stream) const {
    stream << pos1()
           << high1()
           << chr2()
           << pos2()
           << high2()
           << offset()
           << n_people()
           << n_bridges()
           << median_bridges()
           << max_bridges();
  }

  void finalize(std::vector<unsigned int> & counts) {
    sort(counts.begin(), counts.end());
    median_bridges_ = counts[counts.size() / 2];
    max_bridges_ = counts.back();
    if (n_people_ != counts.size()) {
      throw Error("n_people discrepancy");
    }
    if (accumulate(counts.begin(), counts.end(), 0U) != n_bridges_) {
      throw Error("n_bridges discrepancy");
    }
  }

 private:
  uint32_t pos1_;                  // 32  32 4
  uint32_t pos2_;                  // 32  64 8
  uint8_t chr1_;                   //  8  72 9
  uint8_t chr2_;                   //  8  80 10
  uint16_t n_people_;              // 16  96 12
  uint32_t n_bridges_;             // 32 128 16
  uint16_t median_bridges_;        // 16 144 18
  uint16_t max_bridges_;           // 16 160 20
  int16_t offset_;                 // 16 176 22
  bool high1_;                     //  8 184 23
  bool high2_;                     //  8 192 24
};

template <class STREAM>
STREAM & operator<<(STREAM & out, const PopBridgeInfo & bridge) {
  bridge.output(out);
  return out;
}

// Matched allele and reference counts
class Counts {
 public:
  Counts() { }

  unsigned int count{0};
  std::map<std::string, unsigned int> allele_count{};
  unsigned int ref_count{0};
  unsigned int get_allele_count(const std::string & allele) const {
    auto found = allele_count.find(allele);
    if (found == allele_count.end()) {
      return 0;
    } else {
      return found->second;
    }
  }
};

class BridgeCounts {
 public:
  template <class MUMDEX>
  BridgeCounts(const MUMDEX & mumdex,
               const Reference & ref,
               const Mappability & mappability,
               const unsigned int chr[2],
               const unsigned int pos[2],
               const unsigned int high[2],
               const int offset,
               const bool count_dupes = true) {
    const unsigned int abspos[2]{ref.abspos(chr[0], pos[0]),
          ref.abspos(chr[1], pos[1])};
    const unsigned int map[2]{mappability.low_high(high[0], abspos[0]),
          mappability.low_high(high[1], abspos[1])};

    // Collect various counts for each anchor a and b
    for (const bool b : {false, true}) {
      const unsigned int early{152};
      const auto begin_index = mumdex.lower_bound(
          chr[b], pos[b] > early ? pos[b] - early : 0);
      const auto end_index = mumdex.lower_bound(
          chr[b], pos[b] + early);
      std::set<uint64_t> counted_anchors;
      std::set<uint64_t> counted_anchors_ref;
      std::set<uint64_t> counted_any;
      std::set<uint64_t> counted_any_ref;
      std::set<uint64_t> counted_bridges;
      std::set<uint64_t> counted_bridges_ref;
      for (auto index = begin_index; index != end_index; ++index) {
        const auto mum = mumdex.mum(*index);
        const auto pair = mumdex.pair(*index);
        if (pair.dupe() && !count_dupes) continue;
        const uint64_t pair_index{index->pair_index()};
        const uint64_t mum_index{mumdex.mums_start(pair_index) +
              index->mum_in_pair_index()};
        const int read_length(pair.length(mum.read_2()));
        const std::string sequence{
          mumdex.sequences(pair_index)[mum.read_2()]};

        const int gap_sequence_length{offset > 1 ? offset - 1 : 0};
        const int gap_sequence_start_pos{static_cast<int>(pos[b]) +
              (high[b] ? 1 : -gap_sequence_length)};
        const int gap_sequence_stop_pos{gap_sequence_start_pos +
              gap_sequence_length};

        const int gap_sequence_start_offset{
          mum.position0_to_read(gap_sequence_start_pos)};
        const int gap_sequence_stop_offset{
          mum.position0_to_read(gap_sequence_stop_pos)};

        const int start_offset{
          gap_sequence_start_offset < gap_sequence_stop_offset ?
              gap_sequence_start_offset : gap_sequence_stop_offset + 1};
        const int stop_offset{
          gap_sequence_start_offset < gap_sequence_stop_offset ?
              gap_sequence_stop_offset : gap_sequence_start_offset + 1};

        const bool gap_in_read{start_offset >= 0 &&
              stop_offset <= read_length};

        if (gap_in_read) {
          const std::string gap_sequence{[&sequence,
                                     gap_sequence_start_offset,
                                     gap_sequence_stop_offset,
                                     mum]() {
              std::string result;
              const int increment{
                gap_sequence_start_offset < gap_sequence_stop_offset ?
                    1 : -1};
              for (int o{gap_sequence_start_offset};
                   o != gap_sequence_stop_offset; o += increment) {
                const char base{sequence[o]};
                result += mum.flipped() ? complement(base) : base;
              }
              return result;
            }()};

          // Ignore reads with gap sequences containing N
          if (gap_sequence.find('N') != std::string::npos) continue;

          const int mum_anchor_pos{static_cast<int>(mum.position0()) +
                static_cast<int>(high[b] ? mum.length() - 1 : 0)};
          const int mum_anchor_offset{mum.position0_to_read(mum_anchor_pos)};
          // Is MUM an anchor?
          if (mum_anchor_pos == static_cast<int>(pos[b]) &&
              mum_anchor_offset != 0 &&  // redundant since gap is in read
              mum_anchor_offset + 1 != read_length) {
            if (!counted_anchors.count(pair_index)) {
              ++anchor[b].count;
              ++anchor[b].allele_count[gap_sequence];
              counted_anchors.insert(pair_index);
            }
            // Count bridges
            if (b) {
              for (uint64_t m{mumdex.mums_start(pair_index)};
                   m != mumdex.mums_stop(pair_index); ++m) {
                if (m == mum_index) continue;
                const MUM mum2{mumdex.mum(m)};
                if (mum2.read_2() != mum.read_2()) continue;
                const int mum2_anchor_pos{static_cast<int>(mum2.position0()) +
                      static_cast<int>(high[1 - b] ? mum2.length() - 1 : 0)};
                const int mum2_anchor_offset{
                  mum2.position0_to_read(mum2_anchor_pos)};
                const int expected_offset{mum_anchor_offset +
                      (mum.flipped() == high[b] ? -1 : 1) * offset};
                if (mum2_anchor_offset == expected_offset &&
                    mum2_anchor_pos == static_cast<int>(pos[1 - b]) &&
                    !counted_bridges.count(pair_index)) {
                  ++bridge.count;
                  ++bridge.allele_count[gap_sequence];
                  counted_bridges.insert(pair_index);
                }
              }
            }
          }
          if (!counted_any.count(pair_index)) {
            ++any[b].count;
            ++any[b].allele_count[gap_sequence];
            counted_any.insert(pair_index);
          }
          // Count reference allele if seen
          if (start_offset >= static_cast<int>(mum.offset()) &&
              stop_offset <= static_cast<int>(mum.offset() + mum.length())) {
            if (!counted_any_ref.count(pair_index)) {
              ++any[b].ref_count;
              counted_any_ref.insert(pair_index);
            }
            // Correct anchor reference count to match anchor count bias
            const int map_offset{static_cast<int>(map[b]) - 1};
            const int mum_needs_to_cover_pos{static_cast<int>(pos[b]) +
                  map_offset * (high[b] ? -1 : 1)};
            if (mum_needs_to_cover_pos >= static_cast<int>(mum.position0()) &&
                mum_needs_to_cover_pos <
                static_cast<int>(mum.position0() + mum.length()) &&
                !counted_anchors_ref.count(pair_index)) {
              ++anchor[b].ref_count;
              counted_anchors_ref.insert(pair_index);
              // Reference count on par with bridge count
              if (b) {
                const int map_offset2{static_cast<int>(map[1 - b]) - 1};
                const int mum_needs_to_cover_pos2{static_cast<int>(pos[b]) +
                      offset + map_offset2 * (high[b] ? 1 : -1)};
                const int mum_needs_to_cover_offset2{
                  mum.position0_to_read(mum_needs_to_cover_pos2)};
                if (mum_needs_to_cover_offset2 >=
                    static_cast<int>(mum.offset()) &&
                    mum_needs_to_cover_offset2 <
                    static_cast<int>(mum.offset() + mum.length()) &&
                    !counted_bridges_ref.count(pair_index)) {
                  ++bridge.ref_count;
                  counted_bridges_ref.insert(pair_index);
                }
              }
            }
          }
        }
      }
    }
  }

  Counts bridge{};
  Counts anchor[2];
  Counts any[2];
};


}  // namespace paa

#endif  // PAA_BRIDGES_H
```
