## Supplementary material for "G-Graph: An interactive genomic graph viewer": MUMdex Source Code for G-Graph: fasta.h

```
//
// fasta.h
//
// dealing with the reference for suffix array use
//
// Modifications of sparseMEM Copyright Peter Andrews 2013-2016 @ CSHL
//

#ifndef LONGMEM_FASTA_H_
#define LONGMEM_FASTA_H_

#include <algorithm>
#include <iostream>
#include <fstream>
#include <numeric>
#include <sstream>
#include <string>
#include <vector>

#include "error.h"
#include "mumdex.h"
#include "strings.h"

namespace paa {

// Trim a string, giving start and end in trimmed version.
// NOTE: Assumes line.length() > 0!!!!
inline void trim(const std::string & line, uint64_t & start, uint64_t & end) {
  // Trim leading spaces.
  for (uint64_t i = start; i < line.size(); ++i) {
    if (line[i] != ' ') {
      start = i;
      break;
    }
  }
  // Trim trailing spaces.
  for (uint64_t i = line.size() ; i != 1; --i) {
    if (line[i-1] != ' ') {
      end = i;
      break;
    }
  }
}

class RefArgs {
 public:
  RefArgs(const char * ref_fasta_ = nullptr,
          const bool rcref_ = false, const bool verbose_ = false) :
      ref_fasta(ref_fasta_), rcref(rcref_), verbose(verbose_) { }
  const char * ref_fasta;
  bool rcref;
  bool verbose;

 private:
  RefArgs & operator=(const RefArgs &) = delete;
};

class Sequence : public RefArgs {
 public:
  explicit Sequence(const RefArgs & arguments) :
      RefArgs(arguments), using_mapping(false) {
    const time_t start_time = time(nullptr);

    // Reference cache filename
    std::ostringstream saved_reference_stream;
    saved_reference_stream << ref_fasta << ".bin";
    bin_base = saved_reference_stream.str();
    mkdir(bin_base);
    saved_reference_stream << "/rc" << rcref << ".ref";
    bin_base = saved_reference_stream.str();
    saved_reference_stream << ".bin";
    const std::string saved_reference = saved_reference_stream.str();

    const uint64_t fasta_size = file_size(ref_fasta);

    // Load or create reference
    if (readable(saved_reference)) {
      if (verbose) std::cerr << "loading reference binary" << std::endl;

      FILE * reference = fopen(saved_reference.c_str(), "rb");
      if (!reference) throw Error("could not open reference bin file")
                          << saved_reference << "for reading";

      uint64_t fasta_saved_size;
      bread(reference, fasta_saved_size, "fasta_size");
      if (fasta_size != fasta_saved_size)
        throw Error("")
            << "reference fasta size has changed\n"
            << "maybe the reference has changed?\n"
            << "If so, you will need to delete the current "
            << "reference to proceed";
      bread(reference, N, "N");
      using_mapping = true;
      bread(bin_base + ".seq.bin", seq, "seq", N);
      uint64_t descr_size;
      bread(reference, descr_size, "descr_size");
      startpos.resize(descr_size);
      sizes.resize(descr_size);
      descr.resize(descr_size);
      for (unsigned int i = 0; i != descr_size; ++i) {
        bread(reference, startpos[i], "startpos");
        bread(reference, sizes[i], "sizes");
        uint64_t string_size;
        bread(reference, string_size, "string_size");
        descr[i].resize(string_size);
        bread(reference, descr[i][0], "description string", string_size);
      }
      bread(reference, maxdescrlen, "maxdescrlen");
      if (fclose(reference) != 0)
        throw Error("problem closing reference file");
    } else {
      if (verbose) std::cerr << "loading reference from fasta" << std::endl;
      // Reservation assumes fasta line lengths < 100 bases to limit size
      seq_vec.reserve((rcref ? 2 : 1) * 0.99 * fasta_size);
      std::string meta, line;
      uint64_t length = 0;

      // Everything starts at zero.
      startpos.push_back(0);

      std::ifstream data(ref_fasta);

      if (!data.is_open()) throw Error("unable to open") << ref_fasta;

      while (!data.eof()) {
        getline(data, line);  // Load one line at a time.
        if (!data.eof() && line.size() == 0) continue;

        uint64_t start = 0, end = line.size();

        // Meta tag line and start of a new sequence.
        if (data.eof() || line[0] == '>') {
          // Save previous sequence and meta data.
          if (length > 0) {
            const uint64_t this_start = startpos.back();
            if (false && verbose) std::cerr << meta << " " << length
                                            << " " << this_start << std::endl;
            descr.push_back(meta);
            if (rcref || !data.eof()) {
              seq_vec.push_back('`');  // ` character used to separate strings
              startpos.push_back(seq_vec.size());
            }
            sizes.push_back(length);
            if (rcref) {
              descr.push_back(meta);
              sizes.push_back(length);
              //            string reversed(seq_vec.begin() + this_start,
              //                seq_vec.begin() + this_start + length);
              // &* to avoid very valid mac warning:
              // bad difference type for string iterator.  use pointer instead
              std::string reversed(&*seq_vec.begin() + this_start,
                                   &*seq_vec.begin() + this_start + length);
              reverse_complement(&reversed);
              seq_vec.insert(seq_vec.end(), reversed.begin(), reversed.end());
              if (!data.eof()) {
                seq_vec.push_back('`');  // ` character used to separate strings
                startpos.push_back(seq_vec.size());
              }
            }
            if (data.eof()) break;
          }
          // Reset parser state.
          start = 1;
          meta = "";
          length = 0;
        }
        trim(line, start, end);
        // Collect meta data.
        if (line[0] == '>') {
          for (uint64_t i = start; i != end; ++i) {
            if (line[i] == ' ') break;
            meta += line[i];
          }
        } else {  // Collect sequence data.
          length += end - start;
          for (uint64_t i = start; i != end; ++i) {
            const char base = toupper(line[i]);
            seq_vec.push_back(base == 'N' ? 'X' : base);
          }
        }
      }
      seq_vec.push_back('$');
      if (false && verbose)
        std::cerr << "seq_vec.length=" << seq_vec.size() << std::endl;

      N = seq_vec.size();
      seq = &seq_vec[0];

      // Get maximum reference sequence description length.
      maxdescrlen = 0;
      for (uint64_t i = 0; i != descr.size(); ++i) {
        if (maxdescrlen < descr[i].size())  maxdescrlen = descr[i].size();
      }

      if (verbose) std::cerr << "saving reference binary" << std::endl;

      FILE * reference = fopen(saved_reference.c_str(), "wb");
      if (reference == nullptr)
        throw Error("could not open reference")
            << saved_reference << "for writing";
      bwrite(reference, fasta_size, "fasta_size");
      bwrite(reference, N, "N");
      bwrite(bin_base + ".seq.bin", seq[0], "seq", N);
      const uint64_t descr_size = descr.size();
      bwrite(reference, descr_size, "descr_size");
      for (unsigned int i = 0; i != descr_size; ++i) {
        bwrite(reference, startpos[i], "startpos");
        bwrite(reference, sizes[i], "sizes");
        const uint64_t string_size = descr[i].size();
        bwrite(reference, string_size, "string_size");
        bwrite(reference, descr[i][0], "description string", string_size);
      }
      bwrite(reference, maxdescrlen, "maxdescrlen");
      if (fclose(reference) != 0)
        throw Error("problem closing reference file");

      // Create simple reference output for use later after mapping
      const GrowingReference ref(*this,
                                 GrowingReference::CreateFromAlignerSequence());
      ref.save(ref_fasta);
    }

    const time_t end_time = time(nullptr);
    if (verbose) std::cerr << "constructed reference in "
                           << end_time - start_time << " seconds" << std::endl;
  }

  ~Sequence() {
    if (memory_mapped && using_mapping) {
      if (munmap(seq, N * sizeof(*seq)))
        std::cerr << "sequence memory unmap failure" << std::endl;
    } else {
      if (using_mapping) free(seq);
    }
  }
  char operator[] (const uint64_t n) const { return seq[n]; }
  uint64_t bytes() const {
    return sizeof(Sequence) + seq_vec.capacity() +
        sizeof(uint64_t) * (startpos.capacity() + sizes.capacity()) +
        bin_base.capacity() + accumulate(
            descr.begin(), descr.end(), 0ul,
            [](const uint64_t tot, const std::string & name) {
              return tot + name.capacity();
            });
  }
  std::string sam_header() const {
    std::ostringstream out;
    out << "@HD\tVN:1.0\tSO:unsorted" << std::endl;
    for (unsigned int chr = 0; chr < sizes.size();
         chr += (rcref ? 2 : 1)) {
      out << "@SQ\tSN:" << descr[chr] << "\tLN:" << sizes[chr] << std::endl;
    }
    out << "@PG\tID:mumdex\tPN:mumdex\tVN:0.5" << std::endl;
    return out.str();
  }

  uint64_t N{};  // !< Length of the sequence.
  std::vector<char> seq_vec{};
  char * seq{};
  std::vector<std::string> descr{};
  std::vector<uint64_t> startpos{};
  std::vector<uint64_t> sizes{};
  uint64_t maxdescrlen{};
  std::string bin_base{};

 private:
  bool using_mapping{};
  Sequence(const Sequence &) = delete;
  Sequence & operator=(const Sequence &) = delete;
};

}  // namespace paa

#endif  // LONGMEM_FASTA_H_
```
