## Supplementary material for "G-Graph: An interactive genomic graph viewer": MUMdex Source Code for G-Graph: genes.cpp

```
//
// genes.cpp
//
// dealing with gene information
//
// Copyright 2014 Peter Andrews @ CSHL
//

#include "genes.h"

#include <algorithm>
#include <fstream>
#include <iostream>
#include <map>
#include <sstream>
#include <string>
#include <vector>

#include "error.h"
#include "mumdex.h"

using std::cerr;
using std::endl;
using std::ifstream;
using std::iostream;
using std::istream;
using std::istringstream;
using std::ofstream;
using std::ostream;
using std::ostringstream;
using std::map;
using std::sort;
using std::string;
using std::vector;

namespace paa {

GeneXrefs::GeneXrefs(const string & file_name) {
  ifstream file(file_name.c_str());
  if (!file) throw Error("Problem opening GeneXref file") << file_name;
  GeneXref info;
  while (getline(file, info.name, '\t')) {
    getline(file, info.mRNA, '\t');
    getline(file, info.spID, '\t');
    getline(file, info.spDisplayID, '\t');
    getline(file, info.geneSymbol, '\t');
    getline(file, info.refSeq, '\t');
    getline(file, info.protAcc, '\t');
    getline(file, info.description, '\t');
    file.ignore(1000, '\n');
    data[info.name] = info;
  }
  if (data.empty()) throw Error("Problem loading data for GeneXrefs");
}

KnownGene::KnownGene(const string & line, const unsigned int n,
                     const ChromosomeIndexLookup & index) {
  istringstream input(line);
  input >> name >> chr_name >> strand
        >> t_start >> t_stop >> c_start >> c_stop
        >> n_exons;
  // sout << name << chr << n_exons << endl;
  exon_starts.resize(n_exons);
  for (unsigned int i = 0; i != n_exons; ++i) {
    input >> exon_starts[i];
    input.get();
  }
  exon_stops.resize(n_exons);
  for (unsigned int i = 0; i != n_exons; ++i) {
    input >> exon_stops[i];
    input.get();
  }
  if (!input) throw Error("Problem parsing line") << line;
  input.ignore(10000, '\n');

  chr = index[chr_name];

  // Set up junctions
  /*
    junctions.emplace_back(n, junctions.size(), false,
    chr, exon_starts[0], vector<int>());
  */
  for (unsigned int i = 1; i != n_exons; ++i) {
    vector<int> invariants;
    for (unsigned int j = i; j != 0; --j) {
      invariants.push_back(exon_stops[j - 1] - exon_starts[i]);
    }
    /*
      junctions.emplace_back(n, junctions.size(), true,
      chr, exon_stops[i], invariants);
    */
    junctions.emplace_back(n, junctions.size(),
                           chr, exon_starts[i], invariants);
  }
}

KnownGenes::KnownGenes(const std::string & genes_file_name,
                       const std::string & isoforms_file_name,
                       const ChromosomeIndexLookup & index,
                       const Reference & ref_arg) : ref{ref_arg} {
  // Read isoform lookup file
  ifstream isoform_file(isoforms_file_name.c_str());
  if (!isoform_file) throw Error("Problem opening") << isoforms_file_name;
  unsigned int cluster_index;
  string gene_name;
  map<string, unsigned int> isoform_lookup;
  while (isoform_file >> cluster_index >> gene_name) {
    isoform_lookup[gene_name] = cluster_index;
  }
  // Read known gene info to get exon boundaries
  ifstream genes_file(genes_file_name.c_str());
  if (!genes_file) throw Error("Problem opening") << genes_file_name;
  string line;
  genes.reserve(100000);
  while (getline(genes_file, line)) {
    const auto search_to = line.begin() + (line.size() > 30 ? 30 : line.size());
    if (find(line.begin(), search_to, '_') == search_to) {  // Done badly
      genes.emplace_back(line, genes.size(), index);
      KnownGene & gene = genes.back();
      gene.cluster_index = isoform_lookup[gene.name];
    }
  }
  sort(genes.begin(), genes.end());
  for (unsigned int g = 0; g != genes.size(); ++g) {
    for (unsigned int j = 0; j != genes[g].junctions.size(); ++j) {
      KnownGene & gene(genes[g]);
      gene.junctions[j].gene_index = g;
      junctions.push_back(gene.junctions[j]);
    }
  }
  sort(junctions.begin(), junctions.end(), JunctionSorter());
  if (0) cerr << "Loaded " << genes.size() << " genes with "
              << junctions.size() << " junctions" << endl;
}

/*
stringstream & KnownGenes::generate_bed(stringstream & inout) const {
  generate_bed(inout);
  return inout;
}
*/

void KnownGenes::generate_bed(ostream & output) const {
  if (junctions.empty()) throw Error("Empty junctions");
  unsigned int last_chr = 100000;
  unsigned int start_pos = 0;
  unsigned int last_pos = 0;
  for (unsigned int j = 0; j != junctions.size(); ++j) {
    const PseudoGeneJunction & junction = junctions[j];
    if (last_chr != junction.chr ||
        last_pos + junction_window < junction.pos) {
      if (start_pos) {
        output << ref.name(last_chr) << "\t"
               << start_pos << "\t" << last_pos << '\n';
      }
      last_chr = junction.chr;
      start_pos = junction.pos > junction_window + 1 ?
          junction.pos - junction_window : 1;
    }
    last_pos = junction.pos + junction_window;
  }
  output << ref.name(last_chr) << "\t" << start_pos << "\t" << last_pos << '\n';
}

string KnownGenes::generate_bed(const string & name) const {
  ofstream output(name.c_str());
  if (!output) throw Error("Problem opening bed for writing") << name;
  generate_bed(output);
  return name;
}

AllJunctionCounts::AllJunctionCounts(const KnownGenes & genes,
                                     const std::vector<string> & samples) :
    junction_counts(samples.size(), vector<unsigned int>(genes.size())),
    total_counts(samples.size(), vector<unsigned int>(genes.size())) {
  for (unsigned int s = 0; s != samples.size(); ++s) {
    if (0) cerr << s << " " << samples[s] << endl;
    counts.emplace_back(genes);
    ostringstream file_name;
    file_name << "/data/safe/paa/analysis/mums/output/samples/"
              << samples[s] << "/pseudo.out";
    ifstream input(file_name.str().c_str());
    if (!input) throw Error("Cound not open pseudo counts file for sample")
                    << samples[s];
    string line;
    string gene_name;
    unsigned int junction_count;
    unsigned int n_junctions;
    unsigned int total_count;
    unsigned int gene_index;
    string chr;
    unsigned int pos;
    unsigned int junction_index;
    unsigned int junction_hits;
    while (getline(input, line)) {
      istringstream input_line(line.c_str());
      input_line >> gene_name >> junction_count >> n_junctions
                 >> total_count >> gene_index >> chr >> pos;
      ++counts[s].n_genes;
      counts[s].n_junctions += junction_count;
      counts[s].total_count += total_count;
      for (unsigned int j = 0; j != junction_count; ++j) {
        input_line >> junction_index;
        input_line >> junction_hits;
        counts[s][gene_index][junction_index].invariant_count = junction_hits;
      }
      junction_counts[s][gene_index] = junction_count;
      total_counts[s][gene_index] = total_count;
      if (!input_line) throw Error("pseudogene junction count parse error")
                           << line;
    }
  }
  cerr << "Loaded " << samples.size() << " sample count files" << endl;
}

#if 0
      // Gene exon information suitable for searching for interval overlaps
      for (unsigned int g{0}; g != genes.size(); ++g) {
        const KnownGene & gene{genes[g]};
        for (unsigned int e{0}; e != gene.n_exons; ++e) {
          exons.emplace_back(gene.chr,
                             gene.exon_starts[e], gene.exon_stops[e], g);
        }
      }
      sort(exons.begin(), exons.end());
      unsigned int last_chromosome{0};
      unsigned int lowest_position{0};
      for (unsigned int i{0}; i != exons.size(); ++i) {
        const unsigned int e(exons.size() - i - 1);
        Exon & exon{exons[e]};
        if (last_chromosome != exon.chromosome) {
          lowest_position = exon.start_position;
        } else if (lowest_position > exon.start_position) {
          lowest_position = exon.start_position;
        }
        exon.lowest_later_start = lowest_position;
      }
#endif

}  // namespace paa
```
