## Supplementary material for "G-Graph: An interactive genomic graph viewer": MUMdex Source Code for G-Graph: genes.h

```
//
// genes.h
//
// information about genes
//
// Copyright 2014 Peter Andrews @ CSHL
//

#ifndef PAA_GENES_H
#define PAA_GENES_H

#include <algorithm>
#include <fstream>
#include <iostream>
#include <map>
#include <sstream>
#include <string>
#include <utility>
#include <vector>

#include "mumdex.h"

namespace paa {

const unsigned int junction_window = 5;

struct Gap {
  Gap(const unsigned int chr_, const unsigned int start_,
      const unsigned int stop_, const std::string type_) :
      chr{chr_}, start{start_}, stop{stop_}, type{type_} {}

  unsigned int chr;
  unsigned int start;
  unsigned int stop;
  std::string type;
};

class Gaps {
 public:
  Gaps(const std::string & file_name,
       const ChromosomeIndexLookup & lookup,
       const std::string & types) {
    std::ifstream input{file_name.c_str()};
    if (!input) throw Error("Cound not open gap input file") << file_name;
    input.ignore(10000, '\n');
    std::string line;
    unsigned int bin;
    std::string chr_name;
    unsigned int start;
    unsigned int stop;
    unsigned int ix;
    std::string n;
    unsigned int size;
    std::string type;
    std::string bridge;
    while (getline(input, line)) {
      std::istringstream line_str{line.c_str()};
      line_str >> bin >> chr_name >> start >> stop
               >> ix >> n >> size >> type >> bridge;
      if (size && types.find(type) != std::string::npos)
        data.emplace_back(lookup[chr_name], start, stop, type);
    }
    std::cerr << "Loaded " << data.size() << " gaps" << std::endl;
  }

  std::vector<Gap>::const_iterator begin() const { return data.begin(); }
  std::vector<Gap>::const_iterator end() const { return data.end(); }

 private:
  std::vector<Gap> data{};
};

class GeneXref {
 public:
  GeneXref() { }
  std::string name{};
  std::string mRNA{};
  std::string spID{};
  std::string spDisplayID{};
  std::string geneSymbol{};
  std::string refSeq{};
  std::string protAcc{};
  std::string description{};
  bool operator<(const GeneXref & rhs) const { return name < rhs.name; }
};

class GeneXrefs {
 public:
  explicit GeneXrefs(const std::string & file_name);
  const GeneXref & operator[](const std::string & name) const {
    return data.at(name);
  }

 private:
  std::map<std::string, GeneXref> data{};
};

class PseudoGeneJunction : public PosInfo {
 public:
  PseudoGeneJunction(const unsigned int gene_index_,
                     const unsigned int junction_index_,
                     const unsigned int chr_, const unsigned int pos_,
                     const std::vector<int> & invariants_) :
      PosInfo(chr_, pos_), gene_index(gene_index_),
      junction_index(junction_index_),
      invariants(invariants_) {}
  /*
  bool operator<(const PosInfo & right) const {
    return *this < right;
  }
  */
  unsigned int gene_index;
  unsigned int junction_index;
  std::vector<int> invariants;
};

enum class HitType { none, gene, intron, exon };
template <class HIT>
bool hit_less(const HIT lhs, const HIT rhs) {
  if (lhs == rhs) return false;
  if (lhs == HitType::none) return true;
  if (rhs == HitType::none) return false;
  if (lhs == HitType::gene) return true;
  if (rhs == HitType::gene) return false;
  if (lhs == HitType::intron) return true;
  if (rhs == HitType::intron) return false;
  if (lhs == HitType::exon) return true;
  return false;
}


class ChromosomeIndexLookup;
class KnownGene {
 public:
  KnownGene() = default;
  explicit KnownGene(const std::string & line, const unsigned int n,
                     const ChromosomeIndexLookup & index);
  std::string name{};
  std::string chr_name{};
  unsigned int chr{};
  char strand{};
  unsigned int t_start{};
  unsigned int t_stop{};
  unsigned int c_start{};
  unsigned int c_stop{};
  unsigned int n_exons{};
  unsigned int cluster_index{};
  std::vector<unsigned int> exon_starts{};
  std::vector<unsigned int> exon_stops{};
  std::vector<PseudoGeneJunction> junctions{};
  unsigned int length() const {
    return t_stop - t_start;
  }
  bool in_exon(const unsigned int c, const unsigned int p) const {
    if (c != chr) return false;
    for (unsigned int e{0}; e != exon_starts.size(); ++e) {
      if (p >= exon_starts[e] && p < exon_stops[e]) return true;
    }
    return false;
  }
  bool operator<(const KnownGene & rhs) const {
    if (chr == rhs.chr) {
      if (t_start == rhs.t_start) {
        return t_stop < rhs.t_stop;
      } else {
        return t_start < rhs.t_start;
      }
    } else {
      return chr < rhs.chr;
    }
  }
  HitType hit(const unsigned int chr_, const unsigned int pos_) const {
    if (chr != chr_ || pos_ < t_start || pos_ >= t_stop) {
      return HitType::none;
    } else if (pos_ < c_start || pos_ >= c_stop) {
      return HitType::gene;
    } else {
      for (unsigned int e{0}; e != exon_starts.size(); ++e) {
        if (pos_ >= exon_starts[e] && pos_ < exon_stops[e]) {
          return HitType::exon;
        }
      }
      return HitType::intron;
    }
  }
};

struct GeneFinder {
  bool operator()(const KnownGene & lhs, const PosInfo & rhs) {
    if (lhs.chr == rhs.chr) {
      return lhs.t_start < rhs.pos;
    } else {
      return lhs.chr < rhs.chr;
    }
  }
  bool operator()(const PosInfo & lhs, const KnownGene & rhs) {
    if (lhs.chr == rhs.chr) {
      return lhs.pos < rhs.t_start;
    } else {
      return lhs.chr < rhs.chr;
    }
  }
};

class JunctionCount {
 public:
  unsigned int invariant_count = 0;
  // unsigned int anchor_count = 0;
};

class KnownGenes {
 public:
  // Empty gene information in case UCSC files are unavailable
  explicit KnownGenes(const Reference & ref_arg) : ref{ref_arg} {}
  KnownGenes(const std::string & genes_file_name,
             const std::string & isoforms_file_name,
             const ChromosomeIndexLookup & index,
             const Reference & ref_arg);
  const KnownGene & operator[] (const unsigned int i) const {
    return genes[i];
  }
  unsigned int size() const { return static_cast<unsigned int>(genes.size()); }
  std::vector<KnownGene>::const_iterator begin() const { return genes.begin(); }
  std::vector<KnownGene>::const_iterator end() const { return genes.end(); }
  // std::stringstream & generate_bed(std::stringstream & inout) const;
  void generate_bed(std::ostream & output) const;
  std::string generate_bed(const std::string & name) const;
  std::vector<unsigned int> find_genes(const unsigned int chr_arg,
                                       const unsigned int pos_arg) const {
    using ChrPos = std::pair<unsigned int, unsigned int>;
    std::vector<KnownGene>::const_iterator match_end{lower_bound(
        genes.begin(), genes.end(), ChrPos{chr_arg, pos_arg},
        [] (const KnownGene & gene, const ChrPos chrpos) {
          if (gene.chr == chrpos.first) {
            return gene.t_start < chrpos.second + 1;
          }
          return gene.chr < chrpos.first;
        })};
    std::vector<unsigned int> result;
    if (match_end == genes.end()) {
      --match_end;
    }
    while (true) {
      const KnownGene & gene{*match_end};
      if (gene.chr <= chr_arg) {
        if (gene.chr < chr_arg) break;
        if (gene.t_stop + 100000 < pos_arg) break;
        if (gene.t_start < pos_arg && pos_arg < gene.t_stop) {
          result.push_back(
              static_cast<unsigned int>(match_end - genes.begin()));
        }
        if (match_end == genes.begin()) break;
      }
      --match_end;
    }
    return result;
  }

  using GeneOverlap = std::pair<unsigned int, unsigned int>;
  using GeneOverlaps = std::vector<GeneOverlap>;
  GeneOverlaps find_genes(const unsigned int chr_arg,
                          const unsigned int low_pos,
                          const unsigned int high_pos) const {
    using ChrPos = std::pair<unsigned int, unsigned int>;
    std::vector<KnownGene>::const_iterator match_end{lower_bound(
        genes.begin(), genes.end(), ChrPos{chr_arg, high_pos},
        [] (const KnownGene & gene, const ChrPos chrpos) {
          if (gene.chr == chrpos.first) {
            return gene.t_start < chrpos.second + 1;
          }
          return gene.chr < chrpos.first;
        })};
    GeneOverlaps result;
    if (match_end == genes.end()) {
      --match_end;
    }
    while (true) {
      const KnownGene & gene{*match_end};
      if (gene.chr <= chr_arg) {
        if (gene.chr < chr_arg) break;
        if (gene.t_stop + 100000 < low_pos) break;
        if (gene.t_start < high_pos && low_pos < gene.t_stop) {
          GeneOverlap overlap{match_end - genes.begin(), 0};
          for (unsigned int e{0}; e != gene.n_exons; ++e) {
            if (gene.exon_starts[e] < high_pos &&
                gene.exon_stops[e] > low_pos) {
              ++overlap.second;
            }
          }
          result.push_back(overlap);
        }
      }
      if (match_end-- == genes.begin()) break;
    }
    return result;
  }

 private:
  std::vector<KnownGene> genes{};

 public:
  const Reference & ref;
  std::vector<PseudoGeneJunction> junctions{};
};

class GeneLookup {
 public:
  using KGP = const KnownGene *;
  using MMap = std::multimap<std::string, KGP>;
  using MMapI = MMap::value_type;
  using Iter = MMap::const_iterator;
  using ER = std::pair<Iter, Iter>;
  explicit GeneLookup(const KnownGenes & genes_, const GeneXrefs & xrefs) :
      genes{genes_} {
    for (const KnownGene & gene : genes) {
      data.emplace(xrefs[gene.name].geneSymbol, &gene);
    }
  }
  // const GeneXref & operator[](const std::string & name) const {
  //   return data.at(name);
  // }
  ER isoforms(const std::string & name) const {
     return data.equal_range(name);
  }

 private:
  const KnownGenes & genes;
  MMap data{};
};

class JunctionCounts {
 public:
  explicit JunctionCounts(const KnownGenes & genes) {
    for (unsigned int g = 0; g != genes.size(); ++g) {
      const KnownGene & gene = genes[g];
      junctions.push_back(std::vector<JunctionCount>(gene.junctions.size()));
    }
  }
  const std::vector<JunctionCount> & operator[] (const unsigned int i) const {
    return junctions[i];
  }
  std::vector<JunctionCount> & operator[] (const unsigned int i) {
    return junctions[i];
  }
  unsigned int n_genes{0};
  unsigned int n_junctions{0};
  unsigned int total_count{0};

 private:
  std::vector<std::vector<JunctionCount> > junctions{};
};

class AllJunctionCounts {
 public:
  AllJunctionCounts(const KnownGenes & genes,
                    const std::vector<std::string> & samples);
  const JunctionCounts & operator[] (const unsigned int i) const {
    return counts[i];
  }
  unsigned int size() const {
    return static_cast<unsigned int>(counts.size());
  }
 private:
  std::vector<JunctionCounts> counts{};
 public:
  std::vector<std::vector<unsigned int> > junction_counts;
  std::vector<std::vector<unsigned int> > total_counts;
};

class JunctionSorter {
 public:
  bool operator()(const PosInfo * left,
                  const PosInfo * right) const {
    return *left < *right;
  }
  bool operator()(const PosInfo left,
                  const PosInfo * right) const {
    return left < *right;
  }
  bool operator()(const PosInfo * left,
                  const PosInfo right) const {
    return *left < right;
  }
  bool operator()(const PosInfo left,
                  const PosInfo right) const {
    return left < right;
  }
};

struct JunctionSave {
  JunctionSave() { }
  JunctionSave(const unsigned int jind, const unsigned int iind,
               const int inv, const unsigned int pos) :
      junction_index{jind}, invariant_index{iind},
    invariant{inv}, position{pos} { }
  unsigned int junction_index{};
  unsigned int invariant_index{};
  int invariant{};
  unsigned int position{};
  bool operator<(const JunctionSave other) const {
    if (junction_index == other.junction_index) {
      if (position == other.position) {
        if (invariant_index == other.invariant_index) {
          return invariant < other.invariant;
        } else {
          return invariant_index < other.invariant_index;
        }
      } else {
        return position < other.position;
      }
    } else {
      return junction_index < other.junction_index;
    }
  }
};

struct OldJunctionSave {
  unsigned int junction_index{};
  unsigned int invariant_index{};
  int invariant{};
  bool operator<(const OldJunctionSave other) const {
    if (junction_index == other.junction_index) {
      if (invariant_index == other.invariant_index) {
        return invariant < other.invariant;
      } else {
        return invariant_index < other.invariant_index;
      }
    } else {
      return junction_index < other.junction_index;
    }
  }
};

struct JunctionInfo : public JunctionSave {
  struct JunctionReadError { };
  JunctionInfo(std::istream & in, uint64_t sample_arg) :
      sample(sample_arg) {
    in >> junction_index >> invariant_index >> invariant >> position >> count;
    if (!in) throw JunctionReadError();
  }
  uint64_t count{};
  uint64_t sample{};
};

class GeneInfo {
 public:
  GeneInfo(const HitType & hit_, const unsigned int exon_) :
      hit{hit_}, exon{exon_} {}
  HitType hit;
  unsigned int exon{0};
  bool operator<(const GeneInfo & other) const {
    if (exon == other.exon) {
      return hit_less(hit, other.hit);
    } else {
      return exon < other.exon;
    }
  }
};

class TempGeneInfo {
 public:
  TempGeneInfo(const unsigned int gene_, const unsigned int pos_,
               const GeneInfo info_) :
      gene{gene_}, pos{pos_}, info{info_} {}

  unsigned int gene;
  unsigned int pos;
  GeneInfo info;
  bool operator<(const TempGeneInfo & other) const {
    if (pos == other.pos) {
      if (gene == other.gene) {
        return info < other.info;
      } else {
        return gene < other.gene;
      }
    } else {
      return pos < other.pos;
    }
  }
};

class GeneInfoInterval {
 public:
  GeneInfoInterval(const unsigned int start_, const unsigned int stop_) :
      start_pos{start_}, stop_pos{stop_} {}
  unsigned int start_pos;
  unsigned int stop_pos;
  unsigned int size() const {
    return stop_pos - start_pos;
  }
  std::map<unsigned int, GeneInfo> info{};
};

class GeneHitFinder {
 public:
  explicit GeneHitFinder(const KnownGenes & genes) {
    std::vector<TempGeneInfo> info;
    unsigned int last_chr{genes[0].chr};
    for (unsigned int g{0}; g <= genes.size(); ++g) {
      if (g == genes.size() || genes[g].chr != last_chr) {
        if (info.size()) {
          sort(info.begin(), info.end());
          GeneInfoInterval current_info{0, 0};
          std::vector<GeneInfoInterval> & chr_intervals{intervals[last_chr]};
          chr_intervals.push_back(current_info);
          for (const TempGeneInfo & this_info : info) {
            chr_intervals.back().stop_pos = this_info.pos;
            if (chr_intervals.back().size() == 0) {
              chr_intervals.pop_back();
            }
            current_info.info.erase(this_info.gene);
            current_info.start_pos = this_info.pos;
            switch (this_info.info.hit) {
              case HitType::none:
                break;
              case HitType::gene:
                current_info.info.emplace(this_info.gene,
                                          GeneInfo{HitType::gene, 0});
                break;
              case HitType::exon:
                current_info.info.emplace(
                    this_info.gene,
                    GeneInfo{HitType::exon, this_info.info.exon});
                break;
              case HitType::intron:
              default:
                throw Error("unexpected intron");
                break;
            }
            chr_intervals.push_back(current_info);
          }
          chr_intervals.back().stop_pos = genes.ref.size(last_chr);
          info.clear();
        }
        if (g == genes.size()) break;
      }
      const KnownGene & gene{genes[g]};
      last_chr = gene.chr;
      info.emplace_back(g, gene.t_start, GeneInfo{HitType::gene, 0});
      // info.emplace_back(g, gene.c_start, GeneInfo{HitType::intron, 0});
      // info.emplace_back(g, gene.c_stop, GeneInfo{HitType::gene, 0});
      for (unsigned int e{0}; e != gene.n_exons; ++e) {
        info.emplace_back(g, gene.exon_starts[e], GeneInfo{HitType::exon, e});
        info.emplace_back(g, gene.exon_stops[e], GeneInfo{HitType::gene, e});
      }
      info.emplace_back(g, gene.t_stop, GeneInfo{HitType::none, gene.n_exons});
    }
    for (unsigned int c{0}; c != genes.ref.n_chromosomes(); ++c) {
      if (intervals.count(c) == 0) {
        intervals[c].emplace_back(0, genes.ref.size(c));
      }
    }
  }
  const GeneInfoInterval &
  lookup(const unsigned int chr, const unsigned int pos) const {
    auto found = upper_bound(intervals.at(chr).begin(),
                             intervals.at(chr).end(),
                             pos,
                             [](const unsigned int p,
                                const GeneInfoInterval & i){
                               return p < i.stop_pos;
                             });
    if (found == intervals.at(chr).end())
      throw Error("Gene interval lookup error");
    return *found;
  }
  std::vector<const GeneInfoInterval *>
  lookup(const unsigned int chr,
         const unsigned int start_pos,
         const unsigned int stop_pos) const {
    auto start = upper_bound(intervals.at(chr).begin(),
                             intervals.at(chr).end(),
                             start_pos,
                             [](const unsigned int p,
                                const GeneInfoInterval & i){
                               return p < i.stop_pos;
                             });
    std::vector<const GeneInfoInterval *> result;
    while (start != intervals.at(chr).end()) {
      if (start->start_pos >= stop_pos) {
        break;
      }
      result.push_back(&*start);
      ++start;
    }
    return result;
  }

  std::map<unsigned int, std::vector<GeneInfoInterval>> intervals{};
};


}  // namespace paa


#endif  // PAA_GENES_H
```
