## Supplementary material for "G-Graph: An interactive genomic graph viewer": MUMdex Source Code for G-Graph: longSA.h

```
//
// longSA.h
//
// suffix array mapping
//
// Modifications of sparseMEM Copyright Peter Andrews 2013-2017 @ CSHL
//

#ifndef LONGMEM_LONGSA_H_
#define LONGMEM_LONGSA_H_

#include <limits.h>

#include <algorithm>
#include <deque>
#include <iostream>
#include <limits>
#include <sstream>
#include <string>
#include <utility>
#include <vector>

#include "error.h"
#include "fasta.h"
#include "files.h"
#include "mumdex.h"
#include "qsufsort.h"
#include "threads.h"
#include "utility.h"

namespace paa {

class SimpleHit {
 public:
  SimpleHit(const Sequence & ref, uint64_t rpos,
            const uint64_t off_, const uint64_t len_) :
      off(static_cast<unsigned int>(off_)),
      len(static_cast<unsigned int>(len_)) {
    bool flipped{false};
    if (rpos >= ref.N) {  // for flipped non rcref
      rpos -= ref.N;
      flipped = true;
    }
    auto chrit = upper_bound(ref.startpos.begin(), ref.startpos.end(), rpos);
    chr = static_cast<unsigned int>(chrit - ref.startpos.begin() - 1);
    pos = static_cast<unsigned int>(rpos - *--chrit);
    if (ref.rcref) {
      if ((chr % 2) == 1) {
        pos = static_cast<unsigned int>(ref.sizes[chr] - pos - len);
        flipped = true;
      }
      chr /= 2;
    }
    dir = (flipped ? '-' : '+');
  }
  std::ostream & out(std::ostream & stream) const {
    return stream << len << '\t'
                  << off << '\t'
                  << '('
                  << pos + 1 << ','
                  << chr << ','
                  << dir
                  << ')';
  }
  unsigned int chr{};
  unsigned int pos{};
  unsigned int off{};
  unsigned int len{};
  char dir{};
  bool operator==(const SimpleHit & rhs) const {
    return off == rhs.off && len == rhs.len &&
        pos == rhs.pos && chr == rhs.chr &&
        dir == rhs.dir;
  }
};
inline std::ostream & operator<<(std::ostream & out, const SimpleHit & hit) {
  return hit.out(out);
}

// Stores the LCP array in an unsigned char (0-255).  Values larger
// than or equal to 255 are stored in a sorted array.
// Simulates a vector<int> LCP;
template <class ANINT>
struct vec_uchar {
  struct item_t {
    item_t() {}
    item_t(ANINT i, ANINT v) {
      idx = i;
      val = v;
    }
    ANINT idx{};
    ANINT val{};
    bool operator < (const item_t & t) const { return idx < t.idx;  }
  };
  vec_uchar() : N_vec(0), vec(nullptr), cap_M(0), N_M(0), M(nullptr),
                using_mapping(false) {}
  ~vec_uchar() {
    if (memory_mapped && using_mapping) {
      if (munmap(vec, N_vec * sizeof(unsigned char)))
        std::cerr << "vec memory unmap failure" << std::endl;
      if (N_M)
        if (munmap(M, N_M * sizeof(item_t)))
          std::cerr << "M memory unmap failure" << std::endl;
    } else {
      free(vec);
      free(M);
    }
  }

  // Vector X[i] notation to get LCP values.
  ANINT operator[] (const ANINT idx) const {
    if (vec[idx] == std::numeric_limits<unsigned char>::max())
      return std::lower_bound(M, M + N_M, item_t(idx, 0))->val;
    else
      return vec[idx];
  }
  // Actually set LCP values, distingushes large and small LCP
  // values.
  void set(const ANINT idx, const ANINT v) {
    if (v >= std::numeric_limits<unsigned char>::max()) {
      vec[idx] = std::numeric_limits<unsigned char>::max();
      if (N_M == cap_M) {
        set_M_capacity(N_M * 2);
      }
      M[N_M++] = item_t(idx, v);
    } else {
      vec[idx] = static_cast<unsigned char>(v);
    }
  }

  void set_M_capacity(ANINT new_M) {
    if (new_M == 0) new_M = 1024;
    cap_M = new_M;
    if (cap_M > std::numeric_limits<ANINT>::max()) {
      throw Error("LCP M array too big");
    }
    if ((M = reinterpret_cast<item_t *>(realloc(M, sizeof(item_t) * cap_M)))
        == nullptr) throw Error("M realloc failed");
  }

  // Once all the values are set, call init. This will assure the
  // values >= 255 are sorted by index for fast retrieval.
  void init() {
    std::sort(M, M + N_M);
  }
  void load(const std::string & base, FILE * index) {
    using_mapping = true;
    bread(index, N_vec, "N_vec");
    bread(base + ".lcp.vec.bin", vec, "vec", N_vec);
    bread(index, N_M, "N_M");
    bread(base + ".lcp.m.bin", M, "M", N_M);
  }
  void save(const std::string & base, FILE * index) const {
    bwrite(index, N_vec, "N_vec");
    bwrite(base + ".lcp.vec.bin", vec[0], "vec", N_vec);
    bwrite(index, N_M, "N_M");
    bwrite(base + ".lcp.m.bin", M[0], "M", N_M);
  }
  void resize(const ANINT N) {
    N_vec = N;
    if ((vec = reinterpret_cast<unsigned char *>(
            malloc(sizeof(unsigned char) * N))) == nullptr)
      throw Error("malloc error for lcp");
  }
  uint8_t uchar(const ANINT idx) const {
    return vec[idx];
  }

  uint64_t bytes() const {
    return sizeof(vec_uchar) + N_vec + cap_M * sizeof(item_t);
  }

 private:
  ANINT N_vec;
  unsigned char * vec;
  uint64_t cap_M;
  ANINT N_M;
  item_t * M;
  bool using_mapping;
  vec_uchar(const vec_uchar &) = delete;
  vec_uchar & operator=(const vec_uchar &) = delete;
};

// depth : [start...end]
struct interval_t {
  interval_t() : depth(-1), start(1), end(0) { }
  interval_t(const uint64_t s, const uint64_t e,
             const uint64_t d) : depth(d), start(s), end(e) {}
  void reset(const uint64_t e) {
    depth = 0;
    start = 0;
    end = e;
  }
  uint64_t size() const { return end - start + 1; }
  uint64_t depth, start, end;
};

// Match find by find_mams.
struct match_t {
  match_t() : ref{0}, offset{0}, len{0} { }
  match_t(const uint64_t r, const unsigned int o, const unsigned int l) :
      ref{r}, offset{o}, len{l} { }
  bool operator<(const match_t & rhs) const {
    if (offset == rhs.offset) {
      return len > rhs.len;
    } else {
      return offset < rhs.offset;
    }
  }
  unsigned int rcoffset(const unsigned int read_length) const {
    return read_length - len - offset;
  }
  uint64_t ref;         // position in reference sequence
  unsigned int offset;  // position in query
  unsigned int len;     // length of match
};

class Args;
class SAArgs {
 public:
  SAArgs(const bool verbose_ = false, const bool mappability_ = false,
         const RefArgs ref_args_ = RefArgs(),
         const unsigned int mappability_threads_ = 4) :
      verbose(verbose_), mappability(mappability_), ref_args(ref_args_),
      mappability_threads(mappability_threads_) {}
  operator const RefArgs & () const { return ref_args; }
  bool verbose;
  bool mappability;
  RefArgs ref_args;
  unsigned int mappability_threads;

 private:
  SAArgs & operator=(const SAArgs &) = delete;
  friend class Args;
};

template <class ANINT>
struct longSA_t : public SAArgs {
  uint64_t size() const { return N; }

  bool using_mapping{};

  const Sequence ref;
  const uint64_t N{};  // !< Length of the sequence.

  const uint64_t logN{};  // ceil(log(N))
  const uint64_t Nm1{};  // N - 1

  ANINT * SA{};
  ANINT * ISA{};
  vec_uchar<ANINT> LCP{};  // Simulates a vector<int> LCP.

  // Constructor builds suffix array.
  void common_longSA_initialization();
  longSA_t(const std::string & fasta, const bool rcref, const bool mmap,
           const bool verbose_ = true) :
      SAArgs(verbose_, false, RefArgs(fasta.c_str(), rcref, verbose_)),
      using_mapping(mmap), ref(*this), N(ref.N),  // mmap = false may be best?
      logN(static_cast<uint64_t>(ceil(log(N) / log(2.0)))), Nm1(N - 1) {
    common_longSA_initialization();
  }
  explicit longSA_t(const SAArgs & arguments) :
      SAArgs(arguments), using_mapping(false),
      ref(arguments), N(ref.N),  // S(ref.seq),
      logN(static_cast<uint64_t>(ceil(log(N) / log(2.0)))), Nm1(N - 1) {
    common_longSA_initialization();
  }

  ~longSA_t() {
    if (memory_mapped && using_mapping) {
      if (munmap(SA, N * sizeof(ANINT)))
        std::cerr << "SA Memory unmap failure" << std::endl;
      if (munmap(ISA, N * sizeof(ANINT)))
        std::cerr << "ISA Memory unmap failure" << std::endl;
    } else {
      free(SA);
      free(ISA);
    }
  }

  // Modified Kasai et all for LCP computation.
  void computeLCP() {
    uint64_t h = 0;
    uint64_t n_high = 0;
    // First determine how many values are high
    for (uint64_t i = 0; i < N; ++i) {
      uint64_t m = ISA[i];
      if (m) {
        const uint64_t j = SA[m-1];
        while (i+h < N && j+h < N && ref[i+h] == ref[j+h]) ++h;
        if (h >= std::numeric_limits<unsigned char>::max()) {
          ++n_high;
        }
      }
      h = std::max(static_cast<int64_t>(0), static_cast<int64_t>(h - 1));
    }
    // Set M vector capacity to avoid overgrowing array
    LCP.set_M_capacity(static_cast<ANINT>(n_high));
    h = 0;
    // Fill LCP
    for (uint64_t i = 0; i < N; ++i) {
      uint64_t m = ISA[i];
      if (m == 0) {
        LCP.set(static_cast<ANINT>(m), 0);  // LCP[m] = 0;
      } else {
        const uint64_t j = SA[m-1];
        while (i+h < N && j+h < N && ref[i+h] == ref[j+h]) ++h;
        LCP.set(static_cast<ANINT>(m), static_cast<ANINT>(h));  // LCP[m] = h;
      }
      h = std::max(static_cast<int64_t>(0), static_cast<int64_t>(h - 1));
    }
  }

  // Simple suffix array search.
  bool search(const std::string &P, uint64_t &start, uint64_t &end) const {
    start = 0;
    end = N - 1;
    uint64_t i = 0;
    while (i < P.length()) {
      if (top_down_faster(P[i], i, start, end) == false) {
        return false;
      }
      ++i;
    }
    return true;
  }

  // Given SA interval apply binary search to match character c at
  // position i in the search string. Adapted from the C++ source code
  // for the wordSA implementation from the following paper: Ferragina
  // and Fischer. Suffix Arrays on Words. CPM 2007.
  inline bool top_down_faster(const char c, const uint64_t i,
                              uint64_t &start, uint64_t &end) const {
    uint64_t l, r, m, r2 = end, l2 = start;
    int64_t vgl;
    bool found = false;
    const int64_t cmp_with_first = static_cast<int64_t>(c) -
        static_cast<int64_t>(ref[SA[start]+i]);
    const int64_t cmp_with_last = static_cast<int64_t>(c) -
        static_cast<int64_t>(ref[SA[end]+i]);
    if (cmp_with_first < 0) {
      l = start + 1;
      l2 = start;  // pattern doesn't occur!
    } else if (cmp_with_last > 0) {
      l = end + 1;
      l2 = end;
      // pattern doesn't occur!
    } else {
      // search for left border:
      l = start;
      r = end;
      if (cmp_with_first == 0) {
        found = true;
        r2 = r;
      } else {
        while (r > l + 1) {
          m = (l+r) / 2;
          vgl = static_cast<int64_t>(c) - static_cast<int64_t>(ref[SA[m] + i]);
          if (vgl <= 0) {
            if (!found && vgl == 0) {
              found = true;
              l2 = m;
              r2 = r;  // search interval for right border
            }
            r = m;
          } else {
            l = m;
          }
        }
        l = r;
      }
      // search for right border (in the range [l2:r2])
      if (!found) {
        l2 = l - 1;  // pattern not found => right border to the left of 'l'
      }
      if (cmp_with_last == 0) {
        l2 = end;  // right border is the end of the array
      } else {
        while (r2 > l2 + 1) {
          m = (l2 + r2) / 2;
          vgl = static_cast<int64_t>(c) - static_cast<int64_t>(ref[SA[m] + i]);
          if (vgl < 0)
            r2 = m;
          else
            l2 = m;
        }
      }
    }
    start = l;
    end = l2;
    return l <= l2;
  }

  // Traverse pattern P starting from a given prefix and interval
  // until mismatch or min_len characters reached.
  inline void traverse(const std::string &P, const uint64_t prefix,
                       interval_t &cur, const ANINT min_len) const {
    if (cur.depth >= min_len) return;

    while (prefix+cur.depth < P.length()) {
      uint64_t start = cur.start;
      uint64_t end = cur.end;
      // If we reach a mismatch, stop.
      if (top_down_faster(P[prefix+cur.depth], cur.depth, start, end) == false)
        return;

      // Advance to next interval.
      cur.depth += 1;
      cur.start = start;
      cur.end = end;

      // If we reach min_len, stop.
      if (cur.depth == min_len) return;
    }
  }

  // Expand ISA/LCP interval. Used to simulate suffix links.
  inline bool expand_link(interval_t * link) const {
    const uint64_t thresh = 2 * link->depth * logN;
    uint64_t exp = 0;  // Threshold link expansion.
    uint64_t start = link->start;
    uint64_t end = link->end;
    while (LCP[static_cast<ANINT>(start)] >= link->depth) {
      if (++exp >= thresh) return false;
      --start;
    }
    while (end < Nm1 && LCP[static_cast<ANINT>(end + 1)] >= link->depth) {
      if (++exp >= thresh) return false;
      ++end;
    }
    link->start = start;
    link->end = end;
    return true;
  }

  std::vector<SimpleHit> find_mams(const std::string & query,
                                   const unsigned int min_len = 2) const {
    if (ref.rcref) {
      return find_mams_in_this_sa(query, min_len);
    } else {
      static thread_local std::vector<match_t> matches;
      matches.clear();
      find_mams(matches, query, min_len);
      std::vector<SimpleHit> mams;
      for (const match_t & match : matches) {
        mams.emplace_back(ref, match.ref, match.offset, match.len);
      }
      return mams;
    }
  }

  std::vector<SimpleHit> find_mams_in_this_sa(
      const std::string & query,
      const unsigned int min_len = 0) const {
    std::vector<SimpleHit> mams;
    interval_t cur(0, N - 1, 0);
    uint64_t prefix = 0;
    while (prefix < query.length()) {
      // Traverse SA top down until mismatch or full string is matched.
      traverse(query, prefix, cur, query.length());
      if (cur.depth <= 1) {
        cur.depth = 0;
        cur.start = 0;
        cur.end = N - 1;
        ++prefix;
        continue;
      }
      if (cur.size() == 1 && cur.depth >= min_len) {
        if (is_leftmaximal(query, prefix, SA[cur.start])) {
          mams.emplace_back(ref, SA[cur.start], prefix, cur.depth);
        }
      }
      do {
        cur.depth = cur.depth-1;
        cur.start = ISA[SA[cur.start] + 1];
        cur.end = ISA[SA[cur.end] + 1];
        ++prefix;
        if (cur.depth == 0 || expand_link(&cur) == false) {
          cur.depth = 0;
          cur.start = 0;
          cur.end = N - 1;
          break;
        }
      } while (cur.depth > 0 && cur.size() == 1);
    }
    return mams;
  }

  void find_mams(std::vector<match_t> & matches,
                 const std::string & query,
                 const unsigned int min_len = 2) const {
    find_mams_in_this_sa(matches, query, min_len);
    if (!ref.rcref) {
      const uint64_t regular_stop{matches.size()};
      static thread_local std::string rcquery;
      rcquery.assign(query);
      reverse_complement(&rcquery);
      find_mams_in_this_sa(matches, rcquery, min_len);
      for (uint64_t m{regular_stop}; m < matches.size(); ++m) {
        match_t & match{matches[m]};
        match.ref += ref.N;
        match.offset = match.rcoffset(static_cast<unsigned int>(query.size()));
      }
      if (regular_stop == 0) {
        reverse(matches.begin(), matches.end());
      } else {
        sort(matches.begin(), matches.end());
      }
      uint64_t nm{1};
      // Eliminate non-MUMs
      for (uint64_t m{0}; m != matches.size();) {
        match_t & match{matches[m]};
        if (match.offset == query.size()) {
          ++m;
          nm = m + 1;
          continue;
        }
        if (nm < matches.size()) {
          match_t & next_match{matches[nm]};
          if (next_match.offset == query.size()) {
            ++nm;
            continue;
          }
          if (match.offset == next_match.offset) {
            if (match.len == next_match.len) {
              match.offset = static_cast<unsigned int>(query.size());
              next_match.offset = static_cast<unsigned int>(query.size());
              ++m;
              nm = m + 1;
              continue;
            } else {
              next_match.offset = static_cast<unsigned int>(query.size());
              ++nm;
              continue;
            }
          } else if (next_match.offset < match.offset + match.len) {
            if (match.offset + match.len >=
                next_match.offset + next_match.len) {
              next_match.offset = static_cast<unsigned int>(query.size());
              ++nm;
              continue;
            } else {
              ++nm;
              continue;
            }
          }
        }
        static thread_local std::string reversed_mum;
        if (match.ref >= ref.N) {
          reversed_mum.assign(query, match.offset, match.len);
        } else {
          reversed_mum.assign(rcquery, match.rcoffset(static_cast<unsigned int>(
              query.size())), match.len);
        }
        if (n_forward_copies(reversed_mum)) {
          match.offset = static_cast<unsigned int>(query.size());
        }
        ++m;
        nm = m + 1;
      }
      sort(matches.begin(), matches.end());
      while (matches.size() && matches.back().offset == query.size()) {
        matches.pop_back();
      }
    }
  }

  // Maximal Almost-Unique Match (MAM). Match is unique in the indexed
  // sequence S. as computed by MUMmer version 2 by Salzberg
  // et. al. Note this is a "one-sided" query. It "streams" the query
  // P throught he index.  Consequently, repeats can occur in the
  // pattern P.
  // NOTE: min_len must be > 1
  void find_mams_in_this_sa(std::vector<match_t> & matches,
                            const std::string & query,
                            const unsigned int min_len = 2) const {
    interval_t cur(0, N - 1, 0);
    uint64_t prefix = 0;
    while (prefix < query.length()) {
      // Traverse SA top down until mismatch or full string is matched.
      traverse(query, prefix, cur, static_cast<unsigned int>(query.length()));
      if (cur.depth <= 1) {
        cur.depth = 0;
        cur.start = 0;
        cur.end = N - 1;
        ++prefix;
        continue;
      }
      if (cur.size() == 1 && cur.depth >= min_len) {
        if (is_leftmaximal(query, prefix, SA[cur.start])) {
          matches.emplace_back(SA[cur.start], prefix, cur.depth);
        }
      }
      do {
        cur.depth = cur.depth-1;
        cur.start = ISA[SA[cur.start] + 1];
        cur.end = ISA[SA[cur.end] + 1];
        ++prefix;
        if (cur.depth == 0 || expand_link(&cur) == false) {
          cur.depth = 0;
          cur.start = 0;
          cur.end = N - 1;
          break;
        }
      } while (cur.depth > 0 && cur.size() == 1);
    }
  }

  // Does not include reverse complement if not rcref
  unsigned int n_forward_copies(const std::string & query) const {
    interval_t cur(0, N - 1, 0);
    // Traverse SA top down until mismatch or full string is matched.
    traverse(query, 0, cur, static_cast<ANINT>(query.length()));
    if (cur.depth == query.length()) {
      return static_cast<unsigned int>(cur.size());
    }
    return 0;
  }
  unsigned int n_copies(const std::string & query) const {
    unsigned int result{n_forward_copies(query)};
    if (!ref.rcref) {
      std::string rcquery{query};
      reverse_complement(&rcquery);
      return result + n_forward_copies(rcquery);
    }
    return result;
  }
  bool exists(const std::string & query) const {
    unsigned int result{n_forward_copies(query)};
    if (result) return true;
    if (!ref.rcref) {
      std::string rcquery{query};
      reverse_complement(&rcquery);
      result = n_forward_copies(rcquery);
    }
    return result;
  }

  // Returns true if the position p1 in the query pattern and p2 in the
  // reference is left maximal.
  inline bool is_leftmaximal(const std::string &P, const uint64_t p1,
                             const uint64_t p2) const {
    if (p1 == 0 || p2 == 0)
      return true;
    else
      return P[p1-1] != ref[p2-1];
  }

  uint64_t bytes() const {
    return sizeof(longSA_t) + ref.bytes() - sizeof(Sequence) + LCP.bytes() -
        sizeof(vec_uchar<ANINT>) + 2 * N * sizeof(ANINT);
  }

 private:
  longSA_t(const longSA_t &) = delete;
  longSA_t & operator=(const longSA_t &) = delete;
};

using longSA = longSA_t<uint64_t>;
using shortSA = longSA_t<uint32_t>;

class MappabilityBuilder {
  // Definition of the mappability values
  // 255 - no uniqueness even if we go to the end of the chromosome
  // 254 - no uniqueness even if we go 253 positions left or right
  // 0 < n <= 253 uniquenes if we go n positions left or right
  // and no uniqueness if we go n-1
  // should never be '0'
  explicit MappabilityBuilder(const std::string & ref_fasta_,
                              const std::string save_name_ = "") :
      ref_fasta{ref_fasta_},
    save_name{save_name_.size() ? save_name_ : ref_fasta + ".bin/"},
    low_name{save_name + "map.low.bin"},
    high_name{save_name + "map.high.bin"} { }

 public:
  MappabilityBuilder(const longSA & sa, const unsigned int) :
      MappabilityBuilder{sa.ref.ref_fasta, ""} {
    ref_size = [&sa]() {
      uint64_t rsize = 0;
      for (unsigned int chr = 0; chr != sa.ref.sizes.size(); chr += 2) {
        rsize += sa.ref.sizes[chr];
      }
      return rsize;
    }();

    std::vector<uint8_t> low(ref_size);
    std::vector<uint8_t> high(ref_size);

    unsigned int iMp = 0;
    for (unsigned int iSA_Cp = 0; iSA_Cp != sa.ref.sizes.size(); iSA_Cp += 2) {
      const unsigned int chr_start = iMp;
      unsigned int iSA_Cn = iSA_Cp + 1;
      unsigned int chr_len = static_cast<unsigned int>(sa.ref.sizes[iSA_Cp]);

      uint64_t iSA_Rp = sa.ref.startpos[iSA_Cp];
      uint64_t iSA_Rn = sa.ref.startpos[iSA_Cn] + sa.ref.sizes[iSA_Cn] - 1;
      for (unsigned int p = 0; p != chr_len; ++p, ++iMp, ++iSA_Rp, --iSA_Rn) {
        const auto iSA_Sp = sa.ISA[iSA_Rp];
        const auto iSA_Sn = sa.ISA[iSA_Rn];

        if (sa.verbose && (iMp % 100000000) == 0) {
          std::cout << "MappabilityBuilder progress:" << iMp << "\n";
        }
        low[iMp] = std::min(std::max(sa.LCP.uchar(iSA_Sp),
                                     sa.LCP.uchar(iSA_Sp + 1)) + 1, 254);
        high[iMp] = std::min(std::max(sa.LCP.uchar(iSA_Sn),
                                      sa.LCP.uchar(iSA_Sn + 1)) + 1, 254);
      }
      for (unsigned int cp = 0; cp < std::min(chr_len, 253u); ++cp) {
        auto ap = chr_start + cp;
        if (high[ap] > cp + 1) {
          high[ap] = 255;
        }
      }
      for (unsigned int cp=0; cp < std::min(chr_len - 1, 253u); ++cp) {
        auto ap = chr_start + chr_len - cp - 1;
        if (low[ap] > cp + 1) {
          low[ap] = 255;
        }
      }
    }
    bwrite(low_name, low[0], "low", ref_size);
    bwrite(high_name, high[0], "high", ref_size);
  }

  MappabilityBuilder(const shortSA & sa,
                     const unsigned int n_threads = 12,
                     const std::string save_name_ = "",
                     const void * mappability = nullptr) :
      MappabilityBuilder{sa.ref.ref_fasta, save_name_} {
    using FileInfo = std::pair<unsigned int, std::string>;
    auto mappability_block = [this, &sa](
        const Mappability * map,
        const unsigned int iSA_Cp,
        unsigned int p_start,
        unsigned int p_stop,
        const unsigned int chr_len,
        unsigned int iMp,
        uint64_t iSA_Rp,
        uint64_t & completed) {
      static thread_local std::vector<uint8_t> low(p_stop - p_start);
      low.clear();
      low.resize(p_stop - p_start);
      static thread_local std::vector<uint8_t> high(p_stop - p_start);
      high.clear();
      high.resize(p_stop - p_start);
      iMp = 0;
      for (unsigned int p{p_start}; p != p_stop; ++p, ++iMp, ++iSA_Rp) {
        const auto iSA_Sp = sa.ISA[iSA_Rp];
        low[iMp] = std::min(std::max(sa.LCP.uchar(iSA_Sp),
                                     sa.LCP.uchar(iSA_Sp + 1)) + 1, 254);
        if (p != p_start && low[iMp - 1] < 254)
          low[iMp] = std::max(low[iMp],
                               static_cast<uint8_t>(low[iMp - 1] - 1));
        do {
          if (low[iMp] >= 254 || p + low[iMp] > chr_len) {
            break;
          }
          const auto seq_start = sa.ref.seq + sa.ref.startpos[iSA_Cp] + p;
          std::string to_test(seq_start, seq_start + low[iMp]);
          reverse_complement(&to_test);
          if (!sa.n_forward_copies(to_test)) {
            break;
          }
          ++low[iMp];
        } while (true);
        const unsigned int end_dist{chr_len - p};
        if (end_dist <= 253 && low[iMp] > end_dist) {
          low[iMp] = 255;
        }
        if (low[iMp] <= 254 && p + low[iMp] - 1 < p_stop)
          high[iMp + low[iMp] - 1] = low[iMp];
        if (iMp) {
          if (high[iMp]) {
            high[iMp] = std::min(high[iMp],
                                 static_cast<uint8_t>(high[iMp - 1] + 1));
          } else {
            high[iMp] = high[iMp - 1] + 1;
          }
        } else {
          high[iMp] = 254;
        }
        if (p == p_start || high[iMp] == 255 || high[iMp] == 0) {
          high[iMp] = 254;
        }
        do {
          if (high[iMp] == 1) break;
          const unsigned int trial_length{
            std::min(p + 1, static_cast<unsigned int>(high[iMp] - 1))};
          const char * seq_stop{
            sa.ref.seq + sa.ref.startpos[iSA_Cp] + p + 1};
          const char * seq_start{seq_stop - trial_length};
          std::string to_test(seq_start, seq_stop);
          std::string orig{to_test};
          const unsigned int n_normal{sa.n_forward_copies(to_test)};
          if (n_normal > 1) {
            break;
          }
          reverse_complement(&to_test);
          const unsigned int n_reverse{sa.n_forward_copies(to_test)};
          if (n_normal + n_reverse < 1) {
            throw Error("No copies found - strange")
                << trial_length << " " << to_test.size() << " " << p
                << " '" << to_test << "'" << " '" << orig << "'";
          }
          if (n_normal + n_reverse > 1) {
            break;
          }
          --high[iMp];
        } while (true);
        if (p <= 253 && high[iMp] > p + 1) {
          high[iMp] = 255;
        }
        if (map) {
          if (map->low(iMp) != low[iMp]) {
            throw Error("Low mappability mismatch")
            << sa.ref.descr[iSA_Cp] << " " << p << " "
            << chr_len - p - 1 << " " << chr_len << " "
            << map->low(iMp) << " "
            << static_cast<unsigned int>(low[iMp]);
          }
          if (map->high(iMp) != high[iMp]) {
            throw Error("High mappability mismatch")
            << sa.ref.descr[iSA_Cp] << " " << p << " "
            << chr_len - p - 1 << " " << chr_len << " "
            << map->high(iMp) << " "
            << static_cast<unsigned int>(high[iMp]);
          }
        }
      }

      const std::string file_base{ref_fasta + ".bin/map_temp/map." +
                                  std::to_string(iSA_Rp)};
      bwrite(file_base + ".low.bin", low[0], "low", low.size());
      bwrite(file_base + ".high.bin", high[0], "high", high.size());

      static std::mutex completed_mutex;
      std::unique_lock<std::mutex> completed_guard{completed_mutex};
      completed += p_stop - p_start;
      return FileInfo{iSA_Rp, file_base};
    };

    const Mappability * map{
      reinterpret_cast<const Mappability *>(mappability)};
    ref_size = [&sa]() {
      uint64_t rsize = 0;
      for (unsigned int chr = 0; chr != sa.ref.sizes.size(); ++chr) {
        rsize += sa.ref.sizes[chr];
      }
      return rsize;
    }();

    ThreadPool pool{n_threads};
    ThreadPool::Results<FileInfo> results;

    const std::string temp_dir{ref_fasta + ".bin/map_temp/"};
    mkdir(temp_dir);

    unsigned int iMp = 0;
    const bool do_jump{false};
    const time_t start_time{time(nullptr)};
    uint64_t positions_to_do{0};
    uint64_t positions_completed{0};
    for (unsigned int iSA_Cp = 0; iSA_Cp != sa.ref.sizes.size(); ++iSA_Cp) {
      unsigned int chr_len = static_cast<unsigned int>(sa.ref.sizes[iSA_Cp]);

      uint64_t iSA_Rp = sa.ref.startpos[iSA_Cp];
      for (unsigned int p = 0; p != chr_len;) {
        const unsigned int this_block_size{p + block_size < chr_len ?
              block_size : chr_len - p};

        pool.run(results, mappability_block,
                 map, iSA_Cp, p, p + this_block_size,
                 chr_len, iMp, iSA_Rp,
                 std::ref(positions_completed));
        positions_to_do += this_block_size;
        const uint64_t jump_dist{(do_jump && block_size * 2 + 100 < chr_len) ?
              (p ? this_block_size : chr_len - this_block_size) :
              this_block_size};
        iMp += jump_dist;
        iSA_Rp += jump_dist;
        p += jump_dist;
      }
    }
    std::cerr << "waiting for " << results.size()
              << " tasks to construct mappability values" << std::endl;

    time_t stop_time{time(nullptr)};
    const char csi[]{27, '[', 0};

    std::deque<FileInfo> to_concatenate;
    while (results.size()) {
      to_concatenate.emplace_back(results.get());
      stop_time = time(nullptr);
      const int64_t elapsed{stop_time - start_time};
      const double remaining{1.0 * (positions_to_do - positions_completed) *
            elapsed / positions_completed};
      std::cout << '\r'
                << 100.0 * positions_completed / positions_to_do
                << "% positions done "
                << (stop_time - start_time) / 3600.0 << " hours elapsed "
                << remaining / 3600 << " hours to go "
                << csi << "K"
                << std::flush;
    }
    std::cout << '\r' << "100% done. Elapsed "
              << (stop_time - start_time) / 3600.0 << " hours"
              << csi << "K" << std::endl;

    sort(to_concatenate.begin(), to_concatenate.end(),
         [](const FileInfo & lhs, const FileInfo & rhs) {
           return lhs.first < rhs.first;
         });
    FILE * low_file = fopen(low_name.c_str(), "wb");
    if (low_file == nullptr) {
      throw Error("could not open low mappability file")
          << low_name << "for writing";
    }
    FILE * high_file = fopen(high_name.c_str(), "wb");
    if (high_file == nullptr) {
      throw Error("could not open high mappability file")
          << high_name << "for writing";
    }
    Progress progress(to_concatenate.size(), 0.1,
                           "Mappability file concatenation");
    while (to_concatenate.size()) {
      progress();
      const FileInfo & file_info{to_concatenate.front()};
      {
        const std::string low_name_{file_info.second + ".low.bin"};
        const MappedVector<char> low{low_name_};
        bwrite(low_file, *low.begin(), "low", low.size());
        unlink(low_name_);
      }
      const std::string high_name_{file_info.second + ".high.bin"};
      const MappedVector<char> high{high_name_};
      bwrite(high_file, *high.begin(), "high", high.size());
      unlink(high_name_);
      to_concatenate.pop_front();
    }
    fclose(low_file);
    fclose(high_file);
    rmdir(temp_dir.c_str());
  }
  uint64_t size() const { return ref_size; }

 private:
  const unsigned int block_size{100000};
  const std::string ref_fasta{};
  const std::string save_name{};
  const std::string low_name{};
  const std::string high_name{};
  uint64_t ref_size{};
};

template <class ANINT>
void longSA_t<ANINT>::common_longSA_initialization() {
  const time_t start_time = time(nullptr);

  // Index cache filename
  std::ostringstream saved_index_stream;
  saved_index_stream << ref.ref_fasta << ".bin";
  saved_index_stream << "/rc" << ref.rcref;
  saved_index_stream << ".i" << sizeof(ANINT) << ".index";
  const std::string bin_base = saved_index_stream.str();
  saved_index_stream << ".bin";
  const std::string saved_index = saved_index_stream.str();

  const uint64_t fasta_size = file_size(ref.ref_fasta);

  // Load or create index
  if (readable(saved_index)) {
    if (verbose) std::cerr << "loading index binary" << std::endl;

    FILE * index = fopen(saved_index.c_str(), "rb");
    if (index == nullptr) {
      throw Error("could not open index")
          << saved_index << "for reading";
    }

    uint64_t fasta_saved_size;
    bread(index, fasta_saved_size, "fasta_size");
    if (fasta_size != fasta_saved_size) {
      throw Error("saved fasta size used for index does not"
                       "match current fasta size\n"
                       "maybe the reference has changed?\n"
                       "you may need to delete the current index to proceed");
    }
    uint64_t dummy;
    bread(index, dummy, "logN");
    bread(index, dummy, "logN");
    uint64_t SA_size;
    bread(index, SA_size, "SA_size");

    using_mapping = true;
    bread(bin_base + ".sa.bin", SA, "SA", SA_size);
    bread(bin_base + ".isa.bin", ISA, "ISA", SA_size);
    LCP.load(bin_base, index);
    if (fclose(index) != 0) throw Error("problem closing index file");
  } else {
    if (verbose) std::cerr << "creating index from reference" << std::endl;

    if ((SA = reinterpret_cast<ANINT *>(malloc(sizeof(ANINT) * N))) ==
        nullptr) throw Error("SA malloc error");
    if ((ISA = reinterpret_cast<ANINT *>(malloc(sizeof(ANINT) * N))) ==
        nullptr) throw Error("ISA malloc error");

    ANINT char2int[UCHAR_MAX + 1];  // Map from char to integer alphabet.

    // Zero char2int mapping.
    for (ANINT i = 0; i != UCHAR_MAX; ++i) char2int[i] = 0;

    // Determine which characters are used in the string S.
    for (ANINT i = 0; i != N; ++i) char2int[static_cast<uint8_t>(ref[i])] = 1;

    // Count the size of the alphabet.
    ANINT alphasz = 0;
    for (ANINT i = 0; i != UCHAR_MAX; ++i) {
      if (char2int[i])
        char2int[i] = alphasz++;
      else
        char2int[i] = static_cast<ANINT>(-1);
    }

    // Remap the alphabet.
    for (ANINT i = 0; i != N; ++i) {
      ISA[i] = char2int[static_cast<uint8_t>(ref[i])] + 1;
    }
    // First "character" equals 1 because of above plus one,
    // l=1 in suffixsort().
    const ANINT alphalast = alphasz + 1;

    // Use LS algorithm to construct the suffix array.
    const suffixsort<ANINT> sorter{&ISA[0], &SA[0],
          N - 1, alphalast, 1, verbose};

    // Use algorithm by Kasai et al to construct LCP array.
    if (verbose) std::cerr << "creating LCP array" << std::endl;
    LCP.resize(static_cast<ANINT>(N));
    computeLCP();  // SA + ISA -> LCP
    LCP.init();

    if (verbose) std::cerr << "saving index" << std::endl;

    FILE * index = fopen(saved_index.c_str(), "wb");
    if (index == nullptr)
      throw Error("could not open index")
          << saved_index << "for writing";
    bwrite(index, fasta_size, "fasta_size");
    bwrite(index, logN, "logN");
    bwrite(index, Nm1, "Nm1");
    bwrite(index, N, "SA_size");
    bwrite(bin_base + ".sa.bin", SA[0], "SA", N);
    bwrite(bin_base + ".isa.bin", ISA[0], "ISA", N);
    LCP.save(bin_base, index);
    if (fclose(index) != 0) throw Error("problem closing index file");

    // Create mappability binary output only if it does not yet exist
    if (!readable(std::string() + ref.ref_fasta + ".bin/map.low.bin")) {
      MappabilityBuilder map(*this, mappability_threads);
    }
  }

  if (verbose) {
    const time_t end_time = time(nullptr);
    std::cerr << "constructed index in "
              << end_time - start_time << " seconds" << std::endl;
  }
}

}  // namespace paa

#endif  // LONGMEM_LONGSA_H_
```
