## Supplementary material for "G-Graph: An interactive genomic graph viewer": MUMdex Source Code for G-Graph: mapper.h

```
//
// mapper.h
//
// MUM based mapping
//
// Copyright Peter Andrews 2015 @ CSHL
//

#ifndef MAPPER_H_
#define MAPPER_H_

#include <algorithm>
#include <string>
#include <queue>
#include <utility>
#include <vector>

#include "encode.h"
#include "error.h"
#include "longSA.h"
#include "mumdex.h"
#include "pstream.h"
#include "query.h"
#include "sort.h"
#include "threads.h"

namespace paa {

class PairMergeHelper {
 public:
  explicit PairMergeHelper(const MUMdex & mumdex_arg,
                           const unsigned int id__) :
      mumdex_(&mumdex_arg), id_{id__} { }
  bool operator<(const PairMergeHelper & right_) const {
    const Pair left{pair()};
    const Pair right{right_.pair()};
    const MUM * left_mums{mumdex_->mums_begin()};
    const MUM * right_mums{right_.mumdex_->mums_begin()};
    return right.lessPair(right_mums, left, left_mums);
  }
  PairMergeHelper & operator++() {
    ++n_;
    return *this;
  }
  operator bool() const { return n_ != mumdex_->n_pairs(); }
  Pair pair() const { return mumdex_->pair(n_); }
  const MUMdex & mumdex() const { return *mumdex_; }
  uint64_t n() const { return n_; }
  unsigned int id() const { return id_; }

 private:
  const MUMdex * mumdex_;
  uint64_t n_{0};
  unsigned int id_;
};

class IndexMergeHelper {
 public:
  explicit IndexMergeHelper(const MappedVector<TempMUMindex> & index_arg) :
      index_(&index_arg) { }
  bool operator<(const IndexMergeHelper & right_) const {
    return right_.index() < index();
  }
  IndexMergeHelper & operator++() {
    ++n_;
    return *this;
  }
  operator bool() const { return n_ != index_->size(); }
  TempMUMindex index() const { return (*index_)[n_]; }
  uint64_t n() const { return n_; }

 private:
  const MappedVector<TempMUMindex> * index_;
  uint64_t n_{0};
};

std::pair<int, void *> mmap_create_file(const std::string & file_name,
                                              const uint64_t size) {
  const int file{open(file_name.c_str(), O_RDWR | O_CREAT | O_TRUNC,
                      S_IRUSR | S_IWUSR | S_IRGRP | S_IROTH)};
  if (file == -1) {
    perror("open error");
    throw Error("Problem opening index file for memory mapped i/o")
        << file_name;
  }
  if (lseek(file, size - 1, SEEK_SET) == static_cast<off_t>(-1)) {
    perror("seek end error");
    throw Error("lseek to end error in") << file_name;
  }
  if (::write(file, "", 1) < 1) {
    perror("write error");
    throw Error("Problem extending index file") << file_name;
  }
  if (lseek(file, 0, SEEK_SET) == static_cast<off_t>(-1)) {
    perror("seek begin error");
    throw Error("lseek to origin error in") << file_name;
  }
  void * memory{mmap(nullptr, size, PROT_READ | PROT_WRITE,
                     MAP_SHARED, file, 0)};
  if (memory == MAP_FAILED) {
    perror("mmap error");
    throw Error("Memory mapping failed for file") << file_name;
  }
  return std::make_pair(file, memory);
}

#if 0
class MUMdexMerger {
 public:
  explicit MUMdexMerger(const std::string & mumdex_name,
                        const unsigned int n_threads,
                        const bool mark_dupes) {
    // Get mumdex part names
    const std::vector<std::string> part_names{[&mumdex_name]() {
        redi::ipstream part_names_process{
          std::string("ls -d ") + mumdex_name + "/part.*"};
        if (!part_names_process)
          throw Error("Could not open process to read part names in")
              << mumdex_name;
        std::vector<std::string> names;
        std::string name;
        while (getline(part_names_process, name)) {
          names.push_back(name);
        }
        if (names.empty()) throw Error("No mumdex parts found");
        return names;
      }()};

    std::cerr << "Merging " << part_names.size() << " index parts" << std::endl;
    const time_t start_time{time(nullptr)};

    {  // In a block so mumdexs are freed before directories are removed
      // Load mumdex parts
      const std::string reference_name{saved_ref_name(part_names.front())};
      const Reference ref{reference_name};
      const std::vector<MUMdex> mumdexs{[&part_names, &ref]() {
          std::vector<MUMdex> mumdexs_;
          mumdexs_.reserve(part_names.size());
          for (const std::string & name : part_names) {
            mumdexs_.emplace_back(name, ref);
          }
          return mumdexs_;
        }()};

      // Load format strings describing optional metadata passthroughs
      const std::vector<std::string>
          optional_formats{read_optional_formats(mumdex_name)};

      // Load optional savers
      OptionalSavers saver{optional_formats};
      const std::vector<OptionalSavers> savers{
        [&optional_formats, &mumdexs, &part_names]() {
          std::vector<OptionalSavers> savers_;
          savers_.reserve(part_names.size());
          for (unsigned int f{0}; f != mumdexs.size(); ++f) {
            const std::string & name{part_names[f]};
            savers_.emplace_back(optional_formats);
            savers_.back().load(name, mumdexs[f].n_pairs() * 2);
          }
          return savers_;
        }()};

      // Output files for optional info
      const std::vector<FILE *> optional_files{
        [&mumdex_name, &savers]() {
          std::vector<FILE *> optional_files_;
          if (savers.front().size() == 0) return optional_files_;
          optional_files_.reserve(savers.front().size());
          for (unsigned int s{0}; s != savers.front().size(); ++s) {
            const std::string file_name{
              savers.front()[s].file_name(mumdex_name)};
            std::FILE * file{std::fopen(file_name.c_str(), "wb+")};
            if (file == nullptr) throw Error("Problem opening file")
                                     << file_name;
            optional_files_.push_back(file);
          }
          return optional_files_;
        }()};

      // How many mums total?
      const uint64_t n_mums{[&mumdexs]() {
          uint64_t n{0};
          for (const MUMdex & mumdex : mumdexs) {
            n += mumdex.n_mums();
          }
          return n;
        }()};

      // Get memory requirements for index sorting
      const uint64_t sort_size{n_mums * sizeof(TempMUMindex)};

      // Open memory mapped file for index read/write and expand to size
      const std::string sortName{index_name(mumdex_name) + ".temp"};
      std::pair<int, void *> sort_map{mmap_create_file(sortName, sort_size)};

      // Set up pointers for sorting of index
      TempMUMindex * const sort_begin =
          reinterpret_cast<TempMUMindex *>(sort_map.second);
      TempMUMindex * const sort_end{sort_begin + n_mums};
      TempMUMindex * sort_ptr{sort_begin};

      // Prepare to populate index for sorting
      sequential(sort_begin, sort_end);

      // Merge and mark duplicates and prepare index for sort
      MUMdex_build_base all;
      all.save_reference_name(mumdex_name, reference_name);
      std::vector<std::FILE *> files{all.open_for_write(mumdex_name)};
      const unsigned int max_n{1000000};
      uint64_t n_pairs{0};
      uint64_t mums_offset{0};
      uint64_t extra_bases_offset{0};
      const MUMdex * last_mumdex{nullptr};
      uint64_t last_p{0};
      std::priority_queue<PairMergeHelper> merge_helpers{[&mumdexs]() {
          std::priority_queue<PairMergeHelper> helpers;
          for (unsigned int f{0}; f != mumdexs.size(); ++f) {
            const MUMdex & mumdex{mumdexs[f]};
            helpers.emplace(mumdex, f);
          }
          return helpers;
        }()};
      while (merge_helpers.size()) {
        PairMergeHelper top{merge_helpers.top()};
        const MUMdex & mumdex{top.mumdex()};
        const uint64_t n{top.n()};
        const unsigned int id{top.id()};
        all.emplace_back(mumdex, n, mums_offset, extra_bases_offset,
                         last_mumdex, last_p, mark_dupes);
        saver.copy(savers[id], 2 * n);
        saver.copy(savers[id], 2 * n + 1);
        for (unsigned int m{0}; m != mumdex.n_mums(n); ++m) {
          *(sort_ptr++) = TempMUMindex{
            MUMindex(n_pairs, m), mumdex.mum(mumdex.pair(n).mums_start() + m)};
        }
        merge_helpers.pop();
        if (++top) merge_helpers.push(top);
        if ((++n_pairs % max_n) == 0 || merge_helpers.empty()) {
          saver.write_and_reduce(optional_files, merge_helpers.empty());
          const ExtraBasesSaveInfo info{all.write(files)};
          mums_offset += all.n_mums();
          all.clear();
          extra_bases_offset += info.n_saved_bases();
          const std::vector<uint64_t> & last_words{info.last_words()};
          if (last_words.size()) {
            if (merge_helpers.empty()) {
              bwritec(files[3], &last_words[0], "Last Words",
                      last_words.size() * sizeof(uint64_t));
            } else {
              all.bases().extra().add_excess_bases(info);
            }
          }
        }
      }
      all.close_files(files);
      for (FILE * file : optional_files) {
        if (fclose(file)) {
          throw Error("problem closing optional file");
        }
      }
      const time_t sort_start_time{time(nullptr)};

      // Prepare index output file
      const std::string indexName{index_name(mumdex_name)};
      const uint64_t index_size{n_mums * sizeof(MUMindex)};
      std::pair<int, void *> index_map{mmap_create_file(indexName, index_size)};
      MUMindex * const index_begin{
        reinterpret_cast<MUMindex *>(index_map.second)};
      MUMindex * const index_end{index_begin + n_mums};

      // avoids unused variable warnings for alternate compilation paths
      if (0) std::cerr << n_threads << index_end;

      // Sort index and output
      random(sort_begin, sort_end);
#ifdef SPLIT___
      TempMUMindex * const  middle{sort_begin + (sort_end - sort_begin) / 2};
      std::sort(sort_begin, middle);
      std::sort(middle, sort_end);
      sequential(sort_begin, sort_end);
      sequential(index_begin, index_end);
      std::merge(sort_begin, middle, middle, sort_end, index_begin);
#else
#ifdef PARALLEL
      ParallelSortMove(sort_begin, sort_end, index_begin, n_threads);
#else
      PeterSort(sort_begin, sort_end, index_begin);
#endif
#endif

      // Index sort clean up
      if (false && munmap(sort_map.second, sort_size)) {
        perror("sort unmap error");
        throw Error("Memory unmap map failed for file") << sortName;
      }
      if (close(sort_map.first) == -1) {
        perror("close error");
        throw Error("File close error for") << sortName;
      }
      unlink(sortName);

      if (msync(index_map.second, index_size, MS_SYNC) == -1) {
        perror("msync error");
        throw Error("Memory sync failure for file") << indexName;
      }
      if (false && munmap(index_map.second, index_size) == -1) {
        perror("munmap error");
        throw Error("Memory unmap failure for file") << indexName;
      }
      if (close(index_map.first) == -1) {
        perror("close error");
        throw Error("File close error for") << indexName;
      }

      const time_t stop_time{time(nullptr)};
      std::cerr << "Merge took " << sort_start_time - start_time
                << " seconds and sort took "
                << stop_time - sort_start_time << " seconds." << std::endl;
      // Test sort
#if 0
      const MUMdex test{mumdex_name};
      if (!std::is_sorted(test.index().begin(), test.index().end(),
                          lessMUM_index<UnMappedVector>(
                              test.mums(), test.pairs()))) {
        throw Error("Not sorted");
      } else {
        std::cerr << "Sort ok" << std::endl;
      }
#endif
    }

    // Remove parts and change mumdex to write protected for safety
    if (false) {
      for (const std::string & part : part_names) {
        if (system((std::string("rm -R ") + part).c_str()))
          throw Error("Problem removing mumdex parts");
      }
      if (system((std::string("chmod a-w ") + mumdex_name + "/*.bin").c_str()))
          throw Error("Problem making mumdex write protected");
    }
  }
};
#endif

class MUMdexMergerNew {
 public:
  explicit MUMdexMergerNew(const std::string & mumdex_name,
                           const unsigned int n_gb,
                           const unsigned int n_threads,
                           const bool mark_dupes) {
    // Get mumdex part names
    const std::vector<std::string> part_names{[&mumdex_name]() {
        redi::ipstream part_names_process{
          std::string("ls -d ") + mumdex_name + "/part.*"};
        if (!part_names_process)
          throw Error("Could not open process to read part names in")
              << mumdex_name;
        std::vector<std::string> names;
        std::string name;
        while (getline(part_names_process, name)) {
          names.push_back(name);
        }
        if (names.empty()) throw Error("No mumdex parts found");
        return names;
      }()};

    std::cerr << "Merging " << part_names.size() << " index parts" << std::endl;
    const time_t start_time{time(nullptr)};
    std::vector<std::string> sort_files;

    {  // In a block so mumdexes are freed before directories are removed
      // Load mumdex parts
      const std::string reference_name{saved_ref_name(part_names.front())};
      const Reference ref{reference_name};
      const std::vector<MUMdex> mumdexs{[&part_names, &ref]() {
          std::vector<MUMdex> mumdexs_;
          mumdexs_.reserve(part_names.size());
          for (const std::string & name : part_names) {
            mumdexs_.emplace_back(name, ref);
          }
          return mumdexs_;
        }()};

      // Load format strings describing optional metadata passthroughs
      const std::vector<std::string>
          optional_formats{read_optional_formats(mumdex_name)};

      // Load optional savers
      OptionalSavers saver{optional_formats};
      const std::vector<OptionalSavers> savers{
        [&optional_formats, &mumdexs, &part_names]() {
          std::vector<OptionalSavers> savers_;
          savers_.reserve(part_names.size());
          for (unsigned int f{0}; f != mumdexs.size(); ++f) {
            const std::string & name{part_names[f]};
            savers_.emplace_back(optional_formats);
            savers_.back().load(name, mumdexs[f].n_pairs() * 2);
          }
          return savers_;
        }()};

      // Output files for optional info
      const std::vector<FILE *> optional_files{
        [&mumdex_name, &savers]() {
          std::vector<FILE *> optional_files_;
          if (savers.front().size() == 0) return optional_files_;
          optional_files_.reserve(savers.front().size());
          for (unsigned int s{0}; s != savers.front().size(); ++s) {
            const std::string file_name{
              savers.front()[s].file_name(mumdex_name)};
            std::FILE * file{std::fopen(file_name.c_str(), "wb+")};
            if (file == nullptr) throw Error("Problem opening file")
                                     << file_name;
            optional_files_.push_back(file);
          }
          return optional_files_;
        }()};

      // How many mums total?
#if 0
      const uint64_t n_mums{[&mumdexs]() {
          uint64_t n{0};
          for (const MUMdex & mumdex : mumdexs) {
            n += mumdex.n_mums();
          }
          return n;
        }()};
#endif
      // Sort vectors and threads
      const uint64_t gb{1024*1024*1024};
      const uint64_t per_sort_bytes{n_gb * gb / n_threads};
      const uint64_t per_sort_elements{per_sort_bytes / sizeof(TempMUMindex)};
      std::vector<std::vector<TempMUMindex>> sort_vectors(n_threads);
      for (std::vector<TempMUMindex> & vec : sort_vectors) {
        vec.reserve(per_sort_elements);
      }
      unsigned int current_sorter{0};
      unsigned int sort_id{0};
      ThreadPool pool{n_threads};
      ThreadPool::Results<std::pair<unsigned int, std::string>> sorters;

      // Merge and mark duplicates and start sort of index
      MUMdex_build_base all;
      all.save_reference_name(mumdex_name, reference_name);
      std::vector<std::FILE *> files{all.open_for_write(mumdex_name)};
      const unsigned int max_n{1000000};
      uint64_t n_pairs{0};
      uint64_t mums_offset{0};
      uint64_t extra_bases_offset{0};
      const MUMdex * last_mumdex{nullptr};
      uint64_t last_p{0};
      std::priority_queue<PairMergeHelper> merge_helpers{[&mumdexs]() {
          std::priority_queue<PairMergeHelper> helpers;
          for (unsigned int f{0}; f != mumdexs.size(); ++f) {
            const MUMdex & mumdex{mumdexs[f]};
            helpers.emplace(mumdex, f);
          }
          return helpers;
        }()};
      while (merge_helpers.size()) {
        PairMergeHelper top{merge_helpers.top()};
        const MUMdex & mumdex{top.mumdex()};
        const uint64_t n{top.n()};
        const unsigned int id{top.id()};
        all.emplace_back(mumdex, n, mums_offset, extra_bases_offset,
                         last_mumdex, last_p, mark_dupes);
        saver.copy(savers[id], 2 * n);
        saver.copy(savers[id], 2 * n + 1);
        merge_helpers.pop();
        if (++top) merge_helpers.push(top);

        for (unsigned int m{0}; m != mumdex.n_mums(n); ++m) {
          sort_vectors[current_sorter].emplace_back(
              MUMindex(n_pairs, m),
              mumdex.mum(mumdex.pair(n).mums_start() + m));
        }

        // Flush sort vector, start new one
        if (sort_vectors[current_sorter].size() + 150 >=
            per_sort_elements || merge_helpers.empty()) {
          pool.run(sorters, [&sort_vectors, &mumdex_name]
                   (const unsigned int current, const unsigned int sortid) {
                     std::vector<TempMUMindex> & sorter{sort_vectors[current]};
                     sort(sorter.begin(), sorter.end());
                     std::ostringstream out_name;
                     out_name << mumdex_name << "/index.sorted." << sortid
                              << ".bin";
                     bwrite(out_name.str(), sorter[0], "Temporary Index",
                            sorter.size());
                     sorter.clear();
                     return std::pair<unsigned int, std::string>{
                       current, out_name.str()};
                   }, current_sorter, ++sort_id);
          if (sorters.size() == n_threads) {
            const std::pair<unsigned int, std::string> result{sorters.get()};
            current_sorter = result.first;
            sort_files.push_back(result.second);
          } else {
            ++current_sorter;
          }
        }
        if ((++n_pairs % max_n) == 0 || merge_helpers.empty()) {
          saver.write_and_reduce(optional_files, merge_helpers.empty());
          const ExtraBasesSaveInfo info{all.write(files)};
          mums_offset += all.n_mums();
          all.clear();
          extra_bases_offset += info.n_saved_bases();
          const std::vector<uint64_t> & last_words{info.last_words()};
          if (last_words.size()) {
            if (merge_helpers.empty()) {
              bwritec(files[3], &last_words[0], "Last Words",
                      last_words.size() * sizeof(uint64_t));
            } else {
              all.bases().extra().add_excess_bases(info);
            }
          }
        }
      }
      all.close_files(files);
      for (FILE * file : optional_files) {
        if (fclose(file)) {
          throw Error("problem closing optional file");
        }
      }

      // Wait for all sort jobs to finish
      while (sorters.size()) {
        const std::pair<unsigned int, std::string> result{sorters.get()};
        sort_files.push_back(result.second);
      }
      // Clear sort memory
      for (std::vector<TempMUMindex> & vec : sort_vectors) {
        std::vector<TempMUMindex> empty;
        empty.swap(vec);
      }

      // Merge sort files
      const time_t sort_start_time{time(nullptr)};
      const std::vector<MappedVector<TempMUMindex>> sorted_indexes{
        [&sort_files]() {
          std::vector<MappedVector<TempMUMindex> > result;
          result.reserve(sort_files.size());
          for (const std::string & file : sort_files) {
            result.emplace_back(file);
          }
          return result;
        }()};

      std::priority_queue<IndexMergeHelper> index_mergers{
        [&sorted_indexes]() {
          std::priority_queue<IndexMergeHelper> result;
          for (unsigned int s{0}; s != sorted_indexes.size(); ++s) {
            const MappedVector<TempMUMindex> & sorted{sorted_indexes[s]};
            result.emplace(sorted);
          }
          return result;
        }()};

      const std::string index_name{mumdex_name + "/index.bin"};
      FILE * index_file = fopen(index_name.c_str(), "wb");
      if (index_file == NULL) {
        throw Error("Could not open index file for writing") << index_name;
      }
      std::vector<MUMindex> indexes;
      const uint64_t index_merge_size{n_gb * gb / sizeof(MUMindex)};
      indexes.reserve(index_merge_size);
      while (index_mergers.size()) {
        IndexMergeHelper top{index_mergers.top()};
        indexes.push_back(top.index());
        index_mergers.pop();
        if (++top) index_mergers.push(top);
        if (indexes.size() == index_merge_size || index_mergers.empty()) {
          bwrite(index_file, indexes[0], "MUM Index", indexes.size());
          indexes.clear();
        }
      }
      fclose(index_file);

      const time_t stop_time{time(nullptr)};
      std::cerr << "Merge and index sort took " << sort_start_time - start_time
                << " seconds and index merge took "
                << stop_time - sort_start_time << " seconds." << std::endl;
      // Test sort
#if 0
      const MUMdex test{mumdex_name};
      if (!std::is_sorted(test.index().begin(), test.index().end(),
                          lessMUM_index<UnMappedVector>(
                              test.mums(), test.pairs()))) {
        throw Error("Not sorted");
      } else {
        std::cerr << "Sort ok" << std::endl;
      }
#endif
    }

    // Remove parts and change mumdex to write protected for safety
    if (true) {
      for (const std::string & file : sort_files) {
        if (system((std::string("rm -R ") + file).c_str()))
          throw Error("Problem removing temporary sort files");
      }
      for (const std::string & part : part_names) {
        if (system((std::string("rm -R ") + part).c_str()))
          throw Error("Problem removing mumdex parts");
      }
      if (system((std::string("chmod a-w ") + mumdex_name + "/*.bin").c_str()))
          throw Error("Problem making mumdex write protected");
    }
  }
};

// Class to handle options parsing and to deliver options to classes
class Args : public SAArgs, public ReadPairsArgs, public ReadersArgs {
 public:
  Args(const std::string ref_fasta_,
       const bool read_ahead_ = false,
       const unsigned int n_threads_ = 1,
       const std::vector<std::string> & optional_ =
       std::vector<std::string>()) :
      SAArgs(), ReadPairsArgs(), ReadersArgs(), verbose(true) {
    ref_args.verbose = true;
    SAArgs::verbose = true;
    ReadPairsArgs::verbose = true;
    ReadersArgs::verbose = true;
    ref_args.rcref = true;
    sam_in = true;
    read_ahead = read_ahead_;
    memory_mapped = true;
    if (optional_.size()) {
      optional = optional_;
      pass_optional = false;
    }
    fasta = ref_fasta_;
    ref_args.ref_fasta = fasta.c_str();
    n_threads = n_threads_;
    mappability_threads = n_threads;
  }
  void set_sam_files(const std::vector<std::string> & sams) {
    for (unsigned int s{0}; s != sams.size(); ++s) {
      saved_sams[s] = const_cast<char *>(sams[s].c_str());
    }
    input = &saved_sams[0];
    n_input = static_cast<unsigned int>(sams.size());
  }
  bool verbose;

 private:
  void usage(const std::string & prog) const;
  Args(const Args &) = delete;
  Args & operator=(const Args &) = delete;
  char * saved_sams[200];
  std::string fasta{};
};

class Mapper {
 public:
  Mapper(const std::string & fasta,
         const bool read_ahead_ = false,
         const unsigned int n_threads = 8,
         const std::vector<std::string> & optional_ =
         std::vector<std::string>()) :
      args{fasta, read_ahead_, n_threads, optional_}, sa_{args} { }

  void create_mumdex(const std::vector<std::string> & sams,
                     const std::string & mumdex_name,
                     const unsigned int n_threads = 1) {
    args.set_sam_files(sams);
    {
      // Helps read queries
      Readers readers(args);

      // Pairs manages Pair worker threads
      longReadPairs pairs(args, sa_, readers);
    }
    // Merge mumdex and output
    const MUMdexMergerNew merger{"mumdex", 2, n_threads, true};

    // std::cerr << "Done" << std::endl;

    // Change name of mumdex from default
    if (mumdex_name != "mumdex")
      if (system((std::string("mv mumdex ") + mumdex_name).c_str()))
        throw Error("Problem moving mumdex to") << mumdex_name;
  }

  const longSA & sa() const {
    return sa_;
  }

 private:
    Args args;
    longSA sa_;
};

}  // namespace paa

#endif  // MAPPER_H_
```
