## Supplementary material for "G-Graph: An interactive genomic graph viewer": MUMdex Source Code for G-Graph: merge_mumdex.cpp

```
//
// merge_mumdex
//
// takes mumdex parts produced by mummer and merges them
//
// Copyright 2015 Peter Andrews @ CSHL
//

#include <exception>
#include <string>

#include "error.h"
#include "mapper.h"

using std::cerr;
using std::endl;
using std::exception;
using std::string;

using paa::Error;
using paa::MUMdexMergerNew;

int main(int argc, char * argv[]) try {
  --argc;
  if (argc != 2 && argc != 3 && argc != 4)
    throw Error("usage: merge_mumdex mumdex_name max_gb "
                "[n_threads] [mark_dupes]");

  const string mumdex_name{argv[1]};
  const unsigned int n_gb = atoi(argv[2]);
  if (n_gb == 0) throw Error("max GB not set properly");
  const unsigned int n_threads = argc == 3 ? atoi(argv[3]) : 1;
  if (n_threads == 0) throw Error("N threads not set properly");
  const bool mark_dupes{static_cast<bool>(argc == 4 ? atoi(argv[4]) : 1)};
  const MUMdexMergerNew merger{mumdex_name, n_gb, n_threads, mark_dupes};

  return 0;
} catch (Error & e) {
  cerr << "paa::Error:" << endl;
  cerr << e.what() << endl;
  return 1;
} catch (exception & e) {
  cerr << "std::exception" << endl;
  cerr << e.what() << endl;
  return 1;
} catch (...) {
  cerr << "unknown exception was caught" << endl;
  return 1;
}
```
