## Supplementary material for "G-Graph: An interactive genomic graph viewer": MUMdex Source Code for G-Graph: mumdex.h

```
//
// mumdex.h
//
// MUM-based alignment / mapping index
//
// Copyright 2015-2017 Peter Andrews @ CSHL
//

#ifndef PAA_MUMDEX_H
#define PAA_MUMDEX_H

#include <algorithm>
#include <array>
#include <cstdio>
#include <fstream>
#include <map>
#include <sstream>
#include <string>
#include <utility>
#include <vector>

#include "error.h"
#include "files.h"
#include "strings.h"

/*

TODO:

bam input
bam read index
multi-thread merges OK
child table
clip bad q scores
speed up optional passthrough
viewer: half
BedFile in bed.h should use chromosome indices, not strings
add low/high N anchor flags to mum structure by squeezing position 
add maximum exact match length in read
more efficient bases storage
nearby bridges
improve known gene input OK
internal tests: both, mendel, % in pop, technical replicates
merge_mumdex parallel sort, lower memory requirement OK
explaining spurious bridges
whole snp sequence
N check
is exon inside long deletions
population vs family cuts

*/

namespace paa {

inline char complement(const char base) {
  switch (base) {
    case 'x': return 'x';
    case 'X': return 'X';
    case 'n': return 'n';
    case 'N': return 'N';
    case 'a': return 't';
    case 'c': return 'g';
    case 'g': return 'c';
    case 't': return 'a';
    case 'r': return 'y'; /* a or g */
    case 'y': return 'r'; /* c or t */
    case 'm': return 'k'; /* a or c */
    case 'k': return 'm'; /* g or t */
    case 'b': return 'v'; /* c, g or t */
    case 'd': return 'h'; /* a, g or t */
    case 'h': return 'd'; /* a, c or t */
    case 'v': return 'b'; /* a, c or g */
    case 'A': return 'T';
    case 'C': return 'G';
    case 'G': return 'C';
    case 'T': return 'A';
    case 'R': return 'Y'; /* a or g */
    case 'Y': return 'R'; /* c or t */
    case 'M': return 'K'; /* a or c */
    case 'K': return 'M'; /* g or t */
    case 'B': return 'V'; /* c, g or t */
    case 'D': return 'H'; /* a, g or t */
    case 'H': return 'D'; /* a, c or t */
    case 'V': return 'B'; /* a, c or g */
    default:
      throw Error("Unexpected base in reverse_complement") << base;
  }
}

// Return the reverse complement of sequence. This allows searching
// the plus strand of instances on the minus strand.
inline void reverse_complement(std::string * const seq_rc) {
  // Reverse in-place.
  reverse(seq_rc->begin(), seq_rc->end());
  for (uint64_t i = 0; i != seq_rc->size(); ++i) {
    // Adapted from Kurtz code in MUMmer v3.
    char & ch = (*seq_rc)[i];
    switch (ch) {
      case 'x': ch = 'x'; break;
      case 'X': ch = 'X'; break;
      case 'n': ch = 'n'; break;
      case 'N': ch = 'N'; break;
      case 'a': ch = 't'; break;
      case 'c': ch = 'g'; break;
      case 'g': ch = 'c'; break;
      case 't': ch = 'a'; break;
      case 'r': ch = 'y'; break; /* a or g */
      case 'y': ch = 'r'; break; /* c or t */
      case 'm': ch = 'k'; break; /* a or c */
      case 'k': ch = 'm'; break; /* g or t */
      case 'b': ch = 'v'; break; /* c, g or t */
      case 'd': ch = 'h'; break; /* a, g or t */
      case 'h': ch = 'd'; break; /* a, c or t */
      case 'v': ch = 'b'; break; /* a, c or g */
      case 'A': ch = 'T'; break;
      case 'C': ch = 'G'; break;
      case 'G': ch = 'C'; break;
      case 'T': ch = 'A'; break;
      case 'R': ch = 'Y'; break; /* a or g */
      case 'Y': ch = 'R'; break; /* c or t */
      case 'M': ch = 'K'; break; /* a or c */
      case 'K': ch = 'M'; break; /* g or t */
      case 'B': ch = 'V'; break; /* c, g or t */
      case 'D': ch = 'H'; break; /* a, g or t */
      case 'H': ch = 'D'; break; /* a, c or t */
      case 'V': ch = 'B'; break; /* a, c or g */
      default:
        throw Error("Unexpected base in reverse_complement") << ch;
    }
  }
}

inline std::string reverse_complement(std::string sequence) {
  reverse_complement(&sequence);
  return sequence;
}

class PosInfo {
 public:
  PosInfo(const unsigned int chr_, const unsigned int pos_) :
      chr{chr_}, pos{pos_} {}
  unsigned int chr;
  unsigned int pos;
  bool operator<(const PosInfo & other) const {
    if (chr == other.chr) {
      return pos < other.pos;
    } else {
      return chr < other.chr;
    }
  }
  bool operator>=(const PosInfo & other) const {
    if (chr == other.chr) {
      return pos >= other.pos;
    } else {
      return chr >= other.chr;
    }
  }
  bool operator==(const PosInfo & other) const {
    return chr == other.chr && pos == other.pos;
  }
};

template <template <class ...> class VECTOR>
class TReference {
 public:
  using const_iterator = typename VECTOR<char>::const_iterator;
  TReference() {}
  TReference(const TReference &) = delete;
  TReference(TReference &&) = delete;
  TReference & operator=(const TReference &) = delete;
  TReference & operator=(TReference &&) = delete;
  TReference(const std::string & fasta, bool) = delete;  // just in case
  TReference(const std::string & fasta, double) = delete;  // just in case

  // To distinguish constructor usage
  class ReadFromBinary {};
  class CreateFromAlignerSequence {};

  // construct from mummer rc ref aligner reference
  template <class Sequence>
  TReference(const Sequence & mref, const CreateFromAlignerSequence) :
      seq{mref.rcref ? mref.N / 2 : mref.N} {
    for (unsigned int c{0}; c != mref.descr.size();
         (mref.rcref ? c += 2 : ++c)) {
      chr_len.push_back(static_cast<unsigned int>(mref.sizes[c]));
      for (unsigned int b{0}; b != chr_len.back(); ++b) {
        const char base{mref[mref.startpos[c] + b]};
        seq.push_back(base == 'X' ? 'N' : base);
      }
      chr_name.push_back(mref.descr[c]);
    }
    uint64_t pos{0};
    for (unsigned int c{0}; c != chr_len.size(); ++c) {
      chr.push_back(seq.begin() + pos);
      chrs.push_back(c);
      pos += chr_len[c];
    }
  }

  // read from fasta file or binary - and do not use too much physical memory
  explicit TReference(const std::string & fasta,
                      const size_t max_bytes = 4906) :
      seq{"/dev/null", false},
    chr_len{"/dev/null", false},
    fasta_file_{fasta} {
      const std::string ref_bin_name{fasta + ".bin/ref.seq.bin"};
      if (readable(ref_bin_name)) {
        new (this) TReference{fasta, ReadFromBinary()};  // placement new
        return;
      }
      std::cerr << "Creating binary reference cache from "
                << fasta << std::endl;
      {
        mkdir(fasta + ".bin/");
        FILE * output = fopen(ref_bin_name.c_str(), "wb");
        TReference<GrowingVector> ref;
        ref.read_and_write_from_fasta(fasta, output, max_bytes);
        if (fclose(output) != 0)
          throw Error("problem closing output file") << ref_bin_name;
        ref.save(fasta, false);
      }
      new (this) TReference{fasta};
    }

  void read_and_write_from_fasta(const std::string & fasta,
                                 FILE * out_file,
                                 const uint64_t max_bytes) {
    std::ifstream input{fasta.c_str()};
    if (!input) {
      throw Error("Could not open fasta file") << fasta;
    }
    std::string line;
    unsigned int chr_end{0};
    uint64_t seq_size{0};
    while (input >> line) {
      if (line[0] == '>') {
        if (chr_name.size()) {
          chr_len.push_back(static_cast<unsigned int>(seq_size - chr_end));
        }
        chr_end = static_cast<unsigned int>(seq_size);
        chrs.push_back(static_cast<unsigned int>(chr_name.size()));
        chr_name.push_back(line.substr(1));
        input.ignore(10000, '\n');
      } else {
        for (unsigned int b{0}; b != line.size(); ++b) {
          seq.push_back(toupper(line[b]));
          ++seq_size;
        }
      }
      if (seq.size() > max_bytes) {
        seq.write(out_file);
        seq.clear();
      }
    }
    if (seq.size()) {
      seq.write(out_file);
      seq.clear();
    }
    chr_len.push_back(static_cast<unsigned int>(seq_size - chr_end));
  }

  // output fasta - only used for validation or if fasta was deleted
  void fasta_out(const std::string & out_file,
                 const uint64_t line_length = 50) const {
    std::ofstream out(out_file.c_str());
    if (!out)
      throw Error("Could not open fasta file for writing") << out_file;
    for (unsigned int c{0}; c != n_chromosomes(); ++c) {
      out << '>' << name(c);
      for (unsigned int b{0}; b != size(c); ++b) {
        if ((b % line_length == 0)) out << '\n';
        out << (*this)[c][b];
      }
      out << '\n';
    }
  }
  std::string subseq(const unsigned int chromosome,
                     const unsigned int start_pos,
                     const unsigned int stop_pos) const {
    std::string out;
    for (unsigned int b{start_pos}; b != stop_pos; ++b) {
      out += chr[chromosome][b];
    }
    return out;
  }

  const std::string name() const {
    return remove_including_initial(
        remove_including_final(fasta_file_, '/'), '.');
  }

  // direct access
  uint64_t size() const { return seq.size(); }
  const_iterator begin() const { return seq.begin(); }
  const_iterator end() const { return seq.end(); }

  // chromosome access
  const std::vector<unsigned int> & chromosomes() const {
    return chrs;
  }
  unsigned int n_chromosomes() const {
    return static_cast<unsigned int>(chr.size()); }
  const std::string & name(const uint64_t chromosome) const {
    return chr_name[chromosome];
  }
  unsigned int offset(const uint64_t chromosome) const {
    return static_cast<unsigned int>(chr[chromosome] - begin());
  }
  unsigned int size(const uint64_t chromosome) const {
    return chr_len[chromosome];
  }
  const_iterator operator[](const uint64_t chromosome) const {
    return chr[chromosome];
  }
  const std::string & fasta_file() const {
    return fasta_file_;
  }
  unsigned int abspos(const unsigned int chromosome,
                      const unsigned int position) const {
    return offset(chromosome) + position;
  }
  using ChrPos = std::pair<unsigned int, unsigned int>;
  unsigned int abspos(const ChrPos chrpos_arg) const {
    return offset(chrpos_arg.first) + chrpos_arg.second;
  }
  ChrPos chrpos(const unsigned int abspos_arg) const {
    typename std::vector<const_iterator>::const_iterator upper_chr{
      std::upper_bound(chr.begin(), chr.end(), chr.front() + abspos_arg)};
    const unsigned int chr_{static_cast<unsigned int>(
        --upper_chr - chr.begin())};
    const unsigned int pos_{abspos_arg -
          static_cast<unsigned int>(chr[chr_] - chr[0])};
    return {chr_, pos_};
  }

  // X and Y chromosomes
  unsigned int find_x_chromosome() const {
    for (unsigned int c{0}; c != chr_name.size(); ++c) {
      if (chr_name[c].back() == 'X') {
        return c;
      }
    }
    throw Error("Could not determine X chromosome");
  }
  unsigned int find_y_chromosome() const {
    for (unsigned int c{0}; c != chr_name.size(); ++c) {
      if (chr_name[c].back() == 'Y') {
        return c;
      }
    }
    throw Error("Could not determine Y chromosome");
  }

  // save to binary file
  void save(const std::string & fasta, const bool save_seq = true) const {
    if (save_seq) {
      seq.save(fasta + ".bin/ref.seq.bin");
    }

    chr_len.save(fasta + ".bin/ref.chr_len.bin");
    std::string chr_names_filename{fasta + ".bin/ref.chr_name.bin"};
    std::ofstream chr_names_file{chr_names_filename.c_str()};
    if (!chr_names_file)
      throw Error("Could not open chromosome names file for writing")
          << chr_names_filename;
    for (const std::string & name_ : chr_name) {
      chr_names_file << name_ << '\n';
    }
  }

 private:
  // read from binary file (not from fasta)
  explicit TReference(const std::string & fasta, const ReadFromBinary) :
      seq{fasta + ".bin/ref.seq.bin"},
    chr_len{fasta + ".bin/ref.chr_len.bin"},
    fasta_file_{fasta} {
      std::string chr_names_filename{fasta_file_ + ".bin/ref.chr_name.bin"};
      std::ifstream chr_names_file{chr_names_filename.c_str()};
      if (!chr_names_file) {
        throw Error("Could not open chromosome names file for reading")
            << chr_names_filename;
      }
      std::string line;
      uint64_t pos{0};
      for (unsigned int c{0}; c != chr_len.size(); ++c) {
        if (!std::getline(chr_names_file, line)) {
          throw Error("Problem reading name for chromosome") << c;
        }
        chr_name.push_back(line);
        chr.push_back(seq.begin() + pos);
        chrs.push_back(c);
        pos += chr_len[c];
      }
    }

  VECTOR<char> seq{};
  VECTOR<unsigned int> chr_len{};
  std::vector<std::string> chr_name{};
  std::vector<const_iterator> chr{};
  std::vector<unsigned int> chrs{};
  std::string fasta_file_{"unknown"};
};

using PreMappedReference = TReference<PreMappedVector>;
using UnMappedReference = TReference<UnMappedVector>;
using MemoryReference = TReference<MemoryVector>;
using FileReference = TReference<FileVector>;
using GrowingReference = TReference<GrowingVector>;
using Reference = PreMappedReference;

class ChromosomeIndexLookup {
 public:
  template <class Ref>
  explicit ChromosomeIndexLookup(const Ref & ref) {
    for (unsigned int c{0}; c != ref.n_chromosomes(); ++c) {
      lookup_table.emplace(ref.name(c), c);
    }
  }
  // complaining lookup
  unsigned int operator[](const std::string & name) const {
    try {
      return lookup_table.at(name);
    } catch(...) {
      std::cerr << "Problem looking up chromsosome \""
                << name << "\"" << std::endl;
      throw;
    }
  }
  // quiet lookup (still throws)
  unsigned int operator()(const std::string & name) const {
    return lookup_table.at(name);
  }

  bool exists(const std::string & chr) const {
    return lookup_table.find(chr) != lookup_table.end();
  }

  unsigned int size() const {
    return static_cast<unsigned int>(lookup_table.size());
  }

 private:
  std::map<std::string, unsigned int> lookup_table{};
};

class RefPlus : public Reference {
 public:
  explicit RefPlus(const std::string & fasta_file_name) :
      Reference{fasta_file_name}, chr_lookup{*this} { }
  ~RefPlus() {}

  const ChromosomeIndexLookup chr_lookup;
};

inline std::string ref_name(const std::string & name) {
  return name + "/ref.txt";
}

inline std::string saved_ref_name(const std::string & mumdex_name) {
  std::ifstream ref_name_file{ref_name(mumdex_name).c_str()};
  if (!ref_name_file)
    throw Error("Problem opening reference name file for mumdex")
        << mumdex_name;
  std::string name;
  std::getline(ref_name_file, name);
  if (name.empty() || !ref_name_file)
    throw Error("Problem reading reference name file for mumdex")
        << mumdex_name;
  return name;
}

template <template <class ...> class VECTOR>
class TMappability {
  // Definition of the mappability values
  // 255 - no uniqueness even if we go to the end of the chromosome
  // 254 - no uniqueness even if we go 253 positions left or right
  // 0 < n <= 253 uniqueness if we go n positions left or right
  // and no uniqueness if we go n-1
  // should never be '0'

 public:
  TMappability(const std::string & fasta_name, bool) :
      low_{fasta_name + ".bin/map.low.bin"},
    high_{fasta_name + ".bin/map.high.bin"} {
      if (low_.size() != high_.size()) {
        throw Error("Mappability size mismatch for fasta_name");
      }
    }
  explicit TMappability(const std::string & mumdex_name) :
      TMappability{saved_ref_name(mumdex_name), true} {
  }
  template <class Ref>
  explicit TMappability(const Ref & ref) :
      TMappability{ref.fasta_file(), true} {
  }
  uint64_t size() const { return low_.size(); }
  unsigned int low(const uint64_t pos) const { return low_[pos]; }
  unsigned int high(const uint64_t pos) const { return high_[pos]; }
  unsigned int low_high(const bool is_high, const uint64_t pos) const {
    return is_high ? high_[pos] : low_[pos];
  }
  unsigned int min(const unsigned int pos, const unsigned int length) const {
    unsigned int result{255};
    const unsigned int stop{pos + length};
    for (unsigned int p{pos}; p != stop; ++p) {
      result = std::min(result, low(p));
      if (p + result > stop) break;
    }
    return result;
  }

 private:
  VECTOR<uint8_t> low_;
  VECTOR<uint8_t> high_;
};

using MemoryMappability = TMappability<MemoryVector>;
using PreMappedMappability = TMappability<PreMappedVector>;
using UnMappedMappability = TMappability<UnMappedVector>;
using FileMappability = TMappability<FileVector>;
using Mappability = PreMappedMappability;

// bit field lengths
constexpr unsigned int mums_bits{40};  // allows for 1,099,511,627,776 mums
constexpr unsigned int position_bits{32};  // min 28 for chr 1
constexpr unsigned int chromosome_bits{8};  // min 7 for chrAll (84 seq)
constexpr unsigned int read_bits{10};  // min 8 for up to 255 bp
constexpr unsigned int pair_mums_bits{read_bits + 1};
constexpr unsigned int bool_bits{1};

const char base_chars[]{"0ATCGNX--"};
inline uint64_t base_index(char base) {
  switch (base) {
    case 'A':
      return 1;
    case 'T':
      return 2;
    case 'C':
      return 3;
    case 'G':
      return 4;
    case 'N':
      return 5;
    case 'Z':
      return 5;
    case 'X':
      return 6;
    default:
      throw Error("Bad base error") << base;
  }
}

inline std::string extra_name(const std::string & name) {
  return name + "/bases.extra.bin";
}

class ExtraBasesSaveInfo {
 public:
  ExtraBasesSaveInfo(const std::vector<uint64_t> & last_words_arg,
                     const uint64_t n_bases_arg)
      : last_words_(last_words_arg), n_bases_(n_bases_arg) {}
  const std::vector<uint64_t> & last_words() const { return last_words_; }
  uint64_t n_bases() const { return n_bases_; }
  unsigned int n_excess_bases() const { return n_bases_ % 64; }
  uint64_t n_saved_bases() const { return n_bases_ - (n_bases_ % 64); }

 private:
  std::vector<uint64_t> last_words_;
  uint64_t n_bases_;
};

template <template <class ...> class VECTOR>
class TExtraBases {
 public:
  // construction
  TExtraBases() = default;
  TExtraBases(const TExtraBases &) = delete;
  TExtraBases(TExtraBases &&) = default;
  TExtraBases & operator=(const TExtraBases &) = delete;
  TExtraBases & operator=(TExtraBases &&) = delete;

  explicit TExtraBases(const uint64_t initial_size) :
      data{initial_size}, n_bases_{0} {}
  explicit TExtraBases(const std::string & name)
      : data{extra_name(name)}, n_bases_{data.size() * 64 / 3} {
        uint64_t last_index{n_bases_};
    if (last_index) {
      while (retrieve(--last_index) == base_chars[0]) --n_bases_;
    }
  }

  // info
  uint64_t bytes() const {
    return sizeof(TExtraBases) + data.bytes();
  }

  // grow
  uint64_t add(const std::string & bases_) {
    const uint64_t starting_index{n_bases_};
    uint64_t current_bases{n_bases_};
    n_bases_ += bases_.size() + 1;
    const uint64_t required_bits{n_bases_ * 3};
    const uint64_t required_words{(required_bits % 64 ? 1 : 0) +
          required_bits / 64};
    while (data.size() < required_words) data.push_back(0);
    for (uint64_t i{0}; i != bases_.size() + 1; ++i) {
      const uint64_t value{base_index(i == bases_.size() ? 'X' : bases_[i])};
      for (unsigned int bit{0}; bit != 3; ++bit) {
        const uint64_t bit_value{(value >> bit) & 1};
        if (bit_value) {
          const uint64_t bit_index{current_bases * 3 + bit};
          const uint64_t word_index{bit_index / 64};
          const uint64_t bit_in_word_index{bit_index % 64};
          data[word_index] |= (bit_value << bit_in_word_index);
        }
      }
      ++current_bases;
    }
    return starting_index;
  }
  void add_excess_bases(const ExtraBasesSaveInfo & info) {
    for (const uint64_t word : info.last_words()) {
      data.push_back(word);
    }
    n_bases_ += info.n_excess_bases();
  }

  // access
  char retrieve(const uint64_t base_index) const {
    uint64_t return_value{0};
    for (unsigned int bit{0}; bit != 3; ++bit) {
      const uint64_t bit_index{base_index * 3 + bit};
      const uint64_t word_index{bit_index / 64};
      const uint64_t bit_in_word_index{bit_index % 64};
      return_value |= (((data[word_index] >> bit_in_word_index) & 1) << bit);
    }
    return base_chars[return_value];
  }

  // saving
  void save(const std::string & name) const { data.save(name); }
  ExtraBasesSaveInfo write(std::FILE * file) const {
    const uint64_t n_complete_words{n_bases_ / 64 * 3};
    data.write_n(file, n_complete_words);
    return ExtraBasesSaveInfo{
      std::vector<uint64_t>(data.begin() + n_complete_words, data.end()),
          n_bases_};
  }
  void clear() {
    data.clear();
    n_bases_ = 0;
  }

 private:
  VECTOR<uint64_t> data{};
  uint64_t n_bases_{0};
};

class SplitBases {
 public:
  // construction
  template <class ExtraBases>
  SplitBases(const std::string & new_bases, ExtraBases & extra,
             const uint64_t offset = 0) {
    if (new_bases.size() < max_bases_in_word) {
      data = 0;
      for (uint64_t i{0}; i != new_bases.size(); ++i) {
        data |= (base_index(new_bases[i]) << (i * 3));
      }
      data |= (base_index('X') << (new_bases.size() * 3));
    } else {
      data = extra.add(new_bases) + offset;
      data |= index_bit;
    }
  }

  // access
  template <class ExtraBases>
  std::string bases(const ExtraBases & extra) const {
    std::string result;
    uint64_t i{0};
    while (true) {
      const char b{base(i++, extra)};
      if (b == 'X') return result;
      result += b;
    }
  }
  template <class ExtraBases>
  void bases(const ExtraBases & extra, std::string & result) const {
    result.clear();
    uint64_t i{0};
    while (true) {
      const char b{base(i++, extra)};
      if (b == 'X') return;
      result += b;
    }
  }

 private:
  SplitBases() = default;
  // access
  template <class ExtraBases>
  char base(const uint64_t base_index, const ExtraBases & extra) const {
    if (is_index()) return extra.retrieve(index() + base_index);
    return (*this)[base_index];
  }
  char operator[](const uint64_t base_index) const {
    return base_chars[(data >> (base_index * 3)) & 7UL];
  }

  // internal use
  static constexpr unsigned int max_bases_in_word{21};
  // mac and linux have different UL behaviors
  // so cast to make sure, ignore warning that is no big deal
#pragma GCC diagnostic push
#pragma GCC diagnostic ignored "-Wunknown-pragmas"
#pragma GCC diagnostic ignored "-Wuseless-cast"
  static constexpr uint64_t index_bit{
    static_cast<uint64_t>(9223372036854775808UL)};  // 1UL << 63
  static constexpr uint64_t data_bits{
    static_cast<uint64_t>(9223372036854775807UL)};  // ~index_bit
#pragma GCC diagnostic pop
  bool is_index() const { return data & index_bit; }
  uint64_t index() const { return data & data_bits; }
  uint64_t data{};
};

inline std::string bases_name(const std::string & name) {
  return name + "/bases.bin";
}

template <template <class ...> class VECTOR>
class TAllBases {
 public:
  using ExtraBases = TExtraBases<VECTOR>;
  // construct and assign
  TAllBases() = default;
  TAllBases(TAllBases &&) = default;
  explicit TAllBases(const uint64_t initial_size) :
      bases_{initial_size}, extra_{initial_size} {}
  explicit TAllBases(const std::string & base_name) :
      bases_{bases_name(base_name)}, extra_{base_name} {}

  // deleted
  TAllBases(const TAllBases &) = delete;
  TAllBases & operator=(const TAllBases &) = delete;

  // info and access
  uint64_t size() const { return bases_.size(); }
  uint64_t bytes() const {
    return sizeof(TAllBases) + bases_.bytes() + extra_.bytes();
  }
  std::string bases(const uint64_t pair) const {
    return bases_[pair].bases(extra_);
  }
  void bases(const uint64_t pair, std::string & result) const {
    bases_[pair].bases(extra_, result);
  }

  const VECTOR<SplitBases> & bases() const { return bases_; }
  VECTOR<SplitBases> & bases() { return bases_; }
  const ExtraBases & extra() const { return extra_; }
  ExtraBases & extra() { return extra_; }

  // grow
  void push_back(const std::string & val, const uint64_t offset = 0) {
    bases_.emplace_back(val, extra_, offset);
  }

  // save
  void save(const std::string & bases_name,
            const std::string & extra_name) const {
    bases_.save(bases_name);
    extra_.save(extra_name);
  }
  ExtraBasesSaveInfo write(std::FILE * bases_file,
                           std::FILE * extra_file) const {
    bases_.write(bases_file);
    return extra_.write(extra_file);
  }
  void clear() {
    bases_.clear();
    extra_.clear();
  }

 private:
  VECTOR<SplitBases> bases_{};
  ExtraBases extra_{};
};

class Invariant {
 public:
  template <class PAIR, class MUM>
  Invariant(const PAIR pair, const MUM left, const MUM right) :
      extra{0}, ordinary_{false} {
    if (left.offset() >= right.offset())
      throw Error("MUM offset misordering in Invariant");
    if (left.read_2() != right.read_2())
      throw Error("Calculating invariant for mums on different reads");
    const unsigned int length{pair.length(left.read_2())};
    low_chromosome_ = left.chromosome() < right.chromosome() ?
        left.chromosome() : right.chromosome();
    high_chromosome_ = left.chromosome() < right.chromosome() ?
        right.chromosome() : left.chromosome();
    if (left.flipped() == right.flipped()) {
      ordinary_ = true;
      value_ = left.read_position0(length) - right.read_position0(length);
    } else {
      value_ = left.read_position0(length) + right.read_position0(length) +
          length;
    }
    if (left.flipped()) value_ = -value_;
  }
  bool operator<(const Invariant rhs) const {
    if (low_chromosome_ == rhs.low_chromosome_) {
      if (high_chromosome_ == rhs.high_chromosome_) {
        if (ordinary_ == rhs.ordinary_) {
          return value_ < rhs.value_;
        } else {
          return ordinary_ < rhs.ordinary_;
        }
      } else {
        return high_chromosome_ < rhs.high_chromosome_;
      }
    } else {
      return low_chromosome_ < rhs.low_chromosome_;
    }
  }
  bool operator==(const Invariant rhs) const {
    return low_chromosome_ == rhs.low_chromosome_ &&
        high_chromosome_ == rhs.high_chromosome_ &&
        ordinary_ == rhs.ordinary_ &&
        value_ == rhs.value_;
  }
  std::string string() const {
    std::ostringstream out;
    out << ordinary_ << ':'
        << low_chromosome_ << ':'
        << high_chromosome_ << ':'
        << value_;
    return out.str();
  }
  bool ordinary() const { return ordinary_; }
  int value() const { return value_; }
  unsigned int abs_value() const { return value_ > 0 ? value_ : -value_; }
  int16_t low_chromosome() const { return low_chromosome_; }
  int16_t high_chromosome() const { return high_chromosome_; }
  void hack() { extra = 0; }

 private:
  int value_;
  uint16_t low_chromosome_: chromosome_bits;
  uint16_t high_chromosome_: chromosome_bits;
  uint16_t extra: 15;
  uint16_t ordinary_: 1;
};

class Pair {
 public:
  // construct
  Pair(const uint64_t mums_start_arg,
       const uint64_t read_1_length_arg,
       const bool read_1_bad_arg,
       const uint64_t read_2_length_arg,
       const bool read_2_bad_arg,
       const bool has_mums_arg) :
      mums_start_{mums_start_arg},
    read_1_length_{static_cast<unsigned int>(read_1_length_arg)},
    read_2_length_{static_cast<unsigned int>(read_2_length_arg)},
    read_1_bad_{read_1_bad_arg},
    read_2_bad_{read_2_bad_arg},
    has_mums_{has_mums_arg},
    dupe_{false} {}

  // access
  uint64_t mums_start() const { return mums_start_; }
  unsigned int read_1_length() const { return read_1_length_; }
  unsigned int read_2_length() const { return read_2_length_; }
  bool read_1_bad() const { return read_1_bad_; }
  bool read_2_bad() const { return read_2_bad_; }
  bool dupe() const { return dupe_; }
  bool has_mums() const { return has_mums_; }

  // invariant
  template <class MUM>
  Invariant invariant(const MUM left, const MUM right) const {
    return Invariant(*this, left, right);
  }

  // access by read 0/1
  unsigned int length(const uint64_t i) const {
    return i ? read_2_length_ : read_1_length_;
  }
  bool bad(const uint64_t i) const { return i ? read_2_bad_ : read_1_bad_; }

  // modify
  void mums_start(const uint64_t new_start) { mums_start_ = new_start;}
  void read_1_bad(const bool new_bad) { read_1_bad_ = new_bad; }
  void read_2_bad(const bool new_bad) { read_2_bad_ = new_bad; }
  void bad(const uint64_t i, const bool new_bad) {
    if (i) {
      read_2_bad_ = new_bad;
    } else {
      read_1_bad_ = new_bad;
    }
  }
  void dupe(const bool new_dupe) { dupe_ = new_dupe; }

  // ordering
  bool empty_less(const Pair right) const {
    if (dupe() == right.dupe()) {
      if (read_1_bad() == right.read_1_bad()) {
        if (read_2_bad() == right.read_2_bad()) {
          if (read_1_length() == right.read_1_length()) {
            if (read_2_length() == right.read_2_length()) {
              return false;
            } else {
              return read_2_length() > right.read_2_length();
            }
          } else {
            return read_1_length() > right.read_1_length();
          }
        } else {
          return read_2_bad() < right.read_2_bad();
        }
      } else {
        return read_1_bad() < right.read_1_bad();
      }
    } else {
      return dupe() < right.dupe();
    }
  }

  template <class MUM>
  std::array<const MUM * const, 2> ordered_first_mums(const MUM * mums) const {
    std::array<const MUM *, 2> ordered{{nullptr, nullptr}};
    if (has_mums()) {
      for (const MUM * mum_{mums + mums_start()}; ; ++mum_) {
        if (!ordered[mum_->read_2()]) {
          ordered[mum_->read_2()] = mum_;
          if (mum_->read_2()) break;
        }
        if (mum_->last_hit()) break;
      }
    }
    if (ordered[1]) {
      if (!ordered[0] ||
          ordered[1]->lessReadPos(*ordered[0],
                                  read_2_length(), read_1_length())) {
        using std::swap;
        swap(ordered[0], ordered[1]);
      }
    }
    return std::array<const MUM * const, 2>{{ordered[0], ordered[1]}};
  }

  template <class MUM>
  bool lessPair(const MUM * const left_mums,
                const Pair right, const MUM * const right_mums) const {
    using OrderedMums = std::array<const MUM * const, 2>;
    const OrderedMums left_ordered = ordered_first_mums(left_mums);
    const OrderedMums right_ordered = right.ordered_first_mums(right_mums);

    const MUM * left_mum{left_ordered[0]};
    const MUM * right_mum{right_ordered[0]};
    if (!right_mum) {
      if (!left_mum) {
        return empty_less(right);
      }
      return true;
    }
    if (!left_mum) {
      return false;
    }
    if (left_mum->chromosome() == right_mum->chromosome() &&
        left_mum->read_position0(length(left_mum->read_2())) ==
        right_mum->read_position0(right.length(right_mum->read_2())) &&
        read_1_length_ == right.read_1_length_ &&
        read_2_length_ == right.read_2_length_) {
      const MUM * left_mum_2{left_ordered[1]};
      const MUM * right_mum_2{right_ordered[1]};
      if (!right_mum_2) {
        if (!left_mum_2) {
          return empty_less(right);
        }
        return true;
      }
      if (!left_mum_2) {
        return false;
      }
      if (left_mum_2->chromosome() == right_mum_2->chromosome() &&
          left_mum_2->read_position0(length(left_mum_2->read_2())) ==
          right_mum_2->read_position0(right.length(right_mum_2->read_2()))) {
        return empty_less(right);
      }
      return left_mum_2->lessReadPos(*right_mum_2,
                                     length(left_mum_2->read_2()),
                                     right.length(right_mum_2->read_2()));
    }
    return left_mum->lessReadPos(*right_mum,
                                 length(left_mum->read_2()),
                                 right.length(right_mum->read_2()));
  }

 private:
  Pair() = default;
  uint64_t mums_start_: mums_bits;
  unsigned int read_1_length_: read_bits;
  unsigned int read_2_length_: read_bits;
  unsigned int read_1_bad_: bool_bits;
  unsigned int read_2_bad_: bool_bits;
  unsigned int has_mums_: bool_bits;
  unsigned int dupe_: bool_bits;
};

// instead of pairs, mums, index
// have pairs, mums, sorted mums, sorted pairs?

class MUM {
 public:
  // construction
  MUM(const uint64_t chromosome_arg,
      const uint64_t position_arg,  // up to 2^31, due to occasional int
      const uint64_t offset_arg,
      const uint64_t length_arg,
      const bool flipped_arg,
      const bool read_2_arg,
      const bool last_hit_arg,
      const bool touches_end_arg) :
      position_{static_cast<unsigned int>(position_arg)},
    chromosome_{static_cast<unsigned int>(chromosome_arg)},
    offset_{static_cast<unsigned int>(offset_arg)},
    length_{static_cast<unsigned int>(length_arg)},
    flipped_{flipped_arg},
    read_2_{read_2_arg},
    last_hit_{last_hit_arg},
    touches_end_{touches_end_arg} {}

  // this one is for lookup purposes only
  MUM(const uint64_t chromosome_arg,
      const uint64_t position_arg) :
      position_{static_cast<unsigned int>(position_arg)},
    chromosome_{static_cast<unsigned int>(chromosome_arg)},
    offset_{0},
    length_{0},
    flipped_{0},
    read_2_{0},
    last_hit_{0},
    touches_end_{0} {}

  // access
  unsigned int chromosome() const { return chromosome_; }
  unsigned int position0() const { return position_; }
  unsigned int position1() const { return position_ + 1; }
  int read_position0(const uint64_t read_length) const {
    if (flipped_) {
      return static_cast<int>(position_) - static_cast<int>(read_length) +
          static_cast<int>(offset_) + static_cast<int>(length_);
    } else {
      return static_cast<int>(position_) - static_cast<int>(offset_);
    }
  }
  int read_position1(const uint64_t read_length) const {
    return read_position0(read_length) + 1;
  }
  unsigned int offset() const { return offset_; }
  unsigned int stop_offset() const { return offset_ + length_; }
  unsigned int length() const { return length_; }
  bool flipped() const { return flipped_; }
  bool read(const bool read_2_arg) const { return read_2_ == read_2_arg; }
  bool read_2() const { return read_2_; }
  bool last_hit() const { return last_hit_; }
  bool touches_end() const { return touches_end_; }

  // anchor position and offset, zero based
  unsigned int low_position() const { return position_; }
  unsigned int high_position() const { return position_ + length_ - 1; }
  unsigned int anchor_position(const bool high) const {
    return high ? high_position() : low_position();
  }
  unsigned int low_offset() const {
    return flipped_ ? length_ - offset_ - 1 : offset_;
  }
  unsigned int high_offset() const {
    return flipped_ ? offset_ : length_ - offset_ - 1;
  }
  unsigned int anchor_offset(const bool high) const {
    return high ? high_offset() : low_offset();
  }
  int read_start_position() const {
    return flipped_ ? static_cast<int>(position_) + static_cast<int>(offset_) +
        static_cast<int>(length_) - 1 :
        static_cast<int>(position_) - static_cast<int>(offset_);
  }

  // coordinate transformation
  int read_to_position0(const int base) const {
    if (flipped_) {
      return static_cast<int>(position_) + static_cast<int>(length_) +
          static_cast<int>(offset_) - base - 1;
    } else {
      return static_cast<int>(position_) - static_cast<int>(offset_) + base;
    }
  }
  int position0_to_read(const int pos) const {
    if (flipped_) {
      return static_cast<int>(position_) + static_cast<int>(length_) +
          static_cast<int>(offset_) - pos - 1;
    } else {
      return pos - static_cast<int>(position_) + static_cast<int>(offset_);
    }
  }

  // MUM equality
  bool operator==(const MUM right) const {
    return static_cast<uint64_t>(*this) == static_cast<uint64_t>(right);
    return chromosome_ == right.chromosome_ &&
        position_ == right.position_ &&
        offset_ == right.offset_ &&
        length_ == right.length_ &&
        flipped_ == right.flipped_ &&
        read_2_ == right.read_2_ &&
        last_hit_ == right.last_hit_ &&
        touches_end_ == right.touches_end_;
  }

  // MUM ordering
  bool operator<(const MUM right) const {
    if (chromosome() == right.chromosome()) {
      if (position0() == right.position0()) {
        if (read_2() == right.read_2()) {
          if (offset() == right.offset()) {
            if (length() == right.length()) {
              if (flipped() == right.flipped()) {
                if (last_hit() == right.last_hit()) {
                  if (touches_end() == right.touches_end()) {
                    return false;
                  } else {
                    return touches_end() < right.touches_end();
                  }
                } else {
                  return last_hit() < right.last_hit();
                }
              } else {
                return flipped() < right.flipped();
              }
            } else {
              return length() < right.length();
            }
          } else {
            return offset() < right.offset();
          }
        } else {
          return read_2() < right.read_2();
        }
      } else {
        return position0() < right.position0();
      }
    } else {
      return chromosome() < right.chromosome();
    }
  }
  bool operator<=(const MUM right) const {
    if (chromosome() == right.chromosome()) {
      if (position0() == right.position0()) {
        if (read_2() == right.read_2()) {
          if (offset() == right.offset()) {
            if (length() == right.length()) {
              if (flipped() == right.flipped()) {
                if (last_hit() == right.last_hit()) {
                  if (touches_end() == right.touches_end()) {
                    return false;
                  } else {
                    return touches_end() <= right.touches_end();
                  }
                } else {
                  return last_hit() <= right.last_hit();
                }
              } else {
                return flipped() <= right.flipped();
              }
            } else {
              return length() <= right.length();
            }
          } else {
            return offset() <= right.offset();
          }
        } else {
          return read_2() <= right.read_2();
        }
      } else {
        return position0() <= right.position0();
      }
    } else {
      return chromosome() <= right.chromosome();
    }
  }
  bool operator>(const MUM right) const {
    if (chromosome() == right.chromosome()) {
      if (position0() == right.position0()) {
        if (read_2() == right.read_2()) {
          if (offset() == right.offset()) {
            if (length() == right.length()) {
              if (flipped() == right.flipped()) {
                if (last_hit() == right.last_hit()) {
                  if (touches_end() == right.touches_end()) {
                    return false;
                  } else {
                    return touches_end() > right.touches_end();
                  }
                } else {
                  return last_hit() > right.last_hit();
                }
              } else {
                return flipped() > right.flipped();
              }
            } else {
              return length() > right.length();
            }
          } else {
            return offset() > right.offset();
          }
        } else {
          return read_2() > right.read_2();
        }
      } else {
        return position0() > right.position0();
      }
    } else {
      return chromosome() > right.chromosome();
    }
  }

  // Pair ordering
  bool lessReadPos(const MUM right,
                   const uint64_t left_length,
                   const uint64_t right_length) const {
    if (chromosome() == right.chromosome()) {
      if (read_position0(left_length) == right.read_position0(right_length)) {
        if (length() == right.length()) {
          if (offset() == right.offset()) {
            if (read_2() == right.read_2()) {
              if (flipped() == right.flipped()) {
                if (last_hit() == right.last_hit()) {
                  if (touches_end() == right.touches_end()) {
                    return false;
                  } else {
                    return touches_end() < right.touches_end();
                  }
                } else {
                  return last_hit() < right.last_hit();
                }
              } else {
                return flipped() < right.flipped();
              }
            } else {
              return read_2() < right.read_2();
            }
          } else {
            return offset() < right.offset();
          }
        } else {
          return length() < right.length();
        }
      } else {
        return read_position0(left_length) <
            right.read_position0(right_length);
      }
    } else {
      return chromosome() < right.chromosome();
    }
  }

  // proximity
  bool close_by(const MUM other, const uint64_t distance) const {
    if (chromosome_ != other.chromosome_) return false;
    if (position_ + distance < other.position_ ||
        other.position_ + distance < position_)
      return false;
    return true;
  }

  int pos_to_base(const int pos) const {
    if (flipped()) {
      return position0() + offset() + length() - pos - 1;
    } else {
      return pos + offset() - position0();
    }
  }

  MUM() : position_{0}, chromosome_{0},
    offset_{0}, length_{0}, flipped_{0},
    read_2_{0}, last_hit_{0}, touches_end_{0} { }

 private:
  // look at bits
  operator uint64_t() const {
    return *reinterpret_cast<const uint64_t *>(this);
  }

  unsigned int position_: position_bits;
  unsigned int chromosome_: chromosome_bits;
  unsigned int offset_: read_bits;
  unsigned int length_: read_bits;
  unsigned int flipped_: bool_bits;
  unsigned int read_2_: bool_bits;
  unsigned int last_hit_: bool_bits;  // last in pair
  unsigned int touches_end_: bool_bits;
};

class MUMindex {
 public:
  MUMindex(const uint64_t pair_index_arg, const uint16_t mum_index_arg = 0,
           const uint16_t second_mum_index_arg = 0) :
      pair_index_{pair_index_arg}, mum_index_{mum_index_arg},
    second_mum_index_{second_mum_index_arg}, filler{0} {}

  operator uint64_t() const { return pair_index_; }
  uint64_t pair_index() const { return pair_index_; }
  template <class MUMDEX>
  uint64_t mum_index(const MUMDEX & mumdex) const {
    return mumdex.mums_start(pair_index_) + mum_index_;
  }
  unsigned int mum_in_pair_index() const { return mum_index_; }
  unsigned int second_mum_in_pair_index() const { return second_mum_index_; }
  bool index_less(const MUMindex rhs) const {
    if (pair_index_ == rhs.pair_index_) {
      // ignore second_mum_index_
      return mum_index_ < rhs.mum_index_;
    } else {
      return pair_index_ < rhs.pair_index_;
    }
  }
  void hack() { filler = 0; }

 private:
  MUMindex() = default;
  uint64_t pair_index_: mums_bits;
  uint64_t mum_index_ : pair_mums_bits;
  uint64_t second_mum_index_ : pair_mums_bits;
  uint64_t filler : 64 - mums_bits - 2 * pair_mums_bits;
};

class TempMUMindex {
 public:
  TempMUMindex(const MUMindex index_arg, const MUM mum_arg) :
      index_{index_arg}, mum_{mum_arg} {}
  bool operator<(const TempMUMindex & rhs) const {
    if (mum_ == rhs.mum_) return index_.index_less(rhs.index_);
    return mum_ < rhs.mum_;
  }
  bool operator<=(const TempMUMindex & rhs) const {
    if (mum_ == rhs.mum_) return index_.index_less(rhs.index_);
    return mum_ <= rhs.mum_;
  }
  bool operator>(const TempMUMindex & rhs) const {
    if (mum_ == rhs.mum_) return index_.index_less(rhs.index_);
    return mum_ > rhs.mum_;
  }
  operator MUMindex() const { return index_; }

  // private:
  MUMindex index_;
  MUM mum_;
};

template <template <class ...> class VECTOR>
class lessMUM_index {
 public:
  explicit lessMUM_index(const VECTOR<MUM> & mums_,
                         const VECTOR<Pair> & pairs_) :
      mums{&mums_}, pairs{&pairs_} {}
  bool operator()(const MUMindex lhs, const MUMindex rhs) const {
    const MUM lmum{(*mums)[(*pairs)[lhs.pair_index()].mums_start() +
                           lhs.mum_in_pair_index()]};
    const MUM rmum{(*mums)[(*pairs)[rhs.pair_index()].mums_start() +
                           rhs.mum_in_pair_index()]};
    if (lmum == rmum) return lhs.index_less(rhs);
    return lmum < rmum;
  }
  bool operator()(const MUM lhs, const MUMindex rhs) const {
    const MUM rmum{(*mums)[(*pairs)[rhs.pair_index()].mums_start() +
                           rhs.mum_in_pair_index()]};
    return lhs < rmum;
  }
  bool operator()(const MUMindex lhs, const MUM rhs) const {
    const MUM lmum{(*mums)[(*pairs)[lhs.pair_index()].mums_start() +
                           lhs.mum_in_pair_index()]};
    return lmum < rhs;
  }

 private:
  const VECTOR<MUM> * const mums;
  const VECTOR<Pair> * const pairs;
};

template <template <class ...> class VECTOR>
class lessPair_index {
 public:
  explicit lessPair_index(const VECTOR<Pair> & pairs_,
                          const VECTOR<MUM> & mums_) :
      pairs{&pairs_}, mums{&mums_} {}
  bool operator()(const uint64_t left_, const uint64_t right_) const {
    const Pair left{(*pairs)[left_]};
    const Pair right{(*pairs)[right_]};
    return left.lessPair(mums->begin(), right, mums->begin());
  }

 private:
  const VECTOR<Pair> * const pairs;
  const VECTOR<MUM> * const mums;
};

inline std::string pairs_name(const std::string & name) {
  return name + "/pairs.bin";
}
inline std::string mums_name(const std::string & name) {
  return name + "/mums.bin";
}
inline std::string index_name(const std::string & name) {
  return name + "/index.bin";
}

class mumdex_names {
 public:
  explicit mumdex_names(const std::string & name) :
      names{pairs_name(name),
        mums_name(name),
        bases_name(name),
        extra_name(name)} {}
  const std::string & operator[](const uint64_t i) const {
    return names[i];
  }
  uint64_t size() const { return names.size(); }
  std::vector<std::string>::const_iterator begin() const {
    return names.begin();
  }
  std::vector<std::string>::const_iterator end() const { return names.end(); }

 private:
  std::vector<std::string> names;
};

class PositionRegion {
 public:
  // 1-based input
  PositionRegion(const std::string & region,
                 const ChromosomeIndexLookup & chr_lookup) {
    std::istringstream region_stream{region.c_str()};
    std::string chromosome_string;
    getline(region_stream, chromosome_string, ':');
    start_chromosome = stop_chromosome = chr_lookup[chromosome_string];
    region_stream >> start_position;
    start_position -= 1;
    region_stream.get();
    region_stream >> stop_position;
    stop_position -= 1;
    if (!region_stream)
      throw Error("Problem parsing region string") << region;
  }
#if 0
  PositionRegion(const uint16_t start_chromosome_,
                 const unsigned int start_position_,
                 const uint16_t stop_chromosome_,
                 const unsigned int stop_position_) :
      start_position{start_position_}, stop_position{stop_position_},
      start_chromosome{start_chromosome_}, stop_chromosome{stop_chromosome_} {}
#endif

  unsigned int start_position{0};
  unsigned int stop_position{0};
  uint16_t start_chromosome{0};
  uint16_t stop_chromosome{0};
};

template <template <class ...> class VECTOR>
class TMUMRegion {
 public:
  using const_iterator = typename VECTOR<MUMindex>::const_iterator;
  TMUMRegion(const const_iterator begin_arg,
             const const_iterator end_arg) :
      begin_{begin_arg}, end_{end_arg} {}

  const_iterator begin() const { return begin_; }
  const_iterator end() const { return end_; }
  uint64_t size() const { return end() - begin(); }

 private:
  const_iterator begin_;
  const_iterator end_;
};

template <template <class ...> class VECTOR>
class TMUMdex_base {
 public:
  // typedefs
  using AllBases = TAllBases<VECTOR>;
  using ExtraBases = typename AllBases::ExtraBases;
  using MUMRegion = TMUMRegion<VECTOR>;
  using cMUMIter = typename VECTOR<MUM>::const_iterator;

  // construction
  TMUMdex_base() = default;
  TMUMdex_base(TMUMdex_base &&) = default;
  explicit TMUMdex_base(const uint64_t initial_pairs,
                        const uint64_t initial_mums) :
      pairs_{initial_pairs}, mums_{initial_mums}, bases_{initial_pairs},
    index_{initial_pairs} {}
  TMUMdex_base(const uint64_t pairs_size, const uint64_t mums_size_,
           const uint64_t bases_size, const uint64_t index_size) :
      pairs_{pairs_size}, mums_{mums_size_}, bases_{bases_size},
    index_{index_size} {}
  explicit TMUMdex_base(const std::string & name) :
      pairs_{pairs_name(name)}, mums_{mums_name(name)}, bases_(name),
    index_{index_name(name), false} {}

  // deleted
  TMUMdex_base(const TMUMdex_base &) = delete;
  TMUMdex_base & operator=(const TMUMdex_base &) = delete;

  // pairs
  uint64_t n_pairs() const { return pairs_.size(); }
  Pair pair(const uint64_t pair_index) const { return pairs_[pair_index]; }
  Pair pair(const MUMindex m) const { return pairs_[m.pair_index()]; }
  const VECTOR<Pair> & pairs() const { return pairs_; }
  const Pair * begin() const { return pairs_.begin(); }
  const Pair * end() const { return pairs_.end(); }

  // mums
  uint64_t n_mums() const { return mums_.size(); }
  const VECTOR<MUM> & mums() const { return mums_; }
  MUM mum(const uint64_t m) const { return mums_[m]; }
  MUM mum(const MUMindex m) const {
    return mums_[pairs_[m.pair_index()].mums_start() + m.mum_in_pair_index()];
  }
  uint64_t mum_index(const MUMindex m) const {
    return pairs_[m.pair_index()].mums_start() + m.mum_in_pair_index();
  }
  MUM sorted_mum(const uint64_t m) const { return mums_[index_[m]]; }
  const MUM * mums_begin() const { return mums_.begin(); }
  const MUM * mums_end() const { return mums_.end(); }
  const MUM * mums_begin(const uint64_t pair_index) const {
    return mums_.begin() + mums_start(pair_index);
  }
  const MUM * mums_end(const uint64_t pair_index) const {
    return mums_.begin() + mums_stop(pair_index);
  }
  const MUM * mums_begin(const Pair pair_) const {
    return mums_.begin() + pair_.mums_start();
  }
  uint64_t mums_start(const uint64_t pair_index) const {
    return pairs_[pair_index].mums_start();
  }
  uint64_t mums_stop(const uint64_t pair_index) const {
    if (pair_index + 1 == pairs_.size()) {
      return mums_.size();
    }
    return pairs_[pair_index + 1].mums_start();
  }
  unsigned int n_mums(const uint64_t pair_index) const {
    return static_cast<unsigned int>(
        mums_stop(pair_index) - mums_start(pair_index));
  }
  const MUM * first_mum_read_1(const uint64_t pair_index) const {
    if (pairs_[pair_index].has_mums()) {
      return mums_begin(pair_index);
    }
    return nullptr;
  }
  const MUM * first_mum_read_2(const uint64_t pair_index) const {
    if (pairs_[pair_index].has_mums()) {
      for (const MUM * mum_{mums_begin(pair_index)};
           mum_ != mums_end(pair_index); ++mum_) {
        if (mum_->read_2()) return mum_;
      }
    }
    return nullptr;
  }

  // bases
  const AllBases & bases() const { return bases_; }
  const ExtraBases & extra_bases() const { return bases_.extra(); }

  // index
  const VECTOR<MUMindex> & index() const { return index_; }
  typename VECTOR<MUMindex>::const_iterator lower_bound(
      const uint64_t chromosome,
      const uint64_t position) const {
    const MUM search_for{chromosome, position};
    return std::lower_bound(index_.begin(), index_.end(), search_for,
                            lessMUM_index<VECTOR>(mums_, pairs_));
  }
  typename VECTOR<MUMindex>::const_iterator lower_bound(
      const PosInfo position) const {
    return lower_bound(position.chr, position.pos);
  }
  MUMRegion region() const { return MUMRegion{index_.begin(), index_.end()}; }
  MUMRegion region(const std::string & region_arg,
                   const ChromosomeIndexLookup & chr_lookup) const {
    const PositionRegion region_{region_arg, chr_lookup};
    return MUMRegion{lower_bound(region_.start_chromosome,
                                 region_.start_position),
          lower_bound(region_.stop_chromosome, region_.stop_position)};
  }
  MUMRegion region(const unsigned int start_chromosome,
                   const unsigned int start_position,
                   const unsigned int stop_chromosome,
                   const unsigned int stop_position) const {
    return MUMRegion{lower_bound(start_chromosome, start_position),
          lower_bound(stop_chromosome, stop_position)};
  }

  // saving
  void save_reference_name(const std::string & mumdex_name,
                           const std::string & ref_file_name) const {
    std::string file_name{ref_name(mumdex_name)};
    std::ofstream ref_name_file(file_name.c_str());
    if (!ref_name_file)
      throw Error("Problem saving reference name in") << file_name;
    ref_name_file << ref_file_name << std::endl;
  }
  void clear() {
    pairs_.clear();
    mums_.clear();
    bases_.clear();
    index_.clear();
  }

  void save(const std::string & name) const {
    mkdir(name);
    const mumdex_names names{mumdex_names(name)};
    pairs_.save(names[0]);
    mums_.save(names[1]);
    bases_.save(names[2], names[3]);
  }
  // rearrange according to pair sorting
  void rearrange(const TMUMdex_base & mumdex) {
    for (uint64_t i{0}; i != mumdex.index_.size(); ++i) {
      const uint64_t p{mumdex.index_[i]};
      const Pair & pair_{mumdex.pairs_[p]};
      pairs_.push_back(pair_);
      pairs_.back().mums_start(mums_.size());
      bases_.push_back(mumdex.bases_.bases(p));
      for (uint64_t m{pair_.mums_start()}; m != mumdex.mums_stop(p); ++m) {
        mums_.push_back(mumdex.mums_[m]);
      }
    }
  }

 private:
  friend class MUMdexMerger;
  friend class MUMdexMergerNew;
  // friend class MUMdex_build;
  VECTOR<Pair> & pairs() { return pairs_; }
  AllBases & bases() { return bases_; }
  static void close_files(const std::vector<std::FILE *> & files) {
    for (std::FILE * file : files) {
      if (fclose(file)) {
        throw Error("problem closing mumdex file");
      }
    }
  }
  ExtraBasesSaveInfo write(const std::vector<std::FILE *> & files) const {
    pairs_.write(files[0]);
    mums_.write(files[1]);
    return bases_.write(files[2], files[3]);
  }
  static std::vector<std::FILE *> open_for_write(const std::string & name) {
    const mumdex_names names{name};
    std::vector<std::FILE *> files;
    files.reserve(names.size());
    for (const std::string & file_name : names) {
      std::FILE * file{std::fopen(file_name.c_str(), "wb+")};
      if (file == nullptr) throw Error("Problem opening file") << file_name;
      files.push_back(file);
    }
    return files;
  }
  template <class MUMDEX>
  void emplace_back(const MUMDEX & mumdex, const uint64_t p,
                    const uint64_t mum_offset, const uint64_t extra_offset,
                    const MUMDEX * & last_mumdex, uint64_t & last_p,
                    const bool mark_dupes) {
    using OrderedMums = std::array<const MUM * const, 2>;
    const OrderedMums last_ordered_mums = last_mumdex ?
        last_mumdex->pair(last_p).ordered_first_mums(
            last_mumdex->mums_begin()) :
        std::array<const MUM * const, 2>{{nullptr, nullptr}};
    const MUM * const last_mum_1{last_mumdex ? last_ordered_mums[0] : nullptr};
    const MUM * const last_mum_2{last_mumdex ? last_ordered_mums[1] : nullptr};

    Pair pair_{mumdex.pair(p)};
    const OrderedMums ordered_mums =
        pair_.ordered_first_mums(mumdex.mums_begin());
    const MUM * const mum_1{ordered_mums[0]};
    const MUM * const mum_2{ordered_mums[1]};

    if (mum_1 && last_mum_1) {
      if (mum_2 && last_mum_2) {
        const Pair last_pair{last_mumdex->pair(last_p)};
        if (last_pair.length(0) == pair_.length(0) &&
            last_pair.length(1) == pair_.length(1) &&
            mum_1->read_position0(pair_.length(mum_1->read_2())) ==
            last_mum_1->read_position0(
                last_pair.length(last_mum_1->read_2())) &&
            mum_2->read_position0(pair_.length(mum_2->read_2())) ==
            last_mum_2->read_position0(
                last_pair.length(last_mum_2->read_2())) &&
            mum_1->chromosome() == last_mum_1->chromosome() &&
            mum_2->chromosome() == last_mum_2->chromosome()) {
          if (mark_dupes) pair_.dupe(true);
        }
      }
    }
    last_mumdex = &mumdex;
    last_p = p;
    pair_.mums_start(mum_offset + mums_.size());
    pairs_.push_back(pair_);
    bases_.push_back(mumdex.bases_.bases(p), extra_offset);
    mums_.insert_back(mumdex.mums_begin(p), mumdex.mums_end(p));
  }

 protected:
  VECTOR<Pair> pairs_{};
  VECTOR<MUM> mums_{};
  AllBases bases_{};
  VECTOR<MUMindex> index_{};  // pairs for parts, mums when complete
};

using MUMdex_build_base = TMUMdex_base<GrowingVector>;

// template <template <class ...> class VECTOR>
template <class Sequence>
class MUMdex_build : public MUMdex_build_base {
 public:
  // construction
  MUMdex_build(const Sequence & ref_arg, const std::string & name_arg,
               const uint64_t max_pairs_arg = 1000000,
               const uint64_t max_mums_arg = 16000000) :
      MUMdex_build_base{max_pairs_arg, max_mums_arg},
    ref_{&ref_arg}, name{name_arg}, n_out{0} {}

  MUMdex_build(const MUMdex_build &) = delete;
  MUMdex_build & operator=(const MUMdex_build &) = delete;

  // growing
  template<class match_t>
  void emplace_back(const std::vector<match_t> & matches_1,
                    const std::string & query_1,
                    const bool bad_1,
                    const std::vector<match_t> & matches_2,
                    const std::string & query_2,
                    const bool bad_2) {
    index_.emplace_back(pairs_.size());
    const uint64_t n_matches{matches_1.size() + matches_2.size()};
    pairs_.emplace_back(mums_.size(),
                        query_1.size(), bad_1,
                        query_2.size(), bad_2,
                        n_matches);

    const std::vector<match_t> * matches[2]{&matches_1, &matches_2};
    const std::string * queries[2]{&query_1, &query_2};
    std::string extra_bases_;
    for (const bool read_2 : {false, true}) {
      unsigned int query_pos{0};
      for (unsigned int m{0}; m != matches[read_2]->size(); ++m) {
        const match_t & mum_{(*matches[read_2])[m]};
        bool flipped{false};
        uint64_t rpos{mum_.ref};
        if (rpos >= ref_->N) {  // for flipped non rcref
          rpos -= ref_->N;
          flipped = true;
        }
        std::vector<uint64_t>::const_iterator it =
            upper_bound(ref_->startpos.begin(), ref_->startpos.end(), rpos);
        unsigned int chromosome{static_cast<unsigned int>(
            it - ref_->startpos.begin() - 1)};
        unsigned int position{static_cast<unsigned int>(rpos - *--it)};
        if (ref_->rcref) {
          if ((chromosome % 2) == 1) {
            position = static_cast<unsigned int>(
                ref_->sizes[chromosome] - position - mum_.len);
            flipped = true;
          }
          chromosome /= 2;
        }
        const unsigned int offset{static_cast<unsigned int>(mum_.offset)};
        const bool last_hit{(m + 1 == matches[read_2]->size()) &&
              (read_2 || matches[1]->empty())};
        const bool touches_end{offset + mum_.len == queries[read_2]->size()};
        mums_.emplace_back(chromosome, position,
                           offset, mum_.len, flipped,
                           read_2, last_hit, touches_end);
        while (query_pos < mum_.offset) {
          extra_bases_ += (*queries[read_2])[query_pos++];
        }
        query_pos = static_cast<unsigned int>(mum_.offset + mum_.len);
      }
      while (query_pos != (*queries[read_2]).size()) {
        extra_bases_ += (*queries[read_2])[query_pos++];
      }
    }
    bases_.push_back(extra_bases_);
  }

  // saving
  void save() const { MUMdex_build_base::save(name); }
  void save(const std::string & name_arg) const {
    MUMdex_build_base::save(name_arg);
  }
  std::string sort_and_save_part(MUMdex_build_base & sorted) {
    std::sort(index_.begin(), index_.end(),
              lessPair_index<GrowingVector>(pairs_, mums_));
    ++n_out;
    std::ostringstream part_name;
    part_name << name << "." << n_out;
    sorted.rearrange(*this);
    sorted.save(part_name.str());
    save_reference_name(part_name.str(), ref_->ref_fasta);
    sorted.clear();
    return part_name.str();
  }

 private:
  const Sequence * const ref_;
  std::string name{""};
  uint64_t n_out{0};
};

template <class MUMdex>
class TAnchors {
 public:
  using MUMRegion = typename MUMdex::MUMRegion;
  class AnchorIterator {
   public:
    using const_iterator = typename MUMRegion::const_iterator;
    AnchorIterator(const TAnchors & anchors_, const const_iterator start) :
        anchors{anchors_}, current{start} {
          if (current != anchors.region.end() && !anchors.pass(current)) {
            ++(*this);
          }
        }
    bool operator!=(const AnchorIterator & other) const {
      return current != other.current;
    }
    AnchorIterator & operator++() {
      while (++current != anchors.region.end()) {
        if (anchors.pass(current)) {
          break;
        }
      }
      return *this;
    }
    MUMindex operator*() const {
      return *current;
    }

   private:
    const TAnchors & anchors;
    const_iterator current;
  };
  using const_iterator = AnchorIterator;

  TAnchors(const MUMdex & mumdex_,
           const MUMRegion region_,
           const unsigned int chromosome_,
           const unsigned int position_,
           const bool high_) :
      mumdex{mumdex_}, region{region_},
    chromosome{chromosome_}, position{position_}, high{high_} {
      if (0) std::cerr << "counting anchors at "
                       << mumdex_.reference().name(chromosome) << " "
                       << position << " " << high << " in "
                       << region.end() - region.begin() << " mums"
                       << std::endl;
    }

  AnchorIterator begin() const { return AnchorIterator{*this, region.begin()}; }
  AnchorIterator end() const { return AnchorIterator{*this, region.end()}; }

  // template just to avoid redefinition rule for stand-alone function
  // that must be defined out of class (due to using mumdex defined below)
  template <class Iter>
  bool pass(const Iter current) const {
    const MUM mum{mumdex.mum(*current)};
    if (mum.chromosome() != chromosome) return false;
    if (mum.anchor_position(high) != position) return false;
    // Make sure anchor is not at end of read
    if (high) {
      if (mum.flipped()) {
        if (!mum.offset()) {
          return false;
        }
      } else {
        if (mum.touches_end()) {
          return false;
        }
      }
    } else {
      if (mum.flipped()) {
        if (mum.touches_end()) {
          return false;
        }
      } else {
        if (!mum.offset()) {
          return false;
        }
      }
    }
#if 0
      // make sure anchor is not an N anchor
      const std::array<std::string, 2> sequences(
          mumdex.sequences(current->pair_index()));
      const bool end_of{high != mum.flipped()};
      const unsigned int next_base{mum.offset() + (end_of ? mum.length() : -1)};
      if (sequences[mum.read_2()][next_base] == 'N') return false;
#endif
    return true;
  }

 private:
  const MUMdex & mumdex;
  const MUMRegion region;
  const unsigned int chromosome;
  const unsigned int position;
  const bool high;
};

template <template <class ...> class VECTOR>
class TMUMdex : public TMUMdex_base<VECTOR> {
 public:
  using Base = TMUMdex_base<VECTOR>;
  using Base::pairs_;
  using Base::mums_;
  using Base::bases_;
  using Base::index_;
  using MUMRegion = TMUMRegion<VECTOR>;
  using Anchors = TAnchors<TMUMdex<VECTOR>>;

  // construction
  explicit TMUMdex(const std::string & name_arg,
                   const Reference * ref_arg = nullptr) :
      TMUMdex_base<VECTOR>{name_arg},
    ref_{ref_arg ? ref_arg : new Reference(saved_ref_name(name_arg))},
    own_ref{ref_arg ? false : true} {}
  TMUMdex(const std::string & name_arg, const Reference & ref_arg) :
      TMUMdex_base<VECTOR>{name_arg}, ref_{&ref_arg} {}
  TMUMdex(TMUMdex && other) noexcept : TMUMdex_base<VECTOR>{std::move(other)},
    ref_{other.ref_}, own_ref{other.own_ref} {
    other.own_ref = false;
  }
  ~TMUMdex() { if (own_ref) delete ref_; }

  TMUMdex(const TMUMdex &) = delete;
  TMUMdex & operator=(const TMUMdex &) = delete;

  // info
  uint64_t bytes() const {
    return sizeof(TMUMdex) +
        pairs_.bytes() + mums_.bytes() + bases_.bytes() + index_.bytes();
  }

  const Reference & reference() const { return *ref_; }

  const MUM * longest_MUM(const uint64_t p) const {
    const Pair pair_{this->pair(p)};
    if (pair_.dupe()) return nullptr;
    if (!pair_.has_mums()) return nullptr;
    uint64_t longest{pair_.mums_start()};
    for (uint64_t m{pair_.mums_start() + 1}; m != this->mums_stop(p); ++m) {
      if (this->mum(m).length() > this->mum(longest).length()) {
        longest = m;
      }
    }
    return mums_.begin() + longest;
  }

  template <class MAPPABILITY>
  const MUM * cn_MUM(const uint64_t p,
                     const MAPPABILITY & mappability,
                     const unsigned int min_length,
                     const unsigned int min_excess,
                     const unsigned int max_mappability) const {
    const MUM * longest_mum{longest_MUM(p)};
    if (longest_mum == nullptr) return nullptr;
    const MUM mum_{*longest_mum};
    if (mum_.length() < min_length) return nullptr;
    const unsigned int abspos{reference().abspos(mum_.chromosome(),
                                                 mum_.position0())};
    const unsigned int min_map{mappability.min(abspos, mum_.length())};
    if (min_map > mum_.length()) {
      throw Error("Unexpected min map bad");
    }
    if (min_map > max_mappability) return nullptr;
    const unsigned int excess_map{mum_.length() - min_map};
    if (excess_map < min_excess) return nullptr;
    return longest_mum;
  }

  using Annotation = std::pair<MUM, std::string>;
  void pair_view(std::ostream & out, const uint64_t p,
                 const std::vector<Annotation> & annotations =
                 std::vector<std::pair<MUM, std::string>>()) const {
    const Pair pair_{pairs_[p]};
    std::array<std::string, 2> seqs(sequences(p));
    reverse_complement(&seqs[1]);
    const MUM * best1{mums_.end()};
    const MUM * best2{mums_.end()};
    unsigned int best_length{0};
    for (const MUM * mum1{this->mums_begin(p)};
         mum1 != this->mums_end(p); ++mum1) {
      if (mum1->last_hit()) break;
      if (mum1->read_2()) break;
      for (const MUM * mum2{mum1 + 1}; mum2 != this->mums_end(p); ++mum2) {
        if (!mum2->read_2()) continue;
        if (mum1->close_by(*mum2, 1000)) {
          const unsigned int length{mum1->length() + mum2->length()};
          if (length > best_length) {
            best1 = mum1;
            best2 = mum2;
            best_length = length;
          }
        }
      }
    }
    std::vector<MUM> to_show;
    for (const MUM * mump{this->mums_begin(p)};
         mump != this->mums_end(p); ++mump) {
      if (mump->read_2()) break;
      to_show.push_back(*mump);
    }
    while (to_show.size()) {
      unsigned int written{0};
      std::vector<MUM> to_save;
      for (unsigned int m{0}; m != to_show.size(); ++m) {
        const MUM smum{to_show[m]};
        if (smum.offset() >= written) {
          while (smum.offset() > written) {
            out << ' ';
            ++written;
          }
          std::ostringstream pos;
          pos << '[';
          for (const Annotation & annotation : annotations) {
            if (annotation.first == smum) {
              pos << annotation.second;
            }
          }
          pos << ((best_length && smum == *best1) ? "*" : "")
              << reference().name(smum.chromosome())
              << (smum.flipped() ? '-' : '+')
              << smum.read_position1(pair_.length(0))
              << " " << smum.offset() << " " << smum.length();
          std::string pos_string{pos.str()};
          if (pos_string.size() < smum.length()) {
            while (pos_string.size() + 1 < smum.length()) {
              if (pos_string.size() % 2) {
                pos_string += ' ';
              } else {
                pos_string.insert(1, 1, ' ');
              }
            }
            pos_string += ']';
          } else {
            pos_string.insert(smum.length() - 1, 1, ']');
          }
          out << pos_string;
          written += pos_string.size();
        } else {
          to_save.push_back(smum);
        }
      }
      if (written) out << std::endl;
      to_save.swap(to_show);
    }
    int offset{0};
    for (unsigned int r{0}; r != seqs.size(); ++r) {
      if (r) {
        if (best_length) {
          if (best1->flipped() != best2->flipped()) {
            offset = (best1->flipped() ? -1 : 1) *
                (best2->read_position0(pair_.length(1)) -
                 best1->read_position0(pair_.length(0)));
            out << " " << offset << std::endl;
            if (offset >= 0 && offset < 150) {
              std::ostringstream disagreement;
              bool disagree{false};
              for (int o{0}; o != offset; ++o) {
                disagreement << ' ';
              }
              // sign problem below with offset?
              for (unsigned int b = offset; b < seqs[0].size(); ++b) {
                if (b - offset >= seqs[1].size()) break;
                if (seqs[0][b] != seqs[1][b - offset]) {
                  disagree = true;
                  disagreement << 'X';
                } else {
                  disagreement << ' ';
                }
              }
              if (disagree) {
                out << disagreement.str();
                out << std::endl;
              }
              for (int o{0}; o != offset; ++o) {
                out << ' ';
              }
            } else {
              offset = 0;
            }
          } else {
            out << " ?" << std::endl;
          }
        } else {
          out << " ?" << std::endl;
        }
      }
      if (best_length) {
        const MUM best{r ? *best2 : *best1};
        const bool flip{static_cast<bool>(r) != best.flipped()};
        const unsigned int length{pair_.length(r)};
        const int rpos{best.read_position0(length)};
        for (unsigned int b{0}; b != seqs[r].size(); ++b) {
          // narrowing sign problem?
          const int pos = flip ? rpos + length - b - 1 : rpos + b;
          const char base{flip ? complement(seqs[r][b]) : seqs[r][b]};
          if (pos >= 0 && static_cast<unsigned int>(pos) <
              reference().size(best.chromosome()) &&
              reference()[best.chromosome()][pos] != base) {
            out << seqs[r][b];
          } else {
            out << static_cast<char>(tolower(seqs[r][b]));
          }
        }
      } else {
        out << seqs[r];
      }
    }
    out << std::endl;
    for (const MUM * mump{this->mums_begin(p)};
         mump != this->mums_end(p); ++mump) {
      if (!mump->read_2()) continue;
      to_show.push_back(*mump);
    }
    reverse(to_show.begin(), to_show.end());
    while (to_show.size()) {
      unsigned int written{0};
      std::vector<MUM> to_save;
      for (unsigned int m{0}; m != to_show.size(); ++m) {
        const MUM smum{to_show[m]};
        const unsigned int moffset{offset + pair_.length(1) - smum.offset() -
              smum.length()};
        if (moffset >= written) {
          while (moffset > written) {
            out << ' ';
            ++written;
          }
          std::ostringstream pos;
          pos << '[';
          for (const Annotation & annotation : annotations) {
            if (annotation.first == smum) {
              pos << annotation.second;
            }
          }
          pos << ((best_length && smum == *best2) ? "*" : "")
              << reference().name(smum.chromosome())
              << (smum.flipped() ? '+' : '-')
              << smum.read_position1(pair_.length(1))
              << " " << smum.offset() << " " << smum.length();

          std::string pos_string{pos.str()};
          if (pos_string.size() < smum.length()) {
            while (pos_string.size() + 1 < smum.length()) {
              if (pos_string.size() % 2) {
                pos_string += ' ';
              } else {
                pos_string.insert(1, 1, ' ');
              }
            }
            pos_string += ']';
          } else {
            pos_string.insert(smum.length() - 1, 1, ']');
          }
          out << pos_string;
          written += pos_string.size();
        } else {
          to_save.push_back(smum);
        }
      }
      if (to_save.size() && written) out << std::endl;
      to_save.swap(to_show);
    }
  }
  std::string pair_view(const uint64_t p) const {
    std::ostringstream out;
    pair_view(out, p);
    return out.str();
  }


  Anchors low_anchors(const unsigned int chromosome,
                      const unsigned int position) const {
    return Anchors{*this, MUMRegion{this->lower_bound(chromosome, position),
            this->lower_bound(chromosome, position + 1)},
          chromosome, position, false};
  }
  Anchors high_anchors(const unsigned int chromosome,
                      const unsigned int position,
                      const unsigned int max_read_length = 155,
                      const unsigned int min_mum_length = 1) const {
    const unsigned int min_position{position > max_read_length ?
          position - max_read_length : 0};
    const unsigned int max_position{position > min_mum_length ?
          position - min_mum_length : 0};
    return Anchors{*this, MUMRegion{this->lower_bound(chromosome, min_position),
            this->lower_bound(chromosome, max_position + 1)},
          chromosome, position, true};
  }
  Anchors anchors(const unsigned int chromosome,
                  const unsigned int position,
                  const bool high,
                  const unsigned int max_read_length = 155,
                  const unsigned int min_mum_length = 1) const {
    if (high) {
      return high_anchors(chromosome, position, max_read_length,
                          min_mum_length);
    } else {
      return low_anchors(chromosome, position);
    }
  }

  std::array<std::string, 2> sequences(const uint64_t p) const {
    std::array<std::string, 2> result;
    const std::string extra_bases_{bases_.bases(p)};
    unsigned int extra_i{0};
    const Pair & pair_{pairs_[p]};
    uint64_t mum_index_{pair_.mums_start()};
    const uint64_t mum_stop_index{this->mums_stop(p)};
    for (const bool read_2 : {false, true}) {
      unsigned int read_i{0};
      for (; mum_index_ != mum_stop_index; ++mum_index_) {
        const MUM mum_{mums_[mum_index_]};
        if (mum_.read_2() != read_2) break;
        const unsigned int offset{mum_.offset()};
        while (read_i < offset) {
          result[read_2] += extra_bases_[extra_i++];
          ++read_i;
        }
        for (; read_i < offset + mum_.length(); ++read_i) {
          if (mum_.flipped()) {
            result[read_2] += complement(
                (*ref_)[mum_.chromosome()]
                [mum_.position0() + offset + mum_.length() - read_i - 1]);
          } else {
            result[read_2] += (*ref_)[mum_.chromosome()]
                [mum_.position0() + read_i - offset];
          }
        }
      }
      while (read_i != pair_.length(read_2)) {
        result[read_2] += extra_bases_[extra_i++];
        ++read_i;
      }
    }
    return result;
  }

  std::string adjacent_sequence(const uint64_t pair_index,
                                const MUM mum_, const bool high,
                                const unsigned int adj_len) const {
    std::string seq;
    const std::array<std::string, 2> seqs(sequences(pair_index));
    for (unsigned int a{0}; a != adj_len; ++a) {
      const bool right{high != mum_.flipped()};
      const unsigned int offset{mum_.offset()};  // to save space below
      const char base{right ?
            (offset + mum_.length() + a <
             this->pair(pair_index).length(mum_.read_2()) ?
             seqs[mum_.read_2()][offset + mum_.length() + a] : 'x') :
            (offset > adj_len - a ?
             seqs[mum_.read_2()][offset - adj_len + a] : 'x')};
      seq += base;
    }
    if (mum_.flipped()) reverse_complement(&seq);
    return seq;
  }

 private:
  const Reference * const ref_;
  bool own_ref{false};
};

using MemoryMUMdex = TMUMdex<MemoryVector>;
using PreMappedMUMdex = TMUMdex<PreMappedVector>;
using UnMappedMUMdex = TMUMdex<UnMappedVector>;
using FileMUMdex = TMUMdex<FileVector>;
using MUMdex = UnMappedMUMdex;

}  // namespace paa

#endif  // PAA_MUMDEX_H
```
