## Supplementary material for "G-Graph: An interactive genomic graph viewer": MUMdex Source Code for G-Graph: mummer.cpp

```
//
// mummer
//
// map reads to produce mumdex parts
//
// Modifications of sparseMEM Copyright Peter Andrews 2013-2015 @ CSHL
//

#include <unistd.h>
#include <limits.h>
#include <getopt.h>

#include <exception>
#include <iostream>
#include <string>

#include "error.h"
#include "longSA.h"
#include "query.h"
#include "strings.h"

using std::cerr;
using std::endl;
using std::exception;
using std::string;

using paa::Error;
using paa::Readers;
using paa::ReadersArgs;
using paa::ReadPairsArgs;
using paa::SAArgs;
using paa::longReadPairs;
using paa::longSA;
using paa::shortReadPairs;
using paa::shortSA;

// Class to handle options parsing and to deliver options to classes
class Args : public SAArgs, public ReadPairsArgs, public ReadersArgs {
 public:
  Args(int argc_, char * argv_[]);
  bool verbose;
 private:
  void check_integer_sizes() const;
  void usage(const string & prog) const;
  int argc;
  char * * argv;
  Args(const Args &) = delete;
  Args & operator=(const Args &) = delete;
};

int main(int argc, char* argv[])  try {
  // Process options and args.
  const Args args(argc, argv);

  if (args.ref_args.rcref) {
    // Create suffix array
    const longSA sa(args);

    if (args.n_input == 0) return 0;

    // Readers read queries and pass them off to a ReadPair in pairs
    Readers readers(args);

    // ReadPairs manages ReadPair worker threads
    longReadPairs pairs(args, sa, readers);
  } else {
    // Create suffix array
    const shortSA sa(args);

    if (args.n_input == 0) return 0;

    // Readers read queries and pass them off to a ReadPair in pairs
    Readers readers(args);

    // ReadPairs manages ReadPair worker threads
    shortReadPairs pairs(args, sa, readers);
  }
  return 0;
} catch (Error & e) {
  cerr << "paa::Error:" << endl;
  cerr << e.what() << endl;
  return 1;
} catch (exception & e) {
  cerr << "std::exception" << endl;
  cerr << e.what() << endl;
  return 1;
} catch (...) {
  cerr << "unknown exception was caught" << endl;
  return 1;
}

Args::Args(int argc_, char * argv_[])
    : SAArgs(), ReadPairsArgs(), ReadersArgs(),
      verbose(false), argc(argc_), argv(argv_) {
  // Collect arguments from the command line. These options are allowed.
  struct option long_options[] = {
    {"l", 1, nullptr, 0},  // 0
    {"rcref", 0, nullptr, 0},  // 1
    {"samin", 0, nullptr, 0},  // 2
    {"fastq", 0, nullptr, 0},  // 3
    {"fastqs", 0, nullptr, 0},   // 4
    {"threads", 1, nullptr, 0},  // 5
    {"verbose", 0, nullptr, 0},  // 6
    {"uncached", 0, nullptr, 0},  // 7
    {"normalmem", 0, nullptr, 0},  // 8
    {"maxmem", 1, nullptr, 0},  // 9 not implemented
    {"optional", 1, nullptr, 0},  // 10
    {"qthreads", 1, nullptr, 0},  // 11
    {nullptr, 0, nullptr, 0}
  };
  fastq = true;
  fastqs = true;
  while (true) {
    int longindex = -1;
    int c = getopt_long_only(argc, argv, "", long_options, &longindex);
    if (c == -1) {
      break;  // Done parsing flags.
    } else if (c == '?') {  // If the user entered junk, let him know.
      cerr << "Invalid arguments." << endl;
      usage(argv[0]);
    } else {
      // Branch on long options.
      switch (longindex) {
        case 0: min_len = atoi(optarg); break;
        case 1: ref_args.rcref = true; break;
        case 2: sam_in = true; fastq = false; fastqs = false; break;
        case 3: fastq = true; fastqs = false; break;
        case 4: fastqs = true; fastq = true; break;
        case 5:
        case 11:
          max_n_threads = atoi(optarg);
          mappability_threads = max_n_threads;
          break;
        case 6:  // This is messy - a design flaw
          ref_args.verbose = true;
          SAArgs::verbose = true;
          ReadPairsArgs::verbose = true;
          ReadersArgs::verbose = true;
          verbose = true;
          break;
        case 7: paa::read_ahead = false; break;
        case 8: paa::memory_mapped = false; break;
        case 9:
          max_mem = atof(optarg); break;
        case 10:
          optional.push_back(optarg);
          pass_optional = true;
          break;
        default: break;
      }
    }
  }

  // Validate arguments
  argc -= optind;
  if (argc < 1) {
    cerr << "There are too few arguments" << endl;
    usage(argv[0]);
  }
  if (pass_optional && fastq)
    throw Error("-optional cannot be used with -fastq");
  if (!fastq && !sam_in)
    throw Error("You must use either -fastq or -fastqs or -samin");
  if (fastq && sam_in) throw Error("-fastq cannot be used with -samin");
  char * * args = argv + optind;
  ref_args.ref_fasta = *args;
  n_input = argc - 1;
  input = args + 1;
  // check_integer_sizes();
  n_threads = max_n_threads;  // do something better later
}

// Display proper command line usage
void Args::usage(const string & prog) const {
  cerr << "Usage: " << prog <<
      " [options] <reference-file> <query-file> ...\n"
      ""
      /*
      "-mum         "
      "  compute maximal matches that are unique in both sequences\n"
      "-mumreference"
      "  compute maximal matches that are unique in the reference-\n"
      "             "
      "  sequence but not necessarily in the query-sequence (default)\n"
      "-mumcand       same as -mumreference\n"
      "-maxmatch    "
      "  compute all maximal matches regardless of their uniqueness\n"
      */
      "-l N           set the minimum length of a match to N\n"
      "               if not set, the default value is 1\n"
      "-verbose       output diagnostics and progress to stderr\n"
      "-samin         input in SAM format\n"
      "-fastq         input in fastq format (one file read1/read2 alternate)\n"
      "-fastqs        input in fastq format two or more files read1/read2 ...\n"
      "               fastqs is the default\n"
      "-threads N     number of threads to use\n"
      "-rcref         reverse complement reference (you should use this)\n"
      "-cached        when memory mapped, does not pre-fault pages\n"
      "-normalmem     turn off memory mapping\n"
      "-optional FMT  pass optional SAM/fastq fields through\n"
      "  Accepted FMT strings ([xyz] means xyz is optional):\n"
      "  (note format string must be properly quoted if you use a shell)\n"
      "  ERR|N[|S|R]     quality scores with maximum read length N, \n"
      "                  starting scale S (33), range R (42)\n"
      "  FIX|N|KEY|ALPHA String of length N with alphabet ALPHA, field KEY\n"
      "  VAR|N|KEY|ALPHA String up to length N with alphabet ALPHA, field KEY\n"
      "  TAG|N|KEY|S|O   Tag of length N, start S, read 2 start O, field KEY\n";
  // "  INT|N|KEY       Integer up to but excluding N, field KEY\n";

  exit(1);
}
```
