## Supplementary material for "G-Graph: An interactive genomic graph viewer": MUMdex Source Code for G-Graph: mview.cpp

```
//
// mview.cpp
//
// gui to explore MUMdex mappings
//
// Copyright 2017 Peter Andrews @ CSHL
//

#include <exception>
#include <fstream>
#include <iostream>
#include <map>
#include <sstream>
#include <string>
#include <vector>

#include "error.h"
#include "mumdex.h"
#include "mview.h"
#include "x11plot.h"

using std::cerr;
using std::endl;
using std::exception;
using paa::Error;
using paa::Geometry;

int main(int argc, char* argv[]) try {
  // Process optional command line arguments
  --argc;
  int width{1000};
  int height{800};
  int x_off{10};
  int y_off{40};
  while (argc) {
    if (argv[1][0] == '-') {
      const std::string option{argv[1]};
      if (option == "--geometry") {
        const std::string geometry{argv[2]};
        std::istringstream geometry_stream{geometry.c_str()};
        char dummy;
        geometry_stream >> width >> dummy >> height >> x_off >> y_off;
        if (!geometry_stream) {
          throw Error("Problem parsing geometry") << geometry;
        }
        if (false)
          std::cerr << "Geometry set to width " << width << " height " << height
                    << " x offset " << x_off << " y offset " << y_off
                    << std::endl;
        argv += 2;
        argc -= 2;
      } else {
        throw Error("Unrecognized command line option") << option;
      }
    } else {
      break;
    }
  }

  if (argc < 3)
    throw Error("usage: mview chr pos mumdex .."
                "   or  mview cand_list samples_dir pop_file");

  if (atoi(argv[2])) {
    // Process arguments
    const std::string chr{argv[1]};
    const unsigned int pos{static_cast<unsigned int>(atoi(argv[2]))};
    argv += 2;
    argc -= 2;

    const std::vector<std::string> mumdex_names{[&argv, &argc]() {
        std::vector<std::string> result;
        result.reserve(argc);
        while (argc) {
          result.emplace_back(argv[1]);
          --argc;
          ++argv;
        }
        return result;
      }()};

    const std::vector<paa::MUMDEX> mumdexes{[&mumdex_names]() {
        std::vector<paa::MUMDEX> result;
        result.reserve(mumdex_names.size());
      for (const std::string & name : mumdex_names) {
        result.emplace_back(name);
      }
      return result;
      }()};

    // App to display multiple windows
    paa::X11App app;

    // Read viewer class
    paa::X11MUMdexViewer & viewer{app.create<paa::X11MUMdexViewer>(
        mumdex_names, mumdexes,
        Geometry{{width, height}, {x_off, y_off}})};

    viewer.set_position(chr, pos);

    // Run the app
    app.run();

  } else {
    const std::string cand_file_name{argv[1]};
    const std::string samples_dir{argv[2]};
    const std::string pop_file_name{argv[3]};
    paa::X11CandidateViewer viewer{cand_file_name, samples_dir, pop_file_name,
      {{width, height}, {x_off, y_off}}};
  }

  return 0;
} catch (Error & e) {
  cerr << "paa::Error:" << endl;  cerr << e.what() << endl;
  return 1;
} catch (exception & e) {
  cerr << "std::exception" << endl;
  cerr << e.what() << endl;
  return 1;
} catch (...) {
  cerr << "unknown exception was caught" << endl;
  return 1;
}
```
