## Supplementary material for "G-Graph: An interactive genomic graph viewer": MUMdex Source Code for G-Graph: mview.h

```
//
// mview.h
//
// X11 MUMdex viewing
//
// Copyright 2017 Peter Andrews @ CSHL
//

#ifndef PAA_MVIEW_H
#define PAA_MVIEW_H

#include <algorithm>
#include <array>
#include <deque>
#include <exception>
#include <future>
#include <iostream>
#include <map>
#include <mutex>
#include <random>
#include <set>
#include <sstream>
#include <string>
#include <utility>
#include <vector>

#include "bridges.h"
#include "error.h"
#include "mumdex.h"
#include "strings.h"
#include "threads.h"
#include "x11plot.h"

namespace paa {

using MUMDEX = PreMappedMUMdex;
// using MUMDEX = MemoryMUMdex;
// using MUMDEX = FileMUMdex;
// using MUMDEX = MUMdex;

class X11MUMdexViewer : public X11Win {
 public:
  static constexpr int border_width{3};

  X11MUMdexViewer(X11App & app__,
                  const std::vector<std::string> & mumdex_names__,
                  const std::vector<MUMDEX> & mumdexes__,
                  const Geometry geometry__ = {{1200, 1000}, {0, 0}}) :
      X11Win{app__, geometry__},
    mumdex_names_{mumdex_names__},
    mumdexes_{mumdexes__},
    mumdex_{&mumdexes_.front()},
    ref{mumdex_->reference()}, chr_lookup{ref}, mappability{ref},
    read_mutexes{mumdexes_.size()},
    pool{static_cast<unsigned int>(mumdexes_.size())} {
        // Events to watch out for
        XSelectInput(display(), window, StructureNotifyMask | ExposureMask |
                     KeyPressMask | EnterWindowMask | LeaveWindowMask |
                     ButtonPressMask | PointerMotionMask | ButtonReleaseMask);

        // Graphics contexts
        border_gc = create_gc(app.black, app.white, border_width,
                              LineSolid, CapButt, JoinMiter);
        border_fill_gc = create_gc(app.white, app.black, border_width,
                                   LineSolid, CapButt, JoinMiter);
        minor_gc = create_gc(app.black, app.white, 1, LineOnOffDash,
                             CapButt, JoinMiter);
        major_gc = create_gc(app.black, app.white, 2, LineOnOffDash,
                             CapButt, JoinMiter);

        // colors
        std::vector<std::string> levels{"00", "66", "bb", "ff"};
        color_names.push_back("dd/dd/dd");
        for (const std::string & r : levels) {
          for (const std::string & g : levels) {
            for (const std::string & b : levels) {
              if (r == g && g == b) continue;
              color_names.push_back(r + "/" + g + "/" + b);
            }
          }
        }
        X_colors.resize(color_names.size());
        color_gcs.resize(color_names.size());
        for (unsigned int c{0}; c != color_names.size(); ++c) {
          const std::string & color_name{std::string("rgb:") + color_names[c]};
          XColor & color{X_colors[c]};
          if (!XAllocNamedColor(display(), app.colormap, color_name.c_str(),
                                &color, &color))
            throw Error("Could not get color") << color_name;

          color_gcs[c] = create_gc(color.pixel, app.white, 1,
                                   LineSolid, CapButt, JoinMiter);
        }
        char_gcs.resize(256);
        char_gcs['a'] = char_gcs['A'] = color_gcs[7];
        char_gcs['c'] = char_gcs['C'] = color_gcs[59];
        char_gcs['g'] = char_gcs['G'] = color_gcs[23];
        char_gcs['t'] = char_gcs['T'] = color_gcs[46];
        char_gcs['n'] = char_gcs['N'] = gc;

        XColor grey;
        if (!XAllocNamedColor(display(), app.colormap, "rgb:bb/bb/bb",
                              &grey, &grey)) throw Error("Could not get grey");
        mum_gc = create_gc(grey.pixel, app.white);

        // Add radios to master radio list
        for (std::vector<Radio> * radio_vec : {&unnamed_radios})
          for (Radio & radio : (*radio_vec)) radios.push_back(&radio);

        std::cout << "Window id is " << window << std::endl;
        return;
#if 0
        // continuous data pre-loading
        for (unsigned int m{0}; m != mumdexes_.size(); ++m) {
          futures.push_back(
              pool.run([this](const MUMDEX &, std::mutex & mutex_) {
                  while (true) {
                    {
                      std::lock_guard<std::mutex> lock(exited_mutex);
                      if (exited) return;
                    }
                    std::lock_guard<std::mutex> lock(mutex_);
                  }
                }, std::ref(mumdexes_[m]), std::ref(read_mutexes[m])));
        }
#endif
    }

  ~X11MUMdexViewer() {
    {
      std::lock_guard<std::mutex> lock(exited_mutex);
      exited = true;
    }
  }

  X11MUMdexViewer(const X11MUMdexViewer &) = delete;
  X11MUMdexViewer & operator=(const X11MUMdexViewer &) = delete;

  virtual void button_press(const XButtonEvent & event) {
    last_motion = last_press = event;
    moved = false;
    for (Radio * radio : radios) { if (radio->press(event)) return; }

    const int64_t old_index{offset_index};
    XEvent next_event;
    XButtonEvent & this_event{next_event.xbutton};
    do {
      this_event = event;
      bool scroll_event{false};
      if (this_event.button == Button4) {
        --offset_index;
        scroll_event = true;
      }
      if (this_event.button == Button5) {
        ++offset_index;
        scroll_event = true;
      }
      if (!scroll_event) break;
    } while (XCheckTypedWindowEvent(display(), window,
                                   ButtonPressMask, &next_event));

    if (offset_index != old_index) draw();
  }

  virtual void button_release(const XButtonEvent &) {
    for (Radio * radio : radios) { if (radio->release(last_press)) return; }
  }

  virtual void motion(const XMotionEvent & event) {
    moved = true;
    if (XPending(display())) return;

    Event motion_event{Event::X, &event};
    const Point point{event};

    // Status text
    std::string status;
    for (Radio * radio : radios) {
      if (radio->contains(event)) {
        status = radio->description;
        if (!radio->visible()) status += " (inactive)";
        draw_status(status);
        draw_controls();
        return;
      }
    }

    if (invariants_radio) return;
    if (inside) {
      std::ostringstream coordinates;
      const int chrpos{x_to_chrpos(point[0])};
      if (point[1] >= bounds[1][0] &&
          point[1] <
          bounds[1][2] / pixels_per_read * pixels_per_read + bounds[1][0]) {
        const int64_t map_index{y_to_map_index(point[1])};
        if (map_index >= static_cast<int64_t>(displayed_mums.size())) return;
        coordinates << std::setprecision(12)
                    << "position "
                    << ref.name(chromosome) << " " << chrpos
                    << " " << ref[chromosome][chrpos]
                    << " line " << map_index + 1
                    << " of " << std::min(
                        static_cast<int64_t>(bounds[1][2] / pixels_per_read),
                        static_cast<int64_t>(displayed_mums.size()));
        const MUMindex mindex{displayed_mums[map_index]};
        const Pair pair{mumdex_->pair(mindex)};
        const MUM mum{mumdex_->mum(mindex)};
        const int read_length(pair.length(mum.read_2()));
        const int read_pos{mum.read_position0(read_length)};
        if (chrpos >= read_pos &&
            chrpos < read_pos + read_length) {
          coordinates << " read " << (mum.read_2() + 1)
                      << " flipped " << mum.flipped();

          // Determine mum in read that is pointed at
          const std::vector<MUM> mums{read_mums(mindex)};
          std::vector<uint64_t> mum_ids(read_length, mums.size());
          for (uint64_t m{0}; m != mums.size(); ++m) {
            const MUM rm{mums[m]};
            for (unsigned int b{rm.offset()}; b != rm.offset() + rm.length();
                 ++b) {
              if (mum_ids[b] == mums.size()) {
                mum_ids[b] = m;
              }
            }
          }
          const int l_base{chrpos - read_pos};
          const int base{mum.flipped() ? read_length - l_base - 1 : l_base};
          const uint64_t mum_i{mum_ids[base]};
          if (mum_i != mums.size()) {
            const MUM omum{mums[mum_i]};
            const OneBridgeInfo this_bridge(bridge(mum, omum));
            coordinates << " MUM pos " << ref.name(omum.chromosome())
                        << " " << omum.position0()
                        << " offset " << omum.offset()
                        << " length " << omum.length()
                        << " strand " << (omum.flipped() ? '-' : '+');
            if (!(mum == omum)) {
              coordinates << " invariant " << this_bridge.invariant()
                          << " overlap " << this_bridge.offset();
            }
          }
        }
        status = coordinates.str();
      }
    }
    draw_status(status);
    draw_controls();
    last_motion = point;
  }

  void draw_status(const std::string & status) const {
    XFillRectangle(display(), window, fill_gc, bounds[0][0], 0,
                   bounds[0][1] - bounds[0][0], bounds[1][0] - border_width);
    const double avail_height{bounds[1][0] * 0.65};
    static X11Font * status_font{nullptr};
    X11Font * fits{app.fonts.fits(status, bounds[0][2], avail_height)};
    if (fits != status_font) {
      status_font = fits;
      XSetFont(display(), gc, fits->id());
    }
    XDrawString(display(), window, gc, bounds[0][0],
                fits->centered_y((bounds[1][0] - border_width) / 2),
                const_cast<char *>(status.c_str()),
                static_cast<unsigned int>(status.size()));
  }

  virtual void enter(const XCrossingEvent &) {
    inside = true;
  }

  virtual void leave(const XCrossingEvent &) {
    inside = false;
    if (destroyed) return;
    draw_status("");
  }

  bool display_read(const MUMindex mindex) const {
    const MUM mum{mumdex_->mum(mindex)};
    if (mum.length() < min_mum_len) return false;
    const unsigned int abspos{ref.abspos(mum.chromosome(), mum.position0())};
    const unsigned int min_map{mappability.min(abspos, mum.length())};
    const unsigned int excess{mum.length() - min_map};
    if (excess < min_excess) return false;
    return mum.chromosome() == chromosome &&
        read_position(mindex) >= start_position;
  }

  int read_position(const MUMindex & mindex) const {
    const MUM mum{mumdex_->mum(mindex)};
    return mum.read_position0(mumdex_->pair(mindex).length(mum.read_2()));
    // should be equivalent to...
    const Pair pair{mumdex_->pair(mindex)};
    const unsigned int read_length{pair.length(mum.read_2())};
    return mum.read_position0(read_length);
  }

  unsigned int calc_border() const {
    return static_cast<unsigned int>(std::min(width(), height()) / 20);
  }

  virtual void client_message(const XClientMessageEvent & event) {
    X11Win::client_message(event);
    if (static_cast<uint64_t>(event.data.l[0]) == *app.wmDeleteMessage()) {
      return;
    }
    // Note segmentation fault caused by this code somehow when exiting app
    const unsigned int c{static_cast<unsigned int>(event.data.l[1])};
    const unsigned int p{static_cast<unsigned int>(event.data.l[2])};
    std::cout << "Setting view to position "
              << ref.name(c) << " " << p << std::endl;
    set_position(c, p);
    draw();
    XFlush(display());
  }

  virtual void draw() {
    const unsigned int border{calc_border()};
    set_bounds(border, width() - border, border, height() - border);

    // White background
    XFillRectangle(display(), window, fill_gc, 0, 0, width(), height());

    // Optional invariants legend page
    if (invariants_radio) {
      const unsigned int n_x{8};
      const unsigned int n_y{8};
      for (unsigned int i{0}; i != color_names.size(); ++i) {
        const unsigned int swidth{bounds[0][2] / n_x};
        const unsigned int sheight{bounds[1][2] / n_y};
        const unsigned int x{bounds[0][0] + (i % n_x) * swidth};
        const unsigned int y{bounds[1][0] + (i / n_y) * sheight};
        const unsigned int edge{std::min(swidth, sheight) / 3};
        const unsigned int bwidth{swidth - edge};
        const unsigned int bheight{sheight - edge};
        // rectangle(cgc(i), x, y, bwidth, bheight, true);
        const GC color_gc{color_gcs[i]};
        rectangle(color_gc, x, y, bwidth, bheight, true);
        std::ostringstream text_stream;
        for (const bool positive : {false, true}) {
          for (unsigned int j{0}; j != color_names.size(); ++j) {
            const int invariant{(positive ? 1 : -1) *
                  static_cast<int>(j)};
            if (positive && j == 0) continue;
            if (cgc(invariant) == color_gc) {
              if (positive) text_stream << " ";
              text_stream << invariant;
            }
          }
        }
        std::string text{text_stream.str()};
        static X11Font * status_font{nullptr};
        X11Font * fits{app.fonts.fits(text, bwidth, bheight)};
        if (fits != status_font) {
          status_font = fits;
          XSetFont(display(), gc, fits->id());
        }
        XDrawString(display(), window, gc,
                    fits->centered_x(text, x + bwidth / 2),
                    fits->centered_y(y + bheight / 2),
                    const_cast<char *>(text.c_str()),
                    static_cast<unsigned int>(text.size()));
      }
      draw_controls();
      return;
    }

    displayed_mums.clear();

    // Adjust start index to respond to offset_index
    while (offset_index != 0) {
      if ((offset_index < 0 && start_index == 0) ||
          (offset_index > 0 && start_index + 1 ==
           static_cast<int64_t>(mumdex_->n_mums()))) {
        offset_index = 0;
      } else {
        start_index += (offset_index > 0 ? 1 : -1);
        const MUMindex mindex{mumdex_->index()[start_index]};
        const MUM mum{mumdex_->mum(mindex)};
        chromosome = mum.chromosome();
        start_position = mum.position0();
        if (display_read(mindex)) offset_index += (offset_index > 0 ? -1 : 1);
      }
    }

    lowest_pos = start_position;

    const MUMindex smindex{mumdex_->index()[start_index]};
    const Pair spair{mumdex_->pair(smindex)};
    const MUM smum{mumdex_->mum(smindex)};
    const unsigned int sread_length{spair.length(smum.read_2())};
    chromosome = smum.chromosome();
    start_position = smum.read_position0(sread_length);

    // Choose reads to display
    std::set<uint64_t> seen;
    int64_t stop_index{0};
    for (uint64_t index = start_index; index != mumdex_->n_mums(); ++index) {
      stop_index = index;
      const MUMindex mindex{mumdex_->index()[index]};
      const Pair pair{mumdex_->pair(mindex)};
      if (pair.dupe()) continue;
      const MUM mum{mumdex_->mum(mindex)};
      const unsigned int read_length{pair.length(mum.read_2())};
      const int64_t mum_x{chrpos_to_x(mum.read_position0(read_length))};
      if (mum_x >= width()) break;

      if (displayed_mums.size() &&
          mum.chromosome() != mumdex_->mum(displayed_mums.front()).chromosome())
        break;
      if (!display_read(mindex)) continue;

      const uint64_t read_id{mindex.pair_index() +
            mum.read_2() * mumdex_->n_pairs()};
      if (!dupes_radio && !seen.insert(read_id).second) continue;
      displayed_mums.push_back(mindex);
      lowest_pos = std::min(lowest_pos, mum.read_position0(read_length));
    }

    auto index_less = [this](const MUMindex & lhs, const MUMindex & rhs) {
      return read_position(lhs) < read_position(rhs);
    };
    stable_sort(displayed_mums.begin(), displayed_mums.end(), index_less);

    const int used_read_spacing{read_spacing >= pixels_per_read ?
          pixels_per_read - 1 : read_spacing};
    const int height__{pixels_per_read - used_read_spacing};

#if 0
    std::set<int> page_invariants;
    std::vector<unsigned int> last_bases(displayed_reads.size());
    const double avail_height{bounds[1][0] * 0.5};
    static X11Font * status_font{nullptr};
    static GC inv_gc{create_gc(app.black, app.white)};
    X11Font * fits{app.fonts.fits(status, bounds[0][2], avail_height)};
    if (fits != status_font) {
      status_font = fits;
      XSetFont(display(), inv_gc, fits->id());
    }
#endif

    // if (displayed_mums.empty()) throw Error("No reads to show");

    // Mark selected position
    const int selected_x{chrpos_to_x(selected_position) + pixels_per_base / 2};
    XDrawLine(display(), window, gc,
              selected_x, bounds[1][0], selected_x, bounds[1][1]);


    for (uint64_t i{0}; i != displayed_mums.size(); ++i) {
      const int y{map_index_to_y(i)};
      // need to view later reads too!
      if (y + pixels_per_read + bounds[1][0] >= height()) break;
      const MUMindex mindex{displayed_mums[i]};
      const Pair pair{mumdex_->pair(mindex)};
      const MUM mum{mumdex_->mum(mindex)};
      const unsigned int read_length{pair.length(mum.read_2())};
      // std::cout << "Read " << i << endl;

      const int read_pos{mum.read_position0(read_length)};
      const int read_x{chrpos_to_x(read_pos)};
      if (read_x >= width()) break;
#if 0
      if (legend_radio) {
        first_bases[i] = read_x;
        last_bases[i] = read_x + pixels_per_base * read_length;
      }
#endif
      // Other MUMs in read
      const std::vector<MUM> other_mums{read_mums(mindex)};
      std::vector<unsigned int> base_seen(read_length);
      unsigned int total_n_bases{0};
      for (const MUM omum : other_mums) {
        unsigned int first_hit{0};
        unsigned int last_hit{0};
        unsigned int n_bases{0};
        for (unsigned int b{omum.offset()}; b != omum.offset() + omum.length();
             ++b) {
          if (!base_seen[b]) {
            base_seen[b] = 1;
            if (!n_bases) first_hit = b;
            last_hit = b;
            ++n_bases;
            ++total_n_bases;
          }
        }
        if (n_bases == 0) continue;
        if (0) std::cout << " (" << omum.offset() << ", "
                         << omum.length() << ", "
                         << n_bases << ")";

        const unsigned int first_base{mum.flipped() ?
              read_length - last_hit - 1 : first_hit};
        const int first_pos{mum.read_position0(read_length) +
              static_cast<int>(first_base)};
        const int omum_x{chrpos_to_x(first_pos)};
        // if (bridge.invariant() != 0) continue;
        const int invariant{static_cast<int>(bridge(mum, omum).invariant())};
#if 0
        auto found = page_invariants.insert(invariant);
        if (found.second) {
          // Draw legend nearby
          const bool left_;
          rectangle(cgc(invariant), x, fits->below(),
                    avail_height, avail_height, true);
        }
#endif
        rectangle(cgc(invariant), omum_x, y,
                  pixels_per_base * n_bases,
                  //  std::max(3U, pixels_per_read) - used_read_spacing, true);
                  height__, true);
      }

      // Draw unseen bases
      if (!bases_radio) {
        if (total_n_bases != read_length) {
          const std::array<std::string, 2> sequences(
              mumdex_->sequences(mindex.pair_index()));
          const std::string & sequence{sequences[mum.read_2()]};
          for (unsigned int b{0}; b != base_seen.size(); ++b) {
            const bool empty{!base_seen[b]};
            if (empty) {
              const char base{mum.flipped() ? complement(sequence[b]) :
                    sequence[b]};
              const unsigned int base_offset{mum.flipped() ?
                    read_length - b - 1 : b};
              const int64_t pos{read_pos + static_cast<int>(base_offset)};
              const GC base_gc{base == ref[mum.chromosome()][pos] ?
                    color_gcs[0] : char_gcs[base]};
              rectangle(base_gc, read_x + base_offset * pixels_per_base, y,
                        pixels_per_base,
                        height__, true);
            }
          }
        }
      } else {
        unsigned int low_(static_cast<unsigned int>(base_seen.size()));
        for (unsigned int b{0}; b != base_seen.size(); ++b) {
          const bool empty{!base_seen[b]};
          if (empty && low_ == base_seen.size()) low_ = b;
          if ((!empty && low_ != base_seen.size()) ||
              (empty && b + 1 == base_seen.size())) {
            const unsigned int high_{b + (empty ? 1 : 0)};
            const unsigned int wid{pixels_per_base * (high_ - low_)};
            const unsigned int off{mum.flipped() ?
                  read_length - high_ : low_};
            rectangle(gc, read_x + off * pixels_per_base, y, wid,
                      height__, true);
            low_ = static_cast<unsigned int>(base_seen.size());
          }
        }
      }
    }
    draw_controls();

    // Read ahead if not busy
    if (0) {
      const unsigned int read_ahead_n{10000};
      // const uint64_t read_ahead_block{100};
      const int64_t new_smallest_index{std::min(
          smallest_index,
          start_index > read_ahead_n ?
          start_index - read_ahead_n : 0)};
      const int64_t new_biggest_index{std::min(
          static_cast<int64_t>(mumdex_->n_mums()),
          std::max(stop_index + read_ahead_n, biggest_index))};
      // threads.run()
      smallest_index = new_smallest_index;
      biggest_index = new_biggest_index;
    }
    XFlush(display());
  }

  OneBridgeInfo bridge(const MUM mumA, const MUM mumB) const {
    const MUM mum1{mumA.offset() < mumB.offset() ? mumA : mumB};
    const MUM mum2{mumA.offset() < mumB.offset() ? mumB : mumA};
    const bool mum1_is_high{!mum1.flipped()};
    const bool mum2_is_high{mum2.flipped()};
    return OneBridgeInfo{mum1, mum1_is_high, mum2, mum2_is_high};
  }

  GC cgc(const int invariant) const {
    if (invariant == 0) return color_gcs[0];
    return color_gcs[(((abs_invariant_radio ? abs(invariant) : invariant) - 1) %
                      (color_gcs.size() - 1)) + 1];
  }

  void draw_controls() {
    for (const Radio * radio : radios) radio->draw();
  }

  void rectangle(GC gc_, const int x, const int y,
                 const unsigned int w, const unsigned int h,
                 bool fill = false) {
    if (fill) {
      XFillRectangle(display(), window, gc_, x, y, w, std::max(2U, h));
      if (h > 3 && border_radio)
        XDrawRectangle(display(), window, gc, x, y, w, h);
    } else {
      XDrawRectangle(display(), window, gc_, x, y, w, h);
    }
  }

  int map_index_to_y(const int64_t index) const {
    return static_cast<int>(index * pixels_per_read + bounds[1][0]);
  }
  int64_t y_to_map_index(int y) const {
    return (static_cast<int64_t>(y) - bounds[1][0]) / pixels_per_read;
  }
  int chrpos_to_x(const int pos) const {
    return (pos - lowest_pos) * pixels_per_base +
        bounds[0][0];
  }
  int x_to_chrpos(const int x) const {
    return (x - bounds[0][0]) / pixels_per_base + lowest_pos;
  }

  bool inside{true};

  void set_mumdex(const unsigned int m) {
    mumdex_ = &*mumdexes_.begin() + m;
    XStoreName(display(), window, mumdex_names_[m].c_str());
  }

  void set_position(const std::string & chr, const unsigned int position,
                    const bool center = true) {
    chromosome = chr_lookup[chr];
    set_position(chromosome, position, center);
  }

  void set_position(const unsigned int chr, const unsigned int position,
                    const bool center = true) {
    chromosome = chr;
    if (center) selected_position = position;
    start_position = position;
    const unsigned int border{calc_border()};
    const int n_bases((width() - 2 * border) / pixels_per_base / 2);
    if (center && start_position > n_bases) start_position -= n_bases;
    start_index = mumdex_->lower_bound(chromosome, start_position) -
        mumdex_->index().begin();
    if (start_index == static_cast<int64_t>(mumdex_->index().size()))
      start_index = 0;
    while (read_position(mumdex_->index()[start_index]) < start_position &&
           start_index + 1 != static_cast<int64_t>(mumdex_->index().size()) &&
           mumdex_->mum(mumdex_->index()[start_index]).chromosome() ==
           chromosome) {
      ++start_index;
    }
    start_position = read_position(mumdex_->index()[start_index]);
    offset_index = 0;
    smallest_index = start_index;
    biggest_index = start_index;
  }

  std::vector<MUM> read_mums(const MUMindex mindex) const {
    std::vector<MUM> other_mums;
    const uint64_t pair_index{mindex.pair_index()};
    const Pair pair{mumdex_->pair(mindex)};
    const MUM mum{mumdex_->mum(mindex)};
    other_mums.push_back(mum);
    for (uint64_t mi{mumdex_->mums_start(pair_index)};
         mi != mumdex_->mums_stop(pair_index); ++mi) {
      if (mi == pair.mums_start() + mindex.mum_in_pair_index()) continue;
      const MUM omum{mumdex_->mum(mi)};
      if (omum.read_2() == mum.read_2()) other_mums.push_back(omum);
    }
    if (0) {
      stable_sort(other_mums.begin() + 1, other_mums.end(),
                  [](const MUM lhs, const MUM rhs) {
                    if (lhs.length() == rhs.length()) {
                      return lhs.offset() < rhs.offset();
                    } else {
                      return lhs.length() > rhs.length();
                    }
                  });
    } else {
      stable_sort(other_mums.begin(), other_mums.end(),
                  [mum, this](const MUM lhs, const MUM rhs) {
                    const int linv{abs(mum_invariant(mum, lhs))};
                    const int rinv{abs(mum_invariant(mum, rhs))};
                    if (linv == rinv) {
                      return lhs.length() > rhs.length();
                    } else {
                      return linv < rinv;
                    }
                  });
    }

    return other_mums;
  }

  int mum_invariant(const MUM mum1, const MUM mum2) const {
    return static_cast<int>(bridge(mum1, mum2).invariant());
  }

  void report_mum_len() const {
    std::ostringstream out;
    out << "Minimum main MUM length is now " << min_mum_len <<
        " and minimum excess mappability is " << min_excess;
    draw_status(out.str());
  }

 private:
  const std::vector<std::string> & mumdex_names_;
  const std::vector<MUMDEX> & mumdexes_;
  const MUMDEX * mumdex_;
  const Reference & ref;
  const ChromosomeIndexLookup chr_lookup;
  const Mappability mappability;

  std::vector<std::string> color_names{};
  std::vector<XColor> X_colors{};
  std::vector<GC> color_gcs{};
  std::vector<GC> char_gcs{};

  std::vector<std::mutex> read_mutexes{};
  unsigned int chromosome{0};
  int selected_position{0};
  int start_position{0};

  bool exited{false};
  std::mutex exited_mutex{};
  ThreadPool pool;
  std::vector<std::future<void>> futures{};

  std::vector<MUMindex> displayed_mums{};
  int64_t start_index{0};
  int64_t smallest_index{0};
  int64_t biggest_index{0};
  int64_t offset_index{0};
  int lowest_pos{0};

  int read_spacing{0};
  int pixels_per_read{5};  // height
  int pixels_per_base{5};  // width

  unsigned int min_mum_len{20};
  unsigned int min_excess{10};

  // Graphics contexts
  GC border_gc{}, border_fill_gc{}, minor_gc{}, major_gc{}, mum_gc{};

  // Radios
  Radio mums_radio{"Toggle mum display", this, {-1, 1},
    {[this]() { draw(); }}, true, false};
  Radio dupes_radio{"Toggle pair de-duping", this, {-1, 2},
    {[this]() { draw(); }}, true, false};
  Radio border_radio{"Toggle borders around reads", this, {-1, 3},
    {[this]() { draw(); }}, true, false};
  Radio bases_radio{"Toggle bases display", this, {-1, 4},
    {[this]() { draw(); }}, true, false};
  Radio invariants_radio{"Toggle invariants legend display", this, {-1, 6},
    {[this]() { draw(); }}, true, false};
  Radio abs_invariant_radio{"Toggle absolute value of invariant for colors",
        this, {-1, 7}, {[this]() { draw(); }}, true, false};
  Radio rand_invariant_radio{"Randomize invariant colors", this, {-1, 8},
    {[this]() {
        static std::random_device rd;
        static auto mersenne = std::mt19937_64(rd());
        shuffle(color_gcs.begin() + 1, color_gcs.end(), mersenne);
        draw();
      }}};
  Radio legend_radio{"Toggle in-screen legend display", this, {-1, 9},
    {[this]() { draw(); }}, true, false};

  Radio sample_radio{"Switch to next sample", this, {1, -1},
    {[this]() {
        ++mumdex_;
        if (mumdex_ == &*mumdexes_.end()) mumdex_ = &*mumdexes_.begin();
        const unsigned int m{static_cast<unsigned int>(
            mumdex_ - &*mumdexes_.begin())};
        XStoreName(display(), window, mumdex_names_[m].c_str());
        set_position(ref.name(chromosome), start_position, false);
        draw();
        draw_status(mumdex_names_[mumdex_ - &*mumdexes_.begin()]);
      }, [this]() { return mumdex_names_.size() > 1; }}};

  std::vector<Radio *> radios{&dupes_radio,
        &border_radio, &bases_radio, &invariants_radio,
        &abs_invariant_radio, &rand_invariant_radio, &sample_radio};
  std::vector<Radio> create_unnamed_radios() {
  return std::vector<Radio>{
    {"Go to next read", this, {1, 2}, {[this]() {
          offset_index += 1; draw(); }}},
    {"Go to previous read", this, {1, 3}, {[this]() {
          offset_index -= 1; draw(); }}},

    {"Jump half a page forward", this, {1, 4.5}, {[this]() {
          offset_index += std::max(1, height() / pixels_per_read / 2);
          draw(); }}},
    {"Jump half a page backwards", this, {1, 5.5}, {[this]() {
          offset_index -= std::max(1, height() / pixels_per_read / 2);
          draw(); }}},

    {"Increase read spacing", this, {-1, -4.5},
      {[this]() { read_spacing += 1; draw(); },
            [this]() {return read_spacing + 1 < pixels_per_read; }}},
    {"Decrease read spacing", this, {-1, -5.5},
      {[this]() { read_spacing -= 1; draw(); },
            [this]() {return read_spacing; }}},

    {"Increase minimum main MUM excess mappability", this, {-7, -1},
      {[this]() { min_excess += 1; draw(); report_mum_len(); }}},
    {"Decrease minimum main MUM excess mappability", this, {-8, -1},
      {[this]() { min_excess -= 1; draw(); report_mum_len(); },
            [this]() {return min_excess; }}},

    {"Increase minimum main MUM length", this, {-4.5, -1},
      {[this]() { min_mum_len += 1; draw(); report_mum_len(); }}},
    {"Decrease minimum main MUM length", this, {-5.5, -1},
      {[this]() { min_mum_len -= 1; draw(); report_mum_len(); },
            [this]() {return min_mum_len; }}},

    {"Make reads wider", this, {-2, -1},
      {[this]() { pixels_per_base += 1; draw(); }}},
    {"Make reads narrower", this, {-3, -1},
      {[this]() { pixels_per_base -= 1; draw(); },
            [this]() {return pixels_per_base > 1; }}},

    {"Make reads taller", this, {-1, -2},
      {[this]() { pixels_per_read += 1; draw(); }}},
    {"Make reads shorter", this, {-1, -3},
      {[this]() { pixels_per_read -= 1; draw(); },
            [this]() { return pixels_per_read > 1; }}},

    {"Make reads taller and wider", this, {-1, -1},
      {[this]() {
          (pixels_per_read *= 1.15) += 1;
          (pixels_per_base *= 1.1) += 1;
          draw(); }}},
    {"Make reads shorter and narrower", this, {-2, -2},
      {[this]() {
          if (pixels_per_read > 1) pixels_per_read /= 1.1;
          if (pixels_per_base > 1) pixels_per_base /= 1.1;
          draw();
        }, [this]() { return pixels_per_read > 1 || pixels_per_base > 1; }}},
        };
  }

  std::vector<Radio> unnamed_radios{create_unnamed_radios()};

  Point last_press{}, last_motion{};
  bool moved{false};
};

class X11CandidateViewer : public Geometry {
 public:
  X11CandidateViewer(const std::string & cand_file_name,
                     const std::string & samples_dir,
                     const std::string & pop_file_name,
                     const Geometry & geometry_ = {{1200, 1000}, {0, 0}}) :
      Geometry{geometry_}, pop{pop_file_name} {
    std::ifstream cand_list_file{cand_file_name.c_str()};
    if (!cand_list_file) throw Error("Could not open candidates file")
                             << cand_file_name;

    std::string line;
    std::vector<std::string> the_rest;
    while (getline(cand_list_file, line)) {
      std::string name;
      std::string chr;
      unsigned int p;
      std::string rest;
      std::istringstream line_stream{line.c_str()};
      line_stream >> name >> chr >> p;
      if (!line_stream) throw Error("Problem parsing candidate list line")
                            << line;
      getline(line_stream, rest);
      names.push_back(name);
      chrs.push_back(chr);
      pos.push_back(p);
      the_rest.push_back(rest);
      if (!mumdex_lookup.count(name)) {
        const Family family{pop.family(pop.sample(name))};
        const std::vector<Sample> family_samples{pop.samples(family)};
        for (unsigned int s{0}; s != family_samples.size(); ++s) {
          const Sample fsample{family_samples[s]};
          const std::string sample{pop.sample(fsample)};
          mumdex_lookup.emplace(sample, mumdexes.size());
          mumdexes.emplace_back(samples_dir + "/" + sample + "/mumdex");
          mumdex_names.push_back(sample + " " + pop.member(fsample));
        }
      }
    }

    bool rest_used{false};
    for (const std::string & r : the_rest) {
      if (r.size()) {
        rest_used = true;
        break;
      }
    }

    std::vector<std::vector<std::string>> text{
      {"Sample"}, {"Type"}, {"Family"}, {"Location"}};
    if (rest_used) {
      text.push_back({"Extra"});
    }
    std::vector<unsigned int> exclusive_rows{0};
    for (unsigned int n{0}; n != names.size(); ++n) {
      const Sample sample{pop.sample(names[n])};
      const Family family{pop.family(sample)};
      text[0].push_back(names[n]);
      text[1].push_back(pop.member(sample));
      text[2].push_back(pop.family(family));
      text[3].push_back(chrs[n] + " " + std::to_string(pos[n]));
      if (rest_used) text[4].push_back(the_rest[n]);
      exclusive_rows.push_back(n + 1);
    }

    X11TextGrid::CallBack call_back{
      std::bind(&X11CandidateViewer::launch_graph, this,
                std::placeholders::_1)};
    X11TextGrid::CallBack cell_test{
      std::bind(&X11CandidateViewer::launch_ready, this,
                std::placeholders::_1)};
    using Vec = std::vector<unsigned int>;
    using Cells = std::vector<std::pair<unsigned int, unsigned int>>;

    app.create<X11TextGrid>(text, Cells{}, Cells{},
                            Vec{0, 1, 3}, Vec{0}, Vec{0, 2},
                            exclusive_rows, call_back, nullptr, cell_test,
                            *this);
    launch_graph2(0);
    app.run();
  }
#if 0
  bool close_views() {
    for (auto i = next(app.windows.begin()); i != app.windows.end(); ++i) {
      app.close_window((*i)->window);
    }
    return true;
  }
#endif
  bool launch_graph(const X11TextGrid::CellStatus & status) {
    const unsigned int selected_cand{static_cast<unsigned int>(
        std::find(status[2].begin(), status[2].end(), 1) -
        status[2].begin() - 1)};
    // last_selected = selected_cand;
    return launch_graph2(selected_cand);
  }

  bool launch_graph2(const unsigned int selected_cand) {
    const std::string sample_name{names[selected_cand]};
    const Sample sample{pop.sample(sample_name)};
    const Family family{pop.family(sample)};
    const std::vector<Sample> family_samples{pop.samples(family)};
    const int n_samples{static_cast<int>(family_samples.size())};
    const int n_x{n_samples < 4 ? 1 : (n_samples < 9 ? 2 : 3)};
    const int n_y{(n_samples + (n_x - 1)) / n_x};
    const int sx{width() / n_x};
    const int sy{height() / n_y};
    for (unsigned int s{0}; s != family_samples.size(); ++s) {
      const unsigned int xi{n_x > 1 ? (s % n_x) : 0};
      const unsigned int yi{s / n_x};
      const Sample fsample{family_samples[s]};
      const unsigned int mumdex_id{mumdex_lookup[pop.sample(fsample)]};
      std::cerr << "launched mumdex " << mumdex_id
                << " of " << mumdexes.size()
                << " and cand " << selected_cand
                << " of " << names.size()
                << " " << xi << " " << yi << " " << n_x << " " << n_y
                << std::endl;
      X11MUMdexViewer & viewer{app.create<X11MUMdexViewer>(
          mumdex_names, mumdexes,
          Geometry{{sx - x_offset(), sy - y_offset()},
            {static_cast<int>(xi * sx), static_cast<int>(yi * sy)}})};
        viewer.set_mumdex(mumdex_id);
        viewer.set_position(chrs[selected_cand], pos[selected_cand]);
      }
      return true;
  }
  bool launch_ready(const X11TextGrid::CellStatus & status) const {
    return std::find(status[2].begin(), status[2].end(), 1) != status[2].end();
  }

  X11App app{};
  std::vector<std::string> names{};
  std::vector<std::string> chrs{};
  std::vector<unsigned int> pos{};
  std::vector<MUMDEX> mumdexes{};
  std::vector<std::string> mumdex_names{};
  std::map<std::string, unsigned int> mumdex_lookup{};
  Population pop;
};

}  // namespace paa

#endif  // PAA_MVIEW
```
