## Supplementary material for "G-Graph: An interactive genomic graph viewer": MUMdex Source Code for G-Graph: population.h

```
//
// population.h
//
// define a population structure
//
// Copyright 2016 Peter Andrews @ CSHL
//

#ifndef PAA_POPULATION_H
#define PAA_POPULATION_H

#include <algorithm>
#include <fstream>
#include <map>
#include <sstream>
#include <utility>
#include <vector>
#include <string>

#include "error.h"

namespace paa {

class Population {
 public:
  enum class Relation { none, parent, child, before, after };
  enum class Member { normal,
      mother, father, proband, sibling,
      matched, cancer, source, edited };

  class Tags {
   public:
    explicit Tags(const std::string member_arg) { *this = tags(member_arg); }
    Tags(const Member member_arg, const Relation relation_arg) :
        member_{member_arg}, relation_{relation_arg} {}
    Member member() const { return member_; }
    Relation relation() const { return relation_; }

    // members
    Tags tags(const std::string & member_arg) const {
      if (member_arg == "mother")  return { Member::mother,  Relation::parent };
      if (member_arg == "father")  return { Member::father,  Relation::parent };
      if (member_arg == "proband") return { Member::proband, Relation::child  };
      if (member_arg == "sibling") return { Member::sibling, Relation::child  };
      if (member_arg == "matched") return { Member::matched, Relation::before };
      if (member_arg == "cancer")  return { Member::cancer,  Relation::after  };
      if (member_arg == "source")  return { Member::source,  Relation::before };
      if (member_arg == "edited")  return { Member::edited,  Relation::after  };
      if (member_arg == "normal")  return { Member::normal,  Relation::none   };
      throw Error("Unknown member") << member_arg;
    }

   private:
    // Ignore spurious warning here - not sure why triggered
#pragma GCC diagnostic push
#pragma GCC diagnostic ignored "-Wuseless-cast"
    Member member_{Member::normal};
    Relation relation_{Relation::none};
#pragma GCC diagnostic pop
  };

  class Family {
   public:
    Family() { }
    explicit Family(const uint64_t id_arg) :
        id{static_cast<unsigned int>(id_arg)} { }
    operator unsigned int() const { return id; }
    Family operator*() const {
      return *this;
    }
    Family & operator++() {
      ++id;
      return *this;
    }

   private:
    unsigned int id{0};
  };

  class Sample {
   public:
    Sample() { }
    explicit Sample(const uint64_t id_arg) :
        id{static_cast<unsigned int>(id_arg)} { }
    operator unsigned int() const { return id; }
    Sample operator*() const {
      return *this;
    }
    Sample & operator++() {
      ++id;
      return *this;
    }

   private:
    unsigned int id{0};
  };

  class FamilyRange {
   public:
    explicit FamilyRange(const uint64_t n_samples) :
        stop{static_cast<unsigned int>(n_samples)} { }
    FamilyRange(const Family start_arg, const Family stop_arg) :
        start{start_arg}, stop{stop_arg} { }
    Family begin() const { return start; }
    Family end() const { return stop; }
    unsigned int size() const { return stop - start; }

   private:
    Family start{0};
    Family stop{0};
  };

  class SampleRange {
   public:
    explicit SampleRange(const uint64_t n_samples) :
        stop{static_cast<unsigned int>(n_samples)} { }
    SampleRange(const Sample start_arg, const Sample stop_arg) :
        start{start_arg}, stop{stop_arg} { }
    Sample begin() const { return start; }
    Sample end() const { return stop; }

   private:
    Sample start{0};
    Sample stop{0};
  };

  explicit Population(const std::string & family_info) {
    std::ifstream family_file{family_info.c_str()};
    if (!family_file)
      throw Error("Could not open family file") << family_info;
    std::string line;
    std::string family_string;
    std::string member_string;
    std::string sample_string;
    std::string sex_string;
    while (family_file) {
      getline(family_file, line);
      if (line.empty()) break;
      if (line[0] == '#') continue;
      std::istringstream line_stream{line};
      if (line_stream) {
        line_stream >> family_string;
        if (!line_stream) throw Error("Problem reading family") << n_families();

        const Family f{n_families()};
        families_[family_string] = f;
        family_string_.push_back(family_string);
        family_sample_.push_back(std::vector<Sample>());
        family_normal_.push_back(std::vector<Sample>());
        family_mother_.push_back(std::vector<Sample>());
        family_father_.push_back(std::vector<Sample>());
        family_proband_.push_back(std::vector<Sample>());
        family_sibling_.push_back(std::vector<Sample>());
        family_matched_.push_back(std::vector<Sample>());
        family_cancer_.push_back(std::vector<Sample>());
        family_source_.push_back(std::vector<Sample>());
        family_edited_.push_back(std::vector<Sample>());
        family_unrelated_.push_back(std::vector<Sample>());
        family_parent_.push_back(std::vector<Sample>());
        family_child_.push_back(std::vector<Sample>());
        family_before_.push_back(std::vector<Sample>());
        family_after_.push_back(std::vector<Sample>());
        family_unaffected_.push_back(std::vector<Sample>());
        family_affected_.push_back(std::vector<Sample>());
        while (line_stream) {
          line_stream >> member_string >> sample_string >> sex_string;
          if (line_stream) {
            const Sample s{n_samples()};
            samples_[family_string].push_back(s);
            samples_[sample_string].push_back(s);
            family_sample_.back().push_back(s);
            sample_family_.push_back(f);
            member_string_.push_back(member_string);
            sample_string_.push_back(sample_string);
            sex_string_.push_back(sex_string);
            nX_.push_back(static_cast<unsigned int>(
                count(sex_string.begin(), sex_string.end(), 'X')));
            nY_.push_back(static_cast<unsigned int>(
                count(sex_string.begin(), sex_string.end(), 'Y')));

            const Tags tags{member_string};
            member_.push_back(tags.member());
            relation_.push_back(tags.relation());
            affected_.push_back(is_proband(s) || is_cancer(s) || is_edited(s));

            if (is_normal(s)) family_normal_.back().push_back(s);
            if (is_mother(s)) family_mother_.back().push_back(s);
            if (is_father(s)) family_father_.back().push_back(s);
            if (is_proband(s)) family_proband_.back().push_back(s);
            if (is_sibling(s)) family_sibling_.back().push_back(s);
            if (is_matched(s)) family_matched_.back().push_back(s);
            if (is_cancer(s)) family_cancer_.back().push_back(s);
            if (is_source(s)) family_source_.back().push_back(s);
            if (is_edited(s)) family_edited_.back().push_back(s);
            if (is_unrelated(s)) family_unrelated_.back().push_back(s);
            if (is_parent(s)) family_parent_.back().push_back(s);
            if (is_child(s)) family_child_.back().push_back(s);
            if (is_before(s)) family_before_.back().push_back(s);
            if (is_after(s)) family_after_.back().push_back(s);
            if (is_unaffected(s)) family_unaffected_.back().push_back(s);
            if (is_affected(s)) family_affected_.back().push_back(s);
          }
        }
        if (family_sample_.back().size() == 0)
          throw Error("Empty family") << family_string;
      }
    }
    if (n_families() == 0) throw Error("No families loaded");
    if (n_samples() == 0) throw Error("No samples loaded");
  }

  // returning a number
  unsigned int n_families() const {
    return static_cast<unsigned int>(family_string_.size());
  }
  unsigned int n_samples() const {
    return static_cast<unsigned int>(sample_string_.size());
  }
  unsigned int n_members(const Family f) const {
    return static_cast<unsigned int>(family_sample_[f].size());
  }
  unsigned int nX(const Sample s) const { return nX_[s]; }
  unsigned int nY(const Sample s) const { return nY_[s]; }

  // returning a string
  const std::string & family(const Family f) const { return family_string_[f]; }
  const std::string & sample(const Sample s) const { return sample_string_[s]; }
  const std::string & member(const Sample s) const { return member_string_[s]; }
  const std::string & sex(const Sample s) const { return sex_string_[s]; }

  // returning a range of families or samples
  FamilyRange families() const { return FamilyRange{n_families()}; }
  SampleRange samples() const { return SampleRange{n_samples()}; }

  // returning family or sample ids
  Family family(const Sample s) const { return sample_family_[s]; }
  Family family(const std::string & name) const {
    return families_.at(name);
  }
  Sample sample(const std::string & name) const {
    const auto & sample_returned = samples(name);
    if (sample_returned.size() != 1)
      throw Error("One sample was not returned for") << name;
    return sample_returned.front();
  }
  const std::vector<Sample> & samples(const std::string & name) const {
    return samples_.at(name);
  }
  const std::vector<Sample> & samples(const Family f) const {
    return family_sample_[f];
  }
  const std::vector<Sample> & normal(const Family f) const {
    return family_normal_[f];
  }
  const std::vector<Sample> & mother(const Family f) const {
    return family_mother_[f];
  }
  const std::vector<Sample> & father(const Family f) const {
    return family_father_[f];
  }
  const std::vector<Sample> & proband(const Family f) const {
    return family_proband_[f];
  }
  const std::vector<Sample> & sibling(const Family f) const {
    return family_sibling_[f];
  }
  const std::vector<Sample> & matched(const Family f) const {
    return family_matched_[f];
  }
  const std::vector<Sample> & cancer(const Family f) const {
    return family_cancer_[f];
  }
  const std::vector<Sample> & source(const Family f) const {
    return family_source_[f];
  }
  const std::vector<Sample> & edited(const Family f) const {
    return family_edited_[f];
  }
  const std::vector<Sample> & unrelated(const Family f) const {
    return family_unrelated_[f];
  }
  const std::vector<Sample> & parent(const Family f) const {
    return family_parent_[f];
  }
  const std::vector<Sample> & child(const Family f) const {
    return family_child_[f];
  }
  const std::vector<Sample> & before(const Family f) const {
    return family_before_[f];
  }
  const std::vector<Sample> & after(const Family f) const {
    return family_after_[f];
  }
  const std::vector<Sample> & unaffected(const Family f) const {
    return family_unaffected_[f];
  }
  const std::vector<Sample> & affected(const Family f) const {
    return family_affected_[f];
  }

  // returning a bool

  // member
  bool is_normal(const Sample s) const {
    return member_[s] == Member::normal;
  }
  bool is_mother(const Sample s) const {
    return member_[s] == Member::mother;
  }
  bool is_father(const Sample s) const {
    return member_[s] == Member::father;
  }
  bool is_proband(const Sample s) const {
    return member_[s] == Member::proband;
  }
  bool is_sibling(const Sample s) const {
    return member_[s] == Member::sibling;
  }
  bool is_matched(const Sample s) const {
    return member_[s] == Member::matched;
  }
  bool is_cancer(const Sample s) const {
    return member_[s] == Member::cancer;
  }
  bool is_source(const Sample s) const {
    return member_[s] == Member::source;
  }
  bool is_edited(const Sample s) const {
    return member_[s] == Member::edited;
  }
  // relation
  bool is_unrelated(const Sample s) const {
    return relation_[s] == Relation::none;
  }
  bool is_parent(const Sample s) const {
    return relation_[s] == Relation::parent;
  }
  bool is_child(const Sample s) const {
    return relation_[s] == Relation::child;
  }
  bool is_before(const Sample s) const {
    return relation_[s] == Relation::before;
  }
  bool is_after(const Sample s) const {
    return relation_[s] == Relation::after;
  }
  // affected
  bool is_affected(const Sample s) const {
    return affected_[s];
  }
  bool is_unaffected(const Sample s) const {
    return !affected_[s];
  }

  // other
  void output_sample_meta() const {
    std::ofstream sample_meta("samples.txt");
    if (!sample_meta) throw Error("Problem opening samples.txt");
    for (const Family f : families()) {
      for (const Sample s : samples(f)) {
        sample_meta << family(f) << '\t' << n_members(f) << '\t'
                    << sample(s) << '\t' << member(s) << '\t'
                    << sex(s) << std::endl;
      }
    }
  }
  const std::string mumdex_name(const std::string & samples_dir,
                                const Sample s) const {
    return samples_dir + "/" + sample(s) + "/mumdex";
  }

 private:
  // families
  std::vector<std::string> family_string_{};
  std::vector<std::vector<Sample>> family_sample_{};
  std::vector<std::vector<Sample>> family_normal_{};
  std::vector<std::vector<Sample>> family_mother_{};
  std::vector<std::vector<Sample>> family_father_{};
  std::vector<std::vector<Sample>> family_proband_{};
  std::vector<std::vector<Sample>> family_sibling_{};
  std::vector<std::vector<Sample>> family_matched_{};
  std::vector<std::vector<Sample>> family_cancer_{};
  std::vector<std::vector<Sample>> family_source_{};
  std::vector<std::vector<Sample>> family_edited_{};
  std::vector<std::vector<Sample>> family_unrelated_{};
  std::vector<std::vector<Sample>> family_parent_{};
  std::vector<std::vector<Sample>> family_child_{};
  std::vector<std::vector<Sample>> family_before_{};
  std::vector<std::vector<Sample>> family_after_{};
  std::vector<std::vector<Sample>> family_unaffected_{};
  std::vector<std::vector<Sample>> family_affected_{};

  // samples
  std::vector<Family> sample_family_{};
  std::vector<std::string> member_string_{};
  std::vector<std::string> sample_string_{};
  std::vector<std::string> sex_string_{};
  std::vector<unsigned int> nX_{};
  std::vector<unsigned int> nY_{};
  std::vector<Member> member_{};
  std::vector<Relation> relation_{};
  std::vector<bool> affected_{};

  // lookup
  std::map<std::string, std::vector<Sample>> samples_{};
  std::map<std::string, Family> families_{};
};

using Sample = Population::Sample;
using Family = Population::Family;

}  // namespace paa

#endif  // PAA_POPULATION_H
```
