## Supplementary material for "G-Graph: An interactive genomic graph viewer": MUMdex Source Code for G-Graph: population_bridges.cpp

```
//
// population_bridges
//
// bridge information over a population to find candidates
//
// Copyright 2016 Peter Andrews @ CSHL
//

// TO DO
//
// use simpler ThreadPool for threads!
// consider opportunity for mate support cut!
// loosen cuts, except excess mappability
//
//
// N anchors
//
// nearby bridges TODO
// reduce reference requirement for chrY, chrX bridges in Male MAYBE
// overlap cut PROB NOT
// consolidation of bridges - spurious removal NO
// reduction of reference in sample with candidate? NO
// mate support cut OK
// reference count in parents OK
// mappability excess OK
// bridge in parents with no support cuts or mum length cuts OK
// similar anchors in parents with no support cuts or mum length cuts OK
// do not check mother for chrY bridges OK

// cuts:
//
// need to handle family structure, additional normals,
// cancer and matched normal, twins, technical replicates
// edited variants (crispr)
//
// eliminating cuts
//   bridge counts
//   anchor support
//   mate support
//   seen in matched normal or replicate
//   excess mappability - spurious likely
//   parent reference count
//   N-base anchor extension (similar sequence to bridge in parents)
//
// discriminating cuts: classes of candidates
//   seen in population - still possible denovo, rare vs ultra rare
//   seen in parent - denovo, inherited, mendel testing
//     parent bridge check
//     parent anchor check
//

// internal tests
//
// both percentage
// transmission mendel agreement
// percent seen in population
// microsatellite identity
//
// external tests
//
// compare with Ivan
// compare with Yoon-Ha, Chris
// validation
//

#include <algorithm>
#include <array>
#include <chrono>
#include <deque>
#include <exception>
#include <iostream>
#include <list>
#include <future>
#include <mutex>
#include <queue>
#include <sstream>
#include <string>
#include <utility>
#include <vector>

#include "bridges.h"
#include "error.h"
#include "files.h"
#include "mumdex.h"
#include "population.h"
#include "sequence.h"
#include "utility.h"

using std::array;
using std::async;
using std::cerr;
using std::condition_variable;
using std::cout;
using std::deque;
using std::endl;
using std::exception;
using std::future;
using std::list;
using std::lock_guard;
using std::lower_bound;
using std::move;
using std::mutex;
using std::ostringstream;
using std::priority_queue;
using std::string;
using std::unique_lock;
using std::vector;

using paa::serr;
using paa::sout;
using paa::BridgeInfo;
using paa::ChromosomeIndexLookup;
using paa::ConsensusSequence;
using paa::Error;
using paa::Family;
using paa::FileVector;
using paa::Mappability;
using paa::MergeHelper;
using paa::MergeHelperCompare;
using paa::MUM;
using paa::MUMindex;
using paa::Pair;
using paa::PopBridgeInfo;
using paa::Population;
using paa::Reference;
using paa::Sample;
using paa::Similarity;
using paa::SpaceOut;

#if 1
using MUMdex = paa::FileMUMdex;
using Bridges = paa::FileBridges;
#else
using MUMdex = paa::MUMdex;
using Bridges = paa::Bridges;
#endif

using Anchors = MUMdex::Anchors;
using Bridge = Bridges::Bridge;

const bool verbose{false};
const std::chrono::seconds sleep_duration{1};
const unsigned int max_threads{5};

const bool exclude_snps{true};
const bool allow_skips{true};
bool require_two_or_less{true};
bool require_one_family{true};
bool require_just_kids{true};

unsigned int rare_denovo_family_limit{10000};
const unsigned int min_support_length{20};
const unsigned int min_bridge_count{5};
const unsigned int min_mate_count{1};
const unsigned int min_mate_support{5};
const unsigned int min_parent_coverage{8};
const unsigned int min_parent_mum_length{25};
const unsigned int min_excess_mappability{2};
const unsigned int adjacent_length{10};
const unsigned int adjacent_zero_length{5};
const unsigned int max_pop_seen{100000};

uint64_t n_no_cut{0};
uint64_t n_not_snp{0};
uint64_t n_two_or_less{0};
uint64_t n_one_family{0};
uint64_t n_children{0};
uint64_t n_children_one_family{0};
uint64_t n_no_rare_inherited{0};
uint64_t n_one_parent{0};
uint64_t n_big_count{0};
uint64_t n_good_support{0};
uint64_t n_good_mate_support{0};
uint64_t n_good_excess{0};
uint64_t n_good_pop{0};
uint64_t n_adjacent_cut{0};
uint64_t n_parent_coverage_cut{0};

class SlowChecks {
 public:
  explicit SlowChecks(const Population & pop_arg,
                      const Reference & ref_arg,
                      const unsigned int chromosome_arg,
                      const string & samples_dir) :
      pop{pop_arg}, ref{ref_arg},
      chromosome{chromosome_arg}, chromosome_offset{ref.offset(chromosome)},
    seen_denovo_samples(pop.n_samples()),
    seen_denovo_families(pop.n_families()) {
      collector = async(std::launch::async, std::ref(*this));
      mumdex_names.reserve(pop.n_samples());
      for (const auto s : pop.samples()) {
        const string mumdex_dir{samples_dir + "/" + pop.sample(s) + "/mumdex"};
        mumdex_names.emplace_back(mumdex_dir);
      }

      // Find name and index for chromosome Y
      bool found = false;
      // if (verbose) cerr << ref.n_chromosomes() << endl;
      for (unsigned int c = 0; c != ref.n_chromosomes(); ++c) {
        // if (verbose) cerr << c << " " << ref.name(c) << endl;
        if (ref.name(c).find('Y') != string::npos) {
          chrY = c;
          found = true;
          break;
        }
      }
      if (!found) {
        unique_lock<mutex> done_lock(done_mutex);
        done = true;
        throw Error("Could not find index for Y chromosome");
      }
  }
  ~SlowChecks() {
    try {
      if (verbose) cerr << "enter bridge destructor" << endl;
      // Wait for slow_check processing threads to finish
      unique_lock<mutex> running_futures_lock(running_futures_mutex);
      unique_lock<mutex> finished_futures_lock{finished_futures_mutex};
      while (running_futures.size() || finished_futures.size()) {
        finished_futures_lock.unlock();
        running_futures_lock.unlock();
        std::this_thread::sleep_for(sleep_duration);
        {
          unique_lock<mutex> done_lock(done_mutex);
          if (done) {
            cerr << "Failure" << endl;
            return;
          }
        }
        running_futures_lock.lock();
        finished_futures_lock.lock();
      }
      finished_futures_lock.unlock();
      running_futures_lock.unlock();

      if (verbose) cerr << "signal finished" << endl;
      unique_lock<mutex> done_lock(done_mutex);
      done = true;
      done_lock.unlock();
      if (verbose) cerr << "done set" << endl;
      collector.get();
      cerr << "all done" << endl;
    } catch (Error & e) {
      cerr << "paa::Error:" << endl;
      cerr << e.what() << endl;
    } catch (exception & e) {
      cerr << "std::exception" << endl;
      cerr << e.what() << endl;
    }
  }

  void process(const vector<BridgeInfo> & bridges,
               const vector<Sample> & samples,
               const unsigned int max_bridge_count,
               const unsigned int n_families,
               const unsigned int n_samples,
               const unsigned int total_bridge_count,
               const unsigned int normal_bridge_count,
               const unsigned int normal_seen,
               const unsigned int max_anchor1_support,
               const unsigned int max_anchor2_support,
               const unsigned int max_mate_anchor1_count,
               const unsigned int max_mate_anchor1_support,
               const unsigned int max_mate_anchor2_count,
               const unsigned int max_mate_anchor2_support,
               const unsigned int mum1_map,
               const unsigned int mum2_map,
               const string & pop_string) {
    {
      // Check if done and break if so
      unique_lock<mutex> done_lock(done_mutex);
      if (done) {
        throw Error("Exception thrown somewhere");
      }
    }

    const BridgeInfo & bridge{bridges.front()};
    static unsigned int n_checked{0};
    ++n_checked;
    if (verbose) serr << n_checked << running_futures.size() + 1
                      << "checking" << bridge.chr1() << bridge.pos1() << endl;

    unique_lock<mutex> running_futures_lock(running_futures_mutex);

    // Make sure not too many processes running first
    while (running_futures.size() >= max_threads) {
      running_futures_lock.unlock();
      // if (verbose) std::cerr << "Too many threads - waiting" << endl;

      // Wait for finished future to be ready, or sleep duration
      // spurious wakeups are no problem, so don't test condition
      unique_lock<mutex> finished_futures_lock(finished_futures_mutex);
      finished_future_condition.wait_for(
          finished_futures_lock, sleep_duration);
      finished_futures_lock.unlock();

      running_futures_lock.lock();
    }

    // Create new future
    const auto new_future = running_futures.insert(running_futures.end(),
                                                   future<string>());
    running_futures_lock.unlock();
    *new_future = async(std::launch::async, std::ref(*this),
                        new_future,
                        bridges, samples, max_bridge_count,
                        n_families, n_samples,
                        total_bridge_count, normal_bridge_count,
                        normal_seen,
                        max_anchor1_support, max_anchor2_support,
                        max_mate_anchor1_count,
                        max_mate_anchor1_support,
                        max_mate_anchor2_count,
                        max_mate_anchor2_support,
                        mum1_map, mum2_map, pop_string);
  }

  string operator()(list<future<string>>::iterator this_future,
                    const vector<BridgeInfo> bridges,
                    const vector<Sample> samples,
                    const unsigned int max_bridge_count,
                    const unsigned int n_families,
                    const unsigned int n_samples,
                    const unsigned int total_bridge_count,
                    const unsigned int normal_bridge_count,
                    const unsigned int normal_seen,
                    const unsigned int max_anchor1_support,
                    const unsigned int max_anchor2_support,
                    const unsigned int max_mate_anchor1_count,
                    const unsigned int max_mate_anchor1_support,
                    const unsigned int max_mate_anchor2_count,
                    const unsigned int max_mate_anchor2_support,
                    const unsigned int mum1_map,
                    const unsigned int mum2_map,
                    const string pop_string) try {
    const BridgeInfo & bridge{bridges.front()};
    const Family family{pop.family(samples.front())};
    const vector<Sample> & family_members{pop.samples(family)};

#if 0
    for (unsigned int m{0}; m != family_members.size(); ++m) {
      const Sample member{family_members[m]};
      cout << "To load " << mumdex_names[member] << endl;
    }
#endif

    // Bridge in parent cut, any mum length
    static thread_local vector<Sample> parents;
    parents.clear();
    for (const Sample parent : pop.parent(family)) {
      // ignore check if we already know it is in the parent
      if (find(samples.begin(), samples.end(), parent) != samples.end()) {
        parents.push_back(parent);
        continue;
      }
      const MUMdex & mumdex{mumdex_names[parent], ref};
      if (Bridges{bridge, mumdex}.exist()) {
        if (require_just_kids || n_families > 1) throw string("");
        parents.push_back(parent);
      }
    }
    if (parents.size() > 1) {
      throw string("");
    } else {
      lock_guard<mutex> count_guard{parent_bridge_count_mutex};
      ++n_parent_no_bridges;
    }

    // See if bridge is in other children, get consensus sequence
    static thread_local vector<Sample> children;
    children.clear();
    ConsensusSequence consensus[2]{adjacent_length, adjacent_length};
    for (const Sample child : pop.child(family)) {
      bool in_child = false;
      const MUMdex & mumdex{mumdex_names[child], ref};

      for (const Bridge sbridge : Bridges{bridge, mumdex}) {
        in_child = true;
        unsigned int i{0};
        for (const string & seq : sbridge.adjacent_sequences(adjacent_length)) {
          consensus[i++].add(seq);
        }
      }

      if (in_child) {
        children.push_back(child);
      } else {
        if (find(samples.begin(), samples.end(), child) != samples.end()) {
          throw Error("Seen in child, then not seen in child");
        }
      }
    }
    if (children.size()) for (const bool a2 : {0, 1}) consensus[a2].determine();

    unsigned int min_parent_coverage_seen(-1);
    vector<array<unsigned int, 2>> n_coverage(family_members.size(),
                                              array<unsigned int, 2>{{0, 0}});
    vector<array<unsigned int, 2>> n_anchor(family_members.size(),
                                            array<unsigned int, 2>{{0, 0}});

    // Count reference and anchor in all family members
    for (unsigned int m{0}; m != family_members.size(); ++m) {
      const Sample member{family_members[m]};
      const MUMdex & mumdex{mumdex_names[member], ref};
      // sleep(1);
      for (const unsigned int anchor : {0, 1}) {
        const unsigned int achr{anchor ? bridge.chr2() : bridge.chr1()};
        const unsigned int apos{anchor ? bridge.pos2() : bridge.pos1()};
        const bool ahigh{anchor ? bridge.high2() : bridge.high1()};
        static thread_local vector<uint64_t> seen_pairs;
        seen_pairs.clear();
        const unsigned int max_read_length{154};  // record in mumdex!
        const auto region = mumdex.region(achr, apos > max_read_length ?
                                          apos - max_read_length :
                                          0, achr, apos + 1);
        for (const MUMindex pm : region) {
          const Pair pair{mumdex.pair(pm)};
          if (pair.dupe()) continue;
          const MUM mum{mumdex.mum(pm)};
          if (pair.bad(mum.read_2())) continue;
          if (mum.position0() > apos) continue;
          if (mum.position0() + mum.length() - 1 < apos) continue;

          // look for similar anchor adjacent sequence in parents
          if (require_just_kids && pop.is_parent(member) &&
              ((ahigh && mum.position0() + mum.length() - 1 == apos) ||
               (!ahigh && mum.position0() == apos))) {
            const std::string seq{mumdex.adjacent_sequence(
                pm.pair_index(), mumdex.mum(pm), ahigh, adjacent_length)};
            const Similarity similarity{consensus[anchor].similarity(seq)};
            if ((similarity.n_mismatches() == 0 &&
                 similarity.n_testable() >= adjacent_zero_length) ||
                (similarity.n_mismatches() == 1 &&
                 similarity.n_testable() + 1 >= adjacent_length)) {
              lock_guard<mutex> adjacent_count_guard{adjacent_count_mutex};
              ++n_adjacent_cut;
              throw string("");
            }
          }

          if (achr == mum.chromosome() &&
              ((ahigh == false && apos == mum.position0()) ||
               (ahigh == true && apos == mum.position0() + mum.length() - 1))) {
            ++n_anchor[m][anchor];
          }
          if (mum.length() < min_parent_mum_length) continue;

          // increment coverage if pair not yet seen
          if (find(seen_pairs.begin(), seen_pairs.end(),
                   pm.pair_index()) != seen_pairs.end()) continue;
          seen_pairs.push_back(pm.pair_index());
          ++n_coverage[m][anchor];
        }
        if (pop.is_parent(member)) {
          if (min_parent_coverage_seen > n_coverage[m][anchor]) {
            min_parent_coverage_seen = n_coverage[m][anchor];
          }
          // quit if low parent coverage when looking just for denovos
          if (require_just_kids &&
              !(achr == chrY && pop.nY(member) == 0) &&
              min_parent_coverage_seen < min_parent_coverage) {
            lock_guard<mutex> parent_count_guard{parent_count_mutex};
            ++n_parent_coverage_cut;
            throw string("");
          }
        }
      }
    }

    {
      lock_guard<mutex> to_output_count_guard{to_output_count_mutex};
      ++n_to_output;
    }

    // Output of candidate information
    ostringstream out;
    SpaceOut<ostringstream> ssout(out);
    ssout << pop.family(family);
    if (parents.size()) {
      ssout << pop.member(parents.front());
    } else {
      ssout << "none";
    }
    if (children.size() > 2) {
      ssout << "many";
    } else if (children.size() == 2) {
      ssout << "both";
    } else if (children.size() == 1) {
      ssout << pop.member(children.front());
    } else {
      ssout << "none";
    }

    ssout << n_families << n_samples << parents.size() + children.size()
          << bridge.chr1()
          << ref.name(bridge.chr1()) << bridge.pos1() << bridge.high1()
          << ref.name(bridge.chr2()) << bridge.pos2() << bridge.high2()
          << bridge.description() << bridge.orientation_char()
          << bridge.invariant() << bridge.offset()
          << max_bridge_count
          << max_anchor1_support << max_anchor2_support
          << max_mate_anchor1_count << max_mate_anchor1_support
          << max_mate_anchor2_count << max_mate_anchor2_support
          << mum1_map << mum2_map << min_parent_coverage_seen;

    for (unsigned int m{0}; m != family_members.size(); ++m) {
      if (m) {
        out << ",";
      } else {
        out << " ";
      }
      const Sample sample{family_members[m]};
      const vector<Sample>::const_iterator found{
        find(samples.begin(), samples.end(), sample)};
      if (found == samples.end()) {
        out << 0;
      } else {
        out << bridges[found - samples.begin()].bridge_count();
      }
    }

    for (const bool anchor2 : {false, true}) {
      out << " ";
      for (unsigned int m{0}; m != family_members.size(); ++m) {
        if (m) out << ",";
        out << n_anchor[m][anchor2];
      }
    }

    for (const bool anchor2 : {false, true}) {
      out << " ";
      for (unsigned int m{0}; m != family_members.size(); ++m) {
        if (m) out << ",";
        out << n_coverage[m][anchor2];
      }
    }

    out << " " << pop_string << " " << total_bridge_count << " "
        << normal_bridge_count << " " << normal_seen;

    lock_guard<mutex> denovo_count_guard{denovo_count_mutex};

    // denovo or shared between siblings
    if (parents.size() == 0) {
      ++n_denovo;
      ++seen_denovo_families[family];
      for (const Sample sample : children) ++seen_denovo_samples[sample];
      if (children.size() == 1) {
        if (pop.is_proband(children.front())) {
          ++n_denovo_proband;
        } else {
          ++n_denovo_sibling;
        }
      } else {
        ++n_denovo_siblings;
      }
    } else if (children.size() == 0) {
      ++n_transmission_none;
    } else if (children.size() >= 2) {
      ++n_transmission_both;
    } else if (pop.is_proband(children.front())) {
      ++n_denovo_proband;
    } else {
      ++n_denovo_sibling;
    }

    throw out.str();
  } catch (const string & out) {
    {
      unique_lock<mutex> running_futures_lock{running_futures_mutex};
      unique_lock<mutex> finished_futures_lock{finished_futures_mutex};
      finished_futures.insert(finished_futures.end(), move(*this_future));
      running_futures.erase(this_future);
    }
    finished_future_condition.notify_all();
    return out;
  } catch (Error & e) {
    cerr << "paa::Error:" << endl;
    cerr << e.what() << endl;
    cerr << "At " << bridges.front().pos1() << endl;
    unique_lock<mutex> done_lock(done_mutex);
    done = true;
    throw;
  } catch (exception & e) {
    cerr << "std::exception" << endl;
    cerr << e.what() << endl;
    unique_lock<mutex> done_lock(done_mutex);
    done = true;
    throw;
  }

  void operator()() {
    while (true) {
      unique_lock<mutex> finished_futures_lock(finished_futures_mutex);

      // Possibly wait for notice that a future is finished
      if (finished_futures.empty()) {
        finished_future_condition.wait_for(
            finished_futures_lock, sleep_duration,
            [this]{return finished_futures.size();});
      }

      // Clear out finished_futures if there are any
      while (finished_futures.size()) {
        finished_futures_lock.unlock();
        const string out{finished_futures.front().get()};
        finished_futures_lock.lock();
        if (out.size()) cout << out << endl;
        finished_futures.pop_front();
      }
      finished_futures_lock.unlock();

      // Check if done and break if so
      unique_lock<mutex> done_lock(done_mutex);
      if (done) break;
    }

    // summary info
    for (const auto s : pop.samples()) {
      if (pop.is_child(s)) {
        const Family f{pop.family(s)};
        serr << s
             << pop.sample(s) << seen_denovo_samples[s]
             << pop.family(f) << seen_denovo_families[f]
             << endl;
      }
    }

    serr << "cuts"
         << n_no_cut
         << n_not_snp
         << n_two_or_less
         << n_good_pop
         << n_one_family
         << n_children
         << n_children_one_family
         << n_no_rare_inherited
         << n_one_parent
         << n_big_count
         << n_good_support
         << n_good_mate_support
         << n_good_excess
         << n_parent_no_bridges
         << n_adjacent_cut
         << n_parent_coverage_cut
         << n_to_output
         << endl;

    serr << "denovos"
         << n_denovo << "all"
         << n_denovo_siblings << "both"
         << n_denovo_proband << "proband"
         << n_denovo_sibling << "sibling"
         << endl;

    serr << "transmission"
         << n_transmission_both << "both"
         << n_transmission_proband << "proband"
         << n_transmission_sibling << "sibling"
         << n_transmission_none << "none" << endl;

    const double genome_ratio{static_cast<double>(ref.size()) /
          ref.size(chromosome)};
    serr << "genome projection denovo "
         << static_cast<unsigned int>(genome_ratio * n_denovo) << endl;
  }

 private:
  const Population & pop;
  const Reference & ref;
  const unsigned int chromosome{};
  const unsigned int chromosome_offset{};
  unsigned int chrY{};
  vector<string> mumdex_names{};
  future<void> collector{};
  mutex parent_bridge_count_mutex{};
  mutex to_output_count_mutex{};
  mutex denovo_count_mutex{};
  mutex adjacent_count_mutex{};
  mutex parent_count_mutex{};

  mutex running_futures_mutex{};
  list<future<string>> running_futures{};

  mutex finished_futures_mutex{};
  list<future<string>> finished_futures{};
  condition_variable finished_future_condition{};

  mutex done_mutex{};
  bool done{false};

  vector<Sample> seen_denovo_samples{};
  vector<Family> seen_denovo_families{};
  uint64_t n_denovo_proband{0};
  uint64_t n_denovo_sibling{0};
  uint64_t n_parent_no_bridges{0};
  uint64_t n_to_output{0};
  uint64_t n_denovo_siblings{0};
  uint64_t n_denovo{0};
  uint64_t n_transmission_none{0};
  uint64_t n_transmission_both{0};
  uint64_t n_transmission_proband{0};
  uint64_t n_transmission_sibling{0};
};

int main(int argc, char* argv[])  try {
  --argc;
  if (argc != 7 && argc != 9) {
    throw Error("usage: population_bridges type family_file bridges_dir "
                "ref samples_dir use_pop chromosome [start stop]");
  }

  // Set analysis type
  const string type{argv[1]};
  if (type == "rare_denovo") {
    require_two_or_less = false;
    require_one_family = false;
    require_just_kids = true;
    rare_denovo_family_limit = 5;
  } else if (type == "ultra_rare_denovo") {
    require_two_or_less = true;
    require_one_family = true;
    require_just_kids = true;
  } else if (type == "denovo") {
    require_two_or_less = false;
    require_one_family = false;
    require_just_kids = true;
  } else if (type == "ultra_rare") {
    require_two_or_less = false;
    require_one_family = true;
    require_just_kids = false;
  } else if (type == "ultra_rare_and_denovo") {
    require_two_or_less = false;
    require_one_family = false;
    require_just_kids = false;
  } else {
    throw Error("Unknown analysis type") << type;
  }

  // Other command line arguments
  const string family_file{argv[2]};
  const Population pop{family_file};
  const string bridges_dir{argv[3]};
  const string reference_file{argv[4]};
  const Reference ref{reference_file};
  const ChromosomeIndexLookup chr_lookup{ref};
  const Mappability map{reference_file, true};
  const string samples_dir{argv[5]};
  const bool use_pop{static_cast<bool>(atoi(argv[6]))};
  const string chromosome_name{argv[7]};
  const unsigned int chromosome{chr_lookup[chromosome_name]};
  const unsigned int chromosome_offset{ref.offset(chromosome)};
  const unsigned int start{argc == 9 ?
        static_cast<unsigned int>(atoi(argv[8])) : 0};
  const unsigned int stop{argc == 9 ?
        static_cast<unsigned int>(atoi(argv[9])) : ref.size(chromosome)};

  // Parallel process candidate checks
  SlowChecks slow_checks(pop, ref, chromosome, samples_dir);

  // Set up queue of bridge files
  deque<MergeHelper> helpers;
  using PQueue = priority_queue<MergeHelper*, vector<MergeHelper *>,
      MergeHelperCompare>;
  PQueue queue{MergeHelperCompare()};
  unsigned int n_samples_skipped{0};
  for (const auto s : pop.samples()) {
    ostringstream bridges_name;
    bridges_name << bridges_dir << "/" << pop.sample(s) << "/"
                 << get_bridge_file_name(ref, chromosome);
    MergeHelper helper{bridges_name.str(), s, start, stop};
    if (helper.valid_start()) {
      helpers.push_back(move(helper));
      queue.push(&helpers.back());
    } else {
      ++n_samples_skipped;
    }
    serr << pop.family(pop.family(s))
         << pop.sample(s)
         << pop.member(s)
         << bridges_name.str()
         << queue.size() << endl;
  }

  if (!allow_skips && n_samples_skipped)
    throw Error("Samples skipped") << n_samples_skipped;

  // Load PopBridgeInfo file, and advance to start location, if called for
  ostringstream pop_file_name;
  pop_file_name << bridges_dir << "/popbridges."
                << chromosome_name << ".bin";
  FileVector<PopBridgeInfo> pop_info{use_pop ?
        pop_file_name.str() : string{"/dev/null"}};
  FileVector<PopBridgeInfo>::const_iterator pop_bridge{use_pop ?
        lower_bound(pop_info.begin(), pop_info.end(), start,
                    [](const PopBridgeInfo & bridge, const unsigned int val) {
                      return bridge.pos1() < val;
                    }) : pop_info.begin()};

  // Document the run
  serr << "Run parameters:" << endl
       << "  type" << type << endl
       << "  family_file" << family_file << endl
       << "  samples_dir" << samples_dir << endl
       << "  bridges_dir" << bridges_dir << endl
       << "  use_pop" << use_pop << endl
       << "  chromosome" << chromosome_name << endl
       << "  start" << start << endl
       << "  stop" << stop << endl
       << "  n_families" << pop.n_families() << endl
       << "  n_samples" << pop.n_samples() << endl
       << "  n_samples_skipped" << n_samples_skipped << endl
       << "  rare_denovo_family_limit" << rare_denovo_family_limit << endl
       << "  require_two_or_less" << require_two_or_less << endl
       << "  require_one_family" << require_one_family << endl
       << "  require_just_kids" << require_just_kids << endl
       << "  exclude_snps" << exclude_snps << endl
       << "  min_support_length" << min_support_length << endl
       << "  min_bridge_count" << min_bridge_count << endl
       << "  min_mate_count" << min_mate_count << endl
       << "  min_mate_support" << min_mate_support << endl
       << "  min_parent_coverage" << min_parent_coverage << endl
       << "  min_parent_mum_length" << min_parent_mum_length << endl
       << "  min_excess_mappability" << min_excess_mappability << endl
       << "  adjacent_length" << adjacent_length << endl
       << "  adjacent_zero_length" << adjacent_zero_length << endl
       << "  max_pop_seen" << max_pop_seen << endl;

    sout << "family" << "parent" << "kids"
         << "nFam" << "nSam" << "nInFam"
         << "chr"
         << "chrA" << "posA" << "highA"
         << "chrB" << "posB" << "highB"
         << "type" << "ori"
         << "invariant" << "offset"
         << "bridge"
         << "supA" << "supB"
         << "mCA" << "mSA"
         << "mCB" << "mSB"
         << "mapA" << "mapB" << "minParCov"
         << "bridges"
         << "anchorsA" << "anchorsB"
         << "coveragesA" << "coveragesB"
         << "pNP" << "pNB" << "pMed" << "pMax" << "tBC" << "nBC" << "nS"
         << endl;

  // Loop, adding one bridge at a time to all_bridges if the same event
  vector<BridgeInfo> all_bridges;
  vector<Sample> all_samples;
  unsigned int total_bridge_count{0};
  unsigned int normal_bridge_count{0};
  unsigned int normal_seen{0};
  bool reset{true};
  while (true) {
    // New bridge expected - start fresh
    if (reset) {
      all_bridges.clear();
      all_samples.clear();
      total_bridge_count = 0;
      normal_bridge_count = 0;
      normal_seen = 0;
      reset = false;
    }

    // Add a bridge to the list if the same event
    bool bridge_done{true};
    if (queue.size()) {
      MergeHelper * top{queue.top()};
      const BridgeInfo current{top->current()};
      if (all_bridges.empty() || !(all_bridges.back() < current)) {
        ++n_no_cut;
        bridge_done = false;
        all_bridges.push_back(current);
        total_bridge_count += current.bridge_count();
        const Sample sample{top->sample()};
        if (pop.is_parent(sample) || pop.is_normal(sample) ||
            pop.is_matched(sample)) {
          normal_bridge_count += current.bridge_count();
          ++normal_seen;
        }
        all_samples.emplace_back(sample);
        queue.pop();
        if (top->advance()) {
          queue.push(top);
        }
      }
      if (queue.empty()) bridge_done = true;
      if (bridge_done) reset = true;
    } else {
      if (all_samples.empty()) {
        if (verbose) cerr << "bridge queue empty" << endl;
        break;
      }
    }

    // No new bridges added, so bridge list is ready to process
    if (bridge_done) {
      // The first bridge, the exemplar since all are the same
      const BridgeInfo & bridge{all_bridges[0]};

      // SNP cut
      const bool is_snp{bridge.chr1() == bridge.chr2() &&
            bridge.invariant() == 0 && bridge.high1() != bridge.high2()};
      if (is_snp) {
        if (exclude_snps) continue;
      } else {
        ++n_not_snp;
      }
      // cerr << n_not_snp << " " << n_good_excess << endl;

      // More than two cut
      if (all_samples.size() <= 2) {
        ++n_two_or_less;
      } else {
        if (require_two_or_less) continue;
      }

      // Count families seen
      const unsigned int n_families{[&all_samples, &pop] {
          unsigned int n{0};
          Family last_family{100000};
          for (const Sample sample : all_samples) {
            const Family family{pop.family(sample)};
            if (family != last_family) {
              ++n;
              last_family = family;
            }
          }
          return n;
        }()};

      // Multiple family cut
      if (n_families == 1) {
        ++n_one_family;
      } else {
        if (require_one_family) continue;
        if (n_families > rare_denovo_family_limit) continue;
      }

      // Check population file for event
      const string default_pop_string{"0 0 0 0"};
      string pop_string;
      if (use_pop) {
        while (pop_bridge != pop_info.end() && *pop_bridge < bridge) {
          ++pop_bridge;
        }
      }
      if (use_pop && pop_bridge != pop_info.end() &&
          !(bridge < *pop_bridge)) {
        // Found matching event
        ostringstream pop_out;
        pop_out << (*pop_bridge).n_people() << " "
                << (*pop_bridge).n_bridges() << " "
                << (*pop_bridge).median_bridges() << " "
                << (*pop_bridge).max_bridges();
        pop_string = pop_out.str();
        if ((*pop_bridge).n_people() > max_pop_seen) {
          continue;
        }
      } else {
        pop_string = default_pop_string;
      }
      ++n_good_pop;

      // Get mappability information
      const unsigned int mum1_abspos{chromosome_offset + bridge.pos1()};
      const unsigned int mum2_abspos{ref.offset(bridge.chr2()) + bridge.pos2()};
      const unsigned int mum1_map{map.low_high(bridge.high1(), mum1_abspos)};
      const unsigned int mum2_map{map.low_high(bridge.high2(), mum2_abspos)};

      // Look at each family separately
      const unsigned int n_samples{static_cast<unsigned int>(
          all_samples.size())};
      while (all_samples.size()) {
        // Gather samples and bridges for family
        static vector<Sample> samples;
        static vector<BridgeInfo> bridges;
        samples.clear();
        bridges.clear();
        while (all_samples.size() &&
               (samples.empty() ||
                pop.family(samples.back()) ==
                pop.family(all_samples.back()))) {
          samples.push_back(all_samples.back());
          all_samples.pop_back();
          bridges.push_back(all_bridges.back());
          all_bridges.pop_back();
        }

        // Parent sample cuts
        bool just_kids{true};
        unsigned int n_parents{0};
        for (const Sample sample : samples) {
          if (pop.is_parent(sample)) {
            just_kids = false;
            ++n_parents;
          }
        }
        if (just_kids) {
          ++n_children;
          if (n_families == 1) {
            ++n_children_one_family;
          }
        } else {
          // Seen in parent
          // Cut if we only want denovos or if not ultra-rare
          if (require_just_kids || n_families > 1) continue;
          ++n_no_rare_inherited;
          if (n_parents > 1) continue;
          ++n_one_parent;
        }

        // Gather max support counts and lengths
        unsigned int max_bridge_count{0};
        unsigned int max_anchor1_support{0};
        unsigned int max_anchor2_support{0};
        unsigned int max_mate_anchor1_count{0};
        unsigned int max_mate_anchor1_support{0};
        unsigned int max_mate_anchor2_count{0};
        unsigned int max_mate_anchor2_support{0};
        for (const BridgeInfo & sample_bridge : bridges) {
          if (max_bridge_count < sample_bridge.bridge_count()) {
            max_bridge_count = sample_bridge.bridge_count();
          }
          if (max_anchor1_support < sample_bridge.anchor1_length()) {
            max_anchor1_support = sample_bridge.anchor1_length();
          }
          if (max_anchor2_support < sample_bridge.anchor2_length()) {
            max_anchor2_support = sample_bridge.anchor2_length();
          }
          if (max_mate_anchor1_count < sample_bridge.mate_anchor1_count()) {
            max_mate_anchor1_count = sample_bridge.mate_anchor1_count();
          }
          if (max_mate_anchor2_count < sample_bridge.mate_anchor2_count()) {
            max_mate_anchor2_count = sample_bridge.mate_anchor2_count();
          }
          if (max_mate_anchor1_support < sample_bridge.mate_anchor1_length()) {
            max_mate_anchor1_support = sample_bridge.mate_anchor1_length();
          }
          if (max_mate_anchor2_support < sample_bridge.mate_anchor2_length()) {
            max_mate_anchor2_support = sample_bridge.mate_anchor2_length();
          }
        }

        // Low bridge count cut
        if (max_bridge_count < min_bridge_count) continue;
        ++n_big_count;

        // Short anchor support cut
        if (max_anchor1_support < min_support_length ||
            max_anchor2_support < min_support_length) continue;
        ++n_good_support;

        // Short mate support cut
        if (max_mate_anchor1_count < min_mate_count ||
            max_mate_anchor2_count < min_mate_count ||
            max_mate_anchor1_support < min_mate_support ||
            max_mate_anchor2_support < min_mate_support) continue;
        ++n_good_mate_support;

        // Short excess mappability cut
        if (max_anchor1_support < mum1_map) throw Error("Mappability");
        if (max_anchor1_support < min_excess_mappability + mum1_map) {
          continue;
        } else {
          if (max_anchor2_support < min_excess_mappability +  mum2_map) {
            continue;
          }
        }
        ++n_good_excess;
        // cout << n_good_excess << endl;

        slow_checks.process(bridges, samples, max_bridge_count,
                            n_families, n_samples, total_bridge_count,
                            normal_bridge_count, normal_seen,
                            max_anchor1_support, max_anchor2_support,
                            max_mate_anchor1_count, max_mate_anchor1_support,
                            max_mate_anchor2_count, max_mate_anchor2_support,
                            mum1_map, mum2_map, pop_string);
      }
    }
  }

  cerr << "finished bridge loop" << endl;

  return 0;
} catch (Error & e) {
  cerr << "paa::Error:" << endl;
  cerr << e.what() << endl;
  return 1;
} catch (exception & e) {
  cerr << "std::exception" << endl;
  cerr << e.what() << endl;
  return 1;
} catch (...) {
  cerr << "unknown exception was caught" << endl;
  return 1;
}
```
