## Supplementary material for "G-Graph: An interactive genomic graph viewer": MUMdex Source Code for G-Graph: population_database.cpp

```
//
// population_database
//
// Create the population database
//
// Copyright 2016 Peter Andrews @ CSHL
//

#include <algorithm>
#include <deque>
#include <exception>
#include <iostream>
#include <future>
#include <limits>
#include <queue>
#include <sstream>
#include <string>
#include <utility>
#include <vector>

#include "bridges.h"
#include "error.h"
#include "files.h"
#include "mumdex.h"
#include "population.h"
#include "utility.h"

using std::async;
using std::cerr;
using std::cout;
using std::deque;
using std::endl;
using std::exception;
using std::future;
using std::lower_bound;
using std::move;
using std::ostringstream;
using std::priority_queue;
using std::ref;
using std::string;
using std::vector;

using paa::bwritec;
using paa::serr;
using paa::sout;
using paa::BridgeInfo;
using paa::ChromosomeIndexLookup;
using paa::Error;
using paa::FileVector;
using paa::MergeHelper;
using paa::MergeHelperCompare;
using paa::PopBridgeInfo;
using paa::Population;
using paa::Reference;
using paa::Sample;

int main(int argc, char* argv[])  try {
  --argc;
  if (argc != 6) {
    throw Error("usage: population_database ref family_file bridges_dir "
                "chromosome start stop");
  }

  const Reference ref{argv[1]};
  const ChromosomeIndexLookup chr_lookup{ref};
  const Population pop{argv[2]};
  const string bridges_dir{argv[3]};
  const string chromosome_name{argv[4]};
  const unsigned int start{static_cast<unsigned int>(atoi(argv[5]))};
  const unsigned int stop{static_cast<unsigned int>(atoi(argv[6]))};
  const unsigned int chromosome{chr_lookup[chromosome_name]};

  vector<string> sample_dirs;
  for (const Sample sample : pop.samples()) {
    if (pop.is_parent(sample) || pop.is_normal(sample) ||
        pop.is_source(sample) || pop.is_matched(sample) ||
        pop.is_before(sample)) {
      sample_dirs.push_back(bridges_dir + "/" + pop.sample(sample));
    }
  }

  deque<MergeHelper> helpers;
  using PQueue = priority_queue<MergeHelper*, vector<MergeHelper *>,
      MergeHelperCompare>;
  PQueue queue{MergeHelperCompare()};
  uint64_t n_saved{0};
  for (const string & dir : sample_dirs) {
    ostringstream bridges_name;
    bridges_name << dir << "/" << get_bridge_file_name(ref, chromosome);
    MergeHelper helper{bridges_name.str(), Sample{0}, start, stop};
    if (helper.valid_start()) {
      helpers.push_back(move(helper));
      queue.push(&helpers.back());
    }
  }

  ostringstream output_file_name;
  output_file_name << "popbridges." << chromosome_name << "."
                   << start << "-" << stop << ".bin";
  FILE * output_file{fopen(output_file_name.str().c_str(), "wb")};
  if (output_file == nullptr) {
    throw Error("Could not open output file") << output_file_name.str();
  }

  // merge bridge info
  if (queue.size()) {
    vector<PopBridgeInfo> merged;
    vector<PopBridgeInfo> to_save;
    vector<unsigned int> counts;
    future<void> saver;
    while (true) {
      MergeHelper * lowest{queue.top()};
      const BridgeInfo current{lowest->current()};
      queue.pop();
      if (lowest->advance()) {
        queue.push(lowest);
      }
      const bool empty{queue.empty()};

      // Finish off completed entries
      if (merged.size() && merged.back() < current && counts.size()) {
        if (merged.back().invariant() == 0) {
          merged.pop_back();
        } else {
          merged.back().finalize(counts);
        }
        counts.clear();
      }
      counts.push_back(current.bridge_count());

      if (merged.empty() || merged.back() < current) {
        if (merged.size() &&
            merged.back().n_people() == merged.back().n_bridges()) {
          merged.pop_back();
        }
        merged.emplace_back(current);
      } else {
        merged.back().combine(current);
      }
      const uint64_t max_merged{1024 * 1024};
      if (merged.size() == max_merged || empty) {
        if (merged.size() && empty) {
          merged.back().finalize(counts);
        }
        if (saver.valid()) {
          saver.get();
        }
        const PopBridgeInfo last_entry{merged.back()};
        if (!empty || last_entry.n_people() == last_entry.n_bridges()) {
          merged.pop_back();
        }
        n_saved += merged.size();
        to_save.swap(merged);
        merged.clear();
        if (!empty) merged.push_back(last_entry);
        saver = async(
            std::launch::async,
            [output_file]
            (const vector<PopBridgeInfo> & tosave) {
              if (tosave.size()) {
                bwritec(output_file, &tosave[0], "bridges",
                        tosave.size() * sizeof(PopBridgeInfo));
              }
            }, std::ref(to_save));
        if (empty) {
          break;
        }
      }
    }
    if (saver.valid()) {
      saver.get();
    }
  }
  fclose(output_file);

  cerr << "saved " << n_saved << " bridges" << endl;

  return 0;
} catch (Error & e) {
  cerr << "paa::Error:" << endl;
  cerr << e.what() << endl;
  return 1;
} catch (exception & e) {
  cerr << "std::exception" << endl;
  cerr << e.what() << endl;
  return 1;
} catch (...) {
  cerr << "unknown exception was caught" << endl;
  return 1;
}
```
