## Supplementary material for "G-Graph: An interactive genomic graph viewer": MUMdex Source Code for G-Graph: population_denovos.cpp

```
//
// population_denovos
//
// examine bridge information over a population to find denovo candidates
//
// Copyright 2018 Peter Andrews @ CSHL
//

// TO DO
//
// consider opportunity for mate support cut!
//
//
// N anchors
//
// nearby bridges TODO
// reduce reference requirement for chrY, chrX bridges in Male MAYBE
// overlap cut PROB NOT
// consolidation of bridges - spurious removal NO
// reduction of reference in sample with candidate? NO
// mate support cut OK
// reference count in parents OK
// mappability excess OK
// bridge in parents with no support cuts or mum length cuts OK
// similar anchors in parents with no support cuts or mum length cuts OK
// do not check mother for chrY bridges OK

// cuts:
//
// need to handle family structure, additional normals,
// cancer and matched normal, twins, technical replicates
// edited variants (crispr)
//
// eliminating cuts
//   bridge counts
//   anchor support
//   mate support
//   seen in matched normal or replicate
//   excess mappability - spurious likely
//   parent reference count
//   N-base anchor extension (similar sequence to bridge in parents)
//
// discriminating cuts: classes of candidates
//   seen in population - still possible denovo, rare vs ultra rare
//   seen in parent - denovo, inherited, mendel testing
//     parent bridge check
//     parent anchor check
//

// internal tests
//
// both percentage
// transmission mendel agreement
// percent seen in population
// microsatellite identity
//
// external tests
//
// compare with Ivan
// compare with Yoon-Ha, Chris
// validation
//

#include <algorithm>
#include <array>
#include <chrono>
#include <exception>
#include <functional>
#include <future>
#include <iostream>
#include <mutex>
#include <sstream>
#include <string>
#include <utility>
#include <vector>

#include "block_reader.h"
#include "bridges.h"
#include "error.h"
#include "files.h"
#include "mumdex.h"
#include "population.h"
#include "sequence.h"
#include "threads.h"
#include "utility.h"

using std::array;
using std::cerr;
using std::cout;
using std::defer_lock;
using std::endl;
using std::exception;
using std::flush;
using std::future;
using std::lock_guard;
using std::lower_bound;
using std::mutex;
using std::ostream;
using std::ostringstream;
using std::pair;
using std::string;
using std::unique_lock;
using std::vector;

using paa::file_size;
using paa::get_block_size;
using paa::serr;
using paa::sout;
using paa::BridgeInfo;
using BlockMerger = paa::BlockMerger<BridgeInfo>;
using BlockReader = paa::BlockReader<BridgeInfo>;
using paa::ChromosomeIndexLookup;
using paa::ConsensusSequence;
using paa::Error;
using paa::Family;
using paa::FileVector;
using paa::Mappability;
using paa::MUM;
using paa::MUMindex;
using paa::Pair;
using paa::Population;
using paa::Reference;
using paa::Sample;
using paa::Similarity;
using paa::ThreadPool;

#define USE_FILES 1

#if USE_FILES
using MUMdex = paa::FileMUMdex;
using Bridges = paa::FileBridges;
#else
using MUMdex = paa::MUMdex;
using Bridges = paa::Bridges;
#endif

using Anchors = MUMdex::Anchors;
using Bridge = Bridges::Bridge;

const bool verbose{false};

const bool exclude_snps{true};

const double denovo_family_fraction{0.02};
const unsigned int denovo_min_limit{5};
unsigned int denovo_family_limit{10000};
const unsigned int min_support_length{25};
const unsigned int min_bridge_count{5};
const unsigned int min_mate_count{1};
const unsigned int min_mate_support{20};
const unsigned int min_parent_coverage{10};
const unsigned int min_parent_mum_length{25};
const unsigned int min_excess_mappability{1};
const unsigned int adjacent_length{10};
const unsigned int adjacent_zero_length{5};

uint64_t n_no_cut{0};
uint64_t n_not_snp{0};
uint64_t n_one_family{0};
uint64_t n_children{0};
uint64_t n_children_one_family{0};
uint64_t n_big_count{0};
uint64_t n_good_support{0};
uint64_t n_good_mate_support{0};
uint64_t n_good_excess{0};
uint64_t n_adjacent_cut{0};
uint64_t n_parent_coverage_cut{0};

class SlowChecks {
 public:
  explicit SlowChecks(const Population & pop_arg,
                      const Reference & ref_arg,
                      const unsigned int chromosome_arg,
                      const string & samples_dir) :
      pop{pop_arg}, ref{ref_arg},
      chromosome{chromosome_arg}, chromosome_offset{ref.offset(chromosome)},
    chrY{ref.find_y_chromosome()},
    seen_denovo_samples(pop.n_samples()),
    seen_denovo_families(pop.n_families()),
    family_locks(pop.n_families()) {
      mumdex_names.reserve(pop.n_samples());
      for (const auto s : pop.samples()) {
        const string mumdex_dir{samples_dir + "/" + pop.sample(s) + "/mumdex"};
        mumdex_names.emplace_back(mumdex_dir);
      }
  }

  void operator()(const vector<BridgeInfo> bridges,
                  const vector<Sample> samples,
                  const unsigned int max_bridge_count,
                  const unsigned int n_families,
                  const unsigned int n_samples,
                  const unsigned int total_bridge_count,
                  const unsigned int parent_bridge_count,
                  const unsigned int parent_seen,
                  const unsigned int child_bridge_count,
                  const unsigned int child_seen,
                  const unsigned int max_anchor1_support,
                  const unsigned int max_anchor2_support,
                  const unsigned int max_mate_anchor1_count,
                  const unsigned int max_mate_anchor1_support,
                  const unsigned int max_mate_anchor2_count,
                  const unsigned int max_mate_anchor2_support,
                  const unsigned int mum1_map,
                  const unsigned int mum2_map,
                  vector<unsigned int> parent_counts,
                  vector<unsigned int> children_counts) {
    const BridgeInfo & bridge{bridges.front()};
    const Family family{pop.family(samples.front())};
    unique_lock<mutex> family_lock{family_locks[family], defer_lock};
    if (USE_FILES) family_lock.lock();
    const vector<Sample> & family_members{pop.samples(family)};

    // Confirm bridge is in child, get consensus sequence
    ConsensusSequence consensus[2]{adjacent_length, adjacent_length};
    for (const Sample child : samples) {
      bool in_child = false;
      const MUMdex & mumdex{mumdex_names[child], ref};
      for (const Bridge sbridge : Bridges{bridge, mumdex}) {
        in_child = true;
        unsigned int i{0};
        for (const string & seq : sbridge.adjacent_sequences(adjacent_length))
          consensus[i++].add(seq);
      }
      if (!in_child) throw Error("Seen in child, then not seen in child");
    }
    for (const bool a2 : {0, 1}) consensus[a2].determine();

    unsigned int min_parent_coverage_seen(-1);
    vector<array<unsigned int, 2>> n_coverage(family_members.size(),
                                              array<unsigned int, 2>{{0, 0}});
    vector<array<unsigned int, 2>> n_anchor(family_members.size(),
                                            array<unsigned int, 2>{{0, 0}});

    // Count reference and anchor in all family members
    for (unsigned int m{0}; m != family_members.size(); ++m) {
      const Sample member{family_members[m]};
      const MUMdex & mumdex{mumdex_names[member], ref};
      for (const unsigned int anchor : {0, 1}) {
        const unsigned int achr{anchor ? bridge.chr2() : bridge.chr1()};
        const unsigned int apos{anchor ? bridge.pos2() : bridge.pos1()};
        const bool ahigh{anchor ? bridge.high2() : bridge.high1()};
        static thread_local vector<uint64_t> seen_pairs;
        seen_pairs.clear();
        const unsigned int max_read_length{154};  // record in mumdex!
        const auto region = mumdex.region(achr, apos > max_read_length ?
                                          apos - max_read_length :
                                          0, achr, apos + 1);
        for (const MUMindex pm : region) {
          const Pair pair{mumdex.pair(pm)};
          if (pair.dupe()) continue;
          const MUM mum{mumdex.mum(pm)};
          if (pair.bad(mum.read_2())) continue;
          if (mum.position0() > apos) continue;
          if (mum.position0() + mum.length() - 1 < apos) continue;

          // look for similar anchor adjacent sequence in parents
          if (pop.is_parent(member) &&
              ((ahigh && mum.position0() + mum.length() - 1 == apos) ||
               (!ahigh && mum.position0() == apos))) {
            const std::string seq{mumdex.adjacent_sequence(
                pm.pair_index(), mumdex.mum(pm), ahigh, adjacent_length)};
            const Similarity similarity{consensus[anchor].similarity(seq)};
            if ((similarity.n_mismatches() == 0 &&
                 similarity.n_testable() >= adjacent_zero_length) ||
                (similarity.n_mismatches() == 1 &&
                 similarity.n_testable() + 1 >= adjacent_length)) {
              lock_guard<mutex> adjacent_count_guard{adjacent_count_mutex};
              ++n_adjacent_cut;
              return;
            }
          }

          if (achr == mum.chromosome() &&
              ((ahigh == false && apos == mum.position0()) ||
               (ahigh == true && apos == mum.position0() + mum.length() - 1)))
            ++n_anchor[m][anchor];
          if (mum.length() < min_parent_mum_length) continue;

          // increment coverage if pair not yet seen
          if (find(seen_pairs.begin(), seen_pairs.end(),
                   pm.pair_index()) != seen_pairs.end()) continue;
          seen_pairs.push_back(pm.pair_index());
          ++n_coverage[m][anchor];
        }
        if (pop.is_parent(member)) {
          if (min_parent_coverage_seen > n_coverage[m][anchor]) {
            min_parent_coverage_seen = n_coverage[m][anchor];
          }
          // quit if low parent coverage when looking just for denovos
          if (!(achr == chrY && pop.nY(member) == 0) &&
              min_parent_coverage_seen < min_parent_coverage) {
            lock_guard<mutex> parent_count_guard{parent_count_mutex};
            ++n_parent_coverage_cut;
            return;
          }
        }
      }
    }

    {
      lock_guard<mutex> to_output_count_guard{to_output_count_mutex};
      ++n_to_output;
    }

    // Output of candidate information
    ostringstream out;

    // individuals info
    out << pop.family(family);
    for (unsigned int s{0}; s != samples.size(); ++s)
      out << (s ? "," : " ") << pop.sample(samples[s]);
    for (unsigned int s{0}; s != samples.size(); ++s)
      out << (s ? "," : " ") << pop.sex(samples[s]);
    out << " ";
    if (samples.size() > 2) {
      out << "many";
    } else if (samples.size() == 2) {
      out << "both";
    } else if (samples.size() == 1) {
      out << pop.member(samples.front());
    } else {
      throw Error("Unexpected no samples");
    }
    out << " " << samples.size() << " ";
    for (unsigned int m{0}; m != family_members.size(); ++m) {
      const Sample member{family_members[m]};
      if (pop.is_child(member)) out << pop.member(member)[0];
    }

    // bridge info
    out << " " << bridge.chr1()
        << " " << ref.name(bridge.chr1())
        << " " << bridge.pos1()
        << " " << bridge.high1()
        << " " << ref.name(bridge.chr2())
        << " " << bridge.pos2()
        << " " << bridge.high2()
        << " " << bridge.invariant()
        << " " << bridge.offset()
        << " " << bridge.description()
        << " " << bridge.orientation_char();

    // anchor info
    out << " " << max_anchor1_support
        << " " << max_anchor2_support
        << " " << max_anchor1_support - mum1_map
        << " " << max_anchor2_support - mum2_map
        << " " << mum1_map
        << " " << mum2_map
        << " " << max_mate_anchor1_count
        << " " << max_mate_anchor2_count
        << " " << max_mate_anchor1_support
        << " " << max_mate_anchor2_support;

    // bridge counts
    out << " " << max_bridge_count;
    vector<unsigned int> kids_counts;
    for (unsigned int m{0}; m != family_members.size(); ++m) {
      out << (m ? "," : " ");
      const Sample sample{family_members[m]};
      const vector<Sample>::const_iterator found{
        find(samples.begin(), samples.end(), sample)};
      if (found == samples.end()) {
        out << 0;
      } else {
        const unsigned int count{
          bridges[found - samples.begin()].bridge_count()};
        kids_counts.push_back(count);
        out << count;
      }
    }

    // anchor counts
    for (const bool anchor2 : {false, true})
      for (unsigned int m{0}; m != family_members.size(); ++m)
        out << (m ? ',' : ' ') << n_anchor[m][anchor2];

    // coverage counts
    out << " " << min_parent_coverage_seen;
    for (const bool anchor2 : {false, true})
      for (unsigned int m{0}; m != family_members.size(); ++m)
        out << (m ? ',' : ' ') << n_coverage[m][anchor2];

    // get max other - the highest count in other families
    sort(parent_counts.begin(), parent_counts.end(),
         std::greater<unsigned int>());
    sort(children_counts.begin(), children_counts.end(),
         std::greater<unsigned int>());
    unsigned int max_other{parent_counts.size() ? parent_counts.front() : 0U};
    for (const unsigned int other : children_counts) {
      bool kid_count{false};
      for (vector<unsigned int>::iterator iter{kids_counts.begin()};
           iter != kids_counts.end(); ++iter) {
        if (*iter == other) {
          kids_counts.erase(iter);
          kid_count = true;
          break;
        }
      }
      if (!kid_count) {
        if (other > max_other) max_other = other;
        break;
      }
    }

    // population bridge counts
    out << " " << n_families
        << " " << n_samples
        << " " << parent_seen
        << " " << child_seen
        << " " << total_bridge_count
        << " " << parent_bridge_count
        << " " << child_bridge_count
        << " " << max_other;

    // parent counts
    if (parent_counts.size()) {
      for (unsigned int p{0}; p != parent_counts.size(); ++p)
        out << (p ? ',' : ' ') << parent_counts[p];
    } else {
      out << " 0";
    }
    // children counts
    if (children_counts.size()) {
      for (unsigned int c{0}; c != children_counts.size(); ++c)
        out << (c ? ',' : ' ') << children_counts[c];
    } else {
      out << " 0";
    }

    {  // This block necessary to prevent deadlock on return and later get()
      lock_guard<mutex> denovo_count_guard{denovo_count_mutex};

      // denovo or shared between siblings
      ++n_denovo;
      ++seen_denovo_families[family];
      for (const Sample sample : samples) ++seen_denovo_samples[sample];
      if (samples.size() == 1) {
        if (pop.is_proband(samples.front())) {
          ++n_denovo_proband;
        } else {
          ++n_denovo_sibling;
        }
      } else {
        ++n_denovo_siblings;
      }

      cout << out.str() << endl;
    }

    if (USE_FILES) family_lock.unlock();

    return;
  }

  void show_stats() {
    // summary info
    for (const auto s : pop.samples()) {
      if (pop.is_child(s)) {
        const Family f{pop.family(s)};
        serr << s
             << pop.sample(s) << seen_denovo_samples[s]
             << pop.family(f) << seen_denovo_families[f]
             << endl;
      }
    }

    serr << "cuts"
         << n_no_cut
         << n_not_snp
         << n_one_family
         << n_children
         << n_children_one_family
         << n_big_count
         << n_good_support
         << n_good_mate_support
         << n_good_excess
         << n_adjacent_cut
         << n_parent_coverage_cut
         << n_to_output
         << endl;

    serr << "denovos"
         << n_denovo << "all"
         << n_denovo_siblings << "both"
         << n_denovo_proband << "proband"
         << n_denovo_sibling << "sibling"
         << endl;
  }

 private:
  const Population & pop;
  const Reference & ref;
  const unsigned int chromosome;
  const unsigned int chromosome_offset;
  unsigned int chrY;
  vector<string> mumdex_names{};
  vector<Sample> seen_denovo_samples{};
  vector<Family> seen_denovo_families{};

  vector<mutex> family_locks{};
  mutex parent_bridge_count_mutex{};
  mutex to_output_count_mutex{};
  mutex denovo_count_mutex{};
  mutex adjacent_count_mutex{};
  mutex parent_count_mutex{};

  uint64_t n_denovo_proband{0};
  uint64_t n_denovo_sibling{0};
  uint64_t n_to_output{0};
  uint64_t n_denovo_siblings{0};
  uint64_t n_denovo{0};
};


void bridge_out(ostream & out, const BridgeInfo & bridge,
                const bool end_line = false) {
  out << " " << bridge.chr1()
      << " " << bridge.pos1()
      << " " << bridge.high1()
      << " " << bridge.chr2()
      << " " << bridge.pos2()
      << " " << bridge.high2()
      << " " << bridge.invariant()
      << " " << bridge.offset()
      << " " << bridge.bridge_count();
  if (end_line) out << endl;
}

int main(int argc, char* argv[])  try {
  if (--argc != 8)
    throw Error("usage: population_denovos ref family_file bridges_dir "
                "samples_dir n_threads chromosome start stop");

  // Other command line arguments
  const string reference_file{argv[1]};
  const Reference ref{reference_file};
  const ChromosomeIndexLookup chr_lookup{ref};
  const Mappability map{reference_file, true};
  const string family_file{argv[2]};
  const Population pop{family_file};
  const string bridges_dir{argv[3]};
  const string samples_dir{argv[4]};
  const unsigned int n_threads{static_cast<unsigned int>(atoi(argv[5]))};
  const string chromosome_name{argv[6]};
  const unsigned int chromosome{chr_lookup[chromosome_name]};
  const unsigned int chromosome_offset{ref.offset(chromosome)};
  const unsigned int start{static_cast<unsigned int>(atoi(argv[7]))};
  const unsigned int stop{static_cast<unsigned int>(atoi(argv[8]))};

  denovo_family_limit = static_cast<unsigned int>(
      std::max(1.0 * denovo_min_limit,
               denovo_family_fraction * pop.n_families()));

  // Parallel process slow candidate checks
  SlowChecks slow_checks(pop, ref, chromosome, samples_dir);

  ThreadPool pool{n_threads};
  ThreadPool::Results<void> results;

  // Set up bridge files to start position, in parallel to save time
  vector<future<BlockReader>> futures;
  futures.reserve(pop.n_samples());
  uint64_t block_size{4096};
  for (const auto s : pop.samples()) {
    ostringstream bridges_name;
    bridges_name << bridges_dir << "/" << pop.sample(s) << "/"
                 << get_bridge_file_name(ref, chromosome);
    if (!s) {
      block_size = std::max(block_size, get_block_size(bridges_name.str()));
      cerr << "Block size is " << block_size << endl;
    }
    futures.push_back(pool.run([start, stop](const string name){
          return BlockReader{name, start, stop};
        }, bridges_name.str()));
  }
  vector<BlockReader> readers;
  // readers.reserve(pop.n_samples());
  unsigned int n_samples_loaded{0};
  for (const auto s : pop.samples()) {
    if (s != n_samples_loaded) throw Error("Sample sanity check");
    readers.push_back(futures[n_samples_loaded].get());
    serr << pop.family(pop.family(s))
         << pop.sample(s)
         << pop.member(s)
         << ++n_samples_loaded
         << endl;
  }

  ThreadPool load_pool{n_threads};
  BlockMerger merger{load_pool, block_size = 32768, readers};

  // Document the run
  serr << "Run parameters:" << endl
       << "  n_threads" << n_threads << endl
       << "  family_file" << family_file << endl
       << "  samples_dir" << samples_dir << endl
       << "  bridges_dir" << bridges_dir << endl
       << "  chromosome" << chromosome_name << endl
       << "  start" << start << endl
       << "  stop" << stop << endl
       << "  n_families" << pop.n_families() << endl
       << "  n_samples" << pop.n_samples() << endl
       << "  denovo_family_limit" << denovo_family_limit << endl
       << "  exclude_snps" << exclude_snps << endl
       << "  min_support_length" << min_support_length << endl
       << "  min_bridge_count" << min_bridge_count << endl
       << "  min_mate_count" << min_mate_count << endl
       << "  min_mate_support" << min_mate_support << endl
       << "  min_parent_coverage" << min_parent_coverage << endl
       << "  min_parent_mum_length" << min_parent_mum_length << endl
       << "  min_excess_mappability" << min_excess_mappability << endl
       << "  adjacent_length" << adjacent_length << endl
       << "  adjacent_zero_length" << adjacent_zero_length << endl;

  sout << "family"
       << "samples"
       << "sex"
       << "kids"
       << "n_kids"
       << "kid_types"

       << "chr"
       << "chrA" << "posA" << "highA"
       << "chrB" << "posB" << "highB"
       << "invariant" << "offset"
       << "type" << "orient"

       << "supA" << "supB"
       << "emapA" << "emapB"
       << "mapA" << "mapB"
       << "mCA" << "mCB"
       << "mSA" << "mSB"

       << "bridge_count"
       << "bridges"
       << "anchorsA" << "anchorsB"
       << "min_parent_coverage"
       << "coveragesA" << "coveragesB"

       << "pop_n_families"
       << "pop_n_samples"
       << "pop_n_parents"
       << "pop_n_children"
       << "pop_total"
       << "pop_parent_total"
       << "pop_child_total"
       << "pop_max_other"
       << "pop_parent_counts"
       << "pop_child_counts"
       << endl;

  // Loop, adding one bridge at a time to all_bridges if the same event
  vector<BridgeInfo> all_bridges;
  vector<Sample> all_samples;
  vector<unsigned int> parent_counts;
  vector<unsigned int> children_counts;
  unsigned int total_bridge_count{0};
  unsigned int parent_bridge_count{0};
  unsigned int parent_seen{0};
  unsigned int child_bridge_count{0};
  unsigned int child_seen{0};
  uint64_t n_bridges{0};
  const uint64_t notify_interval{10000000};
  bool reset{true};
  BridgeInfo last_bridge{};
  while (true) {
    // New bridge expected - start fresh
    if (reset) {
      all_bridges.clear();
      all_samples.clear();
      parent_counts.clear();
      children_counts.clear();
      total_bridge_count = 0;
      parent_bridge_count = 0;
      parent_seen = 0;
      child_bridge_count = 0;
      child_seen = 0;
      reset = false;
    }

    // Add a bridge to the list if the same event
    bool bridge_done{true};
    if (merger.available()) {
      const BlockMerger::Item & item{merger.next()};
      const BridgeInfo current{item.first};
      // if (all_bridges.size() && current < all_bridges.back()) {
      if (current < last_bridge) {
        cerr << "Bridge misordering" << endl;
        bridge_out(cerr, last_bridge);
        cerr << endl;
        bridge_out(cerr, current);
        const Sample sample{item.second};
        cerr << " " << pop.sample(sample);
        cerr << endl;
        exit(1);
      }
      last_bridge = current;
      if (all_bridges.empty() || !(all_bridges.back() < current)) {
        ++n_no_cut;
        bridge_done = false;
        all_bridges.push_back(current);
        total_bridge_count += current.bridge_count();
        const Sample sample{item.second};
        all_samples.emplace_back(sample);
        if (pop.is_parent(sample)) {
          parent_bridge_count += current.bridge_count();
          ++parent_seen;
        } else {
          child_bridge_count += current.bridge_count();
          ++child_seen;
        }
        merger.advance();
      }
    } else {
      if (all_samples.empty()) {
        if (verbose) cerr << "bridge queue empty" << endl;
        break;
      }
    }

    // No new bridges added, so bridge list is ready to process
    if (bridge_done) {
      reset = true;

      // The first bridge, the exemplar since all are the same signature
      const BridgeInfo & bridge{all_bridges[0]};

      if ((++n_bridges % notify_interval) == 0)
        cerr << "progress: " << n_bridges << " " << bridge.pos1() << " "
             << 1.0 * (bridge.pos1() - start) / (stop - start) << endl;

      if (false) {
        bridge_out(cerr, bridge);
        for (unsigned int s{0}; s != all_samples.size(); ++s)
          cerr << " " << all_samples[s] << "." << pop.sample(all_samples[s]);
        cerr << endl;
      }

      // SNP cut
      const bool is_snp{bridge.chr1() == bridge.chr2() &&
            bridge.invariant() == 0 && bridge.high1() != bridge.high2()};
      if (is_snp) {
        if (exclude_snps) continue;
      } else {
        ++n_not_snp;
      }

      // Count families seen
      const unsigned int n_families{[&all_samples, &pop] {
          unsigned int n{0};
          Family last_family{100000};
          for (const Sample sample : all_samples) {
            const Family family{pop.family(sample)};
            if (family != last_family) {
              ++n;
              last_family = family;
            }
          }
          return n;
        }()};

      // Multiple family cut
      if (n_families == 1) {
        ++n_one_family;
      } else {
        if (n_families > denovo_family_limit) continue;
      }

      // Get mappability information
      const unsigned int mum1_abspos{chromosome_offset + bridge.pos1()};
      const unsigned int mum2_abspos{ref.offset(bridge.chr2()) + bridge.pos2()};
      const unsigned int mum1_map{map.low_high(bridge.high1(), mum1_abspos)};
      const unsigned int mum2_map{map.low_high(bridge.high2(), mum2_abspos)};

      for (unsigned int s{0}; s != all_samples.size(); ++s)
        if (pop.is_parent(all_samples[s])) {
          parent_counts.push_back(all_bridges[s].bridge_count());
        } else {
          children_counts.push_back(all_bridges[s].bridge_count());
        }
      // Look at each family separately
      const unsigned int n_samples{static_cast<unsigned int>(
          all_samples.size())};
      while (all_samples.size()) {
        // Gather samples and bridges for family
        static vector<Sample> samples;
        static vector<BridgeInfo> bridges;
        samples.clear();
        bridges.clear();
        while (all_samples.size() &&
               (samples.empty() ||
                pop.family(samples.back()) ==
                pop.family(all_samples.back()))) {
          samples.push_back(all_samples.back());
          all_samples.pop_back();
          bridges.push_back(all_bridges.back());
          all_bridges.pop_back();
        }

        // Parent sample cuts
        bool just_kids{true};
        unsigned int n_parents{0};
        for (const Sample sample : samples) {
          if (pop.is_parent(sample)) {
            just_kids = false;
            ++n_parents;
          }
        }
        if (just_kids) {
          ++n_children;
          if (n_families == 1) ++n_children_one_family;
        } else {
          continue;  // simple denovo cut
        }

        // Gather max support counts and lengths
        unsigned int max_bridge_count{0};
        unsigned int max_anchor1_support{0};
        unsigned int max_anchor2_support{0};
        unsigned int max_mate_anchor1_count{0};
        unsigned int max_mate_anchor1_support{0};
        unsigned int max_mate_anchor2_count{0};
        unsigned int max_mate_anchor2_support{0};
        for (const BridgeInfo & sample_bridge : bridges) {
          if (max_bridge_count < sample_bridge.bridge_count())
            max_bridge_count = sample_bridge.bridge_count();
          if (max_anchor1_support < sample_bridge.anchor1_length())
            max_anchor1_support = sample_bridge.anchor1_length();
          if (max_anchor2_support < sample_bridge.anchor2_length())
            max_anchor2_support = sample_bridge.anchor2_length();
          if (max_mate_anchor1_count < sample_bridge.mate_anchor1_count())
            max_mate_anchor1_count = sample_bridge.mate_anchor1_count();
          if (max_mate_anchor2_count < sample_bridge.mate_anchor2_count())
            max_mate_anchor2_count = sample_bridge.mate_anchor2_count();
          if (max_mate_anchor1_support < sample_bridge.mate_anchor1_length())
            max_mate_anchor1_support = sample_bridge.mate_anchor1_length();
          if (max_mate_anchor2_support < sample_bridge.mate_anchor2_length())
            max_mate_anchor2_support = sample_bridge.mate_anchor2_length();
        }

        // Low bridge count cut
        if (max_bridge_count < min_bridge_count) continue;
        ++n_big_count;

        // Short anchor support cut
        if (max_anchor1_support < min_support_length ||
            max_anchor2_support < min_support_length) continue;
        ++n_good_support;

        // Short mate support cut
        if (max_mate_anchor1_count < min_mate_count ||
            max_mate_anchor2_count < min_mate_count ||
            max_mate_anchor1_support < min_mate_support ||
            max_mate_anchor2_support < min_mate_support) continue;
        ++n_good_mate_support;

        // Short excess mappability cut
        if (max_anchor1_support < mum1_map) {
          cerr << "Mappability error" << endl;
          exit(1);
        }
        if (max_anchor1_support < min_excess_mappability + mum1_map)
          continue;
        if (max_anchor2_support < min_excess_mappability +  mum2_map)
          continue;
        ++n_good_excess;

        if (results.size() == n_threads) results.get();
        pool.run(results, std::ref(slow_checks),
                 bridges, samples, max_bridge_count,
                 n_families, n_samples, total_bridge_count,
                 parent_bridge_count, parent_seen,
                 child_bridge_count, child_seen,
                 max_anchor1_support, max_anchor2_support,
                 max_mate_anchor1_count, max_mate_anchor1_support,
                 max_mate_anchor2_count, max_mate_anchor2_support,
                 mum1_map, mum2_map, parent_counts, children_counts);
      }
    }
  }

  // Wait for all slow checks to finish
  while (results.size()) results.get();

  // Output cut stats
  slow_checks.show_stats();

  cerr << "finished bridge loop" << endl;

  return 0;
} catch (Error & e) {
  cerr << "paa::Error:" << endl;
  cerr << e.what() << endl;
  return 1;
} catch (exception & e) {
  cerr << "std::exception" << endl;
  cerr << e.what() << endl;
  return 1;
} catch (...) {
  cerr << "unknown exception was caught" << endl;
  return 1;
}
```
