## Supplementary material for "G-Graph: An interactive genomic graph viewer": MUMdex Source Code for G-Graph: qsufsort.h

```
//
// qsufsort.h
//
// suffix sorting
//
// Copyright Peter Andrews 2016 @ CSHL
// See bottom of this document for additional copyright notice
//

#ifndef LONGMEM_QSUFSORT_H_
#define LONGMEM_QSUFSORT_H_

#include <stdio.h>
#include <stdlib.h>

#include <algorithm>
#include <cstdint>
#include <limits>
#include <vector>

namespace paa {

/* Transforms the alphabet of x by attempting to aggregate several symbols into
   one, while preserving the suffix order of x. The alphabet may also be
   compacted, so that x on output comprises all integers of the new alphabet
   with no skipped numbers.

   Input: x is an array of size n+1 whose first n elements are positive
   integers in the range l...k-1. p is array of size n+1, used for temporary
   storage. q controls aggregation and compaction by defining the maximum value
   for any symbol during transformation: q must be at least k-l; if q<=n,
   compaction is guaranteed; if k-l>n, compaction is never done; if q is
   INT_MAX, the maximum number of symbols are aggregated into one.

   Output: Returns an integer j in the range 1...q representing the size of the
   new alphabet. If j<=n+1, the alphabet is compacted. The global variable r is
   set to the number of old symbols grouped into one. Only x[n] is 0.*/
template <class UINT>
uint64_t transform_alpha(UINT * const x, UINT * const p,
                         const uint64_t n, const uint64_t k, const uint64_t l,
                         const uint64_t q, uint64_t & qssr) {
  UINT *pi, *pj;
  uint64_t b, c, d, e, i, j, m, s;

  for (s = 0, i = k - l; i; i >>= 1)
    ++s;                           /* s is number of bits in old symbol.*/
  e = std::numeric_limits<UINT>::max() >> (s + 1);  /* e is to check overflow.*/
  for (b = d = qssr = 0; qssr < n && d <= e &&
           (c = d << s | (k - l)) <= q; ++qssr) {
    b = b << s | (x[qssr] - l + 1);  /* b is start of x in chunk alphabet.*/
    d = c;                         /* d is max symbol in chunk alphabet.*/
  }
  m = (1 << (qssr - 1) * s) - 1;   /* m masks off top old symbol from chunk.*/
  x[n] = static_cast<UINT>(l - 1);    /* emulate zero terminator.*/
  if (d <= n) {                    /* if bucketing possible, compact alphabet.*/
    for (pi = p; pi <= p + d; ++pi)
      *pi = 0;                     /* zero transformation table.*/
    for (pi = x + qssr, c = b; pi <= x + n; ++pi) {
      p[c] = 1;                    /* mark used chunk symbol.*/
      c = (c & m) << s | (*pi - l + 1);  /* shift in next old symbol in chunk.*/
    }
    for (i = 1; i < qssr; ++i) {   /* handle last r-1 positions.*/
      p[c] = 1;                    /* mark used chunk symbol.*/
      c = (c & m) << s;            /* shift in next old symbol in chunk.*/
    }
    for (pi = p, j = 1; pi <= p + d; ++pi)
      if (*pi)
        *pi = static_cast<UINT>(j++);            /* j is new alphabet size.*/
    for (pi = x, pj = x + qssr, c = b; pj <= x + n; ++pi, ++pj) {
      *pi = p[c];                  /* transform to new alphabet.*/
      c = (c & m) << s | (*pj - l + 1);  /* shift in next old symbol in chunk.*/
    }
    while (pi < x + n) {           /* handle last r-1 positions.*/
      *pi++ = p[c];                /* transform to new alphabet.*/
      c = (c & m) << s;            /* shift right-end zero in chunk.*/
    }
  } else {                         /* bucketing not possible, don't compact.*/
    for (pi = x, pj = x + qssr, c = b; pj <= x + n; ++pi, ++pj) {
      *pi = static_cast<UINT>(c);       /* transform to new alphabet.*/
      c = (c & m) << s | (*pj - l + 1);  /* shift in next old symbol in chunk.*/
    }
    while (pi < x + n) {           /* handle last r-1 positions.*/
      *pi++ = static_cast<UINT>(c);     /* transform to new alphabet.*/
      c = (c & m) << s;            /* shift right-end zero in chunk.*/
    }
    j = d + 1;                     /* new alphabet size.*/
  }
  x[n] = 0;                        /* end-of-string symbol is zero.*/
  return j;                        /* return new alphabet size.*/
}

// Use signed int, faster algorithms for long integer type
// Use same unsigned int for all other int sizes (typically only uint32_t)
template <class UINT> struct SortInt { using Type = UINT; };
template <> struct SortInt<uint64_t> { using Type = int64_t; };

// Only can be used with uint32_t and uint64_t
template <class UINT>
class suffixsort {
  using SINT = typename SortInt<UINT>::Type;
  using Bool = char;

 public:
  /* Makes suffix array p of x. x becomes inverse of p. p and x are both of size
     n+1. Contents of x[0...n-1] are integers in the range l...k-1. Original
     contents of x[n] is disregarded, the n-th symbol being regarded as
     end-of-string smaller than all other symbols. */
  suffixsort(uint32_t * const x, uint32_t * const p,
             const uint64_t n, const uint64_t k, const uint64_t l,
             const bool verbose_ = true) : V{x}, I{p}, verbose{verbose_} {
    UINT *pi, *pk;
    uint64_t i, j;
    UINT s, sl;

    // Is position a sorted group?
    std::vector<Bool> is_sg(n + 1);

    if (verbose) fprintf(stderr, "alphabet transform\n");
    uint64_t r = 0;
    if (n >= k - l) {                /* if bucketing possible,*/
      j = transform_alpha(x, p, n, k, l, n, r);
      if (verbose) fprintf(stderr, "bucket sort\n");
      bucketsort(is_sg, x, p, n, j);   /* bucketsort on first r positions.*/
    } else {
      // Will never encounter this branch - alphabet bigger than input
      transform_alpha(x, p, n, k, l, std::numeric_limits<UINT>::max() >> 1, r);
      if (verbose) fprintf(stderr, "initialize I\n");
      /* initialize I with suffix numbers. */
      for (i = 0; i <= n; ++i)
        p[i] = static_cast<UINT>(i);
      h = 0;
      if (verbose) fprintf(stderr, "sort split\n");
      sort_split(is_sg, p, n + 1);       /* quicksort on first r positions.*/
    }
    h = r;                  /* number of symbols aggregated by transform.*/

    if (verbose) fprintf(stderr, "main sort\n");
    while (!is_sg[0] || *p <= n) {
      pi = p;                     /* pi is first position of group.*/
      sl = 0;                     /* sl is length of sorted groups.*/
      do {
        s = *pi;   // Added
        if (is_sg[pi - p]) {
          // pi -= s;              /* skip over sorted group.*/
          pi += s;              /* skip over sorted group.*/
          sl += s;              /* add length to sl.*/
        } else {
          if (sl) {
            *(pi - sl) = sl;     /* combine sorted groups before pi.*/
            is_sg[pi - sl - p] = true;  // Added
            sl = 0;
          }
          pk = p + x[s] + 1;   /* pk-1 is last position of unsorted group.*/
          sort_split(is_sg, pi, pk - pi);
          pi = pk;              /* next group.*/
        }
      } while (pi <= p + n);
      if (sl) {                  /* if the array ends with a sorted group.*/
        *(pi - sl) = sl;           /* combine sorted groups at end of I.*/
        is_sg[pi - sl - p] = true;  // Added
      }
      h *= 2;                    /* double sorted-depth.*/
    }

    if (verbose) fprintf(stderr, "reconstruct SA from ISA\n");
    for (i = 0; i <= n; ++i) {
      p[x[i]] = static_cast<UINT>(i);
    }
  }

  suffixsort(uint64_t * const x, uint64_t * const p,
             const uint64_t n, const uint64_t k, const uint64_t l,
             const bool verbose_ = true) :
      V{reinterpret_cast<int64_t *>(x)}, I{reinterpret_cast<int64_t *>(p)},
    verbose{verbose_} {
    int64_t *pi, *pk;
    uint64_t i, j;
    int64_t s, sl;

    if (verbose) fprintf(stderr, "alphabet transform\n");
    uint64_t r = 0;
    if (n >= k-l) {                /* if bucketing possible,*/
      j = transform_alpha(x, p, n, k, l, n, r);
      if (verbose) fprintf(stderr, "bucket sort\n");
      bucketsort(V, I, n, j);   /* bucketsort on first r positions.*/
    } else {
      transform_alpha(x, p, n, k, l, std::numeric_limits<int64_t>::max(), r);
      if (verbose) fprintf(stderr, "initialize I\n");
      for (i = 0; i <= n; ++i)
        I[i] = i;                /* initialize I with suffix numbers. */
      h = 0;
      if (verbose) fprintf(stderr, "sort split\n");
      sort_split(I, n + 1);       /* quicksort on first r positions.*/
    }
    h = r;                  /* number of symbols aggregated by transform.*/

    if (verbose) fprintf(stderr, "main sort\n");
    while (*I >= -static_cast<int64_t>(n)) {
      pi = I;                     /* pi is first position of group.*/
      sl = 0;                     /* sl is negated length of sorted groups.*/
      do {
        if ((s = *pi) < 0) {
          pi -= s;              /* skip over sorted group.*/
          sl += s;              /* add negated length to sl.*/
        } else {
          if (sl) {
            *(pi + sl) = sl;     /* combine sorted groups before pi.*/
            sl = 0;
          }
          pk = I + V[s]+1;   /* pk-1 is last position of unsorted group.*/
          sort_split(pi, pk - pi);
          pi = pk;              /* next group.*/
        }
      } while (pi <= I + n);
      if (sl)                   /* if the array ends with a sorted group.*/
        *(pi + sl) = sl;           /* combine sorted groups at end of I.*/
      h *= 2;                    /* double sorted-depth.*/
    }

    if (verbose) fprintf(stderr, "reconstruct SA from ISA\n");
    for (i = 0; i <= n; ++i) {
      p[V[i]] = i;
    }
  }

 private:
  void update_group(std::vector<Bool> & is_sg,
                    UINT * pl, UINT * const pm) {
    const uint64_t g = pm - I;  /* group number.*/
    V[*pl] = static_cast<SINT>(g);  /* update group number of first position.*/
    if (pl == pm) {
      *pl = 1;                    /* one element, sorted group.*/
      is_sg[pl - I] = true;
    } else {
      do {                         /* more than one element, unsorted group.*/
        V[*++pl] = static_cast<SINT>(g);           /* update group numbers.*/
      } while (pl < pm);
    }
  }
  void update_group(SINT * pl, SINT * const pm) {
    const uint64_t g = pm - I;  /* group number.*/
    V[*pl] = g;                 /* update group number of first position.*/
    if (pl == pm) {
      *pl = -1;                    /* one element, sorted group.*/
    } else {
      do {                         /* more than one element, unsorted group.*/
        V[*++pl] = g;           /* update group numbers.*/
      } while (pl < pm);
    }
  }

  /* Subroutine for select_sort_split and sort_split. Sets group numbers for a
     group whose lowest position in I is pl and highest position is pm.*/
  /* Quadratic sorting method to use for small subarrays. To be able to update
     group numbers consistently, a variant of selection sorting is used.*/
  void select_sort_split(std::vector<Bool> & is_sg,
                         UINT * const p, const uint64_t n) {
    UINT *pa, *pb, *pi, *pn;
    uint64_t f, v;

    pa = p;                        /* pa is start of group being picked out.*/
    pn = p + n - 1;                /* pn is last position of subarray.*/
    while (pa < pn) {
      for (pi = pb = pa + 1, f = KEY(pa); pi <= pn; ++pi)
        if ((v = KEY(pi)) < f) {
          f = v;                 /* f is smallest key found.*/
          SWAP(pi, pa);          /* place smallest element at beginning.*/
          pb = pa + 1;           /* pb is position for elements equal to f.*/
        } else if (v == f) {     /* if equal to smallest key.*/
          SWAP(pi, pb);          /* place next to other smallest elements.*/
          ++pb;
        }
      update_group(is_sg, pa, pb - 1);  /* update group values for new group.*/
      pa = pb;                   /* continue sorting rest of the subarray.*/
    }
    if (pa == pn) {              /* check if last part is single element.*/
      V[*pa] = static_cast<SINT>(pa - I);
      *pa = 1;                  /* sorted group.*/
      is_sg[pa - I] = true;
    }
  }
  void select_sort_split(SINT * const p, const uint64_t n) {
    SINT *pa, *pb, *pi, *pn;
    uint64_t f, v;

    pa = p;                        /* pa is start of group being picked out.*/
    pn = p + n - 1;                /* pn is last position of subarray.*/
    while (pa < pn) {
      for (pi = pb = pa + 1, f = KEY(pa); pi <= pn; ++pi)
        if ((v = KEY(pi)) < f) {
          f = v;                 /* f is smallest key found.*/
          SWAP(pi, pa);          /* place smallest element at beginning.*/
          pb = pa + 1;           /* pb is position for elements equal to f.*/
        } else if (v == f) {     /* if equal to smallest key.*/
          SWAP(pi, pb);          /* place next to other smallest elements.*/
          ++pb;
        }
      update_group(pa, pb - 1);  /* update group values for new group.*/
      pa = pb;                   /* continue sorting rest of the subarray.*/
    }
    if (pa == pn) {              /* check if last part is single element.*/
      V[*pa] = pa - I;
      *pa = -1;                  /* sorted group.*/
    }
  }

  /* Subroutine for sort_split, algorithm by Bentley & McIlroy.*/
  uint64_t choose_pivot(SINT * const p, const uint64_t n) {
    SINT *pl, *pm, *pn;

    pm = p + (n>>1);              /* small arrays, middle element.*/
    if (n > 7) {
      pl = p;
      pn = p + n - 1;
      if (n > 40) {               /* big arrays, pseudomedian of 9.*/
        const uint64_t s = n >> 3;
        pl = MED3(pl, pl + s, pl + s + s);
        pm = MED3(pm - s, pm, pm + s);
        pn = MED3(pn - s - s, pn - s, pn);
      }
      pm = MED3(pl, pm, pn);      /* midsize arrays, median of 3.*/
    }
    return KEY(pm);
  }

  /* Sorting routine called for each unsorted group. Sorts the array of integers
     (suffix numbers) of length n starting at p.
     The algorithm is a ternary-split
     quicksort taken from Bentley & McIlroy, "Engineering a Sort Function",
     Software -- Practice and Experience 23(11), 1249-1265 (November 1993). This
     function is based on Program 7.*/
  void sort_split(std::vector<Bool> & is_sg, UINT * const p, const uint64_t n) {
    UINT *pa, *pb, *pc, *pd, *pl, *pm, *pn;
    UINT s, t;
    uint64_t f, v;

    if (n < 7) {                 /* multi-selection sort smallest arrays.*/
      select_sort_split(is_sg, p, n);
      return;
    }

    v = choose_pivot(p, n);
    pa = pb = p;
    pc = pd = p + n - 1;
    while (1) {                  /* split-end partition.*/
      while (pb <= pc && (f = KEY(pb)) <= v) {
        if (f == v) {
          SWAP(pa, pb);
          ++pa;
        }
        ++pb;
      }
      while (pc >= pb && (f = KEY(pc)) >= v) {
        if (f == v) {
          SWAP(pc, pd);
          --pd;
        }
        --pc;
      }
      if (pb > pc)
        break;
      SWAP(pb, pc);
      ++pb;
      --pc;
    }
    pn = p+n;
    if ((s = static_cast<UINT>(pa - p)) > (t = static_cast<UINT>(pb - pa)))
      s = t;
    for (pl = p, pm = pb - s; s; --s, ++pl, ++pm)
      SWAP(pl, pm);
    if ((s = static_cast<UINT>(pd - pc)) > (t = static_cast<UINT>(pn - pd - 1)))
      s = t;
    for (pl = pb, pm = pn - s; s; --s, ++pl, ++pm)
      SWAP(pl, pm);

    s = static_cast<UINT>(pb - pa);
    t = static_cast<UINT>(pd - pc);
    if (s > 0)
      sort_split(is_sg, p, s);
    update_group(is_sg, p + s, p + n - t - 1);
    if (t > 0)
      sort_split(is_sg, p + n - t, t);
  }
  void sort_split(SINT * const p, const uint64_t n) {
    SINT *pa, *pb, *pc, *pd, *pl, *pm, *pn;
    SINT s, t;
    uint64_t f, v;

    if (n < 7) {                 /* multi-selection sort smallest arrays.*/
      select_sort_split(p, n);
      return;
    }

    v = choose_pivot(p, n);
    pa = pb = p;
    pc = pd = p + n - 1;
    while (1) {                  /* split-end partition.*/
      while (pb <= pc && (f = KEY(pb)) <= v) {
        if (f == v) {
          SWAP(pa, pb);
          ++pa;
        }
        ++pb;
      }
      while (pc >= pb && (f = KEY(pc)) >= v) {
        if (f == v) {
          SWAP(pc, pd);
          --pd;
        }
        --pc;
      }
      if (pb > pc)
        break;
      SWAP(pb, pc);
      ++pb;
      --pc;
    }
    pn = p+n;
    if ((s = pa - p) > (t = pb - pa))
      s = t;
    for (pl = p, pm = pb - s; s; --s, ++pl, ++pm)
      SWAP(pl, pm);
    if ((s = pd - pc) > (t = pn - pd - 1))
      s = t;
    for (pl = pb, pm = pn - s; s; --s, ++pl, ++pm)
      SWAP(pl, pm);

    s = pb - pa;
    t = pd - pc;
    if (s > 0)
      sort_split(p, s);
    update_group(p + s, p + n - t - 1);
    if (t > 0)
      sort_split(p + n - t, t);
  }

  // Bucketsort for first iteration.
  // Input: x[0...n-1] holds integers in the range 1...k-1, all of which appear
  // at least once. x[n] is 0. (This is the corresponding output of transform.)
  // k must be at most n + 1. p is array of size n+1 whose contents
  // are disregarded.

  // Output: x is V and p is I after the initial sorting stage of the refined
  // suffix sorting algorithm.
  void bucketsort(std::vector<Bool> & is_sg, UINT * const x, UINT * const p,
                  const uint64_t n, const uint64_t k) {
    UINT *pi, d;
    uint64_t c, i, g;

    for (pi = p; pi < p + k; ++pi)
      *pi = static_cast<UINT>(-1);  /* mark linked lists empty.*/
    for (i = 0; i <= n; ++i) {
      x[i] = p[c = x[i]];           /* insert in linked list.*/
      p[c] = static_cast<UINT>(i);
    }
    for (pi = p + k - 1, i = n; pi >= p; --pi) {
      d = x[c = *pi];               /* c is position, d is next in list.*/
      x[c] = static_cast<UINT>(g = i);  /* last position equals group number.*/
      if (d != static_cast<UINT>(-1)) {  /* if more than one element in group.*/
        p[i--] = static_cast<UINT>(c);  /* p is permutation for the sorted x.*/
        do {
          d = x[c = d];             /* next in linked list.*/
          x[c] = static_cast<UINT>(g);                 /* group number in x.*/
          p[i--] = static_cast<UINT>(c);               /* permutation in p.*/
        } while (d != static_cast<UINT>(-1));
      } else {
        is_sg[i] = true;
        p[i--] = 1;                /* one element, sorted group.*/
      }
    }
  }
  void bucketsort(SINT * const x, SINT * const p,
                  const uint64_t n, const uint64_t k) {
    SINT *pi, d;
    uint64_t c, i, g;

    for (pi = p; pi < p + k; ++pi)
      *pi = -1;                     /* mark linked lists empty.*/
    for (i = 0; i <= n; ++i) {
      x[i] = p[c = x[i]];           /* insert in linked list.*/
      p[c] = i;
    }
    for (pi = p + k - 1, i = n; pi >= p; --pi) {
      d = x[c = *pi];               /* c is position, d is next in list.*/
      x[c] = g = i;                 /* last position equals group number.*/
      if (d >= 0) {                 /* if more than one element in group.*/
        p[i--] = c;                 /* p is permutation for the sorted x.*/
        do {
          d = x[c = d];             /* next in linked list.*/
          x[c] = g;                 /* group number in x.*/
          p[i--] = c;               /* permutation in p.*/
        } while (d >= 0);
      } else {
        p[i--] = -1;                /* one element, sorted group.*/
      }
    }
  }

  SINT KEY(const SINT * const p) const { return (V[*(p) + h]); }
  static void SWAP(SINT * const p, SINT * const q) { std::swap(*(p), *(q)); }
  SINT * MED3(SINT * const a, SINT * const b, SINT * const c) const {
    return (KEY(a) < KEY(b) ? (KEY(b) < KEY(c) ? (b) :
                               KEY(a) < KEY(c) ? (c) : (a)) :
            (KEY(b) > KEY(c) ? (b) : KEY(a) > KEY(c) ? (c) : (a)));
  }

  SINT * V;         /* inverse array, ultimately inverse of I.*/
  SINT * I;         /* group array, ultimately suffix array.*/
  uint64_t h{0};       /* length of already-sorted prefixes.*/
  bool verbose;
};

}  // namespace paa

#endif  // LONGMEM_QSUFSORT_H_

/* qsufsort.c
   Copyright 1999, N. Jesper Larsson, all rights reserved.

   This file contains an implementation of the algorithm presented in "Faster
   Suffix Sorting" by N. Jesper Larsson and Kunihiko
   Sadakane.

   This software may be used freely for any purpose. However, when distributed,
   the original source must be clearly stated, and, when the source code is
   distributed, the copyright notice must be retained and any alterations in
   the code must be clearly marked. No warranty is given regarding the quality
   of this software.*/
```
