## Supplementary material for "G-Graph: An interactive genomic graph viewer": MUMdex Source Code for G-Graph: query.h

```
//
// query.h
//
// mapping queries for mummer
//
// Copyright Peter Andrews 2015-2016 @ CSHL
//

#ifndef LONGMEM_QUERY_H_
#define LONGMEM_QUERY_H_

#include <pthread.h>

#include <algorithm>
#include <fstream>
#include <string>
#include <vector>

#include "encode.h"
#include "error.h"
#include "longSA.h"
#include "mumdex.h"
#include "sam.h"

namespace paa {

struct Query {
  std::string query{};
  std::string optional{};
  bool bad{};
};

class AlignerArgs {
 public:
  AlignerArgs() : min_len(2) {}
  unsigned int min_len;

 private:
  AlignerArgs & operator=(const AlignerArgs &) = delete;
};

template<class SA>
class Aligner : public Query, public AlignerArgs {
 public:
  // Used by PairRunner class
  Aligner(const AlignerArgs & args, const SA & sa_) :
      Query(), AlignerArgs(args), sa(sa_), matches(0) {}
  void clear() { matches.clear(); }
  void run() { sa.find_mams(matches, query, min_len); }

  const SA & sa;
  std::vector<match_t> matches;

 private:
  Aligner & operator=(const Aligner &) = delete;
};

class ReaderArgs {
 public:
  ReaderArgs() : fastq(false), fastqs(false),
                 sam_in(false), verbose(false) {}
  bool fastq;
  bool fastqs;
  bool sam_in;
  bool verbose;

 private:
  ReaderArgs & operator=(const ReaderArgs &) = delete;
};

class QueryReader {
 public:
  QueryReader() { pthread_mutex_init(&mutex, nullptr); }
  ~QueryReader() { pthread_mutex_destroy(&mutex); }
  template <class Pair>
  bool read_pair(Pair & pair, const bool sam_in) {
    constexpr bool check_names{false};
    pthread_mutex_lock(&mutex);
    if (sam_in) {
      getline(data1, pair.line1);
      getline(data1, pair.line2);
    } else {
      static std::string name1;
      static std::string name2;
      // read 1
      if (check_names) {
        getline(data1, name1);
      } else {
        data1.ignore(10000, '\n');
      }
      getline(data1, pair.line1);
      data1.ignore(10000, '\n');
      data1.ignore(10000, '\n');
      // read 2
      if (check_names) {
        getline(*data2_p, name2);
      } else {
        data2_p->ignore(10000, '\n');
      }
      getline(*data2_p, pair.line2);
      data2_p->ignore(10000, '\n');
      data2_p->ignore(10000, '\n');
      if (check_names) {
        // Make sure names for read 1 and read 2 match
        const size_t pos1{name1.find_first_of("# ")};
        if (pos1 != std::string::npos) {
          name1.erase(pos1);
        }
        const size_t pos2{name2.find_first_of("# ")};
        if (pos2 != std::string::npos) {
          name2.erase(pos2);
        }
        if (data1 && name1 != name2) {
          throw Error("Paired sequence names do not match")
              << name1 << name2;
        }
      }
    }
    pthread_mutex_unlock(&mutex);
    if (data1) return true;
    return false;
  }
  void open(const char * query_input) {
    data1.open(query_input);
    if (!data1) throw Error("unable to open") << query_input;
    data2_p = &data1;
  }
  void open(const char * query_input, const char * query_input2) {
    data1.open(query_input);
    if (!data1) throw Error("unable to open") << query_input;
    data2.open(query_input2);
    if (!data2) throw Error("unable to open") << query_input2;
    data2_p = &data2;
  }

 private:
  pthread_mutex_t mutex{};
  std::ifstream data1{};
  std::ifstream data2{};
  std::ifstream * data2_p{};
  QueryReader(const QueryReader &) = delete;
  QueryReader & operator=(const QueryReader &) = delete;
};

class ReadersArgs : public ReaderArgs {
 public:
  ReadersArgs() : n_input(0), input(nullptr) {}
  unsigned int n_input;
  char * * input;

 private:
  ReadersArgs & operator=(const ReadersArgs &) = delete;
};

class Readers : public ReadersArgs {
 public:
  explicit Readers(const ReadersArgs & args)
      : ReadersArgs(args),
        readers(fastqs ? args.n_input / 2 : args.n_input),
        reader_bad(readers.size()) {
    for (unsigned int r = 0; r != readers.size(); ++r) {
      if (fastqs) {
        readers[r].open(args.input[2 * r], args.input[2 * r + 1]);
      } else {
        readers[r].open(args.input[r]);
      }
    }
    if (verbose) std::cerr << "using " << readers.size() << " query reader"
                           << (readers.size() > 1 ? "s" : "") << std::endl;
    pthread_mutex_init(&mutex, nullptr);
  }
  ~Readers() { pthread_mutex_destroy(&mutex); }
  template <class Pair>
  bool read_pair(Pair & pair) {
    while (current + 1 != readers.size() || !reader_bad[current]) {
      if (readers[current].read_pair(pair, sam_in)) {
        return true;
      } else {
        pthread_mutex_lock(&mutex);
        if (!reader_bad[current]) {
          reader_bad[current] = true;
          if (current + 1 != readers.size()) {
            ++current;
          }
        }
        pthread_mutex_unlock(&mutex);
      }
    }
    return false;
  }

 private:
  std::vector<QueryReader> readers;
  std::vector<bool> reader_bad;
  unsigned int current{0};
  pthread_mutex_t mutex{};
  Readers(const Readers &) = delete;
  Readers & operator=(const Readers &) = delete;
};

class ReadPairArgs : public AlignerArgs {
 public:
  ReadPairArgs() : AlignerArgs(), max_mem{0.0}, pass_optional{false} {}
  double max_mem;
  bool pass_optional;
  std::vector<std::string> optional{};

 private:
  ReadPairArgs & operator=(const ReadPairArgs &) = delete;
};

template<class SA>
class ReadPair : public ReadPairArgs {
 public:
  ReadPair(const ReadPairArgs & args, const SA & sa_, Readers & readers_) :
      ReadPairArgs(args),
      n_queries(0), read1(args, sa_), read2(args, sa_), readers(readers_) {}

  static void * runner_thread(void * obj) {
    try {
      reinterpret_cast<ReadPair *>(obj)->run();
    }
    catch(std::exception & e) {
      std::cerr << e.what() << std::endl;
      exit(1);
    }
    catch(...) {
      std::cerr << "Some exception was caught." << std::endl;
      exit(1);
    }
    pthread_exit(nullptr);
  }

  uint64_t n_queries;
  std::string line1{};
  std::string line2{};

 private:
  void run() {
    std::ostringstream out_name;
    out_name << "mumdex/part." << reinterpret_cast<uint64_t>(this);
    const unsigned int max_pairs = (read1.sa.ref.rcref ? 10 : 1) * 200000;
    const unsigned int max_mums = max_pairs * 16;
    MUMdex_build<Sequence> mumdex{read1.sa.ref, out_name.str(),
          max_pairs, max_mums + 100};
    MUMdex_build_base sorted{max_pairs, max_mums + 100};
    OptionalSavers optional_savers{optional};
    OptionalSavers sorting_savers{optional};

    std::string name, ref, pos, mapq, cigar, mref, mpos, tlen, seq,
        errors_and_optional;
    unsigned int flag;

    std::istringstream input;
    while (readers.read_pair(*this)) {
      for (const bool second : {false, true}) {
        Aligner<SA> & read = second ? read2 : read1;
        std::string & line = second ? line2 : line1;

        if (readers.sam_in) {
          input.clear();
          input.str(line);
          input >> name >> flag >> ref >> pos >> mapq >> cigar
                >> mref >> mpos >> tlen >> read.query;
          if (!input)
            throw Error("Sam format parse error for line") << line;
          if (pass_optional) {
            input.get();
            getline(input, read.optional);
          }
          if (static_cast<bool>(flag & sam::is_second) != second) {
            throw Error("Unpaired read")
                << static_cast<bool>(flag & sam::is_second) + 1;
          }
          read.bad = flag & sam::is_bad_vendor_quality;
        } else {
          // read.query = move(line);
          using std::swap;
          swap(read.query, line);
          read.bad = false;
        }
        ++n_queries;
        read.run();

        if (second) {
          mumdex.emplace_back(read1.matches, read1.query, read1.bad,
                              read2.matches, read2.query, read2.bad);
          optional_savers.extract(read1.optional, read2.optional);
          read1.clear();
          read2.clear();
          if (mumdex.n_pairs() == max_pairs ||
              mumdex.n_mums() >= max_mums) {
            const std::string part_name = mumdex.sort_and_save_part(sorted);
            optional_savers.save_and_clear(part_name, sorting_savers,
                                           mumdex.index());
            mumdex.clear();
          }
        }
      }
    }
    if (mumdex.n_pairs()) {
      const std::string part_name = mumdex.sort_and_save_part(sorted);
      optional_savers.save_and_clear(part_name, sorting_savers, mumdex.index());
      mumdex.clear();
    }
    if ((n_queries % 2) == 1) {
      throw Error("unpaired read found at end");
    }
  }

  Aligner<SA> read1;
  Aligner<SA> read2;
  Readers & readers;

  ReadPair & operator=(const ReadPair &) = delete;
};

class ReadPairsArgs : public ReadPairArgs {
 public:
  ReadPairsArgs() : ReadPairArgs(), max_n_threads(1),
                    n_threads(1), verbose(false) {}
  unsigned int max_n_threads;
  unsigned int n_threads;
  bool verbose;

 private:
  ReadPairsArgs & operator=(const ReadPairsArgs &) = delete;
};

template<class SA>
class ReadPairs : public ReadPairsArgs {
 public:
  ReadPairs(const ReadPairsArgs & args, const SA & sa, Readers & readers)
      : ReadPairsArgs(args), thread_ids(n_threads),
        pairs(n_threads, ReadPair<SA>(args, sa, readers)),
        start_time(time(nullptr)) {
    if (verbose)
      std::cerr << "running " << n_threads << " thread"
           << (n_threads > 1 ? "s" : "") << " to answer queries" << std::endl;

    // Create mumdex directory
    if (readable("mumdex"))
      throw Error("mumdex directory already exists.  Quitting");
    mkdir("mumdex");
    if (optional.size()) {
      optional_format_saver(optional, "mumdex");
    }

    // Initialize pairs and start threads
    pthread_attr_t attr;
    pthread_attr_init(&attr);
    pthread_attr_setdetachstate(&attr, PTHREAD_CREATE_JOINABLE);
    for (unsigned int thread = 0; thread != n_threads; ++thread) {
      if (pthread_create(&thread_ids[thread], &attr,
                         &ReadPair<SA>::runner_thread, &pairs[thread]))
        throw Error("Problem creating runner_thread") << thread;
    }
  }

  ~ReadPairs() {
    uint64_t n_processed = 0;
    for (unsigned int t = 0; t != n_threads; ++t) {
      if (pthread_join(thread_ids[t], nullptr)) {
        std::cerr << "Problem joining ReadPairs thread " << t << std::endl;
      }
      n_processed += pairs[t].n_queries;
    }
    const time_t elapsed_time = time(nullptr) - start_time;
    if (verbose) std::cerr << "ran " << n_processed << " queries in "
                      << time(nullptr) - start_time << " seconds";
    if (verbose && elapsed_time) {
      std::cerr << " for a rate of "
           << 60E-6 * n_processed / elapsed_time
           << " million queries per minute" << std::endl;
    } else if (verbose) {
      std::cerr << std::endl;
    }
  }

 private:
  std::vector<pthread_t> thread_ids;
  std::vector<ReadPair<SA>> pairs;
  const time_t start_time;
  ReadPairs(const ReadPairs &) = delete;
  ReadPairs & operator=(const ReadPairs &) = delete;
};
using shortReadPairs = ReadPairs<shortSA>;
using longReadPairs = ReadPairs<longSA>;

}  // namespace paa

#endif  // LONGMEM_QUERY_H_
```
