## Supplementary material for "G-Graph: An interactive genomic graph viewer": MUMdex Source Code for G-Graph: sam.h

```
//
// sam.h
//
// sam info
//
// Copyright 2016 Peter Andrews @ CSHL
//

#ifndef PAA_SAM_H
#define PAA_SAM_H

namespace sam {

enum MapFlag {
  is_paired = 1 << 0,
  is_proper = 1 << 1,
  is_unmapped = 1 << 2,
  is_mate_unmapped = 1 << 3,
  is_reversed = 1 << 4,
  is_mate_reversed = 1 << 5,
  is_first = 1 << 6,
  is_second = 1 << 7,
  is_not_primary = 1 << 8,
  is_bad_vendor_quality = 1 << 9,
  is_a_duplicate = 1 << 10
};

}  // namespace sam


#endif  // PAA_SAM_H
```
