## Supplementary material for "G-Graph: An interactive genomic graph viewer": MUMdex Source Code for G-Graph: sequence.h

```
//
// sequence.h
//
// various sequence classes and functions
//
// Copyright 2016 Peter Andrews @ CSHL
//

#ifndef PAA_SEQUENCE_H
#define PAA_SEQUENCE_H

#include <algorithm>
#include <array>
#include <sstream>
#include <string>
#include <utility>
#include <vector>

#include "error.h"
#include "mumdex.h"
#include "pstream.h"

namespace paa {

class Similarity {
 public:
  Similarity(const std::string & seq1, const std::string & seq2) {
    if (seq1.size() != seq2.size())
      throw Error("Bad sequence size in Similarity");
    for (unsigned int b{0}; b != seq1.size(); ++b) {
      if (seq1[b] != 'x' && seq1[b] != 'N' &&
          seq1[b] != 'x' && seq2[b] != 'N') {
        if (seq1[b] == seq2[b]) {
          ++n_matches_;
        } else {
          ++n_mismatches_;
        }
      }
    }
  }

  unsigned int n_matches() const { return n_matches_; }
  unsigned int n_testable() const { return n_matches_ + n_mismatches_; }
  unsigned int n_mismatches() const { return n_mismatches_; }

 private:
  unsigned int n_matches_{0};
  unsigned int n_mismatches_{0};
};

class ConsensusSequence {
 public:
  class BaseId {
   public:
    explicit BaseId(const char base) {
      switch (base) {
        case 'A':
          index = 0;
          break;
        case 'C':
          index = 1;
          break;
        case 'G':
          index = 2;
          break;
        case 'T':
          index = 3;
          break;
        case 'N':
          index = 4;
          break;
        case 'x':
          index = 5;
          break;
        default:
          throw Error("Unexpected base seen") << base;
      }
    }
    operator unsigned int() const {
      return index;
    }
    unsigned int value() const {
      return index;
    }

   private:
    unsigned int index{};
  };

  class Base {
   public:
    explicit Base(const unsigned int index) {
      switch (index) {
        case 0:
          base = 'A';
          break;
        case 1:
          base = 'C';
          break;
        case 2:
          base = 'G';
          break;
        case 3:
          base = 'T';
          break;
        case 4:
          base = 'N';
          break;
        case 5:
          base = 'x';
          break;
        default:
          throw Error("Unexpected index seen") << index;
      }
    }
    operator char() const {
      return base;
    }
    char value() const {
      return base;
    }

    static constexpr unsigned int n_bases{6};
    static constexpr unsigned int n_good_bases{4};

   private:
    char base{};
  };

  ConsensusSequence(const unsigned int length_arg) :  // NOLINT
      base_counts(length_arg, std::array<unsigned int, Base::n_bases>{
          {0, 0, 0, 0, 0, 0}}) { }
  void add(const std::string & seq) {
    if (seq.size() != base_counts.size())
      throw Error("Bad sequence size in ConsensusSequence::add");
    for (unsigned int b{0}; b != seq.size(); ++b) {
      ++base_counts[b][BaseId{seq[b]}];
    }
  }
  // Need to call sequences() before similarity() will work
  const std::string & sequence() const {
    determine();
    return consensus;
  }

  Similarity similarity(const std::string & seq) const {
    return Similarity{seq, consensus};
  }
  unsigned int size() const {
    return static_cast<unsigned int>(base_counts.size());
  }
  unsigned int count(const unsigned int b, const char base_) const {
    return base_counts[b][BaseId{base_}];
  }
  unsigned int count(const unsigned int b, const unsigned int index) const {
    return base_counts[b][index];
  }

  void determine() const {
    consensus.clear();
    for (const auto counts : base_counts) {
      using entry = std::pair<unsigned int, unsigned int>;
      std::array<entry, Base::n_good_bases> sorted;
      for (unsigned int b{0}; b != Base::n_good_bases; ++b) {
        sorted[b] = {b, counts[b]};
      }
      sort(sorted.begin(), sorted.end(),
           [](const entry left, const entry right) {
             return left.second > right.second;
           });
      const char base_{[&sorted] {
          if (sorted.front().first < Base::n_good_bases &&
              sorted.front().second &&
              sorted[0].second != sorted[1].second) {
            return static_cast<char>(Base{sorted.front().first});
          } else {
            return 'x';
          }
        }()};
      consensus += base_;
    }
  }

 private:
  std::vector<std::array<unsigned int, Base::n_bases>> base_counts{};
  mutable std::string consensus{};
};
using Base = ConsensusSequence::Base;

class Repeatness {
 public:
  template <class longSA>
  explicit Repeatness(const std::string & sequence_arg, const longSA & sa,
                      const Reference & ref, const Mappability & mappability) :
      sequence_{sequence_arg}, mismatches_(4) {
    const auto mams = sa.find_mams(sequence_);
    n_mams_ = static_cast<unsigned int>(mams.size());
    for (const auto mam : mams) {
      if (mam.len > max_length_) max_length_ = mam.len;
    }
    for (const auto mam : mams) {
      const unsigned int chr{mam.chr};
      const unsigned int pos{mam.pos};
      const unsigned int abspos_start{ref.offset(chr) + pos};
      for (unsigned int b{0}; b != mam.len; ++b) {
        const unsigned int low_map{mappability.low(abspos_start + b)};
        if (low_map <= b + 1) {
          if (min_mappability_ > mam.len - low_map) {
            min_mappability_ = mam.len - low_map;
          }
          if (max_mappability_ < mam.len - low_map) {
            max_mappability_ = mam.len - low_map;
          }
        }
        const unsigned int high_map{mappability.high(abspos_start + b)};
        if (high_map <= mam.len - b) {
          if (min_mappability_ > mam.len - high_map) {
            min_mappability_ = mam.len - high_map;
          }
          if (max_mappability_ < mam.len - high_map) {
            max_mappability_ = mam.len - high_map;
          }
        }
      }
    }
    const std::string bowtie_command{
      "/data/software/bowtie/bowtie-0.12.8/bowtie "
          "--mm -a --best -v 3 hg19 --suppress 1,2,3,4,5,6,7 --quiet -c "};
    const std::string filter{"| perl -pe 's/,/ /g'"};
    redi::ipstream bowtie{bowtie_command + sequence_ + filter};
    std::string alignment;
    while (getline(bowtie, alignment)) {
      std::istringstream line_to_parse{alignment};
      std::string mismatch;
      unsigned int n_mismatches{0};
      while (line_to_parse >> mismatch) {
        ++n_mismatches;
      }
      ++mismatches_[n_mismatches];
    }
  }

  const std::string & sequence() const {
    return sequence_;
  }
  unsigned int n_mams() const {
    return n_mams_;
  }
  const std::vector<unsigned int> & mismatches() const {
    return mismatches_;
  }
  unsigned int max_length() const {
    return max_length_;
  }
  unsigned int min_mappability() const {
    return min_mappability_;
  }
  unsigned int max_mappability() const {
    return max_mappability_;
  }

  std::string summary() const {
    std::ostringstream out;
    out << "sequence " << sequence() << " of length " << sequence().size()
        << " has " << n_mams() << " mum" << (n_mams() == 1 ? "" : "s");
    if (n_mams()) {
      out << " with max length " << max_length()
          << " and excess mappability "
          << "min: " << min_mappability() << ", "
          << "max: " << max_mappability();
    }
    const unsigned int alignments{mismatches()[0] + mismatches()[1] +
          mismatches()[2] + mismatches()[3]};
    out << " and "
        << alignments
        << " bowtie alignments";
    if (alignments) {
      out << " with mismatches "
          << 0 << ":" << mismatches()[0] << ", "
          << 1 << ":" << mismatches()[1] << ", "
          << 2 << ":" << mismatches()[2] << ", "
          << 3 << ":" <<  mismatches()[3];
    }
    return out.str();
  }

  std::string minimal() const {
    std::ostringstream out;
    out << n_mams() << " ";
    if (n_mams()) {
      out << max_mappability();
    } else {
      out << 0;
    }
    out << " " << mismatches()[0]
        << " " << mismatches()[1]
        << " " << mismatches()[2]
        << " " << mismatches()[3];
    return out.str();
  }

 private:
  std::string sequence_{};
  unsigned int n_mams_{};
  std::vector<unsigned int> mismatches_{};
  unsigned int max_length_{0};
  unsigned int min_mappability_{255};
  unsigned int max_mappability_{0};
};

class GCATscore {
 public:
  explicit GCATscore(const std::string & sequence) {
    for (const char base : sequence) {
      switch (base) {
        case 'A':
        case 'T':
        case 'N':
            score += 2;
        break;
        case 'C':
        case 'G':
            score += 4;
        break;
        default:
          throw Error("Unknown base seen") << base;
      }
    }
  }
  operator unsigned int() const { return score; }
  unsigned int value() const { return score; }

 private:
  unsigned int score{0};
};

class Mismatches {
 public:
  static constexpr uint64_t max_mismatch{2};
  static constexpr uint64_t n_values{max_mismatch + 1};
  static constexpr unsigned int block_size{1000000};
  static constexpr uint64_t min_len{10};
  static constexpr uint64_t max_len{254};

  // Load from cache
  explicit Mismatches(const Reference & ref_) :
      ref{ref_}, data{cache_filename()} { }

  // Load from text or cache, depending
  Mismatches(const Reference & ref_,
             const std::string & text_input_dir) :
      ref{ref_} {
    if (readable(cache_filename())) {
      new (this) Mismatches{ref};
      return;
    }

    std::cerr << "Building mismatch cache at " << cache_filename() << std::endl;

    std::string c;  // chr
    unsigned int p;  // pos
    unsigned int m;  // mappability
    unsigned int s;  // skipped

    // Construct from text files
    std::vector<unsigned int> data_(n_values * ref.size());
    uint64_t n_probs{0};
    for (unsigned int chr{0}; chr != ref.n_chromosomes(); ++chr) {
      const std::string chr_name{ref.name(chr)};
      unsigned int start{0};
      const unsigned int size{ref.size(chr)};
      while (start < size) {
        const unsigned int stop{std::min(size, start + block_size)};
        const std::string file_name{text_input_dir + "/chr" + chr_name +
              "_" + std::to_string(start) + ".txt"};
        std::cerr << "Loading " << file_name << std::endl;

        std::ifstream input_file{file_name.c_str()};
        if (!input_file) {
          throw Error("Cannot open file") << file_name;
        } else {
          for (unsigned int pos{start}; pos != stop; ++pos) {
            // if (input_file.peek() == 'R') input_file.ignore(100000000, '\n');
            input_file >> c >> p >> m >> s;
            if (p != pos) throw Error("Position mismatch")
                              << p << pos << "in" << file_name;
            if (c != chr_name) throw Error("Chromosome mismatch")
                                   << c << " " << chr_name << "in" << file_name;
            std::array<unsigned int, n_values> & cs{
              *reinterpret_cast<std::array<unsigned int, n_values> *>(
                  &data_[index(ref.abspos(chr, pos))])};
            for (unsigned int & v : cs) input_file >> v;
            if (cs[0] != 1 && !s) {  // check for unexpected zero-mismatch  ma
              const std::string sequence{ref.subseq(chr, pos, pos + m)};
              std::cerr << "Unexpected zero mismatch count"
                        << chr_name << " " << pos << " "  << cs[0]
                        << " "  << s << " " << sequence.size()
                        << " " << sequence << std::endl;;
            }
            if (!input_file) throw Error("Problem reading file")
                                 << file_name << "at position" << pos << "of"
                                 << start << "to" << stop;
            input_file.get();
          }
        }
        start = stop;
      }
    }
    std::cerr << n_probs << " problems encountered" << std::endl;
    bwrite(cache_filename(), data_[0], "Mismatch binary", data_.size());
    new (this) Mismatches{ref};
  }

  uint64_t index(const unsigned int abspos,
                 const uint64_t n_mismatches = 0) const {
    return abspos * n_values + n_mismatches;
  }
  unsigned int count(const unsigned int abspos) const {
    const uint64_t start_index{index(abspos)};
    unsigned int result{0};
    for (uint64_t i{start_index}; i != start_index + n_values; ++i)
      result += data[i];
    return result;
  }
  unsigned int count(const unsigned int abspos,
                     const uint64_t n_mismatches) const {
    return data[index(abspos, n_mismatches)];
  }
  unsigned int count(const unsigned int chr, const unsigned int pos,
                     const uint64_t n_mismatches) const {
    const unsigned int abspos{ref.abspos(chr, pos)};
    return count(abspos, n_mismatches);
  }

 private:
  std::string cache_filename() const {
    return ref.fasta_file() + ".bin/mismatches.bin";
  }
  const Reference & ref;
  MappedVector<unsigned int> data{"/dev/null", false};
};

struct Repeat {
 public:
  uint64_t chromosome: 8;     // 8
  uint64_t position: 28;      // 36
  uint64_t total_length: 25;  // 61
  uint64_t start_base: 28;    // 28
  uint64_t n_copies: 28;      // 56
  uint64_t stop_base: 28;     // 28
  std::string motif;
  bool operator<(const Repeat & rhs) const {
    if (chromosome == rhs.chromosome) {
      return position < rhs.position;
    } else {
      return chromosome < rhs.chromosome;
    }
  }
};

class Repeats {
 public:
  explicit Repeats(const Reference & ref_) :
      ref{ref_} {
    const ChromosomeIndexLookup chr_lookup{ref};
    const std::string repeats_file_name{ref.fasta_file() + ".bin/repeats.txt"};
    std::ifstream repeats_file{repeats_file_name.c_str()};
    if (!repeats_file)
      throw Error("Problem opening repeats file") << repeats_file_name;
    std::string chromosome_name;
    unsigned int position;
    unsigned int total_length;
    unsigned int motif_length;
    unsigned int start_base;
    unsigned int n_copies;
    unsigned int stop_base;
    std::string motif;
    repeats.reserve(670000000);
    while (repeats_file >> chromosome_name >> position >>  total_length
           >> motif_length >> start_base >> n_copies >> stop_base >> motif) {
      repeats.push_back(Repeat{
          chr_lookup[chromosome_name], position, total_length,
              start_base, n_copies, stop_base, motif});
    }
    std::cerr << "Loaded " << repeats.size() << " repeat lines" << std::endl;
  }

  const Repeat * operator()(const unsigned int chromosome_,
                            const unsigned int position_) const {
    const Repeat to_look_up{chromosome_, position_, 0, 0, 0, 0, ""};
    auto found = upper_bound(repeats.begin(), repeats.end(), to_look_up);
    unsigned int best_length{0};
    auto best = found;
    while (found-- != repeats.begin() &&
           found->chromosome == chromosome_ &&
           found->position <= position_ &&
           position_ <
           static_cast<unsigned int>(found->position + found->total_length)) {
      if (best_length < found->total_length) {
        best_length = found->total_length;
        best = found;
      }
    }
    if (best_length) {
      return &*best;
    } else {
      return nullptr;
    }
  }

 private:
  const Reference & ref;
  std::vector<Repeat> repeats{};
};

}  // namespace paa

#endif  // PAA_SEQUENCE_H
```
