## Supplementary material for "G-Graph: An interactive genomic graph viewer": MUMdex Source Code for G-Graph: all_copyright_info.txt

** I try to only use third-party software if it has a stable interface for at least 20 years, or if I can legally and easily modify it for my needs and distribute it as part of my codebase.  I distribute source and not binaries.

>> A limited amount of my code includes non-standard (not c/c++ standard) header files that users need to have installed on their systems if they want to compile certain programs.  I currently limit myself to including these headers:

my x11plot.h includes X11/Xlib.h, X11/Xutil.h and X11/keysym.h

my cngsl.h includes gsl/gsl_cdf.h

my haha.h includes gsl/gsl_{cdf,randist}.h (does not exist yet in published code)

my python_mumdex.cpp includes Python.h

** The rest of my borrowed code is included with my package download, and has been heavily edited to suit my needs and tastes and to survive my harsh compiler warning options.  I distribute the modified code and the original licenses:

>> some suffix array code in my fasta.h, longSA.h, qsufsort.h, mummer.cpp was taken from the sparseMEM package.  Thir original license is:

* Copyright (c) 2009, Princeton University
* All rights reserved.
*
* Redistribution and use in source and binary forms, with or without
* modification, are permitted provided that the following conditions are met:
*     * Redistributions of source code must retain the above copyright
*       notice, this list of conditions and the following disclaimer.
*     * Redistributions in binary form must reproduce the above copyright
*       notice, this list of conditions and the following disclaimer in the
*       documentation and/or other materials provided with the distribution.
*     * Neither the name of the Princeton University nor the
*       names of its contributors may be used to endorse or promote products
*       derived from this software without specific prior written permission.
*
* THIS SOFTWARE IS PROVIDED BY Princeton University ''AS IS'' AND ANY
* EXPRESS OR IMPLIED WARRANTIES, INCLUDING, BUT NOT LIMITED TO, THE IMPLIED
* WARRANTIES OF MERCHANTABILITY AND FITNESS FOR A PARTICULAR PURPOSE ARE
* DISCLAIMED. IN NO EVENT SHALL Princeton University BE LIABLE FOR ANY
* DIRECT, INDIRECT, INCIDENTAL, SPECIAL, EXEMPLARY, OR CONSEQUENTIAL DAMAGES
* (INCLUDING, BUT NOT LIMITED TO, PROCUREMENT OF SUBSTITUTE GOODS OR SERVICES;
* LOSS OF USE, DATA, OR PROFITS; OR BUSINESS INTERRUPTION) HOWEVER CAUSED AND
* ON ANY THEORY OF LIABILITY, WHETHER IN CONTRACT, STRICT LIABILITY, OR TORT
* (INCLUDING NEGLIGENCE OR OTHERWISE) ARISING IN ANY WAY OUT OF THE USE OF THIS
* SOFTWARE, EVEN IF ADVISED OF THE POSSIBILITY OF SUCH DAMAGE.

and qsufsort.h has this embedded original copyright message:

/* qsufsort.c
   Copyright 1999, N. Jesper Larsson, all rights reserved.

   This file contains an implementation of the algorithm presented in "Faster
   Suffix Sorting" by N. Jesper Larsson and Kunihiko
   Sadakane.

   This software may be used freely for any purpose. However, when distributed,
   the original source must be clearly stated, and, when the source code is
   distributed, the copyright notice must be retained and any alterations in
   the code must be clearly marked. No warranty is given regarding the quality
   of this software.*/

>> I redistribute "pstream.h" as part of my package.  It has this original license:

Copyright (C) 2001,2002,2003,2004,2005,2006,2007,2008 Jonathan Wakely

This file is part of PStreams.

PStreams is free software; you can redistribute it and/or modify
it under the terms of the GNU Lesser General Public License as published by
the Free Software Foundation; either version 3 of the License, or
(at your option) any later version.

PStreams is distributed in the hope that it will be useful,
but WITHOUT ANY WARRANTY; without even the implied warranty of
MERCHANTABILITY or FITNESS FOR A PARTICULAR PURPOSE.  See the
GNU Lesser General Public License for more details.

You should have received a copy of the GNU Lesser General Public License
along with this program.  If not, see <http://www.gnu.org/licenses/>.
*/

>> I redistribute "threads.h" and indicate that I have altered it from the original.  It has this original license, which I redistribute with threads.h:

/*
  Copyright (c) 2012 Jakob Progsch, Václav Zeman
  This software is provided 'as-is', without any express or implied
  warranty. In no event will the authors be held liable for any damages
  arising from the use of this software.
  Permission is granted to anyone to use this software for any purpose,
  including commercial applications, and to alter it and redistribute it
  freely, subject to the following restrictions:

  1. The origin of this software must not be misrepresented; you must not
     claim that you wrote the original software. If you use this software
     in a product, an acknowledgment in the product documentation would be
     appreciated but is not required.
  2. Altered source versions must be plainly marked as such, and must not be
     misrepresented as being the original software.
  3. This notice may not be removed or altered from any source
     distribution.
 */

>> I redistribute cpplint.py, with this original copyright message:

# Copyright (c) 2009 Google Inc. All rights reserved.
#
# Redistribution and use in source and binary forms, with or without
# modification, are permitted provided that the following conditions are
# met:
#
#    * Redistributions of source code must retain the above copyright
# notice, this list of conditions and the following disclaimer.
#    * Redistributions in binary form must reproduce the above
# copyright notice, this list of conditions and the following disclaimer
# in the documentation and/or other materials provided with the
# distribution.
#    * Neither the name of Google Inc. nor the names of its
# contributors may be used to endorse or promote products derived from
# this software without specific prior written permission.
#
# THIS SOFTWARE IS PROVIDED BY THE COPYRIGHT HOLDERS AND CONTRIBUTORS
# "AS IS" AND ANY EXPRESS OR IMPLIED WARRANTIES, INCLUDING, BUT NOT
# LIMITED TO, THE IMPLIED WARRANTIES OF MERCHANTABILITY AND FITNESS FOR
# A PARTICULAR PURPOSE ARE DISCLAIMED. IN NO EVENT SHALL THE COPYRIGHT
# OWNER OR CONTRIBUTORS BE LIABLE FOR ANY DIRECT, INDIRECT, INCIDENTAL,
# SPECIAL, EXEMPLARY, OR CONSEQUENTIAL DAMAGES (INCLUDING, BUT NOT
# LIMITED TO, PROCUREMENT OF SUBSTITUTE GOODS OR SERVICES; LOSS OF USE,
# DATA, OR PROFITS; OR BUSINESS INTERRUPTION) HOWEVER CAUSED AND ON ANY
# THEORY OF LIABILITY, WHETHER IN CONTRACT, STRICT LIABILITY, OR TORT
# (INCLUDING NEGLIGENCE OR OTHERWISE) ARISING IN ANY WAY OUT OF THE USE
# OF THIS SOFTWARE, EVEN IF ADVISED OF THE POSSIBILITY OF SUCH DAMAGE.

>> I redistribute lowess.h, with this original message:

//
// Copyright (c) 2015, Hannes Roest
// All rights reserved.
//
// This software is released under a three-clause BSD license:
// Redistribution and use in source and binary forms, with or without
// modification, are permitted provided that the following conditions are met:
//  * Redistributions of source code must retain the above copyright
//    notice, this list of conditions and the following disclaimer.
//  * Redistributions in binary form must reproduce the above copyright
//    notice, this list of conditions and the following disclaimer in the
//    documentation and/or other materials provided with the distribution.
//  * Neither the name of any author or any participating institution
//    may be used to endorse or promote products derived from this software
//    without specific prior written permission.
// --------------------------------------------------------------------------
// THIS SOFTWARE IS PROVIDED BY THE COPYRIGHT HOLDERS AND CONTRIBUTORS "AS IS"
// AND ANY EXPRESS OR IMPLIED WARRANTIES, INCLUDING, BUT NOT LIMITED TO, THE
// IMPLIED WARRANTIES OF MERCHANTABILITY AND FITNESS FOR A PARTICULAR PURPOSE
// ARE DISCLAIMED. IN NO EVENT SHALL ANY OF THE AUTHORS OR THE CONTRIBUTING
// INSTITUTIONS BE LIABLE FOR ANY DIRECT, INDIRECT, INCIDENTAL, SPECIAL,
// EXEMPLARY, OR CONSEQUENTIAL DAMAGES (INCLUDING, BUT NOT LIMITED TO,
// PROCUREMENT OF SUBSTITUTE GOODS OR SERVICES; LOSS OF USE, DATA, OR PROFITS;
// OR BUSINESS INTERRUPTION) HOWEVER CAUSED AND ON ANY THEORY OF LIABILITY,
// WHETHER IN CONTRACT, STRICT LIABILITY, OR TORT (INCLUDING NEGLIGENCE OR
// OTHERWISE) ARISING IN ANY WAY OUT OF THE USE OF THIS SOFTWARE, EVEN IF
// ADVISED OF THE POSSIBILITY OF SUCH DAMAGE.
