## Supplementary material for "G-Graph: An interactive genomic graph viewer": MUMdex Source Code for G-Graph: add_abspos.cpp

```
//
// add_abspos
//
// add an abspos column to a file - for CN_abspos only (chrs 1..22,X,Y only)
//
// Copyright 2017 Peter Andrews @ CSHL
//

#include <exception>
#include <iostream>
#include <sstream>
#include <string>

#include "cn.h"
#include "error.h"
#include "mumdex.h"

using std::cin;
using std::cout;
using std::cerr;
using std::endl;
using std::exception;
using std::istringstream;
using std::string;

using paa::ChromosomeIndexLookup;
using paa::CN_abspos;
using paa::Error;
using paa::Reference;

int main(int argc, char * argv[]) try {
  if (--argc != 3)
    throw Error("usage: add_abspos ref chr_col_idx pos_col_idx");

  const Reference ref{argv[1]};
  const ChromosomeIndexLookup chr_lookup{ref};
  const CN_abspos cn_abspos{ref};
  const unsigned int chr_col{static_cast<unsigned int>(atoi(argv[2]))};
  const unsigned int pos_col{static_cast<unsigned int>(atoi(argv[3]))};
  string line;
  string chr_name;
  getline(cin, line);
  const bool uses_tabs{line.find('\t') != string::npos};
  const char sep_char{uses_tabs ? '\t' : ' '};
  cout << line << sep_char << "abspos" << endl;
  while (getline(cin, line)) {
    istringstream line_stream{line.c_str()};
    string value;
    unsigned int index{0};
    bool saw_chr{false};
    bool saw_pos{false};
    unsigned int chr{0};
    unsigned int pos{0};
    while (line_stream >> value) {
      if (!saw_chr && index == chr_col) {
        chr = chr_lookup(value);
        saw_chr = true;
      }
      if (!saw_pos && index == pos_col) {
        pos = atoi(value.c_str());
        saw_pos = true;
      }
      if (saw_chr && saw_pos) break;
      ++index;
    }
    if (!saw_chr) throw Error("Missing chr column") << line;
    if (!saw_pos) throw Error("Missing pos column") << line;
    const unsigned int abspos{cn_abspos(chr, pos)};
    cout << line << sep_char << abspos << endl;
  }

  return 0;
} catch (Error & e) {
  cerr << "paa::Error:" << endl;
  cerr << e.what() << endl;
  return 1;
} catch (exception & e) {
  cerr << "std::exception" << endl;
  cerr << e.what() << endl;
  return 1;
} catch (...) {
  cerr << "unknown exception was caught" << endl;
  return 1;
}
```
