## Supplementary material for "G-Graph: An interactive genomic graph viewer": MUMdex Source Code for G-Graph: agre_family.pl

```
#! /usr/bin/perl

# cat metaData.txt | ~/mumdex/agre_family.pl | grep mom | grep dad | grep -e prb -e sib | sed 's/mom/mother/g ; s/dad/father/g ; s/prb/proband/g ; s/sib/sibling/g'

use strict;

my %sample_to_ind;
my %ind_to_sample;
my %sample_gender;
my %sample_role;
my %family_samples;

while (<>) {
    chomp;
    my ($atts, $bam, $sample, $fam, $ind, $mom, $dad, $gender, $role) = split /\t/;

    $sample_to_ind{$sample} = $ind;
    $ind_to_sample{$ind} = $sample;
    $sample_gender{$sample} = $gender;
    $sample_role{$sample} = $role;
    push @{$family_samples{$fam}}, $sample;

    # print "$sample $fam $gender\n";

}

for my $family (keys %family_samples) {

    print "$family";
    for my $sample (@{$family_samples{$family}}) {
        print " $sample_role{$sample} $sample $sample_gender{$sample}";
    }
    print "\n";

}
```
