## Supplementary material for "G-Graph: An interactive genomic graph viewer": MUMdex Source Code for G-Graph: analysis_bed.sh

```
#! /bin/bash

max=$1
if [ "$max" = "" ] ; then
    echo specify maximum size 1>&2
    exit 1
fi
echo max is $max 1>&2
last_chr=chr1
n_base=0
n_lines=0
begin=0
end=0

while read chr start stop ; do
    
    if [ $chr != $last_chr ] || [ $n_base -gt $max ] ; then
        echo $last_chr $begin $end | sed 's/ /\t/g'

        last_chr=$chr
        n_base=0
        n_lines=0
        begin=$start

    fi

    n_base=$((n_base + stop - start + 1))
    n_lines=$((n_lines + 1))
    end=$stop

done
echo $last_chr $begin $end | sed 's/ /\t/g'
```
