## Supplementary material for "G-Graph: An interactive genomic graph viewer": MUMdex Source Code for G-Graph: analyze_transmission.cpp

```
//
// analyze_transmission
//
// study transmission results from transmission.cpp
//
// Copyright 2017 Peter Andrews @ CSHL
//

#include <algorithm>
#include <exception>
#include <iostream>
#include <fstream>
#include <map>
#include <memory>
#include <sstream>
#include <string>
#include <utility>
#include <vector>

#include "error.h"
#include "mumdex.h"
#include "population.h"
#include "psplot.h"
#include "stats.h"
#include "utility.h"

using std::cerr;
using std::cout;
using std::endl;
using std::exception;
using std::ifstream;
using std::istringstream;
using std::make_unique;
using std::map;
using std::ofstream;
using std::ostringstream;
using std::pair;
using std::string;
using std::to_string;
using std::unique_ptr;
using std::vector;

using paa::serr;
using paa::sout;
using paa::tout;
using paa::Bounds;
using paa::Error;
using paa::Family;
using paa::LinearRegression;
using paa::Marker;
using paa::Population;
using paa::PSDoc;
using paa::PSPage;
using paa::PSGraph;
using paa::PSHSeries;
using paa::PSXYSeries;
using paa::Reference;
using paa::Sample;
using paa::SpaceOut;

using MUMdex = paa::MUMdex;

int main(int argc, char* argv[])  try {
  --argc;
  if (argc != 4) {
    throw Error("usage: analyze_transmission ref family_file age_file "
                "results_file");
  }

  // Other command line arguments
  const string reference_file{argv[1]};
  const Reference ref{reference_file};

  const string family_file{argv[2]};
  const Population pop{family_file};

  const string ages_name{argv[3]};
  ifstream ages_file{ages_name.c_str()};
  if (!ages_file) throw Error("Could not open age file") << ages_name;

  string line;
  getline(ages_file, line);
  string family;
  unsigned int age;
  map<string, unsigned int> ages_map;
  while (ages_file >> family >> age) {
    istringstream family_stream{family.c_str()};
    string fid;
    string mem;
    getline(family_stream, fid, '.');
    family_stream >> mem;
    if (mem == "p2") continue;
    ages_map[fid + mem[0]] = age;
  }
  vector<string> samples;
  vector<unsigned int> ages;
  vector<unsigned int> is_male;
  vector<char> sample_member_types;
  map<string, unsigned int> sample_lookup;
  map<char, map<unsigned int, unsigned int>> member_sex_counts;
  for (const Sample s : pop.samples()) {
    const Family f{pop.family(s)};
    const string sample_id{pop.family(f) + pop.member(s)[0]};
    if (0)
      cout << pop.family(f) << " " << pop.sample(s) << " "
           << pop.member(s) << " "
           << ages_map[sample_id] << endl;
    sample_lookup[sample_id] = static_cast<unsigned int>(samples.size());
    ages.push_back(ages_map[sample_id]);
    samples.push_back(sample_id);
    is_male.push_back(pop.nY(s));
    sample_member_types.push_back(pop.member(s)[0]);
    ++member_sex_counts[pop.member(s)[0]][pop.nY(s)];
    ++member_sex_counts[pop.member(s)[0]][2];
  }

  const string results_name{argv[4]};
  ifstream results_file{results_name.c_str()};
  if (!results_file) throw Error("Could not open results file") << results_name;
  vector<string> event_info;
  map<char, vector<pair<unsigned int, unsigned int>>> member_events;
  // map<char, vector<pair<unsigned int, unsigned int>>> sex_member_events[2];
  map<char, map<string, unsigned int>> member_types;
  map<char, map<string, unsigned int>> sex_member_types[2];
  vector<vector<pair<unsigned int, unsigned int>>>
      sample_events(samples.size());
  while (getline(results_file, line)) {
    istringstream line_stream{line.c_str()};
    string info;
    for (unsigned int n{0}; n != 23; ++n) {
      string field;
      line_stream >> field;
      if (n) info += ' ';
      info += field;
    }
    if (info.find_first_of("XY") != string::npos) {
      continue;
    }
    string genes;
    line_stream.get();
    getline(line_stream, genes, '\t');
    info += ' ';
    info += genes;

    string all_members;
    // line_stream.get();
    getline(line_stream, all_members, '\t');
    const uint64_t n_families(count(
        all_members.begin(), all_members.end(), ' ') + 1);
    istringstream member_stream{all_members.c_str()};
    string members;
    while (member_stream >> members) {
      if (members.find(':') == string::npos)
        throw Error("Problem parsing family") << members << line;
      istringstream family_stream{members.c_str()};
      getline(family_stream, family, ':');
      info += ' ';
      info += family + ":";
      string these_members;
      family_stream >> these_members;
      istringstream these_stream{these_members.c_str()};
      char member;
      while (these_stream >> member) {
        unsigned int count;
        line_stream >> count;
        if (!line_stream) throw Error("Line stream parse error");
        info += to_string(count) + member;
        const string sample_id{family + member};
        try {
          const unsigned int sample_index{sample_lookup.at(sample_id)};
          pair<unsigned int, unsigned int> event_details{
            event_info.size(), count};
          const bool male(is_male[sample_index]);
          if (count >= 10 && n_families == 1) {
            ++member_types[member][these_members];
            ++sex_member_types[male][member][these_members];
            member_events[member].push_back(event_details);
            sample_events[sample_index].push_back(event_details);
          }
        } catch (...) {
          cerr << "Sample id not found " << sample_id << " "
               << line << endl;
        }
      }
    }

    event_info.push_back(info);
    // cout << info << endl;
  }

  for (const auto & meminfo : member_types) {
    cout << "Counts for member " << meminfo.first << endl;
    vector<pair<string, unsigned int>> sorted;
    for (const auto & memcounts : meminfo.second) {
      sorted.push_back(memcounts);
    }
    sort(sorted.begin(), sorted.end(),
         [](const pair<string, unsigned int> & lhs,
            const pair<string, unsigned int> & rhs) {
              if (lhs.second == rhs.second) {
                return lhs.first < rhs.first;
              } else {
                return lhs.second < rhs.second;
              }
            });
    uint64_t total{0};
    for (const auto & memcounts : sorted) {
      cout << memcounts.first << "\t" << memcounts.second << endl;
      total += memcounts.second;
    }
    cout << "TOTAL\t" << total << endl;
    cout << "N ind\t" << member_sex_counts[meminfo.first][2] << endl;
    cout << "per\t" << 1.0 * total / member_sex_counts[meminfo.first][2]
         << endl;
  }

  for (const char mem_char : {'p', 's'}) {
    for (const bool male : {false, true}) {
      cout << "Counts for member " << mem_char
           << " only " << (male ? "males" : "females") << endl;
      vector<pair<string, unsigned int>> sorted;
      for (const auto & memcounts : sex_member_types[male][mem_char]) {
        sorted.push_back(memcounts);
      }
      sort(sorted.begin(), sorted.end(),
           [](const pair<string, unsigned int> & lhs,
              const pair<string, unsigned int> & rhs) {
             if (lhs.second == rhs.second) {
               return lhs.first < rhs.first;
             } else {
               return lhs.second < rhs.second;
             }
           });
      uint64_t total{0};
      for (const auto & memcounts : sorted) {
        cout << memcounts.first << "\t" << memcounts.second << endl;
        total += memcounts.second;
      }
      cout << "TOTAL\t" << total << endl;
      cout << "N ind\t" << member_sex_counts[mem_char][male] << endl;
      cout << "per\t" << 1.0 * total / member_sex_counts[mem_char][male]
           << endl;
    }
  }

  // Examine age effect and sex effect
  PSDoc ps{"transmission"};
  PSGraph age_effect{ps, "Age and Sex effect;Age;N event"};
  age_effect.log_y(true);
  Marker male_marker{paa::circle(), 0.2, "0 0 1", 1, true};
  Marker female_marker{paa::circle(), 0.2, "1 0 0", 1, true};
  Marker * fm_markers[2]{&female_marker, &male_marker};

  PSXYSeries age_male{age_effect, male_marker};
  PSXYSeries age_female{age_effect, female_marker};
  PSXYSeries * age_graphs[2]{&age_female, &age_male};

  map<char, vector<std::unique_ptr<PSGraph>>> graphs;
  map<char, vector<std::unique_ptr<PSHSeries<unsigned int, unsigned int>>>>
      series;

  map<char, vector<vector<unsigned int>>> ranked_counts;

  const string sexes[2]{"female", "male"};
  for (const char mem : {'m', 'f', 'p', 's'}) {
    for (const unsigned int x : {0, 1, 2}) {
      ranked_counts[mem].resize(3);
      if (x != 2 && (mem == 'm' || mem == 'f')) {
        graphs[mem].push_back(nullptr);
        series[mem].push_back(nullptr);
        continue;
      }
      ostringstream title;
      title << mem << " sample event counts";
      if (x != 2) {
        title << " " << sexes[x] << " only";
      }
      title << ";Count;N";
      graphs[mem].push_back(make_unique<PSGraph>(ps, title.str(),
                                                 Bounds{0.0, 10000.0}));
      series[mem].push_back(
          make_unique<PSHSeries<unsigned int, unsigned int> >(
              *graphs[mem].back(), 100, "1 0 0", false));
    }
  }


  uint64_t sex_totals[2]{0, 0};
  uint64_t sex_events[2]{0, 0};
  map<char, uint64_t> member_bridge_counts;
  map<char, uint64_t> member_n_events;
  vector<double> sample_n_events;

  for (unsigned int s{0}; s != samples.size(); ++s) {
    uint64_t total_count{0};
    for (auto & ei : sample_events[s]) {
      total_count += ei.second;
    }
    member_bridge_counts[sample_member_types[s]] += total_count;
    member_n_events[sample_member_types[s]] += sample_events[s].size();

    sex_totals[is_male[s]] += total_count;
    sex_events[is_male[s]] += sample_events[s].size();
    // const double average_count{1.0 * total_count / sample_events[s].size()};
    sample_n_events.push_back(sample_events[s].size());
    age_graphs[is_male[s]]->add_point(
        ages[s] / 12.0,
        sample_events[s].size());
    series[sample_member_types[s]][2]->add_point(
        static_cast<unsigned int>(sample_events[s].size()));
    if (sample_member_types[s] == 'p' || sample_member_types[s] == 's') {
      series[sample_member_types[s]][is_male[s]]->add_point(
          static_cast<unsigned int>(sample_events[s].size()));
    }
    ranked_counts[sample_member_types[s]][2].push_back(
        static_cast<unsigned int>(sample_events[s].size()));
    ranked_counts[sample_member_types[s]][is_male[s]].push_back(
        static_cast<unsigned int>(sample_events[s].size()));
  }

  map<char, std::unique_ptr<PSGraph>> frac_graphs;
  map<char, vector<std::unique_ptr<PSXYSeries>>> frac_series;

  for (const char mem : {'m', 'f', 'p', 's'}) {
    ostringstream title;
    title << mem << " sample event counts";
    title << ";Rank;Count";
    frac_graphs[mem] = make_unique<PSGraph>(
        ps, title.str(), Bounds{0.0, 1.0});
    for (const unsigned int x : {0, 1}) {
      vector<unsigned int> & counts{ranked_counts[mem][x]};
      sort(counts.begin(), counts.end());
      if ((mem == 'm' && x == 0) ||
          (mem == 'f' && x == 1) ||
          mem == 'p' || mem == 's') {
        frac_series[mem].push_back(make_unique<PSXYSeries>(
            *frac_graphs[mem], *fm_markers[x]));
        for (unsigned int c{0}; c != counts.size(); ++c) {
          frac_series[mem].back()->add_point(1.0 * c / counts.size(),
                                            counts[c]);
        }
      }
    }
  }
  frac_graphs['f']->add(&*frac_series['m'].back());

  for (const char member : {'m', 'f', 'p', 's'}) {
    cerr << member << " average autosome count is "
         << 1.0 * member_bridge_counts[member] / member_n_events[member]
         << endl;
  }

  for (const bool male : {false, true}) {
    cerr << sexes[male] << " average autosome count is "
         << 1.0 * sex_totals[male] / sex_events[male] << endl;
  }

  cerr << "Sex ratio is "
       << 1.0 * sex_totals[1] / sex_events[1] /
      (1.0 * sex_totals[0] / sex_events[0]) << endl;
  uint64_t n_bases[2]{0, 0};
  for (unsigned int c{0}; c!= ref.n_chromosomes(); ++c) {
    if (ref.name(c) != "X" && ref.name(c) != "Y") {
      for (const bool male : {false, true}) {
        n_bases[male] += 2 * ref.size(c);
      }
    } else {
      n_bases[1] += ref.size(c);
      if (ref.name(c) == "X") {
        n_bases[0] += 2 * ref.size(c);
      }
    }
  }
  cerr << "Theory gives " << 1.0 * n_bases[0] / n_bases[1] << endl;
  const LinearRegression age_line{ages, sample_n_events};
  cerr << "N event = "
       << age_line.intercept << " +/- " << age_line.intercept_error << " + "
       << age_line.slope << " +/- " << age_line.slope_error << " * age" << endl;

  ofstream family_table{"family_info.txt"};
  for (const Family fam : pop.families()) {
    const string & family_name{pop.family(fam)};
    family_table << family_name;
    for (const char member : {'m', 'f', 'p', 's'}) {
      const string sample_id{family_name + member};
      const unsigned int sample_index{sample_lookup.at(sample_id)};
      family_table << "\t" << sample_events[sample_index].size();
    }
    family_table << "\t" << is_male[sample_lookup.at(family_name + 'p')];
    family_table << "\t" << is_male[sample_lookup.at(family_name + 's')];
    family_table << endl;
  }

  return 0;
} catch (Error & e) {
  cerr << "paa::Error:" << endl;
  cerr << e.what() << endl;
  return 1;
} catch (exception & e) {
  cerr << "std::exception" << endl;
  cerr << e.what() << endl;
  return 1;
} catch (...) {
  cerr << "unknown exception was caught" << endl;
  return 1;
}
```
