## Supplementary material for "G-Graph: An interactive genomic graph viewer": MUMdex Source Code for G-Graph: anchor_mismatches.cpp

```
//
// anchor_mismatches
//
// measure the number of mismatches in genome for anchors in bridges from stdin
// by counting mappings with N mismatches for N in 0..3
//
// Copyright 2018 Peter Andrews @ CSHL
//

#include <exception>
#include <iostream>
#include <sstream>
#include <string>
#include <vector>

#include "error.h"
#include "mumdex.h"
#include "pstream.h"

using redi::ipstream;

using std::cerr;
using std::cin;
using std::cout;
using std::endl;
using std::exception;
using std::istringstream;
using std::string;
using std::to_string;
using std::vector;

using paa::ChromosomeIndexLookup;
using paa::Error;
using paa::Reference;

int main(int argc, char* argv[])  try {
  --argc;
  if (argc != 2) throw Error("anchor_mismatches n_mismatches ref_fasta");

  const unsigned int n_mismatch{static_cast<unsigned int>(atoi(argv[1]))};
  const string ref_fasta{argv[2]};
  const Reference ref{ref_fasta};
  const ChromosomeIndexLookup chr_lookup{ref};

  string chr_name;
  unsigned int pos;
  unsigned int high;
  unsigned int support;
  unsigned int n{0};
  unsigned int total{0};
  while (cin >> chr_name >> pos >> high >> support) {
    if ((n % 2) == 0) cout << n / 2 + 1;
    const unsigned int chr{chr_lookup[chr_name]};
    const unsigned int start{high ? pos - support + 1 : pos};
    const unsigned int stop{high ? pos + 1 : pos + support};
    const string sequence{ref.subseq(chr, start, stop)};
    if (sequence.find_first_not_of("ACGTacgt") != string::npos)
      throw Error("Unexpected base found in") << sequence;
    vector<unsigned int> counts(n_mismatch + 1);
    const std::string bowtie_base{
      "/data/software/bowtie/bowtie-0.12.8/bowtie "
          "--quiet --mm -a -v " + to_string(n_mismatch) +
          " --suppress 1,2,3,4,5,6,7 "
          "/data/software/bowtie/bowtie-1.2.2/indexes/g1k -c "};
    const std::string filter{"| perl -pe 's/,/ /g'"};

    // Run bowtie command and get output
    const std::string bowtie_command{
      bowtie_base + sequence + filter};
    if (false) cout << "Running: " << bowtie_command << endl;
    redi::ipstream bowtie{bowtie_command};
    if (!bowtie) throw Error("Bowtie failed on ") << bowtie_command;
    string alignment;
    string mismatch;
    while (getline(bowtie, alignment)) {
      if (false) cout << "alignment " << alignment << endl;
      unsigned int n_mismatches{0};
      istringstream align_stream{alignment.c_str()};
      while (align_stream >> mismatch) ++n_mismatches;
      ++counts[n_mismatches];
    }
    if (counts[0] != 1) throw Error("Unexpected count for zero mismatches");
    for (unsigned int m{1}; m != n_mismatch + 1; ++m) {
      cout << "\t" << counts[m];
      total += counts[m];
    }
    if ((++n % 2) == 0) {
      cout << "\t" << total<< "\n";
      total = 0;
    }
  }
  cerr << "done" << endl;

  return 0;
} catch (Error & e) {
  cerr << "paa::Error:" << endl;
  cerr << e.what() << endl;
  return 1;
} catch (exception & e) {
  cerr << "std::exception" << endl;
  cerr << e.what() << endl;
  return 1;
} catch (...) {
  cerr << "unknown exception was caught" << endl;
  return 1;
}
```
