## Supplementary material for "G-Graph: An interactive genomic graph viewer": MUMdex Source Code for G-Graph: anchor_reads.cpp

```
//
// anchor_reads
//
// show reads with a given anchor
//
// Copyright Peter Andrews 2015 @ CSHL
//

#include <algorithm>
#include <exception>
#include <iostream>
#include <fstream>
#include <limits>
#include <set>
#include <sstream>
#include <string>
#include <vector>

#include "error.h"
#include "genes.h"
#include "mumdex.h"
#include "utility.h"

using std::exception;
using std::cout;
using std::cerr;
using std::endl;
using std::ifstream;
using std::ofstream;
using std::istringstream;
using std::numeric_limits;
using std::set;
using std::string;
using std::vector;

using paa::base_chars;
using paa::base_index;
using paa::complement;
using paa::reverse_complement;
using paa::ChromosomeIndexLookup;
using paa::Error;
using paa::MUMdex;
using paa::GeneXrefs;
using paa::KnownGenes;
using paa::Mappability;
using paa::MUM;
using paa::PosInfo;
using paa::Reference;
using paa::serr;
using paa::sout;

int main(int argc, char* argv[]) try {
  paa::exit_on_pipe_close();

  --argc;
  if (argc != 5) throw Error("usage: anchor_reads search_info_file"
                             "ref_fa genes isoforms xrefs");

  const int max_read_length = 120;
  const bool show_mums = false;

  ifstream search_info{argv[1]};
  if (!search_info) throw Error("Could not open search_info file") << argv[1];

  const string ref_name{argv[2]};
  const string genes_name{argv[3]};
  const string isoforms_name{argv[4]};
  const string kgXrefs_name{argv[5]};

  const Mappability map{ref_name, true};

  const Reference ref{ref_name};
  const ChromosomeIndexLookup chr_lookup{ref};

  const KnownGenes genes{genes_name, isoforms_name, chr_lookup, ref};
  const GeneXrefs xref{kgXrefs_name};

  ofstream sequences("sequences.txt");
  unsigned int last_gene_index = static_cast<unsigned int>(-1);
  string last_sample;
  string line;
  set<uint64_t> pairs_for_gene;
  while (true) {
    getline(search_info, line);
    if (line.empty()) break;
    istringstream in{line.c_str()};
    string sample;
    string mumdex_name;
    unsigned int gene_index;
    string gene_symbol;
    unsigned int junction_index;
    string chromosome;
    unsigned int invariant_index;
    int used_invariant;
    unsigned int position;
    int invariant;
    vector<unsigned int> positions;
    in >> sample >> mumdex_name >> gene_index >> gene_symbol
       >> junction_index >> chromosome >> invariant;
    if (!in) throw Error("Parse error");
    if (invariant >= 0) throw Error("Bad invariant") << invariant;
    const string gene_sample = gene_symbol + mumdex_name;
    if (gene_index != last_gene_index || sample != last_sample) {
      if (last_sample.size()) sequences << endl << endl;
      pairs_for_gene.clear();
      sequences << sample << " " << gene_index << " "
                << gene_symbol << " " << mumdex_name << endl;
    }
    last_sample = sample;
    last_gene_index = gene_index;
    const MUMdex mumdex{mumdex_name};
    const auto gene = genes[gene_index];
    const unsigned int chr{chr_lookup[chromosome]};
    unsigned int n = 0;
    set <uint64_t> pairs;
    unsigned int n_low_anchors = 0;
    unsigned int n_high_anchors = 0;
    set <unsigned int> low_positions;
    set <unsigned int> high_positions;
    while (in >> invariant_index >> used_invariant >> position) {
      if (invariant_index != 0) {
        cerr << "ignoring skipped exon junction" << endl;
        continue;
      }
      low_positions.insert(position);
      if (show_mums) sout << "Looking for anchors at"
                          << chromosome << position << used_invariant << endl;
      const PosInfo posinfo{chr, position};
      for (auto mum_iter = mumdex.lower_bound(posinfo);
           mum_iter != mumdex.index().end(); ++mum_iter) {
        const auto mum = mumdex.mum(*mum_iter);
        const auto pair = mumdex.pair(*mum_iter);
        if (pair.dupe()) continue;
        if (mum.chromosome() != chr || mum.position0() != position) break;
        if (mum.flipped()) {
          if (mum.touches_end()) continue;
        } else {
          if (mum.offset() == 0) continue;
        }
        const auto pair_index = mum_iter->pair_index();
        if (show_mums)
          sout << "low mum" << ++n << chromosome << position
               << mum.flipped() << mum.length()
               << mum.offset() << mum.touches_end()
               << pair_index << endl;
        ++n_low_anchors;
        pairs.insert(pair_index);
        for (auto other_iter = mumdex.mums_begin(pair_index);
             other_iter != mumdex.mums_end(pair_index); ++other_iter) {
          const auto other = *other_iter;
          if (other == mum) continue;
          if (other.flipped() != mum.flipped()) continue;
          if (other.chromosome() != mum.chromosome()) continue;
          if (other.read_2() != mum.read_2()) continue;
          if (other.read_position0(pair.length(other.read_2())) -
              mum.read_position0(pair.length(mum.read_2())) != used_invariant)
            continue;
          high_positions.insert(other.position0() + other.length());
        }
      }
      const PosInfo low_posinfo{chr, static_cast<int>(position) >
            max_read_length - used_invariant ?
            position - max_read_length + used_invariant : 0};
      const PosInfo high_posinfo{chr, low_posinfo.pos + 2 * max_read_length};
      for (auto mum_iter = mumdex.lower_bound(low_posinfo);
           mum_iter != mumdex.lower_bound(high_posinfo); ++mum_iter) {
        const auto mum = mumdex.mum(*mum_iter);
        const auto pair = mumdex.pair(*mum_iter);
        if (pair.dupe()) continue;
        if (!mum.flipped()) {
          if (mum.touches_end()) continue;
        } else {
          if (mum.offset() == 0) continue;
        }
        const auto pair_index = mum_iter->pair_index();
        // if (pairs.count(pair_index)) continue;
        bool matches = false;
        for (const auto pos : high_positions) {
          if (mum.position0() + mum.length() == pos) {
            matches = true;
          }
        }
        if (!matches) continue;
        pairs.insert(pair_index);
        if (show_mums)
          sout << "high mum" << ++n << chromosome
               << mum.position0() + mum.length()
               << mum.flipped() << mum.length()
               << mum.offset() << mum.touches_end()
               << pair_index << endl;
        ++n_high_anchors;
      }
    }
    if (low_positions.empty()) {
      continue;
    }
    sout << "Index file" << mumdex_name << endl;
    sout << "Gene" << gene_symbol
         << "isoform" << genes[gene_index].name
         << "junction" << genes.junctions[junction_index].junction_index
         << "on" << chromosome << "for invariant" << invariant << endl;
    sout << "found" << n_low_anchors << "low anchors at";
    for (const auto pos : low_positions) {
      sout << pos;
    }
    sout << "and" << n_high_anchors << "high anchors at";
    for (const auto pos : high_positions) {
      sout << pos;
    }
    sout << "in" << pairs.size() << "pairs" << endl;

    unsigned int low_pos = numeric_limits<unsigned int>::max();
    unsigned int high_pos = 0;
    for (const auto pair_index : pairs) {
      for (auto mum_iter = mumdex.mums_begin(pair_index);
           mum_iter != mumdex.mums_end(pair_index); ++mum_iter) {
        const auto mum = *mum_iter;
        if (mum.chromosome() != chr) continue;
        bool in_exon = false;
        unsigned int E = 0;
        for (unsigned int e = 0; e != gene.n_exons; ++e) {
          if ((mum.position0() >= gene.exon_starts[e] &&
               mum.position0() < gene.exon_stops[e]) ||
              (mum.position0() + mum.length() >= gene.exon_starts[e] &&
               mum.position0() + mum.length() < gene.exon_stops[e]) ||
              (mum.position0() < gene.exon_starts[e] &&
               mum.position0() + mum.length() >= gene.exon_stops[e])) {
            in_exon = true;
            E = e;
            break;
          }
        }
        if (!in_exon) continue;
        const auto low = mum.position0() < gene.exon_starts[E] ?
            gene.exon_starts[E] : mum.position0();
        const auto high = mum.position0() + mum.length() >= gene.exon_stops[E] ?
            gene.exon_stops[E] : mum.position0() + mum.length();
        if (low < low_pos) low_pos = low;
        if (high > high_pos) high_pos = high;
      }
    }
    unsigned int distance = 0;
    unsigned int skipped = 0;
    for (unsigned int e = 0; e != gene.n_exons; ++e) {
      if (low_pos >= gene.exon_stops[e]) continue;
      if (low_pos >= gene.exon_starts[e] &&
          low_pos < gene.exon_stops[e] &&
          high_pos >= gene.exon_starts[e] &&
          high_pos < gene.exon_stops[e]) {
        throw Error("Unexpected in-exon");
        distance += high_pos - low_pos;
        break;
      }
      if (low_pos >= gene.exon_starts[e] &&
          low_pos < gene.exon_stops[e]) {
        distance += gene.exon_stops[e] - low_pos;
      } else if (high_pos >= gene.exon_starts[e] &&
                 high_pos < gene.exon_stops[e]) {
        distance += high_pos - gene.exon_starts[e];
        ++skipped;
      } else if (high_pos >= gene.exon_stops[e]) {
        distance += gene.exon_stops[e] - gene.exon_starts[e];
        ++skipped;
      } else {
        break;
      }
    }
    sout << "Length" << high_pos - low_pos
         << "minus up to" << skipped << "skipped introns =" << distance
         << "low pos" << low_pos << "high pos" << high_pos << endl;
    vector<vector<char>> scaffold(distance);
    int distance_in[2]{0, 0};
    for (const auto pair_index : pairs) {
      const auto pair = mumdex.pair(pair_index);
      unsigned int max_lengths[2]{0, 0};
      MUM best_mums[2];
      int offsets[2]{0, 0};
      for (auto mum_iter = mumdex.mums_begin(pair_index);
           mum_iter != mumdex.mums_end(pair_index); ++mum_iter) {
        const auto mum = *mum_iter;
        const auto read = mum.read_2();
        if (mum.chromosome() != chr) continue;
        bool in_exon = false;
        const auto pos = mum.position0();
        for (unsigned int e = 0; e != gene.n_exons; ++e) {
          if ((mum.position0() >= gene.exon_starts[e] &&
               mum.position0() < gene.exon_stops[e]) ||
              (mum.position0() + mum.length() >= gene.exon_starts[e] &&
               mum.position0() + mum.length() < gene.exon_stops[e]) ||
              (mum.position0() < gene.exon_starts[e] &&
               mum.position0() + mum.length() >= gene.exon_stops[e])) {
            in_exon = true;
            break;
          }
        }
        if (!in_exon) continue;
        if (mum.length() > max_lengths[read]) {
          unsigned int pos_in = 0;
          unsigned int offset = 0;
          for (unsigned int e = 0; e != gene.n_exons; ++e) {
            if (pos < low_pos) {
              pos_in = 0;
              offset = low_pos - pos;
              break;
            }
            if (low_pos >= gene.exon_stops[e]) continue;
            if (low_pos >= gene.exon_starts[e] &&
                low_pos < gene.exon_stops[e]) {
              if (pos >= gene.exon_starts[e] &&
                  pos < gene.exon_stops[e]) {
                pos_in += pos - low_pos;
              } else {
                pos_in += gene.exon_stops[e] - low_pos;
              }
            } else if (pos >= gene.exon_starts[e] &&
                       pos < gene.exon_stops[e]) {
              pos_in += pos - gene.exon_starts[e];
            } else if (pos >= gene.exon_stops[e]) {
              pos_in += gene.exon_stops[e] - gene.exon_starts[e];
            } else if (pos < gene.exon_starts[e] &&
                       pos + mum.length() >= gene.exon_starts[e]) {
              offset = gene.exon_starts[e] - pos;
              break;
            }
          }
          max_lengths[read] = mum.length();
          best_mums[read] = mum;
          distance_in[read] = pos_in;
          offsets[read] = offset;
        }
      }
      auto seqs = mumdex.sequences(pair_index);
      for (const bool r : {false, true}) {
        if (!max_lengths[r]) continue;
        const auto mum = best_mums[r];
        const auto & sequence = seqs[r];
        const auto read_start = mum.read_position0(pair.length(r));
        for (auto p = read_start;
             p != read_start + static_cast<int>(pair.length(r)); ++p) {
          if (p < static_cast<int>(low_pos)) continue;
          if (p >= static_cast<int>(high_pos)) break;
          const auto b = mum.pos_to_base(p);
          const auto base = sequence[b];
          const auto rcbase = mum.flipped() ? complement(base) : base;
          const auto index = distance_in[r] - offsets[r] + p -
              static_cast<int>(mum.position0());
          if (0) serr << scaffold.size() << low_pos << high_pos
                      << p << distance_in[r] << mum.position0()
                      << index << rcbase << endl;
          if (index >=0 && index < static_cast<int>(scaffold.size()))
            scaffold[index].push_back(rcbase);
        }
      }
      if (max_lengths[0] == 0 && max_lengths[1] == 0)
        throw Error("No best mum found");
      const auto best_read = max_lengths[0] >= max_lengths[1] ? 0 : 1;
      if (best_mums[best_read].flipped()) {
        reverse_complement(&seqs[best_read]);
      } else {
        reverse_complement(&seqs[1 - best_read]);
      }
      for (const bool r : {false, true}) {
        cout << seqs[r] << endl;
      }
      if (pairs_for_gene.count(pair_index) == 0) {
        for (const bool r : {false, true}) {
          sequences << seqs[r] << endl;
        }
        pairs_for_gene.insert(pair_index);
      }
    }
    unsigned int d = 0;
    for (unsigned int e = 0; e != gene.n_exons; ++e) {
      if (gene.exon_stops[e] <= low_pos) continue;
      if (gene.exon_starts[e] > high_pos) break;
      for (unsigned int p = gene.exon_starts[e]; p != gene.exon_stops[e];
           ++p ) {
        const bool show_scaffolding = true;
        if (p >= low_pos && p < high_pos) {
          if (show_scaffolding)
            sout << d << chromosome << p << e << ref[chr][p] << "";
          auto & bases = scaffold[d];
          sort(bases.begin(), bases.end());
          if (show_scaffolding) {
            for (const auto base : bases) {
              cout << base;
            }
            if (bases.empty()) cout << '-';
          }
          bases.push_back(ref[chr][p]);
          vector<unsigned int> base_counts(6);
          for (const auto base : bases) {
            ++base_counts[base_index(base)];
          }
          unsigned int most_base = 0;
          for (unsigned int b = 1; b != 5; ++b) {
            if (base_counts[b] > base_counts[most_base]) {
              most_base = b;
            }
          }
          unsigned int next_most_base = 0;
          for (unsigned int b = 1; b != 5; ++b) {
            if (b == most_base) continue;
            if (base_counts[b] > base_counts[next_most_base]) {
              next_most_base = b;
            }
          }
          const auto best_base_tmp = most_base ?
              (base_counts[most_base] == base_counts[next_most_base] ? '!' :
               base_chars[most_base]) : ' ';
          const auto best_base = bases.size() == 1 ?
              static_cast<char>(tolower(best_base_tmp)) : best_base_tmp;
          if (show_scaffolding) {
            sout << best_base;
            if (best_base == '!') {
              sout << "UNDECIDED";
            } else {
              if (best_base_tmp != ref[chr][p]) sout << "REF_MISMATCH";
            }
            sout << endl;
          }
          ++d;
        }
      }
    }
    // mumdex.pair_view(cout, pair_index, {{mum, "E"}});
  }

  return 0;
} catch (Error & e) {
  cerr << "paa::Error:" << endl;
  cerr << e.what() << endl;
  return 1;
} catch (exception & e) {
  cerr << "std::exception" << endl;
  cerr << e.what() << endl;
  return 1;
} catch (...) {
  cerr << "unknown exception was caught" << endl;
  return 1;
}
```
