## Supplementary material for "G-Graph: An interactive genomic graph viewer": MUMdex Source Code for G-Graph: anchor_repeatness.cpp

```
//
// anchor_repeatness
//
// are anchors sufficiently unique in their sequence
//
// Copyright 2016 Peter Andrews @ CSHL
//

#include <exception>
#include <iostream>
#include <string>
#include <vector>

#include "error.h"
#include "longSA.h"
#include "mapper.h"
#include "mumdex.h"
#include "sequence.h"

using std::cerr;
using std::cout;
using std::endl;
using std::exception;
using std::string;
using std::vector;

using paa::longSA;
using paa::ChromosomeIndexLookup;
using paa::Error;
using paa::Mappability;
using paa::Reference;
using paa::Repeatness;

int main(int argc, char* argv[])  try {
  --argc;
  if (argc != 7 && argc != 9)
    throw Error("usage: anchor_repeatness ref_fasta "
                "chr1 pos1 high1 chr2 pos2 high2 [len1 len2]");

  const string ref_fasta{argv[1]};
  const longSA sa{ref_fasta.c_str(), true, true, false};
  const Reference ref{ref_fasta};
  const Mappability mappability{ref_fasta, true};
  const ChromosomeIndexLookup lookup{ref};

  const unsigned int len1{argc == 9 ?
        static_cast<unsigned int>(atoi(argv[8])) : 0};
  const unsigned int len2{argc == 9 ?
        static_cast<unsigned int>(atoi(argv[9])) : 0};

  ++argv;
  for (const bool pos2 : {false, true}) {
    // Get anchor positions
    const string achr_string{argv[1]};
    const unsigned int achr{lookup[achr_string]};
    const unsigned int apos{static_cast<unsigned int>(atoi(argv[2]))};
    const bool high{static_cast<bool>(atoi(argv[3]))};
    argv += 3;

    // Mappability at anchor
    const unsigned int mapp{mappability.low_high(
        high, ref.abspos(achr, apos))};

    const unsigned int max_len{pos2 ? len2 : len1};
    const unsigned int len{max_len ? max_len : mapp};

    const unsigned int start_pos{high ?
          (apos + 1 > len ? apos - len + 1 : 0) : apos};
    const unsigned int stop_pos{high ? apos + 1 : apos + len};
    const string sequence{ref.subseq(achr, start_pos, stop_pos)};
    const Repeatness repeatness{sequence, sa, ref, mappability};
    if (0) cout << achr << " "
                << apos << " "
                << high << " "
                << sequence << " "
                << sequence.size() << " "
                << len << " "
                << repeatness.n_mams() << " ";
    cout << repeatness.max_mappability() << " "
         << repeatness.mismatches()[0] << " "
         << repeatness.mismatches()[1] << " "
         << repeatness.mismatches()[2] << " "
         << repeatness.mismatches()[3] << endl;
  }

  return 0;
} catch (Error & e) {
  cerr << "paa::Error:" << endl;
  cerr << e.what() << endl;
  return 1;
} catch (exception & e) {
  cerr << "std::exception" << endl;
  cerr << e.what() << endl;
  return 1;
} catch (...) {
  cerr << "unknown exception was caught" << endl;
  return 1;
}
```
