## Supplementary material for "G-Graph: An interactive genomic graph viewer": MUMdex Source Code for G-Graph: annotate_candidates.cpp

```
//
// annotate_candidates
//
// Add repeat and mismatch information to a candidate list
//
// Copyright 2018 Peter Andrews @ CSHL
//

#include <algorithm>
#include <exception>
#include <iostream>
#include <sstream>
#include <string>
#include <vector>

#include "error.h"
#include "mumdex.h"
// #include "repeats.h"
#include "sequence.h"

using std::cin;
using std::cout;
using std::cerr;
using std::endl;
using std::exception;
using std::istringstream;
using std::ostringstream;
using std::string;
using std::vector;

using paa::ChromosomeIndexLookup;
using paa::Error;
using paa::Reference;
using paa::Repeat;
using paa::Repeats;
using paa::Mismatches;

int main(int argc, char * argv[]) try {
  if (--argc < 3)
    throw Error("usage: annotate_candidates ref pos_offset chr_col ..");

  // process arguments
  const Reference ref{argv[1]};
  const ChromosomeIndexLookup chr_lookup{ref};
  const int pos_offset{atoi(argv[2])};
  argc -= 2;
  argv += 2;
  vector<unsigned int> chrs;
  while (argc--) {
    chrs.push_back(atoi(argv++[1]));
  }
  sort(chrs.begin(), chrs.end());
  if (chrs.empty()) throw Error("No chromosome columns");

  // data sources
  const Mismatches mismatches{ref};
  const Repeats repeats{ref};

  // output header
  string line;
  getline(cin, line);
  cout << line;
  for (unsigned int c{0}; c != chrs.size(); ++c) {
    const char letter{static_cast<char>('A' + c)};
    cout << " echo_1" << letter
         << " echo_2" << letter
         << " rep_total_length" << letter
         << " rep_motif_length" << letter
         << " rep_motif_copies" << letter
         << " rep_motif_start" << letter
         << " rep_motif_stop" << letter
         << " rep_motif" << letter;
  }
  cout << endl;

  string entry;
  unsigned int chr{0};
  unsigned int pos{0};
  while (getline(cin, line)) {
    istringstream in{line.c_str()};
    unsigned int col{0};
    unsigned int cn{0};
    ostringstream annotation;
    while (in >> entry) {
      const unsigned int ccol{chrs[cn]};
      const unsigned int pcol{static_cast<unsigned int>(
          static_cast<int>(ccol) + pos_offset)};
      bool ready{false};
      if (col == ccol) {
        chr = chr_lookup[entry];
        if (pos_offset < 0) ready = true;
      } else if (col == pcol) {
        pos = static_cast<unsigned int>(stoul(entry));
        if (pos_offset > 0) ready = true;
      }
      if (ready) {
        for (unsigned int m{1}; m != 3 ; ++m)
          annotation << " " << mismatches.count(chr, pos, m);
        const Repeat * found{repeats(chr, pos)};
        if (found) {
          annotation << " " << found->total_length
                     << " " << found->motif.size()
                     << " " << found->n_copies
                     << " " << found->start_base
                     << " " << found->stop_base
                     << " " << found->motif;
        } else {
          annotation << " 0 0 0 0 0 0";
        }
        ++cn;
        if (cn == chrs.size()) {
          cout << line << annotation.str() << endl;
          break;
        }
      }
      ++col;
    }
  }

  return 0;
} catch (Error & e) {
  cerr << "paa::Error:" << endl;
  cerr << e.what() << endl;
  return 1;
} catch (exception & e) {
  cerr << "std::exception" << endl;
  cerr << e.what() << endl;
  return 1;
} catch (...) {
  cerr << "unknown exception was caught" << endl;
  return 1;
}
```
