## Supplementary material for "G-Graph: An interactive genomic graph viewer": MUMdex Source Code for G-Graph: annotate_repeats.cpp

```
//
// annotate_repeats
//
// Add repeat information to a candidate list
//
// Copyright 2016 Peter Andrews @ CSHL
//

#include <algorithm>
#include <exception>
#include <iostream>
#include <sstream>
#include <string>
#include <vector>

#include "error.h"
#include "mumdex.h"
#include "repeats.h"

using std::cin;
using std::cout;
using std::cerr;
using std::endl;
using std::exception;
using std::istringstream;
using std::ostringstream;
using std::string;
using std::vector;

using paa::ChromosomeIndexLookup;
using paa::Error;
using paa::Reference;
using paa::MaskerRepeat;
using paa::RepeatMasker;

int main(int argc, char * argv[]) try {
  if (--argc < 4)
    throw Error("usage: annotate_repeats repeats_file "
                "ref pos_offset chr_col ..");

  const Reference ref{argv[2]};
  const ChromosomeIndexLookup chr_lookup{ref};
  const RepeatMasker repeats{argv[1], chr_lookup};
  const int pos_offset{atoi(argv[3])};
  argc -= 2;
  argv += 2;

  vector<unsigned int> chrs;
  while (--argc) {
    ++argv;
    chrs.push_back(atoi(argv[1]));
  }
  sort(chrs.begin(), chrs.end());
  if (chrs.empty()) throw Error("No chromosome columns");

  string line;
  string entry;
  unsigned int chr{0};
  unsigned int pos{0};
  while (getline(cin, line)) {
    istringstream in{line.c_str()};
    unsigned int col{0};
    unsigned int cn{0};
    ostringstream annotation;
    while (in >> entry) {
      const unsigned int ccol{chrs[cn]};
      const unsigned int pcol{static_cast<unsigned int>(
          static_cast<int>(ccol) + pos_offset)};
      bool ready{false};
      if (col == ccol) {
        chr = chr_lookup[entry];
        if (pos_offset < 0) ready = true;
      } else if (col == pcol) {
        pos = static_cast<unsigned int>(stoul(entry));
        if (pos_offset > 0) ready = true;
      }
      if (ready) {
        if (cn) {
          annotation << ";";
        }
        const vector<const MaskerRepeat *> result{repeats(chr, pos)};
        for (unsigned int r{0}; r != result.size(); ++r) {
          const MaskerRepeat * repeat{result[r]};
          if (r) {
            annotation << ",";
          }
          annotation << repeat->repeat << "-" << repeat->family;
        }
        ++cn;
        if (cn == chrs.size()) {
          cout << line << " " << annotation.str() << endl;
          break;
        }
      }
      ++col;
    }
  }

  return 0;
} catch (Error & e) {
  cerr << "paa::Error:" << endl;
  cerr << e.what() << endl;
  return 1;
} catch (exception & e) {
  cerr << "std::exception" << endl;
  cerr << e.what() << endl;
  return 1;
} catch (...) {
  cerr << "unknown exception was caught" << endl;
  return 1;
}
```
