## Supplementary material for "G-Graph: An interactive genomic graph viewer": MUMdex Source Code for G-Graph: assembler.cpp

```
//
// assembler
//
// debruijn read pair assembly and viewing
//
// Copyright Peter Andrews 2018 @ CSHL
//

#include <unistd.h>

#include <algorithm>
#include <exception>
#include <iostream>
#include <fstream>
#include <limits>
#include <sstream>
#include <set>
#include <string>
#include <unordered_map>
#include <unordered_set>
#include <utility>
#include <vector>

#include "error.h"
#include "utility.h"

namespace paa {

using Index = uint64_t;

template <class String>
class StringView {
 public:
  explicit StringView(const String & string__) :
      string_{string__}, start_{0}, size_{string_.size()} { }
  StringView(const StringView & string) = delete;
  // explicit StringView(const StringView & string__) :
  //    string_{string__.string_}, start_{0}, size_{string_.size()} { }

  StringView(const String & string__, const Index start__) :
      string_{string__}, start_{start__}, size_{string_.size() - start_} { }
  StringView(const StringView & string__, const Index start__) :
      string_{string__.string_},
    start_{start__ + string__.start_}, size_{string_.size() - start_} { }

  StringView(const String & string__, const Index start__, const Index size__) :
      string_{string__}, start_{start__}, size_{size__} { }
  StringView(const StringView & string__,
             const Index start__, const Index size__) :
      string_{string__.string_},
    start_{start__ + string__.start_}, size_{size__} { }

  Index size() const { return size_; }
  char operator[](const Index index) const {
    return string_[start_ + index];
  }
  template <class String2>
  bool operator!=(const String2 & rhs) const {
    for (Index i{0}; i != size_; ++i) {
      if ((*this)[i] != rhs[i]) return true;
    }
    return false;
  }

 private:
  const String & string_;
  Index start_{0};
  Index size_{0};
};

template<class String>
std::ostream & operator<<(std::ostream & out,
                          const StringView<String> & string) {
  for (Index index{0}; index != string.size(); ++index) {
    out << string[index];
  }
  return out;
}

template<class String>
class RCView {
 public:
  explicit RCView(const String & string__,
                  const Index start__, const Index size__,
                  const bool rc__ = true) :
      string_{string__}, start_{start__}, size_{size__}, rc_{rc__} { }

  bool rc() const { return rc_; }
  void rc(const bool rc__) { rc_ = rc__; }
  void flip() { rc_ = !rc_; }
  Index size() const { return size_; }
  char operator[](const Index index) const {
    return rc_ ?
        complement(string_[start_ + size_ - index - 1]) :
        string_[start_ + index];
  }
  template <class String2>
  bool operator!=(const String2 & rhs) const {
    for (Index i{0}; i != size(); ++i) {
      if ((*this)[i] != rhs[i]) return true;
    }
    return false;
  }
  static void test();

 private:
  static char complement(const char c) {
    switch (c) {
      case 'A':
        return 'T';
      case 'C':
        return 'G';
      case 'G':
        return 'C';
      case 'T':
        return 'A';
      default:
        throw Error("Unknown base in reverse complement");
    }
  }
  const String & string_;
  Index start_{0};
  Index size_{0};
  bool rc_;
};

template<class String>
std::ostream & operator<<(std::ostream & out,
                          const RCView<String> & string) {
  for (Index index{0}; index != string.size(); ++index) {
    out << string[index];
  }
  return out;
}

template<class String>
void RCView<String>::test() {
  std::cout << "RCView test" << std::endl;
  const std::string long_bases{"ACGTA"};
  const std::string bases{long_bases.substr(1, 4)};
  RCView<std::string> rc_bases{long_bases, 1, 4};
  std::cout << bases << " -> " << rc_bases << std::endl;
  rc_bases.flip();
  std::cout << bases << " == " << rc_bases << std::endl;
}

template<class String>
bool operator!=(const std::string & lhs,
                const StringView<String> & rhs) {
  return rhs != lhs;
}

constexpr Index c_str_length(const char * const str) {
  return *str ? 1UL + c_str_length(str + 1UL) : 0UL;
}

constexpr char c_bases[]{"ACGT"};
constexpr char c_lc_bases[]{"acgt"};
constexpr char c_letters[]{"abcdefg"};

constexpr Index bits_in_uint(const Index uint) {
  return uint ? 1 + bits_in_uint(uint >> 1) : 0;
}

template <class Word>
void show_binary(const Word word) {
  constexpr Word n_bits_word{8 * sizeof(Word)};
  for (unsigned int i{0}; i != n_bits_word; ++i) {
    std::cout << ((word >> (n_bits_word - i - 1)) & 1);
  }
  std::cout << std::endl;
}

template <const char * alphabet>
class CharacterEncoding {
 public:
  using Encoded = uint64_t;
  static constexpr Encoded size{c_str_length(alphabet)};
  static constexpr Encoded zero{0};
  static constexpr Encoded one{zero + 1};
  static constexpr Encoded all_ones{~zero};
  static constexpr Encoded n_chars{one << 8};
  static constexpr Encoded bits_per_char{bits_in_uint(size - 1)};
  static constexpr Encoded bits_per_encoded{sizeof(Encoded) * 8};
  static constexpr Encoded bits_after_encoded{bits_per_encoded - bits_per_char};
  static constexpr Encoded read_mask{all_ones >> bits_after_encoded};
  static constexpr Encoded bad_input{all_ones};

  explicit constexpr CharacterEncoding(const bool) {}
  CharacterEncoding() {
    for (unsigned int c{0}; c != n_chars; ++c) {
      const Encoded found{static_cast<Encoded>(
          std::find(alphabet, alphabet + size, c) - alphabet)};
      lookup[c] = found == size ? bad_input : found;
    }
  }

  Encoded operator()(const char c) const {
    return lookup[static_cast<uint8_t>(c)];
  }
  char operator()(const Encoded index) const {
    return alphabet[index];
  }

  static void test() {
    std::cout << "CharacterEncoding test" << std::endl;
    constexpr CharacterEncoding<alphabet> empty_encoding{true};
    if (false) std::cerr << empty_encoding('a');
    const CharacterEncoding<alphabet> base_encoding{};
    std::cout << alphabet
              << " " << size
              << " " << n_chars
              << " " << bits_per_char
              << " " << read_mask
              << " " << zero
              << " " << all_ones
              << std::endl;
    for (const char * pc{alphabet}; *pc; ++pc) {
      const Encoded e{base_encoding(*pc)};
      const char d{base_encoding(e)};
      if (*pc != d) throw Error("Roundtrip decoding failed") << *pc << d;
      std::cout << *pc << " " << e << " " << d << " "
                << static_cast<Encoded>(d) << std::endl;
    }
    if (alphabet == c_bases) {
      show_binary(zero);
      show_binary(one);
      show_binary(all_ones);
      show_binary(read_mask);
    }
  }

 private:
  Encoded lookup[n_chars]{};
};

template <const char * alphabet>
class CompressedString {
 public:
  using Encoding = CharacterEncoding<alphabet>;
  using View = StringView<CompressedString>;
  using String = std::string;
  using Word = uint64_t;
  using AddResult = std::pair<Index, Index>;

  static constexpr Index chars_per_word{
    sizeof(Word) * 8 / Encoding::bits_per_char};
  static constexpr Index bits_per_word{
    chars_per_word * Encoding::bits_per_char};
  static constexpr Index bad_index{Encoding::bad_input};

  explicit CompressedString(const Index bytes_initial_capacity = 256) :
      encoding{} {
    data.reserve(bytes_initial_capacity);
  }

  Index size() const { return size_; }
  char operator[](const Index index) const {
    return get_char(index);
  }
  View operator()(const Index index, const Index length) const {
    return View{*this, index, length};
  }

  Index add(const char c) {
    if (put_char(c)) {
      return size_ - 1;
    } else {
      return bad_index;
    }
  }

  // returns start index, n_chars added
  AddResult add(const String & input) {
    Index n_added{0};
    for (const char c : input) {
      if (put_char(c)) {
        ++n_added;
      } else {
        break;
      }
    }
    return {size_ - n_added, n_added};
  }

  bool in_alphabet(const char c) const {
    return encoding(c) != Encoding::bad_input;
  }

  void put_int(const Index char_index) {
    if (size_ * chars_per_word == data.size())
      data.resize(data.size() ? data.size() * 2 : 1);
    Word & saved_word{word(size_)};
    saved_word |= (char_index << shift(size_));
    ++size_;
  }

  void downsize(const Index new_size) {
    for (Index index{new_size}; index != size_; ++index) {
      // Can entire words be cleared?
      if (shift(index) == 0) {
        const Index start_word{word_index(index)};
        const Index stop_word{word_index(size_) + (shift(size_) ? 1 : 0)};
        for (Index wi{start_word}; wi != stop_word; ++wi) {
          data[wi] = 0;
        }
        break;
      }
      Word & saved_word{word(index)};
      saved_word &= ~(Encoding::read_mask << shift(index));
    }
    size_ = new_size;
  }

  static void test() {
    std::cout << "CompressedString test" << std::endl;
    using CompressedTest = CompressedString<c_bases>;
    CompressedTest test_{};
    std::cout << "Chars per word is "
              << CompressedTest::chars_per_word << std::endl;
    const std::vector<String> test_strings{"ACGTACACTG", "ACAGATTTTA"};
    for (const String & test_string : test_strings) {
      std::cout << "Add " << test_string << std::endl;
      const AddResult add_result{test_.add(test_string)};
      test_.show_data();
      if (add_result.second != test_string.size())
        throw Error("Could not add whole string")
            << test_string << add_result.second;
      std::cout << "Retrieve " << add_result.first << " "
                << add_result.second << std::endl;
      const View retrieved{test_, add_result.first, add_result.second};
      std::cout << retrieved << std::endl;
      if (test_string != retrieved)
        throw Error("Strings do not match") << test_string << retrieved;
    }
    std::cout << "String view test" << std::endl;
    const StringView<CompressedTest> view{test_, 5, 10};
    for (unsigned int i{0}; i != view.size(); ++i) {
      std::cout << view[i];
    }
    std::cout << std::endl;
  }

 private:
  void show_data() const {
    Index n_show{size_};
    for (Word w : data) {
      for (Index c{0}; c != chars_per_word; ++c) {
        if (c) std::cout << " ";
        const Word one_char{w & Encoding::read_mask};
        std::cout << one_char;
        w >>= Encoding::bits_per_char;
        if (--n_show == 0) break;
      }
    }
    std::cout << std::endl;
  }
  String to_string(const Index index, const Index length) const {
    String result{""};
    for (Index i{0}; i != length; ++i) {
      result += get_char(index + i);
    }
    return result;
  }
  Index word_index(const Index index) const {
    return index / chars_per_word;
  }
  Word word(const Index index) const {
    return data[word_index(index)];
  }
  Word & word(const Index index) {
    return data[word_index(index)];
  }
  Index shift(const Index index) const {
    return (index % chars_per_word) * Encoding::bits_per_char;
  }
  bool put_char(const char c) {
    const Index char_index{encoding(c)};
    if (char_index == Encoding::bad_input) return false;
    put_int(char_index);
    const char dc{get_char(size_ - 1)};
    if (dc != c) throw Error("Store error") << c << dc;
    return true;
  }
  char get_char(const Index index) const {
    return encoding((word(index) >> shift(index)) & Encoding::read_mask);
  }

  Encoding encoding;
  using Data = std::vector<Word>;
  Data data{};
  Index size_{0};
};


class LinksCounts {
 public:
  using Count = unsigned int;
  static constexpr Count zero{0};
  static constexpr Count one{zero + 1};
  static constexpr Count all_ones{~zero};
  static constexpr Count n_bits{sizeof(Count) * 8};
  static constexpr Count n_bits_dir{zero + 4};
  static constexpr Count n_bits_links{2 * n_bits_dir};
  static constexpr Count n_bits_count{n_bits - n_bits_links};
  static constexpr Count count_mask{all_ones >> n_bits_links};
  static constexpr Count links_mask{all_ones << n_bits_count};
  static constexpr Count out_links_mask{links_mask << n_bits_dir};
  static constexpr Count in_links_mask{out_links_mask >> n_bits_dir};

  LinksCounts & operator++() {
    if (count() != count_mask) ++data_;
    return *this;
  }
  Count count() const { return data_ & count_mask; }
  bool might_be_linked() const { return data_ & links_mask; }
  bool might_be_linked_out() const { return data_ & out_links_mask; }
  bool might_be_linked_in() const { return data_ & in_links_mask; }
  bool might_be_linked(const bool out, const unsigned int i) const {
    return (data_ >> (out * 4 + i + n_bits_count)) & one;
  }
  void set_unlinked(const bool out, const unsigned int i) {
    data_ &= ~(one << (out * 4 + i + n_bits_count));
  }
  static void test() {
    std::cout << "LinksCounts test" << std::endl;
    std::cout << n_bits
              << " " << n_bits_links
              << " " << n_bits_count
              << " " << count_mask
              << " " << bits_in_uint(count_mask)
              << " " << links_mask
              << " " << bits_in_uint(links_mask)
              << " " << (links_mask >> n_bits_count)
              << std::endl;
    LinksCounts test_value;
    ++++test_value;
    test_value.set_unlinked(1, 2);
    std::cout << test_value.count()
              << " " << test_value.data_
              << " " << test_value.might_be_linked(0, 0)
              << " " << test_value.might_be_linked(0, 1)
              << " " << test_value.might_be_linked(0, 2)
              << " " << test_value.might_be_linked(0, 3)
              << " " << test_value.might_be_linked(1, 0)
              << " " << test_value.might_be_linked(1, 1)
              << " " << test_value.might_be_linked(1, 2)
              << " " << test_value.might_be_linked(1, 3)
              << std::endl;
    show_binary(zero);
    show_binary(one);
    show_binary(all_ones);
    show_binary(count_mask);
    show_binary(links_mask);
    show_binary(in_links_mask);
    show_binary(out_links_mask);
  }

 private:
  Count data_{links_mask};
};

template <class Index>
class HashMapBase {
 public:
  using SequenceData = CompressedString<c_bases>;
  using Sequence = StringView<SequenceData>;
  using Node = std::pair<const Index, LinksCounts>;
  using HashMap = std::unordered_map<Index, LinksCounts>;
  using NodeIter = typename HashMap::iterator;
  using ConstNodeIter = typename HashMap::const_iterator;
  using NodeInsert = std::pair<NodeIter, bool>;

 private:
  SequenceData string{};
  HashMap nodes{};
};
#if 0
template <bool do_rc> class HashMap {};
template <> class HashMap<false> : public HashMapBase<Index> {
  Count count{0};
};
template <> class HashMap<true> : public HashMapBase<int64_t> {
  bool count{0};
};
#endif

template <class Count>
class CountsT {
 public:
  // Count operator[](const bool rc) const { return counts[rc]; }
  // Count & operator[](const bool rc) { return counts[rc]; }
  Count count() const { return count_; }
  Count & count() { return count_; }

 private:
  unsigned int count_{0};
  // Count counts[2]{0, 0};
};

using Sequence = std::string;

inline Sequence reverse_complement(const Sequence & input) {
  Sequence result;
  result.reserve(input.size());
  for (Sequence::const_reverse_iterator ci{input.rbegin()};
       ci != input.rend(); ++ci) {
    switch (*ci) {
      case 'A':
        result += 'T';
        break;
      case 'C':
        result += 'G';
        break;
      case 'G':
        result += 'C';
        break;
      case 'T':
        result += 'A';
        break;
      default:
        throw Error("Unknown base in reverse complement");
    }
  }
  return result;
}

#if 1
const Sequence bases{"ACGT"};
template <bool do_rc>
class BaseAssembler {
 public:
  using Count = unsigned int;
  using Counts = CountsT<Count>;
  using Node = std::pair<const Sequence, Counts>;
  using HashMap = std::unordered_map<Sequence, Counts>;
  using NodeIter = HashMap::iterator;
  using ConstNodeIter = HashMap::const_iterator;
  using NodeInsert = std::pair<NodeIter, bool>;
  using StoredSequence = std::pair<Sequence, bool>;
  using StoredSequences = std::vector<StoredSequence>;

  explicit BaseAssembler(const std::string & sequence_file_name,
                         const uint64_t max_lines_ = 0,
                         const uint64_t kmer_ = 30,
                         const uint64_t clip_ = 2) :
      max_lines{max_lines_},
    kmer{kmer_},
    km1{kmer - 1},
    clip{clip_} {
      std::cerr << "Creating graph " << std::endl;
      std::ifstream sequence_file{sequence_file_name};
      if (!sequence_file)
        throw Error("Could not open sequence file") << sequence_file_name;
      char c;
      Sequence sequence;
      while (sequence_file.get(c)) {
        if (c == '\n') {
          sequence.clear();
          if (++n_lines == max_lines) break;
          continue;
        }
        const uint64_t char_index{bases.find(c)};  // Can speed up find
        if (char_index == Sequence::npos) {
          sequence.clear();
          continue;
        }
        sequence += c;
        if (sequence.size() >= kmer) {
          const StoredSequence stored_result{get_canonical(sequence.substr(
              sequence.size() - kmer))};
          const Sequence & canonical_sequence{stored_result.first};
          Node * const this_node{&*nodes.emplace(
              canonical_sequence, Counts{}).first};
          ++this_node->second.count();
        }
      }
    }

  void do_clip() {
    // Clip graph nodes
    if (!clip) return;
    std::cerr << "Size before clip " << nodes.size() << std::endl;
    for (NodeIter node{nodes.begin()}; node != nodes.end();) {
      Count & count{node->second.count()};
      if (count <= clip) {
        node = nodes.erase(node);
      } else {
        ++node;
      }
    }
    std::cerr << "Size after clip " << nodes.size() << std::endl;
  }

  using EdgeCount = std::pair<const Count, const char>;
  using EdgeCounts = std::vector<EdgeCount>;
  EdgeCounts get_edge_counts(Sequence sequence, const bool out,
                             const bool quit_at_two = false) const {
    if (out) {
      sequence += ' ';
    } else {
      sequence = std::string(" ") + sequence;
    }
    if (sequence.size() != kmer)
      throw Error(std::string("Expected different sequence size in get_") +
                  (out ? "out" : "in")+ "_counts");
    EdgeCounts result;
    for (const char base : bases) {
      sequence[out ? sequence.size() - 1 : 0] = base;
      const StoredSequence stored_result{get_canonical(sequence)};
      const Sequence & canonical_sequence{stored_result.first};
      ConstNodeIter edge_node{nodes.find(canonical_sequence)};
      if (edge_node == nodes.end()) continue;
      // const bool stored_rc{stored_result.second};
      const Count matching_count{edge_node->second.count()};
      if (matching_count <= clip) continue;
      result.emplace_back(matching_count, base);
      if (quit_at_two && result.size() == 2) return result;
    }
    return result;
  }
  EdgeCounts get_out_counts(const Sequence & sequence) const {
    return get_edge_counts(sequence, true);
  }
  EdgeCounts get_in_counts(const Sequence & sequence) const {
    return get_edge_counts(sequence, false);
  }
  EdgeCounts up_to_two_out(const Sequence & sequence) const {
    return get_edge_counts(sequence, true, true);
  }
  EdgeCounts up_to_two_in(const Sequence & sequence) const {
    return get_edge_counts(sequence, false, true);
  }
  bool is_in_and_return_single(const Sequence & sequence) const {
    if (up_to_two_in(sequence).size() != 1 ||
        up_to_two_out(sequence).size() != 1) {
      return false;
    } else {
      return true;
    }
  }
  bool is_out_return_single(const Sequence & sequence) const {
    if (up_to_two_in(sequence).size() != 1) {
      return false;
    } else {
      return true;
    }
  }

  const StoredSequence get_canonical(const Sequence & seen_sequence) const {
    const Sequence reversed_sequence{reverse_complement(seen_sequence)};
    if (seen_sequence < reversed_sequence) {
      return {seen_sequence, false};
    } else {
      return {reversed_sequence, true};
    }
  }

  void do_join_nodes() {
    for (NodeIter node{nodes.begin()}; node != nodes.end();) {
    }
  }

  uint64_t hash(const Sequence & sequence) const {
    return std::hash<Sequence>{}(sequence);
  }

  using Seen = std::unordered_set<Sequence>;
  void output_run(const Sequence & sequence, Seen & seen,
                  const uint64_t run_size = 0,
                  uint64_t total_count = 0,
                  Count min_count = 0) const {
    // First see if sequence is good
    if (seen.count(sequence)) return;
    const StoredSequence stored_result{get_canonical(sequence)};
    const Sequence & canonical_sequence{stored_result.first};
    const ConstNodeIter node{nodes.find(canonical_sequence)};
    const Count count{node->second.count()};
    if (count <= clip) return;
    if (min_count == 0) min_count = count;
    min_count = count < min_count ? count : min_count;
    total_count += count;

    const Sequence prefix{sequence.substr(0, km1)};
    const EdgeCounts in_edges{up_to_two_in(prefix)};
    const EdgeCounts in_return_edges{up_to_two_out(prefix)};
    const bool is_in_single{in_edges.size() == 1 &&
          in_return_edges.size() == 1};
    if (!run_size) {
      if (is_in_single) return;
      std::cout << "Run: ";
    }
    seen.emplace(sequence);
    std::cout << sequence[0];

    const Sequence suffix{sequence.substr(1)};
    const EdgeCounts out_edges{up_to_two_out(suffix)};
    const EdgeCounts out_return_edges{up_to_two_in(suffix)};
    const bool is_out_single{out_edges.size() == 1 &&
          out_return_edges.size() == 1};
    if (is_out_single) {
      output_run(suffix + out_edges.front().second, seen, run_size + 1,
                 total_count, min_count);
    } else {
      std::cout << suffix << " " << run_size + suffix.size()
                << " " << total_count
                << " " << min_count << std::endl;
    }
  }

  void output_runs() const {
    Seen seen;
    for (const Node & node : nodes) {
      for (const bool rc : {false, true}) {
        if (node.second.count() <= clip) continue;
        const Sequence & canonical_sequence{node.first};
        Sequence sequence{rc ? reverse_complement(canonical_sequence) :
              canonical_sequence};
        output_run(sequence, seen);
      }
    }
  }

  const Node * output_dot(const Sequence & sequence,
                          std::ostream & out, Seen & seen,
                          const bool reversed = false) const {
    // Get node count
    const StoredSequence stored{get_canonical(sequence)};
    const Sequence & canonical{stored.first};
    const ConstNodeIter node{nodes.find(canonical)};
    if (node == nodes.end()) return nullptr;
    const Count count{node->second.count()};
    if (count <= clip) return nullptr;

    // Check if already processed
    if (!seen.emplace(canonical).second) return &*node;

    // Define node
    const uint64_t this_id{hash(canonical)};

    unsigned int n_right{100};
    for (const bool right : {true, false}) {
      const Sequence suffix{right ? sequence.substr(sequence.size() - km1) :
            reverse_complement(sequence.substr(0, km1))};
      const bool next_reversed{right == reversed};
      for (const char c : bases) {
        const Sequence next_sequence{suffix + c};
        const Node * next_node{output_dot(next_sequence, out, seen,
                                          next_reversed)};
        if (!next_node) continue;
        // if (right != reversed) ++n_right;
        const Sequence & next_canonical{next_node->first};
        if (canonical > next_canonical) continue;
        // Show link
        if (next_reversed) {
          out << "\"" << hash(next_canonical) << "\" -> \"" << this_id << '"';
        } else {
          out << "\"" << this_id << "\" -> \"" << hash(next_canonical) << '"';
        }
        out << ";" << std::endl;
      }
    }

    out << "\"" << this_id << "\"";
    out << " [label=\""
        << (n_right ? (Sequence() + sequence.front()) : sequence)
        << "\"];" << std::endl;
    return &*node;
  }

  void output_dot(const std::string & name) const {
    const std::string dot_name{name + ".dot"};
    std::ofstream out{dot_name.c_str()};
    if (!out) throw Error("Could not open dot file for writing") << dot_name;
    out << "digraph pseudogene {" << std::endl;
    out << "rankdir=LR;" << std::endl;
    out << "node [fontname=Courier, shape=box];" << std::endl;
    out << "edge [";
    out << "color=blue, ";
    out << "];" << std::endl;
    Seen seen;
    for (const Node & node : nodes) output_dot(node.first, out, seen);
    out << "}" << std::endl;
  }

  void create_graph(std::string name, const bool show = true) const {
    if (nodes.size() > 100000) {
      std::cerr << "Too many nodes to render " << nodes.size() << std::endl;
      return;
    }
    std::ostringstream suffix{};
    suffix << "-" << max_lines << "-" << kmer << "-" << clip;
    name += suffix.str();
    std::cerr << "Outputting graph " << name << std::endl;
    output_dot(name);
    std::ostringstream render;
    std::vector<std::string> renderers{"neato"};
    // renderers.push_back("dot");
    // renderers.push_back("fdp");
    std::vector<std::string> pdf_names;
    const std::string dot_name{name + ".dot"};
    const std::string pdf_name{name + ".pdf"};
    for (const std::string & renderer : renderers) {
      std::string pdf_type_name{name};
      if (renderers.size() > 1) {
        pdf_type_name += ".";
        pdf_type_name += renderer;
      }
      render << "echo Rendering graph " << renderer << " 1>&2 ; ";
      pdf_type_name += ".pdf";
      render << renderer << " -Tpdf -o "
             << pdf_type_name << " " << dot_name
             << " ;";
      if (show) {
        // render << "echo Displaying graph " << renderer << " ; ";
        if (0) render << "xpdf -z "
                      << (renderer == "dot" ? "50%" : "page")
                      << " -g 1200x800+0+0 " << pdf_type_name << "& ";
      }
      pdf_names.push_back(std::move(pdf_type_name));
    }
    // render << "echo Waiting for exit ; wait";
    if (system(render.str().c_str()) == -1) {
      std::cerr << "Problem rendering graph" << std::endl;
    }
    unlink(dot_name.c_str());
  }

 private:
  uint64_t max_lines{0};
  uint64_t n_lines{0};
  HashMap nodes{};
  const uint64_t kmer;
  const uint64_t km1;

 public:
  uint64_t clip;
};
#endif
}  // namespace paa

using std::cerr;
using std::endl;
using std::exception;
using paa::Error;

int main(int argc, char* argv[]) try {
  if (false) {
    paa::CharacterEncoding<paa::c_bases>::test();
    paa::CharacterEncoding<paa::c_lc_bases>::test();
    paa::CharacterEncoding<paa::c_letters>::test();
    paa::CompressedString<paa::c_bases>::test();
    paa::RCView<std::string>::test();
    paa::LinksCounts::test();
    return 0;
  }
  if (--argc < 5)
    throw Error("usage: debruijn out_name sequences_file n kmer clip ...");

  // Output name
  std::string out_name{argv[1]};

  // Input sequence file
  std::string sequences_file{argv[2]};

  // Max lines to read from file (0 unlimited)
  const uint64_t max_lines{static_cast<uint64_t>(atol(argv[3]))};

  // Kmer choice
  const unsigned int kmer{static_cast<unsigned int>(atoi(argv[4]))};

  // Clipping
  const unsigned int clip{static_cast<unsigned int>(atoi(argv[5]))};

  // Assemble
  paa::BaseAssembler<true> assembler{sequences_file, max_lines, kmer, clip};

  // Output
  argc -= 4;
  argv += 4;
  while (argc--) {
    const unsigned int output_clip{static_cast<unsigned int>(atoi(argv++[1]))};
    assembler.clip = output_clip;
    assembler.do_clip();
    assembler.output_runs();
    assembler.create_graph(out_name);
  }

  return 0;
} catch (Error & e) {
  cerr << "paa::Error:" << endl;
  cerr << e.what() << endl;
  return 1;
} catch (exception & e) {
  cerr << "std::exception" << endl;
  cerr << e.what() << endl;
  return 1;
} catch (...) {
  cerr << "unknown exception was caught" << endl;
  return 1;
}
```
