## Supplementary material for "G-Graph: An interactive genomic graph viewer": MUMdex Source Code for G-Graph: averages.cpp

```
//
// averages
//
// calculate averages over named columns of data
//
// Copyright 2018 Peter Andrews @ CSHL
//

#include <exception>
#include <iostream>
#include <sstream>
#include <string>
#include <vector>

#include "error.h"

using std::cerr;
using std::cin;
using std::cout;
using std::endl;
using std::exception;
using std::istringstream;
using std::string;
using std::vector;

using paa::Error;

int main(int argc, char**)  try {
  --argc;
  if (argc != 0) throw Error("data_file | averages");

  string line;
  getline(cin, line);
  istringstream header_stream{line.c_str()};
  vector<string> header;
  while (header_stream >> line) header.push_back(line);
  vector<double> data(header.size());
  uint64_t n{0};
  while (getline(cin, line)) {
    istringstream line_stream{line.c_str()};
    for (double & val : data) {
      double input;
      line_stream >> input;
      if (!line_stream) throw Error("Bad input from line") << n << line;
      val += input;
    }
    ++n;
  }
  for (double & val : data) val /= n;
  cout << "name";
  for (const string & name : header) cout << "\t" << name;
  cout << "\naverage";
  for (double & val : data) cout << "\t" << val;
  cout << "\n";

  return 0;
} catch (Error & e) {
  cerr << "paa::Error:" << endl;
  cerr << e.what() << endl;
  return 1;
} catch (exception & e) {
  cerr << "std::exception" << endl;
  cerr << e.what() << endl;
  return 1;
} catch (...) {
  cerr << "unknown exception was caught" << endl;
  return 1;
}
```
