## Supplementary material for "G-Graph: An interactive genomic graph viewer": MUMdex Source Code for G-Graph: balance_samples.sh

```
#! /bin/bash

while true ; do

    # Get space available for each node
    for node in wigclust{1..24} ; do
        echo $node $(df -k /mnt/$node/data |
            grep -v Available | awk '{print $4}')
    done | sort -k2n > space.txt

    # Find node with least space and data and return smallest data dir there
    for node in $(cat space.txt | awk '{print $1}') ; do
        smallest_dir=$(du -ks /mnt/$node/data/unsafe/paa/mums/wg-samples/*/ \
            2> /dev/null | sort -k1n | head -n 1)
        if [ "$smallest_dir" != "" ] ; then
            least_node=$node
            least_space=$(cat space.txt | grep "$node " | awk '{print $2}')
            move_size=$(echo $smallest_dir | awk '{print $1}')
            move_dir=$(echo $smallest_dir | awk '{print $2}')
            break
        fi
    done

    # Node with the most space
    most_node=$(tail -n 1 space.txt | awk '{print $1}')
    most_space=$(tail -n 1 space.txt | awk '{print $2}')

    # Reality checks
    if [ ! -e "$move_dir" ] || [ "$move_dir" = "" ] ; then
        echo could not find move dir $move_dir
        exit 1
    fi
    if [ "$least_space" = "" ] ; then
        echo could not find node with least space
        exit 1
    fi
    if [ "$most_node" = "$least_node" ] ; then
        echo most and least nodes are the same
        exit 1
    fi
    
    # Move if not balanced already
    if [ $((most_space - least_space)) -gt $((2 * move_size)) ] ; then
        sleep 10 # just in case a move in progress
        move_dir=${move_dir%/}
        sample=${move_dir##*/}
        if [ "$sample" = "" ] ; then
            echo sample undefined
            exit 1
        fi
        echo moving $sample at $move_dir $move_size from $least_node $least_space to $most_node $most_space
        to_dir=/mnt/$most_node/data/unsafe/paa/mums/wg-samples/$sample
        link_dir=/data/safe/paa/analysis/mums/wg-output/samples/$sample
        rm -Rf $to_dir &&
        mv $move_dir $to_dir &&
        rm -f $link_dir &&
        ln -sf $to_dir $link_dir
    else
        echo space appears well balanced
        cat space.txt
        exit 0
    fi

done

rm space.txt
exit 0
```
