## Supplementary material for "G-Graph: An interactive genomic graph viewer": MUMdex Source Code for G-Graph: bam_genotyper.cpp

```
//
// bam_genotyper.cpp
//
// returns genotype from bam file
//
// Copyright 2016 Peter Andrews @ CSHL
//

#include <exception>
#include <iostream>
#include <sstream>
#include <string>
#include <map>
#include <utility>
#include <vector>

#include "pstream.h"

#include "error.h"
#include "mumdex.h"
#include "utility.h"

using std::cerr;
using std::cout;
using std::endl;
using std::exception;
using std::istringstream;
using std::pair;
using std::string;
using std::map;
using std::vector;

using redi::ipstream;

using paa::serr;
using paa::sout;
using paa::ChromosomeIndexLookup;
using paa::Error;
using paa::Reference;

// Output and filter settings
const unsigned int min_count{1};  // minimum base count to output position
const unsigned int min_mapq{20};  // check if reasonable for mapper used
const char min_baseq{20 + '!'};   // Illimuna 1.8+ quality = phred + 33

// Holds essential read information
struct ReadData {
  unsigned int pos{};
  string cigar{};
  string bases{};
  string errors{};
};

// One base and quality score for one position
struct BaseQual {
  BaseQual(const char base_, const char qual_) :
      base{base_}, qual{qual_} { }
  char base{};
  char qual{};
};

// Processes a read or two reads based on cigar string
// and gets consensus for overlapping mates
class Consensus {
 public:
  void process(const ReadData & data, const bool clear = true) {
    if (clear) positions.clear();
    unsigned int pos{data.pos};
    unsigned int base{0};
    istringstream cigar{data.cigar.c_str()};
    unsigned int cigar_bases;
    char cigar_char;
    while (cigar >> cigar_bases >> cigar_char) {
      switch (cigar_char) {
        case 'S':
          base += cigar_bases;
          break;
        case 'M':
        case '=':
        case 'X':
          n_bases_ += cigar_bases;
          for (unsigned int b{0}; b != cigar_bases; ++b, ++base, ++pos) {
            const BaseQual base_qual{data.bases[base], data.errors[base]};
            const pair<unsigned int, BaseQual> pos_base{pos, base_qual};
            const auto stored = positions.insert(pos_base);
            // If insertion failed, position is already present in map
            if (stored.second == false) {
              ++n_overlap_;
              BaseQual & stored_base_qual = stored.first->second;
              if (stored_base_qual.qual < base_qual.qual) {
                stored_base_qual.base = base_qual.base;
                stored_base_qual.qual = base_qual.qual;
              } else if (stored_base_qual.qual == base_qual.qual &&
                         stored_base_qual.base != base_qual.base) {
                stored_base_qual.base = 'X';
              }
            }
          }
          break;
        case 'D':
        case 'N':
          pos += cigar_bases;
          break;
        case 'I':
          base += cigar_bases;
          break;
        case 'H':
          break;
        default:
          throw Error("Unknown cigar character") << cigar_char;
      }
    }
    if (base != data.bases.size()) {
      throw Error("Wrong number of bases read in cigar") << cigar.str();
    }
  }

  void process(const ReadData & data1, const ReadData & data2) {
    process(data1);
    process(data2, false);
  }

  map<unsigned int, BaseQual>::const_iterator begin() {
    return positions.begin();
  }
  map<unsigned int, BaseQual>::const_iterator end() {
    return positions.end();
  }
  uint64_t n_overlap() const { return n_overlap_; }
  uint64_t n_bases() const { return n_bases_; }

 private:
  map<unsigned int, BaseQual> positions{};
  uint64_t n_overlap_{0};
  uint64_t n_bases_{0};
};

// Reads bam using samtools, outputs Consensus for single or paired reads
class BamReader {
 public:
  BamReader(const string & bam_name, const string & chromosome) :
      sam{string("samtools view ") + bam_name + " " + chromosome} { }

  bool operator>>(Consensus & consensus) {
    pair<string, ReadData> read;
    string & name = read.first;
    ReadData & data = read.second;
    unsigned int flag;
    string chr;
    unsigned int mapq;
    string mchr;
    unsigned int mpos;
    unsigned int tlen;
    string optional;
    while (sam && (sam >> name >> flag >> chr >> data.pos
                   >> mapq >> data.cigar >> mchr >> mpos >> tlen
                   >> data.bases >> data.errors)) {
      // Ignore optional sam fields
      sam.ignore(10000, '\n');
      data.pos -= 1;
      ++all_reads_;

      // Filter reads
      if (mapq < min_mapq ||  // low map quality
          flag & 0x4 ||       // unmapped
          flag & 0x100 ||     // secondary alignment
          flag & 0x200 ||     // bad vendor quality
          flag & 0x400 ||     // PCR or optical dupe
          flag & 0x800) {     // supplementary alignment
        continue;
      }
      ++n_reads_;

      // Base quality cutoff - just transform base to Q
      for (unsigned int b{0}; b != data.errors.size(); ++b) {
        if (data.errors[b] < min_baseq) {
          data.bases[b] = 'Q';
        }
      }

      // Decide whether to output now or to look for mate first
      if ((mchr != "=" && mchr != chr) ||  // mate on different chromosome
          flag & 0x8) {                    // mate unmapped
        consensus.process(data);
        ++n_joined_;
        return true;
      }

      // Try to insert into map for later processing with mate
      // If does not insert, mate already seen so report both
      const auto stored = reads.insert(read);
      if (stored.second == false) {
        const auto & other_data = stored.first->second;
        consensus.process(data, other_data);
        reads.erase(stored.first);
        ++n_joined_;
        return true;
      }
    }
    if (reads.empty()) {
      return false;
    }
    consensus.process(reads.begin()->second);
    reads.erase(reads.begin());
    ++n_joined_;
    return true;
  }

  uint64_t all_reads() const { return all_reads_; }
  uint64_t n_reads() const { return n_reads_; }
  uint64_t n_joined() const { return n_joined_; }

 private:
  ipstream sam;
  map<string, ReadData> reads{};
  uint64_t all_reads_{0};
  uint64_t n_reads_{0};
  uint64_t n_joined_{0};
};

inline unsigned int base_to_int(const char base) {
  if (base == 'A') return 0;
  if (base == 'C') return 1;
  if (base == 'G') return 2;
  if (base == 'T') return 3;
  if (base == 'N') return 4;
  if (base == 'X') return 5;
  if (base == 'Q') return 6;
  throw Error("Unexpected base") << base;
}

int main(int argc, char ** argv) try {
  const time_t start_time{time(nullptr)};

  // Check for proper command line arguments
  --argc;
  if (argc != 2 && argc != 3) throw Error("usage: bam_genotyper ref bam [chr]");

  // Read command line arguments
  const Reference reference{argv[1]};
  const string bam_name{argv[2]};

  // Get start and stop chromosome range
  const ChromosomeIndexLookup chr_lookup{reference};
  const string chr_arg{argc == 3 ? argv[3] : ""};
  const unsigned int start_chr{argc == 3 ? chr_lookup[chr_arg]: 0};
  const unsigned int stop_chr{argc == 3 ? chr_lookup[chr_arg] + 1 :
        reference.n_chromosomes()};

  // Header
  const string bases{"ACGTNXQ"};
  sout << "chr pos ref";
  for (unsigned int base{0}; base != bases.size(); ++base) {
    sout << "n_";
    cout << bases[base];
  }
  sout << endl;

  // Loop over chromosomes
  Consensus consensus;
  uint64_t n_reads{0};
  for (unsigned int chr{start_chr}; chr != stop_chr; ++chr) {
    const string & chr_name{reference.name(chr)};
    const unsigned int chr_len{reference.size(chr)};

    // Base counts vector - avoid repeated memory allocs
    static vector<vector<unsigned int>> counts(bases.size(),
                                               vector<unsigned int>(chr_len));
    for (unsigned int base{0}; base != bases.size(); ++base) {
      counts[base].clear();
      counts[base].resize(chr_len);
    }

    // Custom bam reader object
    BamReader bam{bam_name, chr_name};

    // Process count of all pairs
    while (bam >> consensus) {
      for (const pair<unsigned int, BaseQual> & pos_base_qual : consensus) {
        ++counts[base_to_int(pos_base_qual.second.base)][pos_base_qual.first];
      }
    }

    // Report counts for all positions exceeding min_count total for A C G T
    uint64_t total_good_count{0};
    uint64_t total_count{0};
    for (unsigned int pos{0}; pos != chr_len; ++pos) {
      const unsigned int total_base_count{
        counts[0][pos] + counts[1][pos] + counts[2][pos] + counts[3][pos]};
      total_good_count += total_base_count;
      total_count += total_base_count +
          counts[4][pos] + counts[5][pos] + counts[6][pos];
      if (total_base_count >= min_count) {
        sout << chr_name << pos + 1 << reference[chr][pos];
        if (total_base_count) {
          cout << " ";
          for (unsigned int base{0}; base != 4; ++base) {
            if (counts[base][pos]) {
              cout << bases[base];
            }
          }
        } else {
          sout << "X";
        }
        for (unsigned int base{0}; base != bases.size(); ++base) {
          sout << counts[base][pos];
        }
        sout << endl;
      }
    }

    // Report chromosome stats
    serr << "Chr" << chr_name
         << "joined" << bam.n_joined()
         << "good" << bam.n_reads()
         << "total" << bam.all_reads()
         << "used" << total_good_count
         << "counts" << total_count
         << "overlaps" << consensus.n_overlap()
         << "bases" << consensus.n_bases()
         << "consensus" << consensus.n_bases() - consensus.n_overlap()
         << endl;
    n_reads += bam.all_reads();
  }

  // Report timing
  const time_t stop_time{time(nullptr)};
  const int64_t elapsed{stop_time - start_time};
  serr << "Processed" << n_reads << "in" << elapsed << "seconds at"
       << n_reads / elapsed  << "reads per second" << endl;
  cerr << "All done" << endl;

  return 0;
} catch (Error & e) {
  cerr << "paa::Error:" << endl;
  cerr << e.what() << endl;
  return 1;
} catch (exception & e) {
  cerr << "std::exception" << endl;
  cerr << e.what() << endl;
  return 1;
} catch (...) {
  cerr << "unknown exception was caught" << endl;
  return 1;
}
```
