## Supplementary material for "G-Graph: An interactive genomic graph viewer": MUMdex Source Code for G-Graph: bin_events.cpp

```
//
// mumdex_cn.cpp
//
// Copy number from a mumdex file
//
// Copyright 2016 Peter Andrews @ CSHL
//

#include <algorithm>
#include <exception>
#include <iostream>
#include <memory>
#include <random>
#include <sstream>
#include <string>
#include <vector>

#include "cn.h"
#include "error.h"
#include "genes.h"
#include "mumdex.h"
#include "psplot.h"

using std::cin;
using std::cerr;
using std::endl;
using std::exception;
using std::istringstream;
using std::make_unique;
using std::mt19937_64;
using std::random_device;
using std::string;
using std::uniform_real_distribution;
using std::unique_ptr;
using std::vector;

using paa::Bin;
using paa::ChromosomeIndexLookup;
using paa::CN_abspos;
using paa::Error;
using paa::Gaps;
using paa::Marker;
using paa::PSCNGraph;
using paa::PSDoc;
using paa::PSXYSeries;

using paa::Reference;

int main(int argc, char * argv[]) try {
  paa::exit_on_pipe_close();

  --argc;
  if (argc != 3) throw Error("usage: bin_events ref bins title");

  const Reference ref{argv[1]};
  const ChromosomeIndexLookup lookup{ref};
  const CN_abspos cn_abspos{ref};
  const string bins_name{argv[2]};
  const vector<Bin> bins{load_bins(bins_name, ref)};
  const string title{argv[3]};

  vector<unsigned int> counts(bins.size());
  string line;
  while (getline(cin, line)) {
    istringstream event{line.c_str()};
    string chr_name;
    unsigned int pos;
    unsigned int high;
    for (const bool anchor2 : {false, true}) {
      if (0) cerr << anchor2;
      event >> chr_name >> pos >> high;
      const unsigned int chr{lookup[chr_name]};
      const unsigned int abspos{cn_abspos(chr, pos)};
      if (abspos >= cn_abspos.bad_abspos()) continue;
      vector<Bin>::const_iterator found{
        upper_bound(bins.begin(), bins.end(), abspos)};
      if (found-- != bins.begin()) {
        if (abspos >= found->abspos_start() &&
            abspos < found->abspos_start() + found->length()) {
          ++counts[found - bins.begin()];
        }
      }
    }
  }

  const std::vector<unsigned int> used_chrs{[&bins]() {
      std::vector<unsigned int> result;
      for (unsigned int b{0}; b != bins.size(); ++b) {
        const Bin & bin{bins[b]};
        if (result.empty() || result.back() != bin.chromosome())
          result.push_back(bin.chromosome());
      }
      return result;
    }()};

  random_device rd;
  auto mersenne = mt19937_64(rd());
  uniform_real_distribution<double> udist{-0.25, 0.25};

  const Gaps gaps{ref.fasta_file() + ".bin/gaps.txt",
        lookup, "centromere,telomere"};

  const std::string dark{"1 0 0"};
  const Marker dark_circle_marker{paa::circle(), 0.4, dark, 0.2, false};
  PSDoc ps{title};
  ps.pdf(false);
  PSCNGraph counts_graph{ps, ref, used_chrs, title};
  counts_graph.y_label("Number of Events");
  PSXYSeries counts_series{counts_graph, dark_circle_marker};
  std::vector<std::unique_ptr<PSCNGraph>> chr_graphs(ref.n_chromosomes());
  std::vector<std::unique_ptr<PSXYSeries>> chr_series(ref.n_chromosomes());
  for (const unsigned int chr : used_chrs) {
    chr_graphs[chr] = std::make_unique<PSCNGraph>(
        ps, ref, chr, title);
    chr_graphs[chr]->y_label("Number of Events");
    add_gaps(*chr_graphs[chr], gaps, chr);
    chr_series[chr] = std::make_unique<PSXYSeries>(
        *chr_graphs[chr], dark_circle_marker);
  }
  for (unsigned int b{0}; b != bins.size(); ++b) {
    if (counts[b]) {
      const double count{counts[b] ?
            counts[b] + udist(mersenne) :
            0.5 + 0.1 * udist(mersenne)};
      counts_series.add_point(bins[b].abspos_start(), count);
      chr_series[bins[b].chromosome()]->add_point(
          bins[b].start_position(), count);
    }
  }

  cerr << "Done" << endl;

  return 0;
} catch (Error & e) {
  cerr << "paa::Error:" << endl;
  cerr << e.what() << endl;
  return 1;
} catch (exception & e) {
  cerr << "std::exception" << endl;
  cerr << e.what() << endl;
  return 1;
} catch (...) {
  cerr << "unknown exception was caught" << endl;
  return 1;
}
```
