## Supplementary material for "G-Graph: An interactive genomic graph viewer": MUMdex Source Code for G-Graph: bridge_counts.cpp

```
//
// bridge_counts
//
// look for a specific bridge in one or more samples
// output bridge and coverage counts
//
// Copyright 2017 Peter Andrews @ CSHL
//

#include <exception>
#include <iomanip>
#include <iostream>
#include <string>
#include <vector>

#include "bridges.h"
#include "error.h"
#include "fasta.h"
#include "mumdex.h"
#include "sequence.h"
#include "utility.h"

using std::cerr;
using std::cout;
using std::endl;
using std::exception;
using std::right;
using std::setw;
using std::string;
using std::vector;

using paa::sout;
using paa::Base;
using paa::Bridge;
using paa::Bridges;
using paa::ChromosomeIndexLookup;
using paa::Error;
using paa::Mappability;
using paa::MUM;
using paa::MUMdex;
using paa::Pair;
using paa::Reference;

int main(int argc, char* argv[])  try {
  // Check usage
  if (--argc < 9) {
    throw Error("usage: bridge_counts "
                "chr1 pos1 high1 chr2 pos2 high2 inv offset mumdex ...");
  }

  // Process command line arguments
  const string chr1s{argv[1]};
  const unsigned int pos1{static_cast<unsigned int>(atoi(argv[2]))};
  const bool high1{static_cast<bool>(atoi(argv[3]))};
  const string chr2s{argv[4]};
  const unsigned int pos2{static_cast<unsigned int>(atoi(argv[5]))};
  const bool high2{static_cast<bool>(atoi(argv[6]))};
  const int64_t invariant{atol(argv[7])};
  const int64_t offset{atol(argv[8])};
  argc -= 8;
  argv += 8;

  const vector<string> mumdex_names{[argv, argc]() {
      vector<string> result;
      for (int a{0}; a != argc; ++a) {
        result.push_back(argv[a + 1]);
      }
      return result;
    }()};

  const vector<MUMdex> mumdexes{[&mumdex_names]() {
      vector<MUMdex> result;
      result.reserve(mumdex_names.size());
      for (const string & name : mumdex_names) {
        result.emplace_back(name);
      }
      return result;
    }()};

  // Load reference information
  const Reference & ref{mumdexes.front().reference()};
  const ChromosomeIndexLookup chr_lookup{ref};
  const Mappability mappability{ref};
  const unsigned int chr1{chr_lookup[chr1s]};
  const unsigned int chr2{chr_lookup[chr2s]};

  cout << chr1s << " " << pos1 << " " << high1 << " "
       << chr2s << " " << pos2 << " " << high2 << " "
       << invariant << " " << offset;

  // const string chrs[2]{chr1s, chr2s};
  const unsigned int chr[2]{chr1, chr2};
  const unsigned int pos[2]{pos1, pos2};
  const unsigned int high[2]{high1, high2};
  const int npos[2]{static_cast<int>(pos1) + (high[0] ? 1 : -1),
        static_cast<int>(pos2) + (high[1] ? 1 : -1)};
  const unsigned int early{152};
  const unsigned int abspos[2]{ref.abspos(chr[0], pos[0]),
        ref.abspos(chr[1], pos[1])};
  const unsigned int map[2]{mappability.low_high(high[0], abspos[0]),
        mappability.low_high(high[1], abspos[1])};

  for (unsigned int m{0}; m != mumdexes.size(); ++m) {
    const string name{mumdex_names[m]};
    const MUMdex & mumdex{mumdexes[m]};

    // Count bridges
    unsigned int nb{0};  // bridge count
    for (const Bridge & bridge : Bridges{chr1, pos1, high1,
            chr2, pos2, high2, invariant, mumdex}) {
      // Bridge components
      const Pair pair{bridge.pair()};
      // const MUM mum1{bridge.mum1()};
      // const MUM mum2{bridge.mum2()};

      // Count non-dupes and long enough support anchors
      if (!pair.dupe()) {
        ++nb;
      }
    }
    cout << " " << nb;

    unsigned int tc{0};
    unsigned int ctc{0};
    unsigned int rtc{0};
    for (const bool a : {false, true}) {
      // Count coverage for each anchor
      const auto begin_index = mumdex.lower_bound(
          chr[a], pos[a] > early ? pos[a] - early : 0);
      const auto end_index = mumdex.lower_bound(
          chr[a], pos[a] + 1);
      unsigned int rc{0};
      unsigned int cc{0};
      unsigned int ccc{0};
      for (auto index = begin_index; index != end_index; ++index) {
        const auto mum = mumdex.mum(*index);
        const auto pair = mumdex.pair(*index);
        if (pair.dupe()) continue;
        const int read_length(pair.length(mum.read_2()));
        if (pos[a] >= mum.position0() &&
            pos[a] < mum.position0() + mum.length()) {
          ++cc;
          // Bridge limits in mum space
          const int bp[2]{static_cast<int>(pos[a]) + (high[a] ? -1 : 1) *
                static_cast<int>(map[a] - 1),
                static_cast<int>(pos[a]) + (high[a] ? 1 : -1) *
                static_cast<int>(offset + map[1 - a] - 1)};
          bool good{true};
          for (const bool e : {false, true}) {
            const int p{mum.position0_to_read(bp[e])};
            if (p < 0 || p >= read_length) {
              good = false;
            }
          }
          if (good) {
            ++ccc;
            // Is reference continuous at anchor position?
            if (npos[a] >= static_cast<int>(mum.position0()) &&
                npos[a] < static_cast<int>(mum.position0() + mum.length())) {
              ++rc;
            }
          }
        }
      }
      tc += cc;
      ctc += ccc;
      rtc += rc;
      cout << " " << rc << " " << cc << " " << ccc;
    }
    cout << " " << 2.0 * nb / tc << " " << 2.0 * nb / ctc
         << " " << nb / (nb + rtc / 2.0);
  }
  cout << endl;

  return 0;
} catch (Error & e) {
  cerr << "paa::Error:" << endl;
  cerr << e.what() << endl;
  return 1;
} catch (exception & e) {
  cerr << "std::exception" << endl;
  cerr << e.what() << endl;
  return 1;
} catch (...) {
  cerr << "unknown exception was caught" << endl;
  return 1;
}
```
