## Supplementary material for "G-Graph: An interactive genomic graph viewer": MUMdex Source Code for G-Graph: bridge_effect.cpp

```
//
// bridge_effect
//
// assess bridge for effect
//
// Copyright 2016 Peter Andrews @ CSHL
//

#include <array>
#include <exception>
#include <iostream>
#include <set>
#include <sstream>
#include <string>
#include <vector>

#include "bridges.h"
#include "error.h"
#include "fasta.h"
#include "genes.h"
#include "mumdex.h"
#include "population.h"
#include "sequence.h"
#include "utility.h"
#include "pstream.h"

using std::array;
using std::cerr;
using std::cout;
using std::endl;
using std::exception;
using std::ostringstream;
using std::right;
using std::set;
using std::string;
using std::vector;

using paa::saved_ref_name;
using paa::sout;
using paa::Base;
using paa::Bridge;
using paa::Bridges;
using paa::ChromosomeIndexLookup;
using paa::ConsensusSequence;
using paa::Error;
using paa::GeneXrefs;
using paa::KnownGenes;
using paa::MUMdex;
using paa::Population;
using paa::Reference;

using redi::pstream;

int main(int argc, char* argv[])  try {
  // Check usage
  if (--argc != 10) {
    throw Error("usage: find_bridges samples_dir pop_file "
                "chr1 pos1 high1 chr2 pos2 high2 inv sample");
  }

  const bool run_snp_command{true};
  const string tab{run_snp_command ? "\t" : "\\t"};

  // Process command line arguments
  const string samples_dir{argv[1]};
  const Population pop{argv[2]};
  const string chr1s{argv[3]};
  const unsigned int pos1{static_cast<unsigned int>(atoi(argv[4]))};
  const bool high1{static_cast<bool>(atoi(argv[5]))};
  const string chr2s{argv[6]};
  const unsigned int pos2{static_cast<unsigned int>(atoi(argv[7]))};
  const bool high2{static_cast<bool>(atoi(argv[8]))};
  const int64_t invariant{atol(argv[9])};
  argc -= 9;
  argv += 9;

  // Load reference information
  const string sample_name{argv[1]};
  const string mumdex_dir{samples_dir + "/" + sample_name + "/mumdex"};
  const MUMdex mumdex{mumdex_dir};
  const Reference & ref{mumdex.reference()};
  const ChromosomeIndexLookup chr_lookup{ref};
  const unsigned int chr1{chr_lookup[chr1s]};
  const unsigned int chr2{chr_lookup[chr2s]};

  const array<string, 2> chrs{{chr1s, chr2s}};
  const array<unsigned int, 2> chr{{chr1, chr2}};
  const array<unsigned int, 2> pos{{pos1, pos2}};

  if (chr1 == chr2 && high1 != high2 && invariant == 0 && !run_snp_command) {
    sout << "echo";
  }

  sout << chr1s << pos1 << high1 << chr2s << pos2 << high2 << invariant;

  if (chr1 == chr2 && high1 != high2 && invariant == 0) {
    // Deal with snp
    const array<unsigned int, 2> snp_pos{{pos1 + (high1 ? 1 : -1),
            pos2 + (high2 ? 1 : -1)}};

    // Consensus sequence adjacent to anchors
    const unsigned int adj_len{1};
    ConsensusSequence consensus[2]{adj_len, adj_len};

    // Loop over bridges found
    for (const Bridge & bridge :
             Bridges{chr1, pos1, high1, chr2, pos2, high2, invariant, mumdex}) {
      const array<string, 2> sequences = bridge.adjacent_sequences(adj_len);
      for (const bool anchor2 : {false, true}) {
        const string & sequence{sequences[anchor2]};
        consensus[anchor2].add(sequence);
      }
    }

    // Output consensus sequences
    for (const bool anchor2 : {false, true}) {
      const string & seq{consensus[anchor2].sequence()};
      const unsigned int snp_chr{chr[anchor2]};

      if (anchor2 && chr[0] == chr[1] && snp_pos[0] == snp_pos[1]) {
        break;
      }

      ostringstream annotate_command;
      const string pipeline_dir{"/mnt/wigclust8/data/safe/autism/SeqPipeline"};
      const string annotate_program{"python/DAE/tools/annotate_variants.py"};
      annotate_command << pipeline_dir << "/" << annotate_program
                       << " -H -c 1 -p 2 -r 3 -a 4 "
                       << " <(echo "
                       << chrs[anchor2].substr(3) << " "
                       << snp_pos[anchor2] + 1 << " "
                       << ref[snp_chr][snp_pos[anchor2]] << " "
                       << seq << " | perl -pe 's/ /" << tab
                       << "/g') 2> /dev/null"
                       << " | perl -ne 'print unless /^#/'"
                       << " | cut -f 5-"
                       << " | perl -pe 's/" << tab << "/ /g'";
      if (run_snp_command) {
        redi::ipstream annotate{annotate_command.str()};
        unsigned int n_lines{0};
        while (annotate) {
          string line;
          getline(annotate, line);
          if (line.size()) {
            if (++n_lines != 1) throw Error("Too many output lines");
            sout << ref[snp_chr][snp_pos[anchor2]];
            cout << "->" << seq;
            sout << line;
          }
        }
      } else {
        sout << " $(" << annotate_command.str() << ")";
      }
    }
  } else {
    // Load gene info
    const string reference_file{saved_ref_name(mumdex_dir)};
    const string genes_name{reference_file + ".bin/knownGene.txt"};
    const string isoforms_name{reference_file + ".bin/knownIsoforms.txt"};
    const string kgXrefs_name{reference_file + ".bin/kgXref.txt"};
    const KnownGenes genes{genes_name, isoforms_name, chr_lookup, ref};
    const GeneXrefs xref{kgXrefs_name};

    for (const bool anchor2 : {false, true}) {
      const vector<unsigned int> pos_genes{
        genes.find_genes(chr[anchor2], pos[anchor2])};
      bool in_exon{false};
      set<string> gene_names;
      for (const unsigned int gene : pos_genes) {
        gene_names.insert(xref[genes[gene].name].geneSymbol);
        if (genes[gene].in_exon(chr[anchor2], pos[anchor2])) {
          in_exon = true;
        }
      }
      if (in_exon) sout << "exon";
      unsigned int n{0};
      for (const string & name : gene_names) {
        if (n++) {
          cout << "," << name;
        } else {
          sout << name;
        }
      }
      if (gene_names.empty()) {
        sout << "intergenic";
      }
    }

    if (chr1 == chr2 && high1 != high2 && ((invariant % 3) == 0) &&
        labs(invariant) < 100) {
      sout << "short multiple of 3";
    }
  }
  sout << endl;

  return 0;
} catch (Error & e) {
  cerr << "paa::Error:" << endl;
  cerr << e.what() << endl;
  return 1;
} catch (exception & e) {
  cerr << "std::exception" << endl;
  cerr << e.what() << endl;
  return 1;
} catch (...) {
  cerr << "unknown exception was caught" << endl;
  return 1;
}
```
