## Supplementary material for "G-Graph: An interactive genomic graph viewer": MUMdex Source Code for G-Graph: bridge_gene2.cpp

```
//
// bridge_gene
//
// does bridge impact a gene?
//
// Copyright 2016 Peter Andrews @ CSHL
//

#include <algorithm>
#include <array>
#include <exception>
#include <iostream>
#include <map>
#include <set>
#include <sstream>
#include <string>
#include <utility>
#include <vector>

#include "bridges.h"
#include "error.h"
#include "fasta.h"
#include "genes.h"
#include "mumdex.h"
#include "population.h"
#include "sequence.h"
#include "utility.h"
#include "pstream.h"

using std::array;
using std::cerr;
using std::cin;
using std::cout;
using std::endl;
using std::exception;
using std::map;
using std::ostringstream;
using std::pair;
using std::right;
using std::set;
using std::string;
using std::vector;

using paa::saved_ref_name;
using paa::sout;
using paa::Base;
using paa::Bridge;
using paa::Bridges;
using paa::ChromosomeIndexLookup;
using paa::ConsensusSequence;
using paa::Error;
using paa::Family;
using paa::GeneXrefs;
using paa::KnownGenes;
using paa::MUM;
using paa::MUMdex;
using paa::Pair;
using paa::Population;
using paa::Reference;
using paa::Sample;

using redi::pstream;

int main(int argc, char* argv[])  try {
  // Check usage
  if (--argc != 1) {
    throw Error("usage: bridge_gene ref");
  }

  // Process command line arguments
  const Reference ref{argv[1]};
  const ChromosomeIndexLookup chr_lookup{ref};

  // sout << chr1s << pos1 << high1 << chr2s << pos2 << high2 << invariant;

  // Load gene info
  const string reference_file{argv[1]};
  const string genes_name{reference_file + ".bin/knownGene.txt"};
  const string isoforms_name{reference_file + ".bin/knownIsoforms.txt"};
  const string kgXrefs_name{reference_file + ".bin/kgXref.txt"};
  const KnownGenes genes{genes_name, isoforms_name, chr_lookup, ref};
  const GeneXrefs xref{kgXrefs_name};

  string chr1s;
  unsigned int pos1;
  unsigned int high1;
  string chr2s;
  unsigned int pos2;
  unsigned int high2;
  int invariant;

  while (cin >> chr1s >> pos1 >> high1 >> chr2s >> pos2 >> high2 >> invariant) {
    const unsigned int chr1{chr_lookup[chr1s]};
    const unsigned int chr2{chr_lookup[chr2s]};

    const array<string, 2> chrs{{chr1s, chr2s}};
    const array<unsigned int, 2> chr{{chr1, chr2}};
    const array<unsigned int, 2> pos{{pos1, pos2}};

    if (chr1 == chr2 && high1 != high2 && ((invariant % 3) == 0) &&
        abs(invariant) < 100) {
      sout << "1";
    } else {
      sout << "0";
    }

    // anchor genes
    for (const bool anchor2 : {false, true}) {
      const vector<unsigned int> pos_genes{
        genes.find_genes(chr[anchor2], pos[anchor2])};
      set<string> gene_names;
      for (const unsigned int gene : pos_genes) {
        const string symbol{xref[genes[gene].name].geneSymbol};
        if (genes[gene].in_exon(chr[anchor2], pos[anchor2])) {
          gene_names.insert("exon-" + symbol);
        } else {
          gene_names.insert(symbol);
        }
      }
      unsigned int n{0};
      for (const string & name : gene_names) {
        if (n++) {
          cout << "," << name;
        } else {
          sout << name;
        }
      }
      if (gene_names.empty()) {
        sout << "intergenic";
      }
    }

    // genes in range
    if (chr1 == chr2) {
      const unsigned int low_pos{std::min(pos1, pos2)};
      const unsigned int high_pos{std::max(pos1, pos2)};
      const KnownGenes::GeneOverlaps gene_overlaps{genes.find_genes(
          chr1, low_pos, high_pos)};
      map<string, unsigned int> named_overlaps;
      for (const KnownGenes::GeneOverlap & overlap : gene_overlaps) {
        const string symbol{xref[genes[overlap.first].name].geneSymbol};
        const unsigned int old_value{named_overlaps[symbol]};
        if (old_value < overlap.second) {
          named_overlaps[symbol] = overlap.second;
        }
      }
      using Overlap = pair<string, unsigned int>;
      vector<Overlap> overlaps(named_overlaps.begin(), named_overlaps.end());
      sort(overlaps.begin(), overlaps.end(),
           [](const Overlap & lhs, const Overlap & rhs) {
             if (lhs.second == rhs.second) {
               return lhs.first < rhs.first;
             } else {
               return lhs.second > rhs.second;
             }
           });
      unsigned int total_exons{0};
      for (unsigned int o{0}; o != overlaps.size(); ++o)
        total_exons += overlaps[o].second;
      cout << " " << overlaps.size() << " " << total_exons << " ";
      for (unsigned int o{0}; o != overlaps.size(); ++o) {
        const Overlap & overlap{overlaps[o]};
        if (o) cout << ",";
        if (overlap.second)
          cout << overlap.second << "-exon"
               << (overlap.second > 1 ? "s" : "") << "-";
        cout << overlap.first;
      }
      if (overlaps.empty()) {
        cout << "intergenic";
      }
    } else {
      sout << "0 0 intergenic";
    }
    sout << endl;
  }

  return 0;
} catch (Error & e) {
  cerr << "paa::Error:" << endl;
  cerr << e.what() << endl;
  return 1;
} catch (exception & e) {
  cerr << "std::exception" << endl;
  cerr << e.what() << endl;
  return 1;
} catch (...) {
  cerr << "unknown exception was caught" << endl;
  return 1;
}
```
