## Supplementary material for "G-Graph: An interactive genomic graph viewer": MUMdex Source Code for G-Graph: bridge_properties.cpp

```
//
// bridge_properties
//
// examine properties of bridges
//
// Copyright 2018 Peter Andrews @ CSHL
//

#include <algorithm>
#include <exception>
#include <iostream>
#include <string>

#include "bridges.h"
#include "error.h"
#include "files.h"
#include "mumdex.h"
#include "utility.h"

using std::cerr;
using std::cout;
using std::endl;
using std::exception;
using std::lower_bound;

using paa::BridgeInfo;
using paa::Error;
using paa::Mappability;
using paa::MappedVector;
using paa::Reference;

int main(int argc, char* argv[])  try {
  paa::exit_on_pipe_close();  // Handle pipe close signal properly

  if (argc != 3 && argc != 6) {
    throw Error("usage: bridge_properties ref bridge_file [chr start stop]");
  }

  const Reference ref{argv[1]};
  const Mappability map{ref};

  const MappedVector<BridgeInfo> bridges{argv[2]};

  MappedVector<BridgeInfo>::const_iterator begin{bridges.begin()};
  MappedVector<BridgeInfo>::const_iterator end{bridges.end()};

  if (argc == 6) {
    auto cmp = [](const BridgeInfo & lhs, const unsigned int lpos) {
      return lhs.pos1() < lpos;
    };
    begin = lower_bound(begin, end, atoi(argv[4]), cmp);
    end = lower_bound(begin, end, atoi(argv[5]), cmp);
  }

  cout << "chr\tpos\thigh\tchr2\tpos2\thigh2\tinv\tioff"
       << "\tal\tbl\taml\tbml\tbc\tamc\tbmc\tem1\tem2"
       << endl;
  for (MappedVector<BridgeInfo>::const_iterator bridge{begin};
       bridge != end ; ++bridge) {
    const unsigned int map1{map.low_high(bridge->high1(),
                                         ref.abspos(bridge->chr1(),
                                                    bridge->pos1()))};
    const unsigned int map2{map.low_high(bridge->high2(),
                                         ref.abspos(bridge->chr2(),
                                                    bridge->pos2()))};
    cout << ref.name(bridge->chr1())
         << '\t' << bridge->pos1()
         << '\t' << bridge->high1()
         << '\t' << ref.name(bridge->chr2())
         << '\t' << bridge->pos2()
         << '\t' << bridge->high2()
         << '\t' << bridge->invariant()
         << '\t' << bridge->offset()
         << '\t' << bridge->anchor1_length()
         << '\t' << bridge->anchor2_length()
         << '\t' << bridge->mate_anchor1_length()
         << '\t' << bridge->mate_anchor2_length()
         << '\t' << bridge->bridge_count()
         << '\t' << bridge->mate_anchor1_count()
         << '\t' << bridge->mate_anchor2_count()
         << '\t' << bridge->anchor1_length() - map1
         << '\t' << bridge->anchor2_length() - map2
         << '\n';
  }

  cerr << "All done" << endl;
  return 0;
} catch (Error & e) {
  cerr << "paa::Error:" << endl;
  cerr << e.what() << endl;
  return 1;
} catch (exception & e) {
  cerr << "std::exception" << endl;
  cerr << e.what() << endl;
  return 1;
} catch (...) {
  cerr << "unknown exception was caught" << endl;
  return 1;
}
```
