## Supplementary material for "G-Graph: An interactive genomic graph viewer": MUMdex Source Code for G-Graph: candidate_stats.cpp

```
//
// candidate_stats.cpp
//
// Copyright 2016 Peter Andrews @ CSHL
//

#include <algorithm>
#include <iostream>
#include <fstream>
#include <functional>
#include <memory>
#include <random>
#include <sstream>
#include <string>
#include <vector>

#include "error.h"
#include "psplot.h"
#include "utility.h"

using std::cout;
using std::cerr;
using std::endl;
using std::ifstream;
using std::istringstream;
using std::function;
using std::mt19937_64;
using std::random_device;
using std::string;
using std::uniform_real_distribution;
using std::vector;

using paa::sout;
using paa::Bounds;
using paa::Error;
using paa::Marker;
using paa::PSDoc;
using paa::PSPage;
using paa::PSGraph;
using paa::PSSeries;
using paa::PSXYSeries;
using paa::PSXYMSeries;

void parse_counts(const string & in_string, unsigned int vals[4]) {
  istringstream in{in_string.c_str()};
  for (const unsigned int i : {0, 1, 2, 3}) {
    in >> vals[i];
    in.get();
  }
}

class CandInfo {
 public:
  class ParseError {};
  explicit CandInfo(const string & line) {
    istringstream input{line.c_str()};
    string bridges_;
    string anchorsA_;
    string anchorsB_;
    string coveragesA_;
    string coveragesB_;
    input >> family >> parent >> kids >> nFam >> nSam >> nInFam
          >> chr >> chrA >> posA >> highA >> chrB >> posB >> highB
          >> type >> ori >> invariant >> offset >> bridge
          >> supA >> supB >> mCA >> mSA >> mCB >> mSB >> mapA >> mapB
          >> minParCov >> bridges_ >> anchorsA_ >> anchorsB_
          >> coveragesA_ >> coveragesB_
          >> pNP >> pNB >> pMed >> pMax >> tBC >> nBC >> nS;
    if (!input) {
      throw ParseError{};
    }
    parse_counts(bridges_, bridges);
    parse_counts(anchorsA_, anchorsA);
    parse_counts(anchorsB_, anchorsB);
    parse_counts(coveragesA_, coveragesA);
    parse_counts(coveragesB_, coveragesB);
  }
  string family{};
  string parent{};
  string kids{};
  unsigned int nFam{};
  unsigned int nSam{};
  unsigned int nInFam{};
  unsigned int chr{};
  string chrA{};
  unsigned int posA{};
  bool highA{};
  string chrB{};
  unsigned int posB{};
  bool highB{};
  string type{};
  char ori{};
  int invariant{};
  int offset{};
  unsigned int bridge{};
  unsigned int supA{};
  unsigned int supB{};
  unsigned int mCA{};
  unsigned int mSA{};
  unsigned int mCB{};
  unsigned int mSB{};
  unsigned int mapA{};
  unsigned int mapB{};
  unsigned int minParCov{};
  unsigned int bridges[4]{0, 0, 0, 0};
  unsigned int anchorsA[4]{0, 0, 0, 0};
  unsigned int anchorsB[4]{0, 0, 0, 0};
  unsigned int coveragesA[4]{0, 0, 0, 0};
  unsigned int coveragesB[4]{0, 0, 0, 0};
  unsigned int pNP{};
  unsigned int pNB{};
  unsigned int pMed{};
  unsigned int pMax{};
  unsigned int tBC{};
  unsigned int nBC{};
  unsigned int nS{};
};

int main(int argc, char ** argv)  try {
  // Check arguments
  if (--argc != 1)
    throw Error("usage: candidate_stats cand");

  // Input file
  const string cand_file_name{argv[1]};
  ifstream cand_file{cand_file_name.c_str()};
  if (!cand_file) throw Error("Could not open file") << cand_file_name;

  // Read Input file into candidate objects
  string line;
  vector<CandInfo> info;
  while (cand_file) {
    getline(cand_file, line);
    if (line.size()) {
      try {
        CandInfo i{line};
        info.push_back(i);
      } catch (CandInfo::ParseError & e) {
        if (line[0] != 'f' || line[1] != 'a') {
          cerr << "parse error for line: " << line << endl;
        }
      }
    }
  }

  // Randomization
  random_device rd;
  mt19937_64 mersenne{rd()};
  function<double()> unitGen{
    bind(uniform_real_distribution<double>(-0.5, 0.5), mersenne)};

  // Document and defaults
  const Marker dot{paa::circle(), 0.2, "0 0 0", 1, true};
  PSDoc ps{"stats", "Candidate Stats"};

  // Establish some bounds
  const double max_parent{1020};

  // Graphs and histograms to fill
  PSXYSeries pNP_vs_pNB{ps, "Population;"
        "Total Parent Bridge Count;Number of Parents With Bridge", dot};
  PSXYSeries pNP_vs_pNB_low{ps, "Population;"
        "Total Parent Bridge Count;Number of Parents With Bridge", dot,
        Bounds{0.0, 1000.0, 0.0, 100.0}};

  PSXYSeries pNP_vs_minParCov{ps, "Population;"
        "Minimum Parent Coverage;Number of Parents With Bridge", dot,
        Bounds{0.0, 200.0, 0.0, 1020.0}};

  PSXYSeries pMed_vs_pNP{ps, "Population;"
        "Number of Parents With Bridge;Median Parent Bridge Count", dot};
  PSXYSeries pMax_vs_pNP{ps, "Population;"
        "Number of Parents With Bridge;Maximum Parent Bridge Count (<100)",
        dot};
  PSXYSeries pMax_vs_bridge{ps, "Population;"
        "Candidate Bridge Count;Maximum Parent Bridge Count (<100)",
        dot};

  PSPage pop_page{ps, "Population", "1 2"};
  paa::PSHSeries<int, uint64_t> pNP{pop_page,
        "All;Number of Parents With Bridge;N",
        Bounds{0.0, max_parent}, 102};
  paa::PSHSeries<int, uint64_t> pNP_low{pop_page,
        "Low Range;Number of Parents With Bridge;N",
        Bounds{0.0, 100}, 100};

  paa::PSHSeries<int, uint64_t> bridge{ps,
        "Candidates;Bridge Count (<30);N",
        Bounds{0.0, 30.0}, 30};

  paa::PSHSeries<int, uint64_t> offset{ps,
        "Candidates;Anchor Offset;N",
        Bounds{-150.0, 50.0}, 200};
  PSPage offset_page{ps, "Candidates", "1 2"};
  paa::PSHSeries<int, uint64_t> offset_r{offset_page,
        "1 to 5 Parents;Anchor Offset;N",
        Bounds{-150.0, 50.0}, 200};
  paa::PSHSeries<int, uint64_t> offset_ur{offset_page,
        "One Family Only;Anchor Offset;N",
        Bounds{-150.0, 50.0}, 200};

  // Enter data to graphs and histograms
  for (const CandInfo & i : info) {
    pNP_vs_pNB.add_point(i.pNB + unitGen(), i.pNP + unitGen());
    pNP_vs_pNB_low.add_point(i.pNB + unitGen(), i.pNP + unitGen());
    pNP_vs_minParCov.add_point(i.minParCov + unitGen(), i.pNP + unitGen());
    pMed_vs_pNP.add_point(i.pNP + unitGen(), i.pMed + unitGen());
    if (i.pMax < 100) {
      pMax_vs_pNP.add_point(i.pNP + unitGen(), i.pMax + unitGen());
      pMax_vs_bridge.add_point(i.bridge + unitGen(), i.pMax + unitGen());
    }
    pNP.add_point(i.pNP);
    pNP_low.add_point(i.pNP);
    bridge.add_point(i.bridge);
    offset.add_point(i.offset);
    if (!i.pNP)
      offset_ur.add_point(i.offset);
    if (i.pNP && i.pNP <= 5)
      offset_r.add_point(i.offset);
  }

  std::cerr << "done" << endl;

  return 0;
} catch (Error & e) {
  cerr << "paa::Error:" << endl;
  cerr << e.what() << endl;
  return 1;
} catch(std::exception & e) {
  cerr << e.what() << '\n';
  return 1;
} catch(...) {
  cerr << "Some exception was caught.\n";
  return 1;
}
```
