## Supplementary material for "G-Graph: An interactive genomic graph viewer": MUMdex Source Code for G-Graph: check_bridges.cpp

```
//
// check_bridges
//
// look in a sample for many bridges
//
// Copyright 2016 Peter Andrews @ CSHL
//

#include <algorithm>
#include <exception>
#include <iostream>
#include <sstream>
#include <string>
#include <vector>

#include "bridges.h"
#include "error.h"
#include "files.h"
#include "mumdex.h"
#include "population.h"
#include "utility.h"

using std::cerr;
using std::cin;
using std::cout;
using std::endl;
using std::equal_range;
using std::exception;
using std::ostringstream;
using std::string;
using std::vector;

using paa::readable;
using paa::sout;
using paa::BridgeInfo;
using paa::ChromosomeIndexLookup;
using paa::Error;
using paa::Family;
using paa::FileVector;
using paa::MUM;
using paa::MUMindex;
using paa::OneBridgeInfo;
using paa::Pair;
using paa::Population;
using paa::Reference;
using paa::Sample;

using Bridges = paa::FileBridges;
using MUMdex = paa::FileMUMdex;
using Anchors = MUMdex::Anchors;
using Bridge = Bridges::Bridge;

int main(int argc, char* argv[])  try {
  // Check usage
  if (--argc != 4) {
    throw Error("usage: check_bridges samples_dir bridges_dir pop_file sample");
  }

  // Process command line arguments
  const string samples_dir{argv[1]};
  const string bridges_dir{argv[2]};
  const Population pop{argv[3]};
  const string family_name{argv[4]};

  const Family family{pop.family(family_name)};
  const vector<Sample> samples{pop.samples(family)};
  const vector<string> sample_names{[&samples, &pop]() {
    vector<string> names;
    names.reserve(samples.size());
    for (const Sample sample : samples) {
      names.push_back(pop.sample(sample));
    }
    return names;
    }()};

  const vector<MUMdex> mumdexes{[&sample_names, &samples_dir]() {
      vector<MUMdex> mumdex;
      mumdex.reserve(sample_names.size());
      for (const string & name : sample_names) {
        mumdex.emplace_back(samples_dir + "/" + name + "/mumdex");
      }
      return mumdex;
    }()};

  const Reference & ref{mumdexes.front().reference()};
  const ChromosomeIndexLookup chr_lookup{ref};

  const vector<vector<FileVector<BridgeInfo>>> bridges{
    [&ref, &bridges_dir, &sample_names]() {
      vector<vector<FileVector<BridgeInfo>>> b;
      b.resize(ref.n_chromosomes());
      for (unsigned int c{0}; c != ref.n_chromosomes(); ++c) {
        b[c].reserve(sample_names.size());
        for (const string & name : sample_names) {
          ostringstream bridges_name;
          bridges_name << bridges_dir << "/" << name << "/"
                       << get_bridge_file_name(ref, c);
          if (!readable(bridges_name.str()))
            throw Error("Could not open bridges file")
                << bridges_name.str() << paa::bridges_bad_message();
          b[c].emplace_back(bridges_name.str());
        }
      }
      return b;
    }()};

  // Read in bridge information and process each
  string chrs[2];
  unsigned int pos[2];
  bool high[2];
  int64_t invariant;
  int32_t offset;
  vector<uint64_t> seen_pairs;
  while (cin >> chrs[0] >> pos[0] >> high[0]
         >> chrs[1] >> pos[1] >> high[1]
         >> invariant >> offset) {
    const unsigned int chr[2]{chr_lookup[chrs[0]], chr_lookup[chrs[1]]};

    // Get bridge counts for samples in family
    const OneBridgeInfo bridge{chr[0], pos[0], high[0],
          chr[1], pos[1], high[1], offset};
    const vector<unsigned int> bridge_counts{
      [bridge, &bridges]() {
        vector<unsigned int> counts;
        const unsigned int c{bridge.chr1()};
        counts.reserve(bridges[c].size());
        for (const FileVector<BridgeInfo> & b : bridges[c]) {
          const auto range = equal_range(b.begin(), b.end(), bridge);
          if (range.first == range.second) {
            counts.push_back(0);
          } else {
            counts.push_back((*range.first).bridge_count());
          }
        }
        return counts;
      }()};

    if (*max_element(bridge_counts.begin(), bridge_counts.end()) == 0) {
      continue;
    }

    for (unsigned int s{0}; s != samples.size(); ++s) {
      const Sample sample{samples[s]};
      const string & sample_name{sample_names[s]};
      const MUMdex & mumdex{mumdexes[s]};

      // Get coverage
      unsigned int coverage[2]{0, 0};
      unsigned int anchor[2]{0, 0};
      unsigned int big_anchor[2]{0, 0};
      const unsigned int min_coverage_support{25};
      for (const unsigned int anchor2 : { false, true }) {
        const unsigned int achr{chr[anchor2]};
        const unsigned int apos{pos[anchor2]};
        const bool ahigh{high[anchor2]};
        seen_pairs.clear();
        const unsigned int max_read_length{154};  // record in mumdex!
        const auto region = mumdex.region(achr, apos > max_read_length ?
                                          apos - max_read_length :
                                          0, achr, apos + 1);
        for (const MUMindex pm : region) {
          const Pair pair{mumdex.pair(pm)};
          if (pair.dupe()) continue;
          const MUM mum{mumdex.mum(pm)};
          if (pair.bad(mum.read_2())) continue;
          if (mum.position0() > apos) continue;
          if (mum.position0() + mum.length() - 1 < apos) continue;
          if (mum.length() < min_coverage_support) continue;

          if (achr == mum.chromosome() &&
              ((ahigh == false && apos == mum.position0()) ||
               (ahigh == true && apos == mum.position0() + mum.length() - 1))) {
            ++anchor[anchor2];
          }

          // increment coverage if pair not yet seen
          if (find(seen_pairs.begin(), seen_pairs.end(),
                   pm.pair_index()) != seen_pairs.end()) continue;
          seen_pairs.push_back(pm.pair_index());

          if (achr == mum.chromosome() &&
              ((ahigh == false && apos == mum.position0()) ||
               (ahigh == true && apos == mum.position0() + mum.length() - 1))) {
            ++big_anchor[anchor2];
          }
          ++coverage[anchor2];
        }
      }

      // Output information for bridge
      sout << sample_name << pop.member(sample) << family_name
           << bridge_counts[s];
      for (const unsigned int anchor2 : { false, true }) {
        sout << coverage[anchor2];
      }
      for (const unsigned int anchor2 : { false, true }) {
        sout << big_anchor[anchor2];
      }
      for (const unsigned int anchor2 : { false, true }) {
        sout << anchor[anchor2];
      }
      for (const unsigned int anchor2 : { false, true }) {
        sout << chrs[anchor2] << pos[anchor2] << high[anchor2];
      }
      sout << invariant << offset << endl;
    }
  }
  cerr << "all done" << endl;

  return 0;
} catch (Error & e) {
  cerr << "paa::Error:" << endl;
  cerr << e.what() << endl;
  return 1;
} catch (exception & e) {
  cerr << "std::exception" << endl;
  cerr << e.what() << endl;
  return 1;
} catch (...) {
  cerr << "unknown exception was caught" << endl;
  return 1;
}
```
