## Supplementary material for "G-Graph: An interactive genomic graph viewer": MUMdex Source Code for G-Graph: check_bridges.py

```
#! /usr/bin/env python

import sys
import mumdex

sample=sys.argv[1]
mumdex_dir="/data/safe/paa/analysis/mums/wg-output/samples/" + sample + "/mumdex"
mums = mumdex.MUMdex(mumdex_dir)
ref = mums.Reference() 

chr1s=sys.argv[2]
pos1=int(sys.argv[3]) + 1
out1=bool(int(sys.argv[4]))

chr2s=sys.argv[5]
pos2=int(sys.argv[6]) + 1
out2=bool(int(sys.argv[7]))

chr1 = ref.index(chr1s)
chr2 = ref.index(chr2s)

pms1=dict()
p1s=set()
for pm in mums.anchors_at(chr1, pos1, out1):
    if mums.MUM(pm[1]).length() >= 20:
        if not pm[0] in pms1:
            pms1[pm[0]] = set()
        pms1[pm[0]].add(pm[1])
        p1s.add(pm[0])

pms2=dict()
p2s=set()
for pm in mums.anchors_at(chr2, pos2, out2):
    if mums.MUM(pm[1]).length() >= 20:
        if not pm[0] in pms2:
            pms2[pm[0]] = set()
        pms2[pm[0]].add(pm[1])
        p2s.add(pm[0])

def invariant(mums, p, m1, out1, m2, out2):
    pair = mums.Pair(p)
    mum1 = mums.MUM(m1)
    mum2 = mums.MUM(m2)

    read_length = pair.length(mum1.read_2())
    mum1_readpos = mums.readpos((p,m1))
    mum2_readpos = mums.readpos((p,m2))
    if mum1.flipped() == mum2.flipped():
        orientation = "="
        if out1:
            sign = -1
        else:
            sign = 1
        invariant = sign * (mum2_readpos - mum1_readpos);
    else:
        if out1:
            orientation = "i"
        else:
            orientation = "o"
        invariant = read_length + mum1_readpos + mum2_readpos;
    return (orientation, invariant)

ps=p1s.intersection(p2s)
for p in sorted(ps):
    seqs = mums.sequences(p)
    pair = mums.Pair(p)
    print "dupe", "seq1", "seq2"
    print pair.dupe(), seqs[0], seqs[1]
    print
    print "chr", "pos", "out", "flipped", "read2", "offset", "length"
    for m in pms1[p]:
        mum = mums.MUM(m)
        print chr1s, pos1 - 1, out1, mum.flipped(), mum.read_2(), mum.offset(), mum.length(), 
        for m2 in pms2[p]:
            mum2 = mums.MUM(m2)
            print invariant(mums, p, m, out1, m2, out2),
        print

    print "chr", "pos", "out", "flipped", "read2", "offset", "length"
    for m in pms2[p]:
        mum = mums.MUM(m)
        print chr2s, pos2 - 1, out2, mum.flipped(), mum.read_2(), mum.offset(), mum.length()
    print
print len(ps)
```
