## Supplementary material for "G-Graph: An interactive genomic graph viewer": MUMdex Source Code for G-Graph: check_bridges_result.cpp

```
//
// check_bridges_result
//
// analyze result of check_bridges
//
// Copyright 2016 Peter Andrews @ CSHL
//

#include <algorithm>
#include <exception>
#include <fstream>
#include <functional>
#include <iomanip>
#include <iostream>
#include <map>
#include <random>
#include <sstream>
#include <string>
#include <utility>
#include <vector>

#include "error.h"
#include "mumdex.h"
#include "population.h"
#include "psplot.h"
#include "utility.h"

using std::cerr;
using std::cin;
using std::cout;
using std::endl;
using std::exception;
using std::function;
using std::ifstream;
using std::istringstream;
using std::map;
using std::min;
using std::max;
using std::mt19937_64;
using std::ostringstream;
using std::pair;
using std::random_device;
using std::setprecision;
using std::string;
using std::uniform_real_distribution;
using std::vector;

using dist_real = uniform_real_distribution<double>;

using paa::graph_defaults;
using paa::sout;
using paa::Bounds;
using paa::ChromosomeIndexLookup;
using paa::Error;
using paa::Family;
using paa::Marker;
using paa::Population;
using paa::PSDoc;
using paa::PSPage;
using paa::PSGraph;
using paa::PSXYSeries;
using paa::PSXYMSeries;
using paa::Sample;

using Reference = paa::UnMappedReference;
using Mappability = paa::UnMappedMappability;

template <class Type>
double avg(const vector<Type> & vals) {
  if (vals.empty()) return 0.0;
  return accumulate(vals.begin(), vals.end(), 0.0) / vals.size();
}

template <class Type>
Type median(vector<Type> vals) {
  if (vals.empty()) return 0.0;
  sort(vals.begin(), vals.end());
  return vals[(vals.size() - 1) / 2];
}

const unsigned int pre{4};
template <class Type>
string avg_median(const vector<Type> & vals) {
  ostringstream out;
  out << setprecision(pre) << avg(vals) << " "
      << setprecision(pre) << median(vals);
  return out.str();
}


int main(int argc, char* argv[])  try {
  // Check usage
  if (--argc < 4) {
    throw Error("usage: check_bridges_result "
                "ref pop_file bridges_file result_file ...");
  }

  random_device rd;
  mt19937_64 mersenne{rd()};
  function<double()> unitGen{
    bind(dist_real(-0.5, 0.5), mersenne)};

  // Process command line arguments
  const Reference ref{argv[1]};
  const Population pop{argv[2]};
  const string bridges_name{argv[3]};
  ifstream bridges_file{bridges_name.c_str()};
  if (!bridges_file)
    throw Error("Problem opening bridges file") << bridges_name;

  const ChromosomeIndexLookup lookup{ref};
  const Mappability mappability{ref};

  // Read in bridges information
  const int Lr{151};
  vector<string> bridges;
  map<string, unsigned int> bridges_lookup;
  vector<vector<unsigned int>> mappabilities;
  vector<int> offsets;
  vector<int> coverage_sums;
  vector<int> bridge_sums;
  vector<int> repeat_lengths;
  map<string, bool> in_zone;

  ifstream lgds{"lgd_pos.txt"};
  double score;
  string chr_s;
  unsigned int bpos;
  unsigned int epos;
  using APP = pair<unsigned int, unsigned int>;
  vector<APP> lgd_abspos;
  while (lgds >> score >> chr_s >> bpos >> epos) {
    if (chr_s == "23") chr_s = "X";
    if (chr_s == "24") chr_s = "Y";
    if (chr_s == "-1") continue;
    const unsigned int c{lookup[chr_s]};
    if (score < 0.1)
      lgd_abspos.emplace_back(ref.abspos(c, bpos), ref.abspos(c, epos));
  }
  auto cpos = [](const APP lhs, const APP rhs) {
    return lhs.second < rhs.second;
  };
  auto dpos = [](const APP lhs, const unsigned int rhs) {
    return lhs.second < rhs;
  };
  sort(lgd_abspos.begin(), lgd_abspos.end(), cpos);
  vector<bool> in_lgd;

  string bridge;
  unsigned int n_in_lgd{0};
  while (getline(bridges_file, bridge)) {
    size_t last{bridge.find_last_of(' ')};
    istringstream bridge_stream{bridge.substr(last).c_str()};
    unsigned int rlen;
    bridge_stream >> rlen;
    bridge.erase(last);
    bridges_lookup[bridge] = static_cast<unsigned int>(bridges.size());
    bridges.push_back(bridge);
    string chrs[2];
    unsigned int pos[2];
    bool h[2];
    int invariant;
    int offset;
    istringstream bin{bridge.c_str()};
    bin >> chrs[0] >> pos[0] >> h[0]
        >> chrs[1] >> pos[1] >> h[1]
        >> invariant >> offset;
    if (!bin) throw Error("Problem interpreting bridge string");
    int coverage_sum{2 + 2 * Lr};
    int bridge_sum{Lr + 2 - offset};
    offsets.push_back(offset);
    mappabilities.push_back(vector<unsigned int>());
    bool inLGD{false};
    for (const bool anchor2 : { false, true }) {
      const unsigned int chr{lookup[chrs[anchor2]]};
      for (unsigned int x{pos[anchor2]};
           x != min(ref.size(chr), pos[anchor2]) + Lr; ++x) {
        const unsigned int abspos{ref.abspos(chr, x)};
        coverage_sum += (mappability.high(abspos) <= Lr);
      }
      const unsigned int abspos{ref.abspos(chr, pos[anchor2])};
      auto found = lower_bound(lgd_abspos.begin(), lgd_abspos.end(),
                               abspos, dpos);
      while (found != lgd_abspos.end() &&
             abspos <= found->second) {
        if (abspos < found->second && abspos >= found->first) {
          inLGD = true;
          break;
        }
        if (abspos < found->first) break;
        ++found;
      }
      const unsigned int mappa{mappability.low_high(h[anchor2], abspos)};
      mappabilities.back().push_back(mappa);
      bridge_sum -= min(20u, mappa);
    }
    if (inLGD) ++n_in_lgd;
    in_lgd.push_back(inLGD);
    coverage_sums.push_back(coverage_sum);
    bridge_sums.push_back(bridge_sum);
    repeat_lengths.push_back(rlen);
  }
  cerr << "Read " << bridges.size() << " bridges" << endl;
  cerr << "In LGD " << n_in_lgd << " n " << lgd_abspos.size() << endl;

  // Read in input files
  argc -= 2;
  argv += 2;
  string sample_name;
  string member;
  string family_name;
  struct Info {
    Info() :
        sample{0}, bridge_count{0},
      coverage{0, 0}, anchors{0, 0}, anchors_all{0, 0},
      bridge{0} { }
    Sample sample;
    unsigned int bridge_count;
    unsigned int coverage[2];
    unsigned int anchors[2];
    unsigned int anchors_all[2];
    unsigned int bridge;
  };
  vector<vector<vector<Info>>> info(bridges.size());
  vector<vector<double>> all_sample_corrs(pop.n_samples());
  unsigned int n_lines{0};
  while (--argc) {
    ++argv;
    const string input_name{argv[1]};
    ifstream input{input_name.c_str()};
    if (!input) throw Error("Problem opening input file") << input_name;
    Info i;
    while (input >> sample_name >> member >> family_name
           >> i.bridge_count
           >> i.coverage[0] >> i.coverage[1]
           >> i.anchors[0] >> i.anchors[1]
           >> i.anchors_all[0] >> i.anchors_all[1]) {
      i.sample = pop.sample(sample_name);
      input.get();
      getline(input, bridge);
      // cout << bridge << endl;
      const unsigned int b{bridges_lookup.at(bridge)};
      i.bridge = b;
      vector<vector<Info>> & bridge_info{info[b]};
      if (bridge_info.empty() || bridge_info.back().size() == 4) {
        bridge_info.push_back(vector<Info>());
      }
      vector<Info> & family_info{bridge_info.back()};
      family_info.push_back(i);
      all_sample_corrs[i.sample].push_back(
          1.0 * (i.coverage[0] + i.coverage[1]) / coverage_sums[b]);
      ++n_lines;
    }
  }
  const vector<double> sample_corrs{[all_sample_corrs]() {
      vector<double> sc;
      for (vector<double> asc : all_sample_corrs) {
        if (asc.size()) {
          sort(asc.begin(), asc.end());
          sc.push_back(asc[(asc.size() - 1) / 2]);
        } else {
          sc.push_back(0);
        }
      }
      return sc;
    }()};

  cerr << "Read " << n_lines << " lines for "
       << info.size() << " bridges" << endl;

  vector<vector<double>> all_expected;
  vector<vector<unsigned int>> all_mappability;
  vector<vector<int>> all_offsets;
  vector<vector<int>> all_coverage;

  Marker marker{paa::circle(), 0.2, "0 0 0", 1, true};
  PSDoc doc1{"coverage"};

  graph_defaults.title_size(30).label_size(24).legend_size(24).
      tick_size(20).border_width(2).grid_width(2).tick_width(2);

  PSXYSeries bcounts{"bridge_counts", ";"
        "Candidate Bridge Count;"
        "Max Other Family Bridge Count",
        Bounds{-1, 45, -1, 35}, marker};

  PSPage offset_page("offset", "", "1 2");
  PSGraph offset_hist{offset_page, ";Offset;Frequency", Bounds{-60, 20}};
  offset_hist.add_text(0.95, 0.85, paa::Text{"A", 40});
  paa::PSHSeries<int, uint64_t> offset1{offset_hist, 80, "0 0 1", true,
        "N Family == 1"};
  paa::PSHSeries<int, uint64_t> offset2{offset_hist, 80, "1 0 0", true,
        "N Family > 1"};
  PSGraph trans_offset_hist{offset_page, ";Offset;Frequency", Bounds{-60, 20}};
  trans_offset_hist.add_text(0.95, 0.85, paa::Text{"B", 40});
  trans_offset_hist.add(&offset1);
  paa::PSHSeries<int, uint64_t> offsett{trans_offset_hist, 80, "1 0 0", true,
        "Transmitted"};

  graph_defaults.reset();

  PSXYSeries trans{"transmission", ";"
        "Mean Members;"
        "Max Other Family Bridge Count",
        Bounds{0.5, 4.5, 0, 35}, marker};

  PSGraph offset_both_hist{doc1, "Anchor Position Read Offsets;"
        "Offset;Frequency", Bounds{-60, 20}};
  offset_both_hist.add(&offset1);
  paa::PSHSeries<int, uint64_t> offsetbb{offset_both_hist, 80, "1 0 0", true,
        "Sibling and Proband"};

  PSGraph repeat_hist{"repeat", ";Repeat Length;Frequency", Bounds{0, 100}};
  paa::PSHSeries<int, uint64_t> repeat1{repeat_hist, 20, "0 0 1", true,
        "N Family == 1"};
  paa::PSHSeries<int, uint64_t> repeat2{repeat_hist, 20, "1 0 0", true,
        "N Family > 1"};


  // PSPage apage{doc1};

  PSDoc doc{"coverage2", "coverage"};

  // fix this one to max other??
  PSXYMSeries counts{doc, "Counts (red is longer mappability);"
        "Candidate Bridge Count;"
        "Other Family Bridge Count",
        Bounds{-1, 35, -1, 35}, marker};

#if 0
  PSPage vsmap_page{doc, "Mappability", "1 2"};
  Bounds vsmap_bounds{0, 2, 0, 140};
  PSGraph vsmapw_graph{vsmap_page,
        "Weak Cuts;Measured / Expected Bridge Count;Mappability",
        vsmap_bounds};
  PSXYSeries vsmapw{vsmapw_graph, marker};
  PSGraph vsmaps_graph{vsmap_page,
        "Strong Cuts;Measured / Expected Bridge Count;Mappability",
        vsmap_bounds};
  PSXYSeries vsmaps{vsmaps_graph, marker};


  PSPage hvscount_page{doc, "Expected Bridge Count Comparison", "1 2"};
  PSGraph hvscountw_graph{hvscount_page,
        "Weak Cuts;Measured / Expected Bridge Count;Expected Bridge Count",
        vsmap_bounds};
  hvscountw_graph.hist(1).add(&vsmapw);
  PSGraph hvscounts_graph{hvscount_page,
        "Strong Cuts;Measured / Expected Bridge Count;Expected Bridge Count",
        vsmap_bounds};
  hvscounts_graph.hist(1).add(&vsmaps);


  PSPage hvsmap_page{doc, "Mappability", "1 2"};
  PSGraph hvsmapw_graph{hvsmap_page,
        "Weak Cuts;Measured / Expected Bridge Count;Mappability",
        vsmap_bounds};
  hvsmapw_graph.yhist(1).add(&vsmapw);
  PSGraph hvsmaps_graph{hvsmap_page,
        "Strong Cuts;Measured / Expected Bridge Count;Mappability",
        vsmap_bounds};
  hvsmaps_graph.yhist(1).add(&vsmaps);


  PSPage vsrep_page{doc, "Repeat Length", "1 2"};
  Bounds vsrep_bounds{0, 2, 0, 85};
  PSGraph vsrepw_graph{vsrep_page,
        "Weak Cuts;Measured / Expected Bridge Count;Repeat Length",
        vsrep_bounds};
  PSXYSeries vsrepw{vsrepw_graph, marker};
  PSGraph vsreps_graph{vsrep_page,
        "Strong Cuts;Measured / Expected Bridge Count;Repeat Length",
        vsrep_bounds};
  PSXYSeries vsreps{vsreps_graph, marker};


  PSPage hvsrep_page{doc, "Repeat Length", "1 2"};
  PSGraph hvsrepw_graph{hvsrep_page,
        "Weak Cuts;Measured / Expected Bridge Count;Repeat Length",
        vsrep_bounds};
  hvsrepw_graph.yhist(1).add(&vsrepw);
  PSGraph hvsreps_graph{hvsrep_page,
        "Strong Cuts;Measured / Expected Bridge Count;Repeat Length",
        vsrep_bounds};
  hvsreps_graph.yhist(1).add(&vsreps);


  PSPage vsoff_page{doc, "Bridge Offset", "1 2"};
  Bounds vsoff_bounds{0, 2, -50, 50};
  PSGraph vsoffw_graph{vsoff_page,
        "Weak Cuts;Measured / Expected Bridge Count;Bridge Offset",
        vsoff_bounds};
  PSXYSeries vsoffw{vsoffw_graph, marker};
  PSGraph vsoffs_graph{vsoff_page,
        "Strong Cuts;Measured / Expected Bridge Count;Bridge Offset",
        vsoff_bounds};
  PSXYSeries vsoffs{vsoffs_graph, marker};

  PSPage hvsoff_page{doc, "Bridge Offset", "1 2"};
  PSGraph hvsoffw_graph{hvsoff_page,
        "Weak Cuts;Measured / Expected Bridge Count;Bridge Offset",
        vsoff_bounds};
  hvsoffw_graph.yhist(1).add(&vsoffw);
  PSGraph hvsoffs_graph{hvsoff_page,
        "Strong Cuts;Measured / Expected Bridge Count;Bridge Offset",
        vsoff_bounds};
  hvsoffs_graph.yhist(1).add(&vsoffs);
#endif

  PSXYSeries map_vs_cov{doc, "Mappability;Coverage;Mappability",
        Bounds{-1, 35, 0, 120}, marker};
  PSXYSeries offset_vs_cov{doc, "Offset;Coverage;Offset",
        Bounds{-1, 35, -90, 30}, marker};
  PSXYMSeries ratios{doc, "Ratios (red is longer mappability);"
        "Candidate Measured / Expected;"
        "Other Families Measured / Expected",
        Bounds{0, 2, 0, 1.5}, marker};
  PSXYSeries others{doc, "Other People;Candidate Measured / Expected;N people",
        Bounds{0.0}, marker};
  PSXYSeries mapg{doc, "Mappability;Candidate Measured / Expected;Mappability",
        Bounds{0.0}, marker};
  PSXYSeries rleng{doc, "Repeat Length;Candidate Measured / Expected;"
        "Repeat Length", Bounds{0.0}, marker};
  PSXYSeries maxg{doc, "Other People Max;Candidate Measured / Expected;"
        "Maximum Other Measured / expected", Bounds{0.0}, marker};


  PSPage blank_page{doc};

#if 0
  PSPage page{doc};
  PSGraph graph{page};
  graph.add(&ratios);
#endif

  sout << "formike cand_expected max_other other_expected repeat_length "
      "chr pos high chr pos high invariant offset" << endl;

  unsigned int ps{0};
  unsigned int ks{0};
  unsigned int pc{0};
  unsigned int kc{0};
  unsigned int parent_seen{0};
  unsigned int kids_seen{0};
  unsigned int parent_count{0};
  unsigned int kids_count{0};
  unsigned int kids_only_seen{0};
  unsigned int one_kid_only_seen{0};
  unsigned int n_aut{0};
  unsigned int n_sib{0};
  unsigned int n_lgd_aut{0};
  unsigned int n_lgd_sib{0};
  // unsigned int kids_only_count{0};
  for (const vector<vector<Info>> & bridge_info : info) {
    if (bridge_info.empty()) continue;
    const bool use_bridge{
      bridges[bridge_info[0][0].bridge].find_first_of("XYMG") == string::npos};
    unsigned int n_family{0};
    unsigned int mkf{0};
    for (const vector<Info> & family_info : bridge_info) {
      // No X Y, etc
      if (use_bridge) {
        for (const unsigned int parent : { 0, 1 }) {
          if (family_info[parent].bridge_count) {
            ++ps;
            pc += family_info[parent].bridge_count;
          }
        }
        for (const unsigned int kid : { 2, 3 }) {
          if (family_info[kid].bridge_count) {
            ++ks;
            kc += family_info[kid].bridge_count;
          }
        }
        if (family_info[0].bridge_count + family_info[1].bridge_count == 0 &&
            max(family_info[2].bridge_count, family_info[3].bridge_count) >
            max(bridge_info[mkf][2].bridge_count,
                bridge_info[mkf][3].bridge_count)) {
          mkf = n_family;
        }

        if ((family_info[0].bridge_count + family_info[1].bridge_count)) {
          if (!family_info[0].bridge_count || !family_info[1].bridge_count) {
            ++parent_seen;
            parent_count += family_info[0].bridge_count +
                family_info[1].bridge_count;
            for (const unsigned int kid : { 2, 3 }) {
              if (family_info[kid].bridge_count) {
                ++kids_seen;
                kids_count += family_info[kid].bridge_count;
              }
            }
          }
        } else {
          if (family_info[2].bridge_count + family_info[3].bridge_count == 0) {
            throw Error("Expected a child count");
          }
          ++kids_only_seen;
          if (family_info[2].bridge_count == 0 ||
              family_info[3].bridge_count == 0) {
            ++one_kid_only_seen;
          }
        }
      }
      ++n_family;
      for (const Info & i : family_info) {
        const unsigned int b{i.bridge};
        const double expected{sample_corrs[i.sample] * bridge_sums[b]};
        if (use_bridge) {
          if (i.bridge_count + 1 > all_expected.size()) {
            all_expected.resize(i.bridge_count + 1);
            all_mappability.resize(i.bridge_count + 1);
            all_offsets.resize(i.bridge_count + 1);
            all_coverage.resize(i.bridge_count + 1);
          }
          map_vs_cov.add_point(i.bridge_count + unitGen(),
                               max(mappabilities[b][0],
                                   mappabilities[b][1]) + unitGen());
          offset_vs_cov.add_point(i.bridge_count + unitGen(),
                                  offsets[b] + unitGen());
          all_mappability[i.bridge_count].push_back(mappabilities[b][0]);
          all_mappability[i.bridge_count].push_back(mappabilities[b][1]);
          all_expected[i.bridge_count].push_back(expected);
          all_offsets[i.bridge_count].push_back(offsets[b]);
          all_coverage[i.bridge_count].push_back(i.coverage[0]);
          all_coverage[i.bridge_count].push_back(i.coverage[1]);
        }
        sout << pop.family(pop.family(i.sample))
             << pop.sample(i.sample)
             << pop.member(i.sample)
             << n_family
             << i.bridge_count;
        for (const bool anchor2 : { false, true }) {
          sout << i.coverage[anchor2];
        }
        for (const bool anchor2 : { false, true }) {
          sout << i.anchors[anchor2];
        }
        for (const bool anchor2 : { false, true }) {
          sout << i.anchors_all[anchor2];
        }
        sout << bridges.at(i.bridge) << endl;
      }
    }
    if (use_bridge) {
      const unsigned int b{bridge_info[0][0].bridge};
      double max_count{0};
      double max_expected{0};
      double expected_1{0};
      unsigned int max_other{0};
      for (const unsigned int kid : { 2, 3 }) {
        const Info & i{bridge_info[mkf][kid]};
        if (max_count < i.bridge_count) {
          max_count = i.bridge_count;
          expected_1 = sample_corrs[i.sample] * bridge_sums[i.bridge];
        }
        if (i.bridge_count) {
          --ks;
          kc -= i.bridge_count;
        }
      }
      unsigned int nfamily{0};
      unsigned int n_people{0};
      for (const vector<Info> & family_info : bridge_info) {
        double family_max_count{0};
        double expected_2{0};
        if (nfamily != mkf) {
          for (const Info & i : family_info) {
            if (bridge_info[mkf][2].bridge_count) {
              if (family_info[0].bridge_count ||
                  family_info[1].bridge_count) {
                n_aut += (family_info[2].bridge_count > 0);
                n_sib += (family_info[3].bridge_count > 0);
                if (in_lgd[b]) {
                  n_lgd_aut += (family_info[2].bridge_count > 0);
                  n_lgd_sib += (family_info[3].bridge_count > 0);
                }
              }
            }

            if (i.bridge_count) ++n_people;
            if (family_max_count < i.bridge_count) {
              family_max_count = i.bridge_count;
              expected_2 = sample_corrs[i.sample] * bridge_sums[i.bridge];
            }
            if (max_other < i.bridge_count) {
              max_other = i.bridge_count;
              max_expected = expected_2;
            }
          }
          const unsigned int max_map{max(mappabilities[b][0],
                                         mappabilities[b][1])};
          const double color{(max_map > 70 ? 70 : max_map) / 70.0};
          ostringstream color_string;
          color_string << color << " 0 0";
          // cout << "color " << color_string.str() << endl;
          marker.scale(0.3);
          marker.color(color_string.str());
          marker.fill_color(color_string.str());
          counts.add_point(max_count + unitGen(),
                           family_max_count + unitGen(), marker);
          if (expected_1 > 0 && expected_2 > 0) {
            ratios.add_point((max_count + unitGen()) / expected_1,
                             (family_max_count + unitGen()) / expected_2,
                             marker);
          }
        }
        ++nfamily;
      }
#if 0
      if (n_family == 1) {
        vsmaps.add_point((max_count + unitGen()) / expected_1,
                         max(mappabilities[b][0], mappabilities[b][1])
                         + unitGen());
      } else {
        vsmapw.add_point((max_count + unitGen()) / expected_1,
                         max(mappabilities[b][0], mappabilities[b][1])
                         + unitGen());
      }

      if (n_family == 1) {
        vsreps.add_point((max_count + unitGen()) / expected_1,
                         repeat_lengths[b] + unitGen());
      } else {
        vsrepw.add_point((max_count + unitGen()) / expected_1,
                         repeat_lengths[b] + unitGen());
      }


      if (n_family == 1) {
        vsoffs.add_point((max_count + unitGen()) / expected_1,
                         offsets[b] + unitGen());
      } else {
        vsoffw.add_point((max_count + unitGen()) / expected_1,
                         offsets[b] + unitGen());
      }
#endif

      bcounts.add_point(max(bridge_info[mkf][2].bridge_count,
                            bridge_info[mkf][3].bridge_count) + 0.8 * unitGen(),
                        max_other + 0.8 * unitGen());

      if (max_other > 1) {
        cout << "special " << bridges[b]
             << " " << n_family
             << " " << mappabilities[b][0]
             << " " << mappabilities[b][1]
             << " " << repeat_lengths[b]
             << endl;
      }

      const int n_in_family{(bridge_info[mkf][2].bridge_count > 0) +
            (bridge_info[mkf][3].bridge_count > 0)};
      if (n_family == 1) {
        offset1.add_point(offsets[b]);
        repeat1.add_point(repeat_lengths[b]);
      } else {
        offset2.add_point(offsets[b]);
      }
      if (n_in_family == 2) {
        offsetbb.add_point(offsets[b]);
      }

      if (n_family > 1) {
        repeat2.add_point(repeat_lengths[b]);
        trans.add_point(0.1 * unitGen() + 1.0 * n_people /
                        (n_family - 1),
                        max_other + unitGen());
      }
      if (max_other > 1 && n_family > 1 &&
          n_people / (n_family - 1) > 1) {
        offsett.add_point(offsets[b]);
      }

      // cout << "offset " << offsets[b] << endl;

      others.add_point((max_count + unitGen()) / expected_1,
                       n_people + unitGen());
      mapg.add_point((max_count + unitGen()) / expected_1,
                     max(mappabilities[b][0], mappabilities[b][1])
                     + unitGen());
      rleng.add_point((max_count + unitGen()) / expected_1,
                      repeat_lengths[b] + unitGen());
      maxg.add_point((max_count + unitGen()) / expected_1,
                     (max_other + unitGen()) / expected_1);
      sout << "formike" << expected_1
           << max_other << max_expected << repeat_lengths[b] << bridges[b]
           << endl;
    }
  }
  kids_only_seen -= info.size();
  one_kid_only_seen -= info.size();

  // Result
  sout << parent_seen << kids_seen << 1.0 * kids_seen / parent_seen
       << parent_count << kids_count << 1.0 * kids_count / parent_count
       << 1.0 * kids_seen / (parent_seen - kids_only_seen)
       << 1.0 * kids_seen / (parent_seen - one_kid_only_seen)
       << kids_only_seen
       << info.size()
       << endl;
  sout << ps << ks << 1.0 * ks / ps
       << pc << kc << 1.0 * kc / pc << endl;

  sout << "cov"
       << "nb"
       << "n"
       << "exp" << "med"
       << "ratio" << "med"
       << "map" << "med"
       << "off" << "med"
       << "cov" << "med"
       << endl;
  for (unsigned int b{0}; b != all_expected.size(); ++b) {
    if (all_expected[b].size()) {
      const double expected{avg(all_expected[b])};
      const double med{median(all_expected[b])};
      cout << "cov "
           << b << " "
           << all_expected[b].size() << " "
           << setprecision(pre) << expected << " "
           << setprecision(pre) << med << " "
           << setprecision(pre) << b / expected << " "
           << setprecision(pre) << b / med << " "
           << avg_median(all_mappability[b]) << " "
           << avg_median(all_offsets[b]) << " "
           << avg_median(all_coverage[b]) << " "
           << endl;
    }
  }

  sout << "aut" << n_aut << "sib" << n_sib << endl;
  sout << "lgd aut" << n_lgd_aut << "sib" << n_lgd_sib << endl;
  cerr << "all done" << endl;
  return 0;
} catch (Error & e) {
  cerr << "paa::Error:" << endl;
  cerr << e.what() << endl;
  return 1;
} catch (exception & e) {
  cerr << "std::exception" << endl;
  cerr << e.what() << endl;
  return 1;
} catch (...) {
  cerr << "unknown exception was caught" << endl;
  return 1;
}
```
