## Supplementary material for "G-Graph: An interactive genomic graph viewer": MUMdex Source Code for G-Graph: check_cand_cn.cpp

```
//
// check_cand_cn
//
// See if can see CN signal for SV candidates
//
// Copyright 2017 Peter Andrews @ CSHL
//

#include <exception>
#include <iostream>
#include <fstream>
#include <memory>
#include <sstream>
#include <string>
#include <utility>
#include <vector>

#include "cn.h"
#include "cngsl.h"
#include "error.h"
#include "mumdex.h"
#include "population.h"
#include "psplot.h"

using std::exception;
using std::cout;
using std::cerr;
using std::endl;
using std::ifstream;
using std::make_unique;
using std::move;
using std::ostringstream;
using std::string;
using std::unique_ptr;
using std::vector;

using paa::Bin;
using paa::ChromosomeIndexLookup;
using paa::CN_Bins;
using paa::CNStateCall;
using paa::CNStateCaller;
using paa::Error;
using paa::Family;
using paa::Marker;
using paa::Population;
using paa::PSDoc;
using paa::PSGraph;
using paa::PSXYSeries;
using paa::Reference;
using paa::Sample;

int main(int argc, char* argv[]) try {
  if (--argc != 4)
    throw Error("usage: check_cand_cn ref pop_file bins cand_file");

  const string ref_name{argv[1]};
  const Reference ref{ref_name};
  const ChromosomeIndexLookup chr_lookup{ref};

  const Population pop{argv[2]};

  const string bins_name{argv[3]};
  const vector<Bin> all_bins{load_bins(bins_name, ref)};
  const vector<Bin> bins{[&all_bins]() {
      vector<Bin> result;
      for (const Bin & bin : all_bins) {
        if (!bin.bad()) {
          result.push_back(bin);
        }
      }
      return result;
    }()};

  ifstream candidates{argv[4]};
  if (!candidates) throw Error("Could not open candidates file") << argv[2];

  // Plots
  PSDoc profiles_ps{"cn_denovo"};
  vector<unique_ptr<PSGraph>> profile_graphs;
  vector<unique_ptr<PSXYSeries>> profile_series;
  vector<string> profile_colors{"1 0 0", "0 0 1", "0 1 0", "1 1 0"};

  string family;
  string member;
  string chr_name[2];
  unsigned int pos[2];
  bool high[2];
  int invariant;
  int offset;
  unsigned int n_seen{0};
  unsigned int n_good{0};

  const string finish_line{" 0 0 X X X X X X X X 0 0 0 0 0"};

  while (candidates >> family >> member
         >> chr_name[0] >> pos[0] >> high[0]
         >> chr_name[1] >> pos[1] >> high[1]
         >> invariant >> offset) {
    const unsigned int chr[2]{chr_lookup[chr_name[0]],
          chr_lookup[chr_name[1]]};
    bool is_denovo{false};
    bool in_parents{false};
    bool in_member{false};

    cout << family << " " << member
         << " " << chr_name[0] << " " << pos[0] << " " << high[0]
         << " " << chr_name[1] << " " << pos[1] << " " << high[1]
         << " " << invariant << " " << offset;

    // Only look at bigger deletions
    if (chr[0] == chr[1] && high[0] != high[1] && labs(invariant) > 1000) {
      Family f{0};
      try {
        f = pop.family(family);
      } catch (...) {
        cerr << "Family not found " << family << endl;
        cout << finish_line << endl;
        continue;
      }

      const vector<Sample> & samples{pop.samples(f)};

      const unsigned int start_bin{[&bins, chr, pos]() {
          for (unsigned int b{0}; b != bins.size(); ++b) {
            const Bin & bin{bins[b]};
            if (bin.chromosome() != chr[0]) continue;
            if (bin.stop_position() >= pos[0]) return b;
          }
          throw Error("Start bin not found");
        }()};
      const unsigned int stop_bin{[&bins, chr, pos]() {
          for (unsigned int b{0}; b != bins.size(); ++b) {
            const Bin & bin{bins[b]};
            if (bin.chromosome() < chr[1]) continue;
            if (bin.chromosome() > chr[1]) return b;
            if (bin.start_position() >= pos[1]) return b;
          }
          return static_cast<unsigned int>(bins.size());
        }()};

      const unsigned int n_bins_seg{stop_bin - start_bin};
      if (n_bins_seg == 0) {
        cout << finish_line << endl;
        continue;
      }

      const unsigned int plot_start_bin{start_bin > n_bins_seg ?
            start_bin - n_bins_seg : 0};
      const unsigned int plot_stop_bin{stop_bin + n_bins_seg > bins.size() ?
            static_cast<unsigned int>(bins.size()) : stop_bin + n_bins_seg};

      const double bin_match_frac{1.0 *
            (pos[1] - pos[0]) /
            (bins[stop_bin].stop_position() -
             bins[start_bin].start_position())};

      cout << " " << n_bins_seg << " " << bin_match_frac;

      const string expected_call{invariant < 0 ? "loss" : "gain"};

      unique_ptr<PSGraph> graph{nullptr};

      for (unsigned int s{0}; s != samples.size(); ++s) {
        const Sample sample{samples[s]};
        try {
          ostringstream results_name;
          if (0) {
            results_name
                << "/data/safe/paa/analysis/mums/cn/new/samples_nocut/"
                << pop.sample(sample) << "/" << pop.sample(sample)
                << "_" << all_bins.size() << "_bins_results.txt";
          } else {
            results_name
                << "/data/safe/paa/analysis/mums/breakpoints_cn/cn470"
                << "/" << pop.sample(sample)
                << "_" << all_bins.size() << "_bins_results.txt";
          }

          const CN_Bins profile{results_name.str()};
          if (profile.size() != bins.size())
            throw Error("Profile bins size mismatch") << sample;

          if (graph == nullptr) {
            graph = make_unique<PSGraph>(profiles_ps, family + " " + member +
                                         " " + expected_call + " chr" +
                                         chr_name[0]);
          }
          const Marker marker{paa::circle(), 1, "0 0 0", 1, true,
                profile_colors[s]};

          profile_series.push_back(make_unique<PSXYSeries>(
              *graph, marker));
          PSXYSeries & series{*profile_series.back()};

          for (unsigned int b{plot_start_bin}; b != plot_stop_bin; ++b) {
            series.add_point(bins[b].start_position(), profile[b].ratio());
          }

          const CNStateCaller caller{ref, bins, profile};
          const CNStateCall call{caller.call(start_bin, stop_bin)};

          if (pop.member(sample) == "father" ||
              pop.member(sample) == "mother") {
            if (call.type == expected_call) in_parents = true;
          } else {
            if (call.type == expected_call) in_member = true;
          }
          cout << " " << call.type << " " << call.score;
        } catch (...) {
          cout << " X X";
        }
      }
      if (graph != nullptr) {
        ostringstream line_stream;
        line_stream << "1 lw 0 0 0 c np "
                    << pos[0] << " xc 0 yfc m "
                    << pos[0] << " xc 1 yfc l sp np "
                    << pos[1] << " xc 0 yfc m "
                    << pos[1] << " xc 1 yfc l sp";
        graph->ps(line_stream.str());
        profile_graphs.push_back(move(graph));
      }
      if (in_member && !in_parents) is_denovo = true;
      ++n_seen;
      if (is_denovo) ++n_good;
      cout << " 1"
           << " " << in_parents
           << " " << in_member
           << " " << is_denovo
           << " " << is_denovo;
    } else {
      cout << finish_line;
    }
    cout << endl;
  }
  cerr << n_good << " of " << n_seen << " or " << 1.0 * n_good / n_seen << endl;

  return 0;
} catch (Error & e) {
  cerr << "paa::Error:" << endl;
  cerr << e.what() << endl;
  return 1;
} catch (exception & e) {
  cerr << "std::exception" << endl;
  cerr << e.what() << endl;
  return 1;
} catch (...) {
  cerr << "unknown exception was caught" << endl;
  return 1;
}
```
