## Supplementary material for "G-Graph: An interactive genomic graph viewer": MUMdex Source Code for G-Graph: check_for_pseudogene.cpp

```
//
// check_for_pseudogene
//
// find denovo pseudogenes in candidate list
//
// Copyright 2017 Peter Andrews @ CSHL
//

#include <exception>
#include <iostream>
#include <fstream>
#include <string>

#include "error.h"
#include "genes.h"
#include "mumdex.h"

using std::exception;
using std::cout;
using std::cerr;
using std::endl;
using std::ifstream;
using std::string;

using paa::ChromosomeIndexLookup;
using paa::Error;
using paa::GeneXrefs;
using paa::JunctionInfo;
using paa::KnownGene;
using paa::KnownGenes;
using paa::Reference;

int main(int argc, char* argv[]) try {
  if (--argc != 2) throw Error("usage: check_for_pseudogenes ref cand_file");

  const string ref_name{argv[1]};
  const Reference ref{ref_name};
  const ChromosomeIndexLookup chr_lookup{ref};
  const string genes_name{ref_name + ".bin/knownGene.txt"};
  const string isoforms_name{ref_name + ".bin/knownIsoforms.txt"};
  const string kgXrefs_name{ref_name + ".bin/kgXref.txt"};

  const KnownGenes genes{genes_name, isoforms_name, chr_lookup, ref};
  const GeneXrefs xref{kgXrefs_name};

  ifstream candidates{argv[2]};
  if (!candidates) throw Error("Could not open candidates file") << argv[2];

  string chr_name[2];
  unsigned int pos[2];
  bool high[2];
  int invariant;
  int offset;

  const unsigned int exon_tolerance{20};
  const unsigned int invariant_tolerance{20};

  while (candidates >> chr_name[0] >> pos[0] >> high[0]
         >> chr_name[1] >> pos[1] >> high[1]
         >> invariant >> offset) {
    const unsigned int chr[2]{chr_lookup[chr_name[0]], chr_lookup[chr_name[1]]};
    bool is_pseudogene{false};

    // Only look at deletions for pseudogenes
    if (chr[0] == chr[1] && high[0] != high[1] && invariant < 0) {
      for (const KnownGene & gene : genes) {
        if (chr[0] != gene.chr) continue;
        // cout << "here " << gene.n_exons << " " << gene.name;
        for (unsigned int e{0}; e + 1 != gene.n_exons; ++e) {
          // cout << " " << gene.exon_starts[e] << " " << gene.exon_stops[e];
          if ((labs(static_cast<int64_t>(pos[0]) -
                    static_cast<int64_t>(gene.exon_stops[e])) <
               exon_tolerance ||
               labs(static_cast<int64_t>(pos[1]) -
                    static_cast<int64_t>(gene.exon_stops[e])) <
               exon_tolerance) &&
              labs(invariant +
                   (static_cast<int64_t>(gene.exon_starts[e + 1]) -
                    static_cast<int64_t>(gene.exon_stops[e]))) <
              invariant_tolerance) {
            is_pseudogene = true;
          }
        }
        // cout << endl;
      }
    }

    cout << chr_name[0] << " " << pos[0] << " " << high[0]
         << " " << chr_name[1] << " " << pos[1] << " " << high[1]
         << " " << invariant << " " << offset << " " << is_pseudogene << endl;
  }

  return 0;
} catch (Error & e) {
  cerr << "paa::Error:" << endl;
  cerr << e.what() << endl;
  return 1;
} catch (exception & e) {
  cerr << "std::exception" << endl;
  cerr << e.what() << endl;
  return 1;
} catch (...) {
  cerr << "unknown exception was caught" << endl;
  return 1;
}
```
