## Supplementary material for "G-Graph: An interactive genomic graph viewer": MUMdex Source Code for G-Graph: check_mappability.cpp

```
//
// check_mappability
//
// Create mappability from shortSA and
// check values against other mappability to make sure they are identical
//
// Copyright 2016 Peter Andrews @ CSHL
//

#include <exception>
#include <iostream>
#include <string>

#include "error.h"
#include "files.h"
#include "mumdex.h"
#include "longSA.h"

using std::cout;
using std::cerr;
using std::endl;
using std::exception;
using std::string;

using paa::Error;
using paa::Mappability;
using paa::MappabilityBuilder;
using paa::mkdir;
using paa::shortSA;

int main(int argc, char * argv[]) try {
  // paa::test_thread_pool();

  if (--argc != 1) throw Error("usage: check_mappability ref_fasta");

  const string fasta_name{argv[1]};
  const Mappability map{fasta_name, true};
  const shortSA sa{fasta_name, false, true};
  mkdir("test.bin");
  const MappabilityBuilder builder(sa, 24, "test.bin/", &map);

  return 0;
} catch (Error & e) {
  cerr << "paa::Error:" << endl;
  cerr << e.what() << endl;
  return 1;
} catch (exception & e) {
  cerr << "std::exception" << endl;
  cerr << e.what() << endl;
  return 1;
} catch (...) {
  cerr << "unknown exception was caught" << endl;
  return 1;
}
```
