## Supplementary material for "G-Graph: An interactive genomic graph viewer": MUMdex Source Code for G-Graph: check_mumdex.cpp

```
//
// check_mumdex
//
// See if mumdex file has any recognizable problems
//
// Copyright 2017 Peter Andrews @ CSHL
//

#include <algorithm>
#include <exception>
#include <iostream>
#include <string>

#include "error.h"
#include "mumdex.h"
#include "utility.h"

using std::cout;
using std::cerr;
using std::endl;
using std::exception;
using std::min;
using std::string;

using paa::Error;
using paa::MUMdex;
using paa::Mappability;
using paa::sout;
using paa::tout;

int main(int argc, char * argv[]) try {
  paa::exit_on_pipe_close();

  if (--argc != 1) throw Error("usage: check_mumdex mumdex_name");

  const string mumdex_name{argv[1]};
  const MUMdex mumdex{mumdex_name};
  // const Mappability map{mumdex_name};
  const auto & ref = mumdex.reference();

  for (uint64_t pair_index = 0; pair_index != mumdex.n_pairs(); ++pair_index) {
    const auto pair = mumdex.pair(pair_index);
    if (pair.has_mums()) {
      if (pair.mums_start() >= mumdex.n_mums())
        throw Error("mums_start for pair out of range");
      for (uint64_t m = pair.mums_start();; ++m) {
        const auto mum = mumdex.mum(m);
        auto report = [pair_index, m, &pair, &mum, &mumdex, &ref]
            (const string & message) {
          static bool first{true};
          if (first) {
            first = false;
            cout << "pair n_pair mum_in_pair mums_in_pair mum n_mum "
                "read_2 flipped chrname chr pos chr_size "
                "off len rlen msg\n";
          }
          cout << pair_index << " " << mumdex.n_pairs() << " "
          << m - pair.mums_start() << " "
          << mumdex.mums_stop(pair_index) - pair.mums_start() << " "
          << m << " " << mumdex.n_mums() << " "
          << mum.read_2() << " " << mum.flipped() << " "
          << ref.name(mum.chromosome()) << " "
          << mum.chromosome() << " " << mum.position0() << " "
          << ref.size(mum.chromosome()) << " "
          << mum.offset() << " " << mum.length() << " "
          << pair.length(mum.read_2()) << " "
          << message << endl;
          for (uint64_t m2 = pair.mums_start();
               m2 != mumdex.mums_stop(pair_index); ++m2) {
            const auto mum2 = mumdex.mum(m2);
            if (m2 == m) continue;
            const auto seq = ref.subseq(mum2.chromosome(), mum2.position0(),
                                        mum2.position0() + mum2.length());
            cout << mum2.read_2() << " " << mum2.flipped() << " "
                 << "-e " << seq << " "
                 << "-e " << paa::reverse_complement(seq) << endl;
          }
        };

        if (mum.offset() >= pair.length(mum.read_2()))
          report("mum offset out of range");
        if (mum.length() > pair.length(mum.read_2()))
          report("mum length out of range");
        if (mum.chromosome() >= ref.n_chromosomes())
          report("mum chromosome out of range");
        if (mum.position0() >= ref.size(mum.chromosome()))
          report("mum position out of range ");

        // const auto offset = ref.offset(mum.chromosome());
        // const auto abspos = offset + mum.position0();
        // const auto min_map = min(
        //    map.in(abspos), map.out(abspos + mum.length() - 1));
        if (mum.last_hit()) break;
      }
    }
  }

  return 0;
} catch (Error & e) {
  cerr << "paa::Error:" << endl;
  cerr << e.what() << endl;
  return 1;
} catch (exception & e) {
  cerr << "std::exception" << endl;
  cerr << e.what() << endl;
  return 1;
} catch (...) {
  cerr << "unknown exception was caught" << endl;
  return 1;
}
```
