## Supplementary material for "G-Graph: An interactive genomic graph viewer": MUMdex Source Code for G-Graph: chr_genome_repeats.sh

```
#! /bin/bash

. ~/mumdex/utility/shell.sh

# parse arguments
ref="$1"
[ "$ref" != "" ] || error no reference passed
[ -e $ref ] || error reference $ref does not exist
chr="$2"
[ "$chr" != "" ] || error no chr passed
len="$3"
[ "$len" != "" ] || error no length passed
warn running on $(hostname -s) with args $ref $chr $len
n_threads="$4"
[ "$n_threads" != "" ] || error no n_threads passed
warn running on $(hostname -s) with args $ref $chr $len $n_threads

# compile local code
start=$(date +%s)
local_mumdex=mumdex.$chr
rsync -a ~/mumdex/ $local_mumdex || warn rsync problem
(
    cd $local_mumdex && make clean
    make -j 4 find_repeats
) > /dev/null || error cannot compile mumdex code
export PATH=$local_mumdex:~/bin:$PATH
compile=$(date +%s)
warn compilation finished

# run find_repeats
find_repeats $ref $chr 0 $len $n_threads &&
warn find_repeats finished successfully ||
error find_repeats failed
repeats=$(date +%s)

# clean up local mumdex
rm -Rf $local_mumdex || warn problem cleaning up $local_mumdex
clean=$(date +%s)

# report timing
warn timing compile $((compile - start)) \
    find_repeats $((repeats - compile)) \
    clean $((clean - repeats)) \
    all $((clean - start))

exit 0
```
