## Supplementary material for "G-Graph: An interactive genomic graph viewer": MUMdex Source Code for G-Graph: colsplit.cpp

```
//
// colsplit
//
// split a table by columns
//
// Copyright 2018 Peter Andrews @ CSHL
//

// command to use, all on one line:
// (echo -n chr pos" " ; cat sampleIds.txt | perl -pe 's/,/ /g' ;
//  zcat baseCntStat-chr10.bgz) | time ~/mumdex/colsplit

#include <iostream>
#include <fstream>
#include <memory>
#include <sstream>
#include <string>
#include <vector>

using std::cerr;
using std::cin;
using std::cout;
using std::endl;
using std::exception;
using std::istringstream;
using std::ofstream;
using std::string;
using std::unique_ptr;
using std::vector;

#include "error.h"
#include "utility.h"
using paa::Error;

int main(int argc, char **) try {
  if (--argc != 0) throw Error("usage: colsplit");

  // make i/o faster
  std::ios_base::sync_with_stdio(false);
  cin.tie(nullptr);

  // column names
  const vector<string> column_names{[]() {
      string text;
      getline(cin, text);
      istringstream header_stream{text.c_str()};
      vector<string> result;
      while (header_stream >> text) result.push_back(text);
      return result;
    }()};

  // output files
  vector<unique_ptr<ofstream>> outputs;
  for (const string name : column_names)
    outputs.push_back(std::make_unique<ofstream>(name.c_str()));

  // process data
  string field;
  uint64_t n{0};
  while (cin >> field) (*outputs[n++ % column_names.size()]) << field << '\n';

  if (n % column_names.size() != 0) cerr << "Warning: uneven columns" << endl;

  return 0;
} catch (Error & e) {
  cerr << "paa::Error:" << endl;
  cerr << e.what() << endl;
  return 1;
} catch(exception & e) {
  cerr << e.what() << endl;
  return 1;
} catch(...) {
  cerr << "Some exception was caught." << endl;
  return 1;
}
```
