## Supplementary material for "G-Graph: An interactive genomic graph viewer": MUMdex Source Code for G-Graph: combine_finebins.cpp

```
//
// combine_finebins
//
// load up finebin files and combine them
//
// Copyright 2017 Peter Andrews @ CSHL
//

#include <exception>
#include <iostream>
#include <string>
#include <vector>

#include "cn.h"
#include "error.h"
#include "files.h"
#include "mumdex.h"

using std::cerr;
using std::cout;
using std::endl;
using std::exception;
using std::string;
using std::vector;

using paa::Error;
using paa::FinestBins;
using paa::Reference;

int main(int argc, char* argv[])  try {
  // Check usage
  if (--argc < 3)
    throw Error("usage: combine_finebins ref out_name bin_file ...");

  // Process command line arguments
  const Reference ref{argv[1]};
  const string out_name{argv[2]};

  // Sum of all bin counts
  FinestBins combined_counts{ref};

  argc -= 2;
  argv += 2;

  // Bin file names
  vector<string> bin_names{[argc, argv]() {
      vector<string> result;
      result.reserve(argc);
      for (int s{0}; s != argc; ++s) {
        result.emplace_back(argv[s + 1]);
      }
      return result;
    }()};

  for (const string & name : bin_names) {
    FinestBins bins{ref, name};
    const bool is_female{bins.n_y() == 0};
    if (is_female) {
      if ((bins.n_x() % 2) != 0)
        throw Error("Bad females") << bins.n_x()<< name;
    } else {
      if (bins.n_y() != bins.n_x())
        throw Error("Bad males") << bins.n_x() << bins.n_y() << name;
    }
    cout << "Loaded " << name
         << " " << bins.n_samples()
         << " " << bins.n_x()
         << " " << bins.n_y()
         << " " << is_female
         << endl;
    combined_counts.add(bins);
  }
  cout << "Saving combined bins" << endl;
  combined_counts.save(out_name + ".combined.bin");

  return 0;
} catch (Error & e) {
  cerr << "paa::Error:" << endl;
  cerr << e.what() << endl;
  return 1;
} catch (exception & e) {
  cerr << "std::exception" << endl;
  cerr << e.what() << endl;
  return 1;
} catch (...) {
  cerr << "unknown exception was caught" << endl;
  return 1;
}
```
