## Supplementary material for "G-Graph: An interactive genomic graph viewer": MUMdex Source Code for G-Graph: compare_segmentation.cpp

```
//
// compare_segmentation
//
// Compare segmentation from two duplicated cn runs
//
// Copyright 2017 Peter Andrews @ CSHL
//

#include <climits>

#include <algorithm>
#include <exception>
#include <fstream>
#include <iostream>
#include <map>
#include <memory>
#include <sstream>
#include <string>
#include <utility>
#include <vector>

#include "cn.h"
#include "cngsl.h"
#include "error.h"
#include "mumdex.h"
#include "psplot.h"
#include "strings.h"

using std::cout;
using std::cerr;
using std::endl;
using std::exception;
using std::ifstream;
using std::make_unique;
using std::map;
using std::max;
using std::min;
using std::ofstream;
using std::ostringstream;
using std::pair;
using std::string;
using std::unique_ptr;
using std::vector;

using paa::Bin;
using paa::Bounds;
using paa::ChromosomeIndexLookup;
using paa::CN_Bins;
using paa::CNStateCall;
using paa::CNStateCaller;
using paa::Error;
using paa::Marker;
using paa::PSDoc;
using paa::PSPage;
using paa::PSGraph;
using paa::PSSeries;
using paa::PSXYSeries;
using paa::PSXYDSeries;
using paa::PSXYMSeries;
using paa::Reference;
using paa::Segment;
using paa::Segments;

class OverlapInfo {
 public:
  OverlapInfo(const unsigned int n_bins_,
              const unsigned int chr_,
              const unsigned int start_,
              const double cn1_,
              const double cn2_,
              const bool pass_score_,
              const bool calls_agree_) :
      n_bins{n_bins_}, chr{chr_}, start{start_},
    cn1{cn1_}, cn2{cn2_},
    pass_score{pass_score_}, calls_agree{calls_agree_} { }
  unsigned int n_bins{0};
  unsigned int chr{0};
  unsigned int start{0};
  double cn1{0.0};
  double cn2{0.0};
  bool pass_score{false};
  bool calls_agree{false};
};

int main(int argc, char * argv[]) try {
  --argc;
  if (argc != 6)
    throw Error("usage: compare_segmentation ref bins file1 file2 name1 name2");

  const Reference ref{argv[1]};
  const ChromosomeIndexLookup chr_lookup{ref};
  const string bins_name{argv[2]};
  const vector<Bin> all_bins{load_bins(bins_name, ref)};

  // List of previously determined good bins
  const vector<Bin> bins{[&all_bins]() {
      vector<Bin> result;
      for (const Bin & bin : all_bins) {
        if (!bin.bad()) {
          result.push_back(bin);
        }
      }
      return result;
    }()};

  // Load data
  const Segments segs1{argv[3], argv[5], ref, chr_lookup};
  const Segments segs2{argv[4], argv[6], ref, chr_lookup};
  const bool flip{segs1.n_segs_good_event() > segs2.n_segs_good_event()};
  const Segments & low_segs{flip ? segs2 : segs1};
  const Segments & high_segs{flip ? segs1 : segs2};

  unsigned int bins_good[2]{0, 0};
  unsigned int bins_match[2]{0, 0};
  unsigned int segs_good[2]{0, 0};
  unsigned int segs_match[2]{0, 0};
  for (const bool swap : {false, true}) {
    const Segments & segsA{swap ? segs1 : segs2};
    const Segments & segsB{swap ? segs2 : segs1};
    for (const Segment & segA : segsA) {
      if (segA.called_event()) {
        ++segs_good[swap];
        bins_good[swap] += segA.n_bins;
        unsigned int n_overlap{0};
        for (unsigned int s{segsB.starts()[segA.chr]};
             s != segsB.starts()[segA.chr + 1]; ++s) {
          const Segment & segB{segsB[s]};
          if (segA.type != segB.type) continue;
          if (segB.stop <= segA.start) continue;
          if (segB.start >= segA.stop) break;
          if (segB.start <= segA.start &&
              segB.stop >= segA.stop) {
            n_overlap += segA.n_bins;
          } else if (segB.start <= segA.start &&
              segB.stop <= segA.stop) {
            n_overlap += segB.stop - segA.start;
          } else if (segB.start >= segA.start &&
              segB.stop <= segA.stop) {
            n_overlap += segB.n_bins;
          } else if (segB.start <= segA.stop &&
              segB.stop >= segA.stop) {
            n_overlap += segA.stop - segB.start;
          } else {
            throw Error("Unexpected overlap");
          }
        }
        bins_match[swap] += n_overlap;
        if ((segA.n_bins <= 5 && n_overlap) ||
            (segA.n_bins > 5 && n_overlap * 5 > 3 * segA.n_bins)) {
          ++segs_match[swap];
        }
      }
    }
  }
  cout << "comp2 " << segs1.name() << " " << segs2.name();
  double max_bins{0.0};
  double max_segs{0.0};
  unsigned int max_n_segs{0};
  for (const bool swap : {false, true}) {
    const double frac_bins{1.0 * bins_match[swap] / bins_good[swap]};
    const double frac_segs{1.0 * segs_match[swap] / segs_good[swap]};
    max_bins = max(frac_bins, max_bins);
    if (frac_segs > max_segs) max_n_segs = segs_good[swap];
    max_segs = max(frac_segs, max_segs);
    if (0)
      cout << " " << bins_good[swap]
           << " " << frac_bins
           << " " << segs_good[swap]
           << " " << frac_segs;
  }
  cout << " " << max_segs << " " << max_bins << " " << max_n_segs << endl;

  {
    double avg_bins{0.0};
    double avg_segs{0.0};
    unsigned int n_swap_bins{0};
    unsigned int n_swap_segs{0};
    unsigned int n_segs{0};
    for (const bool swap : {false, true}) {
      if (bins_good[swap]) {
        const double frac_bins{1.0 * bins_match[swap] / bins_good[swap]};
        avg_bins += frac_bins;
        ++n_swap_bins;
      }
      if (segs_good[swap]) {
        n_segs += segs_good[swap];
        const double frac_segs{1.0 * segs_match[swap] / segs_good[swap]};
        avg_segs += frac_segs;
        ++n_swap_segs;
      }
    }
    if (n_swap_bins && n_swap_segs)
      cout << "comp3 " << segs1.name() << " " << segs2.name() << " "
           << avg_segs / n_swap_segs << " " << avg_bins / n_swap_bins << " "
           << 1.0 * n_segs / n_swap_segs << endl;
  }
  // Compare actual bin values...
  const string out_base{string("compare.") +
        low_segs.name() + "_vs_" + high_segs.name()};
  ofstream detailed{out_base + ".detailed.txt"};
  ostringstream comp5;
  double avg_bins{0.0};
  double avg_segs{0.0};
  unsigned int nn_segs{0};
  for (const bool swap : {false, true}) {
    const Segments & segsA{swap ? segs1 : segs2};
    const Segments & segsB{swap ? segs2 : segs1};
    const CNStateCaller caller{ref, bins, segsB.profile};
    unsigned int n_concordant{0};
    unsigned int n_called{0};
    unsigned int n_bins_concordant{0};
    unsigned int n_bins_called{0};
    for (const Segment & segA : segsA) {
      if (segA.called_event()) {
        ++n_called;
        n_bins_called += segA.n_bins;
        const CNStateCall call{caller.call(segA.start, segA.stop)};
        if (call.type == segA.type) {
          ++n_concordant;
          n_bins_concordant += segA.n_bins;
        }
        detailed << ref.name(segA.chr)
                 << " " << segA.start
                 << " " << segA.stop
                 << " " << segA.n_bins
                 << " " << segA.score
                 << " " << segA.count
                 << " " << segA.cn
                 << " " << segA.expected
                 << " " << (call.type == segA.type)
                 << endl;
      }
    }
    if (0)
      comp5 << " " << n_called << " " << n_concordant << " "
            << n_bins_called << " " << n_bins_concordant;
    cout << "comp4 " << segsA.name() << " " << segsB.name() << " "
        // << n_called << " " << n_concordant << " "
        // << n_bins_called << " " << n_bins_concordant << " x x x x "
         << 1.0 * n_concordant / n_called << " "
         << 1.0 * n_bins_concordant / n_bins_called << " "
         << n_called << endl;
    avg_segs += 1.0 * n_concordant / n_called;
    avg_bins += 1.0 * n_bins_concordant / n_bins_called;
    nn_segs += n_called;
  }
  avg_bins /= 2;
  avg_segs /= 2;
  cout << "comp5 " << segs1.name() << " " << segs2.name() << " "
       << avg_segs << " " << avg_bins << " " << nn_segs / 2.0 << endl;

  // Plots and output
  ofstream out_points{(out_base + ".txt").c_str()};
  PSDoc ps{out_base, "concordance"};
  ps.pdf(false);
  PSGraph concordance_graph{ps, string("Concordance;") +
        low_segs.name() + " Segment CN;" +
        high_segs.name() + " Segment CN"};
  PSXYMSeries concordance_series{concordance_graph};
  PSGraph concordance_graph2{ps, string("Concordance;") +
        low_segs.name() + " Segment CN;" +
        high_segs.name() + " Segment CN"};
  PSXYMSeries concordance_series2{concordance_graph2};

  unsigned int s2{0};
  unsigned int overlap{0};
  vector<OverlapInfo> points;
  for (const Segment & seg1 : low_segs) {
    while (s2 != high_segs.size() &&
           high_segs[s2].stop <= seg1.start) {
      ++s2;
    }
    while (s2 != high_segs.size() &&
           high_segs[s2].start < seg1.stop) {
      const unsigned int start{max(high_segs[s2].start, seg1.start)};
      const unsigned int stop{min(high_segs[s2].stop, seg1.stop)};
      points.emplace_back(stop - start,
                          seg1.chr,
                          max(seg1.startpos, high_segs[s2].startpos),
                          seg1.cn, high_segs[s2].cn,
                          seg1.pass_score() && high_segs[s2].pass_score(),
                          seg1.type == high_segs[s2].type);
      overlap += stop - start;
      if (high_segs[s2].stop > seg1.stop) break;
      ++s2;
    }
  }
  if (overlap != low_segs.n_bins())
    throw Error("Overlap does not equal number of bins");

  auto bigger_first = [](const OverlapInfo & lhs, const OverlapInfo & rhs) {
    return lhs.n_bins > rhs.n_bins;
  };

  sort(points.begin(), points.end(), bigger_first);

  const unsigned int x_chr{ref.find_x_chromosome()};
  const unsigned int y_chr{ref.find_y_chromosome()};

  for (const OverlapInfo & conc : points) {
    const double scale{0.3 + log10(conc.n_bins)};
    const string color{conc.calls_agree ? "0 1 0" :
          (conc.pass_score ? "1 0 0" : "0 0 1")};
    const Marker marker{paa::circle(), scale, "0 0 0", 0.6, true, color};
    concordance_series.add_point(conc.cn1, conc.cn2, marker);
    out_points << ref.name(conc.chr) << " " << conc.start << " "
               << conc.n_bins << " " << conc.cn1 << " "
               << conc.cn2 << endl;
    if (conc.chr != x_chr && conc.chr != y_chr) {
      concordance_series2.add_point(conc.cn1, conc.cn2, marker);
    }
  }

  vector<unique_ptr<PSGraph>> graphs;
  vector<unique_ptr<PSXYMSeries>> series;

  for (const bool flip2 : {false, true}) {
    const Segments & segsA{flip2 ? high_segs : low_segs};
    const Segments & segsB{flip2 ? low_segs : high_segs};
    vector<OverlapInfo> segcomp;
    for (const Segment & segA : segsA) {
      double ratios{0.0};
      for (unsigned int b{segA.start}; b != segA.stop; ++b) {
        ratios += segsB.profile[b].ratio();
      }
      ratios *= 2.0 / segA.n_bins;
      segcomp.emplace_back(segA.n_bins,
                           segA.chr,
                           segA.startpos,
                           segA.cn, ratios,
                           segA.pass_score(),
                           false);
    }
    sort(segcomp.begin(), segcomp.end(), bigger_first);

    graphs.emplace_back(make_unique<PSGraph>(
        ps, string() + " Concordance;" +
        segsA.name() + " CN;" + segsB.name() + " CN"));
    series.emplace_back(make_unique<PSXYMSeries>(*graphs.back()));
    for (const OverlapInfo & conc : segcomp) {
      const double scale{0.3 + log10(conc.n_bins)};
      const string color{conc.pass_score ? "1 0 0" : "0 0 1"};
      const Marker marker{paa::circle(), scale, "0 0 0", 0.6, true, color};
      if (conc.chr != x_chr && conc.chr != y_chr) {
        series.back()->add_point(conc.cn1, conc.cn2, marker);
      }
    }
  }

  PSGraph bin_graph{ps, string("Bin Concordance;") +
        low_segs.name() + " Segment CN;" +
        high_segs.name() + " Segment CN", Bounds{0.0, 4.0, 0.0, 4.0}};
  PSXYDSeries bin_series{bin_graph, 50, 50};
  for (unsigned int b{0}; b != low_segs.profile.size(); ++b) {
    bin_series.add_point(low_segs.profile[b].ratio() * 2,
                         high_segs.profile[b].ratio() * 2);
  }

  for (const Segments * const seg : {&low_segs, &high_segs}) {
    cout << *seg << endl;
  }

  unsigned int n_segments_good2{0};
  unsigned int n_bins_good2{0};
  unsigned int n_events_concordant{0};
  unsigned int segment_bins_concordant{0};

  for (const Segment & seg1 : low_segs) {
    if (!seg1.called_event() || fabs(seg1.expected - seg1.cn) < 0.7) continue;
    unsigned int total_overlap{0};
    unsigned int segment_good_overlap{0};
    unsigned int segment_event2_overlap{0};
    for (unsigned int s{high_segs.starts()[seg1.chr]};
         s != high_segs.starts()[seg1.chr + 1]; ++s) {
      const Segment & seg2{high_segs[s]};
      const unsigned int n_overlap{[&seg1, &seg2]() {
          if (seg1.start <= seg2.start) {
            if (seg1.stop >= seg2.stop) {
              return seg2.n_bins;
            } else if (seg1.stop >= seg2.start) {
              return seg1.stop - seg2.start;
            }
          } else if (seg1.start <= seg2.stop) {
            if (seg1.stop >= seg2.stop) {
              return seg2.stop - seg1.start;
            } else {
              return seg1.n_bins;
            }
          }
          return 0U;
        }()};
      if (n_overlap == 0) continue;
      total_overlap += n_overlap;
      if (seg2.pass_score()) {
        segment_good_overlap += n_overlap;
        if (seg1.type == seg2.type) {
          segment_event2_overlap += n_overlap;
        }
      }
    }
    if (total_overlap != seg1.n_bins)
      throw Error("Bad overlaps") << total_overlap << seg1.n_bins;

    if (segment_good_overlap) {
      ++n_segments_good2;
      n_bins_good2 += segment_good_overlap;
      if (2 * segment_event2_overlap >= segment_good_overlap) {
        ++n_events_concordant;
        segment_bins_concordant += segment_event2_overlap;
      }
    }
  }

  if (0)
  cout << "comp"
       << " " << low_segs.name()
       << " " << high_segs.name()
       << " " << static_cast<int64_t>(low_segs.total_count())
       << " " << static_cast<int64_t>(high_segs.total_count())
       << " " << low_segs.n_segs_good_event()
       << " " << n_segments_good2
       << " " << 1.0 * n_segments_good2 / low_segs.n_segs_good_event()
       << " " << n_events_concordant
       << " " << 1.0 * n_events_concordant / n_segments_good2
       << " " << segment_bins_concordant
       << " " << 1.0 * segment_bins_concordant / n_bins_good2
       << endl;

  return 0;
} catch (Error & e) {
  cout << "paa::Error:" << endl;
  cout << e.what() << endl;
  return 1;
}
catch (exception & e) {
  cout << "std::exception" << endl;
  cout << e.what() << endl;
  return 1;
} catch (...) {
  cout << "unknown exception was caught" << endl;
  return 1;
}
```
