## Supplementary material for "G-Graph: An interactive genomic graph viewer": MUMdex Source Code for G-Graph: control_mview.cpp

```
//
// control_mview.cpp
//
// test controlling mview GUI from other app
//
// Copyright 2017 Peter Andrews @ CSHL
//

#include <exception>
#include <iostream>
#include <string>

#include "error.h"
#include "mumdex.h"
#include "x11plot.h"

using std::cerr;
using std::endl;
using std::exception;

using paa::ChromosomeIndexLookup;
using paa::Error;
using paa::Reference;

int main(int argc, char* argv[]) try {
  // Process optional command line arguments
  --argc;

  if (argc != 4)
    throw Error("usage: control_mview ref window chr pos");

  // Process arguments
  const Reference ref{argv[1]};
  const ChromosomeIndexLookup lookup{ref};
  const unsigned int win{static_cast<unsigned int>(atoi(argv[2]))};
  const std::string chr_name{argv[3]};
  const unsigned int chr{lookup[chr_name]};
  const unsigned int pos{static_cast<unsigned int>(atoi(argv[4]))};

  // Open connection to X server
  Display * display{XOpenDisplay(nullptr)};

  XEvent event;
  event.type = ClientMessage;
  event.xclient.serial = 0;
  event.xclient.send_event = true;
  event.xclient.format = 32;
  event.xclient.window = win;
  event.xclient.data.l[0] = 1000;
  event.xclient.data.l[1] = chr;
  event.xclient.data.l[2] = pos;

  XSendEvent(display, win, true, ClientMessage, &event);

  XFlush(display);

  return 0;
} catch (Error & e) {
  cerr << "paa::Error:" << endl;  cerr << e.what() << endl;
  return 1;
} catch (exception & e) {
  cerr << "std::exception" << endl;
  cerr << e.what() << endl;
  return 1;
} catch (...) {
  cerr << "unknown exception was caught" << endl;
  return 1;
}
```
