## Supplementary material for "G-Graph: An interactive genomic graph viewer": MUMdex Source Code for G-Graph: count_anchors.cpp

```
//
// count_anchors
//
// extract reference and anchor counts from a mumdex
//
// Copyright 2015 Peter Andrews @ CSHL
//

#include <exception>
#include <string>

#include "anchors.h"
#include "bed.h"
#include "error.h"
#include "mumdex.h"

using std::exception;
using std::cerr;
using std::endl;
using std::string;

using paa::AnchorCountsCreator;
using paa::BedFile;
using paa::Error;
using paa::MUMdex;

int main(int argc, char* argv[]) try {
  if (--argc != 3) throw Error("usage: count_anchors bed mumdexfile out_name");

  const string bed_name{argv[1]};
  const BedFile bed{bed_name};

  const string mumdex_name{argv[2]};
  const MUMdex mumdex{mumdex_name};

  const string out_name{argv[3]};
  const auto counts_dir = mumdex_name + "/" + out_name;

  const AnchorCountsCreator creator(mumdex, bed, counts_dir);

  return 0;
} catch (Error & e) {
  cerr << "paa::Error:" << endl;
  cerr << e.what() << endl;
  return 1;
} catch (exception & e) {
  cerr << "std::exception" << endl;
  cerr << e.what() << endl;
  return 1;
} catch (...) {
  cerr << "unknown exception was caught" << endl;
  return 1;
}
```
