## Supplementary material for "G-Graph: An interactive genomic graph viewer": MUMdex Source Code for G-Graph: count_genes.cpp

```
//
// count_genes.cpp
//
// How much of the genome do exons and genes cover?
//
// Copyright 2016 Peter Andrews @ CSHL
//

#include <iostream>
#include <string>
#include <vector>

#include "error.h"
#include "genes.h"
#include "mumdex.h"
#include "utility.h"

using std::cerr;
using std::endl;
using std::exception;
using std::string;
using std::vector;

using paa::sout;
using paa::ChromosomeIndexLookup;
using paa::Error;
using paa::GeneXrefs;
using paa::KnownGene;
using paa::KnownGenes;
using paa::Reference;

int main(int argc, char ** argv) {
  try {
    paa::exit_on_pipe_close();
    if (--argc != 1) throw Error("usage: count_genes reference");

    const string reference_file{argv[1]};
    const Reference ref{reference_file};
    const ChromosomeIndexLookup lookup{ref};
    const string genes_name{reference_file + ".bin/knownGene.txt"};
    const string isoforms_name{reference_file + ".bin/knownIsoforms.txt"};
    const string kgXrefs_name{reference_file + ".bin/kgXref.txt"};
    const KnownGenes genes{genes_name, isoforms_name, lookup, ref};
    const GeneXrefs xref{kgXrefs_name};

    vector <uint8_t> gene_cover(ref.size());
    vector <uint8_t> exon_cover(ref.size());
    vector <uint8_t> middle_exon_cover(ref.size());
    for (const KnownGene & gene : genes) {
      for (unsigned int b{ref.abspos(gene.chr, gene.t_start)};
           b != ref.abspos(gene.chr, gene.t_stop); ++b) {
        gene_cover[b] = 1;
      }
      for (unsigned int e{0}; e != gene.exon_starts.size(); ++e) {
        for (unsigned int b{ref.abspos(gene.chr, gene.exon_starts[e])};
           b != ref.abspos(gene.chr, gene.exon_stops[e]); ++b) {
          exon_cover[b] = 1;
          if (e && e + 1 != gene.exon_starts.size()) middle_exon_cover[b] = 1;
        }
      }
    }

    unsigned int n_gene{0};
    unsigned int n_exon{0};
    unsigned int n_middle_exon{0};
    for (unsigned int b{0}; b != ref.size(); ++b) {
      n_gene += gene_cover[b];
      n_exon += exon_cover[b];
      n_middle_exon += middle_exon_cover[b];
    }

    sout << n_gene << n_gene / static_cast<double>(ref.size())
         << n_exon << n_exon / static_cast<double>(ref.size())
         << n_middle_exon << n_middle_exon / static_cast<double>(ref.size())
         << endl;

    return 0;
  }
  catch(exception & e) {
    cerr << e.what() << endl;
    return 1;
  }
  catch(...) {
    cerr << "Some exception was caught." << endl;
    return 1;
  }
  return 0;
}
```
