## Supplementary material for "G-Graph: An interactive genomic graph viewer": MUMdex Source Code for G-Graph: count_pseudogenes.cpp

```
//
// count_pseudogenes
//
// find pseudogenes and count them
//
// Copyright 2015 Peter Andrews @ CSHL
//

#include <algorithm>
#include <exception>
#include <iostream>
#include <map>
#include <string>
#include <vector>

#include "bed.h"
#include "error.h"
#include "genes.h"
#include "mumdex.h"
#include "utility.h"

using std::exception;
using std::cout;
using std::cerr;
using std::endl;
using std::map;
using std::string;
using std::vector;

using paa::BedFile;
using paa::ChromosomeIndexLookup;
using paa::Error;
using paa::MUMdex;
using paa::JunctionCounts;
using paa::JunctionSave;
using paa::JunctionSorter;
using paa::KnownGenes;
using paa::PosInfo;
using paa::Reference;
using paa::serr;
using paa::sout;
using paa::terr;
using paa::tout;

int main(int argc, char* argv[]) try {
  paa::exit_on_pipe_close();
  paa::memory_mapped = false;

  if (--argc != 3)
    throw Error("usage: count_pseudogenes mumdex genes isoforms");

  const string mumdex_name{argv[1]};
  const MUMdex mumdex{mumdex_name};

  const Reference & ref = mumdex.reference();
  const ChromosomeIndexLookup chr_lookup{ref};

  const string genes_name{argv[2]};
  const string isoforms_name{argv[3]};

  const KnownGenes genes{genes_name, isoforms_name, chr_lookup, ref};
  const BedFile bed{genes.generate_bed("junctions.bed")};
  JunctionCounts gene_counts(genes);

  if (0) {
    const bool print = true;
    for (auto jp : genes.junctions) {
      if (print) sout << ref.name(jp.chr) << jp.pos
                      << genes[jp.gene_index].name << jp.junction_index;
      for (auto inv : jp.invariants) {
        if (print) sout << inv;
      }
      if (print) sout << endl;
    }
  }

  int64_t n_mums = 0;
  int64_t n_acceptable_invariants = 0;
  const int64_t min_invariant = -100000;  // -2318857;
  const unsigned int position_window = 10;
  const int invariant_window = 10;
  const int ignore_deletions = -10;
  vector<uint64_t> seen_pairs;
  const auto & junctions = genes.junctions;
  for (unsigned int ji = 0; ji != junctions.size(); ++ji) {
    unsigned int n_junction = 0;
    seen_pairs.clear();
    map<JunctionSave, unsigned int> junctions_seen;

    // Get range of junctions with this junction start
    const unsigned int jb = ji;
    const unsigned int je = [jb, &junctions]() {
      unsigned int test = jb + 1;
      while (test != junctions.size() &&
             junctions[test].pos == junctions[jb].pos &&
             junctions[test].chr == junctions[jb].chr) {
        ++test;
      }
      return test;
    }();
    // cout << ji << " ";
    ji = je - 1;

    // Determine mum search range and find first mum
    const auto jchromosome = junctions[jb].chr;
    const auto jposition = junctions[jb].pos;
    if (0) cout << ji << " " << jb << " " << je << " "
                << jchromosome << " " << jposition << " "
                << junctions.size() << '\n';
    const auto low_pos = jposition >= position_window ?
        jposition - position_window : 0;
    const auto high_pos = jposition + position_window + 1;
    const auto mum_start_iter = mumdex.lower_bound(jchromosome, low_pos);
    for (auto mum_iter = mum_start_iter; mum_iter != mumdex.index().end();
         ++mum_iter) {
      const auto mum = mumdex.mum(*mum_iter);
      if (mum.position0() >= high_pos ||  mum.chromosome() != jchromosome) {
        break;
      }
      const auto pair = mumdex.pair(*mum_iter);
      if (pair.dupe()) continue;
      const auto pair_index = mum_iter->pair_index();
      if ([pair_index, &seen_pairs]() {
          for (const auto seen : seen_pairs) {
            if (seen == pair_index) {
              return true;
            }
          }
          return false;
        }()) {
        continue;
      }
      const auto mums_start = pair.mums_start();
      const auto pair_mum_index = mum_iter->mum_in_pair_index();
      const auto mum_index = mums_start + pair_mum_index;
      const uint64_t start_mum = mums_start +
          (mum.flipped() ? pair_mum_index + 1 : 0);
      const uint64_t stop_mum = mum.flipped() ?
          mumdex.mums_stop(pair_index) : mums_start + pair_mum_index;
      for (uint64_t mi = start_mum; mi != stop_mum; ++mi) {
        const auto other = mumdex.mum(mi);
        if (mum.read_2()) {
          if (!other.read_2()) continue;
        } else {
          if (other.read_2()) break;
        }
        if (other.chromosome() != mum.chromosome()) continue;
        if (other.flipped() != mum.flipped()) continue;
        const auto first = mum_index < mi ? mum : other;
        const auto second = mum_index < mi ? other : mum;
        const int invariant{static_cast<int>(
            first.flipped() ?
            (static_cast<int64_t>(second.position0()) + second.length() +
             second.offset() - first.position0() - first.length() -
             first.offset()) :
            (static_cast<int64_t>(first.position0()) - first.offset() -
             second.position0() + second.offset()))};
        if (invariant >= ignore_deletions) continue;
        if (invariant < min_invariant) continue;
        ++n_acceptable_invariants;
        seen_pairs.push_back(pair_index);
        int closest_invariant = 0;
        unsigned int n_checked = 0;
        for (unsigned int jc = jb; jc != je; ++jc) {
          const auto & junction = junctions[jc];
          for (unsigned int ii = 0; ii != junction.invariants.size(); ++ii) {
            const int inv{junction.invariants[ii]};
            ++n_checked;
            if (invariant >= inv - invariant_window &&
                invariant < inv + invariant_window) {
              const JunctionSave key{jc, ii, invariant, mum.position0()};
              ++junctions_seen[key];
              if (abs(inv - invariant) <
                  abs(invariant - closest_invariant)) {
                closest_invariant = inv;
              }
            }
          }
        }
        if (closest_invariant) {
          if (0)
            serr << "invariant" << invariant << closest_invariant
                 << n_checked << ji << jposition << "at"
                 << ref.name(mum.chromosome()) << mum.position0()
                 << "for pair" << pair_index << "and read" << mum.read_2() + 1
                 << mum.flipped() << endl;
          ++n_junction;
          break;
        }
      }
      ++n_mums;
      if (0)
        sout << ji << jchromosome  << jposition
             << mum.position0() << pair_index
             << pair_mum_index << '\n';
    }
    if (n_junction) {
      for (auto js = junctions_seen.begin(); js != junctions_seen.end();
           ++js) {
        sout << js->first.junction_index << js->first.invariant_index
             << js->first.invariant << js->first.position << js->second
             << endl;
      }
    }
  }
  serr << "pairs" << mumdex.pairs().size()
       << "mums" << mumdex.mums().size()
       << "ok_mums" << n_mums
       << "n_acceptable_invariants" << n_acceptable_invariants
       << '\n';

  return 0;
} catch (Error & e) {
  cerr << "paa::Error:" << endl;
  cerr << e.what() << endl;
  return 1;
} catch (exception & e) {
  cerr << "std::exception" << endl;
  cerr << e.what() << endl;
  return 1;
} catch (...) {
  cerr << "unknown exception was caught" << endl;
  return 1;
}
```
