## Supplementary material for "G-Graph: An interactive genomic graph viewer": MUMdex Source Code for G-Graph: debruijn.cpp

```
//
// debruijn
//
// read pair assembly and viewing
//
// Copyright 2015 Peter Andrews @ CSHL
//

#include <algorithm>
#include <exception>
#include <iostream>
#include <fstream>
#include <functional>
#include <limits>
#include <list>
#include <map>
#include <random>
#include <set>
#include <sstream>
#include <string>
#include <utility>
#include <vector>

#include "error.h"
#include "genes.h"
#include "longSA.h"
#include "mumdex.h"
#include "utility.h"

using std::bind;
using std::cout;
using std::cerr;
using std::endl;
using std::exception;
using std::ifstream;
using std::ofstream;
using std::list;
using std::make_pair;
using std::map;
using std::multimap;
using std::min;
using std::max;
using std::max_element;
using std::mt19937_64;
using std::next;
using std::numeric_limits;
using std::ostringstream;
using std::pair;
using std::set;
using std::string;
using std::to_string;
using std::uniform_int_distribution;
using std::vector;

using paa::ChromosomeIndexLookup;
using paa::Error;
using paa::GeneFinder;
using paa::GeneXrefs;
using paa::KnownGenes;
using paa::Mappability;
using paa::PosInfo;
using paa::Reference;
using paa::SimpleHit;
using paa::longSA;
using paa::mkdir;
using paa::reverse_complement;
using paa::serr;
using paa::sout;

const string bases{"ACTG"};  // NOLINT

vector<string> left_nodes(const string & node) {
  const string substr = node.substr(0, node.size() - 1);
  vector<string> left;
  left.reserve(4);
  for (const auto base : bases) {
    string grow;
    grow += base;
    grow += substr;
    left.push_back(grow);
  }
  return left;
}

vector<string> right_nodes(const string & node) {
  const string substr = node.substr(1, node.size() - 1);
  vector<string> right;
  right.reserve(4);
  for (const auto base : bases) {
    right.push_back(substr + base);
  }
  return right;
}

template <class T>
struct less_ptr {
  bool operator()(const T & lhs, const T & rhs) const {
    return &*lhs < &*rhs;
  }
};

template <class T>
struct more {
  bool operator()(const T & lhs, const T & rhs) const {
    return lhs > rhs;
  }
};

class Node {
 public:
  explicit Node(const string & sequence_arg, const unsigned int count_arg = 0) :
      sequence{sequence_arg}, count{count_arg} {}
  using iter = list<Node>::iterator;
  using sset = set<iter, less_ptr<iter>>;
  using smap = map<iter, iter, less_ptr<iter>>;
  string sequence;
  unsigned int count;
  sset in_nodes{};
  sset out_nodes{};
  unsigned int contig{0};
  bool main_path{false};
  bool operator<(const Node & other) const {
    return sequence < other.sequence;
  }
  unsigned int mark_nodes(const unsigned int value) {
    if (contig) return 0;
    contig = value;
    unsigned int bases_seen{static_cast<unsigned int>(sequence.size())};
    const vector<sset *> all_links{&in_nodes, &out_nodes};
    for (const auto links : all_links) {
      for (const auto link : *links) {
        bases_seen += link->mark_nodes(value);
      }
    }
    return bases_seen;
  }
  unsigned int count_right(sset & seen) const {
    unsigned int bases_seen{static_cast<unsigned int>(sequence.size())};
    for (const auto link : out_nodes) {
      if (seen.count(link)) continue;
      seen.insert(link);
      bases_seen += link->count_right(seen);
    }
    return bases_seen;
  }
};

unsigned int collapse_nodes(list<Node> & nodes) {
  unsigned int n_changes = 0;
  for (auto node_iter = nodes.begin(); node_iter != nodes.end(); ++node_iter) {
    auto & node = *node_iter;
    while (node.out_nodes.size() == 1) {
      auto next_iter = *node.out_nodes.begin();
      if (node_iter == next_iter) break;
      auto & next = *next_iter;
      if (next.in_nodes.size() != 1) break;
      if (next.count > node.count) node.count = next.count;
      node.sequence += next.sequence;
      node.out_nodes = next.out_nodes;
      for (auto following : node.out_nodes) {
        following->in_nodes.erase(next_iter);
        following->in_nodes.insert(node_iter);
      }
      nodes.erase(next_iter);
      ++n_changes;
    }
  }
  return n_changes;
}

void simplify_graph(list<Node> & nodes) {
  unsigned int n_changes = 1;
  while (n_changes) {
    n_changes = 0;
    for (auto node_iter = nodes.begin(); node_iter != nodes.end();
         ++node_iter) {
      auto & node = *node_iter;
      if (node.in_nodes.size() < 2) continue;
      for (auto in1_iter = node.in_nodes.begin();
           in1_iter != node.in_nodes.end(); ++in1_iter) {
        auto in1 = *in1_iter;
        auto & seq1 = in1->sequence;
        if (seq1.empty()) continue;
        bool changed = false;
        for (auto in2_iter = next(in1_iter);
             in2_iter != node.in_nodes.end(); ++in2_iter) {
          auto in2 = *in2_iter;
          if (in1->out_nodes != in2->out_nodes) continue;
          auto & seq2 = in2->sequence;
          if (seq2.empty()) continue;
          unsigned int overlap = 1;
          for (; overlap <= seq1.size() && overlap <= seq2.size();
               ++overlap) {
            if (seq1[seq1.size() - overlap] != seq2[seq2.size() - overlap]) {
              break;
            }
          }
          if (--overlap) {
            Node new_node{seq1.substr(seq1.size() - overlap),
                  max(in1->count, in2->count)};
            seq1.erase(seq1.size() - overlap);
            seq2.erase(seq2.size() - overlap);
            new_node.in_nodes.insert(in1);
            new_node.in_nodes.insert(in2);
            new_node.out_nodes.insert(in1->out_nodes.begin(),
                                      in1->out_nodes.end());
            auto new_iter = nodes.insert(nodes.end(), new_node);
            for (auto out : new_node.out_nodes) {
              out->in_nodes.erase(in1);
              out->in_nodes.erase(in2);
              out->in_nodes.insert(new_iter);
            }
            in1->out_nodes.clear();
            in2->out_nodes.clear();
            in1->out_nodes.insert(new_iter);
            in2->out_nodes.insert(new_iter);
            changed = true;
            break;
          }
        }
        if (changed) break;
      }
    }
    for (auto node_iter = nodes.begin(); node_iter != nodes.end();
         ++node_iter) {
      auto & node = *node_iter;
      if (node.out_nodes.size() < 2) continue;
      for (auto out1_iter = node.out_nodes.begin();
           out1_iter != node.out_nodes.end(); ++out1_iter) {
        auto out1 = *out1_iter;
        auto & seq1 = out1->sequence;
        if (seq1.empty()) continue;
        bool changed = false;
        for (auto out2_iter = next(out1_iter);
             out2_iter != node.out_nodes.end(); ++out2_iter) {
          auto out2 = *out2_iter;
          if (out1->in_nodes != out2->in_nodes) continue;
          auto & seq2 = out2->sequence;
          if (seq2.empty()) continue;
          unsigned int overlap = 0;
          for (; overlap <= seq1.size() && overlap <= seq2.size();
               ++overlap) {
            if (seq1[overlap] != seq2[overlap]) {
              break;
            }
          }
          if (overlap) {
            Node new_node{seq1.substr(0, overlap),
                  max(out1->count, out2->count)};
            seq1.erase(0, overlap);
            seq2.erase(0, overlap);
            new_node.out_nodes.insert(out1);
            new_node.out_nodes.insert(out2);
            new_node.in_nodes.insert(out1->in_nodes.begin(),
                                     out1->in_nodes.end());
            auto new_iter = nodes.insert(nodes.end(), new_node);
            for (auto in : new_node.in_nodes) {
              in->out_nodes.erase(out1);
              in->out_nodes.erase(out2);
              in->out_nodes.insert(new_iter);
            }
            out1->in_nodes.clear();
            out2->in_nodes.clear();
            out1->in_nodes.insert(new_iter);
            out2->in_nodes.insert(new_iter);
            changed = true;
            break;
          }
        }
        if (changed) break;
      }
    }
    for (auto node = nodes.begin(); node != nodes.end(); ++node) {
      if (node->sequence.size()) continue;
      auto & ins = node->in_nodes;
      auto & outs = node->out_nodes;
      for (auto & in : ins) {
        in->out_nodes.erase(node);
      }
      for (auto & out : outs) {
        out->in_nodes.erase(node);
      }
      for (auto & in : ins) {
        for (auto & out : outs) {
          in->out_nodes.insert(out);
          out->in_nodes.insert(in);
        }
      }
    }
    nodes.remove_if([](const Node & node) { return node.sequence.empty(); });
    n_changes += collapse_nodes(nodes);
  }
}

// For contig gene annotation
struct GeneInfo {
  GeneInfo(const string & text_arg, const unsigned int offset_arg,
           const unsigned int length_arg, const bool exon_arg) :
      text{text_arg}, offset{offset_arg},
    length{length_arg}, exon{exon_arg} {}
  string text;
  unsigned int offset;
  unsigned int length;
  bool exon;
};

Node::iter create_main_path(list<Node> & nodes, Node::smap & seen_nodes,
                            const Node::iter previous, const Node::iter node) {
  // Do proper linking for previously seen nodes
  if (seen_nodes.count(node)) {
    if (previous != nodes.end()) {
      seen_nodes[node]->in_nodes.insert(previous);
      previous->out_nodes.insert(seen_nodes[node]);
    }
    return seen_nodes[node];
  }

  // Add new node to graph
  Node new_node = *node;
  new_node.main_path = true;
  new_node.contig = 0;
  new_node.in_nodes.clear();
  if (previous != nodes.end()) new_node.in_nodes.insert(previous);
  new_node.out_nodes.clear();
  const auto old_end = nodes.end();
  const auto new_iter = nodes.insert(nodes.end(), new_node);
  if (previous != old_end) previous->out_nodes.insert(new_iter);

  // Choose best continuation(s) for node
  pair<unsigned int, unsigned int> best_choice;
  vector<Node::iter> choices;
  for (auto out : node->out_nodes) {
    if (out->count >= best_choice.first) {
      Node::sset seen;
      if (out->count > best_choice.first) {
        choices.clear();
        best_choice = make_pair(out->count, out->count_right(seen));
      } else {
        const auto right_count = out->count_right(seen);
        if (right_count < best_choice.second) {
          continue;
        }
        if (right_count > best_choice.second) {
          choices.clear();
        }
        best_choice = make_pair(out->count, right_count);
      }
      choices.push_back(out);
    }
  }

  // Continue choosing process recursively
  for (const auto choice : choices) {
    create_main_path(nodes, seen_nodes, new_iter, choice);
  }
  return new_iter;
}

int main(int argc, char* argv[]) try {
  --argc;
  if (argc != 4 && argc != 7)
    throw Error("usage: debruijn reference sequences kmer clip"
                " [genes isoforms xrefs]");

  std::random_device rd;
  auto mersenne = mt19937_64(rd());
  auto dist = uniform_int_distribution<uint64_t>(33, 126);
  auto rand_char = bind(dist, mersenne);

  // Controls and settings
  const bool show_input = false;
  const bool show_kmers = false;
  const bool show_kmer_counts = false;
  const bool show_collapsed_kmers = false;

  // To find mums in nodes
  const string fasta{argv[1]};
  const Reference ref{fasta};
  const longSA sa{fasta.c_str(), true, true, false};
  const Mappability mappability{fasta, true};

  // Input sequence file
  ifstream sequences_file{argv[2]};

  // Kmer choice
  const unsigned int kmer{static_cast<unsigned int>(atoi(argv[3]))};

  // Clipping
  const unsigned int clip{static_cast<unsigned int>(atoi(argv[4]))};

  // Output directory
  const string out_dir{"assemblies"};
  mkdir(out_dir);

  while (sequences_file) {
    // Gene metadata, optional
    string sample;
    unsigned int gene_index;
    string gene_symbol;
    string mumdex_name;
    if (argc == 7) {
      sequences_file >> sample >> gene_index >> gene_symbol >> mumdex_name;
    }
    sequences_file.ignore(1000, '\n');

    // Read sequence into simple node structures
    string sequence;
    map<string, unsigned int> simple_nodes;
    unsigned int n_sequences = 0;
    while (getline(sequences_file, sequence)) {
      if (sequence.empty()) {
        sequences_file.ignore(1000, '\n');
        break;
      }
      ++n_sequences;
      if (sequence.size() < kmer) {
        serr << "Ignoring too small sequence" << sequence<< endl;
        continue;
      }
      if (show_input) sout << "Sequence read" << sequence << endl;
      if (show_kmers) serr << "Kmers";
      for (unsigned int b = 0; b != sequence.size() - kmer + 1; ++b) {
        const string subseq = sequence.substr(b, kmer);
        if (show_kmers) serr << subseq;
        if (subseq.find('N') != string::npos) {
          continue;
        }
        ++simple_nodes[subseq];
        string rcseq{reverse_complement(subseq)};
        ++simple_nodes[rcseq];
      }
      if (show_kmers) serr << endl;
    }
    serr << "Loaded" << n_sequences << "for" << gene_symbol << sample << endl;

    // Transfer simple nodes to linked node structure
    list<Node> nodes;
    for (const auto node : simple_nodes) {
      const auto seq = node.first;
      const auto count = node.second;
      if (count > clip) nodes.emplace_back(seq, count);
      if (show_kmer_counts) serr << seq << count << endl;
    }

    // Link nodes
    for (auto & node : nodes) {
      const auto left = left_nodes(node.sequence);
      for (const auto & test : left) {
        auto found = equal_range(nodes.begin(), nodes.end(), Node(test));
        if (found.first != found.second) {
          node.in_nodes.insert(found.first);
        }
      }
      const auto right = right_nodes(node.sequence);
      for (const auto & test : right) {
        auto found = equal_range(nodes.begin(), nodes.end(), Node(test));
        if (found.first != found.second) {
          node.out_nodes.insert(found.first);
        }
      }
    }

    // Remove redundant sequence from nodes
    for (auto & node : nodes) {
      if (node.in_nodes.size()) {
        node.sequence = node.sequence.substr(kmer - 1);
      }
    }

    // Collapse and simplify graph structure
    simplify_graph(nodes);

    // Show collapsed results
    if (show_collapsed_kmers) {
      for (const auto node : nodes) {
        const auto seq = node.sequence;
        const auto count = node.count;
        serr << seq << count << endl;
      }
    }

    // Determine contigs, get info for ordering display
    unsigned int contig = 0;
    vector<Node::iter> left_nodes;
    map<unsigned int, unsigned int> contig_sizes;
    map<Node::iter, unsigned int, less_ptr<Node::iter>> right_sizes;
    map<unsigned int, Node::iter> leftmost_nodes;
    for (auto node = nodes.begin(); node != nodes.end(); ++node) {
      if (node->in_nodes.empty() ||
          (node->in_nodes.size() == 1 && *node->in_nodes.begin() == node)) {
        left_nodes.push_back(node);
        Node::sset seen;
        right_sizes[node] = node->count_right(seen);
        if (node->contig == 0) {
          ++contig;
          contig_sizes[contig] = node->mark_nodes(contig);
        }
        if (leftmost_nodes.count(node->contig) == 0 ||
            right_sizes[leftmost_nodes[node->contig]] <
            right_sizes[node]) {
          leftmost_nodes[node->contig] = node;
        }
      }
    }
    multimap<unsigned int, Node::iter, more<unsigned int>> ordered_left;
    for (const auto elem : leftmost_nodes) {
      const auto node = elem.second;
      ordered_left.emplace(contig_sizes[node->contig], node);
    }

    // Extract main L->R path for each multi-node contig as special contigs
    list<Node> main_path;
    if (0) {
      for (const auto elem : leftmost_nodes) {
        const auto node = elem.second;
        Node::smap seen;
        const auto new_node =
            create_main_path(main_path, seen, main_path.end(), node);
        ordered_left.emplace(new_node->mark_nodes(++contig), new_node);
      }
      simplify_graph(main_path);
    }

    vector<list<Node> *> graphs{&nodes};  // , &main_path};
    const string output_name = out_dir + "/" + gene_symbol + "_" +
        sample + "_" + to_string(clip);
    vector<string> graph_names{output_name, output_name + "_main"};
    vector<string> graph_types{"All Pairs", "Main Paths"};
    vector<bool> printed_legend(graphs.size());
    vector<bool> main_path_type{0, 1};

    for (unsigned int g = 0; g != graphs.size(); ++g) {
      const auto & graph = *graphs[g];
      if (graph.empty()) continue;
      const auto graph_name = graph_names[g];

      // Output graph to graphviz format
      const string dot_name{graph_name + ".dot"};
      ofstream dot{dot_name.c_str()};
      ofstream primers_out{(graph_name + ".pri").c_str()};
      // Overall graph info
      dot << "digraph pseudogene {" << endl
          // << "graph[page=\"8.5,11.0\", size=\"7.5,10\",
          // center=1, ratio=\"fill\"]"
          << "rankdir=LR;" << endl
          << "node [fontname=Courier];" << endl
          << endl;

      // Group all contigs
      if (argc != 7) {
        dot << "name [color=blue, label=\"" << graph_types[g] << "\"];" << endl;
      }
      for (const auto elem : ordered_left) {
        const auto & node = *elem.second;
        if (node.main_path == main_path_type[g]) {
          dot << "\"name\" -> \"" << &node
              << "\" [dir=\"none\", weight=" << elem.first
              << ", tailport=e, headport=w, label=\""
              << elem.first << "\", color=blue, fontcolor=blue, style=dotted];"
              << endl;
        }
      }

      // Annotate and output each node
      for (const auto & node : graph) {
        const auto seq = node.sequence;
        auto mams = sa.find_mams(seq);

        vector<string> lines;
        if (argc == 7) {
          static const string genes_name{argv[5]};
          static const string isoforms_name{argv[6]};
          static const string kgXrefs_name{argv[7]};

          static const ChromosomeIndexLookup chr_lookup{ref};
          static const KnownGenes genes{genes_name, isoforms_name,
                chr_lookup, ref};
          static const GeneXrefs xref{kgXrefs_name};

          const auto & gene = genes[gene_index];

          // Output information on sample and gene as special node
          if (!printed_legend[g]) {
            dot << "name [label=\"" << sample << " " << gene_symbol
                << " " << gene.n_exons << " exons\\n"
                << graph_types[g] << "\", color=blue, fontcolor=blue];" << endl
                << endl;
            printed_legend[g] = true;
          }

          // How far into the pseudogene is each base within an exon?
          const auto exon_offsets = [&gene]() {
            vector<unsigned int> offsets;
            unsigned int n_bases = 0;
            for (unsigned int e = 0; e != gene.n_exons; ++e) {
              offsets.push_back(n_bases);
              n_bases += gene.exon_stops[e] - gene.exon_starts[e];
            }
            return offsets;
          }();

          // Determine exonic and intronic regions for node and save info
          vector<unsigned int> distance_in(seq.size());
          vector<unsigned int> exon_numbers(seq.size());
          vector<GeneInfo> gene_info;
          for (const auto & mam : mams) {
            const auto chromosome = mam.chr;
            if (chromosome != gene.chr) continue;
            if (mam.pos >= gene.t_stop) continue;
            if (mam.pos + mam.len < gene.t_start) continue;
            if (mam.dir != '+') continue;
            unsigned int m = 0;
            while (m != mam.len) {
              const auto p = mam.pos + m;
              unsigned int len = 0;
              bool exon = false;
              for (unsigned int e = 0; e != gene.n_exons; ++e) {
                string descr;
                if (mam.dir == '+') {
                  if (p >= gene.exon_starts[e] && p < gene.exon_stops[e]) {
                    len = min(gene.exon_stops[e] - p, mam.len - m);
                    descr = "Exon " + to_string(e + 1);
                    exon = true;
                  } else if (e + 1 != gene.n_exons) {
                    if (p >= gene.exon_stops[e] &&
                        p < gene.exon_starts[e + 1]) {
                      len = min(gene.exon_starts[e + 1] - p, mam.len - m);
                      descr = "Intron " + to_string(e + 1);
                    }
                  }
                }
                if (len) {
                  string text = len == 1 ? "|" :
                      (p == gene.exon_starts[e] ? "[" : "(");
                  text += descr;
                  if (text.size() < len - 1)
                    text.insert(text.end(), len - 1 - text.size(), ' ');
                  if (len > 1) text.insert(len - 1, 1,
                                           (p + len == gene.exon_stops[e] ?
                                            ']' : ')'));
                  gene_info.emplace_back(text, m + mam.off, len, exon);
                  if (exon) {
                    for (unsigned int b = 0; b != len; ++b) {
                      const unsigned int distance = exon_offsets[e] + p + b -
                          gene.exon_starts[e];
                      distance_in[mam.off + m + b] = distance;
                      exon_numbers[mam.off + m + b] = e + 1;
                    }
                  }
                  break;
                }
              }
              if (len) {
                m += len;
              } else {
                ++m;
              }
            }
          }

          // Annotate node with GC score
          const unsigned int primer_length = 25;
          const unsigned int required_excess = 5;
          vector<unsigned int> primer_scores(distance_in.size());
          vector<unsigned int> all_same(distance_in.size());
          vector<bool> exceeds(distance_in.size());
          vector<string> chromosomes(distance_in.size());
          vector<unsigned int> positions(distance_in.size());
          vector<char> strands(distance_in.size());
          for (unsigned int b1 = 0;
               b1 + primer_length < distance_in.size(); ++b1) {
            unsigned int score = 0;
            string seq_bases;
            all_same[b1] = true;
            for (unsigned int b2 = b1; b2 != b1 + primer_length &&
                     b2 != distance_in.size(); ++b2) {
              if (b2 != b1 && exon_numbers[b2] != exon_numbers[b2 - 1]) {
                all_same[b1] = false;
              }
              if (seq[b2] == 'G' || seq[b2] == 'G') {
                score += 4;
              } else {
                score += 2;
              }
              seq_bases += seq[b2];
            }
            primer_scores[b1] = score;
            auto seq_mams = sa.find_mams(seq_bases);
            if (seq_mams.size() == 1) {
              const auto mam = seq_mams.front();
              const auto offset = ref.offset(mam.chr);
              const auto abspos = offset + mam.pos;
              const auto max_map = max(
                  mappability.low(abspos),
                  mappability.high(abspos + mam.len - 1));
              if (mam.len >= max_map + required_excess)
                exceeds[b1] = true;
              chromosomes[b1] = ref.name(mam.chr);
              positions[b1] = mam.pos + 1;
              strands[b1] = mam.dir;
            }
          }

          // Annotate node with some numbers
          const bool primer_annotation = false;
          if (primer_annotation) {
            vector<vector<unsigned int> *> numbers
            {&primer_scores, &distance_in};
            for (const auto num_ptr : numbers) {
              const auto & nums = *num_ptr;
              const unsigned int max_num = *max_element(nums.begin(),
                                                        nums.end());
              const unsigned int n_digits = [](unsigned int value) {
                unsigned int digit = 0;
                while (value) {
                  ++digit;
                  value /= 10;
                }
                return digit;
              }(max_num);
              for (unsigned int d = 0; d != n_digits; ++d) {
                string line;
                const unsigned int place = n_digits - d - 1;
                for (unsigned int b = 0; b != nums.size(); ++b) {
                  const unsigned int digit = [](unsigned int plc,
                                                unsigned int value) {
                    while (plc--) {
                      value /= 10;
                    }
                    return value % 10;
                  }(place, nums[b]);
                  line += '0' + digit;
                }
                lines.push_back(line);
              }
              if (num_ptr == &primer_scores) {
                string line;
                for (unsigned int b1 = 0; b1 != nums.size(); ++b1) {
                  char char_id = rand_char();
                  while (char_id == '"' || char_id == '\\') {
                    char_id = rand_char();
                  }
                  const bool good = all_same[b1] && exceeds[b1];
                  line += good ? char_id : ' ';
                  const int est_distance_in = [b1, &distance_in]() {
                    if (distance_in[b1])
                      return static_cast<int>(distance_in[b1]);
                    for (unsigned int b = 0; b != distance_in.size(); ++b) {
                      if (distance_in[b]) {
                        return static_cast<int>(distance_in[b]) +
                            static_cast<int>(b1) - static_cast<int>(b);
                      }
                    }
                    return 0;
                  }();
                  if (good) {
                    const string complement{reverse_complement(
                        seq.substr(b1, primer_length))};
                    primers_out << char_id << '\t'
                                << gene_symbol << '\t'
                                << chromosomes[b1] << '\t'
                                << positions[b1] << '\t'
                                << strands[b1] << '\t'
                                << seq.substr(b1, primer_length) << '\t'
                                << complement << '\t'
                                << primer_scores[b1] << '\t'
                                << exon_numbers[b1] << '\t'
                                << est_distance_in << '\t'
                                << endl;
                  }
                }
                lines.push_back(line);
              } else {
                lines.push_back("");
              }
            }
          }

          // Eliminate spurious intronic annotation within exonic annotation
          // (due to sequence similarity)
          vector<GeneInfo> kept_info;
          for (const auto & info : gene_info) {
            bool keep = true;
            if (!info.exon) {
              for (const auto & info2 : gene_info) {
                if (!info2.exon) continue;
                if (info2.offset <= info.offset &&
                    info2.offset + info2.length <=
                    info.offset + info.length) {
                  keep = false;
                }
              }
            }
            if (keep) kept_info.push_back(info);
          }

          // Output intron/exon annotation
          while (kept_info.size()) {
            lines.push_back("");
            string & line = lines.back();
            vector<GeneInfo> saved;
            for (const auto & info : kept_info) {
              if (line.size() > info.offset) {
                saved.push_back(info);
                continue;
              }
              while (line.size() < info.offset) line += ' ';
              line += info.text;
            }
            kept_info.swap(saved);
          }
        }

        // The sequence
        lines.push_back(seq);

        // Output MAM annotation, packing MAMs into available display space
        while (mams.size()) {
          lines.push_back("");
          string & line = lines.back();
          vector<SimpleHit> saved;
          for (const auto & mam : mams) {
            if (line.size() > mam.off) {
              saved.push_back(mam);
              continue;
            }
            while (line.size() != mam.off) line += ' ';
            line += '[';
            line += ref.name(mam.chr) + mam.dir + to_string(mam.pos);
            // line += " " + to_string(mam.off);
            while (line.size() < mam.off + mam.len - 1) line += ' ';
            line.insert(mam.off + mam.len - 1, 1, ']');
          }
          mams.swap(saved);
        }

        // Equalize node information line lengths
        const auto id = &node;
        const auto max_length = [&lines]() {
          unsigned int max = 0;
          for (const auto line : lines) {
            if (line.size() > max) max = static_cast<unsigned int>(line.size());
          }
          return max;
        }();
        for (auto & line : lines) {
          line.insert(line.end(), max_length - line.size(), ' ');
        }

        // Output node markup
        dot << "\"" << id << "\" [shape=box"
            << ", label=\"";
        for (unsigned int l = 0; l != lines.size(); ++l) {
          if (l) dot << "\\n";
          dot << lines[l];
        }
        dot << "\"];" << endl;
      }

      // Create graph edges
      for (const auto & node : graph) {
        const auto seq = node.sequence;
        const auto count = node.count;
        const auto id = &node;
        for (const auto & other : node.out_nodes) {
          const auto counts = min(count, other->count);
          dot << "\"" << id << "\" -> \"" << &*other
              << "\" [dir=\"none\", weight=" << counts
              << ", w=" << counts
              << ", label=" << counts
              << ", tailport=e, headport=w" << "];"
              << endl;
        }
      }

      // End graph output and produce display
      dot << "}" << endl;
      dot.close();
      ostringstream render;
      const string pdf_name = graph_name + ".pdf";
      render << "dot -o " << pdf_name << " -Tpdf " << dot_name;
      // << "&& xpdf " << pdf_name << " &";
      if (system(render.str().c_str()) == -1) {
        cerr << "Problem rendering graph" << endl;
      }
    }
  }

  return 0;
} catch (Error & e) {
  cerr << "paa::Error:" << endl;
  cerr << e.what() << endl;
  return 1;
} catch (exception & e) {
  cerr << "std::exception" << endl;
  cerr << e.what() << endl;
  return 1;
} catch (...) {
  cerr << "unknown exception was caught" << endl;
  return 1;
}
```
