## Supplementary material for "G-Graph: An interactive genomic graph viewer": MUMdex Source Code for G-Graph: denovo_cn.cpp

```
//
// denovo_cn
//
// find denovo segments
//
// Copyright 2017 Peter Andrews @ CSHL
//

#include <algorithm>
#include <exception>
#include <fstream>
#include <future>
#include <iostream>
#include <map>
#include <sstream>
#include <string>
#include <utility>
#include <vector>

#include "cn.h"
#include "cngsl.h"
#include "error.h"
#include "mumdex.h"
#include "population.h"
#include "psplot.h"
#include "stats.h"
#include "strings.h"
#include "threads.h"
#include "utility.h"

using std::cin;
using std::cout;
using std::cerr;
using std::endl;
using std::exception;
using std::future;
using std::ifstream;
using std::map;
using std::max;
using std::ofstream;
using std::ostringstream;
using std::pair;
using std::string;
using std::to_string;
using std::vector;

using paa::remove_substring;
using paa::Bin;
using paa::Bounds;
using paa::ChromosomeIndexLookup;
using paa::CN_abspos;
using paa::CN_Bin;
using paa::CN_Bins;
using paa::CNStateCall;
using paa::CNStateCaller;
using paa::Error;
using paa::Family;
using paa::MAD;
using paa::Marker;
using paa::NormalParams;
using paa::Population;
using paa::Progress;
using paa::PSDoc;
using paa::PSGraph;
using paa::PSHSeries;
using paa::PSXYSeries;
using paa::Reference;
using paa::Sample;
using paa::Segment;
using paa::Segments;
using paa::ThreadPool;

int main(int argc, char* argv[]) try {
  if (--argc != 4)
    throw Error("usage: denovo_cn ref pop_file bin_file results_dir");

  // Process arguments
  const Reference ref{argv[1]};
  const ChromosomeIndexLookup lookup{ref};
  const Population pop{argv[2]};
  const string bins_name{argv[3]};
  const vector<Bin> all_bins{load_bins(bins_name, ref)};
  const unsigned int n_bins(static_cast<unsigned int>(all_bins.size()));

  const unsigned int x{ref.find_x_chromosome()};
  const unsigned int y{ref.find_y_chromosome()};

  // List of previously determined good bins
  const vector<Bin> bins{[&all_bins]() {
      vector<Bin> result;
      for (const Bin & bin : all_bins) {
        if (!bin.bad()) {
          result.push_back(bin);
        }
      }
      return result;
    }()};

  const string results_dir{argv[4]};

  // Load all segments, profiles
  const vector<vector<Segments>> segments{
    [&pop, &results_dir, &ref, &lookup, n_bins]() {
      ThreadPool pool{20};
      vector<future<vector<Segments>>> futures;
      for (const Family & family : pop.families()) {
        futures.push_back(pool.run(
            [&pop, &results_dir, n_bins, &ref, &lookup](const Family & f) {
              vector<Segments> result;
              for (const Sample & sample : pop.samples(f)) {
                const string sample_name{pop.sample(sample)};
                const string segments_file{results_dir + "/" + sample_name +
                      "/" + sample_name + "_" + to_string(n_bins) +
                      "_bins_segments.txt"};
                result.emplace_back(segments_file, sample_name,
                                    ref, lookup);
              }
              return result;
            }, family));
      }
      Progress load_progress{pop.families().size(), 0.1, "File load"};
      vector<vector<Segments>> result;
      for (auto & fut : futures) {
        result.push_back(fut.get());
        load_progress();
      }
      return result;
    }()};
  cerr << "Loaded segments" << endl;

  const vector<vector<CNStateCaller>> caller{[&segments, &bins, &ref]() {
      vector<vector<CNStateCaller>> result;
      for (const vector<Segments> & family_segments : segments) {
        result.push_back(vector<CNStateCaller>());
        for (const Segments & segs : family_segments) {
          result.back().emplace_back(ref, bins, segs.profile);
        }
      }
      return result;
    }()};
  cerr << "Loaded state callers" << endl;

  unsigned int n_call{0};
  unsigned int n_denovo{0};
  for (const Family & family : pop.families()) {
    const vector<Sample> & samples{pop.samples(family)};
    for (unsigned int s{0}; s != samples.size(); ++s) {
      const Sample sample{samples[s]};
      if (pop.is_proband(sample) || pop.is_sibling(sample)) {
        for (const auto & segment : segments[family][s]) {
          if (segment.chr == x || segment.chr == y) continue;
          if (segment.pass_score() && segment.int_cn != segment.expected &&
              fabs(segment.cn - segment.expected) > 0.6) {
            ++n_call;
            // Check parent calls
            bool in_parent{false};
            vector<double> cn;
            vector<CNStateCall> calls;
            for (unsigned int p{0}; p != samples.size(); ++p) {
              const CNStateCall call{
                caller[family][p].call(segment.start, segment.stop)};
              cn.push_back(2.0 * call.segment_count /
                           ((segment.stop - segment.start) *
                            caller[family][p].average_autosome_count));
              calls.push_back(call);
            }
            for (unsigned int p{0}; p != samples.size(); ++p) {
              const CNStateCall & call{calls[p]};
              if (pop.is_parent(samples[p]) &&
                  (call.expected_state != call.best_call ||
                   fabs(cn[s] - cn[p]) < 0.4)) {
                in_parent = true;
              }
            }
            if (!in_parent) {
              ++n_denovo;
              // Get population results
              vector<double> vals;
              vector<double> parent_vals;
              for (const Family & f : pop.families()) {
                const vector<Sample> & ss{pop.samples(f)};
                for (unsigned int s2{0}; s2 != ss.size(); ++s2) {
                  if (pop.is_parent(ss[s2])) {
                    const CNStateCall call{
                      caller[f][s2].call(segment.start, segment.stop)};
                    const double val{1.0 * call.segment_count /
                          ((segment.stop - segment.start) *
                           caller[f][s2].average_autosome_count)};
                    if (f == family) {
                      parent_vals.push_back(val);
                    } else {
                      vals.push_back(val);
                    }
                  }
                }
              }
              const NormalParams norm{vals};
              const double sign{(segment.int_cn > segment.expected ?
                                 1.0 : -1.0)};
              cout << pop.family(family)
                   << " " << pop.sample(sample)
                   << " " << pop.member(sample)
                   << " " << ref.name(segment.chr)
                   << " " << segment.startpos
                   << " " << segment.stoppos
                   << " " << segment.int_cn
                   << " " << segment.expected
                   << " " << segment.cn
                   << " " << segment.score
                   << " " << segment.n_bins
                   << " " << vals.size()
                   << " " << norm.mean
                   << " " << norm.stdev
                   << " " << sign * (parent_vals[0] - norm.mean) / norm.stdev
                   << " " << sign * (parent_vals[1] - norm.mean) / norm.stdev
                   << " " << max(
                       sign * (parent_vals[0] - norm.mean) / norm.stdev,
                       sign * (parent_vals[1] - norm.mean) / norm.stdev);
              for (const double & c : cn) {
                cout << " " << c;
              }
              cout << endl;
            }
          }
        }
      }
    }
  }

  cerr << "denovo " << n_denovo << " event " << n_call
       << " " << 1.0 * n_denovo / n_call << endl;

  return 0;


  const vector<CN_Bins> results{[argv, argc, &bins] () {
      vector<CN_Bins> result;
      result.reserve(argc);
      for (int a{0}; a != argc; ++a) {
        result.emplace_back(argv[a]);
        if (result.back().size() != bins.size()) {
          throw Error("Bin size mismatch");
        }
      }
      return result;
    }()};

  const string out_base{remove_substring(bins_name, ".txt") + ".cut"};

  // Graphs
  PSDoc ps{out_base};
  ps.pdf(false);
  const Marker small_red_marker{paa::circle(), 0.1, "1 0 0", 1, true};

  // Ratio MAD histogram
  PSGraph mad_hist_graph{ps, ";Ratio MAD;N", Bounds(0, 0.2)};
  PSHSeries<double, unsigned int> mad_hist{mad_hist_graph, 200, "1 0 0", false};

  // Ratio MAD vs abspos
  PSGraph mad_vs_abspos_graph{ps, ";Absolute Position (GB); Ratio MAD"};
  mad_vs_abspos_graph.log_y(true);
  // mad_vs_abspos_graph.range().yh(0.5);
  PSXYSeries mad_vs_abspos{mad_vs_abspos_graph, small_red_marker};

  // Get result statistics
  vector<MAD> ratio_mads;
  ratio_mads.reserve(bins.size());
  vector<double> ratios;
  ratios.reserve(results.size());
  const unsigned int x_chr{ref.find_x_chromosome()};
  const unsigned int y_chr{ref.find_y_chromosome()};
  vector<vector<double>> mads(3);
  const double billion{1000000000};
  vector<double> position_mads(bins.size());
  vector<unsigned int> position_regions(bins.size());
  for (unsigned int b{0}; b != bins.size(); ++b) {
    ratios.clear();
    for (unsigned int r{0}; r != results.size(); ++r) {
      const CN_Bin & result_bin{results[r][b]};
      ratios.push_back(result_bin.ratio());
    }
    sort(ratios.begin(), ratios.end());
    ratio_mads.emplace_back(ratios);
    const MAD & mad{ratio_mads.back()};
    const Bin & bin{bins[b]};
    // Fill graphs
    mad_hist.add_point(mad.mad());
    mad_vs_abspos.add_point(bin.abspos() / billion, mad.mad());
    const unsigned int region{bin.chromosome() == x_chr ? 1U :
          (bin.chromosome() == y_chr ? 2U : 0U)};
    mads[region].push_back(mad.mad());
    position_mads[b] = mad.mad();
    position_regions[b] = region;
  }

  // Mad of mads...
  vector<MAD> mads_mads;
  const CN_abspos cn_abspos{ref};
  const unsigned int region_chromosome_start[3]{
    cn_abspos.chromosomes().front(), x_chr, y_chr};
  vector<double> cutoffs;
  for (unsigned int c{0}; c != mads.size(); ++c) {
    vector<double> & vals{mads[c]};
    sort(vals.begin(), vals.end());
    mads_mads.emplace_back(vals);
    const unsigned int chr{region_chromosome_start[c]};
    const MAD & mads_mad{mads_mads.back()};
    const unsigned int start{cn_abspos.ref_offset(chr)};
    const unsigned int stop{chr == cn_abspos.chromosomes().front() ?
          cn_abspos.ref_offset(x_chr) : cn_abspos.ref_offset(chr + 1)};
    const double cutoff{mads_mad.median() + 6 * mads_mad.mad()};
    auto lower = lower_bound(vals.begin(), vals.end(), cutoff);
    cout << ref.name(chr) << " " << start << " " << stop << " "
         << mads_mad.median() << " " << cutoff << " "
         << 1.0 * (lower - vals.begin()) / vals.size() << endl;

    ostringstream cutoff_line;
    cutoff_line << "0 0 0 c 2 lw np " << start / billion << " "
                << cutoff << " gc m "
                << stop / billion << " " << cutoff << " gc l sp";
    mad_vs_abspos_graph.ps(cutoff_line.str());
    cutoffs.push_back(cutoff);
  }

  // Output bins
  const string out_name{out_base + ".txt"};
  ofstream out{out_name.c_str()};
  if (!out) throw Error("Problem opening output bins file") << out_name;
  unsigned int gb{0};
  for (unsigned int b{0}; b != all_bins.size(); ++b) {
    const Bin & bin{all_bins[b]};
    if (bin.bad()) {
      bin.output(out, ref);
    } else {
      bin.output(out, ref, position_mads[gb] > cutoffs[position_regions[gb]]);
      ++gb;
    }
  }

  return 0;
} catch (Error & e) {
  cerr << "paa::Error:" << endl;
  cerr << e.what() << endl;
  return 1;
} catch (exception & e) {
  cerr << "std::exception" << endl;
  cerr << e.what() << endl;
  return 1;
} catch (...) {
  cerr << "unknown exception was caught" << endl;
  return 1;
}
```
