## Supplementary material for "G-Graph: An interactive genomic graph viewer": MUMdex Source Code for G-Graph: denovo_pseudogenes.cpp

```
//
// denovo_pseudogenes
//
// find denovo pseudogenes
//
// Copyright 2015 Peter Andrews @ CSHL
//

#include <algorithm>
#include <exception>
#include <iomanip>
#include <iostream>
#include <fstream>
#include <map>
#include <memory>
#include <set>
#include <sstream>
#include <string>
#include <utility>
#include <vector>

#include "bed.h"
#include "error.h"
#include "genes.h"
#include "mumdex.h"
#include "utility.h"

using std::exception;
using std::cout;
using std::cerr;
using std::endl;
using std::ifstream;
using std::ofstream;
using std::istringstream;
using std::ostringstream;
using std::make_unique;
using std::map;
using std::max;
using std::min;
using std::pair;
using std::set;
using std::setw;
using std::string;
using std::unique_ptr;
using std::vector;

using paa::base_chars;
using paa::base_index;
using paa::complement;
using paa::BedFile;
using paa::ChromosomeIndexLookup;
using paa::CompressedInts;
using paa::Error;
using paa::MUMdex;
using paa::GeneXrefs;
using paa::JunctionInfo;
using paa::KnownGenes;
using paa::Mappability;
using paa::MUM;
using paa::PosInfo;
using paa::read_ahead;
using paa::Reference;
using paa::serr;
using paa::sout;

struct FamilyJunctionSorter {
  explicit FamilyJunctionSorter(const vector<unsigned int> & ids) :
      family_ids{ids} {}
  bool operator()(const JunctionInfo & left, const JunctionInfo & right) const {
    if (left < right) return true;
    if (right < left) return false;
    if (family_ids[left.sample] == family_ids[right.sample])
      return left.sample < right.sample;
    return family_ids[left.sample] < family_ids[right.sample];
  }
  const vector<unsigned int> & family_ids;
};


class SavedOutput {
 public:
  SavedOutput() : pos{0, 0} {}
  SavedOutput(const PosInfo pos_arg,
              const string & output_arg, const string & search_arg) :
      pos{pos_arg}, output{output_arg}, search{search_arg} { }
  PosInfo pos;
  string output{};
  string search{};
  bool operator<(const SavedOutput & right) const {
    return pos < right.pos;
  }
};

class FerryInfo {  // A bridge if same read, invariantless if between mates
 public:
  FerryInfo(const uint64_t pair_index_,
            const MUM exon_arg, const MUM remote_arg, const bool right_arg) :
      pair_index{pair_index_}, exon{exon_arg}, remote{remote_arg},
    right{right_arg} {}
  uint64_t pair_index;
  MUM exon;
  MUM remote;
  bool right;
  unsigned int position() const {
    return remote.position0();
  }
  PosInfo posinfo() const {
    return PosInfo(remote.chromosome(), remote.position0());
  }
  bool same_read() const {
    return exon.read_2() == remote.read_2();
  }
  bool same_orientation() const {
    return (!same_read()) ^ (exon.flipped() == remote.flipped());
  }
  bool operator<(const FerryInfo & rhs) const {
    if (right == rhs.right) {
      if (posinfo() == rhs.posinfo()) {
        if (exon.length() == rhs.exon.length()) {
          return remote.length() > rhs.remote.length();
        } else {
          return exon.length() > rhs.exon.length();
        }
      } else {
        return posinfo() < rhs.posinfo();
      }
    } else {
      return right < rhs.right;
    }
  }
};

class SavedCandidate {
 public:
  SavedCandidate() { }
  SavedCandidate(const unsigned int isoform_index_arg,
                 const unsigned int count_arg) :
      isoform_index{isoform_index_arg}, count{count_arg} {}
  unsigned int isoform_index{0};
  unsigned int count{0};
};

class GeneCandidates {
 public:
  map<unsigned int, SavedCandidate> candidates{};
  map<unsigned int, SavedCandidate> also_seen{};
};

int main(int argc, char* argv[]) try {
  paa::exit_on_pipe_close();

  --argc;
  if (argc != 5) throw Error(
          "usage: denovo_pseudogenes family_info genes isoforms xrefs bed");

  const string ref_name{"/data/safe/paa/analysis/mums/hg19/chrAll.fa"};
  const Reference ref{ref_name};
  const ChromosomeIndexLookup chr_lookup{ref};
  const Mappability mappability{ref_name, true};
  const string genes_name{argv[2]};
  const string isoforms_name{argv[3]};
  const string kgXrefs_name{argv[4]};

  const KnownGenes genes{genes_name, isoforms_name, chr_lookup, ref};
  const GeneXrefs xref{kgXrefs_name};

  const string bed_name{argv[5]};
  const BedFile bed{bed_name};

  const auto last_slash = bed_name.find_last_of('/');
  const string bed_basename = last_slash == string::npos ?
      bed_name : bed_name.substr(last_slash + 1);

  const auto & junctions = genes.junctions;

  const vector<string> member_names{"mother", "father", "self", "sibling"};
  const map<string, unsigned int> member_lookup{
    {"mother", 0}, {"father", 1}, {"self", 2}, {"sibling", 3}};
  vector<string> families;
  vector<unsigned int> members;
  vector<string> samples;
  vector<string> sexes;
  vector<string> pseudo_names;
  vector<unsigned int> family_ids;
  vector<unsigned int> n_members;
  vector<unsigned int> members_start;
  vector<unsigned int> members_stop;
  vector<JunctionInfo> counts;
  vector<string> mumdex_filenames;
  vector<string> anchor_filenames;
  {
    const string family_info{argv[1]};
    ifstream family_file{family_info.c_str()};
    if (!family_file)
      throw Error("Could not open family file") << family_info;
    string line;
    string family;
    string member;
    string sample;
    string sex;
    while (family_file) {
      getline(family_file, line);
      if (line.empty()) break;
      istringstream line_stream{line};
      if (line_stream) {
        line_stream >> family;
        if (!line_stream)
          throw Error("Problem reading family") << families.size();
        unsigned int n_member = 0;
        members_start.push_back(static_cast<unsigned int>(samples.size()));
        while (line_stream) {
          line_stream >> member >> sample >> sex;
          if (line_stream) {
            const string sample_dir =
                "/data/safe/paa/analysis/mums/output/samples/" + sample;
            string pseudo_name = sample_dir + "/pseudo2.out";
            ifstream input{pseudo_name.c_str()};
            if (input) {
              while (input) {
                try {
                  JunctionInfo info{input, samples.size()};
                  counts.push_back(info);
                } catch (JunctionInfo::JunctionReadError) {
                  break;
                }
              }
              ++n_member;
              members.push_back(member_lookup.at(member));
              samples.push_back(sample);
              sexes.push_back(sex);
              pseudo_names.push_back(pseudo_name);
              family_ids.push_back(static_cast<unsigned int>(families.size()));
              const auto mumdex_dir = sample_dir + "/mumdex";
              mumdex_filenames.push_back(mumdex_dir);
              const auto anchor_dir = mumdex_dir + "/" + bed_basename;
              anchor_filenames.push_back(anchor_dir);
            } else {
              cerr << "Could not read " << pseudo_name << endl;
            }
          }
        }
        if (n_member < 3)
          cerr << "Incomplete family " << family << " " << n_member << endl;
        members_stop.push_back(static_cast<unsigned int>(samples.size()));
        families.push_back(family);
        n_members.push_back(n_member);
      }
    }
  }
  serr << "Loaded" << families.size() << "families and"
       << samples.size() << "samples with"
       << counts.size() << "junction counts taking"
       << counts.size() * sizeof(JunctionInfo) << "bytes" << endl;
  sort(counts.begin(), counts.end(), FamilyJunctionSorter(family_ids));
  if (0) {
    for (const auto count : counts) {
      sout << count.junction_index << count.invariant_index
           << count.invariant << count.count
           << families[family_ids[count.sample]]
           << samples[count.sample]
           << member_names[members[count.sample]]
           << sexes[count.sample]
           << endl;
    }
  }
  vector<vector<unsigned int>> gene_junctions(genes.size());
  for (unsigned int j = 0; j != junctions.size(); ++j) {
    const auto & junction = junctions[j];
    gene_junctions[junction.gene_index].push_back(j);
  }
  vector<vector<unsigned int>> junction_sample_counts(
      junctions.size(), vector<unsigned int>(samples.size()));
  vector<vector<unsigned int>> gene_sample_counts(
      genes.size(), vector<unsigned int>(samples.size()));
  vector<vector<unsigned int>> gene_sample_n_junctions(
      genes.size(), vector<unsigned int>(samples.size()));
  vector<vector<unsigned int>> gene_family_sample_counts(
      genes.size(), vector<unsigned int>(families.size()));
  vector<vector<unsigned int>> gene_family_n_samples(
      genes.size(), vector<unsigned int>(families.size()));
  vector<unsigned int> gene_n_families(genes.size());
  vector<unsigned int> gene_n_samples(genes.size());
  vector<unsigned int> gene_counts(genes.size());
  vector<unique_ptr<vector<unsigned int>>> count_lookup(
      junctions.size() * samples.size());
  for (unsigned int c = 0; c != counts.size(); ++c) {
    const auto count = counts[c];
    const auto & info = junctions[count.junction_index];
    const unsigned int j{count.junction_index};
    const unsigned int s{static_cast<unsigned int>(count.sample)};
    const unsigned int l{static_cast<unsigned int>(j * samples.size() + s)};
    if (!count_lookup[l]) {
      count_lookup[l] = make_unique<vector<unsigned int>>();
    }
    count_lookup[l]->push_back(c);
    junction_sample_counts[j][s] +=
        count.count;
    gene_sample_counts[info.gene_index][s] +=
        count.count;
  }
  for (unsigned int j = 0; j != junctions.size(); ++j) {
    const auto & junction = junctions[j];
    for (unsigned int s = 0; s != samples.size(); ++s) {
      if (junction_sample_counts[j][s]) {
        ++gene_sample_n_junctions[junction.gene_index][s];
      }
    }
  }
  for (unsigned int g = 0; g != genes.size(); ++g) {
    for (unsigned int s = 0; s != samples.size(); ++s) {
      if (gene_sample_counts[g][s]) {
        ++gene_n_samples[g];
        ++gene_family_n_samples[g][family_ids[s]];
        gene_family_sample_counts[g][family_ids[s]] +=
            gene_sample_counts[g][s];
      }
      gene_counts[g] += gene_sample_counts[g][s];
    }
  }
  for (unsigned int g = 0; g != genes.size(); ++g) {
    for (unsigned int f = 0; f != families.size(); ++f) {
      if (gene_family_sample_counts[g][f]) {
        ++gene_n_families[g];
      }
    }
  }
  const unsigned int max_gene_families = 10;
  const unsigned int min_sample_count = 3;
  const unsigned int max_n_samples = 10;
  const unsigned int max_family_n_samples = 1;
  const unsigned int min_junctions = 1;
  const set<unsigned int> show_members{2, 3};
  map<string, GeneCandidates> gene_candidates;
  sout << "max_gene_families" << max_gene_families
       << "min_sample_count" << min_sample_count
       << "max_n_samples" << max_n_samples
       << "max_family_n_samples" << max_family_n_samples
       << "min_junctions" << min_junctions
       << endl;
  unsigned int n_seen = 0;
  unsigned int n_selected = 0;
  unsigned int n_denovo = 0;
  unsigned int n_saved = 0;
  ofstream search_out("search.txt");
  for (unsigned int g = 0; g != genes.size(); ++g) {
    const auto & gene = genes[g];
    if (gene_n_families[g] <= max_gene_families) {
      for (unsigned int s = 0; s != samples.size(); ++s) {
        const auto f = family_ids[s];
        if (gene_sample_counts[g][s]) {
          ++n_seen;
          const SavedCandidate to_save{g, gene_sample_counts[g][s]};
          const auto symbol = xref[gene.name].geneSymbol;
          GeneCandidates & gene_cand = gene_candidates[symbol];
          auto & candidates = gene_cand.candidates;
          auto & also_seen = gene_cand.also_seen;
          const auto found = candidates.find(s);
          bool denovo = false;
          if (gene_sample_counts[g][s] >= min_sample_count &&
              gene_n_samples[g] <= max_n_samples &&
              gene_family_n_samples[g][f] <= max_family_n_samples &&
              gene_sample_n_junctions[g][s] >= min_junctions) {
            ++n_selected;
            if (show_members.count(members[s])) {
              ++n_denovo;
              denovo = true;
              also_seen.erase(s);
              if (found == candidates.end() ||
                  found->second.count < to_save.count) {
                if (found == candidates.end()) {
                  ++n_saved;
                }
                candidates[s] = to_save;
              }
              for (auto s2 = members_start[f]; s2 != members_stop[f]; ++s2) {
                if (s == s2) continue;
                if (candidates.count(s2)) continue;
                const SavedCandidate to_save_2{g, gene_sample_counts[g][s2]};
                const auto found2 = also_seen.find(s2);
                if (found2 == also_seen.end() ||
                    found2->second.count < to_save_2.count) {
                  also_seen[s2] = to_save_2;
                }
              }
            }
          }
          if (!denovo) {
            if (found == candidates.end()) {
              const auto found2 = also_seen.find(s);
              if (found2 == also_seen.end() ||
                  found2->second.count < to_save.count) {
                also_seen[s] = to_save;
              }
            }
          }
        }
      }
    }
  }
  vector<SavedOutput> ordered_output;
  for (auto gene_cand_info : gene_candidates) {
    const auto & symbol = gene_cand_info.first;
    const auto & gene_cand = gene_cand_info.second;
    if (gene_cand.candidates.empty()) continue;
    vector<pair<pair<unsigned int, SavedCandidate>, bool>> sample_details;
    for (const auto info : gene_cand.candidates) {
      sample_details.emplace_back(info, true);
    }
    for (const auto info : gene_cand.also_seen) {
      sample_details.emplace_back(info, false);
    }
    ostringstream out;
    out << "----------------------------------------------" << endl;
    ostringstream search_out_gene;
    for (const auto info : sample_details) {
      const auto sample_info = info.first;
      const auto s = sample_info.first;
      const auto f = family_ids[s];
      const auto gene_info = sample_info.second;
      const auto is_candidate = info.second;
      const auto g = gene_info.isoform_index;
      const auto & gene = genes[g];
      if (gene_info.count != gene_sample_counts[g][s])
        throw Error("Unexpected count saved")
            << gene_info.count << gene_sample_counts[g][s] << g << s
            << is_candidate;
      unsigned int n_junctions_seen = 0;
      for (const auto j : gene_junctions[g]) {
        if (junction_sample_counts[j][s]) {
          ++n_junctions_seen;
        }
      }
      out << gene.name << " "
          << symbol << " "
          << gene.strand << " "
          << ref.name(gene.chr) << " "
          << gene.t_start << " "
          << gene.t_stop << " "
          << gene_junctions[g].size() << " "
          << n_junctions_seen << " "
          << gene_sample_counts[g][s] << " "
          << gene_n_families[g] << " "
          << gene_n_samples[g] << " "
          << gene_family_n_samples[g][f] << " "
          << families[f] << " "
          << samples[s] << " "
          << member_names[members[s]] << " "
          << sexes[s] << " "
          << (is_candidate ? "" : "NOT A ") << "Candidate"
          << endl;
      const unsigned int ss = members_start[family_ids[s]];
      const unsigned int se = members_stop[family_ids[s]];
      const auto mumdexs = [ss, se, &mumdex_filenames]() {
        vector<MUMdex> mumdexs_;
        mumdexs_.reserve(se - ss);
        for (unsigned int m = ss; m != se; ++m) {
          mumdexs_.emplace_back(mumdex_filenames[m]);
        }
        return mumdexs_;
      }();
      for (const auto j : gene_junctions[g]) {
        if (junction_sample_counts[j][s]) {
          const auto junction = junctions[j];
          search_out_gene << samples[s] << " "
                          << mumdex_filenames[s] << " "
                          << g << " "
                          << symbol << " "
                          << j << " "
                          << ref.name(junction.chr) << " "
                          << junction.invariants[0];
          out << "  "
              << ref.name(junction.chr) << " "
              << junction.pos << " "
              << junction_sample_counts[j][s];
          vector<unsigned int> hit_positions;
          for (const auto c : *count_lookup[j * samples.size() + s]) {
            const auto count = counts[c];
            search_out_gene << " " << count.invariant_index
                            << " " << count.invariant
                            << " " << count.position;
            out << " " << count.invariant_index
                << ":" << count.position << ":" << count.invariant;
            const auto inv = junctions[j].invariants[count.invariant_index];
            if (inv != count.invariant) {
              out << "!=" << inv;
            }
            out << ":" << count.count;
            if (hit_positions.size() &&
                hit_positions.back() == count.position) continue;
            hit_positions.push_back(count.position);
          }
          search_out_gene << endl;
          reverse(hit_positions.begin(), hit_positions.end());
          out << endl;
          unsigned int block_start = 0;
          unsigned int bed_n = 0;
          const unsigned int position_window = 10;  // should match
          const auto sp = junction.pos > position_window ?
              junction.pos - position_window : 0;
          const auto ep = junction.pos + position_window >=
              ref.size(junction.chr) ? ref.size(junction.chr) :
              junction.pos + position_window;
          while (bed_n + 1 != bed.size()) {
            const auto interval = bed[bed_n + 1];
            const auto bed_chr = chr_lookup[interval.chromosome];
            if (bed_chr > junction.chr ||
                (bed_chr == junction.chr && interval.start_pos > sp)) {
              break;
            }
            block_start += bed[bed_n].stop_pos - bed[bed_n].start_pos;
            ++bed_n;
          }
          for (unsigned int m = ss; m != se; ++m) {
            CompressedInts<uint16_t, uint8_t> compressed
            {anchor_filenames[m], block_start * 4, bed_n};
            out << "    " << member_names[members[m]];
            for (unsigned int b = bed_n; b != bed.size(); ++b) {
              const auto interval = bed[b];
              const auto bed_chr = chr_lookup[interval.chromosome];
              if (bed_chr > junction.chr ||
                  (bed_chr == junction.chr &&
                   interval.start_pos >= ep))
                break;
              for (unsigned int p = interval.start_pos;
                   p != interval.stop_pos; ++p) {
                if (bed_chr == junction.chr && p >= ep) break;
                const auto in_ref = compressed.next_int();
                const auto in_anchor = compressed.next_int();
                const auto out_ref = compressed.next_int();
                const auto out_anchor = compressed.next_int();
                if (p < sp) continue;
                out << '\t' << in_ref << "/" << in_anchor;
                if (s == m && p == hit_positions.back()) {
                  out << "*";
                  hit_positions.pop_back();
                }
                if (0)
                  out << "/" << out_ref << "/" << out_anchor;
              }
            }
            out << endl;
          }
          out << "    right";
          for (unsigned int p = sp; p != ep; ++p) {
            out << "   \t" << ref[junction.chr][p];
          }
          out << endl;
          out << "    left";
          for (unsigned int p = sp; p != ep; ++p) {
            const int po = p + junction.invariants[0];
            out << "   \t" << (po >= 0 ? ref[junction.chr][po] : 'X');
          }
          out << endl;
          for (const auto c : *count_lookup[j * samples.size() + s]) {
            const auto count = counts[c];
            const PosInfo pos{junction.chr, count.position};
            for (unsigned int m = ss; m != se; ++m) {
              auto base_counts = vector<vector<unsigned int>>(
                  position_window * 2, vector<unsigned int>(6));
              const MUMdex & mumdex{mumdexs[m - ss]};
              set<uint64_t> pair_skip;
              for (auto mum_iter = mumdex.lower_bound(pos);
                   mum_iter != mumdex.index().end(); ++mum_iter) {
                const auto mum = mumdex.mum(*mum_iter);
                if (mum.position0() != count.position) break;
                const auto pair = mumdex.pair(*mum_iter);
                if (pair.dupe()) continue;
                if (mum.flipped()) {
                  if (mum.touches_end()) continue;
                } else {
                  if (mum.offset() == 0) continue;
                }
                const auto pair_index = mum_iter->pair_index();
                if (pair_skip.count(pair_index)) continue;
                pair_skip.insert(pair_index);
                const string sequence{
                  mumdex.sequences(pair_index)[mum.read_2()]};
                for (unsigned int p = sp; p != ep; ++p) {
                  const int b = mum.pos_to_base(p);
                  if (b >= 0 && b < static_cast<int>(sequence.size())) {
                    ++base_counts[p - sp][base_index(
                        mum.flipped() ? complement(sequence[b]) :
                        sequence[b])];
                  }
                }
              }
              out << "    " << member_names[members[m]];
              for (unsigned int p = 0; p!= base_counts.size(); ++p) {
                out << '\t';
                for (auto b = 1; b != 6; ++b) {
                  if (base_counts[p][b]) {
                    out << base_counts[p][b] << base_chars[b];
                  }
                }
              }
              out << endl;
            }
          }
        }
      }
      // Look for mums that map elsewhere
      const unsigned int gene_avoidance = 100000;
      const unsigned int match_interval = 5000;
      const unsigned int min_excess = 7;
      const unsigned int center_avoid = 5;
      const unsigned int edge_zone = 5000;
      vector<FerryInfo> ferries;
      const PosInfo t_start{gene.chr, gene.t_start};
      const PosInfo t_stop{gene.chr, gene.t_stop};
      const MUMdex & mumdex{mumdexs[s - ss]};
      const auto left_iter = mumdex.lower_bound(t_start);
      const auto right_iter = mumdex.lower_bound(t_stop);
      const PosInfo left_avoid{gene.chr, gene.t_start > gene_avoidance ?
            gene.t_start - gene_avoidance : 0};
      const PosInfo right_avoid{gene.chr, gene.t_stop + gene_avoidance};
      set<uint64_t> pairs_seen;
      for (auto iter = left_iter; iter != right_iter; ++iter) {
        const auto pair_index = iter->pair_index();
        const auto pair_ = mumdex.pair(*iter);
        if (pair_.dupe()) continue;
        if (pairs_seen.count(pair_index)) continue;
        pairs_seen.insert(pair_index);
        auto edge_mum = mumdex.mums_end();
        unsigned int min_edge_distance = 1000000000;
        bool on_right = false;
        // find mum closest to edge of gene in pair
        for (auto miter = mumdex.mums_begin(pair_index);
             miter != mumdex.mums_end(pair_index); ++miter) {
          const auto mum = *miter;
          if (mum.chromosome() != gene.chr) continue;
          if (mum.position0() + mum.length() < gene.t_start) continue;
          if (mum.position0() >= gene.t_stop) continue;
          bool in_exon = false;
          bool edge_exon = false;
          unsigned int distance_in = 0;
          unsigned int total_length = 0;
          for (unsigned int e = 0; e != gene.n_exons; ++e) {
            if (e == 0 || e + 1 == gene.n_exons) {
              if (mum.position0() >= gene.exon_starts[e] &&
                  mum.position0() < gene.exon_stops[e]) {
                in_exon = true;
                distance_in = total_length +
                    mum.position0() - gene.exon_starts[e];
                edge_exon = true;
              } else if (mum.position0() + mum.length() >=
                         gene.exon_starts[e] &&
                         mum.position0() + mum.length() <
                         gene.exon_stops[e]) {
                in_exon = true;
                distance_in = total_length +
                    mum.position0() + mum.length() - gene.exon_starts[e];
                edge_exon = true;
              } else if (mum.position0() < gene.exon_starts[e] &&
                         mum.position0() + mum.length() >=
                         gene.exon_stops[e]) {
                in_exon = true;
                if (e + 1 == gene.n_exons) {
                  distance_in = total_length;
                } else {
                  distance_in = total_length + gene.exon_stops[e] -
                      gene.exon_starts[e];
                }
                edge_exon = true;
              }
            }
            total_length += gene.exon_stops[e] - gene.exon_starts[e];
          }
          if (!in_exon) continue;
          if (!edge_exon) continue;
          if (total_length <= center_avoid) continue;
          const unsigned int used_edge_zone =
              2 * edge_zone + center_avoid < total_length ?
              edge_zone : (total_length - center_avoid) / 2;
          const bool this_on_right = 2 * distance_in >= total_length;
          const bool at_edge = this_on_right ?
              total_length - distance_in < used_edge_zone :
              distance_in < used_edge_zone;
          if (!at_edge) continue;
          const unsigned int edge_distance =
              this_on_right ? total_length - distance_in : distance_in;
          if (edge_distance < min_edge_distance) {
            min_edge_distance = edge_distance;
            edge_mum = miter;
            on_right = this_on_right;
          }
        }
        if (edge_mum == mumdex.mums_end()) continue;
        const auto mum = *edge_mum;
        const auto mum_index = edge_mum - mumdex.mums_begin(pair_index);
        set<MUM> seen_mums;
        for (unsigned int m = 0; m != mumdex.n_mums(pair_index); ++m) {
          if (m == mum_index) continue;
          const auto other_mum = mumdex.mum(mumdex.mums_start(pair_index) + m);
          const PosInfo other_pos{other_mum.chromosome(),
                other_mum.position0()};
          if (other_pos >= left_avoid && other_pos < right_avoid)
            continue;
          if ([other_mum, &seen_mums]() {
              const unsigned int seen_threshold = 5000;
              for (const auto other : seen_mums) {
                if (other_mum.close_by(other, seen_threshold)) return true;
              }
              return false;
            }()) continue;
          seen_mums.insert(other_mum);
          const bool right_side = (mum.read_2() && !other_mum.read_2()) ?
              !mum.flipped() : (mum.flipped() ^ (m > mum_index));
          if (on_right == right_side)
            ferries.emplace_back(pair_index, mum, other_mum, right_side);
        }
      }
      sort(ferries.begin(), ferries.end());
      for (unsigned int l = 0; l != ferries.size(); ++l) {
        const auto left = ferries[l];
        const unsigned int left_remote_excess =
            left.remote.length() -  mappability.low(
                ref.offset(left.remote.chromosome()) +
                left.remote.position0());
        const unsigned int left_exon_excess =
            left.exon.length() -  mappability.low(
                ref.offset(left.exon.chromosome()) +
                left.exon.position0());
        if (left_remote_excess < min_excess ||
            left_exon_excess < min_excess) continue;
        out << '\n' << "Ferry" << '\t'
            << left.pair_index << '\t'
            << ref.name(left.remote.chromosome()) << '\t'
            << setw(9) << left.remote.position0() << '\t'
            << (left.right ? 'r' : 'l') << '\t'
            << (left.same_read() ? 'm' : 'p') << '\t'
            << left_remote_excess << '\t'
            << left_exon_excess << '\t'
            << left.same_orientation() << '\t'
            << ref.name(left.exon.chromosome()) << '\t'
            << setw(9) << left.exon.position0() << '\t'
            << endl;
        mumdex.pair_view(out, left.pair_index,
                      {{left.exon, "E"}, {left.remote, "R"}});
        if (left.right) continue;
        for (unsigned int r = 0; r != ferries.size(); ++r) {
          const auto right = ferries[r];
          if (!right.right) continue;
          if (left.remote.chromosome() !=
              right.remote.chromosome()) continue;
          if (left.same_orientation() != right.same_orientation())
            continue;
          if (left.remote.position0() + match_interval <
              right.remote.position0() ||
              left.remote.position0() >
              right.remote.position0() + match_interval)
            continue;
          const unsigned int right_remote_excess =
              right.remote.length() -  mappability.low(
                  ref.offset(right.remote.chromosome()) +
                  right.remote.position0());
          const unsigned int right_exon_excess =
              right.exon.length() -  mappability.low(
                  ref.offset(right.exon.chromosome()) +
                  right.exon.position0());
          if (right_remote_excess < min_excess ||
              right_exon_excess < min_excess) continue;
          out << "MATCHES" << '\t'
              << ref.name(right.remote.chromosome()) << '\t'
              << setw(9) << right.remote.position0() << '\t'
              << right.same_read() << '\t'
              << right_remote_excess << '\t'
              << right_exon_excess << '\t'
              << right.right << '\t'
              << ref.name(right.exon.chromosome()) << '\t'
              << setw(9) << right.exon.position0() << '\t'
              << endl;
        }
      }
    }
    out << "----------------------------------------------\n\n" << endl;
    const auto g = gene_cand.candidates.begin()->second.isoform_index;
    const auto & gene = genes[g];
    ordered_output.emplace_back(PosInfo(gene.chr, gene.t_start),
                                out.str(), search_out_gene.str());
  }
  sort(ordered_output.begin(), ordered_output.end());
  unsigned int n_kept = 0;
  for (const auto & out : ordered_output) {
    ++n_kept;
    cout << out.output;
    search_out << out.search;
  }
  if (n_kept != n_saved) throw Error("Unexpected n_kept");
  sout << "Seen" << n_seen << "selected" << n_selected
       << "saved" << n_saved << "denovo" << n_denovo << endl;

  return 0;
} catch (Error & e) {
  cerr << "paa::Error:" << endl;
  cerr << e.what() << endl;
  return 1;
} catch (exception & e) {
  cerr << "std::exception" << endl;
  cerr << e.what() << endl;
  return 1;
} catch (...) {
  cerr << "unknown exception was caught" << endl;
  return 1;
}
```
