## Supplementary material for "G-Graph: An interactive genomic graph viewer": MUMdex Source Code for G-Graph: determine_cn_sex.cpp

```
//
// determine_cn_sex
//
// quick and dirty sex determination from copy number results
//
// Copyright 2017 Peter Andrews @ CSHL
//

#include <algorithm>
#include <exception>
#include <sstream>
#include <string>
#include <vector>

#include "error.h"
#include "tsv.h"

using std::cerr;
using std::cout;
using std::endl;
using std::exception;
using std::string;
using std::vector;

using paa::sout;
using paa::Error;
using paa::TSV;

int main(int argc, char* argv[])  try {
  // Check usage
  if (--argc != 1) {
    throw Error("usage: determine_cn_sex cn_results_file");
  }

  const string cn_results_name{argv[1]};
  const TSV tsv{cn_results_name};

  const uint64_t chr_col{tsv.index("chr")};
  const uint64_t ratio_col{tsv.index("ratio")};

  const string regions{"AXY"};

  vector<vector<double> > ratios(3);
  for (uint64_t r{0}; r != tsv.n_rows(); ++r) {
    const string chr{tsv.as_string(chr_col, r)};
    const bool is_x{chr.find('X') != string::npos};
    const bool is_y{chr.find('Y') != string::npos};
    const uint64_t region_index{is_x ? 1U : (is_y ? 2U : 0U)};
    ratios[region_index].push_back(tsv.as_real(ratio_col, r));
  }

  cout << cn_results_name;
  for (uint64_t r{0}; r != ratios.size(); ++r) {
    vector<double> & region_ratios{ratios[r]};
    sort(region_ratios.begin(), region_ratios.end());
    cout << " " << regions[r] << " "
         << region_ratios[region_ratios.size() / 2];
  }
  cout << endl;

  return 0;
} catch (Error & e) {
  cerr << "paa::Error:" << endl;
  cerr << e.what() << endl;
  return 1;
} catch (exception & e) {
  cerr << "std::exception" << endl;
  cerr << e.what() << endl;
  return 1;
} catch (...) {
  cerr << "unknown exception was caught" << endl;
  return 1;
}
```
