## Supplementary material for "G-Graph: An interactive genomic graph viewer": MUMdex Source Code for G-Graph: determine_sex.cpp

```
//
// determine_sex
//
// quick and dirty sex determination from mumdex
//
// Copyright 2016 Peter Andrews @ CSHL
//

#include <exception>
#include <sstream>
#include <string>
#include <vector>

#include "error.h"
#include "mumdex.h"
#include "population.h"
#include "threads.h"
#include "utility.h"

using std::cerr;
using std::cout;
using std::endl;
using std::exception;
using std::ostringstream;
using std::string;
using std::vector;

using paa::sout;
using paa::ChromosomeIndexLookup;
using paa::Error;
using paa::Family;
using paa::Mappability;
using paa::MUM;
using paa::MUMdex;
using paa::Pair;
using paa::Population;
using paa::Reference;
using paa::Sample;
using paa::ThreadPool;

int main(int argc, char* argv[])  try {
  // Check usage
  if (--argc < 4) {
    throw Error("usage: determine_sex ref samples_dir pop_file "
                "[sample|family] ...");
  }

  // Process command line arguments
  const Reference ref{argv[1]};
  const Mappability mappability{ref};
  const ChromosomeIndexLookup chr_lookup{ref};
  const string samples_dir{argv[2]};
  const Population pop{argv[3]};

  argc -= 3;
  argv += 3;

  const unsigned int x{ref.find_x_chromosome()};
  const unsigned int y{ref.find_y_chromosome()};

  // Determine expected mappable positions
  const unsigned int min_length{100};
  vector<unsigned int> expected(ref.n_chromosomes());
  for (unsigned int c{0}; c != ref.n_chromosomes(); ++c) {
    for (unsigned int p{0}; p != ref.size(c); ++p) {
      const unsigned int abspos{ref.abspos(c, p)};
      const unsigned int mapr{mappability.low(abspos)};
      if (mapr <= min_length) {
        ++expected[c];
      }
    }
  }

  ThreadPool pool{4};
  ThreadPool::Results<string> unordered_results;

  while (argc--) {
    // Get samples for sample or family name
    const string sample_or_family{argv++[1]};
    for (const Sample s : pop.samples(sample_or_family)) {
      pool.run(unordered_results,
               [&pop, &samples_dir, &ref, &mappability, expected, x, y]
               (const Sample & sample) {
          // Get sample info
          const Family family{pop.family(sample)};
          const string sample_name{pop.sample(sample)};
          const string family_name{pop.family(family)};

          // Open mumdex file
          ostringstream mumdex_name;
          mumdex_name << samples_dir << "/"
                      << sample_name << "/mumdex";
          const MUMdex mumdex{mumdex_name.str(), &ref};

          // Count mappings
          vector<uint64_t> counts(ref.n_chromosomes());
          for (uint64_t p{0}; p != mumdex.n_pairs(); ++p) {
            const Pair pair{mumdex.pair(p)};
            if (pair.dupe()) continue;
            for (auto mi = mumdex.mums_begin(p); mi != mumdex.mums_end(p);
                 ++mi) {
              const MUM mum{*mi};
              const unsigned int abspos{ref.abspos(mum.chromosome(),
                                                   mum.position0())};
              const unsigned int mapp{mappability.low(abspos)};
              if (mum.length() >= min_length + 25 && mapp <= min_length) {
                ++counts[mum.chromosome()];
              }
            }
          }

          // Autosome map density
          uint64_t auto_count{0};
          uint64_t auto_expected{0};
          for (unsigned int c{0}; c != ref.n_chromosomes(); ++c) {
            if (c == x || c == y) continue;
            auto_count += counts[c];
            auto_expected += expected[c];
          }
          const double auto_density{1.0 * auto_count / auto_expected};

          // X and Y map density
          unsigned int chrs[2]{x, y};
          double ratio[2]{0, 0};
          for (const unsigned int c : {0, 1}) {
            const unsigned int chr{chrs[c]};
            ratio[c] = 1.0 * counts[chr] / expected[chr] / auto_density;
          }

          const string sex{[ratio]() {
              string result;
              const string chars{"XY"};
              for (const unsigned int c : {0, 1}) {
                for (unsigned int i{1}; i < 2.2 * ratio[c]; ++i) {
                  result += chars[c];
                }
              }
              return result;
            }()};

          ostringstream out;
          out << family_name << " "
              << sample_name << " "
              << pop.member(sample) << " "
              << pop.sex(sample) << " "
              << ratio[0] << " "
              << ratio[1] << " "
              << sex;

          return out.str();
               }, s);
    }
  }

  while (unordered_results.size()) {
    const string result{unordered_results.get()};
    cout << result << endl;
  }

  cerr << "done" << endl;

  return 0;
} catch (Error & e) {
  cerr << "paa::Error:" << endl;
  cerr << e.what() << endl;
  return 1;
} catch (exception & e) {
  cerr << "std::exception" << endl;
  cerr << e.what() << endl;
  return 1;
} catch (...) {
  cerr << "unknown exception was caught" << endl;
  return 1;
}
```
