## Supplementary material for "G-Graph: An interactive genomic graph viewer": MUMdex Source Code for G-Graph: distinct_colors.cpp

```
//
// distinct_colors.cpp
//
// Get a list of distinct colors
//
// Copyright 2018 Peter Andrews @ CSHL
//

#include <exception>
#include <iostream>
#include <string>
#include <vector>

#include "error.h"
#include "x11plot.h"

using std::cerr;
using std::cout;
using std::endl;
using std::exception;
using std::string;
using std::vector;

using paa::Error;
using paa::X11App;
using paa::X11Colors;
using paa::X11Win;

int main(int argc, char * argv[]) try {
  if (--argc != 1) throw Error("usage: distinct_colors n_colors");

  // Target colors
  const unsigned int n_colors{static_cast<unsigned int>(atoi(argv[1]))};

  // Starting colors
  vector<string> starting_colors{
    "rgb:e5/00/00", "rgb:25/00/9e", "rgb:00/b7/00", "rgb:e5/be/00"};
  vector<string> extra_colors{
    "rgb:06/56/93", "rgb:b7/dd/00", "rgb:e5/83/00", "rgb:95/00/95"};
  if (true) starting_colors.insert(starting_colors.end(),
                                   extra_colors.begin(), extra_colors.end());

  // The App
  X11App app;
  X11Colors & colors{app.create<X11Colors>(starting_colors, n_colors)};
  app.create<X11Colors>(colors.color_names, 0, true);
  if (true) colors.print_names();
  if (true) colors.print_fracs();
  app.run();

  return 0;
} catch (Error & e) {
  cerr << "Error:" << endl;
  cerr << e.what() << endl;
  return 1;
} catch (exception & e) {
  cerr << "exception" << endl;
  cerr << e.what() << endl;
  return 1;
} catch (...) {
  cerr << "unknown exception was caught" << endl;
  return 1;
}
```
