## Supplementary material for "G-Graph: An interactive genomic graph viewer": MUMdex Source Code for G-Graph: encode.cpp

```
//
// encode
//
// encode.cpp tests operation of encode.h
//
// Copyright 2015 Peter Andrews @ CSHL
//

#include "encode.h"

#include <exception>
#include <iostream>
#include <fstream>
#include <memory>
#include <sstream>
#include <string>
#include <vector>

#include "error.h"

using std::cerr;
using std::cout;
using std::endl;
using std::exception;
using std::ifstream;
using std::make_unique;
using std::ostringstream;
using std::string;
using std::unique_ptr;
using std::vector;

using paa::EncodedString;
using paa::Encoder;
using paa::Error;
using paa::FixedLengthOptionalSaver;
using paa::FixedLengthStrings;
using paa::IntPacker;
using paa::LCM;
using paa::OptionalFinder;
using paa::OptionalSaver;
using paa::OptionalSavers;
using paa::ReadSubstringExtractor;
using paa::ReadTagOptionalSaver;
using paa::VariableLengthOptionalSaver;

int main(int argc, char * argv[]) try {
  if (--argc > 1) throw Error("usage: encode [input]");

  if (argc == 1) {
    const string input_name{argv[1]};
    ifstream input_file{input_name.c_str()};

    string input;
    char inc;
    while (input_file.get(inc)) {
      input += inc;
    }
    EncodedString encoding{input};
    const string input_decoded = encoding();
    if (input != input_decoded)
      throw Error("EncodedString decode error");
    cout << "Packing is " << encoding.packing() << endl;
  }

  string alphabet{"ACGTN"};
  Encoder encoder{alphabet};
  encoder.show_encoding(cout);

  // Test packer for proper operation for many setups
  vector<uint64_t> inputs{0, 9, 1, 2, 99, 22, 32, 54, 65, 77, 222, 18, 29,
        45, 76, 38, 27, 37, 46, 59, 87, 99, 2020, 10, 2, 3, 4, 5, 20, 11, 276};
  for (uint64_t n_ints = 2; n_ints != 1000; ++n_ints) {
    IntPacker packer(n_ints);
    for (uint64_t i = 0; i != inputs.size(); ++i) {
      packer.push_back(inputs[i] % n_ints);
    }
    for (uint64_t i = 0; i != inputs.size(); ++i) {
      if (packer[i] != (inputs[i] % n_ints)) {
        throw Error("disagreement") << n_ints << i << (inputs[i] % n_ints);
      }
    }
  }

  for (uint64_t length = 1; length != 20; ++length) {
    string input_string{"ATTTTGTAGNAAGGAACATCTGTGCNNNNGNAGAACCATAGGCCNNNNNCCCCT"
          "GCCCCNGCTNTTNNNNNCTGTGACCACTGTTNNNNTCCCCNGCTNTTNNNNNCTGTGACCACCNGC"};
    vector<string> input_strings = [&input_string, length]() {
      vector<string> input_str;
      for (uint64_t i = 0; i + length < input_string.size(); ++i) {
        input_str.push_back(input_string.substr(i, length));
      }
      return input_str;
    }();
    FixedLengthStrings strings{"ACGTN", length};
    for (const auto & str : input_strings) {
      strings.push_back(str);
    }
    if (strings.size() != input_strings.size())
      throw Error("Unexpected stored strings size mismatch");
    for (uint64_t i = 0; i != strings.size(); ++i) {
      if (strings[i] != input_strings[i])
        throw Error("Problem retrieving stored string") << i;
      for (uint64_t c = 0; c != length; ++c) {
        if (strings(i, c) != input_strings[i][c])
          throw Error("Problem retrieving stored character") << i << c;
      }
    }
  }

  ReadSubstringExtractor tag_extractor{"ZV:Z:", 0, 15, 32};
  const OptionalFinder finder("ZV:Z:");
  const string optional{"XT:A:U\tNM:i:1\tSM:i:29\tAM:i:29\tX0:i:1\tX1:i:0"
        "\tXM:i:1\tXO:i:0\tXG:i:0\tMD:Z:75T1\tZR:Z:ACGACAGCT;61;BCR0"
        "03-2.6;GAGTGCT;65;BCR003-2.6\tZO:Z:ACGACAGCT;@?@7DDFFG;GAGTGCT;B@CDD"
        "FF\tZF:Z:111;1437\tZV:Z:CTCATAGCAAACATG/GFFDHIGFIIIGGEC/CTCTTATAA"
        "CGCATG/?HHHGDIJJIJIJIJ mooo\t"};
  cout << finder(optional) << endl;
  cout << tag_extractor(optional, 0) << " "
       << tag_extractor(optional, 1) << endl;
  const OptionalFinder finder2("");
  cout << finder2(optional) << endl;

  const vector<uint16_t> numbers{1, 15000, 42, 65000, 0};
  FixedLengthStrings numbers1{16};
  FixedLengthStrings numbers2{16, true};
  for (const uint16_t number : numbers) {
    numbers1.push_back_binary(number);
    numbers2.push_back(number);
  }
  for (uint64_t n = 0; n != numbers.size(); ++n) {
    cout << numbers1[n] << " " << numbers[n] << " "
         << numbers1.binary_get_int(n) << " "
         << numbers2.get_int(n) << endl;
  }
  cout << numbers2.binary_get_int(3) << endl;
  numbers1.push_back("0011101010011000");
  cout << numbers1[5] << " " << numbers1.binary_get_int(5) << endl;
  cout << endl;

  IntPacker bitPacker{1, true};
  bitPacker.push_back(0);
  bitPacker.push_back(1);
  bitPacker.push_back(1);
  bitPacker.push_back(0);
  bitPacker.push_back(0);
  bitPacker.push_back(1);
  for (uint64_t n = 0; n != bitPacker.size(); ++n) {
    cout << bitPacker[n] << endl;
  }
  cout << endl;

  ifstream sample_input{"sample.opt.txt"};
  string line;
  vector<string> read_info;
  while (getline(sample_input, line)) {
    read_info.push_back(line);
  }

  const string quality_chars =
      "!\"#$%&'()*+,-./0123456789:;<=>?@ABCDEFGHIJ";
  const string other_bases = "TN";
  const string bases = "ACGTN";
  const string qualities_and_bases = quality_chars + other_bases;
  vector<unique_ptr<OptionalSaver>> savers;
  savers.push_back(make_unique<VariableLengthOptionalSaver>(
      "err", "", quality_chars, 79));
  savers.push_back(make_unique<FixedLengthOptionalSaver>(
      "fix.XT", "XT:A:", "UMR", 1));
  savers.push_back(make_unique<FixedLengthOptionalSaver>(
      "fix.ZV", "ZV:Z:", qualities_and_bases, 63));
  savers.push_back(make_unique<ReadTagOptionalSaver>(
      "tag.ZV", "ZV:Z:", bases, 15, 0, 32));

  for (unsigned int i = 0; i != read_info.size(); ++i) {
    const auto & read = read_info[i];
    for (auto & saver : savers) {
      saver->extract(read, i % 2);
    }
  }

  vector<string> saved;
  for (unsigned int i = 0; i != savers.front()->size(); ++i) {
    ostringstream out;
    for (const auto & saver : savers) {
      out << (*saver)[i] << " ";
    }
    out << endl;
    saved.push_back(out.str());
    cout << out.str();
  }

  const vector<string> formats = {
    "ERR|79",
    "FIX|1|XT:A:|UMR",
    "FIX|63|ZV:Z:|!\"#$%&'()*+,-./0123456789:;<=>?@ABCDEFGHIJTN",
    "TAG|15|ZV:Z:|0|32" };

  OptionalSavers saver{formats};
  for (unsigned int r = 0; r != read_info.size(); r += 2) {
    saver.extract(read_info[r], read_info[r + 1]);
  }

  vector<string> reconstructed;
  for (unsigned int r = 0; r != read_info.size(); ++r) {
    ostringstream out;
    for (unsigned int s = 0; s != saver.size(); ++s) {
      out << saver[s][r] << " ";
    }
    out << endl;
    reconstructed.push_back(out.str());
  }
  if (saved != reconstructed) {
    throw Error("mismatch saver");
  }

  std::vector<FILE *> optional_files;
  for (unsigned int i = 0; i != saver.size(); ++i) {
    const std::string file_name{saver[i].file_name("mumdex")};
    std::FILE * file = std::fopen(file_name.c_str(), "wb+");
    if (file == nullptr) throw Error("Problem opening file") << file_name;
    optional_files.push_back(file);
  }
  saver.write_and_reduce(optional_files, true);
  for (auto file : optional_files) {
    if (fclose(file)) {
      throw Error("problem closing optional file");
    }
  }

  OptionalSavers reader{formats};
  reader.load("mumdex", read_info.size());

  reconstructed.clear();
  for (unsigned int r = 0; r != read_info.size(); ++r) {
    ostringstream out;
    for (unsigned int s = 0; s != reader.size(); ++s) {
      out << reader[s][r] << " ";
    }
    out << endl;
    reconstructed.push_back(out.str());
  }
  if (saved != reconstructed) {
    throw Error("mismatch reader");
  }
  cout << endl;

  cout << LCM(2, 3) << endl;
  cout << LCM(3, 3) << endl;
  cout << LCM(4, 3) << endl;
  cout << LCM(2, 6) << endl;
  cout << LCM(20, 12) << endl;

  return 0;
} catch (Error & e) {
  cerr << "paa::Error:" << endl;
  cerr << e.what() << endl;
  return 1;
} catch (exception & e) {
  cerr << "std::exception" << endl;
  cerr << e.what() << endl;
  return 1;
} catch (...) {
  cerr << "unknown exception was caught" << endl;
  return 1;
}
```
