## Supplementary material for "G-Graph: An interactive genomic graph viewer": MUMdex Source Code for G-Graph: event_histogram.cpp

```
//
// event_histogram.cpp
//
// look for concentrations of events in genome space
//
// Copyright 2016 Peter Andrews @ CSHL
//

#include <algorithm>
#include <iostream>
#include <map>
#include <set>
#include <sstream>
#include <string>
#include <utility>
#include <vector>

#include "error.h"
#include "genes.h"
#include "mumdex.h"
#include "utility.h"

using std::cerr;
using std::cin;
using std::cout;
using std::endl;
using std::exception;
using std::istringstream;
using std::map;
using std::pair;
using std::set;
using std::string;
using std::vector;

using paa::sout;
using paa::Error;
using paa::ChromosomeIndexLookup;
using paa::GeneXrefs;
using paa::KnownGenes;
using paa::Reference;

struct PosTagInfo {
  PosTagInfo(const unsigned int chr_arg, const unsigned int pos_arg,
             const string & tag_arg) :
      chr{chr_arg}, pos{pos_arg}, tag{tag_arg} {}
  unsigned int chr;
  unsigned int pos;
  string tag;
};

class BinInfo {
 public:
  void add(const unsigned int chromosome, const unsigned int position,
           const string & tag) {
    positions_.emplace_back(chromosome, position, tag); }
  const vector<PosTagInfo> & positions() const { return positions_; }

 private:
  vector<PosTagInfo> positions_{};
};

int main(int argc, char ** argv) {
  try {
    paa::exit_on_pipe_close();
    if (--argc != 2) throw Error("usage: event_histogram reference bin_size");

    const string reference_file{argv[1]};
    const Reference ref{reference_file};
    const ChromosomeIndexLookup lookup{ref};
    const string genes_name{reference_file + ".bin/knownGene.txt"};
    const string isoforms_name{reference_file + ".bin/knownIsoforms.txt"};
    const string kgXrefs_name{reference_file + ".bin/kgXref.txt"};
    const KnownGenes genes{genes_name, isoforms_name, lookup, ref};
    const GeneXrefs xref{kgXrefs_name};

    const unsigned int bin_size{static_cast<unsigned int>(atoi(argv[2]))};
    const uint64_t n_bins{ref.size() / bin_size + 1};
    vector<BinInfo> bin_counts(n_bins);

    string line;
    while (getline(cin, line)) {
      string chromosome_name;
      unsigned int chromosome_position;
      string tag;
      istringstream in{line.c_str()};
      in >> chromosome_name >> chromosome_position;
      if (!in) throw Error("Problem reading line") << line;
      getline(in, tag);
      const unsigned int chromosome{lookup[chromosome_name]};
      const unsigned int abspos{ref.offset(chromosome) + chromosome_position};
      bin_counts[abspos / bin_size].add(chromosome, chromosome_position, tag);
    }

    if (1) {
      sort(bin_counts.begin(), bin_counts.end(),
           [](const BinInfo & left, const BinInfo & right) {
             return left.positions().size() < right.positions().size();
           });
    }

    for (const BinInfo & bin : bin_counts) {
      if (bin.positions().size()) {
        sout << bin.positions().size() << endl;
        set<string> tags;
        const vector<PosTagInfo> anchors{bin.positions()};
        for (const PosTagInfo anchor : anchors) {
          tags.emplace(anchor.tag);
        }
        for (const string & tag : tags) {
          cout << tag << endl;
        }
      }
    }

#if 0
    using chrpos = pair<unsigned int, unsigned int>;

        sort(anchors.begin(), anchors.end(),
             [] (const PosTagInfo & left, const PosTagInfo & right) {
               if (left.chr == right.chr) {
                 return left.pos < right.pos;
               } else {
                 return left.chr < right.chr;
               }
             });
        set<chrpos> positions;
        set<string> tags;
        map<string, set<string>> gene_tags;
        for (const PosTagInfo anchor : anchors) {
          positions.emplace(anchor.chr, anchor.pos);
          tags.emplace(anchor.tag);
          const vector<unsigned int> pos_genes{
            genes.find_genes(position.first, position.second)};
          for (const unsigned int gene : pos_genes) {
            const string name{xref[genes[gene].name].geneSymbol};
          }
        }
        sout << tags.size();

        // output positions
        string last_chromosome = "";
        vector<unsigned int> bin_genes;
        set<unsigned int> in_exon;
        for (const chrpos position : positions) {
          const string chromosome{ref.name(position.first)};
          if (chromosome != last_chromosome) {
            sout << chromosome;
            last_chromosome = chromosome;
          }
          sout << position.second;
          const vector<unsigned int> pos_genes{
            genes.find_genes(position.first, position.second)};
          bin_genes.insert(bin_genes.end(), pos_genes.begin(), pos_genes.end());
          for (const unsigned int gene : pos_genes) {
            for (const string & tag : pos_tags[position]) {
              gene_tags[gene].insert(tag);
            }
            if (genes[gene].in_exon(position.first, position.second)) {
              in_exon.emplace(gene);
            }
          }
        }

        // output tags
        if (tags.size()) sout << "in";
        for (const string & tag : tags) {
          sout << tag;
        }

        sort(bin_genes.begin(), bin_genes.end());
        bin_genes.erase(unique(bin_genes.begin(), bin_genes.end()),
                        bin_genes.end());
        set<string> bin_names;
        for (const unsigned int gene : bin_genes) {
          if (in_exon.count(gene)) {
            bin_names.insert(string("EXON-") +
                             xref[genes[gene].name].geneSymbol);
          } else {
            bin_names.insert(xref[genes[gene].name].geneSymbol);
          }
        }
        if (bin_names.size()) {
          sout << "in gene";
        }
        for (const string & gene : bin_names) {
          sout << gene;
        }

        if (bin_names.size()) {
          sout << ".  ";
        }
        for (const unsigned int gene : bin_genes) {
          cout << xref[genes[gene].name].geneSymbol << ": ";
          for (const string & tag : gene_tags[gene]) {
            cout << tag  << " ";
          }
        }
        sout << endl;
      }
    }
#endif
    return 0;
  }
  catch(exception & e) {
    cerr << e.what() << endl;
    return 1;
  }
  catch(...) {
    cerr << "Some exception was caught." << endl;
    return 1;
  }
  return 0;
}
```
