## Supplementary material for "G-Graph: An interactive genomic graph viewer": MUMdex Source Code for G-Graph: family_count.cpp

```
//
// family_count.cpp
//
// numerical simulation to determine
// for N families of 4 members each
// how many family coincidences are expected
// if M people are randomly selected
//
// Copyright 2016 Peter Andrews @ CSHL
//

#include <algorithm>
#include <exception>
#include <iostream>
#include <random>
#include <set>
#include <vector>

#include "error.h"
#include "stats.h"
#include "utility.h"

using std::cout;
using std::cerr;
using std::endl;
using std::exception;
using std::set;
using std::vector;

using paa::Error;
using paa::NormalParams;
using paa::sout;

struct Person {
  Person(const unsigned int f, const unsigned int i) :
      family(f), individual(i) { }
  uint16_t family;
  uint16_t individual;
};

int main(int argc, char* argv[], char * []) try {
  if (--argc != 3)
    throw Error("usage: family_count n_families n_trials n_repeats");

  const unsigned int n_families{static_cast<unsigned int>(atoi(argv[1]))};
  const unsigned int n_trials{static_cast<unsigned int>(atoi(argv[2]))};
  const unsigned int n_repeats{static_cast<unsigned int>(atoi(argv[3]))};
  const unsigned int n_in_family{4};
  const unsigned int n_people{n_families * n_in_family};

  vector<Person> people;
  people.reserve(n_people);
  for (unsigned int f{0}; f != n_families; ++f) {
    for (unsigned int m{0}; m != n_in_family; ++m) {
      people.emplace_back(f, m);
    }
  }

  auto mersenne = std::mt19937_64();
  mersenne.seed(time(nullptr));

  vector<vector<unsigned int>> counts(n_people,
                                      vector<unsigned int>(n_repeats));
  for (unsigned int r{0}; r != n_repeats; ++r) {
    for (unsigned int t{0}; t != n_trials; ++t) {
      shuffle(people.begin(), people.end(), mersenne);
      set<unsigned int> families;
      unsigned int repeats_seen{0};
      for (unsigned int p{0}; p != n_people; ++p) {
        repeats_seen += !families.insert(people[p].family).second;
        counts[p][r] += repeats_seen;
      }
    }
  }

  for (unsigned int p{1}; p != n_people; ++p) {
    NormalParams norm(counts[p]);
    sout << p + 1
         << norm.mean / n_trials
         << norm.stdev / n_trials / sqrt(n_repeats)
         << norm.stdev / norm.mean / sqrt(n_repeats);
    if (0) {
      for (unsigned int r{0}; r != n_repeats; ++r) {
        sout << 1.0 * counts[p][r] / n_trials;
      }
    }
    sout  << endl;
  }

  return 0;
} catch (Error & e) {
  cerr << "paa::Error:" << endl;
  cerr << e.what() << endl;
  return 1;
} catch (exception & e) {
  cerr << "std::exception" << endl;
  cerr << e.what() << endl;
  return 1;
} catch (...) {
  cerr << "unknown exception was caught" << endl;
  return 1;
}
```
