## Supplementary material for "G-Graph: An interactive genomic graph viewer": MUMdex Source Code for G-Graph: family_figure.cpp

```
//
// family_figure
//
// plot invariants over a region for a family, considering population
//
// Copyright 2017 Peter Andrews @ CSHL
//

#include <algorithm>
#include <deque>
#include <exception>
#include <functional>
#include <iostream>
#include <queue>
#include <random>
#include <sstream>
#include <string>
#include <utility>
#include <vector>

#include "bridges.h"
#include "error.h"
#include "files.h"
#include "mumdex.h"
#include "psplot.h"

using std::bind;
using std::cout;
using std::cerr;
using std::deque;
using std::endl;
using std::exception;
using std::function;
using std::lower_bound;
using std::move;
using std::mt19937_64;
using std::ostringstream;
using std::priority_queue;
using std::string;
using std::uniform_real_distribution;
using std::vector;

using paa::nunset;
using paa::unset;
using paa::Bounds;
using paa::BridgeInfo;
using paa::ChromosomeIndexLookup;
using paa::Error;
using paa::Family;
using paa::Mappability;
using paa::MappedVector;
using paa::Marker;
using paa::MergeHelper;
using paa::MergeHelperCompare;
using paa::PopBridgeInfo;
using paa::Population;
using paa::PSDoc;
using paa::PSGraph;
using paa::PSPage;
using paa::PSXYMSeries;
using paa::PSXYSeries;
using paa::Reference;
using paa::Sample;

using Info = PopBridgeInfo;

const Reference * paa::ref_ptr;
using paa::ref_ptr;

int main(int argc, char* argv[], char * []) try {
  if (--argc != 7)
    throw Error("usage: family_figure family_file bridges_dir "
                "ref chr start stop family");

  const unsigned int min_support_length{25};
  const unsigned int min_mate_count{1};
  const unsigned int min_mate_support{20};
  const unsigned int min_excess_mappability{1};

  // Arguments
  const Population pop{argv[1]};
  const string bridges_dir{argv[2]};
  const Reference ref{argv[3]};
  ref_ptr = &ref;
  const Mappability map{ref};
  const ChromosomeIndexLookup lookup{ref};
  const string chromosome_name{argv[4]};
  const unsigned int chromosome{lookup[chromosome_name]};
  const unsigned int chromosome_offset{ref.offset(chromosome)};
  const unsigned int start{static_cast<unsigned int>(atol(argv[5]))};
  const unsigned int stop{static_cast<unsigned int>(atol(argv[6]))};
  const string family_name{argv[7]};

  // Bridges file
  ostringstream pop_bridges_file_name;
  pop_bridges_file_name << bridges_dir << "/popbridges."
                        << chromosome_name
                        << ".bin";
  const MappedVector<Info> pop_bridges{pop_bridges_file_name.str()};
  // Bridge range limits
  auto pop_less = [](const Info & lhs, const unsigned int pos) {
    return lhs.pos1() < pos;
  };
  const Info * const pop_lower{lower_bound(
      pop_bridges.begin(), pop_bridges.end(), start, pop_less)};
  const Info * const pop_upper{lower_bound(
      pop_bridges.begin(), pop_bridges.end(), stop, pop_less)};
  const Info * pop_current{pop_lower};

  // Plots
  const Marker negative_marker{paa::circle(), 0.2, "1 0 0", 1, true, "1 0 0"};
  const Marker positive_marker{paa::circle(), 0.2, "0 0 1", 1, true, "0 0 1"};
  const Marker bnegative_marker{paa::circle(), 0.5, "1 0 0", 1, true, "1 0 0"};
  const Marker bpositive_marker{paa::circle(), 0.5, "0 0 1", 1, true, "0 0 1"};

  const double miny{1.0};
  const double maxy{1000000000};

  PSDoc doc{"family_invariants"};
  doc.pdf(false);
  const string title{1 ? string("") :string("Indel Invariants for region on ") +
        chromosome_name + " for SSC family " + family_name};
  PSPage page{doc, title, "1 2"};
  PSGraph dirty_graph{page, "Unfiltered;Position;|Invariant|",
        Bounds{1.0 * start, 1.0 * stop, miny, maxy}};
  dirty_graph.log_y(true);
  PSXYMSeries dirty{dirty_graph};
  PSPage page2{doc, title, "1 2"};
  PSGraph map_graph{page, "Map Length Cut;Position;|Invariant|",
        Bounds{1.0 * start, 1.0 * stop, miny, maxy}};
  map_graph.log_y(true);
  PSXYMSeries map_series{map_graph};
  PSGraph mate_graph{page2,
        "Mate Cut;Position;|Invariant|",
        Bounds{1.0 * start, 1.0 * stop, miny, maxy}};
  mate_graph.log_y(true);
  PSXYMSeries mate{mate_graph};
  PSGraph popcut_graph{page2,
        "Population Filtered (ultra-rare only);Position;|Invariant|",
        Bounds{1.0 * start, 1.0 * stop, miny, maxy}};
  popcut_graph.log_y(true);
  PSXYMSeries popcut{popcut_graph};

  PSPage page3{doc, title, "1 2"};
  PSGraph strong_graph{page3, "Many Bridges Cut;Position;|Invariant|",
        Bounds{1.0 * start, 1.0 * stop, miny, maxy}};
  strong_graph.log_y(true);
  PSXYMSeries strong_series{strong_graph};
  PSGraph final_graph{page3, "Not in Parent Cut;Position;|Invariant|",
        Bounds{1.0 * start, 1.0 * stop, miny, maxy}};
  final_graph.log_y(true);
  PSXYMSeries final_series{final_graph};

  // Random scatter
  auto mersenne = std::mt19937_64();
  mersenne.seed(time(nullptr));
  uniform_real_distribution<double> dist{0, 1};
  function<double()> gen{bind(dist, mersenne)};

  cout << "chr1\tpos1\thigh1\tchr2\tpos2\thigh2\tinvariant\toffset\t"
       << "n_in_family\tn_in_parents\tmax_in_kid\t"
       << "pop_people\tpop_bridges\tpop_median\tpop_max\t"
       << "support1\tsupport2\tmate_count1\tmate_count2\t"
       << "mate_support1\tmate_support2\tmap1\tmap2" << endl;

  // Set up queue of bridge files
  deque<MergeHelper> helpers;
  using PQueue = priority_queue<MergeHelper*, vector<MergeHelper *>,
      MergeHelperCompare>;
  PQueue queue{MergeHelperCompare()};
  unsigned int n_samples_skipped{0};
  for (const auto s : pop.samples(pop.family(family_name))) {
    ostringstream bridges_name;
    bridges_name << bridges_dir << "/" << pop.sample(s) << "/"
                 << get_bridge_file_name(ref, chromosome);
    MergeHelper helper{bridges_name.str(), s, start, stop};
    if (helper.valid_start()) {
      helpers.push_back(move(helper));
      queue.push(&helpers.back());
    } else {
      ++n_samples_skipped;
    }
    cerr << pop.family(pop.family(s)) << " "
         << pop.sample(s) << " "
         << pop.member(s) << " "
         << bridges_name.str() << " "
         << queue.size() << endl;
  }

  vector<BridgeInfo> all_bridges;
  vector<Sample> all_samples;
  bool reset{true};
  while (true) {
    // New bridge expected - start fresh
    if (reset) {
      all_bridges.clear();
      all_samples.clear();
      reset = false;
    }

    // Add a bridge to the list if the same event
    bool bridge_done{true};
    if (queue.size()) {
      MergeHelper * top{queue.top()};
      const BridgeInfo current{top->current()};
      if (all_bridges.empty() || !(all_bridges.back() < current)) {
        bridge_done = false;
        all_bridges.push_back(current);
        const Sample sample{top->sample()};
        all_samples.emplace_back(sample);
        queue.pop();
        if (top->advance()) {
          queue.push(top);
        }
      }
      if (queue.empty()) bridge_done = true;
      if (bridge_done) reset = true;
    } else {
      if (all_samples.empty()) {
        break;
      }
    }

    // No new bridges added, so bridge list is ready to process
    if (bridge_done) {
      // The first bridge, the exemplar since all are the same
      const BridgeInfo & bridge{all_bridges[0]};
      if (bridge.chr1() != bridge.chr2()) continue;
      if (bridge.invariant() == 0) continue;
      if (bridge.high1() == bridge.high2()) continue;

      dirty.add_point(bridge.pos1() + gen(),
                      labs(bridge.invariant()),
                      bridge.invariant() > 0 ?
                      positive_marker: negative_marker);

      const unsigned int mum1_abspos{chromosome_offset + bridge.pos1()};
      const unsigned int mum2_abspos{ref.offset(bridge.chr2()) + bridge.pos2()};
      const unsigned int mum1_map{map.low_high(bridge.high1(), mum1_abspos)};
      const unsigned int mum2_map{map.low_high(bridge.high2(), mum2_abspos)};

      unsigned int max_anchor1_support{0};
      unsigned int max_anchor2_support{0};
      unsigned int max_mate_anchor1_count{0};
      unsigned int max_mate_anchor1_support{0};
      unsigned int max_mate_anchor2_count{0};
      unsigned int max_mate_anchor2_support{0};
      for (const BridgeInfo & sample_bridge : all_bridges) {
        if (max_anchor1_support < sample_bridge.anchor1_length()) {
          max_anchor1_support = sample_bridge.anchor1_length();
        }
        if (max_anchor2_support < sample_bridge.anchor2_length()) {
          max_anchor2_support = sample_bridge.anchor2_length();
        }
        if (max_mate_anchor1_count < sample_bridge.mate_anchor1_count()) {
          max_mate_anchor1_count = sample_bridge.mate_anchor1_count();
        }
        if (max_mate_anchor2_count < sample_bridge.mate_anchor2_count()) {
          max_mate_anchor2_count = sample_bridge.mate_anchor2_count();
        }
        if (max_mate_anchor1_support < sample_bridge.mate_anchor1_length()) {
          max_mate_anchor1_support = sample_bridge.mate_anchor1_length();
        }
        if (max_mate_anchor2_support < sample_bridge.mate_anchor2_length()) {
          max_mate_anchor2_support = sample_bridge.mate_anchor2_length();
        }
      }

      unsigned int n_parents{0};
      unsigned int max_in_kid{0};
      for (unsigned int s{0}; s != all_samples.size(); ++s) {
        const Sample sample{all_samples[s]};
        if (pop.is_parent(sample)) {
          ++n_parents;
        } else {
          if (all_bridges[s].bridge_count() > max_in_kid) {
            max_in_kid = all_bridges[s].bridge_count();
          }
        }
      }

      while (*pop_current < bridge && pop_current != pop_upper) {
        ++pop_current;
      }

      const bool in_pop{!(pop_current == pop_upper ||
                          *pop_current < bridge || bridge < *pop_current)};

      cout << ref.name(bridge.chr1())
           << "\t" << bridge.pos1()
           << "\t" << bridge.high1()
           << "\t" << ref.name(bridge.chr2())
           << "\t" << bridge.pos2()
           << "\t" << bridge.high2()
           << "\t" << bridge.invariant()
           << "\t" << bridge.offset()
           << "\t" << all_samples.size()
           << "\t" << n_parents
           << "\t" << max_in_kid
           << "\t" << (in_pop ? pop_current->n_people() : 0)
           << "\t" << (in_pop ? pop_current->n_bridges() : 0)
           << "\t" << (in_pop ? pop_current->median_bridges() : 0)
           << "\t" << (in_pop ? pop_current->max_bridges() : 0)
           << "\t" << max_anchor1_support
           << "\t" << max_anchor2_support
           << "\t" << max_mate_anchor1_count
           << "\t" << max_mate_anchor2_count
           << "\t" << max_mate_anchor1_support
           << "\t" << max_mate_anchor2_support
           << "\t" << mum1_map
           << "\t" << mum2_map
           << endl;

      // Anchor support
      if (max_anchor1_support < min_support_length ||
          max_anchor2_support < min_support_length) continue;

      // Short excess mappability cut
      if (max_anchor1_support < mum1_map) throw Error("Mappability");
      if (max_anchor1_support < min_excess_mappability + mum1_map) {
        continue;
      } else {
        if (max_anchor2_support < min_excess_mappability + mum2_map) {
          continue;
        }
      }

      map_series.add_point(bridge.pos1() + gen(),
                           labs(bridge.invariant()),
                           bridge.invariant() > 0 ?
                           positive_marker: negative_marker);

      // Short mate support cut
      if (max_mate_anchor1_count < min_mate_count ||
          max_mate_anchor2_count < min_mate_count ||
          max_mate_anchor1_support < min_mate_support ||
          max_mate_anchor2_support < min_mate_support) continue;

      mate.add_point(bridge.pos1() + gen(),
                     labs(bridge.invariant()),
                     bridge.invariant() > 0 ?
                     positive_marker: negative_marker);

      if (!in_pop ||
          pop_current->n_people() <= n_parents) {
        popcut.add_point(bridge.pos1() + gen(),
                         labs(bridge.invariant()),
                         bridge.invariant() > 0 ?
                         bpositive_marker: bnegative_marker);
        if (max_in_kid >= 5) {
          strong_series.add_point(bridge.pos1() + gen(),
                                 labs(bridge.invariant()),
                                 bridge.invariant() > 0 ?
                                 bpositive_marker: bnegative_marker);
          if (n_parents == 0) {
            final_series.add_point(bridge.pos1() + gen(),
                                   labs(bridge.invariant()),
                                   bridge.invariant() > 0 ?
                                   bpositive_marker: bnegative_marker);
          }
        }
      }
    }
  }

  return 0;
} catch (Error & e) {
  cerr << "paa::Error:" << endl;
  cerr << e.what() << endl;
  return 1;
} catch (exception & e) {
  cerr << "std::exception" << endl;
  cerr << e.what() << endl;
  return 1;
} catch (...) {
  cerr << "unknown exception was caught" << endl;
  return 1;
}
```
