## Supplementary material for "G-Graph: An interactive genomic graph viewer": MUMdex Source Code for G-Graph: fastq_kmer_counter.cpp

```
//
// fastq_kmer_counter
//
// counts kmers
//
// Copyright 2017 Peter Andrews @ CSHL
//

#include <exception>
#include <fstream>
#include <iostream>
#include <map>
#include <string>
#include <sstream>
#include <vector>

#include "error.h"
#include "mumdex.h"

using std::cerr;
using std::cout;
using std::endl;
using std::exception;
using std::ifstream;
using std::istringstream;
using std::map;
using std::string;
using std::vector;

using paa::reverse_complement;
using paa::Error;

int main(int argc, char* argv[])  try {
  if (--argc != 3)
    throw Error("usage: fastq_kmer_counter kmer_size fastq1 fastq2");

  const unsigned int kmer_size{static_cast<unsigned int>(atoi(argv[1]))};

  ifstream fastq1{argv[2]};
  ifstream fastq2{argv[3]};
  if (!fastq1 || !fastq2) throw Error("Could not open fastq");
  ifstream * fastqs[2]{&fastq1, &fastq2};

  string line;
  string name;
  string read;
  char c;
  map<string, unsigned int> counts;
  while (fastq1 && fastq2) {
    for (const bool read2 : {false, true}) {
      ifstream & fastq{*fastqs[read2]};
      fastq >> c;
      if (!fastq) continue;
      getline(fastq, line);
      istringstream line_str{line.c_str()};
      line_str >> name;
      fastq >> read >> c >> line;
      for (unsigned int b{0}; b != read.size(); ++b) {
        const string kmer{read.substr(b, kmer_size)};
        if (kmer.size() < kmer_size) break;
        ++counts[kmer];
        ++counts[reverse_complement(kmer)];
      }
    }
  }

  for (const auto elem : counts) {
    cout << elem.first << " " << elem.second << endl;
  }

  return 0;
} catch (Error & e) {
  cerr << "paa::Error:" << endl;
  cerr << e.what() << endl;
  return 1;
} catch (exception & e) {
  cerr << "std::exception" << endl;
  cerr << e.what() << endl;
  return 1;
} catch (...) {
  cerr << "unknown exception was caught" << endl;
  return 1;
}
```
