## Supplementary material for "G-Graph: An interactive genomic graph viewer": MUMdex Source Code for G-Graph: fastq_mapper.cpp

```
//
// fastq_mapper
//
// simple mum mapper maps sequence from fastq
//
// Copyright 2017 Peter Andrews @ CSHL
//

#include <exception>
#include <fstream>
#include <iostream>
#include <string>
#include <sstream>
#include <vector>

#include "error.h"
#include "longSA.h"
#include "mapper.h"
#include "mumdex.h"
#include "utility.h"

using std::cerr;
using std::cin;
using std::cout;
using std::endl;
using std::exception;
using std::ifstream;
using std::istringstream;
using std::string;
using std::vector;

using paa::longSA;
using paa::read_ahead;
using paa::sout;
using paa::Error;
using paa::Reference;
using paa::SimpleHit;

int main(int argc, char* argv[])  try {
  paa::memory_mapped = false;
  read_ahead = false;
  std::ios_base::sync_with_stdio(false);
  cin.tie(nullptr);

  unsigned int min_len{1};
  if (--argc > 3) {
    const std::string option_name{argv[1]};
    if (option_name == "-l") {
      min_len = atoi(argv[2]);
      argc -= 2;
      argv += 2;
    }
  }
  if (argc != 3)
    throw Error("usage: mapper [-l min_len] ref_fasta fastq1 fastq2");

  const string ref_fasta{argv[1]};
  const longSA sa{ref_fasta.c_str(), true, paa::memory_mapped, false};
  const Reference ref{ref_fasta};

  ifstream fastq1{argv[2]};
  ifstream fastq2{argv[3]};
  if (!fastq1 || !fastq2) throw Error("Could not open fastq");
  ifstream * fastqs[2]{&fastq1, &fastq2};

  string line;
  string name;
  string read;
  char c;
  while (fastq1 && fastq2) {
    for (const bool read2 : {false, true}) {
      ifstream & fastq{*fastqs[read2]};
      fastq >> c;
      if (!fastq) continue;
      getline(fastq, line);
      istringstream line_str{line.c_str()};
      line_str >> name;
      fastq >> read >> c >> line;
      const vector<SimpleHit> mums{sa.find_mams(read, min_len)};
      for (const auto mum : mums) {
        sout << name << read2 << ref.name(mum.chr) << mum.pos
             << mum.off << mum.len << mum.dir << endl;
      }
    }
  }

  return 0;
} catch (Error & e) {
  cerr << "paa::Error:" << endl;
  cerr << e.what() << endl;
  return 1;
} catch (exception & e) {
  cerr << "std::exception" << endl;
  cerr << e.what() << endl;
  return 1;
} catch (...) {
  cerr << "unknown exception was caught" << endl;
  return 1;
}
```
