## Supplementary material for "G-Graph: An interactive genomic graph viewer": MUMdex Source Code for G-Graph: fastq_sequences.cpp

```
//
// fastq_sequences
//
//
// Copyright 2018 Peter Andrews @ CSHL
//

#include <iostream>
#include <string>

#include "error.h"
#include "utility.h"

using std::cerr;
using std::cin;
using std::cout;
using std::endl;
using std::exception;
using std::string;

using paa::Error;

int main(int argc, char **) try {
  if (--argc != 0) throw Error("usage: fastq_sequences");

  std::ios_base::sync_with_stdio(false);
  cin.tie(nullptr);

  uint64_t n{0};
  string line;
  while (getline(cin, line)) {
    if (n++ % 4 == 1) cout << line << '\n';
  }

  return 0;
} catch (Error & e) {
  cerr << "paa::Error:" << endl;
  cerr << e.what() << endl;
  return 1;
} catch(exception & e) {
  cerr << e.what() << endl;
  return 1;
} catch(...) {
  cerr << "Some exception was caught." << endl;
  return 1;
}
```
