## Supplementary material for "G-Graph: An interactive genomic graph viewer": MUMdex Source Code for G-Graph: fastq_vt_mapper.cpp

```
//
// fastq_vt_mapper
//
// simple mum mapper maps sequence from fastq
// after processing varietal tag
//
// Copyright 2017 Peter Andrews @ CSHL
//

#include <exception>
#include <fstream>
#include <iostream>
#include <string>
#include <sstream>
#include <vector>

#include "error.h"
#include "longSA.h"
#include "mapper.h"
#include "mumdex.h"
#include "utility.h"

using std::cerr;
using std::cin;
using std::cout;
using std::endl;
using std::exception;
using std::ifstream;
using std::istringstream;
using std::string;
using std::vector;

using paa::longSA;
using paa::read_ahead;
using paa::sout;
using paa::Error;
using paa::Reference;
using paa::SimpleHit;

int main(int argc, char* argv[])  try {
  read_ahead = true;
  if (--argc != 3)
    throw Error("usage: fastq_vt_mapper ref_fasta fastq1 fastq2");

  const string ref_fasta{argv[1]};
  const longSA sa{ref_fasta.c_str(), true, true, false};
  const Reference ref{ref_fasta};

  ifstream fastq1{argv[2]};
  ifstream fastq2{argv[3]};
  if (!fastq1 || !fastq2) throw Error("Could not open fastq");
  ifstream * fastqs[2]{&fastq1, &fastq2};

  string line;
  string name;
  string read;
  char c;
  const string vt_pattern{"NNNWNNNWNNNWNNNTGACT"};
  while (fastq1 && fastq2) {
    vector<string> sequence(2);
    vector<string> vts(2);
    vector<string> names(2);
    vector<unsigned int> vt_mismatches(2);
    for (const bool read2 : {false, true}) {
      ifstream & fastq{*fastqs[read2]};
      fastq >> c;
      if (!fastq) continue;
      getline(fastq, line);
      istringstream line_str{line.c_str()};
      line_str >> name;
      fastq >> read >> c >> line;
      if (read.size() > vt_pattern.size()) {
        sequence[read2] = read.substr(vt_pattern.size());
        vts[read2] = read.substr(0, vt_pattern.size());
        names[read2] = name;
        for (unsigned int b{0}; b != vt_pattern.size(); ++b) {
          switch (vt_pattern[b]) {
            case 'N':
              if (read[b] == 'N') {
                ++vt_mismatches[read2];
              }
              break;
            case 'W':
              if (read[b] == 'C' || read[b] == 'G') {
                ++vt_mismatches[read2];
              }
              break;
            default:
              if (vt_pattern[b] != read[b]) {
                ++vt_mismatches[read2];
              }
              break;
          }
        }
      }
    }
    if (0) {
      sout << vt_mismatches[0] << vt_mismatches[1] << endl;
      continue;
    }
    for (const bool read2 : {false, true}) {
      const vector<SimpleHit> mums{sa.find_mams(sequence[read2])};
      for (const auto mum : mums) {
        sout << names[read2]
             << read2
             << vt_mismatches[read2] << vt_mismatches[1 - read2]
             << vts[0] << vts[1]
             << sequence[read2] << sequence[1 - read2]
             << ref.name(mum.chr) << mum.pos
             << mum.off << mum.len << mum.dir << endl;
      }
    }
  }

  return 0;
} catch (Error & e) {
  cerr << "paa::Error:" << endl;
  cerr << e.what() << endl;
  return 1;
} catch (exception & e) {
  cerr << "std::exception" << endl;
  cerr << e.what() << endl;
  return 1;
} catch (...) {
  cerr << "unknown exception was caught" << endl;
  return 1;
}
```
