## Supplementary material for "G-Graph: An interactive genomic graph viewer": MUMdex Source Code for G-Graph: fastqs2fastq.pl

```
#! /usr/bin/perl -w

use strict;

my $f1n = $ARGV[0];
my $f2n = $ARGV[1];

open(my $f1, "<", $f1n);
open(my $f2, "<", $f2n);

my $l;
while (1) {
    $l = <$f1>;
    if (!defined $l) {last};
    print $l;
    for (my $n = 0; $n < 3; ++$n) {
        $l = <$f1>;
        print $l;
    }
    for (my $n = 0; $n < 4; ++$n) {
        $l = <$f2>;
        print $l;
    }
}
```
