## Supplementary material for "G-Graph: An interactive genomic graph viewer": MUMdex Source Code for G-Graph: file_line.sh

```
#!/bin/bash

readonly INPUT_FILE=$1
readonly PROGNAME=$(basename "$0")
readonly LOCKFILE_DIR=/tmp
readonly LOCK_FD=200

lock() {
    local prefix=$1
    local fd=${2:-$LOCK_FD}
    local lock_file=$LOCKFILE_DIR/$prefix.lock

    # create lock file
    eval "exec $fd>$lock_file"

    # acquire the lock
    flock -n $fd \
        && return 0 \
        || return 1
}

main() {
    # keep trying to get lock
    while ! lock $PROGNAME ; do
        sleep 0.$((RANDOM * 3))
    done

    # initialize file line
    count_file=$LOCKFILE_DIR/count${INPUT_FILE//\//-}.txt
    if [ ! -e  $count_file ] ; then
        echo 1 > $count_file
    fi

    # get current file line
    count=$(cat $count_file)
    line=$(tail -n +$count $INPUT_FILE | head -n 1)

    # record new count
    count=$((count + 1))
    echo $count > $count_file

    # deliver line
    if [ "$line" = "" ] ; then
        echo done
    else
        echo "$line"
    fi
}

main

exit 0
```
