## Supplementary material for "G-Graph: An interactive genomic graph viewer": MUMdex Source Code for G-Graph: filter_bridges.cpp

```
//
// filter_bridges
//
// make bridges file smaller to aid specific processing
//
// Copyright 2018 Peter Andrews @ CSHL
//

#include <exception>
#include <iostream>
#include <fstream>
#include <string>

#include "bridges.h"
#include "error.h"
#include "files.h"
#include "mumdex.h"

using std::cerr;
using std::cout;
using std::endl;
using std::exception;
using std::ofstream;
using std::string;

using paa::BridgeInfo;
using paa::Error;
using paa::MappedVector;
using paa::Reference;

int main(int argc, char* argv[])  try {
  if (--argc != 2) throw Error("usage: filter_bridges bridges_in bridges_out");

  // Check for old/new version problem
  const string bridges_name{argv[1]};
  if ((bridges_name.find("chrbridges") != string::npos && NEW_BRIDGE_FORMAT) ||
      (bridges_name.find("newbridges") != string::npos && !NEW_BRIDGE_FORMAT))
    throw Error("new/old bridges file version problem") <<
        paa::bridges_bad_message();
  const MappedVector<BridgeInfo> bridges{bridges_name};

  const string bridges_out_name{argv[2]};
  ofstream bridges_out{bridges_out_name.c_str(), std::ofstream::binary};
  if (!bridges_out)
    throw Error("Problem opening output file") << bridges_out_name;

  for (const BridgeInfo & bridge : bridges) {
    if (bridge.invariant() == 0 || labs(bridge.invariant()) > 1000000 ||
        bridge.chr1() != bridge.chr2() || bridge.high1() == bridge.high2())
      continue;
    bridges_out.write(reinterpret_cast<const char *>(&bridge),
                      sizeof(BridgeInfo));
  }

  cerr << "All done with " << bridges_name << endl;

  return 0;
} catch (Error & e) {
  cerr << "paa::Error:" << endl;
  cerr << e.what() << endl;
  return 1;
} catch (exception & e) {
  cerr << "std::exception" << endl;
  cerr << e.what() << endl;
  return 1;
} catch (...) {
  cerr << "unknown exception was caught" << endl;
  return 1;
}
```
