## Supplementary material for "G-Graph: An interactive genomic graph viewer": MUMdex Source Code for G-Graph: filter_bridges.sh

```
#! /bin/bash

samples_dir="$1" &&
cd $samples_dir &&
indir=$PWD/bridges_new &&
outdir=$PWD/bridges_filtered &&
mkdir $outdir &&
for file in $indir/*.bin ; do
    ~/mumdex/filter_bridges $file $outdir/$(basename $file) || exit 1    
done &&
echo All Done with $samples_dir &&
touch $outdir/done

exit 0
```
