## Supplementary material for "G-Graph: An interactive genomic graph viewer": MUMdex Source Code for G-Graph: find_bridge.cpp

```
//
// find_bridge
//
// look for a specific bridge in one or more samples
//
// Copyright 2016 Peter Andrews @ CSHL
//

#include <exception>
#include <iomanip>
#include <iostream>
#include <map>
#include <string>

#include "bridges.h"
#include "error.h"
#include "fasta.h"
#include "mumdex.h"
#include "population.h"
#include "sequence.h"
#include "utility.h"

using std::cerr;
using std::cout;
using std::endl;
using std::exception;
using std::map;
using std::right;
using std::setw;
using std::string;

using paa::sout;
using paa::Base;
using paa::Bridge;
using paa::Bridges;
using paa::ChromosomeIndexLookup;
using paa::ConsensusSequence;
using paa::Error;
using paa::Family;
using paa::MUM;
using paa::MUMdex;
using paa::Pair;
using paa::Population;
using paa::Reference;
using paa::Sample;

int main(int argc, char* argv[])  try {
  // Check usage
  if (--argc < 10) {
    throw Error("usage: find_bridge samples_dir pop_file "
                "chr1 pos1 high1 chr2 pos2 high2 inv [sample|family] ...");
  }

  // Process command line arguments
  const string samples_dir{argv[1]};
  const Population pop{argv[2]};
  const string chr1s{argv[3]};
  const unsigned int pos1{static_cast<unsigned int>(atoi(argv[4]))};
  const bool high1{static_cast<bool>(atoi(argv[5]))};
  const string chr2s{argv[6]};
  const unsigned int pos2{static_cast<unsigned int>(atoi(argv[7]))};
  const bool high2{static_cast<bool>(atoi(argv[8]))};
  const int64_t invariant{atol(argv[9])};
  argc -= 9;
  argv += 9;

  // Load reference information
  const string first_sample_name{pop.sample(pop.samples(argv[1]).front())};
  const MUMdex first_mumdex{samples_dir + "/" + first_sample_name + "/mumdex"};
  const Reference & ref{first_mumdex.reference()};
  const ChromosomeIndexLookup chr_lookup{ref};
  const unsigned int chr1{chr_lookup[chr1s]};
  const unsigned int chr2{chr_lookup[chr2s]};

  // Bridge count summary
  map<Sample, unsigned int> sample_counts;

  // Consensus sequence adjacent to anchors
  const unsigned int adj_len{20};
  ConsensusSequence consensus[2]{adj_len, adj_len};

  while (argc--) {
    // Get samples for sample or family name
    const string sample_or_family{argv++[1]};
    for (const Sample sample : pop.samples(sample_or_family)) {
      // Get sample info
      const Family family{pop.family(sample)};
      const string sample_name{pop.sample(sample)};
      const string family_name{pop.family(family)};

      // Load the mumdex for the sample
      const MUMdex mumdex{samples_dir + "/" + sample_name + "/mumdex"};

      // Counts
      unsigned int nb{0};  // bridge count
      unsigned int nu{0};  // non-dupe bridge count
      unsigned int nl{0};  // non-dupe and long enough bridge count

      // Loop over bridges found
      for (const Bridge & bridge : Bridges{chr1, pos1, high1,
              chr2, pos2, high2, invariant, mumdex}) {
        // Bridge components
        const Pair pair{bridge.pair()};
        const MUM mum1{bridge.mum1()};
        const MUM mum2{bridge.mum2()};

        // Count non-dupes and long enough support anchors
        if (!pair.dupe()) {
          ++nu;
          if (mum1.length() >= 25 && mum2.length() >= 25) {
            ++nl;
          }
        }

        if (true) cout << mumdex.pair_view(bridge.pair_index()) << endl;

        // Output bridge information
        cout << endl
             << "-------------------------------------------------------------"
             << endl << endl;

        cout << sample_name << " "
             << setw(3) << right << ++nb << " "
             << setw(3) << right << nu << " "
             << setw(3) << right << nl << " "
             << pair.dupe() << " "
             << setw(3) << right << mum1.length() << " "
             << setw(3) << right << mum2.length();
        unsigned int i{0};
        for (const string & sequence : bridge.adjacent_sequences(adj_len)) {
          consensus[i++].add(sequence);
          cout << " " << sequence;
        }
        cout<< endl;

        // Update counts
        ++sample_counts[sample];
      }
      if (sample_counts.count(sample)) {
        cerr << endl;
      }
    }
  }

  // Output consensus sequences
  if (sample_counts.size()) {
    const unsigned int width{3};
    for (const auto & con : consensus) {
      const string & seq{con.sequence()};
      cout << seq << endl;
      cerr << endl;
      cout << "b";
      for (unsigned int b{0}; b != Base::n_bases; ++b) {
        cout << " " << setw(width) << right << Base{b}.value();
      }
      cout << endl;
      for (unsigned int i{0}; i != con.size(); ++i) {
        cout << seq[i];
        for (unsigned int b{0}; b != Base::n_bases; ++b) {
          cout << " " << setw(width) << right << con.count(i, b);
        }
        cout << endl;
      }
      cerr << endl;
    }
  }

  // Sample bridge count summary
  for (const auto & sample_count : sample_counts) {
    const Sample sample{sample_count.first};
    const Family family{pop.family(sample)};
    const unsigned int count{sample_count.second};
    sout << pop.sample(sample)
         << pop.family(family)
         << pop.member(sample)
         << count << endl;
  }

  // Output how many samples bridge was found in
  if (sample_counts.size()) cerr << endl;
  cout << sample_counts.size() << endl;

  return 0;
} catch (Error & e) {
  cerr << "paa::Error:" << endl;
  cerr << e.what() << endl;
  return 1;
} catch (exception & e) {
  cerr << "std::exception" << endl;
  cerr << e.what() << endl;
  return 1;
} catch (...) {
  cerr << "unknown exception was caught" << endl;
  return 1;
}
```
