## Supplementary material for "G-Graph: An interactive genomic graph viewer": MUMdex Source Code for G-Graph: find_indel.cpp

```
//
// find_indel
//
// Try to find a specific indel, with some slop allowed
//
// Copyright 2018 Peter Andrews @ CSHL
//

#include <algorithm>
#include <exception>
#include <iostream>
#include <sstream>
#include <string>

#include "bridges.h"
#include "error.h"
#include "files.h"
#include "mumdex.h"
#include "population.h"
#include "utility.h"

using std::cerr;
using std::cout;
using std::endl;
using std::exception;
using std::lower_bound;
using std::ostringstream;
using std::string;

using paa::readable;
using paa::sout;
using paa::BridgeInfo;
using paa::ChromosomeIndexLookup;
using paa::Error;
using paa::Family;
using paa::FileVector;
using paa::MUM;
using paa::Population;
using paa::Reference;
using paa::Sample;

int main(int argc, char* argv[])  try {
  // Check usage
  if (--argc < 10) {
    throw Error("usage: find_indel ref bridges_dir pop_file chr pos invariant "
                "pos_slop pos_range inv_slop sample|family ...");
  }

  // Process command line arguments
  const Reference ref{argv[1]};
  const ChromosomeIndexLookup chr_lookup{ref};
  const string bridges_dir{argv[2]};
  const Population pop{argv[3]};
  const string chr_name{argv[4]};
  const int pos{atoi(argv[5])};
  const int invariant{atoi(argv[6])};
  const int pos_slop{atoi(argv[7]) + abs(invariant)};
  const int pos_range{atoi(argv[8])};
  const int invariant_slop{atoi(argv[9])};

  argc -= 9;
  argv += 9;

  const unsigned int chr{chr_lookup[chr_name]};

  while (argc--) {
    // Get samples for sample or family name
    const string sample_or_family{argv++[1]};
    for (const Sample sample : pop.samples(sample_or_family)) {
      // Get sample info
      const Family family{pop.family(sample)};
      const string sample_name{pop.sample(sample)};
      const string family_name{pop.family(family)};

      // Open bridges file
      ostringstream bridges_name;
      bridges_name << bridges_dir << "/" << sample_name << "/"
                   << get_bridge_file_name(ref, chr);
      if (!readable(bridges_name.str()))
        throw Error("Could not open bridges file")
            << bridges_name.str() << paa::bridges_bad_message();
      const FileVector<BridgeInfo> bridges{bridges_name.str()};

      // Advance to start position
      const uint64_t pos_start{pos > pos_slop + pos_range ?
            (pos > pos_slop - pos_range ? pos - pos_slop - pos_range : 0) :
            0UL};
      FileVector<BridgeInfo>::const_iterator start{
        lower_bound(bridges.begin(), bridges.end(), pos_start,
                    [](const BridgeInfo & lhs, const unsigned int lpos) {
                      return lhs.pos1() < lpos;
                    })};

      // Check for matching bridge
      for (FileVector<BridgeInfo>::const_iterator test_iter{start};
           test_iter != bridges.end(); ++test_iter) {
        const BridgeInfo test{*test_iter};
        if (static_cast<int>(test.pos1()) > pos + pos_slop + pos_range) break;
        if (pos + pos_slop >= static_cast<int>(test.pos1()) &&
            pos <= static_cast<int>(test.pos2()) + pos_slop &&
            test.invariant() + invariant_slop >= invariant &&
            test.invariant() <= invariant + invariant_slop &&
            test.chr1() == test.chr2() &&
            test.high1() != test.high2()) {
          if (test.invariant() != 0) {
            sout << family_name << sample_name << pop.member(sample)
                 << ref.name(test.chr1()) << test.pos1() << test.high1()
                 << ref.name(test.chr2()) << test.pos2() << test.high2()
                 << test.invariant() << test.offset() << test.bridge_count()
                 << test.anchor1_length() << test.anchor2_length()
                 << test.mate_anchor1_length() << test.mate_anchor2_length()
                 << test.mate_anchor1_count() << test.mate_anchor1_count()
                 << endl;
          }
        }
      }
    }
  }

  return 0;
} catch (Error & e) {
  cerr << "paa::Error:" << endl;
  cerr << e.what() << endl;
  return 1;
} catch (exception & e) {
  cerr << "std::exception" << endl;
  cerr << e.what() << endl;
  return 1;
} catch (...) {
  cerr << "unknown exception was caught" << endl;
  return 1;
}
```
