## Supplementary material for "G-Graph: An interactive genomic graph viewer": MUMdex Source Code for G-Graph: find_microsatellite.cpp

```
//
// find_microsatellite
//
// find microsatellite at a position
//
// Copyright 2016 Peter Andrews @ CSHL
//

#include <exception>
#include <iostream>
#include <set>
#include <sstream>
#include <string>
#include <vector>

#include "error.h"
#include "mumdex.h"
#include "utility.h"

using std::cout;
using std::cerr;
using std::cin;
using std::endl;
using std::exception;
using std::ostringstream;
using std::set;
using std::string;
using std::vector;

using paa::ChromosomeIndexLookup;
using paa::Error;
using paa::Reference;
using paa::SpaceOut;

using paa::MUMdex;

class MicroSatellite {
 public:
  MicroSatellite(const unsigned int chr_arg,
                 const unsigned int start_arg,
                 const unsigned int stop_arg,
                 const string & motif_arg) :
      chromosome{chr_arg}, start_pos{start_arg}, stop_pos{stop_arg},
    motif{motif_arg} { }
  unsigned int chromosome;
  unsigned int start_pos;
  unsigned int stop_pos;
  string motif;
  unsigned int n_repeat() const { return n_bases() / motif.size(); }
  unsigned int n_bases() const { return stop_pos - start_pos; }
  // less means better!
  bool operator<(const MicroSatellite & rhs) const {
    if (n_bases() == rhs.n_bases()) {
      return motif.size() < rhs.motif.size();
    } else {
      return n_bases() > rhs.n_bases();
    }
  }
};

int main(int argc, char * argv[]) try {
  paa::exit_on_pipe_close();
  if (--argc != 1) throw Error("usage: (echo chr pos) ... | "
                               "find_microsatellite ref");

  const Reference ref{argv[1]};
  const ChromosomeIndexLookup chr_lookup{ref};

  const unsigned int max_motif_length{100};

  string chrs;
  unsigned int chr;
  unsigned int pos;
  unsigned int last_chr{0};
  unsigned int last_end{0};
  while (cin >> chrs >> pos) {
    set<MicroSatellite> repeats;
    chr = chr_lookup[chrs];
    if (chr == last_chr && pos < last_end) continue;
    if (pos >= ref.size(chr)) break;
    for (unsigned int motif_length{1}; motif_length != max_motif_length + 1;
         ++motif_length) {
      for (unsigned int offset{0}; offset != 2 * max_motif_length; ++offset) {
        const unsigned int start_pos{pos + offset > max_motif_length ?
              pos + offset - max_motif_length : 0};
        const unsigned int stop_pos{start_pos + motif_length};
        const string motif{ref.subseq(chr, start_pos, stop_pos)};
        unsigned int n_repeat{1};
        // go up in position
        unsigned int high_pos{stop_pos};
        for (unsigned int repeat{1}; ; ++repeat) {
          const unsigned int repeat_start{start_pos + repeat * motif_length};
          const unsigned int repeat_stop{repeat_start + motif_length};
          if (repeat_stop > ref.size(chr)) break;
          const string repeat_motif{ref.subseq(chr, repeat_start, repeat_stop)};
          if (repeat_motif != motif) break;
          ++n_repeat;
          high_pos = repeat_stop;
        }
        // go down in position
        unsigned int low_pos{start_pos};
        for (unsigned int repeat{1}; ; ++repeat) {
          if (start_pos < repeat * motif_length) break;
          const unsigned int repeat_start{start_pos - repeat * motif_length};
          const unsigned int repeat_stop{repeat_start + motif_length};
          const string repeat_motif{ref.subseq(chr, repeat_start, repeat_stop)};
          if (repeat_motif != motif) break;
          ++n_repeat;
          low_pos = repeat_start;
        }
        // was a repeat found?
        if (n_repeat > 1 && low_pos <= pos && high_pos > pos) {
          repeats.emplace(chr, low_pos, high_pos, motif);
        }
      }
    }
    vector<MicroSatellite> reported;
    for (const MicroSatellite & repeat : repeats) {
      bool skip{false};
      for (const MicroSatellite & before : reported) {
        if (repeat.chromosome == before.chromosome &&
            repeat.start_pos >= before.start_pos &&
            repeat.stop_pos <= before.stop_pos) {
          skip = true;
          break;
        }
      }
      if (skip) continue;
      if (repeat.n_bases() < 3) break;
      ostringstream sout;
      SpaceOut<ostringstream> ssout(sout);
      ssout  // << reported.size()
            << ref.name(repeat.chromosome)
            << repeat.start_pos << repeat.stop_pos
            << repeat.n_bases() << repeat.n_repeat()
            << repeat.motif.size() << repeat.motif;
      // << ref.subseq(repeat.chromosome, repeat.start_pos, repeat.stop_pos);
      last_chr = chr;
      last_end = repeat.stop_pos;
      cout << sout.str() << endl;
      break;
      reported.push_back(repeat);
    }
  }

  cerr << " Done" << endl;
  return 0;
} catch (Error & e) {
  cerr << "paa::Error:" << endl;
  cerr << e.what() << endl;
  return 1;
} catch (exception & e) {
  cerr << "std::exception" << endl;
  cerr << e.what() << endl;
  return 1;
} catch (...) {
  cerr << "unknown exception was caught" << endl;
  return 1;
}
```
