## Supplementary material for "G-Graph: An interactive genomic graph viewer": MUMdex Source Code for G-Graph: find_repeats.cpp

```
//
// find_repeats
//
// find repeat structure
//
// Copyright 2018 Peter Andrews @ CSHL
//

#include <exception>
#include <iostream>
#include <sstream>
#include <string>
#include <vector>

#include "error.h"
#include "mumdex.h"
#include "threads.h"

using std::cout;
using std::cerr;
using std::cin;
using std::endl;
using std::future;
using std::exception;
using std::ostringstream;
using std::string;
using std::vector;

using paa::ChromosomeIndexLookup;
using paa::Error;
using paa::Reference;
using paa::ThreadPool;

constexpr unsigned int max_motif_length{10000};

struct RepeatReturn {
  unsigned int chr;
  unsigned int pos;
  unsigned int bases;
  unsigned int start;
  string motif;
};
using RepeatReturns = vector<RepeatReturn>;

int main(int argc, char * argv[]) try {
  if (--argc != 5)
    throw Error("usage: find_repeats ref chr start stop n_threads");

  const Reference ref{argv[1]};
  const ChromosomeIndexLookup chr_lookup{ref};
  const string chr_name{argv[2]};
  const unsigned int chr{chr_lookup[chr_name]};

  auto block_fun = [&ref, chr]
      (const unsigned int start, const unsigned int stop) {
    RepeatReturns result;
    for (unsigned int pos{start}; pos != stop; ++pos) {
      unsigned int motif_length{max_motif_length};
      unsigned int repeat_bases{0};
      for (; motif_length >= 1; --motif_length) {
        repeat_bases = 0;
        for (unsigned int copy{1}; ; ++copy) {
          for (unsigned int base{0}; base != motif_length; ++base) {
            if (pos + copy * motif_length + base >= ref.size(chr)) break;
            if (ref[chr][pos + base] ==
                ref[chr][pos + base + copy * motif_length]) {
              ++repeat_bases;
            } else {
              break;
            }
          }
          if (repeat_bases < copy * motif_length) break;
        }
        if (repeat_bases >= motif_length) break;
      }
      if (motif_length && repeat_bases >= motif_length) {
        repeat_bases += motif_length;
        for (unsigned int small_motif_length{1};
             small_motif_length != motif_length / 2 + 1; ++small_motif_length) {
          if (motif_length % small_motif_length) continue;
          bool matches{true};
          for (unsigned int base{pos + small_motif_length};
               base != pos + motif_length; ++base) {
            if (ref[chr][pos + (base - pos) % small_motif_length] !=
                ref[chr][base]) {
              matches = false;
              break;
            }
          }
          if (matches) {
            motif_length = small_motif_length;
            break;
          }
        }

        string motif{ref.subseq(chr, pos, pos + motif_length)};
        string best_motif{motif};
        unsigned int motif_start{0};
        for (unsigned int roll{1}; roll != motif_length; ++roll) {
          motif =  motif.back() + motif.substr(0, motif.size() - 1);
          if (motif < best_motif) {
            best_motif = motif;
            motif_start = roll;
          }
        }

        result.push_back(
            RepeatReturn{chr, pos, repeat_bases, motif_start, best_motif});
      }
    }
    return result;
  };

  const unsigned int start{static_cast<unsigned int>(atoi(argv[3]))};
  const unsigned int stop_arg{static_cast<unsigned int>(atoi(argv[4]))};
  const unsigned int stop{(stop_arg == 0 || stop_arg > ref.size(chr)) ?
        ref.size(chr) : stop_arg};
  const unsigned int n_threads{static_cast<unsigned int>(atoi(argv[5]))};

  const unsigned int block_size{1000};
  ThreadPool pool{n_threads};
  vector<future<RepeatReturns>> futures;
  for (unsigned int pos{start}; pos < stop; pos += block_size)
    futures.push_back(pool.run(
        block_fun, pos, pos + block_size > stop ? stop : pos + block_size));
  unsigned int last_base{0};
  for (future<RepeatReturns> & future : futures) {
    const RepeatReturns repeats{future.get()};
    for (const RepeatReturn & repeat : repeats) {
      if (repeat.pos + repeat.bases <= last_base) continue;
      const unsigned int motif_length{static_cast<unsigned int>(
          repeat.motif.size())};
      cout << ref.name(repeat.chr)
           << " " << repeat.pos
           << " " << repeat.bases
           << " " << motif_length
           << " " << repeat.start
           << " " << repeat.bases / motif_length
           << " " << repeat.bases % motif_length
           << " " << repeat.motif
           << "\n";
      last_base = repeat.pos + repeat.bases;
    }
  }

  cerr << "done" << endl;
  return 0;
} catch (Error & e) {
  cerr << "paa::Error:" << endl;
  cerr << e.what() << endl;
  return 1;
} catch (exception & e) {
  cerr << "std::exception" << endl;
  cerr << e.what() << endl;
  return 1;
} catch (...) {
  cerr << "unknown exception was caught" << endl;
  return 1;
}
```
