## Supplementary material for "G-Graph: An interactive genomic graph viewer": MUMdex Source Code for G-Graph: gene_view.cpp

```
//
// gene_view
//
// looks at reads in a gene
//
// Copyright 2015 Peter Andrews @ CSHL
//

#include <array>
#include <algorithm>
#include <exception>
#include <iostream>
#include <fstream>
#include <set>
#include <sstream>
#include <string>

#include "error.h"
#include "genes.h"
#include "mumdex.h"
#include "utility.h"

using std::array;
using std::cout;
using std::cerr;
using std::endl;
using std::exception;
using std::min;
using std::ofstream;
using std::ostringstream;
using std::set;
using std::string;

using paa::reverse_complement;
using paa::ChromosomeIndexLookup;
using paa::Error;
using paa::MUMdex;
using paa::GeneXrefs;
using paa::KnownGenes;
using paa::Mappability;
using paa::MUM;
using paa::PosInfo;
using paa::Reference;
using paa::saved_ref_name;
using paa::sout;

int main(int argc, char * argv[]) try {
  paa::exit_on_pipe_close();

  --argc;
  if (argc != 6 && argc != 9)
    throw Error("usage: gene_view mumdex_name isoform sample kmer clip credit "
                "[genes isoforms xrefs]");

  const string mumdex_name{argv[1]};
  const string isoform{argv[2]};
  const string sample{argv[3]};
  const unsigned int kmer{static_cast<unsigned int>(atoi(argv[4]))};
  const unsigned int clip{static_cast<unsigned int>(atoi(argv[5]))};
  const unsigned int credit{static_cast<unsigned int>(atoi(argv[6]))};

  const string genes_name{argc == 9 ? argv[7] : "knownGene.txt"};
  const string isoforms_name{argc == 9 ? argv[8] : "knownIsoforms.txt"};
  const string kgXrefs_name{argc == 9 ? argv[9] : "kgXref.txt"};

  const MUMdex mumdex{mumdex_name};
  const string ref_name{saved_ref_name(mumdex_name)};
  const Reference & ref{mumdex.reference()};
  const Mappability map{mumdex_name};
  const ChromosomeIndexLookup chr_lookup{ref};
  const KnownGenes genes{genes_name, isoforms_name, chr_lookup, ref};
  const GeneXrefs xref{kgXrefs_name};

  // Look up isoform
  string gene_symbol;
  unsigned int isoform_index = genes.size();
  for (unsigned int i = 0; i != genes.size(); ++i) {
    const auto gene = genes[i];
    if (gene.name == isoform) {
      gene_symbol = xref[isoform].geneSymbol;
      isoform_index = i;
      break;
    }
  }
  if (isoform_index == genes.size())
    throw Error("Could not find isoform") << isoform;

  const auto gene = genes[isoform_index];

  const string search_name{"seq.txt"};
  ofstream search_file{search_name.c_str()};
  search_file << sample << " " << isoform_index << " "
              << gene_symbol << " " << mumdex_name << endl;

  set<uint64_t> seen_pairs;
  const unsigned int start{gene.t_start};
  const unsigned int stop{gene.t_stop};

  const auto lower = mumdex.lower_bound(PosInfo(gene.chr, start));
  const auto upper = mumdex.lower_bound(PosInfo(gene.chr, stop));
  unsigned int n_pairs = 0;
  for (auto iter = lower; iter != upper; ++iter) {
    const auto index(*iter);
    const auto pair_index(index.pair_index());
    if (seen_pairs.count(pair_index)) continue;
    seen_pairs.insert(pair_index);

    unsigned int max_lengths[2]{0, 0};
    MUM best_mums[2];

    for (auto mum_iter = mumdex.mums_begin(pair_index);
         mum_iter != mumdex.mums_end(pair_index); ++mum_iter) {
      const auto mum = *mum_iter;
      const auto read = mum.read_2();
      if (mum.chromosome() != gene.chr) continue;
      if (mum.length() > max_lengths[read] &&
          mum.position0() >= start &&
          mum.position0() < stop) {
        const auto offset = ref.offset(mum.chromosome());
        const uint64_t abspos = offset + mum.position0();
        const auto min_map = min(map.low(abspos),
                                 map.high(abspos + mum.length() - 1));
        if (mum.length() >= min_map + credit) {
          max_lengths[read] = mum.length();
          best_mums[read] = mum;
        }
      }
    }

    if (max_lengths[0] == 0 && max_lengths[1] == 0)
      continue;

    ++n_pairs;
    array<string, 2> sequences(mumdex.sequences(pair_index));
    const auto best_read = max_lengths[0] >= max_lengths[1] ? 0 : 1;
    if (best_mums[best_read].flipped()) {
      reverse_complement(&sequences[best_read]);
    } else {
      reverse_complement(&sequences[1 - best_read]);
    }
    for (const auto & sequence : sequences) {
      search_file << sequence << '\n';
    }
  }
  search_file.close();
  sout << "Output" << n_pairs << "pairs" << endl;
  ostringstream command;
  command << "debruijn " << ref_name << " " << search_name << " "
          << kmer << " " << clip << " "
          << genes_name << " " << isoforms_name << " " << kgXrefs_name;
  cout << command.str() << endl;
  if (system(command.str().c_str()) == -1) {
    cerr << "Problem creating debruijn output" << endl;
  }

  return 0;
} catch (Error & e) {
  cerr << "paa::Error:" << endl;
  cerr << e.what() << endl;
  return 1;
} catch (exception & e) {
  cerr << "std::exception" << endl;
  cerr << e.what() << endl;
  return 1;
} catch (...) {
  cerr << "unknown exception was caught" << endl;
  return 1;
}
```
