## Supplementary material for "G-Graph: An interactive genomic graph viewer": MUMdex Source Code for G-Graph: generate_genome_repeats_parallel.sh

```
#! /bin/bash

. ~/mumdex/utility/shell.sh

ref="$1"
[ "$ref" != "" ] || error no reference passed
[ -e $ref ] || error reference $ref does not exist

n_threads="${2:-20}"
[ "$n_threads" -gt 0 ] || error n_threads must be positive

outdir=$(basename $ref).jobs
mkdir -p $outdir
cd $outdir

[ -e $ref.fai ] || samtools faidx $ref || error could not generate fai for $ref

n=0
while read chr len rest ; do
    n=$((n + 1))
    jname=r.$n.$chr.$(basename $ref | perl -pe 's/(....).*/$1/')
    if [ ! -e $jname.out.txt ] ; then
        qsub -cwd -N $jname -l vf=4G -l h_vmem=4G -pe threads $n_threads \
            -o $jname.out.txt -e $jname.err.txt \
            ~/mumdex/unstable/chr_genome_repeats.sh $ref $chr $len $n_threads
    fi
done < $ref.fai
```
