## Supplementary material for "G-Graph: An interactive genomic graph viewer": MUMdex Source Code for G-Graph: get_population_data_text.sh

```
#! /bin/bash

# for chr in {1..22} X Y ; do start=20000000 ; stop=$((start+1000000)) ; ~/mumdex/unstable/get_population_data_text.sh ~/bridges ~/human_g1k_v37.fasta $chr $start $stop ; done

bridges_dir=$1
ref=$2
chr=$3
start=$4
stop=$5

index=$(perl -ne '$n++ ; print $n - 1 if /^'$chr'\s/' $ref.fai)

echo Info is $ref $bridges_dir $chr $start $stop $index

window=$chr-$start-$stop
mkdir -p $window
cd $window

rsync -a --exclude .git ~/mumdex/ mumdex
(cd mumdex && make -j 4 bridges2txt)

n=0
for sample in $(cd $bridges_dir/ ; ls -d $(echo SSC*/) | perl -pe 's+/++g') ; do
    bridge_dir=$bridges_dir/$sample
    bridge_file=$bridge_dir/chrbridges.$index.bin
    out=$sample.txt.gz
    n=$((n+1))
    if [ ! -e $out ] ; then
        echo $window $n 1>&2
        echo "./mumdex/bridges2txt $ref $bridge_file $chr $start $stop | gzip -c > $out && echo finished $sample $n"
    fi
done | qsub -l h_vmem=20G -cwd -N test -o out.txt -e err.txt 
```
