## Supplementary material for "G-Graph: An interactive genomic graph viewer": MUMdex Source Code for G-Graph: hists.cpp

```
//
// hists
//
// make histograms for input columns
//
// Copyright 2018 Peter Andrews @ CSHL
//

#include <algorithm>
#include <exception>
#include <iostream>
#include <sstream>
#include <string>
#include <vector>

#include "error.h"
#include "psplot.h"

using std::cerr;
using std::cin;
using std::cout;
using std::endl;
using std::exception;
using std::istringstream;
using std::string;
using std::vector;

using paa::Bounds;
using paa::Error;
using paa::PSDoc;
using paa::PSHSeries;
using paa::PSPage;

int main(int argc, char* argv[])  try {
  --argc;
  if (argc != 2) throw Error("data_file | hists n_bins out_name");

  const unsigned int n_bins{static_cast<unsigned int>(atoi(argv[1]))};
  const string out_name{argv[2]};

  string line;
  getline(cin, line);
  istringstream header_stream{line.c_str()};
  vector<string> header;
  while (header_stream >> line) header.push_back(line);
  vector<vector<double>> data(header.size());

  while (getline(cin, line)) {
    istringstream line_stream{line.c_str()};
    for (uint64_t c{0}; c != data.size(); ++c) {
      vector<double> & vals{data[c]};
      double input;
      line_stream >> input;
      if (!line_stream) throw Error("Bad input from line") << line;
      vals.push_back(input);
    }
    if (!line_stream) throw Error("Parse Error");
    line_stream.get();
    if (line_stream) throw Error("unexpected character");
  }

  PSDoc plots{out_name};
  plots.pdf(true);

  using Hist = PSHSeries<double, uint64_t>;
  for (uint64_t c{0}; c != data.size(); ++c) {
    vector<double> & values{data[c]};
    auto minmax = minmax_element(values.begin(), values.end());
    const double min{*minmax.first};
    const double max{*minmax.second};
    PSPage * page{PSPage::create(plots)};
    // const unsigned int n_bins{)};
    const unsigned int used_n_bins{n_bins ? n_bins :
          static_cast<unsigned int>(sqrt(values.size()))};
    const double edge{(max - min) / 20};
    Hist * hist{Hist::create(*page, header[c] + ";" + header[c] + ";N",
                             Bounds{min - edge, max + edge}, used_n_bins)};
    for (unsigned int v{0}; v != values.size(); ++v)
        hist->add_point(values[v]);
  }

  return 0;
} catch (Error & e) {
  cerr << "paa::Error:" << endl;
  cerr << e.what() << endl;
  return 1;
} catch (exception & e) {
  cerr << "std::exception" << endl;
  cerr << e.what() << endl;
  return 1;
} catch (...) {
  cerr << "unknown exception was caught" << endl;
  return 1;
}
```
