## Supplementary material for "G-Graph: An interactive genomic graph viewer": MUMdex Source Code for G-Graph: hmm.h

```
//
// hmm.h
//
// simple hmm hidden markov model
//
// Copyright 2018 Peter Andrews @ CSHL
//

#ifndef PAA_HMM_H_
#define PAA_HMM_H_

#include <algorithm>
#include <cmath>
#include <iostream>
#include <functional>
#include <limits>
#include <memory>
#include <random>
#include <vector>

#include "psplot.h"

namespace paa {

template <class FLOAT>
bool within_tolerance(const FLOAT lhs, const FLOAT rhs,
                      const FLOAT tolerance) {
  const FLOAT maxv{fabs(std::max(lhs, rhs))};
  if (maxv <= std::numeric_limits<FLOAT>::min()) return true;
  return fabs(lhs - rhs) < tolerance * maxv;
}

template <class FLOAT>
FLOAT lnproduct(const FLOAT lhs, const FLOAT rhs) {
  if (lhs > std::numeric_limits<FLOAT>::lowest() &&
      lhs < std::numeric_limits<FLOAT>::max() &&
      rhs > std::numeric_limits<FLOAT>::lowest() &&
      rhs < std::numeric_limits<FLOAT>::max()) {
    return lhs + rhs;
  } else {
    return std::numeric_limits<FLOAT>::lowest();
  }
}

template <class FLOAT>
FLOAT lnproduct(const FLOAT a1, const FLOAT a2, const FLOAT a3) {
  return lnproduct(a1, lnproduct(a2, a3));
}

template <class FLOAT>
FLOAT lnsum(const FLOAT lhs, const FLOAT rhs) {
  if (lhs <= std::numeric_limits<FLOAT>::lowest() ||
      lhs >= std::numeric_limits<FLOAT>::max())
    return rhs;
  if (rhs <= std::numeric_limits<FLOAT>::lowest() ||
      rhs >= std::numeric_limits<FLOAT>::max())
    return lhs;
  if (lhs > rhs)
    return lhs + log(1 + exp(rhs - lhs));
  return rhs + log(1 + exp(lhs - rhs));
}

template <class FLOAT>
class HMMparams_t {
 public:
  static constexpr FLOAT SMALL{std::numeric_limits<FLOAT>::lowest()};
  using Vector = std::vector<FLOAT>;
  using Matrix = std::vector<Vector>;

  HMMparams_t(const Matrix & emissions__,
              const Matrix & transitions__,
              const Vector initial_probs__ = Vector()) :
      emissions_{emissions__},
    transitions_{transitions__},
    initial_probs_{initial_probs__} {
      // Quick and not complete reality check on vector and matrix sizes
      if (n_states() <= 1) throw Error("Too few states in HMM");
      if (emissions_.size() != n_states())
        throw Error("Emissions / Transitions matrix size problem in HMM");
      if (initial_probs_.empty())
        initial_probs_.assign(n_states(), 1.0 / n_states());
      if (initial_probs_.size() != n_states())
        throw Error("Initial probs size is not n_states()");
    }

  // Random starting values
  HMMparams_t(const unsigned int n_states__,
              const unsigned int n_symbols__) :
      HMMparams_t{Matrix(n_states__, Vector(n_symbols__, 1.0 / n_symbols__)),
        Matrix(n_states__, Vector(n_states__, 1.0 / n_states__))} {
    std::random_device rd{};
    std::mt19937_64 mersenne{rd()};
    std::function<double()> gen{std::bind(
        std::uniform_real_distribution<double>(0.1, 1), mersenne)};
    FLOAT den{0};
    for (unsigned int i{0}; i != n_states(); ++i) {
      const FLOAT val{gen()};
      initial_probs_[i] = val;
      den += val;
    }
    for (unsigned int i{0}; i != n_states(); ++i) {
      initial_probs_[i] /= den;
    }
    for (unsigned int i{0}; i != n_states(); ++i) {
      den = 0;
      for (unsigned int j{0}; j != n_states(); ++j) {
        const FLOAT val{gen()};
        transitions_[i][j] = val;
        den += val;
      }
      for (unsigned int j{0}; j != n_states(); ++j) {
        transitions_[i][j] /= den;
      }
      den = 0;
      for (unsigned int k{0}; k != n_states(); ++k) {
        const FLOAT val{gen()};
        emissions_[i][k] = val;
        den += val;
      }
      for (unsigned int k{0}; k != n_states(); ++k) {
        emissions_[i][k] /= den;
      }
    }
  }

  bool within_tolerance(const HMMparams_t & other,
                        const FLOAT tolerance) const {
    for (unsigned int i{0}; i != n_states(); ++i) {
      if (!paa::within_tolerance(initial_probs_[i], other.initial_probs_[i],
                                 tolerance))
        return false;
      for (unsigned int j{0}; j != n_states(); ++j) {
        if (!paa::within_tolerance(transitions_[i][j], other.transitions_[i][j],
                                   tolerance))
          return false;
      }
      for (unsigned int k{0}; k != n_symbols(); ++k) {
        if (!paa::within_tolerance(emissions_[i][k], other.emissions_[i][k],
                                   tolerance))
          return false;
      }
    }
    return true;
  }

  std::ostream & out(std::ostream & stream) const {
    stream << "initial\n";
    for (unsigned int i{0}; i != n_states(); ++i) {
      stream << ' ' << initial_probs_[i];
    }
    stream << "\n";
    stream << "transitions\n";
    for (unsigned int i{0}; i != n_states(); ++i) {
      for (unsigned int j{0}; j != n_states(); ++j) {
        stream << ' ' << transitions_[i][j];
      }
      stream << "\n";
    }
    stream << "emissions\n";
    for (unsigned int i{0}; i != n_states(); ++i) {
      for (unsigned int k{0}; k != n_symbols(); ++k) {
        stream << ' ' << emissions_[i][k];
      }
      stream << "\n";
    }
    return stream;
  }

  unsigned int n_states() const {
    return static_cast<unsigned int>(transitions_.size());
  }
  unsigned int n_symbols() const {
    return static_cast<unsigned int>(emissions_.front().size());
  }
  const Matrix & emissions() const { return emissions_; }
  const Matrix & transitions() const { return transitions_; }
  const Vector & initial_probs() const { return initial_probs_; }
  Matrix & emissions() { return emissions_; }
  Matrix & transitions() { return transitions_; }
  Vector & initial_probs() { return initial_probs_; }

 private:
  Matrix emissions_;
  Matrix transitions_;
  Vector initial_probs_;
};
using HMMparams = HMMparams_t<double>;

template <class FLOAT>
std::ostream & operator<<(std::ostream & stream,
                          const HMMparams_t<FLOAT> & params) {
  return params.out(stream);
}

template <class FLOAT, class OBS>
class HMM_t {
 public:
  using Params = HMMparams_t<FLOAT>;
  using Vector = typename Params::Vector;
  using Matrix = typename Params::Matrix;
  using Matrix3 = std::vector<Matrix>;
  HMM_t(const Params & params__,
        const std::vector<OBS> & observations__) :
      params_{params__},
    observations_{observations__} {
      if (n_observations() <= 1) throw Error("Too few observations in HMM");
    }

  unsigned int n_states() const { return params_.n_states(); }
  unsigned int n_symbols() const { return params_.n_symbols(); }
  unsigned int n_observations() const {
    return static_cast<unsigned int>(observations_.size());
  }

  void forward(Matrix & alpha) const {
    for (unsigned int i{0}; i != n_states(); ++i) {
      alpha[i][0] = lnproduct(log(params_.initial_probs()[i]),
                              log(params_.emissions()[i][observations_[0]]));
    }
    for (unsigned int t{1}; t != n_observations(); ++t) {
      for (unsigned int j{0}; j != n_states(); ++j) {
        FLOAT logalpha{HMMparams::SMALL};
        for (unsigned int i{0}; i != n_states(); ++i) {
          logalpha = lnsum(logalpha, lnproduct(
              alpha[i][t - 1],
              log(params_.transitions()[i][j])));
        }
        alpha[j][t] = lnproduct(logalpha,
                                log(params_.emissions()[j][observations_[t]]));
      }
    }
  }
  void backward(Matrix & beta) const {
    for (unsigned int i{0}; i != n_states(); ++i) {
      beta[i][n_observations() - 1] = 0;
    }
    for (unsigned int tt{1}; tt != n_observations(); ++tt) {
      const unsigned int t{n_observations() - tt - 1};
      for (unsigned int i{0}; i != n_states(); ++i) {
        FLOAT logbeta{HMMparams::SMALL};
        for (unsigned int j{0}; j != n_states(); ++j) {
          logbeta = lnsum(logbeta, lnproduct(
              log(params_.transitions()[i][j]), lnproduct(
                  params_.emissions()[j][observations_[t + 1]],
                  beta[j][t + 1])));
        }
        beta[i][t] = logbeta;
      }
    }
  }
  HMMparams baum_welch_iteration(Matrix & alpha, Matrix & beta,
                                 Matrix & gamma, Matrix3 & xi) const {
    forward(alpha);
    backward(beta);

    // Calculate gamma
    for (unsigned int t{0}; t != n_observations(); ++t) {
      FLOAT norm{HMMparams::SMALL};
      for (unsigned int i{0}; i != n_states(); ++i) {
        gamma[i][t] = lnproduct(alpha[i][t], beta[i][t]);
        norm = lnsum(norm, gamma[i][t]);
      }
      for (unsigned int i{0}; i != n_states(); ++i) {
        gamma[i][t] = lnproduct(gamma[i][t], -norm);
      }
    }

    // Calculate xi
    for (unsigned int t{0}; t + 1 != n_observations(); ++t) {
      FLOAT norm{HMMparams::SMALL};
      for (unsigned int i{0}; i != n_states(); ++i) {
        for (unsigned int j{0}; j != n_states(); ++j) {
          xi[i][j][t] = lnproduct(alpha[i][t], lnproduct(
              log(params_.transitions()[i][j]), lnproduct(
                  log(params_.emissions()[j][observations_[t + 1]]),
                  beta[j][t + 1])));
          norm = lnsum(norm, xi[i][j][t]);
        }
      }
      for (unsigned int i{0}; i != n_states(); ++i) {
        for (unsigned int j{0}; j != n_states(); ++j) {
          xi[i][j][t] = lnproduct(xi[i][j][t], -norm);
        }
      }
    }

    HMMparams new_params{params_};
    Vector & initial_probs{new_params.initial_probs()};
    Matrix & emissions{new_params.emissions()};
    Matrix & transitions{new_params.transitions()};

    // Calculate new parameters
    static thread_local Matrix num1(n_states(), Vector(n_states()));
    static thread_local Matrix den1(n_states(), Vector(n_states()));
    static thread_local Matrix num2(n_states(), Vector(n_symbols()));
    static thread_local Matrix den2(n_states(), Vector(n_symbols()));
    FLOAT isum{0};
    for (unsigned int i{0}; i != n_states(); ++i) {
      initial_probs[i] = exp(gamma[i][0]);
      isum += initial_probs[i];
      for (unsigned int j{0}; j != n_states(); ++j) {
        num1[i][j] = HMMparams::SMALL;
        den1[i][j] = HMMparams::SMALL;
      }
      for (unsigned int k{0}; k != n_symbols(); ++k) {
        num2[i][k] = HMMparams::SMALL;
        den2[i][k] = HMMparams::SMALL;
      }
    }
    for (unsigned int i{0}; i != n_states(); ++i) {
      initial_probs[i] /= isum;
    }
    for (unsigned int t{0}; t != n_observations(); ++t) {
      for (unsigned int i{0}; i != n_states(); ++i) {
        if (t + 1 != n_observations()) {
          for (unsigned int j{0}; j != n_states(); ++j) {
            num1[i][j] = lnsum(num1[i][j], xi[i][j][t]);
            den1[i][j] = lnsum(den1[i][j], gamma[i][t]);
          }
        }
        for (unsigned int k{0}; k != n_symbols(); ++k) {
          if (observations_[t] == k) {
            num2[i][k] = lnsum(num2[i][k], gamma[i][t]);
          }
          den2[i][k] = lnsum(den2[i][k], gamma[i][t]);
        }
      }
    }
    for (unsigned int i{0}; i != n_states(); ++i) {
      FLOAT tsum{0};
      for (unsigned int j{0}; j != n_states(); ++j) {
        transitions[i][j] = exp(lnproduct(num1[i][j], -den1[i][j]));
        tsum += transitions[i][j];
      }
      for (unsigned int j{0}; j != n_states(); ++j) {
        transitions[i][j] /= tsum;
      }
      FLOAT esum{0};
      for (unsigned int k{0}; k != n_symbols(); ++k) {
        emissions[i][k] = exp(lnproduct(num2[i][k], -den2[i][k]));
        esum += emissions[i][k];
      }
      for (unsigned int k{0}; k != n_symbols(); ++k) {
        emissions[i][k] /= esum;
      }
    }

    return new_params;
  }
  unsigned int baum_welch(const unsigned int max_iter = 1000,
                          const FLOAT tolerance = 0.00001) {
    series.push_back(std::make_unique<PSXYSeries>(
        doc, "Observation probability; Iteration; per obs prob", marker));
    // series.back()->parents().front()->log_y(true);
    series.back()->graph().log_y(true);

    static thread_local Matrix alpha(n_states(), Vector(n_observations()));
    static thread_local Matrix beta(n_states(), Vector(n_observations()));
    static thread_local Matrix gamma(n_states(), Vector(n_observations()));
    static thread_local Matrix3 xi(n_states(), gamma);
    std::cout << params_ << std::endl;
    unsigned int i{0};
    for (; i != max_iter; ++i) {
      std::cerr << "Iteration " << i << std::endl;
      const HMMparams new_params{baum_welch_iteration(alpha, beta, gamma, xi)};
      const bool complete{new_params.within_tolerance(params_, tolerance)};
      params_ = new_params;
      std::cout << params_ << std::endl;
      FLOAT obs_prob{HMMparams::SMALL};
      for (unsigned int j{0}; j != n_states(); ++j) {
        obs_prob = lnsum(obs_prob, alpha[j].back());
      }
      series.back()->add_point(i, exp(obs_prob / n_observations()));
      if (complete) break;
    }

    // Make plots of state probabilities
    series.push_back(std::make_unique<PSXYSeries>(
        doc, "State probabilities;Observation; State 0 Probability" , marker));
    for (unsigned int o{0}; o != n_observations(); ++o) {
      series.back()->add_point(o, exp(gamma[0][o]));
    }

    return i;
  }

 private:
  Params params_;
  const std::vector<OBS> & observations_;
  PSDoc doc{"hmm", "hmm"};
  std::vector<std::unique_ptr<PSXYSeries>> series{};
  Marker marker{paa::circle(), 0.3, "0 0 0", 0.2, true};
};

using HMM = HMM_t<double, unsigned int>;

}  // namespace paa

#endif  // PAA_HMM_H_
```
