## Supplementary material for "G-Graph: An interactive genomic graph viewer": MUMdex Source Code for G-Graph: incidental.cpp

```
//
// incidental
//
// use machine learning to identify incidental MUMs
// not finished or well thought out....
//
// Copyright 2018 Peter Andrews @ CSHL
//

#include <exception>
#include <iostream>
#include <string>
#include <vector>

#include "bridges.h"
#include "error.h"
#include "files.h"
#include "mumdex.h"
#include "psplot.h"

using std::cerr;
using std::cout;
using std::endl;
using std::exception;
using std::string;
using std::vector;

using paa::get_bridge_file_name;
using paa::BridgeInfo;
using paa::Error;
using paa::Mappability;
using paa::MappedVector;
using paa::PSDoc;
using paa::PSGraph;
using paa::PSPage;
using paa::PSXYSeries;
using paa::Reference;

int main(int argc, char* argv[])  try {
  // Check usage
  if (--argc < 3)
    throw Error("usage: incidental ref cutoff bridge_files ...");

  // Process command line arguments
  const Reference & ref{argv[1]};
  const Mappability map{ref};
  const int invariant_cutoff{atoi(argv[2])};
  argc -= 2;
  argv += 2;

  // Just a test for using right format
  cerr << get_bridge_file_name(ref, 0) << endl;

  const unsigned int max_length{200};
  vector<vector<uint64_t>> counts(2, vector<uint64_t>(max_length));

  while (argc--) {
    const string bridges_name{argv++[1]};
    const MappedVector<BridgeInfo> bridges{bridges_name};
    cerr << "Loaded " << bridges.size() << " bridges from " << bridges_name
         << endl;
    for (uint64_t b0{0}; b0 + 1 != bridges.size();) {
      const BridgeInfo bridge0{bridges[b0]};
      uint64_t real_bridge{bridges.size()};
      uint64_t b{b0};
      for (; b != bridges.size(); ++b) {
        const BridgeInfo bridge{bridges[b]};
        if (bridge.chr1() == bridge0.chr1() &&
            bridge.pos1() == bridge0.pos1() &&
            bridge.high1() == bridge0.high1()) {
          if (abs(bridge.invariant()) < invariant_cutoff &&
              bridge.high1() != bridge.high2() &&
              (real_bridge == bridges.size() ||
               (abs(bridge.invariant()) <
                abs(bridges[real_bridge].invariant()) ||
                (abs(bridge.invariant()) ==
                 abs(bridges[real_bridge].invariant()) &&
                     bridge.offset() < bridges[real_bridge].offset())))) {
            real_bridge = b;
          }
        } else {
          if (real_bridge != bridges.size()) {
            for (uint64_t b2{b0}; b2 != b; ++b2) {
              const BridgeInfo bridge2{bridges[b2]};
              const bool real{b2 == real_bridge};
              const bool incidental{bridge2.offset() <
                    bridges[real_bridge].offset() &&
                    bridge2.chr2() != bridge2.chr1() &&
                    bridge2.anchor1_length() >=
                    bridges[real_bridge].anchor1_length()};
              if (real || incidental) {
                const unsigned int abspos{ref.abspos(bridge2.chr2(),
                                                     bridge2.pos2())};
                const unsigned int mappability{
                  map.low_high(bridge2.high2(), abspos)};
                const unsigned int excess{
                  bridge2.anchor2_length() - mappability};
                if (bridge2.anchor2_length() < mappability)
                  throw Error("Bad length!");
                ++counts[real][excess];
              }
              if (false)
                cout << b0 << " " << b2 << " " << b
                     << " " << bridge2.chr1()
                     << " " << bridge2.pos1()
                     << " " << bridge2.high1()
                     << " " << bridge2.chr2()
                     << " " << bridge2.pos2()
                     << " " << bridge2.high2()
                     << " " << bridge2.offset()
                     << " " << bridge2.invariant()
                     << " " << real
                     << " " << incidental
                     << endl;
            }
          }
          if (false) cout << "next" << endl;
          b0 = b;
          break;
        }
      }
      if (b == bridges.size()) break;
    }
  }

  PSDoc plots{"incidental"};
  PSGraph excess_graph{plots, "Incidental MUM training set;Excess Mappability;"
        "Probability MUM is not incidental"};
  excess_graph.log_y(true);
  excess_graph.range().yl(0.1);
  PSXYSeries excess_series{excess_graph};

  PSPage good_bad{plots, "Incidental MUM training set", "1 2"};
  PSGraph good_graph{good_bad, "Good MUMs;Excess Mappability;"
        "Number of MUMs"};
  good_graph.log_y(true);
  PSXYSeries good_series{good_graph};
  PSGraph bad_graph{good_bad, "Bad MUMs;Excess Mappability;"
        "Number of MUMs"};
  bad_graph.log_y(true);
  PSXYSeries bad_series{bad_graph};

  for (unsigned int l{0}; l != max_length; ++l) {
    const uint64_t total{counts[0][l] + counts[1][l]};
    if (total) {
      const double prob_good{1.0 * counts[1][l] / total};
      cout << l << " " << counts[0][l] << " " << counts[1][l]
           << " " << prob_good << endl;
      excess_series.add_point(l, prob_good);
      good_series.add_point(l, counts[1][l]);
      bad_series.add_point(l, counts[0][l]);
    }
  }

  cerr << "done" << endl;

  return 0;
} catch (Error & e) {
  cerr << "paa::Error:" << endl;
  cerr << e.what() << endl;
  return 1;
} catch (exception & e) {
  cerr << "std::exception" << endl;
  cerr << e.what() << endl;
  return 1;
} catch (...) {
  cerr << "unknown exception was caught" << endl;
  return 1;
}
```
