## Supplementary material for "G-Graph: An interactive genomic graph viewer": MUMdex Source Code for G-Graph: jackpot.cpp

```
//
// jackpot.cpp
//
// see if there was jackpotting in candidates
//
// Copyright 2016 Peter Andrews @ CSHL
//

#include <algorithm>
#include <exception>
#include <iostream>
#include <random>
#include <string>
#include <sstream>
#include <vector>

#include "error.h"
#include "psplot.h"

using std::cin;
using std::cout;
using std::cerr;
using std::endl;
using std::exception;
using std::istringstream;
using std::max;
using std::vector;

using paa::Bounds;
using paa::Error;
using paa::Marker;
using paa::PSDoc;
using paa::PSGraph;
using paa::PSXYSeries;

int main(int argc, char* argv[]) try {
  if (--argc != 1) throw Error("usage: jackpot input_file");
  cout << argv[1] << endl;
#if 0
  const string type{[&input_name]() {
      istringstream input{input_name.c_str()};
      string result;
      getline(input, result, '_');
      replace(result, '.', ' ');
      return result;
    }()};

  vector<string> column_names;
  vector<string> family_names;
  vector<string> member_types;
  vector<unsigned int> in_40;
  vector<vector<unsigned int>> counts;
  vector<vector<unsigned int>> shared_counts;
  ifstream input_file{input_name.c_str()};
  string dummy;
  input_file >> dummy >> dummy >> dummy;
  for (unsigned int i{0}; i != 5; ++i) {
    getline(input_file, dummy, "_");
    column_names.push_back(dummy);
    input_file >> dummy >> dummy;
  }
  string family;
  string member;
  unsigned int in40;
  unsigned int count;
  unsigned int shared;
  while (input_file >> family >> member >> in40) {
    counts.resize(counts.size() + 1);
    for (unsigned int i{0}; i != 5; ++i) {
      input_file >> count >> shared;
      counts.back().push_back(count);
      shared_counts.back().push_back(shared);
    }
  }

  vector<unique_ptr<PSGraph>> graphs;
  vector<unique_ptr<PSXYSeries>> series;

  for (unsigned int t{0}; t != column_names.size(); ++t) {
    for (bool in : {false, true}) {
    }
  }


  const unsigned int n_trials{10000};

  auto mersenne = std::mt19937_64();
  mersenne.seed(time(nullptr));
  std::poisson_distribution<unsigned int> dist_{1.0 * n_events / n_people};

  Marker big_marker{paa::circle(), 1, "0 0 0", 1, true};
  Marker small_marker{paa::circle(), 1, "1 0 0", 1, false};

  unsigned int n_seen;
  unsigned int n{0};
  vector<unsigned int> counts;
  while (cin >> n_seen) {
    ++n;
    cin.ignore(10000, '\n');
    // cout << n << " " << n_seen << endl;
    if (counts.size() <= n_seen) {
      counts.resize(n_seen + 1);
    }
    ++counts[n_seen];
  }
  counts[0] = n_people - n;

  vector<unsigned int> dist(counts.size());
  for (unsigned int t{0}; t != n_trials; ++t) {
    for (unsigned int p{0}; p != n_people; ++p) {
      const unsigned int result{dist_(mersenne)};
      if (result >= dist.size()) {
        dist.resize(result + 1);
      }
      ++dist[result];
    }
  }

  PSDoc ps{"jackpot"};
  PSGraph jackpot{ps, ";Count;N", Bounds(
      -1.0, counts.size() + 1,
      -10, max(*max_element(counts.begin(), counts.end()),
               *max_element(dist.begin(), dist.end()) / n_trials) + 20)};
  PSGraph jackpot_zoom{ps, ";Count;N",
        Bounds(-1.0, counts.size() + 1, -2, 10)};
  PSXYSeries data{jackpot, big_marker};
  data.add(&jackpot_zoom);
  PSXYSeries dist_data{jackpot, small_marker};
  dist_data.add(&jackpot_zoom);

  for (unsigned int i{0}; i != dist.size(); ++i) {
    dist_data.add_point(i, 1.0 * dist[i] / n_trials);
  }

  for (unsigned int i{0}; i != counts.size(); ++i) {
    data.add_point(i, counts[i]);
  }
#endif
#if 0
  vector<unsigned int> people(n_people);
  for (unsigned int e{0}; e != n_events; ++e) {
  }
#endif
  return 0;
} catch (Error & e) {
  cerr << "paa::Error:" << endl;
  cerr << e.what() << endl;
  return 1;
} catch (exception & e) {
  cerr << "std::exception" << endl;
  cerr << e.what() << endl;
  return 1;
} catch (...) {
  cerr << "unknown exception was caught" << endl;
  return 1;
}
```
