## Supplementary material for "G-Graph: An interactive genomic graph viewer": MUMdex Source Code for G-Graph: job_status.sh

```
#! /bin/bash

(qstat -j '*SSC*map' ; qstat -j '*SSC*calc' )  > qstat.txt


cat qstat.txt | grep hard_queue | sort | uniq -c | sort -k1n > temp4.txt
df -k /mnt/wigclust*/data | sort -k5n | sed 's/\/dev\/sda1/wigclust19:/' > temp5.txt
qstat | grep ' r ' | awk '{print $8}' | sed 's/all.q@//' | sort | uniq -c | sort -k1n > temp3.txt
grep Creating ../run/finished/*.calc/out.txt | awk '{print $4}' | sort | uniq -c | sort -k1n > temp6.txt
grep Creating ../run/failed/*.calc/out.txt | awk '{print $4}' | sort | uniq -c | sort -k1n > temp7.txt

printf "node name\trunning\tqueued\tfull\tdone\tfailed\n"
for node in wigclust{1..24} ; do res=$(grep $node.cshl.edu temp3.txt | awk '{print $2,$1}') ; if [ "$res" = "" ] ; then echo -n $node.cshl.edu 0 ; else echo -n $res ; fi ; echo " "$(grep $node.cshl.edu temp4.txt| awk '{print $1}') $(grep $node: temp5.txt | awk '{print $5}') $(grep $node$ temp6.txt | awk '{print $1}') $(grep $node$ temp7.txt | awk '{print $1}') ; done | sed 's/.cshl.edu// ; s/ /\t/g' ; rm temp{3,4,5,6,7}.txt
echo

echo $(du -ks SSC*/rmdup.bam.bin/*{matches,positions}* | sort -k1n | awk '{tot+=$1;printf "%.1fG average space per sample %dG total space used\n", int(2*tot/NR/100000)/10, tot/1000000}'| tail -n 1 ) $(qacct -j 'SSC*' -d 3 | grep wall | awk '{tot+=$2;print 2*tot/NR}'| tail -n 1) recent seconds per sample
echo
echo

families=($(cd ../families ; ls -d auSSC*))
echo families ${#families[*]} of $(cat ../families.txt|wc -l)

dirs=($(ls -d SSC*))
echo samples ${#dirs[*]}

in_sge=($(cat qstat.txt | grep job_name | awk '{print $2}' | sed 's/\.calc// ; s/\.map// ; s/fix[0-9]\.//' | sort | uniq))
echo in_sge ${#in_sge[*]}

alldone=($(echo ${dirs[*]} ${in_sge[*]} | sed 's/ /\n/g' | sort | uniq -c | awk '{if ($1 == 1) { print $2}}'))
echo alldone ${#alldone[*]} # ${alldone[*]}
echo below here only done samples are counted
echo

success=($(for dir in ${alldone[*]} ; do (cd $dir ; if [ -e success ] ; then echo $dir ; fi) ; done))
echo success ${#success[*]}

bin=($(for dir in ${alldone[*]} ; do (cd $dir ; if [ -e rmdup.bam.bin ] ; then echo $dir ; fi) ; done))
echo bin ${#bin[*]}

no_bam=($(for dir in ${success[*]} ; do (cd $dir ;  if [ ! -e rmdup.bam ] ; then echo $dir ; fi ) done))
echo no_bam ${#no_bam[*]} # ${no_bam[*]}

no_bin=($(for dir in ${success[*]} ; do (cd $dir ;  if [ ! -e rmdup.bam.bin ] ; then echo $dir ; fi ) done))
echo no_bin ${#no_bin[*]} ${no_bin[*]}
echo

no_success=($(for dir in ${alldone[*]} ; do (cd $dir ; if [ ! -e success ] ; then echo $dir ; fi) ; done))
echo no_success ${#no_success[*]} #dirs=\"${no_success[*]}\"
echo

no_map=($(for dir in ${no_success[*]} ; do (cd $dir ; if [ ! -e out.txt ] || ! grep "mumdex has succeeded" out.txt > /dev/null || [ ! -e mapout ] ; then echo $dir ; fi) ; done))
echo no_map ${#no_map[*]} dirs=\"${no_map[*]}\" 
echo for dir in \$dirs \; do rm -Rf \$dir/ \; rm -f \$dir \; done 
echo

map=($(for dir in ${no_success[*]} ; do (cd $dir ; if [ -e out.txt ] && grep "mumdex has succeeded" out.txt > /dev/null && [ -e mapout ] ; then echo $dir ; fi) ; done))
echo map ${#map[*]} dirs=\"${map[*]}\" 
echo


# last=() ; for dir in ... ; do node=$(readlink $dir) ; node=${node#*wigclust} ; node=${node%/data*} ; echo $dir $node ${last[$node]} ; (cd $dir ; echo 'rm -R rmdup.bam.bin ; rm success ; /data/unsafe/paa/mums-new/mumdex/gav quit rmdup.bam || exit 0 ;  rm -R rmdup.bam rmdup.bam.bai ; touch success ' | qsub -q all.q@wigclust$node -cwd -o fix.txt -e fix.err.txt -N fix3.$dir $(if [ "${last[$node]}" != "" ] ; then echo -hold_jid fix3.${last[$node]} ; fi)) ; last[$node]=$dir ; done

rm -f qstat.txt 
```
