## Supplementary material for "G-Graph: An interactive genomic graph viewer": MUMdex Source Code for G-Graph: karyotype.cpp

```
//
// karyotype.cpp
//
// draw a karyotype
//
// Copyright 2016 Peter Andrews @ CSHL
//

#include <algorithm>
#include <exception>
#include <iostream>
#include <fstream>
#include <map>
#include <sstream>
#include <string>
#include <vector>

#include "error.h"
#include "mumdex.h"
#include "repeats.h"

using std::cin;
using std::cout;
using std::cerr;
using std::endl;
using std::exception;
using std::ifstream;
using std::istringstream;
using std::map;
using std::ostringstream;
using std::string;
using std::vector;

using paa::ChromosomeIndexLookup;
using paa::Error;
using paa::Reference;

int main(int argc, char * argv[]) try {
  if (--argc != 4) {
    throw Error("usage: karyotype ref bands_file title scale");
  }

  const Reference ref{argv[1]};
  const ChromosomeIndexLookup chr_lookup{ref};
  ifstream bands_file{argv[2]};
  const string title{argv[3]};
  const double scale{atof(argv[4])};

#if 0
  const vector<string> show_chromosomes{"chr1", "chr2", "chr3", "chr4",
        "chr5", "chr6", "chr7", "chr8", "chr9", "chr10", "chr11", "chr12",
        "chr13", "chr14", "chr15", "chr16", "chr17", "chr18", "chr19", "chr20",
        "chr21", "chr22", "chrX", "chrY"};
#endif

  const vector<string> show_chromosomes{"1", "2", "3", "4",
        "5", "6", "7", "8", "9", "10", "11", "12",
        "13", "14", "15", "16", "17", "18", "19", "20",
        "21", "22", "X", "Y"};

  const map<string, unsigned int> chromosome_index{[&show_chromosomes]() {
      map<string, unsigned int> index;
      for (unsigned int c{0}; c != show_chromosomes.size(); ++c) {
        index[show_chromosomes[c]] = c;
      }
      return index;
    }()};

  const map<string, string> band_colors{
    { "acen", "1 0 1" },
    { "gieStain", "1 1 0" },
    { "gneg", "1 1 1" },
    { "gpos100", "0 0 0" },
    { "gpos25", "0.75 0.75 0.75" },
    { "gpos50", "0.5 0.5 0.5" },
    { "gpos75", "0.25 0.25 0.25" },
    { "gvar", "0 1 1" },
    { "stalk", "1 0 0" },
    { "centromere", "0 0 0" },
    { "telomere", "0 0 0" }};

  const unsigned int page_width{612};
  const unsigned int page_height{792};
  const unsigned int title_height{20};
  const unsigned int padding{20};
  const unsigned int n_chr{static_cast<unsigned int>(show_chromosomes.size())};
  const unsigned int half_chr{(n_chr + 1) / 2};
  const double chr_y_spacing{(page_height - 3.0 * padding - title_height) /
        half_chr + 5};
  const double chr_x_spacing{(page_width - 2.0 * padding) / 2 + 80};
  const double pos_scale{(chr_x_spacing - padding) / 249250621};
  const double chr_height{6};

  cout << R"foo(%!PS-Adobe-3.0
%%BoundingBox: 0 0 )foo" << page_width << " " << page_height
     << R"foo(
%%Pages: (atend)
%%Title: Karyotype
%%Creator: Peter Andrews
%%CreationDate: Today
%%EndComments
%%BeginProlog
%%EndProlog
%%Page: 1 1
)foo";

  cout << "/Helvetica findfont " << title_height
       << " scalefont setfont" << endl;
  cout << page_width / 2 << " " << page_height - title_height - 10
       << " moveto (" << title << ") dup stringwidth pop 2 div neg 0 rmoveto "
       << " show" << endl;

  cout << "/Helvetica findfont 10 scalefont setfont" << endl;


  const auto xpos = [chr_x_spacing, pos_scale, half_chr]
      (const unsigned int chr_index, const unsigned int pos) {
    return (1.3 * padding + chr_index / half_chr * chr_x_spacing +
            pos * pos_scale);
  };

  const auto ypos = [chr_y_spacing, half_chr]
      (const unsigned int chr_index) {
    return padding + (chr_index % half_chr) * chr_y_spacing;
  };

  // Draw chromosome outlines
  for (const string & schr : show_chromosomes) {
    const unsigned int chr{chr_lookup[schr]};
    const unsigned int chr_index{chromosome_index.at(schr)};
    const double xl{xpos(chr_index, 0)};
    const double xh{xpos(chr_index, ref.size(chr))};
    const double yl{ypos(chr_index)};
    const double yh{yl + chr_height};
    cout << "newpath " << xl << " " << yl << " moveto "
         << "gsave (" << schr
         << ") dup stringwidth pop neg 5 sub 0 rmoveto show grestore "
         << xh << " " << yl << " lineto "
         << xh << " " << yh << " lineto "
         << xl << " " << yh << " lineto "
         << "closepath stroke"
         << endl;
    if (0)
    cout << xl << " " << yl << " moveto "
         << "(" << chr
         << ") dup stringwidth pop neg 5 sub 0 rmoveto show"
         << endl;
  }

  bands_file.ignore(1000, '\n');
  unsigned int bin;
  string chr;
  unsigned int start;
  unsigned int stop;
  unsigned int ix;
  string n;
  unsigned int size;
  string name;
  string stain;
  string bridge;
  string last_chr;
  while (bands_file) {
    if (0) {
      bands_file >> chr >> start >> stop >> name >> stain;
    } else {
      bands_file >> bin >> chr >> start >> stop >> ix >> n >> size
                 >> stain >> bridge;
    }
    if (!bands_file) break;
    try {
      const unsigned int chr_index{chromosome_index.at(chr)};
      const double xl{xpos(chr_index, start)};
      const double xh{xpos(chr_index, stop)};
      const double yl{ypos(chr_index)};
      const double yh{yl + chr_height};

      if (last_chr != chr) {
      }
      const string color{band_colors.at(stain)};
      cout << "newpath " << xl << " " << yl << " moveto "
           << xh << " " << yl << " lineto "
           << xh << " " << yh << " lineto "
           << xl << " " << yh << " lineto "
           << "closepath gsave "
           << color << " setrgbcolor fill grestore stroke"
           << endl;
    } catch (std::exception & e) {
      if (string("map::at") == e.what()) {
        continue;
      } else {
        throw;
      }
    }
  }

  vector<vector<bool>> mask(page_width * scale,
                            vector<bool>(page_height * scale));
  string schr;
  unsigned int pos;
  while (cin >> schr >> pos) {
    try {
      const unsigned int chr_index{chromosome_index.at(schr)};
      double off{0};
      const double x{xpos(chr_index, pos)};
      while (true) {
        const double y{ypos(chr_index) + chr_height + off + 2};
        if (x * scale > mask.size()) continue;
        if (y * scale > mask[x * scale].size()) break;
        if (mask[x * scale][y * scale] ||
            mask[(x - 1) * scale][y * scale] ||
            mask[(x + 1) * scale][y * scale] ||
            mask[(x - 2) * scale][y * scale] ||
            mask[(x + 2) * scale][y * scale]) {
          off += 1.9;
        } else {
          mask[x * scale][y * scale] = true;
          cout << "newpath " << x << " " << y << " 0.8 0 360 arc fill" << endl;
          break;
        }
      }
    } catch (std::exception & e) {
      if (string("map::at") == e.what()) {
        continue;
      } else {
        throw;
      }
    }
  }

  cout << "showpage" << endl;
  cout << "%%EndPage: 1" << endl;
  cout << "%%Trailer" << endl;
  cout << "%%Pages: 1" << endl;
  cout << "%%EOF" << endl;

  return 0;
} catch (Error & e) {
  cerr << "paa::Error:" << endl;
  cerr << e.what() << endl;
  return 1;
} catch (exception & e) {
  cerr << "std::exception" << endl;
  cerr << e.what() << endl;
  return 1;
} catch (...) {
  cerr << "unknown exception was caught" << endl;
  return 1;
}
```
