## Supplementary material for "G-Graph: An interactive genomic graph viewer": MUMdex Source Code for G-Graph: local_similarity.cpp

```
//
// local_similarity
//
// a measure of local similarity to a reference region
//
// Copyright 2016 Peter Andrews @ CSHL
//

#include <exception>
#include <iostream>

#include "error.h"
#include "mumdex.h"

using std::cout;
using std::cerr;
using std::endl;
using std::exception;

using paa::ChromosomeIndexLookup;
using paa::Error;
using paa::Reference;

int main(int argc, char * argv[]) try {
  if (--argc != 5)
    throw Error("usage: local_similarity ref chr pos edge ref_size");

  const Reference ref{argv[1]};
  const ChromosomeIndexLookup chr_lookup{ref};
  const unsigned int chr{chr_lookup[argv[2]]};
  const unsigned int pos{static_cast<unsigned int>(atoi(argv[3]))};
  const unsigned int edge{static_cast<unsigned int>(atoi(argv[4]))};
  const unsigned int size{static_cast<unsigned int>(atoi(argv[5]))};
  const unsigned int hsize{size / 2};

  const unsigned int seq_start{pos > edge ? pos - edge : 0};
  const unsigned int seq_stop{pos + edge < ref.size(chr) ?
        pos + edge : ref.size(chr)};
  const unsigned int seq_size{seq_stop - seq_start};

  const unsigned int ref_start{pos > hsize ? pos - hsize : 0};
  const unsigned int ref_stop{pos + hsize < ref.size(chr) ?
        pos + hsize : ref.size(chr)};
  const unsigned int ref_size{ref_stop - ref_start};
  if (ref_size <= seq_size) throw Error("seq size bigger than ref size");

  const char * ref_chr{ref[chr]};
  unsigned int max_similarity{0};
  unsigned int offset{0};
  for (unsigned int r{ref_start}; r + seq_size != ref_stop; ++r) {
    if (r == seq_start) continue;
    unsigned int similarity{0};
    for (unsigned int s{0}; s != seq_size; ++s) {
      similarity += ref_chr[r + s] == ref_chr[seq_start + s];
    }
    if (max_similarity < similarity) {
      max_similarity = similarity;
      offset = r > seq_start ? r - seq_start : seq_start - r;
    }
  }
  cout << 1.0 * max_similarity / seq_size << " " << offset << endl;

  return 0;
} catch (Error & e) {
  cerr << "paa::Error:" << endl;
  cerr << e.what() << endl;
  return 1;
} catch (exception & e) {
  cerr << "std::exception" << endl;
  cerr << e.what() << endl;
  return 1;
} catch (...) {
  cerr << "unknown exception was caught" << endl;
  return 1;
}
```
