## Supplementary material for "G-Graph: An interactive genomic graph viewer": MUMdex Source Code for G-Graph: logs.h

```
//
// logs.h
//
// simple computations with logarithms, mainly for EM methods
//
// Copyright 2019 Peter Andrews @ CSHL
//
//

#ifndef PAA_LOGS_H
#define PAA_LOGS_H

#include <limits>

namespace paa {

constexpr bool log_paranoia{false};

template <class FLOAT>
inline FLOAT lnproduct(const FLOAT lhs, const FLOAT rhs) {
  if (!log_paranoia) return lhs + rhs;
  if (lhs > std::numeric_limits<FLOAT>::lowest() &&
      rhs > std::numeric_limits<FLOAT>::lowest()) {
    return lhs + rhs;
  } else {
    return std::numeric_limits<FLOAT>::lowest();
  }
}

template <class FLOAT>
inline FLOAT lnproduct(const FLOAT a1, const FLOAT a2, const FLOAT a3) {
  return lnproduct(a1, lnproduct(a2, a3));
}
#if 0
template <class FLOAT, class ... ARGS>
inline FLOAT lnsum(const FLOAT lhs, const FLOAT rhs, const ARGS ... rest) {
  return lnsum(lhs, lnsum(rhs, rest...));
}
#endif
template <class FLOAT>
inline FLOAT lnsum(const FLOAT lhs, const FLOAT rhs) {
  if (log_paranoia) {
    if (lhs <= std::numeric_limits<FLOAT>::lowest())
      return rhs;
    if (rhs <= std::numeric_limits<FLOAT>::lowest())
      return lhs;
  }
  if (lhs > rhs)
    return lhs + log(1 + exp(rhs - lhs));
  return rhs + log(1 + exp(lhs - rhs));
}

template <class Vec>
inline double lnsum(const Vec & vec, const uint64_t n) {
  double result{vec[0]};
  for (uint64_t i{1}; i != n; ++i) result = lnsum(vec[i], result);
  return result;
}

}  // namespace paa

#endif  // PAA_LOGS_H
```
