## Supplementary material for "G-Graph: An interactive genomic graph viewer": MUMdex Source Code for G-Graph: make_reference.cpp

```
//
// make_reference
//
// assemble a reference genome from chromosome pieces
//
// Copyright 2018 Peter Andrews @ CSHL
//

#include <exception>
#include <fstream>
#include <iostream>
#include <string>
#include <vector>

#include "error.h"
#include "mumdex.h"
#include "strings.h"

using std::cerr;
using std::cin;
using std::cout;
using std::endl;
using std::exception;
using std::ifstream;
using std::ofstream;
using std::string;
using std::vector;

using paa::remove_substring;
using paa::Error;
using paa::Reference;

int main(int argc, char* argv[])  try {
  if (--argc < 2) throw Error("usage: make_reference out_fa chr_fasta ...");

  const string masked_ref_name{argv[1]};
  ofstream masked_ref_stream{masked_ref_name.c_str()};
  if (!masked_ref_stream)
    throw Error("Problem opening output reference") << masked_ref_name;
  ++argv; --argc;

  const string temp_ref_name{"temp.fa"};
  ofstream temp_ref_stream{temp_ref_name.c_str()};
  if (!temp_ref_stream)
    throw Error("Problem opening temp reference") << temp_ref_name;

  while (argc--) {
    const string chr_fa{argv++[1]};
    const string chr_name{remove_substring(chr_fa, ".fa")};
    cerr << "Working on chromosome " << chr_name << endl;
    ifstream chr_stream{chr_fa.c_str()};
    if (!chr_stream)
      throw Error("Problem opening chromosome reference") << chr_fa;
    string line;
    while (getline(chr_stream, line)) temp_ref_stream << line << "\n";
  }
  temp_ref_stream.close();

  const Reference temp_ref{temp_ref_name};
  const unsigned int y{temp_ref.find_y_chromosome()};
  const bool is_hg38{temp_ref.size(y) == 57227415};
  const bool is_hg19{temp_ref.size(y) == 59373566};
  if (!is_hg38 && !is_hg19)
    throw Error("Cannot identify reference as hg19 or hg38");

  const unsigned int hg19_par_limits[2][2]
  {{10000, 2649520}, {59034049, 59363566}};
  const unsigned int hg38_par_limits[2][2]
  {{10000, 2781479}, {56887902, 57217415}};
  const auto par_limits = is_hg19 ? hg19_par_limits : hg38_par_limits;

  const unsigned int line_length{50};
  unsigned int n_line{0};
  for (unsigned int c{0}; c != temp_ref.n_chromosomes(); ++c) {
    masked_ref_stream << ">" << temp_ref.name(c) << "\n";
    for (unsigned int b{0}; b != temp_ref.size(c); ++b) {
      if (c == y && (
              (b >= par_limits[0][0] && b < par_limits[0][1]) ||
              (b >= par_limits[1][0] && b < par_limits[1][1]))) {
        masked_ref_stream << "N";
      } else {
        masked_ref_stream << temp_ref[c][b];
      }
      if (++n_line == line_length || b + 1 == temp_ref.size(c)) {
        masked_ref_stream << "\n";
        n_line = 0;
      }
    }
    n_line = 0;
  }

  // system((string("rm -Rf ") + temp_ref_name + "*").c_str() );

  return 0;
} catch (Error & e) {
  cerr << "paa::Error:" << endl;
  cerr << e.what() << endl;
  return 1;
} catch (exception & e) {
  cerr << "std::exception" << endl;
  cerr << e.what() << endl;
  return 1;
} catch (...) {
  cerr << "unknown exception was caught" << endl;
  return 1;
}
```
