## Supplementary material for "G-Graph: An interactive genomic graph viewer": MUMdex Source Code for G-Graph: male_cn_stats.cpp

```
//
// male_cn_stats.cpp
//
// Look at a male copy number profile and report stats like signal to noise
//
// Copyright 2016 Peter Andrews @ CSHL
//

#include <algorithm>
#include <exception>
#include <fstream>
#include <iostream>
#include <string>
#include <vector>

#include "error.h"
#include "mumdex.h"
#include "stats.h"

using std::cin;
using std::cout;
using std::cerr;
using std::endl;
using std::exception;
using std::ifstream;
using std::string;
using std::vector;

using paa::AutoCorr;
using paa::ChromosomeIndexLookup;
using paa::Error;
using paa::MAD;
using paa::MADT;
using paa::NormalParams;
using paa::Reference;

using iMAD = paa::MADT<unsigned int>;

int main(int argc, char* argv[]) try {
  if (--argc != 2) throw Error("usage: male_cn_stats ref binned_file");

  const Reference ref{argv[1]};
  const std::string binned_name{argv[2]};
  ifstream binned_file{binned_name.c_str()};
  if (!binned_file) throw Error("Problem opening binned file") << binned_name;

  const ChromosomeIndexLookup lookup{ref};
  const unsigned int x_chr{ref.find_x_chromosome()};
  const unsigned int y_chr{ref.find_y_chromosome()};

  string chr_name;
  unsigned int chrpos;
  unsigned int abspos;
  unsigned int length;
  double gc;
  double map;
  unsigned int count;
  double ratio;
  double seg_ratio;
  double quantal;
  double seg_quantal;

  unsigned int n_bins{0};
  double last_seg_ratio{0};
  unsigned int n_segments{0};
  binned_file.ignore(10000, '\n');
  vector<double> ratios[2];
  vector<double> counts[2];
  double min_ratio{1000000000};
  double max_ratio{0};
  double norm;
  double corr;
  double ncount;
  while (binned_file >> chr_name >> chrpos >> abspos
         >> length >> norm >> corr >> gc >>  map
         >> count >> ncount >> ratio >> seg_ratio >> quantal >> seg_quantal) {
    ++n_bins;

    if (seg_ratio < last_seg_ratio || seg_ratio > last_seg_ratio) {
      ++n_segments;
      last_seg_ratio = seg_ratio;
    }

    const unsigned int chr{lookup[chr_name]};
    if (chr == y_chr) continue;
    if (chr == x_chr &&
        (chrpos < 2781479 + 10000 || chrpos + 10000 > 155701382)) continue;
    if (ratio > max_ratio) max_ratio = ratio;
    if (ratio < min_ratio) min_ratio = ratio;
    ratios[chr == x_chr].push_back(quantal);
    counts[chr == x_chr].push_back(ncount);
  }

  // Actual stats
  const string names[2]{"Autosome", "X chromosome"};
  const NormalParams normals[2]{ratios[0], ratios[1]};
  const AutoCorr autocorrs[2]{{ratios[0], normals[0]},
    {ratios[1], normals[1]}};
  for (const bool x : { false, true }) {
    sort(ratios[x].begin(), ratios[x].end());
  }
  const MAD mads[2]{ratios[0], ratios[1]};
  for (const bool x : { false, true }) {
    sort(counts[x].begin(), counts[x].end());
  }
  const MAD count_mads[2]{counts[0], counts[1]};

  for (const bool x : { false, true }) {
    const NormalParams & normal{normals[x]};
    const AutoCorr & autocorr{autocorrs[x]};
    const MAD & mad{mads[x]};
    const MAD & count_mad{count_mads[x]};
    cout << names[x] << ": "
         << ratios[x].size() << " values, "
         << count_mad.median() << " median count, "
         << "quantal ratio "
         << normal.mean << " mean "
         << normal.stdev << " stdev "
         << mad.median() << " median "
         << mad.mad() << " mad "
         << autocorr.autocorr << " autocorr "
         << endl;
  }

  const double signal{normals[0].mean - normals[1].mean};
  const double noise{sqrt(pow(normals[0].stdev, 2) + pow(normals[1].stdev, 2))};
  const double signal_to_noise{signal / noise};

  const double mad_signal{mads[0].median() - mads[1].median()};
  const double mad_noise{sqrt(pow(mads[0].stdev(), 2) +
                              pow(mads[1].stdev(), 2))};
  const double mad_signal_to_noise{mad_signal / mad_noise};

  // Theoretical stats
  const double a_count_median{count_mads[0].median()};
  const double x_count_median{count_mads[1].median()};
  const double theoretical_signal{a_count_median - x_count_median};
  const double theoretical_noise{sqrt(a_count_median + x_count_median)};
  const double theoretical_signal_to_noise{theoretical_signal /
        theoretical_noise};

  // Summary
  cout << "Summary: "
       << n_bins << " bins, "
       << n_segments << " segments, "
       << "signal " << signal << ", noise " << noise << ", "
       << "signal to noise " << signal_to_noise << ", "
       << "mad signal to noise " << mad_signal_to_noise << ", "
       << "theoretical signal to noise " << theoretical_signal_to_noise << ", "
       << "shortfall " << signal_to_noise / theoretical_signal_to_noise << ", "
       << "mad shortfall " << (mad_signal_to_noise /
                               theoretical_signal_to_noise) << ", "
       << "autosome autocorr " << autocorrs[0].autocorr << " "
       << "minmax " << min_ratio << " " << max_ratio << endl;

  return 0;
} catch (Error & e) {
  cerr << "paa::Error:" << endl;
  cerr << e.what() << endl;
  return 1;
} catch (exception & e) {
  cerr << "std::exception" << endl;
  cerr << e.what() << endl;
  return 1;
} catch (...) {
  cerr << "unknown exception was caught" << endl;
  return 1;
}
```
