## Supplementary material for "G-Graph: An interactive genomic graph viewer": MUMdex Source Code for G-Graph: mapper.cpp

```
//
// mapper
//
// simple mum mapper maps reads from cin
//
// Copyright 2016 Peter Andrews @ CSHL
//

#include "mapper.h"

#include <exception>
#include <iostream>
#include <string>
#include <vector>

#include "error.h"
#include "longSA.h"
#include "mumdex.h"
#include "utility.h"

using std::cerr;
using std::cin;
using std::cout;
using std::endl;
using std::exception;
using std::string;
using std::vector;

using paa::longSA;
using paa::read_ahead;
using paa::sout;
using paa::Error;
using paa::Reference;
using paa::SimpleHit;

int main(int argc, char* argv[])  try {
  read_ahead = true;
  std::ios_base::sync_with_stdio(false);
  cin.tie(nullptr);

  if (--argc != 1) throw Error("usage: mapper ref_fasta");

  const string ref_fasta{argv[1]};
  const longSA sa{ref_fasta, true, true, false};
  const Reference ref{ref_fasta};

  string read;
  while (cin >> read) {
    cout << read << '\n';
    const vector<SimpleHit> mums{sa.find_mams(read)};
    for (const auto mum : mums) {
      sout << ref.name(mum.chr) << mum.pos
           << mum.off << mum.len << mum.dir << '\n';
    }
  }

  return 0;
} catch (Error & e) {
  cerr << "paa::Error:" << endl;
  cerr << e.what() << endl;
  return 1;
} catch (exception & e) {
  cerr << "std::exception" << endl;
  cerr << e.what() << endl;
  return 1;
} catch (...) {
  cerr << "unknown exception was caught" << endl;
  return 1;
}
```
