## Supplementary material for "G-Graph: An interactive genomic graph viewer": MUMdex Source Code for G-Graph: matrix.h

```
//
// matrix.h
//
// simple matrix computations
//
// Copyright 2019 Peter Andrews @ CSHL
//
//

#ifndef PAA_MATRIX_H
#define PAA_MATRIX_H

#include <algorithm>
#include <iostream>
#include <vector>

#include "error.h"

namespace paa {

// Matrix with two compile-time dimensions
template <class Val, uint64_t II = 0, uint64_t JJ = 0>
class Matrix {
 public:
  static constexpr uint64_t I{II};
  static constexpr uint64_t J{JJ};
  Matrix() {}  // Uninitialized!
  Matrix(const std::initializer_list<std::initializer_list<Val> > & input) {
    auto rows = input.begin();
    for (uint64_t i{0}; i != I; ++i, ++rows) {
      auto val = rows->begin();
      for (uint64_t j{0}; j != J; ++j, ++val) (*this)[i][j] = *val;
    }
  }

  const Val * operator[](const uint64_t i) const { return data + i * J; }
  Val * operator[](const uint64_t i) { return data + i * J; }
  void swap(Matrix & other) {
    other.data.swap(data);
  }

 private:
  Val data[I * J];
};

// Base class for dynamic matrices
template <class Val> struct MBase {
  explicit MBase(const uint64_t n_elem) : data(n_elem) { }
  void swap(MBase & other) { other.data.swap(data); }
  std::vector<Val> data;
};
// Matrix with two run-time dimensions
template <class Val>
class Matrix<Val, 0, 0> : private MBase<Val> {
 public:
  const uint64_t I;
  const uint64_t J;
  Matrix(const uint64_t I_, const uint64_t J_) :
      MBase<Val>{I_ * J_}, I{I_}, J{J_} { }
  const Val * operator[](const uint64_t i) const { return &this->data[i * J]; }
  Val * operator[](const uint64_t i) { return &this->data[i * J]; }
  void swap(Matrix & other) {
    if (I != other.I) throw Error("Matrix I size mismatch") << I << other.I;
    if (J != other.J) throw Error("Matrix J size mismatch") << J << other.J;
    MBase<Val>::swap(other);
  }
};
// Matrix with run-time first dimension and compile-time second dimension
template <class Val, uint64_t JJ>
class Matrix<Val, 0, JJ> : private MBase<Val> {
 public:
  const uint64_t I;
  static constexpr uint64_t J{JJ};
  explicit Matrix(const uint64_t I_) : MBase<Val>{I_ * J}, I{I_} { }
  const Val * operator[](const uint64_t i) const { return &this->data[i * J]; }
  Val * operator[](const uint64_t i) { return &this->data[i * J]; }
  template <class Array> void assign(const Array & a) {
    for (uint64_t i{0}; i != I; ++i)
      for (uint64_t j{0}; j != J; ++j)
        this->data[i * J + j] = a[j];
  }
  void swap(Matrix & other) {
    if (I != other.I) throw Error("Matrix I size mismatch") << I << other.I;
    MBase<Val>::swap(other);
  }
};
// Matrix with compile-time first dimension and run-time second dimension
template <class Val, uint64_t II>
class Matrix<Val, II, 0> : private MBase<Val> {
 public:
  static constexpr uint64_t I{II};
  const uint64_t J;
  explicit Matrix(const uint64_t J_) : MBase<Val>{I * J_}, J{J_} { }
  const Val * operator[](const uint64_t i) const { return &this->data[i * J]; }
  Val * operator[](const uint64_t i) { return &this->data[i * J]; }
  void swap(Matrix & other) {
    if (J != other.J) throw Error("Matrix J size mismatch") << J << other.J;
    MBase<Val>::swap(other);
  }
};
// Output for matrices
template <class Val, uint64_t I, uint64_t J>
std::ostream & operator<<(std::ostream & stream,
                        const Matrix<Val, I, J> & matrix) {
  for (uint64_t i{0}; i != matrix.I; ++i) {
    for (uint64_t j{0}; j != matrix.J; ++j) {
      if (j) stream << '\t';
      stream << matrix[i][j];
    }
    if (i + 1 != matrix.I) stream << '\n';
  }
  return stream;
}

// Agrees with eigenvalue-based general method
inline Matrix<double, 2, 2> matrix_power(const Matrix<double, 2, 2> & unit,
                                         const uint64_t power) {
  const double a{unit[0][1]};
  const double b{unit[1][0]};
  const double a_plus_b{a + b};
  const double gamma{pow(1 - a - b, power)};
  return Matrix<double, 2, 2>{
    {(a * gamma + b) / a_plus_b, (a - a * gamma) / a_plus_b},
    {(b - b * gamma) / a_plus_b, (b * gamma + a) / a_plus_b}};
}

#if 0
#include <Eigen/Eigenvalues>
template <int NRows>
class MatrixPowers {
 public:
  using Internal = Eigen::Matrix<double, NRows, NRows>;
  MatrixPowers() {}

  template <class Matrix>
  explicit MatrixPowers(const Matrix & original) {
    initialize(original);
  }

  template <class Matrix>
  void initialize(const Matrix & original) {
    if (original.size() != NRows)  // || original[0].size() != NRows)
      throw Error("Bad Matrix size in MatrixPowers");
    for (unsigned int i{0}; i != NRows; ++i) {
      for (unsigned int j{0}; j != NRows; ++j) {
        input(i, j) = original[i][j];
      }
    }
    std::cerr << "Input matrix:" << std::endl
              << input << std::endl;
    solver.compute(input);
    std::cerr << "Eigenvalues:" << std::endl
              << (eigenvalues = solver.eigenvalues().real()) << std::endl;
    std::cerr << "Eigenvector matrix:" << std::endl
              << (eigenvectors = solver.eigenvectors().real()) << std::endl;
    eigenvalues = solver.eigenvalues().real();
    inverse_eigenvectors = solver.eigenvectors().real().inverse();
  }

  Internal operator()(const double d) const {
    Internal result{Internal::Zero()};
    for (unsigned int r{0}; r != eigenvalues.size(); ++r) {
      result(r, r) = pow(eigenvalues(r), d);
    }
    return eigenvectors * result * inverse_eigenvectors;
  }

  static void test() {
    // Test Matrix powers
    const double a{0.00001};
    const double b{0.00005};
    const Matrix2d<2, 2> trans{{1 - a, a}, {b, 1 - b}};
    const paa::MatrixPowers<2> powers{trans};
    for (const double power : {0, 1, 10, 100, 10000, 100000, 1000000}) {
      std::cout << "Power is " << power << ":" << std::endl;
      std::cout << powers(power) << std::endl;
      std::cout << "new:" << std::endl;
      std::cout << matrix_power(trans, power) << std::endl;
      std::cout << std::endl;
    }
  }

 private:
  Eigen::Matrix<double, NRows, 1> eigenvalues{};
  Internal input{};
  Internal eigenvectors{};
  Internal inverse_eigenvectors{};
  Eigen::EigenSolver<Internal> solver{};
};
#endif

}  // namespace paa

#endif  // PAA_MATRIX_H
```
