## Supplementary material for "G-Graph: An interactive genomic graph viewer": MUMdex Source Code for G-Graph: merger.h

```
//
// merger.cpp
//
// parallel merger - probably not too useful in practice
//
// Copyright 2016 Peter Andrews @ CSHL
//

#include <string>
#include <vector>

template <class TYPE>
class MappedVectorQueue {
 public:
  MappedVectorQueue() { }
  explicit MappedVectorQueue(const string & file_name) :
      file{file_name} { }
  ~MappedVectorQueue() { }
  TYPE top() const {
    return file[current];
  }
  bool pop() {
    if (++current == file.size()) {
      return false;
    } else {
      return true;
    }
  }
  uint64_t size() const {
    return file.size();
  }

 private:
  uint64_t current{0};
  MappedVector<TYPE> file;
};

template <class TYPE>
class ParallelMerger {
 public:
  TYPE top() const {
    if (is_file) {
      return file->top();
    } else {
      if (ready.empty()) throw Error("Empty ready");
      return ready[ready_index];
    }
  }

  bool pop() {
    if (is_file) {
      return file->pop();
    } else {
      if (ready.empty()) throw Error("Empty ready");
      if (++ready_index == ready.size()) {
        if (!fill_thread.valid()) {
          return false;
        }
        if (wait_until_ready()) {
          start_filling();
          return true;
        } else if (ready.empty()) {
          return false;
        } else {
          return true;
        }
      } else {
        return true;
      }
    }
  }

  uint64_t size() const {
    if (is_file) {
      return file->size();
    } else {
      return left->size() + right->size();
    }
  }
  ~ParallelMerger() { }

 public:
  ParallelMerger(const ParallelMerger&) = delete;
  explicit ParallelMerger(const vector<string> & filenames) {
    if (filenames.size() < 1) {
      throw Error("Too few filenames passed to ParallelMerger");
    }
    if (filenames.size() == 1) {
      is_file = true;
      file = make_unique<MappedVectorQueue<TYPE>>(filenames.front());
    } else {
      is_file = false;
      const auto midpoint = filenames.begin() + filenames.size() / 2;
      const vector<string> left_names(filenames.begin(), midpoint);
      const vector<string> right_names(midpoint, filenames.end());
      left = make_unique<ParallelMerger<TYPE>>(left_names);
      right = make_unique<ParallelMerger<TYPE>>(right_names);
      left_good = left->size();
      right_good = right->size();
      const bool nonempty = left_good || right_good;
      if (left_good && !left->is_file && left->wait_until_ready()) {
        left->ready.reserve(max_size);
        left->start_filling();
      }
      if (right_good && !right->is_file && right->wait_until_ready()) {
        right->ready.reserve(max_size);
        right->start_filling();
      }
      if (nonempty) {
        ready.reserve(max_size);
        start_filling();
      }
    }
  }

  void start_filling() {
    fill_thread = async(std::launch::async, ref(*this));
  }
  bool wait_until_ready() {
    const bool result = fill_thread.get();
    swap();
    return result;
  }

  // generate filling vector
  bool operator()() {
    if (is_file) throw Error("file filling");
    while (left_good || right_good) {
      if (!left_good) {
        filling.push_back(right->top());
        right_good = right->pop();
      } else if (!right_good) {
        filling.push_back(left->top());
        left_good = left->pop();
      } else if (right->top() < left->top()) {
        filling.push_back(right->top());
        right_good = right->pop();
      } else {
        filling.push_back(left->top());
        left_good = left->pop();
      }
      if (filling.size() == max_size) break;
    }
    return left_good || right_good;
  }

  void swap() {
    if (is_file) throw Error("file swapping");
    filling.swap(ready);
    filling.clear();
    ready_index = 0;
  }

  uint64_t n_files() {
    if (is_file) {
      return 1;
    } else {
      return left->n_files() + right->n_files();
    }
  }

  uint64_t max_size{10000};
  uint64_t ready_index{0};
  vector<TYPE> ready;
  vector<TYPE> filling;
  future<bool> fill_thread;
  unique_ptr<ParallelMerger<TYPE>> left;
  unique_ptr<ParallelMerger<TYPE>> right;
  bool left_good;
  bool right_good;
  unique_ptr<MappedVectorQueue<TYPE>> file;
  bool is_file;
};

template <class TYPE>
unique_ptr<ParallelMerger<TYPE>>
make_merger(const vector<string> & filenames) {
  if (filenames.empty()) {
    throw Error("No filenames passed to make_merger");
  } else {
    auto merger = make_unique<ParallelMerger<TYPE>>(filenames);
    if (filenames.size() > 1 &&
        merger->size() && merger->wait_until_ready()) {
      merger->start_filling();
    }
    return merger;
  }
}

void test_merger(const uint64_t n) {
  {
    MappedVector<int> one;
    MappedVector<int> two;
    MappedVector<int> three;
    MappedVector<int> four;
    MappedVector<int> five;
    for (unsigned int i = 0; i != n; ++i) {
      one.push_back(i+1);
      two.push_back(i+2);
      three.push_back(i+3);
      four.push_back(i+4);
      five.push_back(i+5);
    }
    one.save("one");
    two.save("two");
    three.save("three");
    four.save("four");
    five.save("five");
  }
  vector<string> names{"one", "two", "three", "four", "five"};
  auto intmerger = make_merger<int>(names);
  int i = 0;
  while (true) {
    const int result = intmerger->top();
    cout << "output " << result << " " << ++i << endl;
    if (!intmerger->pop()) {
      cout << "quit" << endl;
      break;
    }
  }
}
```
