## Supplementary material for "G-Graph: An interactive genomic graph viewer": MUMdex Source Code for G-Graph: mismatches.cpp

```
//
// mismatches
//
// measure the repeatness of sequences in the genome
// by counting mappings with N mismatches for N in 0..3
//
// Copyright 2018 Peter Andrews @ CSHL
//

#include <exception>
#include <iostream>
#include <map>
#include <sstream>
#include <string>
#include <vector>

#include "error.h"
#include "mumdex.h"
#include "pstream.h"
#include "sequence.h"

using redi::ipstream;

using std::array;
using std::cerr;
using std::cout;
using std::endl;
using std::exception;
using std::istringstream;
using std::map;
using std::ostream;
using std::ostringstream;
using std::string;
using std::to_string;
using std::vector;

using paa::reverse_complement;
using paa::ChromosomeIndexLookup;
using paa::Error;
using paa::Mappability;
using paa::Mismatches;
using paa::Reference;

int main(int argc, char* argv[])  try {
  --argc;
  if (argc != 4 && argc != 6)
    throw Error("mismatches [n_mismatch|lookup] ref_fasta "
                "index chr [start stop]");

  const string first_arg{argv[1]};

  const string ref_fasta{argv[2]};
  const Reference ref{ref_fasta};
  const Mappability mappability{ref};
  const ChromosomeIndexLookup chr_lookup{ref};
  const string index{argv[3]};

  const string chr_name{argv[4]};
  const unsigned int chr{chr_lookup[chr_name]};

  const unsigned int start{argc == 4 ? 0 :
        static_cast<unsigned int>(atoi(argv[5]))};
  const unsigned int stop{argc == 4 ? ref.size(chr) :
        static_cast<unsigned int>(atoi(argv[6]))};


  const unsigned int n_mismatch{static_cast<unsigned int>(atoi(argv[1]))};

  if (first_arg == "lookup") {
    const Mismatches mismatches{ref, "out"};
    for (unsigned int pos{start}; pos != stop; ++pos) {
      const unsigned int abspos{ref.abspos(chr, pos)};
      cout << ref.name(chr) << " " << pos << " "
           << mappability.low(abspos);
      for (unsigned int c{0}; c != Mismatches::n_values; ++c)
        cout << " " << mismatches.count(abspos, c);
      cout << endl;
    }
  } else {
    const unsigned int max_queries{500};

    vector<unsigned int> positions;
    vector<unsigned int> mappabilities;
    vector<string> sequences;
    vector<unsigned int> reasons;
    map<string, unsigned int> seq2index;

    for (unsigned int pos{start}; pos != stop; ++pos) {
      positions.push_back(pos);
      const unsigned int length{mappability.low(ref.abspos(chr, pos))};
      mappabilities.push_back(length);
      if (length > Mismatches::min_len && length < Mismatches::max_len) {
        const string sequence{ref.subseq(chr, pos, pos + length)};
        if (sequence.find_first_not_of("ACGTacgt") == string::npos) {
          seq2index[sequence] = static_cast<unsigned int>(sequences.size());
          sequences.push_back(sequence);
          reasons.push_back(0);
        } else {
          reasons.push_back(2);
          sequences.push_back("");
        }
      } else {
        reasons.push_back(1);
        sequences.push_back("");
      }
      if (sequences.size() == max_queries || pos + 1 == stop) {
        // Prepare bowtie command
        ostringstream sequences_stream;
        unsigned int so{0};
        for (const string & seq : sequences) {
          if (seq.size()) {
            if (so++) sequences_stream << ",";
            sequences_stream << seq;
          }
        }
        vector<vector<unsigned int> > counts(
            sequences.size(), vector<unsigned int>(n_mismatch + 1));
        if (sequences_stream.str().size()) {
          const std::string bowtie_base{
            "/data/software/bowtie/bowtie-0.12.8/bowtie "
                "--quiet --mm -a -v " + to_string(n_mismatch) +
                " --suppress 1,3,4,6,7 "
                "/data/software/bowtie/bowtie-1.2.2/indexes/" +
                index + " -c "};
          const std::string filter{"| perl -pe 's/,/ /g'"};

          // Run bowtie command and get output
          const std::string bowtie_command{
            bowtie_base + sequences_stream.str() + filter};
          if (false) cout << "Running: " << bowtie_command << endl;
          redi::ipstream bowtie{bowtie_command};
          if (!bowtie) throw Error("Bowtie failed on ") << bowtie_command;
          string alignment;
          char orient;
          string sequence;
          string mismatch;
          while (getline(bowtie, alignment)) {
            if (false) cout << "alignment " << alignment << endl;
            unsigned int n_mismatches{0};
            istringstream align_stream{alignment.c_str()};
            align_stream >> orient >> sequence;
            while (align_stream >> mismatch) ++n_mismatches;
            if (orient == '-') reverse_complement(&sequence);
            const unsigned int index{seq2index.at(sequence)};
            ++counts[index][n_mismatches];
          }
        }

        for (unsigned int s{0}; s != sequences.size(); ++s) {
          cout << ref.name(chr) << " " << positions[s]
               << " " << mappabilities[s] << " " << reasons[s];
          for (unsigned int m{0}; m != n_mismatch + 1; ++m)
            cout << " " << counts[s][m];
          cout << "\n";
        }

        positions.clear();
        mappabilities.clear();
        sequences.clear();
        seq2index.clear();
        reasons.clear();
      }
    }
  }

  cerr << "done" << endl;
  return 0;
} catch (Error & e) {
  cerr << "paa::Error:" << endl;
  cerr << e.what() << endl;
  return 1;
} catch (exception & e) {
  cerr << "std::exception" << endl;
  cerr << e.what() << endl;
  return 1;
} catch (...) {
  cerr << "unknown exception was caught" << endl;
  return 1;
}
```
