## Supplementary material for "G-Graph: An interactive genomic graph viewer": MUMdex Source Code for G-Graph: missing_chromosome.cpp

```
//
// missing_chromosome
//
// look for missing chromosomes at the 1% level
//
// Copyright 2016 Peter Andrews @ CSHL
//

#include <algorithm>
#include <exception>
#include <iostream>
#include <memory>
#include <string>
#include <vector>

#include "cn.h"
#include "error.h"
#include "mumdex.h"
#include "psplot.h"
#include "utility.h"

using std::cerr;
using std::cout;
using std::endl;
using std::exception;
using std::make_unique;
using std::string;
using std::unique_ptr;
using std::vector;

using paa::Bin;
using paa::Bounds;
using paa::CN_abspos;
using paa::CN_Bin;
using paa::CN_Bins;
using paa::Error;
using paa::Marker;
using paa::PSDoc;
using paa::PSGraph;
using paa::PSHSeries;
using paa::Reference;

int main(int argc, char* argv[])  try {
  // Check usage
  if (--argc < 4) {
    throw Error("usage: missing_chromosome ref bins cn_output_file...");
  }

  const Reference ref{argv[1]};
  const CN_abspos cn_abspos{ref};
  const string bins_name{argv[2]};
  const vector<Bin> all_bins{load_bins(bins_name, ref)};
  argc -= 2;
  argv += 3;

  const vector<CN_Bins> profiles{[argv, argc] () {
      vector<CN_Bins> result;
      result.reserve(argc);
      for (int a{0}; a != argc; ++a) {
        result.emplace_back(argv[a]);
      }
      return result;
    }()};

  // List of previously determined good bins
  const vector<Bin> bins{[&all_bins]() {
      vector<Bin> result;
      for (const Bin & bin : all_bins) {
        if (!bin.bad()) {
          result.push_back(bin);
        }
      }
      return result;
    }()};

  // Bin length sanity check
  for (const CN_Bins & result : profiles) {
    if (result.size() != bins.size()) {
      throw Error("Bin size mismatch");
    }
  }

  const std::vector<unsigned int> & chromosomes{cn_abspos.chromosomes()};
  vector<vector<double>> chromosome_totals(
      chromosomes.size(), vector<double>(profiles.size()));

  const vector<unsigned int> bins_per_chromosome{[&bins]() {
      vector<unsigned int> result;
      unsigned int last_chromosome{1000};
      for (const Bin & bin : bins) {
        if (bin.chromosome() != last_chromosome) {
          result.push_back(0);
          last_chromosome = bin.chromosome();
        }
        ++result.back();
      }
      return result;
    }()};
  if (bins_per_chromosome.size() != chromosomes.size()) {
    throw Error("Chromosome size mismatch");
  }

  unsigned int last_chromosome{bins[0].chromosome()};
  unsigned int chr{0};
  for (unsigned int b{0}; b != bins.size(); ++b) {
    const Bin & bin{bins[b]};
    if (bin.chromosome() != last_chromosome) {
      last_chromosome = bin.chromosome();
      ++chr;
    }
    for (unsigned int s{0}; s != profiles.size(); ++s) {
      const CN_Bin & result_bin{profiles[s][b]};
      chromosome_totals[chr][s] += result_bin.ratio();
    }
  }
  for (unsigned int c{0}; c != chromosomes.size(); ++c) {
    for (unsigned int s{0}; s != profiles.size(); ++s) {
      chromosome_totals[c][s] /= bins_per_chromosome[c];
    }
  }

  const double bad_dist{0.01};

  PSDoc ps{"missing_chromosome", "missing_chromosome"};
  std::vector<std::unique_ptr<PSGraph>> graphs;
  std::vector<std::unique_ptr<PSHSeries<double, unsigned int>>> series;
  const std::string dark{"0 0 0"};
  const Marker dark_circle_marker{paa::circle(), 0.5, dark, 0.2, true};

  for (unsigned int c{0}; c != chromosomes.size(); ++c) {
    const unsigned int chromosome{chromosomes[c]};
    const string name{ref.name(chromosome)};
    const string title{string("Chromosome ") + name};
    graphs.push_back(make_unique<PSGraph>(
        ps, title + ";Average Ratio;N", Bounds(0.9, 1.1)));
    series.push_back(make_unique<PSHSeries<double, unsigned int> >(
        *graphs.back(), 200, "1 0 0", false));
    PSHSeries<double, unsigned int> & hist{*series.back()};

    for (unsigned int s{0}; s != profiles.size(); ++s) {
      hist.add_point(chromosome_totals[c][s]);
    }

    if (name != "chrX" && name != "chrY") {
      for (unsigned int s{0}; s != profiles.size(); ++s) {
        if (fabs(chromosome_totals[c][s] - 1) > bad_dist) {
          cout << argv[s] << " " << name << " " << chromosome_totals[c][s]
               << endl;
        }
      }
    }
  }

  return 0;
} catch (Error & e) {
  cerr << "paa::Error:" << endl;
  cerr << e.what() << endl;
  return 1;
} catch (exception & e) {
  cerr << "std::exception" << endl;
  cerr << e.what() << endl;
  return 1;
} catch (...) {
  cerr << "unknown exception was caught" << endl;
  return 1;
}
```
