## Supplementary material for "G-Graph: An interactive genomic graph viewer": MUMdex Source Code for G-Graph: mumdex_examples.cpp

```
//
// mumdex_examples
//
// examples of mumdex usage
//
// Copyright 2015 Peter Andrews @ CSHL
//

#include <exception>
#include <iostream>
#include <string>

#include "error.h"
#include "mumdex.h"
#include "utility.h"

using std::cout;
using std::cerr;
using std::endl;
using std::exception;
using std::string;

using paa::Error;
using paa::MUMdex;
using paa::sout;
using paa::tout;

int main(int argc, char * argv[]) try {
  if (--argc != 1) throw Error("usage: mumdex_examples mumdex_name");

  const string mumdex_name{argv[1]};
  const MUMdex mumdex{mumdex_name};
  const auto & ref = mumdex.reference();

  const unsigned int max_show = 20;

  sout << "Lists are capped at" << max_show << "lines for clarity" << endl;

  cout << endl << "Loop over mums in pair order." << endl;
  tout << "chr" << "pos" << "read2" << "flipped"
       << "offset" << "length" << "last" << "touches_end" << endl;
  unsigned int n = 0;
  for (auto mum = mumdex.mums().begin(); mum != mumdex.mums().end(); ++mum) {
    tout << ref.name(mum->chromosome()) << mum->position1()
         << mum->read_2() << mum->flipped() << mum->offset() << mum->length()
         << mum->last_hit() << mum->touches_end() << endl;
    if (++n == max_show) break;
  }
  cout << endl;

  cout << "Pair and mum access in pair order" << endl;
  tout << "len_r1" << "len_r2" << "bad_r1" << "bad_r2"
       << "dupe" << "mums?"
       << "chr" << "mumpos" << "readpos" << "read2" << "flipped"
       << "offset" << "length" << "last" << "touches_end" << endl;
  n = 0;
  for (auto pair : mumdex) {
    if (pair.has_mums()) {
      tout << pair.read_1_length() << pair.read_2_length()
           << pair.read_1_bad() << pair.read_2_bad()
           << pair.dupe() << pair.has_mums() << endl;
      for (uint64_t m = pair.mums_start();; ++m) {
        const auto mum = mumdex.mum(m);  // mumdex.mums()[m] also works
        tout << "" << "" << "" << "" << "" << ""
             << ref.name(mum.chromosome()) << mum.position1()
             << mum.read_position1(pair.length(mum.read_2()))
             << mum.read_2() << mum.flipped() << mum.offset()
             << mum.length() << mum.last_hit() << mum.touches_end() << endl;
        if (++n == max_show) break;
        if (mum.last_hit()) break;
      }
      if (n == max_show) break;
    }
  }
  cout << endl;

  // Get [start, stop) position from index of mums so is almost guaranteed to
  // produce output for any sufficiently large mumdex input. In practice
  // desired positions are chosen using other methods like command line input
  const auto n_mums = mumdex.n_mums();
  if (n_mums < 4) throw Error("Expected 4 or more mums for this to work.");
  const auto start_mum = mumdex.mum(mumdex.index()[n_mums * 2 / 4]);
  const auto start_chromosome = start_mum.chromosome();
  const auto start_position = start_mum.position0();  // zero based pos!
  const auto stop_mum = mumdex.mum(mumdex.index()[n_mums * 3 / 4]);
  const auto stop_chromosome = stop_mum.chromosome();
  const auto stop_position = stop_mum.position0();

  cout << "Mum and pair access over a range ordered by mum position." << endl;
  const auto begin_index = mumdex.lower_bound(start_chromosome, start_position);
  const auto end_index = mumdex.lower_bound(stop_chromosome, stop_position);
  if (begin_index == end_index)
    throw Error("begin and end indexes are the same! No mums to show.");
  sout << "Displaying from" << ref.name(start_chromosome) << start_position + 1
       << "to" << ref.name(stop_chromosome) << stop_position + 1 << endl;
  tout << "pair" << "mum"
       << "chr" << "mumpos" << "readpos" << "read2" << "flipped"
       << "offset" << "length" << "last" << "dupe"
       << "readlen" << "touches_end" << endl;
  n = 0;
  for (auto index = begin_index; index != end_index; ++index) {
    const auto pair = mumdex.pair(*index);
    const auto mum = mumdex.mum(*index);
    tout << index->pair_index() << index->mum_in_pair_index()
         << ref.name(mum.chromosome()) << mum.position1()
         << mum.read_position1(pair.length(mum.read_2()))
         << mum.read_2() << mum.flipped() << mum.offset() << mum.length()
         << mum.last_hit() << pair.dupe()
         << pair.length(mum.read_2()) << mum.touches_end() << endl;
    if (++n == max_show) break;
  }
  cout << endl;

  return 0;
} catch (Error & e) {
  cerr << "paa::Error:" << endl;
  cerr << e.what() << endl;
  return 1;
} catch (exception & e) {
  cerr << "std::exception" << endl;
  cerr << e.what() << endl;
  return 1;
} catch (...) {
  cerr << "unknown exception was caught" << endl;
  return 1;
}
```
