## Supplementary material for "G-Graph: An interactive genomic graph viewer": MUMdex Source Code for G-Graph: offset_beds.cpp

```
//
// offset_beds.cpp
//
// Intended to display tracks over a read
//
// Copyright 2019 Peter Andrews @ CSHL
//

#include <fstream>
#include <iostream>
#include <string>
#include <vector>
#include <climits>

#include "error.h"
#include "mumdex.h"
#include "utility.h"

using std::cerr;
using std::cin;
using std::cout;
using std::endl;
using std::exception;
using std::ifstream;
using std::string;
using std::vector;

using paa::ChromosomeIndexLookup;
using paa::Error;
using paa::Reference;

int main(int argc, char ** argv) try {
  if (--argc < 6)
    throw Error("usage: offset_beds ref n_bases [bed char chr offset] ...");

  const Reference ref{argv[1]};
  const ChromosomeIndexLookup chr_lookup{ref};
  const unsigned int n_bases{static_cast<unsigned int>(atoi(argv[2]))};
  argc -= 2;
  argv += 2;

  while (argc) {
    const string bed_name{argv[1]};
    ifstream bed{bed_name.c_str()};
    if (!bed) throw Error("Problem opening bed file") << bed_name;
    const char id{argv[2][0]};
    const string chr_name{argv[3]};
    const unsigned int offset{static_cast<unsigned int>(atoi(argv[4]))};
    string output(n_bases, '.');
    string bed_chr;
    unsigned int bed_start;
    unsigned int bed_stop;
    while (bed >> bed_chr >> bed_start >> bed_stop) {
      if (bed_chr != chr_name) continue;
      if (bed_start > offset + n_bases) continue;
      if (bed_stop < offset) continue;
      for (unsigned int i{bed_start}; i != bed_stop; ++i) {
        if (i < offset) continue;
        if (i >= offset + n_bases) break;
        output[i - offset] = id;
      }
    }
    if (0) cerr << bed_name << " " << id << " " << chr_name << " " << offset
                << " " << output.size() << endl;
    cout << output << endl;
    argc -= 4;
    argv += 4;
  }

  return 0;
} catch (Error & e) {
  cerr << "paa::Error:" << endl;
  cerr << e.what() << endl;
  return 1;
} catch(exception & e) {
  cerr << e.what() << endl;
  return 1;
} catch(...) {
  cerr << "Some exception was caught." << endl;
  return 1;
}
```
