## Supplementary material for "G-Graph: An interactive genomic graph viewer": MUMdex Source Code for G-Graph: optional_test.cpp

```
//
// optional_test
//
// test encoding of mumdex optional information passthrough
//
// Copyright 2015 Peter Andrews @ CSHL
//

#include <array>
#include <exception>
#include <iostream>
#include <string>
#include <vector>

#include "encode.h"
#include "error.h"
#include "mumdex.h"

using std::array;
using std::cout;
using std::cerr;
using std::endl;
using std::exception;
using std::string;
using std::vector;

using paa::Error;
using paa::MUMdex;
using paa::OptionalSavers;
using paa::read_optional_formats;

int main(int argc, char * argv[]) try {
  paa::exit_on_pipe_close();

  if (--argc != 1) throw Error("usage: optional_test mumdex_name");
  const string mumdex_name{argv[1]};

  const MUMdex mumdex{mumdex_name};
  const vector<string> optional_formats{read_optional_formats(mumdex_name)};
  OptionalSavers saver{optional_formats};
  saver.load(mumdex_name, mumdex.n_pairs() * 2);

  for (uint64_t i = 0; i != mumdex.n_pairs(); ++i) {
    const array<string, 2> sequences(mumdex.sequences(i));
    for (const bool r : {false, true}) {
      const auto & sequence = sequences[r];
      cout << sequence;
      for (unsigned int s = 0; s != saver.size(); ++s) {
        cout << " " << saver[s][i * 2 + r];
      }
      cout << endl;
    }
  }

  return 0;
} catch (Error & e) {
  cerr << "paa::Error:" << endl;
  cerr << e.what() << endl;
  return 1;
} catch (exception & e) {
  cerr << "std::exception" << endl;
  cerr << e.what() << endl;
  return 1;
} catch (...) {
  cerr << "unknown exception was caught" << endl;
  return 1;
}
```
