## Supplementary material for "G-Graph: An interactive genomic graph viewer": MUMdex Source Code for G-Graph: pair_view.cpp

```
//
// pair_view
//
// a text view of pairs showing mums that map
//
// Copyright 2015 Peter Andrews @ CSHL
//

#include <exception>
#include <iostream>
#include <string>

#include "error.h"
#include "mumdex.h"
#include "utility.h"

using std::cout;
using std::cerr;
using std::endl;
using std::exception;
using std::string;

using paa::Error;
using paa::UnMappedMUMdex;

int main(int argc, char * argv[]) try {
  paa::exit_on_pipe_close();
  if (--argc != 1) throw Error("usage: pair_view mumdex_name");

  const string mumdex_name{argv[1]};
  const UnMappedMUMdex mumdex{mumdex_name};

  for (uint64_t pair_index = 0; pair_index != mumdex.n_pairs(); ++pair_index) {
    mumdex.pair_view(cout, pair_index);
    cout << endl;
  }

  return 0;
} catch (Error & e) {
  cerr << "paa::Error:" << endl;
  cerr << e.what() << endl;
  return 1;
} catch (exception & e) {
  cerr << "std::exception" << endl;
  cerr << e.what() << endl;
  return 1;
} catch (...) {
  cerr << "unknown exception was caught" << endl;
  return 1;
}
```
