## Supplementary material for "G-Graph: An interactive genomic graph viewer": MUMdex Source Code for G-Graph: parallelize.sh

```
#! /bin/bash

n_threads=$1

if [ "$n_threads" = '' ] ; then
    n_threads=2
fi

command_name=$2

if [ "$command_name" != '' ] ; then
    # Gather job info
    while true ; do
        sleep 10
        ps aux | grep $command_name | grep $USER | grep -v -e grep -e ssh -e parallelize > temp.commands
        n_running=$(cat temp.commands | wc -l)
        if [ $n_running = 0 ] ; then
            rm temp.commands
            break
        else
            echo
            cat temp.commands
        fi
    done &
fi

# implements a process pool!
xargs -I CMD -P $n_threads bash -c CMD

wait

exit 0

running=()
n=0
waitn=-1
while read line ; do
    echo run $n
    $line &
    running[n]=$!
    n=$((n+1))
    if [ $n -ge $n_threads ] ; then
        waitn=$((n-$n_threads))
        echo wait $waitn
        wait ${running[waitn]}
    fi 
    /bin/true # avoids a bash bug!
done

waitn=$((waitn + 1))
while [ $waitn -lt $n ] ; do
        echo end wait $waitn
        wait ${running[waitn]}
        waitn=$((waitn + 1))
done
```
