## Supplementary material for "G-Graph: An interactive genomic graph viewer": MUMdex Source Code for G-Graph: pie.cpp

```
//
// pie.cpp
//
// Make a postscript (or pdf) pie chart
//
// Copyright 2019 Peter Andrews @ CSHL
//

#include <algorithm>
#include <fstream>
#include <iostream>
#include <memory>
#include <sstream>
#include <string>
#include <utility>
#include <vector>

#include "error.h"
#include "utility.h"

using std::cerr;
using std::cin;
using std::cout;
using std::endl;
using std::exception;
using std::ifstream;
using std::istream;
using std::istringstream;
using std::make_unique;
using std::max;
using std::min;
using std::ofstream;
using std::ostringstream;
using std::pair;
using std::string;
using std::unique_ptr;
using std::vector;

using paa::Error;

class PSPie {
 public:
  using Item = pair<double, string>;
  PSPie(const std::string title__, istream & input) :
      title{title__} {
        double value;
        string description;
        while (input >> value) {
          input.get();
          getline(input, description);
          if (!input) throw Error("Problem parsing input");
          if (value <= 0) continue;
          add_item(value, description);
        }
      }
  void add_item(const double value, const std::string & description) {
    items.emplace_back(value, description);
    total += value;
    max_length = max(max_length, description.size());
  }
  void output(const std::string output_prefix,
              bool pdf = true, bool png = true) const {
    const string output_name{output_prefix + ".ps"};
    ofstream outputf{output_name.c_str()};
    if (!outputf) throw Error("Could not open output file") << output_name;
    const double width{792};
    const double height{612};
    const double radius{0.3 * height};
    const double min_edge{width / 2 - radius};
    const double auto_font{min(40.0, 2 * min_edge / max_length)};
    outputf << "%!PS-Adobe-3.0 EPSF-3.0\n"
           << "%%BoundingBox: 0 0 " << width << " " << height << "\n";
    const double light_gray{0.9375};
    outputf << "gsave " << light_gray << " "
           << light_gray << " " << light_gray
           << " setrgbcolor newpath 0 0 moveto " << width << " 0 rlineto "
           << "0 " << height << " rlineto " << -width << " 0 rlineto closepath "
           << " fill stroke grestore\n";  // does nothing????
    outputf << "2 setlinewidth\n";
    double actual_height{height};
    if (title.size()) {
      const double title_font{50};
      actual_height -= title_font * 1.5;
      outputf << "/Helvetica findfont " << title_font << " scalefont setfont\n";
      outputf << width / 2 << " " << height - 1.25 * title_font
             << " moveto (" << title
             << ") dup stringwidth pop 2 div neg 0 rmoveto show\n";
    }
    const double pie_center_height{actual_height / 2.0};
    outputf << "/Helvetica findfont " << auto_font << " scalefont setfont\n";

    const vector<string> colors{[]() {
        const vector<string> test_colors{
          "0.898039 0 0", "0.145098 0 0.619608",
              "0 0.717647 0", "0.898039 0.745098 0",
              "0.0235294 0.337255 0.576471", "0.717647 0.866667 0",
              "0.898039 0.513725 0", "0.584314 0 0.584314",
              "0.972549 0.470588 0.972549", "0 0.0941176 0",
              "0 0.941176 0.533333", "0.564706 0.627451 0.533333",
              "0.972549 0.972549 0.627451", "0 0.658824 0.972549",
              "0.439216 0.313725 0.972549", "0.972549 0.0313725 0.972549",
              "0.470588 0.282353 0.188235", "0.972549 0.25098 0.470588",
              "0.470588 0.972549 0.376471", "0.470588 0.972549 0.972549"};
        vector<string> result;
        for (const string & color : test_colors) {
          istringstream color_stream{color.c_str()};
          double value;
          double total_color{0.0};
          while (color_stream >> value) {
            total_color += value;
          }
          if (total_color > 1) result.push_back(color);
        }
        return result;
      }()};
    double start_angle{0};
    for (uint64_t i{0}; i != items.size(); ++i) {
      const Item & item{items[i]};
      const double value{item.first};
      const string & description{item.second};
      const double percent{round(10000 * value / total)/100};
      const double wedge{value / total * 360};
      const double stop_angle{start_angle + wedge};
      const double center_angle{(start_angle + stop_angle) / 2};
      const double small_angle{0.3};
      cout << value << '\t' << description << '\t' << percent << endl;
      outputf << "newpath "
             << width  << " 2 div " << pie_center_height << " moveto "
             << width  << " 2 div " << pie_center_height << " "
             << radius << " " << start_angle << " "
             << stop_angle << " arc "
             << width  << " 2 div " << pie_center_height << " lineto closepath "
             << "gsave " << colors[i%colors.size()]
             << " setrgbcolor fill grestore stroke\n"
             << "newpath "
             << width  << " 2 div " << pie_center_height << " "
             << radius * 1.05 << " " << center_angle - small_angle << " "
             << center_angle + small_angle << " arc "
             << "(" << description << ")";
      if (center_angle > 90 && center_angle < 270) {
        outputf << " dup stringwidth pop neg 3 sub 0 rmoveto";
      }
      if (center_angle > 180) {
        outputf << " 0 -" << 0.7 * ceil(auto_font) << " rmoveto";
      }
      outputf << " show stroke\n";
      start_angle = stop_angle;
    }
    outputf.close();
    if (pdf || png) {
      ostringstream command;
      command << "ps2pdf -dDEVICEWIDTHPOINTS=" << width
              << " -dDEVICEHEIGHTPOINTS=" << height
              << " " << output_name
              << " " << output_prefix << ".pdf";
      if (png) command << " && convert " << output_prefix << ".pdf "
                       << output_prefix << ".png";
      if (!pdf) command << " && rm -f " << output_prefix << ".pdf ";
      const int result{system(command.str().c_str())};
      if (0) cerr << result;
    }
  }

 private:
  std::string title;
  double total{0};
  size_t max_length{0};
  std::vector<Item> items{};
};

int main(int argc, char ** argv) try {
  --argc;
  const string usage{
    "usage: pie [-pdf] [-png] [-t title] [-o out_prefix] [input]"};
  if (argc > 6) throw Error(usage);
  string title{""};
  bool pdf{false};
  bool png{false};
  string output_prefix{"pie"};
  while (argc) {
    if (argc >= 2 && string(argv[1]) == "-t") {
      title = argv[2];
      argc -= 2;
      argv += 2;
    } else if (argc >= 2 && string(argv[1]) == "-o") {
      output_prefix = argv[2];
      argc -= 2;
      argv += 2;
    } else if (argc >= 1 && string(argv[1]) == "-png") {
      --argc;
      ++argv;
      png = true;
    } else if (argc >= 1 && string(argv[1]) == "-pdf") {
      --argc;
      ++argv;
      pdf = true;
    } else {
      break;
    }
  }
  if (argc > 1) throw Error(usage);
  istream * input{&cin};
  unique_ptr<ifstream> input_file;
  const string input_name{argc == 1 ? argv[1] : "cin"};
  if (input_name != "cin") {
    input_file = make_unique<ifstream>(input_name.c_str());
    if (!*input_file) throw Error("Could not open input") << input_name;
    input = input_file.get();
  }

  const PSPie pie{title, *input};
  pie.output(output_prefix, pdf, png);

  return 0;
} catch (Error & e) {
  cerr << "paa::Error:" << endl;
  cerr << e.what() << endl;
  return 1;
} catch(exception & e) {
  cerr << e.what() << endl;
  return 1;
} catch(...) {
  cerr << "Some exception was caught." << endl;
  return 1;
}
```
