## Supplementary material for "G-Graph: An interactive genomic graph viewer": MUMdex Source Code for G-Graph: pivot_table.cpp

```
//
// pivit_table.cpp
//
// Copyright 2016 Peter Andrews @ CSHL
//

#include <algorithm>
#include <iostream>
#include <map>
#include <sstream>
#include <string>
#include <utility>
#include <vector>

#include "error.h"
#include "utility.h"

using std::cerr;
using std::cin;
using std::cout;
using std::endl;
using std::exception;
using std::istringstream;
using std::map;
using std::pair;
using std::string;
using std::vector;

using paa::Error;

vector<string> get_fields(const string & line) {
  istringstream in{line.c_str()};
  vector<string> fields;
  string field;
  while (in >> field) {
    fields.push_back(field);
  }
  return fields;
}

vector<unsigned int> field_lookup(const vector<string> & header,
                                  const vector<string> & names,
                                  const bool ignore_missing = false) {
  vector<unsigned int> fields;
  for (const string & name : names) {
    const auto found = find(header.begin(), header.end(), name);
    if (found == header.end()) {
      if (!ignore_missing) {
        throw Error("Could not find pivot in header") << name << header.size();
      }
    } else {
      fields.push_back(static_cast<unsigned int>(found - header.begin()));
    }
  }
  return fields;
}

bool is_field_numeric(const string & field) {
  try {
    stod(field);
    return true;
  } catch (...) {
    return false;
  }

  istringstream in{field.c_str()};
  double val;
  in >> val;
  if (!in) return false;
  string excess;
  in >> excess;
  if (excess != "") return false;
  return true;
}

int main(int argc, char ** argv) {
  try {
    paa::exit_on_pipe_close();
    if (--argc > 2) throw Error("usage: pivot_table [pivots] [show]");

    // Header
    const string header_line{[]() {
        string line;
        getline(cin, line);
        return line;
      }()};
    const vector<string> header{get_fields(header_line)};

    // Pivots
    const string pivots_line(argc >= 1 && string() != argv[1] ?
                             argv[1] : header_line);
    const vector<string> pivots{get_fields(pivots_line)};
    // pivot index -> header column index
    const vector<unsigned int> pivot_fields{field_lookup(header, pivots)};

    // Columns to show
    const string show_cols_line(argc >= 2 && string() != argv[2] ?
                                argv[2] : header_line);
    const vector<string> show_cols{get_fields(show_cols_line)};
    // show index -> header column index
    const vector<unsigned int> show_fields{field_lookup(header, show_cols)};
    const vector<unsigned int> show_pivot_fields{
      field_lookup(show_cols, pivots, true)};

    // Counts or values for columns
    vector<bool> is_numeric(header.size());
    vector<unsigned int> column_indices;
    using Doubles = vector<double>;
    using Numbers = pair<uint64_t, Doubles>;
    using Strings = vector<string>;
    using Info = pair<Strings, Numbers>;
    vector<map<string, Info>> all_values{pivots.size()};

    // Read and process data
    string line;
    uint64_t n{0};
    unsigned int n_numeric{0};
    unsigned int n_string{0};
    while (getline(cin, line)) {
      const vector<string> fields{get_fields(line)};
      if (fields.size() != header.size()) {
        throw Error("Fields size does not equal header size");
      }

      // Figure out if columns are numeric or not based on first line only
      if (++n == 1) {
        for (unsigned int f{0}; f != show_fields.size(); ++f) {
          const string & field{fields[show_fields[f]]};
          if (is_field_numeric(field)) {
            is_numeric[f] = true;
            column_indices.push_back(n_numeric++);
          } else {
            is_numeric[f] = false;
            column_indices.push_back(n_string++);
          }
        }
      }

      for (unsigned int pf{0}; pf != pivots.size(); ++pf) {
         map<string, Info> & values{all_values[pf]};
         const string & key{fields[pivot_fields[pf]]};
         map<string, Info>::iterator found{values.lower_bound(key)};
         Info & to_update{
           ((found != values.end() && found->first == key) ?
            found :
            values.insert(found, pair<string, Info>(
                key, Info(Strings(n_string),
                          Numbers(0, Doubles(n_numeric))))))->second};

         Strings & strs{to_update.first};
         Numbers & nums{to_update.second};
         ++nums.first;
         for (unsigned int f{0}; f != show_fields.size(); ++f) {
           const unsigned int i{column_indices[f]};
           const string & field{fields[show_fields[f]]};
           if (is_numeric[f]) {
             nums.second[i] += stod(field);
           } else {
             if (strs[i].empty()) {
               strs[i] = field;
             } else if (strs[i] != field) {
               strs[i] = "-";
             }
           }
         }
      }
    }

    for (unsigned int pf{0}; pf != pivots.size(); ++pf) {
      map<string, Info> & values{all_values[pf]};
      if (pf) cout << endl;
      cout << "p_" << pivots[pf] << endl;

      // Header line
      cout << "count frac";
      for (const string & name : show_cols) {
        cout << " " << name;
      }
      cout << endl;

      vector<pair<string, Info>> field_info(values.begin(), values.end());
      if (is_numeric[show_pivot_fields[pf]]) {
        sort(field_info.begin(), field_info.end(), []
             (const pair<string, Info> & lhs, const pair<string, Info> & rhs) {
               return stod(lhs.first) < stod(rhs.first);
             });
      } else {
        sort(field_info.begin(), field_info.end(), []
             (const pair<string, Info> & lhs, const pair<string, Info> & rhs) {
               return lhs.second.second.first > rhs.second.second.first;
             });
      }

      for (const pair<string, Info> & sinfo : field_info) {
        const Info & info{sinfo.second};
        const Strings & strs{info.first};
        const Numbers & nums{info.second};
        cout << nums.first << " " << 1.0 * nums.first / n;
        for (unsigned int f{0}; f != show_fields.size(); ++f) {
          const unsigned int i{column_indices[f]};
          cout << " ";
          if (is_numeric[f]) {
            cout << nums.second[i] / nums.first;
          } else {
            cout << strs[i];
          }
        }
        cout << endl;
      }
    }

    return 0;
  }
  catch(exception & e) {
    cerr << e.what() << endl;
    return 1;
  }
  catch(...) {
    cerr << "Some exception was caught." << endl;
    return 1;
  }
  return 0;
}
```
