## Supplementary material for "G-Graph: An interactive genomic graph viewer": MUMdex Source Code for G-Graph: plot.cpp

```
//
// plot.cpp
//
// plot series from a text file with format:
//
// x_name y1_name y2_name
// data...
//
// Copyright 2016 Peter Andrews @ CSHL
//

#include <iostream>
#include <fstream>
#include <memory>
#include <sstream>
#include <string>
#include <vector>

#include "error.h"
#include "psplot.h"
#include "utility.h"

using std::cout;
using std::cerr;
using std::endl;
using std::ifstream;
using std::istringstream;
using std::string;
using std::vector;

using paa::sout;
using paa::Bounds;
using paa::Error;
using paa::Marker;
using paa::PSDoc;
using paa::PSPage;
using paa::PSGraph;
using paa::PSXYSeries;

int main(int argc, char ** argv)  try {
  // Check arguments
  --argc;
  if (argc != 2 && argc != 6)
    throw Error("usage: plot data_file title;x;y [xlow xhigh ylow yhigh]");

  // Interpret arguments
  const string input_name{argv[1]};
  ifstream input{input_name};
  if (!input) throw Error("Problem opening data file") << input_name;
  const string title{argv[2]};
  const double x_low{argc == 6 ? strtod(argv[3], nullptr) : paa::unset()};
  const double x_high{argc == 6 ? strtod(argv[4], nullptr) : paa::nunset()};
  const double y_low{argc == 6 ? strtod(argv[5], nullptr) : paa::unset()};
  const double y_high{argc == 6 ? strtod(argv[6], nullptr) : paa::nunset()};

  // Graph
  PSDoc ps{input_name, input_name};
  ps.pdf(false);
  PSGraph graph{ps, title, Bounds{x_low, x_high, y_low, y_high}};
  graph.ps(  // "np 0 0 gfc m 1 0 gfc l 1 1 gfc l 1 0 gfc l cp clip "
           "0 0 0 c 0.5 lw np 0 0 gc m 4 4 gc l sp");
  // graph.log_x(true).log_y(true);
  // graph.log_x(true);
  vector<PSXYSeries> series;
  series.reserve(100);

  // Default markers
  const vector<string> colors{"1 0 0", "0 0 1", "0 1 0"};
  const vector<string (*)()> shapes{paa::circle, paa::square, paa::triangle,
        paa::diamond, paa::plus, paa::xshape, paa::star};
  vector<Marker> markers;

  // Series names
  vector<string> series_names;
  string header;
  getline(input, header);
  istringstream header_stream{header.c_str()};
  header_stream >> header;
  unsigned int color{0};
  unsigned int shape{0};
  while (header_stream >> header) {
    series_names.push_back(header);
    series.emplace_back(graph, Marker{shapes[(shape++) % shapes.size()](),
            0.6, colors[(color++) % colors.size()], 1.0,
            false, "1 1 1"});  // , header);
  }

  // Read data
  double x;
  double y;
  while (input) {
    input >> x;
    if (!input) break;
    for (unsigned int s{0}; s != series.size(); ++s) {
      input >> y;
      // cerr << "add " << s << " " << x << " " << y << endl;
      series[s].add_point(x, y);
    }

    if (!input) throw Error("Problem parsing input");
  }

  return 0;
} catch (Error & e) {
  cerr << "paa::Error:" << endl;
  cerr << e.what() << endl;
  return 1;
} catch(std::exception & e) {
  cerr << e.what() << '\n';
  return 1;
} catch(...) {
  cerr << "Some exception was caught.\n";
  return 1;
}
```
