## Supplementary material for "G-Graph: An interactive genomic graph viewer": MUMdex Source Code for G-Graph: plot_chr_panels.cpp

```
//
// plot_panels
//
// plot a series in several panels
//
// Copyright 2019 Peter Andrews @ CSHL
//

#include <algorithm>
#include <fstream>
#include <iostream>
#include <memory>
#include <random>
#include <sstream>
#include <string>
#include <utility>
#include <vector>
#include <climits>

#include "cn.h"
#include "error.h"
#include "mumdex.h"
#include "psplot.h"
#include "strings.h"
#include "utility.h"

using std::cerr;
using std::cin;
using std::cout;
using std::endl;
using std::exception;
using std::ifstream;
using std::istream;
using std::make_unique;
using std::min;
using std::ostringstream;
using std::pair;
using std::string;
using std::to_string;
using std::unique_ptr;
using std::vector;

using paa::replace_substring;
using paa::CN_abspos;
using paa::Error;
using paa::Marker;
using paa::PSDoc;
using paa::PSGraph;
using paa::PSPage;
using paa::PSXYSeries;
using paa::Reference;

int main(int argc, char ** argv) try {
  paa::exit_on_pipe_close();
  std::ios_base::sync_with_stdio(false);
  cin.tie(nullptr);

  const string usage{"usage: plot_panels [-s scatter] "
        "ref name title n_panels [input_files ...]"};

  // process optional arguments
  --argc;
  double scatter_amount{0.0};
  bool scatter{false};
  while (argc) {
    bool acted{false};
    if (argc > 1 && argv[1] == string("-s")) {
      scatter_amount = atof(argv[2]);
      scatter = true;
      argc -= 2;
      argv += 2;
      acted = true;
    }
    if (!acted) break;
  }

  if (argc < 4) throw Error("Not enough arguments:") << usage;
  const Reference ref{argv[1]};
  const CN_abspos cn_abspos{ref};
  const string out_name{argv[2]};
  const string title{argv[3]};
  const uint64_t n_panels{static_cast<uint64_t>(atoi(argv[4]))};
  argc -= 4;
  argv += 4;

  vector<unique_ptr<ifstream>> input_files;
  vector<istream *> inputs;
  if (argc) {
    while (argc--) {
      const string file_name{(argv++)[1]};
      input_files.push_back(make_unique<ifstream>(file_name.c_str()));
      if (!*input_files.back())
        throw Error("Could not open file for input") << file_name;
      inputs.push_back(input_files.back().get());
    }
  } else {
    inputs.push_back(&cin);
  }

  vector<pair<double, double>> xy;
  {
    double x;
    double y;
    for (istream * in : inputs)
      while ((*in) >> x >> y)
        xy.emplace_back(x, y);
  }

  if (xy.size() < 2) throw Error("Not enough points");

  sort(xy.begin(), xy.end());
  const double min_x{xy.front().first};
  const double max_x{xy.back().first};
  const double width{max_x - min_x};
  if (min_x >= max_x) throw Error("min equals max");

  PSDoc plots{out_name};
  plots.pdf(true);
  PSPage plot_page{plots, "", "1 " + to_string(n_panels)};
  vector<unique_ptr<PSGraph>> graphs;
  vector<unique_ptr<PSXYSeries>> series;
  const Marker marker{paa::circle(), 0.1, "0 0.8 0", 0.1, true, "0 0.8 0"};
  for (uint64_t p{0}; p != n_panels; ++p) {
    graphs.push_back(make_unique<PSGraph>(plot_page, title));
    graphs.back()->do_ticks(false);
    series.push_back(make_unique<PSXYSeries>(*graphs.back(), marker));
    ostringstream ps;
    ps << "2 lw 0 0 0 c 12 sf\n";
    const double min_pos{p * width / n_panels};
    const double max_pos{(p + 1) * width / n_panels};

    for (unsigned int c{0}; c != cn_abspos.chromosomes().size(); ++c) {
      const unsigned int chr{cn_abspos.chromosomes()[c]};
      if (ref.name(chr).find("X") != string::npos) break;
      const unsigned int offset{cn_abspos.offsets()[c]};
      if (offset < min_pos) continue;
      if (offset > max_pos) break;
      ps << "np " << offset << " xc 0 yfc m " << offset << " xc 1 yfc l sp "
         << offset << " xc 1 yfc m 5 -12 rm ("
         << replace_substring(ref.name(chr), "chr", "") << ") s\n";
    }
    graphs.back()->ps(ps.str());
  }
  std::random_device rd{};
  std::mt19937_64 mersenne{rd()};
  std::function<double()> gen{
    std::bind(std::uniform_real_distribution<double>(
        -scatter_amount, scatter_amount), mersenne)};

  for (uint64_t p{0}; p != xy.size(); ++p) {
    const double x{xy[p].first};
    const double y{xy[p].second};
    const uint64_t panel{static_cast<uint64_t>(
        min(1.0 * n_panels - 1, n_panels * (x - min_x) / width))};
    const double scatter_y{(scatter ? gen() : 0.0) + y};
    series[panel]->add_point(x, scatter_y);
  }

  return 0;
} catch (Error & e) {
  cerr << "paa::Error:" << endl;
  cerr << e.what() << endl;
  return 1;
} catch(exception & e) {
  cerr << e.what() << endl;
  return 1;
}
catch(...) {
  cerr << "Some exception was caught." << endl;
  return 1;
}
```
