## Supplementary material for "G-Graph: An interactive genomic graph viewer": MUMdex Source Code for G-Graph: plot_cn_stats.cpp

```
//
// plot_cn_stats.cpp
//
// Copyright 2017 Peter Andrews @ CSHL
//

#include <algorithm>
#include <iostream>
#include <fstream>
#include <functional>
#include <map>
#include <memory>
#include <random>
#include <sstream>
#include <string>
#include <vector>

#include "pstream.h"

#include "error.h"
#include "psplot.h"

using std::cout;
using std::cerr;
using std::endl;
using std::function;
using std::ifstream;
using std::make_unique;
using std::map;
using std::ostringstream;
using std::string;
using std::to_string;
using std::normal_distribution;
using std::unique_ptr;
using std::vector;

using paa::Bounds;
using paa::Error;
using paa::Marker;
using paa::PSDoc;
using paa::PSPage;
using paa::PSGraph;
using paa::PSHSeries;
using paa::PSSeries;
using paa::PSXYSeries;
using paa::PSXYMSeries;

int main(int argc, char ** argv)  try {
  // Check arguments
  --argc;
  const vector<string> colors{"0 1 0", "0 0 1", "1 1 0", "1 0 0", "0 1 1"};
  if (argc < 1 || argc > static_cast<int>(colors.size()))
    throw Error("usage: plot_cn_stats category ...");
  ++argv;

  PSDoc ps{"concordance", "concordance"};
  vector<unique_ptr<PSPage>> pages;
  vector<unique_ptr<PSGraph>> hist_graphs;
  using Hist = PSHSeries<double, unsigned int>;
  vector<unique_ptr<Hist>> hists;
  vector<unique_ptr<Hist>> seg_hists;
  vector<unique_ptr<PSGraph>> graphs;
  vector<unique_ptr<PSXYSeries>> series;
  map<string, unsigned int> graph_ids;
  vector<string> graph_names;
  map<string, map<unsigned int, unsigned int>> series_ids;
  vector<string> categories;
  vector<Marker> markers;
  for (int a{0}; a != argc; ++a) {
    const Marker marker{paa::circle(), 0.7, "0 0 0", 0.6, true, colors[a]};
    markers.push_back(marker);
    const string category{argv[a]};
    categories.push_back(category);
    const string file_name{category + ".txt"};
    ifstream in{file_name.c_str()};
    if (!in) throw Error("Problem opening file") << file_name;
    string dummy;
    string comp;
    double frac_segs;
    double frac_bins;
    vector<unsigned int> n(graphs.size());
    vector<unsigned int> n_pass(graphs.size());
    vector<unsigned int> n_fail(graphs.size());
    vector<unsigned int> n_zero(graphs.size());
    vector<unsigned int> n_one(graphs.size());
    double n_segs;
    while (in
           >> comp
           >> dummy >> dummy
           >> frac_segs >> frac_bins >> n_segs) {
      if (comp == "comp2" || comp == "comp4") continue;
      const string comp_desc{comp == "comp2" ? "Segment Call" :
            (comp == "comp3" ? "Averaged Segment Call" :
             (comp == "comp4" ? "Segment Call - Recall" :
              (comp == "comp5" ? "Averaged Segment Call - Recall" : "")))};
      if (!in) throw Error("parse error");
      if (comp == "comp") continue;
      if (graph_ids.count(comp) == 0) {
        graph_names.push_back(comp);
        graph_ids[comp] = static_cast<unsigned int>(graphs.size());
        pages.push_back(make_unique<PSPage>(
            ps, comp_desc + " Concordance",
            "1 " + to_string(argc)));
        graphs.push_back(make_unique<PSGraph>(
            ps, comp_desc + " Concordance; Percent Segments Concordant;"
            "Percent Bins Concordant"));
       n_pass.push_back(0);
        n_fail.push_back(0);
        n_zero.push_back(0);
        n_one.push_back(0);
        n.push_back(0);
      }
      const unsigned int graph_id{graph_ids[comp]};
      PSGraph & this_graph{*graphs[graph_id]};

      if (series_ids[comp].count(a) == 0) {
        series_ids[comp][a] = static_cast<unsigned int>(series.size());
        series.push_back(make_unique<PSXYSeries>(this_graph, marker));
        PSPage & this_page{*pages[graph_id]};
        hist_graphs.push_back(make_unique<PSGraph>(
            this_page, string(argv[a]) +
            (a + 1 == argc ? ";Percent Segments Concordant" : "")));
        hists.push_back(make_unique<Hist>(
            *hist_graphs.back(), 22, colors[a], false));
        hist_graphs.back()->range().x(0.0, 110);
      }
      const unsigned int series_id{series_ids[comp][a]};
      PSXYSeries & this_series{*series[series_id]};
      this_series.add_point(100* frac_segs, 100 * frac_bins);
      Hist & this_hist{*hists[series_id]};
      this_hist.add_point(frac_segs * 100);
      ++n[graph_id];
      if (frac_segs <= 0 && frac_bins <= 0) ++n_zero[graph_id];
      if (frac_segs > 0.6) ++n_pass[graph_id];
      if (frac_segs > 0.1 && frac_bins > 0.1) ++n_fail[graph_id];
      if (frac_segs >= 1.0) ++n_one[graph_id];
    }
    for (unsigned int i{0}; i != graphs.size(); ++i) {
      cerr << "Plot " << n[i] << " from " << file_name << " for "
           << graph_names[i] << " "
           << 1.0 * n_pass[i] / n[i] << " "
           << 1.0 * n_fail[i] / n[i] << " "
           << 1.0 * n_zero[i] / n[i] << " "
           << 1.0 * n_one[i] / n[i] << endl;
    }
  }
  ostringstream legend;
  const double x_pos{75};
  const double y_pos{35};
  const unsigned int font_size{12};
  legend << "np " << x_pos << " " << y_pos << " gc m ";
  for (unsigned int m{0}; m != categories.size(); ++m) {
    const Marker & marker{markers[m]};
    legend << 0 << " " << font_size << " neg rm gs cxy np m "
           << " gs cxy tr "
           << marker.setup_commands() << marker.draw_commands()
           << " sp gr "
           << 10 << " " << -3 << " rm "
           << " (" << categories[m] << ") s gr ";
  }
  for (unsigned int g{0}; g != graphs.size(); ++g) {
    graphs[g]->ps(legend.str());
  }

  return 0;
} catch (Error & e) {
  cerr << "paa::Error:" << endl;
  cerr << e.what() << endl;
  return 1;
} catch(std::exception & e) {
  cerr << e.what() << '\n';
  return 1;
} catch(...) {
  cerr << "Some exception was caught.\n";
  return 1;
}
```
