## Supplementary material for "G-Graph: An interactive genomic graph viewer": MUMdex Source Code for G-Graph: plot_coverage.cpp

```
//
// plot_coverage.cpp
//
// Copyright 2016 Peter Andrews @ CSHL
//

#include <algorithm>
#include <iostream>
#include <functional>
#include <memory>
#include <random>
#include <string>
#include <vector>

#include "pstream.h"

#include "bed.h"
#include "error.h"
#include "mumdex.h"
#include "psplot.h"
#include "utility.h"

using std::cout;
using std::cerr;
using std::endl;
using std::function;
using std::max;
using std::min;
using std::mt19937_64;
using std::random_device;
using std::string;
using std::to_string;
using std::normal_distribution;
using std::vector;

using redi::ipstream;

using paa::sout;
using paa::BedFile;
using paa::Bounds;
using paa::ChromosomeIndexLookup;
using paa::Error;
using paa::Mappability;
using paa::MUMdex;
using paa::PSDoc;
using paa::PSPage;
using paa::PSGraph;
// using paa::PSHist;
using paa::PSSeries;
using paa::PSXYSeries;
using paa::PSXYMSeries;
using paa::Reference;

int main(int argc, char ** argv)  try {
  // Check arguments
  if (--argc != 6)
    throw Error("usage: plot_coverage mumdex bed counts chr start stop");

  const string mumdex_name{argv[1]};
  const string bed_name{argv[2]};
  const string counts_dir{argv[3]};
  const string chr_name{argv[4]};
  const unsigned int start{static_cast<unsigned int>(atoi(argv[5]))};
  const unsigned int stop{static_cast<unsigned int>(atoi(argv[6]))};

  const MUMdex mumdex{mumdex_name};
  const Reference & ref{mumdex.reference()};
  const ChromosomeIndexLookup chr_lookup{ref};
  const unsigned int chromosome{chr_lookup[chr_name]};
  const Mappability map{ref};
  const BedFile bed{bed_name};
  const uint64_t bed_start{bed.find_bed_line(chr_name, start)};
  const uint64_t bed_stop{bed.find_bed_line(chr_name, stop) + 1};

  const unsigned int chr_offset{ref.offset(chromosome)};
  const unsigned int Lr{[&mumdex]() {
      const uint64_t check_pairs{10000};
      const uint64_t n_pairs{min(check_pairs, mumdex.n_pairs())};
      double result{0};
      for (uint64_t p{0}; p != n_pairs; ++p) {
        for (const bool read2 : { false, true }) {
          result += mumdex.pair(p).length(read2);
        }
      }
      return static_cast<unsigned int>(result / n_pairs / 2);
    }()};
  const double Ch{1.0 * mumdex.n_pairs() * Lr / ref.size()};

  PSDoc ps{"coverage", "coverage"};
  PSPage page1{ps, "", "1 2 =0 1 2 1="};
  PSGraph cover_graph{page1, "Coverage;Position;Coverage",
        Bounds(start - 1, stop, 0, 50)};
  PSXYSeries cover_series{cover_graph, paa::triangle()};
  PSXYSeries exp_cover_series{cover_graph, paa::diamond()};
  Bounds low_high_bounds(start, stop, 0, 30);
  PSGraph low_graph{page1, "Low Coverage;Position;Coverage", low_high_bounds};
  PSXYSeries low_series{low_graph};
  PSXYSeries exp_low_series{low_graph};
  PSGraph high_graph{page1, "High Coverage;Position;Coverage", low_high_bounds};
  PSXYSeries high_series{high_graph};
  PSXYSeries exp_high_series{high_graph};

  const string command{string("~/mumdex/show_counts_special") +
        " " + bed_name + " " + mumdex_name + " " + counts_dir + " " +
        to_string(bed_start) + " " + to_string(bed_stop)};
  // cout << command << endl;
  ipstream coverage{command};
  if (!coverage) throw Error("Problem running command") << command;
  string chr;
  unsigned int pos;
  string rb;
  unsigned int n;
  unsigned int cover;
  unsigned int low_ref;
  unsigned int high_ref;
  unsigned int low_anchor;
  unsigned int high_anchor;
  // sout << Lr << Ch << endl;
  while (coverage >> chr >> pos >> rb >> n
         >> cover
         >> low_ref >> low_anchor
         >> high_ref >> high_anchor) {
    coverage.ignore(10000, '\n');
    if (pos < start) continue;
    if (pos >= stop) break;
    double C{0};
    for (unsigned int p{pos};
         p != min(ref.size(chromosome), pos + Lr); ++p) {
      const unsigned int abspos{chr_offset + p};
      const double Lm{map.high(abspos) + 5.0};
      C += Lm <= Lr;
    }
    C *= 2 * Ch / Lr;
    double Clh[2];
    const unsigned int abspos{chr_offset + pos};
    for (const bool high : { false, true }) {
      const double Lm{map.low_high(high, abspos) + 5.0};
      Clh[high] = cover * max(0.0, (Lr - Lm - 1 - 1.0) / Lr);
    }
    if (0) {
      sout << chr << pos << cover << C
           << low_ref << Clh[0] << high_ref << Clh[1] << endl;
    }
    cover_series.add_point(pos, cover);
    exp_cover_series.add_point(pos, C);
    low_series.add_point(pos, low_ref);
    exp_low_series.add_point(pos, Clh[0]);
    high_series.add_point(pos, high_ref);
    exp_high_series.add_point(pos, Clh[1]);
    // sout << pos << high_ref << Clh[1] << endl;
  }

  std::cerr << "done" << endl;

  return 0;
} catch (Error & e) {
  cerr << "paa::Error:" << endl;
  cerr << e.what() << endl;
  return 1;
} catch(std::exception & e) {
  cerr << e.what() << '\n';
  return 1;
} catch(...) {
  cerr << "Some exception was caught.\n";
  return 1;
}
```
