## Supplementary material for "G-Graph: An interactive genomic graph viewer": MUMdex Source Code for G-Graph: plot_invariants.cpp

```
//
// plot_invariants
//
// plot invariants over a region for a population
//
// Copyright 2016 Peter Andrews @ CSHL
//

#include <algorithm>
#include <exception>
#include <functional>
#include <iostream>
#include <random>
#include <sstream>
#include <string>

#include "bridges.h"
#include "error.h"
#include "files.h"
#include "mumdex.h"
#include "psplot.h"
#include "utility.h"

using std::bind;
using std::cout;
using std::cerr;
using std::endl;
using std::exception;
using std::function;
using std::lower_bound;
using std::mt19937_64;
using std::ostringstream;
using std::string;
using std::uniform_real_distribution;

using paa::Bounds;
using paa::ChromosomeIndexLookup;
using paa::Error;
using paa::MappedVector;
using paa::Marker;
using paa::BridgeInfo;
using paa::PopBridgeInfo;
using paa::PSDoc;
using paa::PSGraph;
using paa::PSXYMSeries;
using paa::PSHSeries;
using paa::Reference;
using paa::sout;

using Info = PopBridgeInfo;

const Reference * paa::ref_ptr;
using paa::ref_ptr;

int main(int argc, char* argv[], char * []) try {
  if (--argc != 5)
    throw Error("usage: plot_invariants pop_dir ref chr start stop");

  // Arguments
  const string pop_dir{argv[1]};
  const Reference ref{argv[2]};
  ref_ptr = &ref;
  const ChromosomeIndexLookup lookup{ref};
  const string chromosome_name{argv[3]};
  // const unsigned int chromosome{lookup[chromosome_name]};
  const unsigned int start{static_cast<unsigned int>(atol(argv[4]))};
  const unsigned int stop{static_cast<unsigned int>(atol(argv[5]))};

  // Bridges file
  ostringstream bridges_file_name;
  bridges_file_name << pop_dir << "/popbridges."
      // << chromosome
                    << chromosome_name
                    << ".bin";
  const MappedVector<Info> bridges{bridges_file_name.str()};

  // Random scatter
  auto mersenne = std::mt19937_64();
  mersenne.seed(time(nullptr));
  uniform_real_distribution<double> dist{0, 1};
  function<double()> gen{bind(dist, mersenne)};

  // Bridge range limits
  auto pop_less = [](const Info & lhs, const unsigned int pos) {
    return lhs.pos1() < pos;
  };
  const Info * const lower{lower_bound(
      bridges.begin(), bridges.end(), start, pop_less)};
  const Info * const upper{lower_bound(
      bridges.begin(), bridges.end(), stop, pop_less)};

  // cerr << "here " << bridges.size() << " " << upper - lower << endl;

  // Plots
  const Marker negative_marker{paa::circle(), 0.2, "1 0 0", 1, true, "1 0 0"};
  const Marker positive_marker{paa::circle(), 0.2, "0 0 1", 1, true, "0 0 1"};
  PSDoc doc{"invariants"};
  PSGraph mixed_graph{doc, string("Invariants for region on chromosome ") +
        chromosome_name + ";Position;Log10(|Invariant|)"};
  PSXYMSeries mixed{mixed_graph};
  PSGraph hist_graph{doc, ";log10(|invariant|);N", Bounds{0.0, 10.0}};
  PSHSeries<double, uint64_t> hist{hist_graph, 100, "0 0 0", false};
  // hist_graph.log_y(true);

  // Loop over range
  for (const Info * b{lower}; b != upper; ++b) {
    const Info & bridge{*b};
    if (bridge.chr1() == bridge.chr2()) {
      if (bridge.invariant() == 0) continue;
      if (bridge.high1() == bridge.high2()) continue;
      // if (bridge.bridge_count() >= 20)
      mixed.add_point(bridge.pos1() + gen(),
                      log10(labs(bridge.invariant())),
                      bridge.invariant() > 0 ?
                      positive_marker: negative_marker);
      hist.add_point(log10(labs(bridge.invariant())));
      // cout << log10(labs(bridge.invariant())) << endl;
    }
  }

#if 0
  const string seq{ref.subseq(chromosome, start, stop)};
  for (unsigned int b{0}; b != seq.size(); ++b) {
    cout << seq[b];
    if (((b + 1) % 80) == 0) cout << "\n";
  }
  cout << endl;
#endif

  return 0;
} catch (Error & e) {
  cerr << "paa::Error:" << endl;
  cerr << e.what() << endl;
  return 1;
} catch (exception & e) {
  cerr << "std::exception" << endl;
  cerr << e.what() << endl;
  return 1;
} catch (...) {
  cerr << "unknown exception was caught" << endl;
  return 1;
}
```
