## Supplementary material for "G-Graph: An interactive genomic graph viewer": MUMdex Source Code for G-Graph: plot_simulate_cn.cpp

```
//
// plot_simulate_cn
//
// plot results from simulate_cn
//
// Copyright 2016 Peter Andrews @ CSHL
//

#include <algorithm>
#include <array>
#include <exception>
#include <fstream>
#include <iostream>
#include <map>
#include <memory>
#include <set>
#include <sstream>
#include <string>
#include <utility>
#include <vector>

#include "error.h"
#include "psplot.h"
#include "threads.h"

using std::array;
using std::cerr;
using std::cout;
using std::cin;
using std::endl;
using std::exception;
using std::ifstream;
using std::istringstream;
using std::make_unique;
using std::map;
using std::max;
using std::min;
using std::ostringstream;
using std::pair;
using std::set;
using std::string;
using std::to_string;
using std::unique_ptr;
using std::vector;

using paa::Error;
using paa::Marker;
using paa::PSDoc;
using paa::PSGraph;
using paa::PSPage;
using paa::PSXYSeries;
using paa::ThreadPool;

int main(int argc, char* argv[])  try {
  // Check usage
  if (--argc != 3) {
    throw Error("usage: plot_simulate_cn out_title level n_bins");
  }

  const string out_title{argv[1]};
  const double level{atof(argv[2])};
  const unsigned int n_bins{static_cast<unsigned int>(atoi(argv[3]))};
  argc -= 3;
  argv += 4;

  // Get data file names, lengths and cpbs used
  vector<double> lengths;
  vector<double> cpbs;
  using DataFiles = map<pair<double, double>, vector<string>>;
  const DataFiles data_files{[&lengths, &cpbs]() {
      set<double> s_lengths;
      set<double> s_cpbs;
      DataFiles result;
      string file_name;
      while (cin >> file_name) {
        istringstream file{file_name.c_str()};
        string prefix;
        getline(file, prefix, '.');
        string s_length;
        getline(file, s_length, '.');
        const double length{strtod(s_length.c_str(), nullptr)};
        s_lengths.insert(length);
        string s_cpb;
        getline(file, s_cpb, '.');
        const double cpb{strtod(s_cpb.c_str(), nullptr)};
        s_cpbs.insert(cpb);
        const pair<double, double> index{length, cpb};
        result[index].push_back(file_name);
        if (length <= 0 || cpb <= 0) cerr << file_name << endl;
      }
      lengths.assign(s_lengths.begin(), s_lengths.end());
      cpbs.assign(s_cpbs.begin(), s_cpbs.end());
      return result;
    }()};

  cout << "Lengths" << endl;
  for (unsigned int l{0}; l != lengths.size(); ++l) {
    cout << l << " " << lengths[l] << endl;
  }
  cout << "Counts per Bin" << endl;
  for (unsigned int c{0}; c != cpbs.size(); ++c) {
    cout << c << " " << cpbs[c] << endl;
  }

  // Graphs
  const std::string dark{"0 0 0"};
  const Marker dark_circle_marker{paa::circle(), 0.3, dark, 0.2, true};
  PSDoc ps{out_title, out_title};
  std::vector<std::unique_ptr<PSPage>> pages;
  std::vector<std::unique_ptr<PSGraph>> graphs;
  std::vector<std::unique_ptr<PSXYSeries>> series;

  // Main result
  using Perf = array<double, 3>;
  using PerfV = vector<Perf>;
  using PerfVV = vector<vector<PerfV> >;
  PerfVV performance(lengths.size(), vector<PerfV>(cpbs.size()));
  vector<vector<unsigned int>> total_n_trials(
      lengths.size(), vector<unsigned int>(cpbs.size()));
  double small{0.5};

  // Parallelization of reading files
  ThreadPool pool{6};
  ThreadPool::Results<void> results;

  // Read data files, calculate performance measures
  // should synchronize small, graph access
  cerr << "Read data" << endl;
  for (const auto file_info_ : data_files) {
    auto data_fun = [&lengths, &cpbs, &small, n_bins, level,
                     &performance, &total_n_trials,
                     &ps, &pages, &graphs, &series, &dark_circle_marker]
        (const DataFiles::value_type & file_info) {
      const double length{file_info.first.first};
      const double cpb{file_info.first.second};

      const unsigned int lengthi{[length, &lengths]() {
          return static_cast<unsigned int>(
              find(lengths.begin(), lengths.end(), length) - lengths.begin());
        }()};
      const unsigned int cpbi{[cpb, &cpbs]() {
          return static_cast<unsigned int>(
              find(cpbs.begin(), cpbs.end(), cpb) - cpbs.begin());
        }()};

      // Read in data
      const vector<string> & file_names{file_info.second};
      vector<pair<double, bool>> min_max_ratios;
      for (const string & file_name : file_names) {
        ifstream data_file{file_name.c_str()};
        unsigned int in_segment;
        double value;
        while (data_file >> in_segment >> value) {
          if (value < 10) {  // To fix a problem
            min_max_ratios.emplace_back(value, in_segment);
          } else {
            min_max_ratios.pop_back();  // To keep even number
          }
        }
      }

      // Process data to get sensitivities and specicifities
      PerfV & perf{performance[lengthi][cpbi]};
      sort(min_max_ratios.begin(), min_max_ratios.end(),
           [](const pair<double, bool> & lhs,
              const pair<double, bool> & rhs) {
             return lhs.first < rhs.first;
           });

      if (min_max_ratios.size() == 0) return;
      const double low{min_max_ratios.front().first};
      const double high{min_max_ratios.back().first};
      unsigned int n_below[2]{0, 0};
      uint64_t li{0};
      const unsigned int n_level_bins{500};
      const uint64_t n_trials{min_max_ratios.size() / 2};
      total_n_trials[lengthi][cpbi] += n_trials;
      const double run_small{1.0 / n_trials};
      small = min(small, run_small);
      bool near_success{false};
      for (unsigned int i{0}; i != n_level_bins + 1; ++i) {
        const double value{low + (high - low) * i / n_level_bins};
        while (li != min_max_ratios.size() &&
               min_max_ratios[li].first <= value)
          ++n_below[min_max_ratios[li++].second];
        const double sens{max(1 - 1.0 * n_below[1] / n_trials, run_small)};
        const double spec{max(1.0 * n_below[0] / n_trials, run_small)};
        if (sens > run_small && sens < 0.3 && spec > run_small && spec < 0.3)
          near_success = true;
        if (sens > run_small && spec > run_small)
          perf.push_back(Perf{{sens, spec, value}});
      }

      const bool do_threshold_plot{false};
      if (do_threshold_plot && near_success) {
        ostringstream title_string;
        title_string << length << " Mb, "
                     << cpb << " counts / bin, "
                     << n_bins << " bins, "
                     << level << " level";

        pages.push_back(make_unique<PSPage>(
            ps, title_string.str(), "1 2"));
        graphs.push_back(make_unique<PSGraph>(
            *pages.back(),
            "Sensitivity;Threshold;Proportion real events missed"));
        graphs.back()->log_y(true);
        series.push_back(make_unique<PSXYSeries>(
            *graphs.back(), dark_circle_marker));
        PSXYSeries * sensitivity{series.back().get()};
        graphs.push_back(make_unique<PSGraph>(
            *pages.back(),
            "Specificity;Threshold;Proportion of trials with fake events"));
        graphs.back()->log_y(true);
        series.push_back(make_unique<PSXYSeries>(
            *graphs.back(), dark_circle_marker));
        PSXYSeries * specificity{series.back().get()};
        for (uint64_t p{0}; p != perf.size(); ++p) {
          sensitivity->add_point(perf[p][2], perf[p][0]);
          specificity->add_point(perf[p][2], perf[p][1]);
        }
      }
    };
    pool.run(results, data_fun, file_info_);
  }
  while (results.size()) results.get();

  cerr << "Make plots" << endl;

  // Final sensitivity - specificity plots
  auto cmp_thresh = [](const Perf & pl, const Perf & pr) {
    return pl[2] < pr[2];
  };
  auto cmp_thresh_val = [](const Perf & pl, const double r) {
    return pl[2] < r;
  };
  const vector<string> colors{
    "0.8 0.8 0.8",  // light grey
        "1 0 1",  // pink
        "0 0.8 0.8",  // teal
        "0.5 0.5 0.5",  // dark grey
        "0 0.9 0",  // light green
        "0 0 1",  // blue
        "0.8 0.8 0",  // dark yellow
        "0 0 0",  // black
        "1 0 0",  // red
        "1 1 0",  // yellow
        "0 0.6 0"  // dark green
        };
  auto color = [&colors](const unsigned int i) {
    return colors[i % colors.size()];
  };
  auto is_filled = [&colors](const unsigned int i) {
    return static_cast<bool>((i / colors.size()) % 2);
  };
  auto fill_color = [color, is_filled](const unsigned int i) {
    return is_filled(i) ? color(i) : "1 1 1";
  };
#if 0
  auto shape = [&colors](const unsigned int) {
    return paa::circle;
    // i < colors.size() * 2 ? paa::circle() : paa::square;
  };
#endif
  for (const bool do_length : {false}) {
    const vector<double> & subjects{do_length ? lengths : cpbs};
    const vector<double> & objects{do_length ? cpbs : lengths};
    for (unsigned int s{0}; s != subjects.size(); ++s) {
      unsigned int n_ok{0};
      unsigned int o1{1000};
      vector<unsigned int> legend_indexes(
          objects.size(), static_cast<unsigned int>(objects.size()));
      for (unsigned int o{0}; o != objects.size(); ++o) {
        PerfV & perf{performance[do_length ? s : o][do_length ? o : s]};
        if (perf.size()) {
          legend_indexes[o] = n_ok;
          ++n_ok;
          if (o1 == 1000) o1 = o;
        }
      }
      if (n_ok < 2) continue;
      const double subject{subjects[s]};
      ostringstream title;
      unsigned int max_trials{0};
      for (unsigned int o{0}; o != objects.size(); ++o) {
        max_trials = max(max_trials,
                         total_n_trials[do_length ? s : o][do_length ? o : s]);
      }
      title << subject << (do_length ? " Mb" : " cpb")
            << ",  " << n_bins << " bins, " << level << " level, "
            << max_trials << " max trials / series;"
            << "Proportion of true events missed;"
            << "Proportion of trials with false events";
      graphs.push_back(make_unique<PSGraph>(ps, title.str()));
      graphs.back()->log_x(true).log_y(true);
      const double min_plot{0.0001};
      const double n_decades{-log10(min_plot)};
      const double x_space{0.08};
      const double y_space{0.000};
      const double extra_x{pow(10, x_space * n_decades)};
      const double extra_y{pow(10, y_space * n_decades)};
      graphs.back()->range().
          xl(min_plot).yl(min_plot).xh(1 + extra_x).yh(1 + extra_y);
      graphs.back()->ps("c2g");
      for (unsigned int o{0}; o != objects.size(); ++o) {
        PerfV & perf{performance[do_length ? s : o][do_length ? o : s]};
        // cerr << s << " " << o << " " << perf.size() << endl;
        if (perf.empty()) continue;
        const unsigned int object(objects[o]);
        const Marker circle_marker{
          paa::circle(), 0.5, color(o), 1, true, fill_color(o)};
        series.push_back(make_unique<PSXYSeries>(
            *graphs.back(), circle_marker));
        graphs.back()->ps(color(o) + " c np 0.92 0.975 " +
                          to_string(legend_indexes[o]) + " [] 0 sd " +
                          " 0.03 mul sub gfc m cxy cxy np 5 0 360 arc " +
                          (is_filled(o) ? "fill" : "sp") + " m "
                          " 10 -3 rm 0 0 0 c (" + to_string(object) + " " +
                          (do_length ? "cpb" : "Mb") + ") show");
        for (const auto & vals :
                 performance[do_length ? s : o][do_length ? o : s]) {
          series.back()->add_point(vals[0], vals[1]);
        }
      }
      // Threshold lines
      const unsigned int n_lines{20};
      graphs.back()->ps("0 0 0 c");
      for (unsigned int i{0}; i <= n_lines; ++i) {
        const double value{level + i * (1 - level) / n_lines};
        graphs.back()->ps("np");
        bool hit_small{false};
        for (unsigned int o{0}; o != objects.size(); ++o) {
          PerfV & perf{performance[do_length ? s : o][do_length ? o : s]};
          sort(perf.begin(), perf.end(), cmp_thresh);
          const auto found = lower_bound(
              perf.begin(), perf.end(), value, cmp_thresh_val);
          if (found == perf.end()) continue;
          ostringstream pss;
          if (!legend_indexes[o])
            pss << " " << ((i % 2) ? "[1 6]" : "[4 4]") << " 0 sd "
                << (*found)[0] << " " << (*found)[1] << " gc m cxy ("
                << value << ") 5 8 rm gsave "
                << ((i % 2) == 0 ? "show " : "pop ") << " grestore "
                << ((i % 2) == 0 ? "l" : "m") << " ";
          if (!hit_small)
            pss << (*found)[0] << " " << (*found)[1] << " gc l ";
          if ((*found)[0] <= small * 5 ||
              (*found)[1] <= small * 5) hit_small = true;
          graphs.back()->ps(pss.str());
        }
        graphs.back()->ps("sp");
      }
    }
  }

  double min_thresh{1000.0};
  double max_thresh{0.0};
  for (unsigned int l{0}; l != lengths.size(); ++l) {
    for (unsigned int c{0}; c != cpbs.size(); ++c) {
      const PerfV & perf{performance[l][c]};
      if (perf.empty()) continue;
      min_thresh = min(min_thresh, perf.front()[2]);
      max_thresh = max(max_thresh, perf.back()[2]);
    }
  }
  cout << "Thresh min " << min_thresh << " max " << max_thresh << endl;

  return 0;
} catch (Error & e) {
  cerr << "paa::Error:" << endl;
  cerr << e.what() << endl;
  return 1;
} catch (exception & e) {
  cerr << "std::exception" << endl;
  cerr << e.what() << endl;
  return 1;
} catch (...) {
  cerr << "unknown exception was caught" << endl;
  return 1;
}
```
