## Supplementary material for "G-Graph: An interactive genomic graph viewer": MUMdex Source Code for G-Graph: plot_tnp.cpp

```
//
// plot_tnp.cpp
//
// plot tumor-normal / population data
//
// Copyright 2017 Peter Andrews @ CSHL
//

#include <algorithm>
#include <exception>
#include <string>
#include <vector>

#include "error.h"
#include "mumdex.h"
#include "tsv.h"

using std::cin;
using std::cout;
using std::cerr;
using std::endl;
using std::exception;
using std::string;
using std::vector;

using paa::nunset;
using paa::unset;
using paa::Bounds;
using paa::ChromosomeIndexLookup;
using paa::Error;
using paa::Reference;
using paa::TSV;

int main(int argc, char* argv[]) try {
  --argc;
  if (argc != 2 && argc != 3) throw Error("usage: plot_tnp ref name [sample]");

  // Genome reference
  const Reference ref{argv[1]};
  const ChromosomeIndexLookup lookup{ref};
  const unsigned int sample{argc == 3 ?
        static_cast<unsigned int>(atoi(argv[3])) : 1};

  // Load data file
  TSV data{argv[2]};
  data.pdf(false);
  data.sample = sample;

  // Add some calculated series to data
  data.add_data("rank2", [&data]() {
      vector<double> result;
      for (unsigned int i{0}; i != data.n_rows(); ++i) {
        result.push_back(i);
      }
      return result;
    }());
  data.add_data("invariant_mod",
                [](const int invariant) {
                  return std::max(-20, std::min(20, invariant));
                }, "invariant");
  data.add_data("bridge_count_co",
                [](const unsigned int c1,
                   const unsigned int c2,
                   const unsigned int c3) noexcept {
                  return c1 + c2 + c3;
                }, "co1", "coM", "coH");
  data.add_data("bridge_count_cm",
                [](const unsigned int c1,
                   const unsigned int c2,
                   const unsigned int c3) noexcept {
                  return c1 + c2 + c3;
                }, "cm1", "cmM", "cmH");
  data.add_data("bridge_count_p",
                [](const unsigned int c1,
                   const unsigned int c2,
                   const unsigned int c3) noexcept {
                  return c1 + c2 + c3;
                }, "p1", "pM", "pH");
  data.add_data_sr("abspos", [&ref, &lookup](
      const string & chromosome, const unsigned int position) {
                     return ref.abspos(lookup[chromosome], position);
                   }, "chrA", "posA");

  // Label the axes better than just column names, and set limits
  data("bridge_count_co").label("Total Bridge Count In Cancer Only");
  data("bridge_count_cm").label("Total Bridge Count In Matched Normal");
  data("bridge_count_p").label("Total Bridge Count In SSC Population");
  data("invariant_mod").label("min(-20, max(Invariant, 20))").range(-21, 21);
  data("abspos").label("Absolute Genome Position");

  data.hist("co1", Bounds{0, 55}, 55);
  data.hist("bridge_count_co", Bounds{0, 55}, 55);
  data.hist("bridge_count_cm", Bounds{0, 55}, 55);
  data.hist("bridge_count_p", Bounds{0, 1021}, 1021);

  data.plot("co1", "p1");
  data.plot("coM", "p1");
  data.plot("coH", "p1");

  data.plot("cm1", "p1");
  data.plot("cmM", "p1");
  data.plot("cmH", "p1");

  data.plot("cm1", "pM");
  data.plot("cmM", "pM");
  data.plot("cmH", "pM");

  data.plot("cm1", "pH");
  data.plot("cmM", "pH");
  data.plot("cmH", "pH");

  data.plot("co1", "pM");
  data.plot("coM", "pM");
  data.plot("coH", "pM");

  data.plot("co1", "pH");
  data.plot("coM", "pH");
  data.plot("coH", "pH");

  data.plot("bridge_count_co", "p1");
  data.plot("bridge_count_co", "pM");
  data.plot("bridge_count_co", "pH");

  data.plot("bridge_count_co", "bridge_count_p");

  data.plot("abspos", "invariant_mod");

#if 0
  data.select("class == LHL");
  data.plot("abspos", "n_people_in");
  data.select("class == HLH");
  data.plot("MS141", "MS267");
  data.plot("rank", "p_value");
  data.plot("abspos", "n_people_in");
  data.plot("n_people_in", "avg_count");
  data.plot("invariant_mod", "n_people_in");
  data.plot("invariant_mod", "n_people_in").range(-21, 21, -1, 11);
  data.plot("rank", "offset");

  // Do plots for different selections of data
  for (const string selection : {"class == HLH", ""}) {
    data.select(selection);

    // Plots
    data.plot("MS141", "MS267");
    data.plot("MS141", "MS149");
    data.plot("MS149", "MS267");
    data.plot("invariant_mod", "n_people_in").range(-21, 21, -1, 2041);
    data.plot("invariant_mod", "n_people_in").range(-21, 21, -1, 11);

    data.plot("max_count", "min_count").range(-1, 50, -1, 11);
    data.plot("avg_count", "max_count").xl(-1).xh(30);
    data.plot("avg_count", "n_people_in").range(-1, 30, -1, 2041);
    data.plot("rank", "p_value");
    if (selection == "") data.plot("rank", "p_value").xl(-1).xh(20);
    // data.plot("total_count", "p_value");
    // data.plot("frac_people", "total_ms_len");
    // data.plot("motif_len", "total_ms_len");
    // data.plot("motif_len", "n_copies");
    // data.plot("motif_len", "total_ms_len");
    data.plot("offset", "total_ms_len");
    // data.plot("offset", "motif_len");
    // data.plot("offset", "n_copies");
    data.plot("abspos", "frac_people");
    data.plot("abspos", "MS141");
    data.plot("abspos", "MS149");
    data.plot("abspos", "MS267");
  }
#endif

  return 0;
} catch (Error & e) {
  cerr << "paa::Error:" << endl;
  cerr << e.what() << endl;
  return 1;
} catch (exception & e) {
  cerr << "std::exception" << endl;
  cerr << e.what() << endl;
  return 1;
} catch (...) {
  cerr << "unknown exception was caught" << endl;
  return 1;
}
```
