## Supplementary material for "G-Graph: An interactive genomic graph viewer": MUMdex Source Code for G-Graph: plot_unequal.cpp

```
//
// plot_unequal.cpp
//
// plot results of unequal_bridges
//
// Copyright 2016 Peter Andrews @ CSHL
//

#include <algorithm>
#include <exception>
#include <string>
#include <vector>

#include "error.h"
#include "mumdex.h"
#include "tsv.h"

using std::cin;
using std::cout;
using std::cerr;
using std::endl;
using std::exception;
using std::string;
using std::vector;

using paa::nunset;
using paa::unset;
using paa::Bounds;
using paa::ChromosomeIndexLookup;
using paa::Error;
using paa::Reference;
using paa::TSV;

int main(int argc, char* argv[]) try {
  if (--argc != 2) throw Error("usage: plot_unequal ref name");

  // Genome reference
  const Reference ref{argv[1]};
  const ChromosomeIndexLookup lookup{ref};

  // Load data file
  TSV data{argv[2]};

  // Add some calculated series to data
  data.add_data("rank2", [&data]() {
      vector<double> result;
      for (unsigned int i{0}; i != data.n_rows(); ++i) {
        result.push_back(i);
      }
      return result;
    }());
  data.add_data("invariant_mod",
                [](const int invariant) {
                  return std::max(-20, std::min(20, invariant));
                }, "invariant");
  data.add_data("total_bridge_count",
                [](const unsigned int c1,
                   const unsigned int c2,
                   const unsigned int c3) noexcept {
                  return c1 + c2 + c3;
                }, "MS141", "MS149", "MS267");
  data.add_data_sr("abspos", [&ref, &lookup](
      const string & chromosome, const unsigned int position) {
                     return ref.abspos(lookup[chromosome], position);
                   }, "chrA", "posA");

  // Label the axes better than just column names, and set limits
  data("rank").label("Event Index, Ordered By P Value");
  data("n_people_in").label("Number of SSC People Seen In").range(-1, 2014);
  data("frac_people").label("Fraction Of SSC Population Event Was Seen In");
  data("total_count").label("Total Bridge Count In SSC Population");
  data("avg_count").
      label("Average Bridge Count In SSC People With Event").high(40);
  data("min_count").label("Minimum Bridge Count In SSC Population");
  data("max_count").label("Maximum Bridge Count In SSC Population").log(false);
  data("offset").label("Signed Distance Between Anchors In Read");
  data("MS141").label("Diagnosis Bridge Count").range(-1, 50);
  data("MS149").label("Remission Bridge Count").range(-1, 50);
  data("MS267").label("Relapse Bridge Count").range(-1, 50);
  data("p_value").label("Event P Value").log(true);
  data("log_p_value").label("Logarithm of Event P Value");
  data("total_ms_len").label("Total Repeat Length").high(50);
  data("n_copies").label("Number Of Copies Of Motif In Repeat");
  data("motif_len").label("Repeat Motif Length");
  data("invariant_mod").label("min(-20, max(Invariant, 20))").range(-21, 21);
  data("total_bridge_count").label("Total Bridge Count");
  data("abspos").label("Absolute Genome Position");

  // For auto-plotting, skip these columns
  data.do_not_plot("rank2", "n_people", "n_people_in",
                   "posA", "hA", "posB", "hB",
                   "invariant", "sA", "sB", "mcA", "mcB", "msA", "msB",
                   "p_value");

  if (0) {
    // vs all others
    data.plot("rank");
  }

  if (0) {
    // Every combo
    data.plot("MS141", "MS149", "MS267");
  }

  if (0) {
    // all vs all
    data.plot();
  }

  if (1) {
    data.plot("MS141", "MS267");
    data.plot("abspos", "n_people_in");

    data.select("class == LHL");
    data.plot("abspos", "n_people_in");

    data.select("class == HLH");

    data.plot("MS141", "MS267");
    data.plot("rank", "p_value");
    data.plot("abspos", "n_people_in");
    data.plot("n_people_in", "avg_count");
    data.plot("invariant_mod", "n_people_in");
    data.plot("invariant_mod", "n_people_in").range(-21, 21, -1, 11);
    data.plot("rank", "offset");

    return 0;
  }

  // Do plots for different selections of data
  for (const string selection : {"class == HLH", ""}) {
    data.select(selection);

    // Plots
    data.plot("MS141", "MS267");
    data.plot("MS141", "MS149");
    data.plot("MS149", "MS267");
    data.plot("invariant_mod", "n_people_in").range(-21, 21, -1, 2041);
    data.plot("invariant_mod", "n_people_in").range(-21, 21, -1, 11);

    data.plot("max_count", "min_count").range(-1, 50, -1, 11);
    data.plot("avg_count", "max_count").xl(-1).xh(30);
    data.plot("avg_count", "n_people_in").range(-1, 30, -1, 2041);
    data.plot("rank", "p_value");
    if (selection == "") data.plot("rank", "p_value").xl(-1).xh(20);
    // data.plot("total_count", "p_value");
    // data.plot("frac_people", "total_ms_len");
    // data.plot("motif_len", "total_ms_len");
    // data.plot("motif_len", "n_copies");
    // data.plot("motif_len", "total_ms_len");
    data.plot("offset", "total_ms_len");
    // data.plot("offset", "motif_len");
    // data.plot("offset", "n_copies");
    data.plot("abspos", "frac_people");
    data.plot("abspos", "MS141");
    data.plot("abspos", "MS149");
    data.plot("abspos", "MS267");
  }

  return 0;
} catch (Error & e) {
  cerr << "paa::Error:" << endl;
  cerr << e.what() << endl;
  return 1;
} catch (exception & e) {
  cerr << "std::exception" << endl;
  cerr << e.what() << endl;
  return 1;
} catch (...) {
  cerr << "unknown exception was caught" << endl;
  return 1;
}
```
