## Supplementary material for "G-Graph: An interactive genomic graph viewer": MUMdex Source Code for G-Graph: poisson.cpp

```
//
// poisson
//
// poisson pdf and cdf tester
//
// Copyright 2016 Peter Andrews @ CSHL
//

#include <exception>
#include <iostream>

#include "error.h"
#include "poisson.h"

using std::cout;
using std::cerr;
using std::endl;
using std::exception;

using paa::Error;
using paa::Poisson;

int main(int argc, char* argv[], char * []) try {
  if (--argc != 2) throw Error("usage: poisson mean max_n");

  const double mean{atof(argv[1])};
  const unsigned int max_n{static_cast<unsigned int>(atol(argv[2]))};

  const Poisson poisson{mean, max_n};

  for (unsigned int n{0}; n <= max_n; ++n) {
    cout << n << " "
         << poisson.pdf(n) << " "
         << poisson.cdf(n) << " "
         << poisson.ucdf(n) << endl;
  }

  return 0;
} catch (Error & e) {
  cerr << "paa::Error:" << endl;
  cerr << e.what() << endl;
  return 1;
} catch (exception & e) {
  cerr << "std::exception" << endl;
  cerr << e.what() << endl;
  return 1;
} catch (...) {
  cerr << "unknown exception was caught" << endl;
  return 1;
}
```
