## Supplementary material for "G-Graph: An interactive genomic graph viewer": MUMdex Source Code for G-Graph: poisson.h

```
//
// poisson.h
//
// Poisson pdf and cdfs
//
// copyright 2016 Peter Andrews
//

#ifndef PAA_POISSON_H_
#define PAA_POISSON_H_

#include <cmath>
#include <vector>

namespace paa {

class Poisson {
 public:
  Poisson(const double mean, const unsigned int max_n) :
      cached_pdf(max_n + 1), cached_cdf(max_n + 1) {
    std::vector<double> log_sums(max_n + 1);
    for (unsigned int n{0}; n <= max_n; ++n) {
      if (n == 0) {
        log_sums[n] = 0;
      } else {
        log_sums[n] = log_sums[n - 1] + log(n);
      }
      cached_pdf[n] = exp(-mean + n * log(mean) - log_sums[n]);
      cached_cdf[n] = (n ? cached_cdf[n - 1] : 0) + cached_pdf[n];
    }
  }
  double pdf(const unsigned int n) const { return cached_pdf[n]; }
  double cdf(const unsigned int n) const { return cached_cdf[n]; }
  double ucdf(const unsigned int n) const {
    return n ? 1 - cached_cdf[n - 1] : 1;
    // return 1 - cached_cdf[n] + cached_pdf[n];
  }

 private:
  std::vector<double> cached_pdf;
  std::vector<double> cached_cdf;
};

}  // namespace paa

#endif  // PAA_POISSON_H_
```
