## Supplementary material for "G-Graph: An interactive genomic graph viewer": MUMdex Source Code for G-Graph: position.h

```
//
// position.h
//
// chromosome position pairs, not used too much
//
// Copyright 2015 Peter Andrews @ CSHL
//

#ifndef PAA_POSITION_H
#define PAA_POSITION_H

namespace paa {

class PosInfo {
 public:
  PosInfo(const unsigned int chr_, const unsigned int pos_) :
      chr{chr_}, pos{pos_} {}
  unsigned int chr;
  unsigned int pos;
  bool operator<(const PosInfo & other) const {
    if (chr == other.chr) {
      return pos < other.pos;
    } else {
      return chr < other.chr;
    }
  }
  bool operator>=(const PosInfo & other) const {
    if (chr == other.chr) {
      return pos >= other.pos;
    } else {
      return chr >= other.chr;
    }
  }
  bool operator==(const PosInfo & other) const {
    return chr == other.chr && pos == other.pos;
  }
};

}  // namespace paa


#endif  // PAA_POSITION_H
```
