## Supplementary material for "G-Graph: An interactive genomic graph viewer": MUMdex Source Code for G-Graph: position_coverage.cpp

```
//
// position_coverage
//
// determine coverage at a position
//
// Copyright 2016 Peter Andrews @ CSHL
//

#include <exception>
#include <iomanip>
#include <iostream>
#include <map>
#include <string>

#include "error.h"
#include "mumdex.h"
#include "population.h"
#include "utility.h"

using std::cerr;
using std::cout;
using std::endl;
using std::exception;
using std::string;

using paa::sout;
using paa::ChromosomeIndexLookup;
using paa::Error;
using paa::Family;
using paa::Mappability;
using paa::MUM;
using paa::MUMdex;
using paa::Pair;
using paa::Population;
using paa::PosInfo;
using paa::Reference;
using paa::Sample;

int main(int argc, char* argv[])  try {
  // Check usage
  if (--argc < 6) {
    throw Error("usage: position_coverage samples_dir pop_file "
                "chr pos mum_len [sample|family] ...");
  }

  // Process command line arguments
  const string samples_dir{argv[1]};
  const Population pop{argv[2]};
  const string chrs{argv[3]};
  const unsigned int pos{static_cast<unsigned int>(atoi(argv[4]))};
  const unsigned int mum_len{static_cast<unsigned int>(atoi(argv[5]))};
  argc -= 5;
  argv += 5;

  // Load reference information
  const string first_sample_name{pop.sample(pop.samples(argv[1]).front())};
  const MUMdex first_mumdex{samples_dir + "/" + first_sample_name + "/mumdex"};
  const Reference & ref{first_mumdex.reference()};
  const ChromosomeIndexLookup chr_lookup{ref};
  const unsigned int chr{chr_lookup[chrs]};
  const Mappability map{ref};
  const unsigned int abspos{ref.abspos(chr, pos)};
  const unsigned int low{map.low(abspos)};
  const unsigned int high{map.high(abspos)};

  const unsigned int read_length{152};

  while (argc--) {
    // Get samples for sample or family name
    const string sample_or_family{argv++[1]};
    for (const Sample sample : pop.samples(sample_or_family)) {
      // Get sample info
      const Family family{pop.family(sample)};
      const string sample_name{pop.sample(sample)};
      const string family_name{pop.family(family)};

      // Load the mumdex for the sample
      const MUMdex mumdex{samples_dir + "/" + sample_name + "/mumdex"};

      // Count MUMs covering position
      const unsigned int start_pos{pos > read_length ?
            pos - read_length : 0};
      const auto lower = mumdex.lower_bound(PosInfo(chr, start_pos));
      const auto upper = mumdex.lower_bound(PosInfo(chr, pos + 1));
      unsigned int n_mums{0};
      for (auto iter = lower; iter != upper; ++iter) {
        const auto index(*iter);
        const Pair pair{mumdex.pair(index)};
        if (pair.dupe()) continue;
        const MUM mum{mumdex.mum(index)};
        if (pair.bad(mum.read_2())) continue;

        if (mum.length() < mum_len) continue;
        if (pos >= mum.position0() &&
            pos < mum.position0() + mum.length()) {
          ++n_mums;
        }
      }
      cout << family_name << " "
           << sample_name << " "
           << pop.member(sample) << " "
           << chrs << " "
           << pos << " "
           << low << " "
           << high << " "
           << n_mums << endl;
    }
  }

  cerr << "done" << endl;

  return 0;
} catch (Error & e) {
  cerr << "paa::Error:" << endl;
  cerr << e.what() << endl;
  return 1;
} catch (exception & e) {
  cerr << "std::exception" << endl;
  cerr << e.what() << endl;
  return 1;
} catch (...) {
  cerr << "unknown exception was caught" << endl;
  return 1;
}
```
