## Supplementary material for "G-Graph: An interactive genomic graph viewer": MUMdex Source Code for G-Graph: primers.cpp

```
//
// primers
//
// find primers for anchors
//
// Copyright 2016 Peter Andrews @ CSHL
//

#include <exception>
#include <iostream>
#include <string>
#include <vector>

#include "error.h"
#include "longSA.h"
#include "mapper.h"
#include "mumdex.h"
#include "sequence.h"

using std::cerr;
using std::cout;
using std::endl;
using std::exception;
using std::string;
using std::vector;

using paa::longSA;
using paa::ChromosomeIndexLookup;
using paa::Error;
using paa::GCATscore;
using paa::Mappability;
using paa::Mapper;
using paa::Reference;
using paa::Repeatness;

int main(int argc, char* argv[])  try {
  if (--argc != 8) throw Error("usage: primers ref_fasta "
                               "chr1 pos1 high1 chr2 pos2 high2 offset");

  const string ref_fasta{argv[1]};
  const longSA sa{ref_fasta.c_str(), true, true, false};
  const Reference ref{ref_fasta};
  const Mappability mappability{ref_fasta, true};
  const ChromosomeIndexLookup lookup{ref};

  const int offset{atoi(argv[8])};
  const unsigned int overlap{static_cast<unsigned int>(
      offset <= 0 ? -offset : 0)};

  ++argv;
  for (const bool pos2 : {false, true}) {
    // Get anchor positions
    const string achr_string{argv[1]};
    const unsigned int achr{lookup[achr_string]};
    const unsigned int apos{static_cast<unsigned int>(atoi(argv[2]))};
    const bool high{static_cast<bool>(atoi(argv[3]))};
    argv += 3;

    bool good{false};
    unsigned int n_tries{0};
    unsigned int n_low_score{0};
    unsigned int n_high_score{0};
    unsigned int n_not_unique{0};
    unsigned int n_matches_0{0};
    unsigned int n_matches_1{0};
    unsigned int n_matches_2{0};
    unsigned int n_matches_3{0};
    // Try score close to 60 first
    for (unsigned int s{0}; s != 5; ++s) {
      // Try many sliding windows
      for (unsigned int i{0}; i != 500; ++i) {
        // Try many sequence lengths
        for (unsigned int j{0}; j != 10; ++j) {
          ++n_tries;

          // Anchor position to avoid overlap
          const unsigned int aopos{high ?
                (apos > overlap + i ? apos - overlap - i : 0) :
                apos + overlap + i};

          // Sequence start position
          const unsigned int best_length{16 + j};
          const unsigned int start_pos{high ?
                (aopos + 1 > best_length ? aopos - best_length + 1 : 0) :
                aopos};
          const unsigned int stop_pos{high ? aopos + 1 : aopos + best_length};
          const unsigned int length{stop_pos - start_pos};
          const string sequence{ref.subseq(achr, start_pos, stop_pos)};
          const GCATscore score{sequence};
          if (score < 60 - s) {
            ++n_low_score;
            continue;
          }
          if (score > 60 + s) {
            ++n_high_score;
            break;
          }
          const auto mams = sa.find_mams(sequence);
          if (mams.size() != 1) {
            ++n_not_unique;
            continue;
          }
          const Repeatness repeatness{sequence, sa, ref, mappability};
          if (repeatness.mismatches()[0] != 1) {
            ++n_matches_0;
            continue;
          }
          if (repeatness.mismatches()[1] != 0) {
            ++n_matches_1;
            continue;
          }
          if (repeatness.mismatches()[2] > 10) {
            ++n_matches_2;
            continue;
          }
          if (repeatness.mismatches()[3] > 100) {
            ++n_matches_3;
            continue;
          }
          cout << pos2 << " "
               << n_tries << " "
               << length << " "
               << sequence << " "
               << repeatness.max_mappability() << " "
               << repeatness.mismatches()[0] << " "
               << repeatness.mismatches()[1] << " "
               << repeatness.mismatches()[2] << " "
               << repeatness.mismatches()[3] << " "
               << score.value() << endl;
          good = true;
          break;
        }
        if (good) break;
      }
      if (good) break;
    }
    if (!good)
      cout << "No good primer found for anchor "
           << n_low_score << " "
           << n_high_score << " "
           << n_not_unique << " "
           << n_matches_0 << " "
           << n_matches_1 << " "
           << n_matches_2 << " "
           << n_matches_3 << endl;
  }

  return 0;
} catch (Error & e) {
  cerr << "paa::Error:" << endl;
  cerr << e.what() << endl;
  return 1;
} catch (exception & e) {
  cerr << "std::exception" << endl;
  cerr << e.what() << endl;
  return 1;
} catch (...) {
  cerr << "unknown exception was caught" << endl;
  return 1;
}
```
