## Supplementary material for "G-Graph: An interactive genomic graph viewer": MUMdex Source Code for G-Graph: print_invariants.cpp

```
//
// print_invariants
//
// Output bridges for a region
//
// Copyright 2016 Peter Andrews @ CSHL
//

#include <algorithm>
#include <exception>
#include <functional>
#include <iostream>
#include <sstream>
#include <string>

#include "bridges.h"
#include "error.h"
#include "files.h"
#include "mumdex.h"
#include "population.h"
#include "utility.h"

using std::bind;
using std::cout;
using std::cerr;
using std::endl;
using std::exception;
using std::lower_bound;
using std::ostringstream;
using std::string;

using paa::readable;
using paa::BridgeInfo;
using paa::ChromosomeIndexLookup;
using paa::Error;
using paa::Family;
using paa::Mappability;
using paa::MappedVector;
using paa::PopBridgeInfo;
using paa::Population;
using paa::Reference;
using paa::Sample;
using paa::sout;

using Info = BridgeInfo;

const Reference * paa::ref_ptr;
using paa::ref_ptr;

int main(int argc, char* argv[], char * []) try {
  if (--argc != 7)
    throw Error("usage: print_invariants family_file samples_dir ref "
                "sample_name chr start stop");

  // Arguments
  const Population pop{argv[1]};
  const string samples_dir{argv[2]};
  const Reference ref{argv[3]};
  const Mappability map{ref};
  ref_ptr = &ref;
  const string sample_name{argv[4]};
  const ChromosomeIndexLookup lookup{ref};
  const string chromosome_name{argv[5]};
  const unsigned int chromosome{lookup[chromosome_name]};
  const unsigned int start{static_cast<unsigned int>(atol(argv[6]))};
  const unsigned int stop{static_cast<unsigned int>(atol(argv[7]))};

  // Family
  const Sample sample{pop.sample(sample_name)};
  const Family family{pop.family(sample)};
  const string family_name{pop.family(family)};

  // Bridges file
  ostringstream bridges_file_name;
  bridges_file_name << samples_dir << "/" << sample_name << "/"
                    << get_bridge_file_name(ref, chromosome);
  if (!readable(bridges_file_name.str()))
    throw Error("Could not open bridges file")
        << bridges_file_name.str() << paa::bridges_bad_message();
  const MappedVector<Info> bridges{bridges_file_name.str()};

  // Bridge range limits
  auto pop_less = [](const Info & lhs, const unsigned int pos) {
    return lhs.pos1() < pos;
  };
  const Info * const lower{lower_bound(
      bridges.begin(), bridges.end(), start, pop_less)};
  const Info * const upper{lower_bound(
      bridges.begin(), bridges.end(), stop, pop_less)};

  // Loop over range
  for (const Info * b{lower}; b != upper; ++b) {
    const Info & bridge{*b};
    if (bridge.chr1() == bridge.chr2() &&
        bridge.invariant() != 0) {
      if (bridge.high1() == bridge.high2()) continue;
      if (bridge.invariant() > -250) continue;
      if (bridge.invariant() < -650) continue;

      // if (bridge.bridge_count() == 1) continue;
      const unsigned int abspos1{ref.abspos(bridge.chr1(), bridge.pos1())};
      const unsigned int mappability1{map.low_high(bridge.high1(), abspos1)};
      const unsigned int excess1{bridge.anchor1_length() - mappability1};
      const unsigned int abspos2{ref.abspos(bridge.chr2(), bridge.pos2())};
      const unsigned int mappability2{map.low_high(bridge.high2(), abspos2)};
      const unsigned int excess2{bridge.anchor2_length() - mappability2};

      bridge.output(sout);
      sout << excess1 << excess2
           << sample_name << family_name << endl;
    }
  }

  return 0;
} catch (Error & e) {
  cerr << "paa::Error:" << endl;
  cerr << e.what() << endl;
  return 1;
} catch (exception & e) {
  cerr << "std::exception" << endl;
  cerr << e.what() << endl;
  return 1;
} catch (...) {
  cerr << "unknown exception was caught" << endl;
  return 1;
}
```
