## Supplementary material for "G-Graph: An interactive genomic graph viewer": MUMdex Source Code for G-Graph: ps2pdf_dir.sh

```
#! /bin/bash

cd $1

find . -name '*.ps' |
while read ps ; do
    pdf=${ps%.ps}
    pdf=${pdf%.eps}.pdf
    if [ ! -e $pdf ] ; then
        head -n 5 $ps |
        grep ^%%BoundingBox |
        awk '{print $4,$5}' |
        grep -v matches |
        while read x y ; do
            ps2pdf -dDEVICEWIDTHPOINTS=$x -dDEVICEHEIGHTPOINTS=$y \
                $ps $pdf
        done
    fi
done
```
