## Supplementary material for "G-Graph: An interactive genomic graph viewer": MUMdex Source Code for G-Graph: random_mumdex_sequences.cpp

```
//
// random_mumdex_sequences
//
// extract read sequences from a mumdex, in random order, allowing repetition
//
// Copyright 2017 Peter Andrews @ CSHL
//

#include <array>
#include <exception>
#include <iostream>
#include <random>
#include <string>

#include "error.h"
#include "mumdex.h"
#include "utility.h"

using std::array;
using std::cout;
using std::cerr;
using std::endl;
using std::exception;
using std::mt19937_64;
using std::random_device;
using std::string;
using std::uniform_int_distribution;

using paa::Error;
using paa::MUMdex;

int main(int argc, char * argv[]) try {
  paa::exit_on_pipe_close();

  if (--argc != 2)
    throw Error("usage: random_mumdex_sequences mumdex_name n");
  const string mumdex_name{argv[1]};
  const MUMdex mumdex(mumdex_name);
  const uint64_t n_seqs{static_cast<uint64_t>(atol(argv[2]))};

  cerr << "Output " << n_seqs
       << " mumdex sequences for " << mumdex_name << endl;

  if (n_seqs > mumdex.n_pairs())
    throw Error("Asking for more sequences than exist in mumdex");

  // Random distribution
  random_device rd;
  uniform_int_distribution<uint64_t> udist(0, mumdex.n_pairs() - 1);
  auto mersenne = mt19937_64(rd());

  for (uint64_t i = 0; i != n_seqs; ++i) {
    const uint64_t j{udist(mersenne)};
    const array<string, 2> sequences(mumdex.sequences(j));
    for (const auto & sequence : sequences) {
      cout << sequence << '\n';
    }
  }

  return 0;
} catch (Error & e) {
  cerr << "paa::Error:" << endl;
  cerr << e.what() << endl;
  return 1;
} catch (exception & e) {
  cerr << "std::exception" << endl;
  cerr << e.what() << endl;
  return 1;
} catch (...) {
  cerr << "unknown exception was caught" << endl;
  return 1;
}
```
