## Supplementary material for "G-Graph: An interactive genomic graph viewer": MUMdex Source Code for G-Graph: random_uniform_distances.cpp

```
//
// random_uniform_distances.cpp
//
// Check properties of distance between points in various dimensions
//
// Copyright 2018 Peter Andrews @ CSHL
//

#include <algorithm>
#include <exception>
#include <iostream>
#include <functional>
#include <random>
#include <string>
#include <vector>

#include "error.h"
#include "psplot.h"

using std::bind;
using std::cout;
using std::cerr;
using std::endl;
using std::exception;
using std::max;
using std::mt19937_64;
using std::random_device;
using std::string;
using std::to_string;
using std::uniform_real_distribution;
using std::vector;

using paa::Bounds;
using paa::Error;
using paa::Marker;
using paa::PSDoc;
using paa::PSGraph;
using paa::PSXYSeries;

template <class Val> inline Val sqr(const Val val) { return val * val; }

int main(int argc, char* argv[]) try {
  // Read command line arguments
  if (--argc != 2)
    throw Error("usage: random_uniform_distances n_points max_dim");
  const unsigned int n_points = static_cast<unsigned int>(
      strtoul(argv[1], nullptr, 10));
  const unsigned int max_dim = static_cast<unsigned int>(
      strtoul(argv[2], nullptr, 10));

  // Set up uniform real random number generator
  random_device rd;
  auto mersenne = mt19937_64(rd());
  auto gen = bind(uniform_real_distribution<double>(0, 1), mersenne);

  // Plots
  PSDoc plots{"distances", "distances", 600, 600};
  plots.pdf(true);
  const Marker marker{paa::circle(), 0.3, "0 0 0", 1, true, "0 0 0"};
  const Marker red_marker{paa::circle(), 0.1, "1 0 0", 1, true, "1 0 0"};

  // Distances
  vector<double> distances;
  const unsigned int n_distances{n_points * n_points / 2};
  distances.reserve(n_distances);

  // loop over dimensions
  for (uint64_t dim{1}; dim <= max_dim; ++dim) {
    for (const bool sphere : {false, true}) {
      cerr << "Dimension " << dim << " sphere " << sphere << endl;

      // Generate positions
      vector<vector<double>> coords(dim, vector<double>(n_points));
      for (uint64_t p{0}; p != n_points;) {
        double dist_from_origin{0};
        for (uint64_t d{0}; d != dim; ++d) {
          const double dist{gen()};
          coords[d][p] = dist;
          dist_from_origin += sqr(dist);
        }
        if (!sphere || dist_from_origin < 1) ++p;
      }

      // Get all distances
      distances.clear();
      for (uint64_t i{0}; i != n_points; ++i) {
        for (uint64_t j{i + 1}; j != n_points; ++j) {
          double dist{0};
          for (uint64_t d{0}; d != dim; ++d)
            dist += sqr(coords[d][i] - coords[d][j]);
          distances.push_back(sqrt(dist));
        }
      }

      // Make sorted distance plots, sampling points to show
      sort(distances.begin(), distances.end());
      PSGraph * const distances_graph{PSGraph::create(
          plots, "Distances for dimension " + to_string(dim) +
          ", " + to_string(n_points) + " points" +
          (sphere ? " (sphere)" : ""),
          Bounds{-0.05 * n_distances, 1.05 * n_distances, -0.05, 1.05})};
      PSXYSeries * const distances_plot{PSXYSeries::create(
          *distances_graph, marker)};
      PSXYSeries * const inv_distances_plot{PSXYSeries::create(
          *distances_graph, red_marker)};
      for (uint64_t p{0}; p < distances.size();
           p += max(n_distances / 1000, 1u)) {
        distances_plot->add_point(p, distances[p] / distances.back());
        inv_distances_plot->add_point(distances.size() - p - 1,
                                      1.0 - distances[p] / distances.back());
      }
    }
  }


  return 0;
} catch (Error & e) {
  cerr << "Error:" << endl;
  cerr << e.what() << endl;
  return 1;
} catch (exception & e) {
  cerr << "exception" << endl;
  cerr << e.what() << endl;
  return 1;
} catch (...) {
  cerr << "unknown exception was caught" << endl;
  return 1;
}
```
