## Supplementary material for "G-Graph: An interactive genomic graph viewer": MUMdex Source Code for G-Graph: rare.cpp

```
//
// rare
//
// study rare variation
//
// Copyright 2017 Peter Andrews @ CSHL
//

#include <algorithm>
#include <deque>
#include <exception>
#include <iostream>
#include <queue>
#include <set>
#include <sstream>
#include <string>
#include <utility>
#include <vector>

#include "bridges.h"
#include "error.h"
#include "files.h"
#include "genes.h"
#include "mumdex.h"
#include "population.h"
#include "utility.h"

using std::cerr;
using std::cout;
using std::deque;
using std::endl;
using std::exception;
using std::lower_bound;
using std::move;
using std::ostringstream;
using std::priority_queue;
using std::set;
using std::string;
using std::vector;

using paa::serr;
using paa::sout;
using paa::tout;
using paa::BridgeInfo;
using paa::ChromosomeIndexLookup;
using paa::Error;
using paa::Family;
using paa::FileVector;
using paa::GeneHitFinder;
using paa::GeneInfoInterval;
using paa::GeneXrefs;
using paa::HitType;
using paa::KnownGene;
using paa::KnownGenes;
using paa::Mappability;
using paa::MergeHelper;
using paa::MergeHelperCompare;
using paa::PopBridgeInfo;
using paa::Population;
using paa::Reference;
using paa::Sample;

using MUMdex = paa::FileMUMdex;
using Bridges = paa::FileBridges;

using Anchors = MUMdex::Anchors;
using Bridge = Bridges::Bridge;

const bool exclude_snps{true};
const bool exclude_indels{false};
const bool exclude_trans{true};
const bool exclude_inversions{true};
const bool allow_skips{true};

const unsigned int min_bridge_count{5};
const unsigned int min_support_length{20};
const unsigned int min_mate_count{1};
const unsigned int min_mate_support{10};
const unsigned int min_excess_mappability{5};
const unsigned int max_n_families{1};
const unsigned int max_pop_seen{0};
const int max_offset{1000};
const int min_offset{-1000};

char hit_char(const HitType info) {
  switch (info) {
    case HitType::none:
      return 'n';
      break;
    case HitType::gene:
      return 'g';
      break;
    case HitType::exon:
      return 'e';
      break;
    case HitType::intron:
      return 'i';
      break;
    default:
      throw Error("unexpected HitType");
  }
}

int main(int argc, char* argv[])  try {
  --argc;
  if (argc != 6) throw Error("usage: rare ref family_file bridges_dir "
                             "chromosome start stop");

  // Other command line arguments
  const string reference_file{argv[1]};
  const Reference ref{reference_file};
  const ChromosomeIndexLookup chr_lookup{ref};
  const Mappability map{reference_file, true};
  const string family_file{argv[2]};
  const Population pop{family_file};
  const string bridges_dir{argv[3]};
  const string chromosome_name{argv[4]};
  const unsigned int chromosome{chr_lookup[chromosome_name]};
  const unsigned int chromosome_offset{ref.offset(chromosome)};
  const unsigned int start{static_cast<unsigned int>(atoi(argv[5]))};
  const unsigned int stop{static_cast<unsigned int>(atoi(argv[6]))};

  // Load gene info
  const string genes_name{reference_file + ".bin/knownGene.txt"};
  const string isoforms_name{reference_file + ".bin/knownIsoforms.txt"};
  const string kgXrefs_name{reference_file + ".bin/kgXref.txt"};
  const KnownGenes genes{genes_name, isoforms_name, chr_lookup, ref};
  const GeneXrefs xref{kgXrefs_name};
  const GeneHitFinder gene_finder{genes};

  // Set up queue of bridge files
  deque<MergeHelper> helpers;
  using PQueue = priority_queue<MergeHelper*, vector<MergeHelper *>,
      MergeHelperCompare>;
  PQueue queue{MergeHelperCompare()};
  unsigned int n_samples_skipped{0};
  for (const auto s : pop.samples()) {
    ostringstream bridges_name;
    bridges_name << bridges_dir << "/" << pop.sample(s) << "/"
                 << get_bridge_file_name(ref, chromosome);
    MergeHelper helper{bridges_name.str(), s, start, stop};
    if (helper.valid_start()) {
      helpers.push_back(move(helper));
      queue.push(&helpers.back());
    } else {
      ++n_samples_skipped;
    }
    serr << pop.family(pop.family(s))
         << pop.sample(s)
         << pop.member(s)
         << bridges_name.str()
         << queue.size() << endl;
  }

  if (!allow_skips && n_samples_skipped)
    throw Error("Samples skipped") << n_samples_skipped;

  // Load PopBridgeInfo file, and advance to start location, if called for
  ostringstream pop_file_name;
  pop_file_name << bridges_dir << "/popbridges."
                << chromosome_name << ".bin";
  FileVector<PopBridgeInfo> pop_info{pop_file_name.str()};
  FileVector<PopBridgeInfo>::const_iterator pop_bridge{
        lower_bound(pop_info.begin(), pop_info.end(), start,
                    [](const PopBridgeInfo & bridge, const unsigned int val) {
                      return bridge.pos1() < val;
                    })};

  if (1) tout << "chrA" << "posA" << "highA"
              << "chrB" << "posB" << "highB"
              << "invariant" << "offset"
              << "mapA" << "mapB"
              << "supA" << "supB"
              << "mCA" << "mCB"
              << "mSA" << "mSB"
              << "nFam" << "nSam"
              << "max_bridge" << "total_bridge"
              << "pNP" << "pNB" << "pMed" << "pMax"
              << "cNT" << "cBT" << "cPT" << "cST" << "cBD" << "cPD" << "cSD"
              << "genes" << "members" << "counts"
              << endl;

  // Loop, adding one bridge at a time to all_bridges if the same event
  vector<BridgeInfo> all_bridges;
  vector<Sample> all_samples;
  unsigned int total_bridge_count{0};
  unsigned int normal_bridge_count{0};
  unsigned int normal_seen{0};
  bool reset{true};
  while (true) {
    // New bridge expected - start fresh
    if (reset) {
      all_bridges.clear();
      all_samples.clear();
      total_bridge_count = 0;
      normal_bridge_count = 0;
      normal_seen = 0;
      reset = false;
    }

    // Add a bridge to the list if the same event
    bool bridge_done{true};
    if (queue.size()) {
      MergeHelper * top{queue.top()};
      const BridgeInfo current{top->current()};
      if (all_bridges.empty() || !(all_bridges.back() < current)) {
        bridge_done = false;
        all_bridges.push_back(current);
        total_bridge_count += current.bridge_count();
        const Sample sample{top->sample()};
        if (pop.is_parent(sample)) {
          normal_bridge_count += current.bridge_count();
          ++normal_seen;
        }
        all_samples.emplace_back(sample);
        queue.pop();
        if (top->advance()) {
          queue.push(top);
        }
      }
      if (queue.empty()) bridge_done = true;
      if (bridge_done) reset = true;
    } else {
      if (all_samples.empty()) {
        break;
      }
    }

    // No new bridges added, so bridge list is ready to process
    if (bridge_done) {
      // The first bridge, the exemplar since all are the same
      const BridgeInfo & bridge{all_bridges[0]};

      // SNP cut
      const bool is_snp{bridge.chr1() == bridge.chr2() &&
            bridge.invariant() == 0 && bridge.high1() != bridge.high2()};
      if (is_snp && exclude_snps) continue;

      // Indel cut
      if (exclude_indels && bridge.chr1() == bridge.chr2() &&
          bridge.high1() != bridge.high2()) continue;

      // Translocation cut
      if (exclude_trans && bridge.chr1() != bridge.chr2()) continue;

      // Inversion cut
      if (exclude_inversions && bridge.chr1() == bridge.chr2() &&
          bridge.high1() == bridge.high2()) continue;

      // Offset cut
      if (bridge.offset() < min_offset || bridge.offset() > max_offset) {
        continue;
      }

      // Count families seen
      const unsigned int n_families{[&all_samples, &pop] {
          unsigned int n{0};
          Family last_family{100000};
          for (const Sample sample : all_samples) {
            const Family family{pop.family(sample)};
            if (family != last_family) {
              ++n;
              last_family = family;
            }
          }
          return n;
        }()};

      if (n_families > max_n_families) continue;

      // Gather max support counts and lengths
      unsigned int max_bridge_count{0};
      unsigned int max_anchor1_support{0};
      unsigned int max_anchor2_support{0};
      unsigned int max_mate_anchor1_count{0};
      unsigned int max_mate_anchor1_support{0};
      unsigned int max_mate_anchor2_count{0};
      unsigned int max_mate_anchor2_support{0};
      for (unsigned int b{0}; b != all_bridges.size(); ++b) {
        const BridgeInfo & sample_bridge{all_bridges[b]};
        if (max_bridge_count < sample_bridge.bridge_count()) {
          max_bridge_count = sample_bridge.bridge_count();
        }
        if (max_anchor1_support < sample_bridge.anchor1_length()) {
          max_anchor1_support = sample_bridge.anchor1_length();
        }
        if (max_anchor2_support < sample_bridge.anchor2_length()) {
          max_anchor2_support = sample_bridge.anchor2_length();
        }
        if (max_mate_anchor1_count < sample_bridge.mate_anchor1_count()) {
          max_mate_anchor1_count = sample_bridge.mate_anchor1_count();
        }
        if (max_mate_anchor2_count < sample_bridge.mate_anchor2_count()) {
          max_mate_anchor2_count = sample_bridge.mate_anchor2_count();
        }
        if (max_mate_anchor1_support < sample_bridge.mate_anchor1_length()) {
          max_mate_anchor1_support = sample_bridge.mate_anchor1_length();
        }
        if (max_mate_anchor2_support < sample_bridge.mate_anchor2_length()) {
          max_mate_anchor2_support = sample_bridge.mate_anchor2_length();
        }
      }

      // Low bridge count cut
      if (max_bridge_count < min_bridge_count) continue;

      // Short anchor support cut
      if (max_anchor1_support < min_support_length ||
          max_anchor2_support < min_support_length) continue;

      // Short mate support cut
      if (max_mate_anchor1_count < min_mate_count ||
          max_mate_anchor2_count < min_mate_count ||
          max_mate_anchor1_support < min_mate_support ||
          max_mate_anchor2_support < min_mate_support) continue;

      // Get mappability information
      const unsigned int mum1_abspos{chromosome_offset + bridge.pos1()};
      const unsigned int mum2_abspos{ref.offset(bridge.chr2()) + bridge.pos2()};
      const unsigned int mum1_map{map.low_high(bridge.high1(), mum1_abspos)};
      const unsigned int mum2_map{map.low_high(bridge.high2(), mum2_abspos)};

      // Short excess mappability cut
      if (max_anchor1_support < mum1_map) throw Error("Mappability");
      if (max_anchor1_support < min_excess_mappability + mum1_map) {
        continue;
      } else {
        if (max_anchor2_support < min_excess_mappability +  mum2_map) {
          continue;
        }
      }

      // Check population file for event
      string pop_string;
      while (pop_bridge != pop_info.end() && *pop_bridge < bridge) {
        ++pop_bridge;
      }
      if (pop_bridge != pop_info.end() && !(bridge < *pop_bridge)) {
        // Found matching event
        ostringstream pop_out;
        pop_out << (*pop_bridge).n_people() << "\t"
                << (*pop_bridge).n_bridges() << "\t"
                << (*pop_bridge).median_bridges() << "\t"
                << (*pop_bridge).max_bridges();
        pop_string = pop_out.str();
        if ((*pop_bridge).n_people() > max_pop_seen) {
          continue;
        }
      } else {
        const string default_pop_string{"0\t0\t0\t0"};
        pop_string = default_pop_string;
      }

      const unsigned int n_samples{static_cast<unsigned int>(
          all_samples.size())};

      unsigned int tot_n_proband{0};
      unsigned int tot_n_sibling{0};
      unsigned int n_non_trans{0};
      unsigned int n_proband_trans{0};
      unsigned int n_sibling_trans{0};
      unsigned int n_both_trans{0};
      unsigned int n_proband_denovo{0};
      unsigned int n_sibling_denovo{0};
      unsigned int n_both_denovo{0};

      // Look at each family separately
      ostringstream mstr;
      ostringstream cstr;
      while (all_samples.size()) {
        // Gather samples and bridges for family
        static vector<Sample> samples;
        static vector<BridgeInfo> bridges;
        samples.clear();
        bridges.clear();
        while (all_samples.size() &&
               (samples.empty() ||
                pop.family(samples.back()) ==
                pop.family(all_samples.back()))) {
          samples.push_back(all_samples.back());
          all_samples.pop_back();
          bridges.push_back(all_bridges.back());
          all_bridges.pop_back();
        }

        mstr << " " << pop.family(pop.family(samples.front())) << ":";

        // Parent sample cuts
        // bool just_kids{true};
        unsigned int n_parents{0};
        unsigned int n_proband{0};
        unsigned int n_sibling{0};
        for (const Sample sample : samples) {
          if (pop.is_parent(sample)) {
            if (pop.is_mother(sample)) {
              mstr << 'm';
            } else {
              mstr << 'f';
            }
            // just_kids = false;
            ++n_parents;
          } else {
            if (pop.is_proband(sample)) {
              mstr << 'p';
              ++n_proband;
            } else {
              mstr << 's';
              ++n_sibling;
            }
          }
        }
        for (const BridgeInfo & info : bridges) {
          cstr << " " << info.bridge_count();
        }
        if (n_parents) {
          if (n_sibling + n_proband == 0) {
            ++n_non_trans;
          } else {
            if (n_proband) {
              ++n_proband_trans;
              if (n_sibling) {
                ++n_both_trans;
              }
            }
            if (n_sibling) {
              ++n_sibling_trans;
            }
          }
        } else {
            if (n_proband) {
              ++n_proband_denovo;
              if (n_sibling) {
                ++n_both_denovo;
              }
            } else {
              ++n_sibling_denovo;
            }
        }
        if (n_parents && (n_proband + n_sibling) == 1) {
          tot_n_proband += n_proband;
          tot_n_sibling += n_sibling;
        }
      }
      // if (tot_n_proband + tot_n_sibling == 0) continue;
      tout << ref.name(bridge.chr1()) << bridge.pos1() << bridge.high1()
           << ref.name(bridge.chr2()) << bridge.pos2() << bridge.high2()
           << bridge.invariant() << bridge.offset()
           << mum1_map << mum2_map
           << max_anchor1_support << max_anchor2_support
           << max_mate_anchor1_count << max_mate_anchor2_count
           << max_mate_anchor1_support << max_mate_anchor2_support
           << n_families << n_samples
           << max_bridge_count << total_bridge_count
           << pop_string
           << n_non_trans << n_both_trans
           << n_proband_trans << n_sibling_trans
           << n_both_denovo << n_proband_denovo << n_sibling_denovo;

      set<string> gene_strings;
      if (bridge.chr1() == bridge.chr2() &&
          bridge.high1() != bridge.high2() &&
          bridge.invariant() < 0 &&
          bridge.invariant() > -100000 &&
          bridge.pos1() < bridge.pos2()) {
        auto intervals = gene_finder.lookup(
            bridge.chr1(), bridge.pos1(), bridge.pos2());
        for (const GeneInfoInterval * i : intervals) {
          const GeneInfoInterval & interval{*i};
          for (const auto & item : interval.info) {
            const unsigned int gene{item.first};
            const string name{xref[genes[gene].name].geneSymbol};
            const char hitchar{hit_char(item.second.hit)};
            string gene_string = name + "." + hitchar;
            if (hitchar == 'e') {
              gene_string += ".";
              gene_string += std::to_string(item.second.exon);
            }
            gene_strings.insert(gene_string);
          }
        }
      } else {
        for (const bool a : {false, true}) {
          const GeneInfoInterval & interval{gene_finder.lookup(
              a ? bridge.chr2() : bridge.chr1(),
              a ? bridge.pos2() : bridge.pos1())};
          for (const auto & item : interval.info) {
            const unsigned int gene{item.first};
            const string name{xref[genes[gene].name].geneSymbol};
            const char hitchar{hit_char(item.second.hit)};
            string gene_string = name + "." + hitchar;
            if (hitchar == 'e') {
              gene_string += ".";
              gene_string += std::to_string(item.second.exon);
            }
            gene_strings.insert(gene_string);
          }
        }
      }
      bool first{true};
      if (gene_strings.empty()) {
        tout << "-";
      } else {
        for (const string & text : gene_strings) {
          if (first) {
            tout << text;
            first = false;
          } else {
            cout << "," << text;
          }
        }
      }

      tout << mstr.str().substr(1) << cstr.str().substr(1) << endl;
    }
  }

  cerr << "finished bridge loop" << endl;

  return 0;
} catch (Error & e) {
  cerr << "paa::Error:" << endl;
  cerr << e.what() << endl;
  return 1;
} catch (exception & e) {
  cerr << "std::exception" << endl;
  cerr << e.what() << endl;
  return 1;
} catch (...) {
  cerr << "unknown exception was caught" << endl;
  return 1;
}
```
