## Supplementary material for "G-Graph: An interactive genomic graph viewer": MUMdex Source Code for G-Graph: repeatness.cpp

```
//
// repeatness
//
// measure the repeatness of sequences
//
// Copyright 2016 Peter Andrews @ CSHL
//

#include <exception>
#include <iostream>
#include <string>

#include "error.h"
#include "longSA.h"
#include "mapper.h"
#include "mumdex.h"
#include "sequence.h"

using std::cerr;
using std::cout;
using std::endl;
using std::exception;
using std::string;

using paa::longSA;
using paa::Error;
using paa::Mappability;
using paa::Mapper;
using paa::Reference;
using paa::Repeatness;

int main(int argc, char* argv[])  try {
  if (--argc < 2)
    throw Error("usage: repeatness ref_fasta [min] sequence ...");

  const string ref_fasta{argv[1]};

  const string second_arg{argv[2]};
  const bool minimal_output{second_arg == "min"};
  if (minimal_output) {
    --argc;
    ++argv;
  }

  // Create suffix array
  const longSA sa{ref_fasta.c_str(), true, true, false};
  const Reference ref{ref_fasta};
  const Mappability mappability{ref_fasta, true};

  while (++argv, --argc) {
    const string sequence{argv[1]};
    const Repeatness repeatness{sequence, sa, ref, mappability};
    if (minimal_output) {
      cout << repeatness.minimal() << endl;
    } else {
      cout << repeatness.summary() << endl;
    }
  }

  return 0;
} catch (Error & e) {
  cerr << "paa::Error:" << endl;
  cerr << e.what() << endl;
  return 1;
} catch (exception & e) {
  cerr << "std::exception" << endl;
  cerr << e.what() << endl;
  return 1;
} catch (...) {
  cerr << "unknown exception was caught" << endl;
  return 1;
}
```
