## Supplementary material for "G-Graph: An interactive genomic graph viewer": MUMdex Source Code for G-Graph: repeats.cpp

```
//
// repeats
//
// Get repeat information for one or more positions
//
// Copyright 2016 Peter Andrews @ CSHL
//

#include "repeats.h"

#include <exception>
#include <iostream>
#include <string>
#include <vector>

#include "error.h"
#include "mumdex.h"

using std::cout;
using std::cerr;
using std::endl;
using std::exception;
using std::vector;

using paa::ChromosomeIndexLookup;
using paa::Error;
using paa::Reference;
using paa::MaskerRepeat;
using paa::RepeatMasker;

int main(int argc, char * argv[]) try {
  if (--argc < 4) throw Error("usage: repeats repeats_file ref chr pos ...");

  const Reference ref{argv[2]};
  const ChromosomeIndexLookup chr_lookup{ref};
  const RepeatMasker repeats{argv[1], chr_lookup};

  argc -= 2;
  argv += 2;
  while (argc) {
    const unsigned int chr{chr_lookup[argv[1]]};
    const unsigned int pos{static_cast<unsigned int>(atoi(argv[2]))};
    const vector<const MaskerRepeat *> result{repeats(chr, pos)};
    for (const MaskerRepeat * repeat : result) {
      cout << argv[1] << " " << argv[2] << " "
           << repeat->repeat << " " << repeat->family << endl;
    }
    argc -= 2;
    argv += 2;
  }

  return 0;
} catch (Error & e) {
  cerr << "paa::Error:" << endl;
  cerr << e.what() << endl;
  return 1;
} catch (exception & e) {
  cerr << "std::exception" << endl;
  cerr << e.what() << endl;
  return 1;
} catch (...) {
  cerr << "unknown exception was caught" << endl;
  return 1;
}
```
