## Supplementary material for "G-Graph: An interactive genomic graph viewer": MUMdex Source Code for G-Graph: repeats.h

```
//
// repeats.h
//
// information about repeats
//
// Copyright 2014 Peter Andrews @ CSHL
//

#ifndef PAA_REPEATS_H
#define PAA_REPEATS_H

#include <algorithm>
#include <iostream>
#include <string>
#include <sstream>
#include <vector>

#include "error.h"
#include "mumdex.h"

namespace paa {

class MaskerRepeat {
 public:
  MaskerRepeat(const std::string & line,
               const ChromosomeIndexLookup & lookup) {
    std::istringstream in{line.c_str()};
    std::string chromosome;
    std::string dummy;
    std::string strand;
    in >> score >> divergence >> deletion >> insertion >> chromosome
       >> start >> stop >> dummy >> strand >> repeat >> family;
    if (!in) throw Error("problem reading first part of line") << line;
    char par;
    char par2;
    if (strand == "+") {
      in >> repeat_start >> repeat_stop >> par >> repeat_left;
    } else {
      in >> par >> repeat_left >> par2 >> repeat_start >> repeat_stop;
    }
    if (!in) throw Error("Problem reading repeats line") << line;
    chr = lookup(chromosome);
  }

  unsigned int chr{};
  unsigned int start{};
  unsigned int stop{};

  unsigned int score{};
  float divergence{};
  float deletion{};
  float insertion{};
  bool flipped{};
  std::string repeat{};
  std::string family{};
  unsigned int repeat_start{};
  unsigned int repeat_stop{};
  unsigned int repeat_left{};
};

class RepeatMasker {
 public:
  RepeatMasker(const std::string & input_file,
               const ChromosomeIndexLookup & lookup) :
      starts{lookup.size()}, stops{lookup.size()} {
    std::ifstream input{input_file.c_str()};
    if (!input) throw Error("Problem opening repeats input file") << input_file;
    std::string line;
    getline(input, line);
    getline(input, line);
    repeats.reserve(5467460);
    unsigned int last_chr(-1);
    while (getline(input, line)) {
      try {
        MaskerRepeat repeat{line, lookup};
        repeats.push_back(repeat);
        if (repeats.back().chr != last_chr) {
          starts[repeats.back().chr] = &repeats.back();
          if (last_chr < stops.size()) {
            stops[last_chr] = &repeats.back();
          }
          last_chr = repeats.back().chr;
        } else {
          // const Repeat & current{repeats.back()};
          // const Repeat & last{*(&repeats.back() - 1)};
          // what goes here?
        }
      } catch (std::exception & e) {
        if (std::string("map::at") != e.what()) {
          throw;
        }
      }
    }
    if (repeats.empty()) throw Error("No repeats loaded");
    stops[last_chr] = &repeats.back() + 1;

    // sort within chromosomes
    for (unsigned int c{0}; c != starts.size(); ++c) {
      std::sort(starts[c], stops[c],
                [](const MaskerRepeat & lhs, const MaskerRepeat & rhs) {
                  if (lhs.stop == rhs.stop) {
                    return lhs.start < rhs.start;
                  } else {
                    return lhs.stop < rhs.stop;
                  }
                });
    }
  }

  const std::vector<const MaskerRepeat *> operator()(
      const unsigned int chr, const unsigned int pos) const {
    std::vector<const MaskerRepeat *> result;
    if (starts[chr] == nullptr) return result;
    auto found(
        std::lower_bound(starts[chr], stops[chr], pos,
                         [](const MaskerRepeat & lhs, const unsigned int p) {
                           return lhs.stop < p;
                         }));
    while (found != stops[chr] && found->start <= pos) {
      result.push_back(found++);
    }
    return result;
  }


  const std::vector<const MaskerRepeat *> operator()(
      const unsigned int chr1, const unsigned int pos1,
      const unsigned int chr2, const unsigned int pos2) const {
    std::vector<const MaskerRepeat *> result;
    if (starts[chr1] != nullptr) {
      auto found(
          std::lower_bound(starts[chr1], stops[chr1], pos1,
                           [](const MaskerRepeat & lhs, const unsigned int p) {
                             return lhs.stop < p;
                           }));
      while (found != stops[chr1] && found->start <= pos1) {
        result.push_back(found++);
      }
    }
    if (starts[chr2] != nullptr) {
      auto found(
          std::lower_bound(starts[chr2], stops[chr2], pos2,
                           [](const MaskerRepeat & lhs, const unsigned int p) {
                             return lhs.stop < p;
                           }));
      while (found != stops[chr2] && found->start <= pos2) {
        result.push_back(found++);
      }
    }
    sort(result.begin(), result.end(), [](const MaskerRepeat * lhs,
                                          const MaskerRepeat * rhs) {
           return lhs->stop - lhs->start > rhs->stop - rhs->start;
         });

    return result;
  }

 private:
  std::vector<MaskerRepeat> repeats{};
  std::vector<MaskerRepeat *> starts{};
  std::vector<MaskerRepeat *> stops{};
};


class ExactRepeat {
 public:
  ExactRepeat(const std::string & line,
               const ChromosomeIndexLookup & lookup) {
    std::istringstream in{line.c_str()};
    std::string chromosome;
    std::string dummy1;
    std::string dummy2;
    in >> chromosome >> start >> stop >> dummy1 >> n_copies >> dummy2 >> motif;
    if (!in) throw Error("problem reading exact repeat line") << line;
    chr = lookup(chromosome);
  }

  unsigned int chr{};
  unsigned int start{};
  unsigned int stop{};
  unsigned int n_copies{};
  std::string motif{};
};

// first generate input data file with generate_genome_repeats.sh
class ExactRepeats {
 public:
  ExactRepeats(const std::string & input_file,
               const ChromosomeIndexLookup & lookup) :
      starts{lookup.size()}, stops{lookup.size()} {
    std::ifstream input{input_file.c_str()};
    if (!input) throw Error("Problem opening repeats input file") << input_file;
    std::string line;
    repeats.reserve(376872423);
    unsigned int last_chr(-1);
    while (getline(input, line)) {
      try {
        ExactRepeat repeat{line, lookup};
        repeats.push_back(repeat);
        if (repeats.back().chr != last_chr) {
          starts[repeats.back().chr] = &repeats.back();
          if (last_chr < stops.size()) {
            stops[last_chr] = &repeats.back();
          }
          last_chr = repeats.back().chr;
        }
      } catch (std::exception & e) {
        if (std::string("map::at") != e.what()) {
          throw;
        }
      }
    }
    if (repeats.empty()) throw Error("No exact repeats loaded");
    stops[last_chr] = &repeats.back() + 1;

    // sort within chromosomes
    for (unsigned int c{0}; c != starts.size(); ++c) {
      std::sort(starts[c], stops[c],
                [](const ExactRepeat & lhs, const ExactRepeat & rhs) {
                  if (lhs.stop == rhs.stop) {
                    return lhs.start < rhs.start;
                  } else {
                    return lhs.stop < rhs.stop;
                  }
                });
    }
  }

  const std::vector<const ExactRepeat *> operator()(
      const unsigned int chr, const unsigned int pos) const {
    std::vector<const ExactRepeat *> result;
    if (starts[chr] == nullptr) return result;
    auto found(
        std::lower_bound(starts[chr], stops[chr], pos,
                         [](const ExactRepeat & lhs, const unsigned int p) {
                           return lhs.stop < p;
                         }));
    while (found != stops[chr] && found->start <= pos) {
      result.push_back(found++);
    }
    return result;
  }

  const std::vector<const ExactRepeat *> operator()(
      const unsigned int chr1, const unsigned int pos1,
      const unsigned int chr2, const unsigned int pos2) const {
    std::vector<const ExactRepeat *> result;
    if (starts[chr1] != nullptr) {
      auto found(
          std::lower_bound(starts[chr1], stops[chr1], pos1,
                           [](const ExactRepeat & lhs, const unsigned int p) {
                             return lhs.stop < p;
                           }));
      while (found != stops[chr1] && found->start <= pos1) {
        result.push_back(found++);
      }
    }
    if (starts[chr2] != nullptr) {
      auto found(
          std::lower_bound(starts[chr2], stops[chr2], pos2,
                           [](const ExactRepeat & lhs, const unsigned int p) {
                             return lhs.stop < p;
                           }));
      while (found != stops[chr2] && found->start <= pos2) {
        result.push_back(found++);
      }
    }
    sort(result.begin(), result.end(), [](const ExactRepeat * lhs,
                                          const ExactRepeat * rhs) {
           return lhs->motif.size() * lhs->n_copies >
               rhs->motif.size() * rhs->n_copies;
         });
    return result;
  }


 private:
  std::vector<ExactRepeat> repeats{};
  std::vector<ExactRepeat *> starts{};
  std::vector<ExactRepeat *> stops{};
};


}  // namespace paa


#endif  // PAA_REPEATS_H
```
