## Supplementary material for "G-Graph: An interactive genomic graph viewer": MUMdex Source Code for G-Graph: run_sample.sh

```
#! /bin/bash

# setup shell environment
. ~/mumdex/utility/shell.sh
rundir=/scratch/${USER}-temp-$$/
function error_delete() {
    echos Error: "$@"
    rm -Rf $rundir
    exit 1
}

#
# process command line arguments
#

# bam file
bam="$1"
[ "$bam" != "" ] || errors no bam file argument passed
[ -e "$bam" ] || errors bam file $bam does not exist

# out dir
outdir="$2"
[ "$outdir" != "" ] || errors no outdir argument passed
[ -e "$outdir" ] || errors outdir $outdir does not exist

# threads
n_threads="${3:-10}"
[ "$n_threads" != "" ] || errors no n_threads argument passed
[ "$n_threads" -gt 0 ] || errors bad n_threads $n_threads

# start processing
echo running MUMdex with arguments $bam $outdir $n_threads on $(hostname -s)
start=$(date +%s)

# directory to run in and copy output to
mkdir -p $rundir || errors cannot create rundir $rundir
cd $rundir || error_delete cannot cd to rundir $rundir

# copy and compile code
(cd ~/mumdex && make clean > /dev/null) || echos make clean problem
rsync -a ~/mumdex/ mumdex-code || echos rsync problem
(
    cd mumdex-code && make clean
    make -j 4 mummer merge_mumdex bridges namepair
) > /dev/null || error_delete cannot compile mumdex code
export PATH=$rundir/mumdex-code:~/bin:$PATH
compile=$(date +%s)

# run mummer
if mummer -normalmem -verbose -rcref -l 20 -qthreads $n_threads -samin \
    ~/hg38/hg38.fa <(
        namepair <(
            samtools view $bam &&
            warn samtools has succeeded || warn samtools has failed
        ) minimal &&
        warn namepair has succeded || warn namepair has failed
    ) ; then
    echo mummer has succeeded
else
    error_delete mummer has failed
fi
mummer=$(date +%s)

# run merge_mumdex
if merge_mumdex mumdex 10 $n_threads ; then
    echo merge_mumex has succeeded
else
    error_delete merge_mumdex has failed
fi
merge=$(date +%s)

# run bridges
if mkdir -p bridges_new &&
    (cd bridges_new && bridges ../mumdex $n_threads) ; then
    echo bridges has succeeded
else
    error_delete bridges has failed
fi
bridges=$(date +%s)

# clean up and copy results to outdir
(cd mumdex-code && make clean > /dev/null) || echos make clean problem
if cp -pr $rundir/* $outdir ; then
    echo copy has succeeded
else
    error_delete copy has failed
fi
rm -Rf $rundir || echos removal of rundir $rundir failed
move=$(date +%s)

# report timing and successful completion
echo timing compile $((compile - start)) \
    mummer $((mummer - compile)) \
    merge $((merge - mummer)) \
    bridges $((bridges - merge)) \
    move $((move - bridges)) \
    all $((move - start)) \
    n_threads $n_threads
echos all done

exit 0
```
