## Supplementary material for "G-Graph: An interactive genomic graph viewer": MUMdex Source Code for G-Graph: run_transmission.sh

```
#! /bin/bash

. ~/mumdex/utility/shell.sh

ref=$1
families=$2
bridges=$3
chr=$4
start=$5
stop=$6
n_threads=$7
if [ "$n_threads" = "" ] || [ $n_threads -lt 1 ] ; then
    error usage run_transmission.sh ref families bridges chr start stop n_threads
fi

node=$(hostname -s)
echo running MUMdex transmission on $node

start_time=$(date +%s)

[ ! -e cand.txt ] || error output file cand.txt already exists

echo rsync
rsync -a ~/mumdex/ mumdex-code > /dev/null

echo compile
(cd mumdex-code ; make clean ; make -j 4 transmission) > /dev/null ||
error problem compiling MUMdex

compile=$(date +%s)

export PATH=mumdex-code:$PATH
set -o pipefail

echo
echo running transmission
if transmission $ref $families $bridges $n_threads $chr $start $stop > cand.txt ; then
    echo transmission succeeded
else
    errors transmission failed
fi

transmission=$(date +%s)

echo
(cd mumdex-code ; make clean > /dev/null)

clean=$(date +%s)

echo timing compile $((compile - start_time)) transmission $((transmission - compile)) clean $((clean-transmission)) all $((clean-start_time))

echo all done with transmission

exit 0
```
