## Supplementary material for "G-Graph: An interactive genomic graph viewer": MUMdex Source Code for G-Graph: run_wg_samples.sh

```
#! /bin/bash

testing=0

# User modifiable configuration
base_dir=/mnt/wigclust19/data/safe/paa/analysis/mums
output_dir=$base_dir/wg-output
mummer_dir=$base_dir/mumdex
local_dir=/data/unsafe/$USER/mums
mkdir -p $output_dir
cd $output_dir

# Which samples to process
ids=$@
if [ "$ids" = '' ] ; then
    echo usage: run_samples.sh autism_id ... 1>&2
    exit 1
fi

# Select nodes
ok_nodes=$(echo wigclust{{1..3},{5..24}})
#ok_nodes=$(echo wigclust{{1..3},6,{8..16}})
queue_list=$(for node in $ok_nodes ; do echo -n all.q@$node, ; done)
queue_list=${queue_list%,}

# Pick reference based on testing status
if [ "$testing" = 1 ] ; then
    reference=chrM.fa
    reference_dir=$base_dir/hg19
    head="| head -n 100000"
    memory=5G
    threads=4
    email=''
else
    reference=chrAll.fa
    reference_dir=$base_dir/hg19
    memory=9G # times threads!
    threads=12
    merge_memory=100G
    # head="| head -n 1000"
fi
main_ref=$reference_dir/$reference
local_ref=$local_dir/$reference

# Commands
samtools=/data/software/samtools/samtools-0.1.19/samtools
echoe='tee >(cat 1>&2)'
set -o pipefail

# sge stuff
qsub=qsub
# qsub=cat
max_threads=20
sge_output='-o out.txt -e err.txt'
max_seconds=$((60 * 60 * 24 * 365))
sge_failure_code=1
print_job_name="perl -p -e 's/.*\"(.*)\".*/$1 /'"
qstat > qstat.txt

# Information about the run (and older runs)
run_id=$(date +%s)
run_dir=$output_dir/runs/run.$run_id
mkdir -p $run_dir
ln -sfT $run_dir run

# Make mummer in run dir as check
make_threads=""
mummer="mumdex/mummer-long -verbose -rcref"
merge="mumdex/merge_fam"
namepair="mumdex/namepair"
source='{Makefile,cpplint.py,*.{h,cpp,cc,mk}}'
if echo compiling mummer &&
    ! (mkdir -p $run_dir/mumdex && cd $run_dir/mumdex &&
        eval rsync -aL $mummer_dir/$source . &&
        make -sj $make_threads mummer-long merge_fam 2>&1 > make.err.tmp &&
        make -s clean) ; then
    echo making of mumdex failed 1>&2
    cat make.err.tmp
    rm -f make.err.tmp
    exit 1
fi

meta=$base_dir/wg-meta.txt

# Build sample list from input of quads and samples
for id in $ids ; do
    if [ "${id/SSC*/SSC}" = "$id" ] ; then
        ids=$(awk '{if ($1 == '$id') print $3}' $meta)
        if [ $(echo $ids | wc -w) -lt 3 ] ; then
            echo Too few samples found for quad $id 1>&2
            continue
        fi
        sample_ids="$sample_ids $ids"
    else
        sample_ids="$sample_ids $id"
    fi
done

echo samples $sample_ids

mkdir -p samples families nodes jobs

# Random
bsnl="\\
    "

# Set up samples to run
echo Trying to run $(echo $sample_ids | wc -w) samples
for sample in $sample_ids ; do

    # Check for reasonable sample
    if [ "$sample" = '' ] ; then
        echo empty sample name 1>&2
        exit 1
    fi
    if [ -e samples/$sample ] ; then
        echo $sample >> skipped.tmp
        continue
    fi

    info=$(awk '{if ($3 == "'$sample'") print}' $meta)

    # Find sample in report
    if [ "$info" = "" ] ; then
        echo sample $sample not found 1>&2
        exit 1
    fi

    read family member sample sex bam rest < <(echo $info)

    if [ "$rest" != "" ] ; then
        echo extra stuff found for sample $sample: $rest
        exit 1
    fi
    
    # Link sample to family directory
    mkdir -p families/$family
    ln -sfT ../../samples/$sample  families/$family/$member
    
    bams="<($namepair <($samtools view $bam $head) minimal)"
    
    # Determine threads needed for mumdex
    nt=$threads

    # Link to samples
    sample_dir=$run_dir/$sample
    rm -Rf $sample_dir
    mkdir -p $sample_dir
    ln -sft $sample_dir $output_dir/families/$family
    ln -sft $output_dir/samples $sample_dir
    
    map_memory=$memory
    
    running_sample=$local_dir/wg-samples-running/$sample
    final_samples=$local_dir/wg-samples
    final_sample=$final_samples/$sample
        
    # mumdex Mapping job
    (tee $sample_dir/mumdex.sh | $qsub) <<EOF
#! /bin/bash
#$ -N $sample.map -pe threads $nt -l vf=$map_memory -p -400
#$ -wd $sample_dir $sge_output
#$ -q $queue_list

node=\$(hostname -s)
echo running mumdex on \$node

rm -Rf $running_sample
mkdir -p $running_sample
cd $running_sample

set -o pipefail

echo syncing reference and code
if ! (rsync -a $run_dir/mumdex/ mumdex ) ; then
    echo rsync of code failed | $echoe
    exit 1
fi

echo compiling code
if ! (cd mumdex &&
      make -sj $make_threads mummer-long merge_fam namepair) ; then
    echo local making of mumdex failed | $echoe
    exit 1
fi

noden=\${node#wigclust}
if [ \$noden -lt 17 ] ; then
    normalmem=-normalmem
fi
echo running mummer
if $mummer -qthreads $nt \$normalmem -samin $local_ref $bams \
        2> map.out ; then
    echo mumdex has succeeded
    touch mapped
else
    echo mumdex has failed | $echoe
    (echo map output: ; cat map.out) 1>&2
    exit 1
fi

echo running merge_fam
if ! $merge fam ; then
    echo merge_fam has failed | $echoe
    exit 1
else
    echo merge_fam has succeeded
    rm mapped
    touch success
    rm -Rf mumdex
    rm -Rf $final_sample
    mkdir $final_samples
    mv $running_sample $final_sample
    ln -sft $output_dir/samples /mnt/\$node$final_sample
    ln -sft $output_dir/samples/$sample $sample_dir/out.txt
    ln -sft $output_dir/samples/$sample $sample_dir/err.txt
    ln -sft $output_dir/samples/$sample $sample_dir/mumdex.sh
    ln -sft $output_dir/samples/$sample $output_dir/families/$family

    exit 0
fi

EOF

done ||
(
    echo An error was detected - Quitting
    echo You may need to delete running jobs
    echo and clean sample directories
    exit 1
) || exit 1

echo Run $run_id started at $(date) >> $output_dir/runs/out.txt

# Report any skipped samples
if [ -e skipped.tmp ] ; then
    echo Skipped $(cat skipped.tmp|wc -l) already completed samples
    rm skipped.tmp
fi

rm -f qstat.txt *.tmp

exit 0
```
