## Supplementary material for "G-Graph: An interactive genomic graph viewer": MUMdex Source Code for G-Graph: setup_bridge_jobs.py

```
#! /bin/env python

import subprocess as sp

n_lines = 1000000

mums_dir = "/data/safe/paa/analysis/mums"
code_dir = "/home/paa/code/mumdex"
out_dir = mums_dir + "/bridges"

command = "cat " + mums_dir + "/genome.bed | " + \
    code_dir + "/split_bed.pl " + str(n_lines)

pipe = sp.Popen(command, shell=True, stdout=sp.PIPE).stdout

n = 0
for line in pipe:
    n += 1
    (chr, start, stop) = line.split()
    n_lines = int(stop) - int(start)
    job_name = "b" + str(n) 
    out = out_dir + "/" + job_name
    print "(echo 'export PATH=/data/software/local/bin:$PATH' ; echo ~/.local/bin/candidate_finder.py 1", chr, int(start) + 1, n_lines, \
        ") | qsub -l vf=10G -o", (out + ".out"), "-e", (out + ".err"), \
        "-N", job_name
```
