## Supplementary material for "G-Graph: An interactive genomic graph viewer": MUMdex Source Code for G-Graph: setup_check_bridges_1.sh

```
#! /bin/bash

#
# run check_bridges script for all candidates
#

n_jobs=${1:-100}
process_memory=20G

# machine-dependent setup
if [ "$SGE_ROOT" = /opt/sge6-2 ] ; then
    # wigclust
    options="-q all.q -l vf=$process_memory"
    base=~/analysis/mums
    main_node=wigclust19
else
    # NYGC
    options="-l h_vmem=$process_memory"
    base=~
    main_node=mumdel.nygenome.org
fi

code=~/mumdex
output=$base/bridges_out/details_1
families=$base/wg-families.txt
samples=$base/samples
bridges=$base/bridges

# make output directory
mkdir -p $output
cd $output

# Jobs directory
jobs=$output/jobs/job.$$
mkdir -p $jobs
subjobs=$jobs/to_run.txt

# Code for all runs
jobscode=$jobs/code
rsync -aL $code/ $jobscode
(cd $jobscode ; make clean > /dev/null)

# Select loci to examine
min_bridge_count=5
min_ref=10
max_families=5
min_support=25
min_mate_support=20
cat $output/../*/*-*/out.txt |
grep -v family |
awk '{if ($18 >= '$min_bridge_count' && $27 >= '$min_ref' && $4 == 1 && $4 <= '$max_families' && $19 >= '$min_support' && $20 >= '$min_support' && $22 >= '$min_mate_support' && $24 >= '$min_mate_support') print }'  | 
sort -k7,7n -k9,9n > candidates_used.txt

# A job for each family
cat $families |
awk '{print $1}' |
while read family ; do
    loci=$output/loci.$family.txt
    cat candidates_used.txt |
    grep "^$family" |
    awk '{print $8,$9,$10,$11,$12,$13,$16,$17}' | 
    uniq > $loci

    out=$family.out.txt
    if [ ! -e $out ] ; then
        echo "cat $loci | check_bridges $samples $bridges $families $family 2> $family.err.txt > $out"
    fi
done > $subjobs

n_subjobs=$(cat $subjobs | wc -l)
if [ $n_subjobs = 0 ] ; then
    echo nothing to do
    exit 0
fi

if [ $n_subjobs -lt $n_jobs ] ; then
    n_jobs=$n_subjobs
fi

# set up sge and job scripts
for jobn in $(eval echo {1..$n_jobs}) ; do
    job=$jobs/job$jobn
    mkdir -p $job

    jobcode=$job/code

    jobscript=$job/job.sh

    sgescript=$job/sge.sh
    name=${job##*/}
    touch $job/sge.err.txt
    touch $job/sge.out.txt
    touch $job/status.txt
    cat <<EOF > $sgescript
open=\$(ulimit -a| grep open | awk '{print \$4}')
start=\$(date +%s)
echo starting bridge_figure job on \$(hostname) at \$(date) >> $job/status.txt
rsync -a $jobscode/ $jobcode
(cd $jobcode ; make check_bridges > /dev/null) || exit 2
export PATH=$jobcode:$PATH
cd $output

while true ; do
  line=\$(ssh -t -t $main_node ~/mumdex/file_line.sh $subjobs | perl -pe 's/\s/ /g; s/\s+$//')
  if [ "\$line" = done ] ; then
    echo done recieved
    break
  fi
  if [ "\$line" = "" ] ; then
    echo no line recieved
    break
  fi
  echo running: "\$line"
  echo \$line | bash
  status=\$?
  echo status was \$status
done

status=\$?
echo finished check_bridges job. status: \$status >> $job/status.txt
stop=\$(date +%s)
echo total time was \$((stop - start)) >> $job/status.txt
exit \$status
EOF
    chmod a+x $sgescript

    echo qsub $options -N njob$jobn -o $job/sge.out.txt -e $job/sge.err.txt $sgescript
done > $jobs/sge.sh

if true ; then
    bash $jobs/sge.sh
else 
    echo To submit jobs, run:
    echo bash $jobs/sge.sh
fi
echo jobdir is $jobs
```
