## Supplementary material for "G-Graph: An interactive genomic graph viewer": MUMdex Source Code for G-Graph: setup_population_bridges.sh

```
#! /bin/bash


#
# run population_bridges program for all candidates
#

n_jobs=${2:-20}
process_memory=40G

# machine-dependent setup
if [ "$SGE_ROOT" = /opt/sge6-2 ] ; then
    # wigclust
    options="-q all.q -l vf=$process_memory"
    base=~/analysis/mums
    main_node=wigclust19
    ref=$base/hg19/chrAll.fa
    bed=$base/hg19/genome10M.bed
else
    # NYGC
    options="-l h_vmem=$process_memory"
    base=~
    main_node=mumdel.nygenome.org
    ref=$base/human_g1k_v37.fasta
    bed=$base/human_g1k_v37.10M.bed
fi

code=~/mumdex
output=$base/population_bridges/
family_file=$base/wg-families.txt
samples=$base/samples
bridges=$base/bridges

# make output directory
mkdir -p $output
cd $output

# Jobs directory
jobs=$output/jobs/job.$$
mkdir -p $jobs
subjobs=$jobs/to_run.txt

# Code for all runs
jobscode=$jobs/code
rsync -aL $code/ $jobscode
(cd $jobscode ; make clean > /dev/null)

# A job for each bed line
cat $bed |
while read chr start stop ; do
    dir=$output/$chr/$start-$stop
    if [ ! -e $dir ] ; then
        mkdir -p $dir
	touch $dir/err.txt
	touch $dir/out.txt
	touch $dir/status.txt

        echo "population_bridges denovo $family_file $bridges $ref $samples 1 $chr $start $stop 2>> $dir/err.txt >> $dir/out.txt"

    fi
done > $subjobs

n_subjobs=$(cat $subjobs | wc -l)
if [ $n_subjobs = 0 ] ; then
    echo nothing to do
    exit 0
fi

if [ $n_subjobs -lt $n_jobs ] ; then
    n_jobs=$n_subjobs
fi

sgescript=$jobs/sge.sh
    cat <<EOF > $sgescript
job=\$1
open=\$(ulimit -a| grep open | awk '{print \$4}')
start=\$(date +%s)
echo starting population_bridges job on \$(hostname) at \$(date) >> \$job/status.txt
rsync -a $jobscode/ \$job/code
(cd \$job/code ; make population_bridges > /dev/null) || exit 2
export PATH=\$job/code:\$PATH
cd $output

while true ; do
  line=\$(ssh -t -t $main_node ~/mumdex/file_line.sh $subjobs | perl -pe 's/\s/ /g; s/\s+$//')
  if [ "\$line" = done ] ; then
    echo done recieved
    break
  fi
  if [ "\$line" = "" ] ; then
    echo no line recieved
    break
  fi
  echo running: "\$line"
  echo "\$line" | bash
  status=\$?
  echo status was \$status
done

status=\$?
echo finished population_bridges job. status: \$status >> \$job/status.txt
stop=\$(date +%s)
echo total time was \$((stop - start)) >> \$job/status.txt
exit \$status
EOF
chmod a+x $sgescript

# set up sge and job scripts
for jobn in $(eval echo {1..$n_jobs}) ; do
    job=$jobs/job$jobn
    mkdir -p $job
    name=${job##*/}
    touch $job/sge.err.txt
    touch $job/sge.out.txt
    touch $job/status.txt

    echo qsub $options -N njob$jobn -o $job/sge.out.txt -e $job/sge.err.txt $sgescript $job
done > $jobs/qsub.sh

if true ; then
    bash $jobs/qsub.sh
else 
    echo To submit jobs, run:
    echo bash $jobs/qsub.sh
fi
echo jobdir is $jobs
```
