## Supplementary material for "G-Graph: An interactive genomic graph viewer": MUMdex Source Code for G-Graph: show_all_counts.cpp

```
//
// show_all_counts
//
// show reference and anchor counts over a population
//
// Copyright 2015 Peter Andrews @ CSHL
//

#include <exception>
#include <iostream>
#include <fstream>
#include <string>

#include "bed.h"
#include "error.h"
#include "mumdex.h"
#include "population.h"
#include "utility.h"

using std::exception;
using std::cout;
using std::cerr;
using std::endl;
using std::flush;
using std::ofstream;
using std::string;

using paa::BedFile;
using paa::ChromosomeIndexLookup;
using paa::CompressedInts;
using paa::Error;
using paa::MUMdex;
using paa::Mappability;
using paa::MappedVector;
using paa::Population;
using paa::Reference;
using paa::saved_ref_name;
using paa::sout;

int main(int argc, char* argv[]) try {
  --argc;
  if (argc < 4 || argc > 6)
    throw Error("usage: show_counts_special bed families pop_dir counts_dir"
                " [bed_begin [bed_end]]");

  // Read command line arguments
  const string bed_name{argv[1]};
  const BedFile bed{bed_name};

  const string family_file{argv[2]};
  const Population info{family_file};

  const string population_dir{argv[3]};

  const string counts_subdir{argv[4]};

  const unsigned int bed_start(argc >= 5 ? atoi(argv[5]) : 0);
  const unsigned int bed_stop(
      argc == 6 ? atoi(argv[6]) : (argc == 5 ? bed_start + 1 : bed.size()));

  // Calculate start position of first analysis block
  unsigned int block_start = 0;  // n loci into dataset for analysis block
  unsigned int bed_n = 0;  // n of current bed interval
  if (bed_start >= bed.size()) throw Error("bed_start too big");
  while (bed_n != bed_start) {
    block_start += bed[bed_n].stop_pos - bed[bed_n].start_pos;
    ++bed_n;
  }
  sout << "Using bed intervals" << bed_start << "to" << bed_stop
       << "and starting at index" << block_start << endl;

  // Output files
  info.output_sample_meta();
  ofstream position_meta("positions.txt");
  if (!position_meta) throw Error("Problem opening position_meta");
  ofstream low_reference("low_reference.txt");
  if (!low_reference) throw Error("Problem opening low_reference");
  ofstream high_reference("high_reference.txt");
  if (!high_reference) throw Error("Problem opening high_reference");
  ofstream low_anchor("low_anchor.txt");
  if (!low_anchor) throw Error("Problem opening low_anchor");
  ofstream high_anchor("high_anchor.txt");
  if (!high_anchor) throw Error("Problem opening high_anchor");
  ofstream low_support("low_support.txt");
  if (!low_support) throw Error("Problem opening low_support");
  ofstream high_support("high_support.txt");
  if (!high_support) throw Error("Problem opening high_support");

  // Loop over samples, output counts for each position
  cout << "sample" << flush;
  for (const auto s : info.samples()) {
    cout << ' ' << s << flush;
    const string mumdex_name = info.mumdex_name(population_dir, s);
    const auto counts_dir = mumdex_name + "/" + counts_subdir;
    const CompressedInts<uint16_t, uint8_t> compressed
    {counts_dir, block_start * 4, bed_start};
    const MappedVector<uint8_t> max_support{counts_dir + "/max_support.bin"};
    unsigned int n = block_start;
    for (unsigned int i = bed_start; i != bed_stop; ++i) {
      const auto & interval = bed[i];
      for (unsigned int pos = interval.start_pos; pos != interval.stop_pos;
           ++pos) {
        if (s == 0) {
          static const Reference ref{saved_ref_name(mumdex_name)};
          static const Mappability map{mumdex_name};
          static const ChromosomeIndexLookup chr_lookup{ref};
          const unsigned int chromosome{chr_lookup[interval.chromosome]};
          const uint64_t abspos = ref.offset(chromosome) + pos;
          position_meta << interval.chromosome << '\t' << pos + 1 << '\t'
                        << ref.offset(chromosome) + pos + 1 << '\t'
                        << i << '\t' << ref[chromosome][pos] << '\t'
                        << map.low(abspos) << '\t' << map.high(abspos) << endl;
        }
        low_reference << compressed.next_int() << ' ';
        low_anchor << compressed.next_int() << ' ';
        high_reference << compressed.next_int() << ' ';
        high_anchor << compressed.next_int() << ' ';
        low_support << static_cast<unsigned int>(max_support[n * 2]) << ' ';
        high_support << static_cast<unsigned int>(max_support[n * 2 + 1])
                    << ' ';
        ++n;
      }
    }
    low_reference << endl;
    low_anchor << endl;
    high_reference << endl;
    high_anchor << endl;
    low_support << endl;
    high_support << endl;
  }
  cout << endl;

  return 0;
} catch (Error & e) {
  cerr << "paa::Error:" << endl;
  cerr << e.what() << endl;
  return 1;
} catch (exception & e) {
  cerr << "std::exception" << endl;
  cerr << e.what() << endl;
  return 1;
} catch (...) {
  cerr << "unknown exception was caught" << endl;
  return 1;
}
```
