## Supplementary material for "G-Graph: An interactive genomic graph viewer": MUMdex Source Code for G-Graph: show_counts.cpp

```
//
// show_counts
//
// show reference and anchor counts (needs update)
//
// Copyright 2015 Peter Andrews @ CSHL
//

#include <exception>
#include <iostream>
#include <string>

#include "bed.h"
#include "error.h"
#include "mumdex.h"

using std::exception;
using std::cerr;
using std::endl;
using std::string;

using paa::BedFile;
using paa::CompressedInts;
using paa::Error;
using paa::sout;

int main(int argc, char* argv[]) try {
  paa::exit_on_pipe_close();

  if (--argc != 2) throw Error("usage: show_counts bed counts_dir");

  const string bed_name{argv[1]};
  const BedFile bed{bed_name};
  const string counts_dir{argv[2]};

  CompressedInts<uint16_t, uint8_t> compressed{counts_dir};

  for (const auto & interval : bed) {
    for (unsigned int pos = interval.start_pos; pos != interval.stop_pos;
         ++pos) {
      const auto coverage = compressed.next_int();
      const auto in_reference = compressed.next_int();
      const auto in_anchor = compressed.next_int();
      compressed.next_int();
      compressed.next_int();
      const auto out_reference = compressed.next_int();
      const auto out_anchor = compressed.next_int();
      compressed.next_int();
      compressed.next_int();
      sout << interval.chromosome << pos << coverage
           << in_reference << in_anchor
           << out_reference << out_anchor << endl;
    }
  }

  return 0;
} catch (Error & e) {
  cerr << "paa::Error:" << endl;
  cerr << e.what() << endl;
  return 1;
} catch (exception & e) {
  cerr << "std::exception" << endl;
  cerr << e.what() << endl;
  return 1;
} catch (...) {
  cerr << "unknown exception was caught" << endl;
  return 1;
}
```
