## Supplementary material for "G-Graph: An interactive genomic graph viewer": MUMdex Source Code for G-Graph: show_counts_special.cpp

```
//
// show_counts_special
//
// show reference and anchor counts and other info for a mumdex
//
// Copyright 2015 Peter Andrews @ CSHL
//

#include <exception>
#include <iostream>
#include <string>

#include "bed.h"
#include "error.h"
#include "mumdex.h"
#include "utility.h"

using std::exception;
using std::cerr;
using std::cout;
using std::endl;
using std::string;

using paa::BedFile;
using paa::ChromosomeIndexLookup;
using paa::CompressedInts;
using paa::Error;
using paa::MUMdex;
using paa::Mappability;
using paa::MappedVector;
using paa::Reference;
using paa::sout;

#define USE_SUPPORT 0

int main(int argc, char* argv[]) try {
  --argc;
  if (argc < 3 || argc > 5)
    throw Error("usage: show_counts_special bed mumdex "
                "counts [bed_begin [bed_end]]");

  const string bed_name{argv[1]};
  const BedFile bed{bed_name};
  const string mumdex_name{argv[2]};
  const string counts_dir{argv[3]};
  const MUMdex mumdex{mumdex_name};
  const Reference & ref = mumdex.reference();
  const Mappability map{mumdex_name};
  const ChromosomeIndexLookup chr_lookup{ref};

  const unsigned int bed_start(argc >= 4 ? atoi(argv[4]) : 0);
  const unsigned int bed_stop(argc == 5 ? atoi(argv[5]) :
                              (argc == 4 ? bed_start + 1 : bed.size()));

  // Jump to start position of first analysis block
  unsigned int block_start = 0;  // n loci into dataset for analysis block
  unsigned int bed_n = 0;  // n of current bed interval
  if (bed_start >= bed.size()) throw Error("bed_start too big");
  while (bed_n != bed_start) {
    block_start += bed[bed_n].stop_pos - bed[bed_n].start_pos;
    ++bed_n;
  }

  const auto last_slash = bed_name.find_last_of('/');
  const string bed_basename = last_slash == string::npos ?
      bed_name : bed_name.substr(last_slash + 1);
  // const auto counts_dir = mumdex_name + "/" + bed_basename;
  CompressedInts<uint16_t, uint8_t> compressed
  {counts_dir, block_start * 4, bed_start};
#if USE_SUPPORT
  MappedVector<uint8_t> max_support{counts_dir + "/max_support.bin"};
#endif
  unsigned int n = block_start;
  for (unsigned int i = bed_start; i != bed_stop; ++i) {
    const auto & interval = bed[i];
    for (unsigned int pos = interval.start_pos; pos != interval.stop_pos;
         ++pos) {
      if (!cout) throw Error("Output stream closed");
      const unsigned int chromosome = chr_lookup[interval.chromosome];
      const uint64_t abspos = ref.offset(chromosome) + pos;
      const auto coverage = compressed.next_int();
      const auto low_reference = compressed.next_int();
      const auto low_anchor = compressed.next_int();
      const auto high_reference = compressed.next_int();
      const auto high_anchor = compressed.next_int();
      sout << interval.chromosome << pos << ref[chromosome][pos]
           << n
           << coverage
           << low_reference << low_anchor
           << high_reference << high_anchor
           << map.low(abspos) << map.high(abspos)
#if USE_SUPPORT
           << static_cast<unsigned int>(max_support[n * 2])
           << static_cast<unsigned int>(max_support[n * 2 + 1])
#endif
           << endl;
      ++n;
    }
  }

  return 0;
} catch (Error & e) {
  cerr << "paa::Error:" << endl;
  cerr << e.what() << endl;
  return 1;
} catch (exception & e) {
  cerr << "std::exception" << endl;
  cerr << e.what() << endl;
  return 1;
} catch (...) {
  cerr << "unknown exception was caught" << endl;
  return 1;
}
```
