## Supplementary material for "G-Graph: An interactive genomic graph viewer": MUMdex Source Code for G-Graph: show_mums.cpp

```
//
// show_mums
//
// output MUM information in a text format
//
// Copyright 2015 Peter Andrews @ CSHL
//

#include <exception>
#include <iostream>
#include <string>
#include <vector>

#include "error.h"
#include "mumdex.h"
#include "utility.h"

using std::cout;
using std::cerr;
using std::endl;
using std::exception;
using std::string;
using std::vector;

using paa::Error;
using paa::MUMdex;
using paa::tout;

int main(int argc, char * argv[]) try {
  paa::exit_on_pipe_close();
  if (--argc != 1) throw Error("usage: show_mums mumdex_name");
  const string mumdex_name{argv[1]};
  const MUMdex mumdex{mumdex_name};
  const auto & ref = mumdex.reference();
  tout << "chr" << "position" << "readpos" << "read2" << "flipped"
       << "offset" << "length" << "last" << "touches" << "dupe" << endl;
  for (const auto index : mumdex.index()) {
    const auto pair = mumdex.pair(index);
    const auto mum = mumdex.mum(index);
    cout << ref.name(mum.chromosome()) << "\t"
         << mum.position1() << "\t"
         << mum.read_position1(pair.length(mum.read_2())) << "\t"
         << mum.read_2() << "\t"
         << mum.flipped() << "\t"
         << mum.offset() << "\t"
         << mum.length() << "\t"
         << mum.last_hit() << "\t"
         << mum.touches_end() << "\t"
         << pair.dupe() << "\n";
  }
  return 0;
} catch (Error & e) {
  cerr << "paa::Error:" << endl;
  cerr << e.what() << endl;
  return 1;
} catch (exception & e) {
  cerr << "std::exception" << endl;
  cerr << e.what() << endl;
  return 1;
} catch (...) {
  cerr << "unknown exception was caught" << endl;
  return 1;
}
```
