## Supplementary material for "G-Graph: An interactive genomic graph viewer": MUMdex Source Code for G-Graph: show_pairs.cpp

```
//
// show_pairs
//
// output pair information in a text format
//
// Copyright 2015 Peter Andrews @ CSHL
//

#include <algorithm>
#include <exception>
#include <iostream>
#include <string>

#include "error.h"
#include "mumdex.h"
#include "utility.h"

using std::cout;
using std::cerr;
using std::endl;
using std::exception;
using std::min;
using std::string;

using paa::Error;
using paa::MUMdex;
using paa::Mappability;
using paa::sout;
using paa::tout;

int main(int argc, char * argv[]) try {
  paa::exit_on_pipe_close();

  if (--argc != 1) throw Error("usage: show_pairs mumdex_name");

  const string mumdex_name{argv[1]};
  const MUMdex mumdex{mumdex_name};
  const Mappability map{mumdex_name};
  const auto & ref = mumdex.reference();

  sout << "id" << "read_len" << "bad" << "dupe" << "has" << "read_2"
       << "chr" << "pos" << "read_pos" << "flipped" << "offset"
       << "length" << "last_hit" << "touches_end" << "excess"
       << "special" << endl;
  for (uint64_t pair_index = 0; pair_index != mumdex.n_pairs(); ++pair_index) {
    const auto pair = mumdex.pair(pair_index);
    // const auto sequences = mumdex.sequences(pair_index);
    bool seen_read[2]{false, false};
    if (pair.has_mums()) {
      for (uint64_t m = pair.mums_start();; ++m) {
        const auto mum = mumdex.mum(m);  // mumdex.mums()[m] also works
        const auto offset = ref.offset(mum.chromosome());
        const auto abspos = offset + mum.position0();
        const auto min_map = map.min(abspos, mum.length());

        const auto first = pair.ordered_first_mums(mumdex.mums().begin());
        const auto special = (first[0] && mum == *first[0] ? 1 :
                              (first[1] && mum == *first[1] ? 2 : 0));

        cout << pair_index << " "
             << pair.length(mum.read_2()) << " "
             << pair.bad(mum.read_2()) << " "
             << pair.dupe() << " "
             << pair.has_mums() << " "
             << mum.read_2() << " "
             << ref.name(mum.chromosome()) << " "
             << mum.position1() << " "
             << mum.read_position1(pair.length(mum.read_2())) << " "
             << mum.flipped() << " "
             << mum.offset() << " "
             << mum.length() << " "
             << mum.last_hit() << " "
             << mum.touches_end() << " "
             << mum.length() - min_map << " "
             << special << "\n";
        seen_read[mum.read_2()] = true;
        if (mum.last_hit()) break;
      }
    }
    if (!seen_read[0]) {
      cout << pair_index << " "
           << pair.length(0) << " "
           << pair.bad(0) << " "
           << pair.dupe() << " "
           << pair.has_mums() << " 0\n";
    }
    if (!seen_read[1]) {
      cout << pair_index << " "
           << pair.length(1) << " "
           << pair.bad(1) << " "
           << pair.dupe() << " "
           << pair.has_mums() << " 1\n";
    }
  }

  return 0;
} catch (Error & e) {
  cerr << "paa::Error:" << endl;
  cerr << e.what() << endl;
  return 1;
} catch (exception & e) {
  cerr << "std::exception" << endl;
  cerr << e.what() << endl;
  return 1;
} catch (...) {
  cerr << "unknown exception was caught" << endl;
  return 1;
}
```
