## Supplementary material for "G-Graph: An interactive genomic graph viewer": MUMdex Source Code for G-Graph: simulate_cn.cpp

```
//
// simulate_cn
//
// Create know profiles from aggregates of individuals
//
// Copyright 2016 Peter Andrews @ CSHL
//

#include <algorithm>
#include <array>
#include <exception>
#include <fstream>
#include <iomanip>
#include <iostream>
#include <limits>
#include <memory>
#include <random>
#include <sstream>
#include <string>
#include <thread>
#include <utility>
#include <vector>

#include "cn.h"
#include "error.h"
#include "files.h"
#include "mumdex.h"
#include "psplot.h"
#include "threads.h"
#include "utility.h"

using std::array;
using std::binomial_distribution;
using std::cerr;
using std::cout;
using std::endl;
using std::exception;
using std::istringstream;
using std::lock_guard;
using std::make_unique;
using std::max;
using std::min;
using std::mt19937_64;
using std::mutex;
using std::numeric_limits;
using std::ofstream;
using std::ostringstream;
using std::pair;
using std::poisson_distribution;
using std::random_device;
using std::setprecision;
using std::string;
using std::to_string;
using std::uniform_int_distribution;
using std::unique_lock;
using std::unique_ptr;
using std::vector;

using paa::Bin;
using paa::Bounds;
using paa::Error;
using paa::FinestBins;
using paa::MappedVector;
using paa::Marker;
using paa::Progress;
using paa::PSDoc;
using paa::PSGraph;
using paa::PSHSeries;
using paa::PSPage;
using paa::PSXYSeries;
using paa::Reference;
using paa::ThreadPool;

#if 0
class Level {
 public:
  Level(const double value_, const bool in_segment_) :
      data(value_, in_segment_) { }
  double value() const { return data.first; }
  bool in_segment() const { return data.second; }

 private:
  pair<float, bool> data;
};
#else
class Level {
 public:
  Level(const double value_, const bool in_segment_) :
      data((in_segment_ ? -1 : 1) * value_) { }
  double value() const { return fabs(data); }
  bool in_segment() const { return data < 0; }

 private:
  float data;
};
#endif

int main(int argc, char* argv[])  try {
  // Check usage
  --argc;
  if (argc != 9) {
    throw Error("usage: simulate_cn ref bins.txt pos_counts bad_positions "
                "level median_cpbs lengths n_trials n_threads");
  }

  const bool show_progress{false};

  //
  // Process command line arguments
  //

  // Genome reference
  const Reference ref{argv[1]};

  // Bins
  const vector<Bin> all_bins{load_bins(argv[2], ref)};
  const vector<Bin> bins{[&all_bins]() {
      vector<Bin> result;
      for (const Bin & bin : all_bins)
        if (!bin.bad()) result.push_back(bin);
      return result;
    }()};
  const unsigned int x_chromosome{ref.find_x_chromosome()};
  const unsigned int x_start{[&bins, x_chromosome]() {
      for (unsigned int b{0}; b != bins.size(); ++b)
        if (bins[b].chromosome() == x_chromosome) return b;
      throw Error("X chromosome not found");
    }()};

  // Position counts
  const string pop_counts_name{argv[3]};

  // Bad positions
  const string bad_pos_name{argv[4]};

  // Segment level to search for (0.995 is 1% cells at copy number 1)
  const string level_str{argv[5]};
  const double level{atof(argv[5])};
  if (level >= 1) throw Error("Assumes level less than 1");

  // Counts per bin
  const vector<double> cpbs{[argv]() {
      istringstream cpbs_stream{argv[6]};
      vector<double> result;
      double value;
      while (cpbs_stream >> value) {
        cpbs_stream.get();
        result.push_back(value);
      }
      return result;
    }()};

  // Segment lengths in Mb
  const vector<double> lengths{[argv]() {
      istringstream lengths_stream{argv[7]};
      vector<double> result;
      double value;
      while (lengths_stream >> value) {
        lengths_stream.get();
        result.push_back(value);
      }
      return result;
    }()};
  const double mb{1000000.0};

  // Number of trials per condition
  const unsigned int n_trials{static_cast<unsigned int>(atoi(argv[8]))};
  // const double small{0.5 / n_trials};

  // Number of threads
  const unsigned int n_threads{static_cast<unsigned int>(atoi(argv[9]))};

  random_device rd;

  // Base name for output
  const string out_title{[&rd]() {
      uniform_int_distribution<uint64_t> udist{
        0, numeric_limits<uint64_t>::max()};
      auto mersenne = mt19937_64(rd());
      return to_string(udist(mersenne));
    }()};

  // Create counts vector - full counts from all samples
  vector<unsigned int> full_counts(x_start);
  ThreadPool pool{n_threads};
  ThreadPool::Results<void> results;
  {
    const FinestBins pop_counts{ref, pop_counts_name};
    const MappedVector<unsigned char> bad{bad_pos_name};
    for (unsigned int b{0}; b != x_start; ++b) {
      const Bin & cbin{bins[b]};
      pool.run(results, [&bad, &full_counts, &pop_counts]
               (const unsigned int abspos_start,
                const unsigned int abspos_stop,
                const unsigned int bin) {
                 for (unsigned int abspos{abspos_start}; abspos != abspos_stop;
                      ++abspos) {
                   if (bad[abspos]) continue;
                   full_counts[bin] += pop_counts[abspos];
                 }
               }, cbin.abspos_start(), cbin.abspos_stop(), b);
    }
    Progress fprogress{results.size(), 0.01, "full counts"};
    while (results.size()) {
      if (show_progress) fprogress();
      results.get();
    }
  }
  const double average_count{std::accumulate(
      full_counts.begin(), full_counts.begin() + x_start, 0.0) / x_start};

#define DO_HIST 0
#if DO_HIST
  // length x cpb x bin
  const unsigned int hist_bins{1000};
  vector<vector<vector<double> > > hist(
      cpbs.size(),
      vector<vector<double>>(lengths.size(), vector<double>(hist_bins)));
  const double edge{fabs(level - 1) * 3};
#endif

  // Graphs
  const std::string dark{"0 0 0"};
  const Marker dark_circle_marker{paa::circle(), 0.3, dark, 0.2, true};
  PSDoc ps{out_title, out_title};
  std::vector<std::unique_ptr<PSPage>> pages;
  std::vector<std::unique_ptr<PSGraph>> graphs;
  std::vector<std::unique_ptr<PSXYSeries>> series;
  using Hist = PSHSeries<double, unsigned int>;
  std::vector<std::unique_ptr<Hist> > hists;

#if 1
  // Bin count distribution - only reflects bin boundary choices
  PSGraph superperson_graph{ps,
        "Total bin count over population;Absolute Position;Ratio"};
  superperson_graph.log_y(true);
  superperson_graph.range().yl(1);
  PSXYSeries superperson_series{superperson_graph, dark_circle_marker};
  for (unsigned int b{0}; b != full_counts.size(); ++b) {
    superperson_series.add_point(bins[b].abspos_start(), full_counts[b]);
  }
#endif

  // Synchronization
  mutex rd_mutex;
  mutex progress_mutex;
  mutex graph_mutex;

  // Simulation progress
  Progress cprogress{cpbs.size() * lengths.size() * n_trials,
        0.01, "simulated counts"};

  // Simulation function for one length and one cpb, many trials
  auto run_fun = [&lengths, &cpbs,
                  n_trials,
                  level, &level_str,
                  &bins, &all_bins, x_start, mb,
                  &full_counts, average_count,
                  &ps, &pages, &graphs, &series,
                  &dark_circle_marker, &graph_mutex,
                  &out_title,
#if DO_HIST
                  &hist, edge,
#endif
                  &rd, &rd_mutex,
                  &cprogress, &progress_mutex]
      (const unsigned int lengthi, const unsigned int cpbi) {
    // Randomization
    unique_lock<mutex> rd_lock(rd_mutex);
    auto mersenne = mt19937_64(rd());
    rd_lock.unlock();

    const double length{lengths[lengthi]};
    // average bins in window
    const unsigned int bins_per_window(1.0 * x_start * length * mb /
                                       bins[x_start].abspos_start());
    uniform_int_distribution<unsigned int> udist{
      0, x_start - bins_per_window - 1};

    const double cpb{cpbs[cpbi]};
    const double downsample_rate{cpb / average_count};

    vector<Level> levels;
    // levels.reserve(x_start * n_trials);
    ostringstream title_string;
    title_string << length << " Mb, "
    << bins_per_window << " / " << all_bins.size() << " bins, "
    << static_cast<unsigned int>(cpb) << " counts / bin, "
    << level_str << " level";

    const unsigned int n_runs{2};
    vector<unsigned int> counts[2][n_runs];
    vector<pair<double, bool>> min_max_ratios;
    min_max_ratios.reserve(n_trials * 2);
    for (unsigned int t{0}; t != n_trials; ++t) {
      if (show_progress) {
        lock_guard<mutex> progress_lock(progress_mutex);
        cprogress();
      }
      const unsigned int bin_start{udist(mersenne)};
      const unsigned int bin_stop{bin_start + bins_per_window};
      const unsigned int abspos_start{bins[bin_start].abspos_start()};
      const unsigned int abspos_stop{bins[bin_stop].abspos_stop()};

      const unsigned int all_in_segment_start{
        bin_start + bins_per_window - 1};
      const unsigned int all_in_segment_stop{bin_stop};
      const unsigned int overlap_segment_start{bin_start};
      const unsigned int overlap_segment_stop{bin_stop + bins_per_window};

      const bool do_basic_page{true && t == 0};
      PSXYSeries * count_series[2]{nullptr, nullptr};
      PSXYSeries * ratio_series{nullptr};
      if (do_basic_page) {
        lock_guard<mutex> series_graph_lock(graph_mutex);
        pages.push_back(make_unique<PSPage>(
            ps, title_string.str() + ", trial " + to_string(t + 1),
            "1 2 =0 0 1 2="));
        for (const bool add_segment : {false, true}) {
          graphs.push_back(make_unique<PSGraph>(
              *pages.back(),
              string(add_segment ? "With" : "Without") + " Segment;"));
          graphs.back()->range().xl(1.0 * abspos_start - length * mb);
          graphs.back()->range().xh(abspos_stop + length * mb);
          if (add_segment) {
            const string start{to_string(abspos_start)};
            const string stop{to_string(abspos_stop)};
            graphs.back()->ps(string("1 0 0 c np ") + start +
                              " xc 0 yfc m " + start + " xc 1 yfc l " +
                              stop + " xc 0 yfc m " + stop + " xc 1 yfc l sp");
          }
          series.push_back(make_unique<PSXYSeries>(
              *graphs.back(), dark_circle_marker));
          count_series[add_segment] = series.back().get();
        }
        graphs.push_back(make_unique<PSGraph>(*pages.back(), ";abspos;ratio"));
        graphs.back()->ps("1 0 0 c np " + to_string(abspos_stop) +
                          " xc 0 yfc m  " + to_string(abspos_stop) +
                          " xc 1 yfc l sp");
        series.push_back(make_unique<PSXYSeries>(
            *graphs.back(), dark_circle_marker));
        ratio_series = series.back().get();
      }

      // Create simulated counts from combined counts
      for (const bool add_segment : {false, true})
        for (unsigned int r{0}; r != n_runs; ++r)
          counts[add_segment][r].clear();
      uint64_t running_total[2][n_runs]{{0, 0}, {0, 0}};
      unsigned int total_n{0};
      double min_max_ratio[2]{1000000.0, 1000000.0};
      for (unsigned int b{0}; b != x_start; ++b) {
        const Bin & bin{bins[b]};

        for (const bool add_segment : {false, true}) {
          const bool in_segment{add_segment && b >= bin_start && b < bin_stop};
          for (unsigned int r{0}; r != n_runs; ++r) {
            binomial_distribution<unsigned int> dist{
              full_counts[b], (in_segment ? level : 1.0) * downsample_rate};
            const unsigned int count{dist(mersenne)};
            counts[add_segment][r].push_back(count);
            running_total[add_segment][r] += count;
            if (do_basic_page && !r) {
              const bool near_segment{
                bin.abspos_start() + length * mb >= abspos_start ||
                    bin.abspos_stop() <= abspos_stop + length * mb};
              if (near_segment) {
                count_series[add_segment]->add_point(bin.abspos_start(), count);
              }
            }
          }
        }

        if (++total_n > bins_per_window) {
          --total_n;
          for (const bool add_segment : {false, true}) {
            for (unsigned int r{0}; r != n_runs; ++r) {
              running_total[add_segment][r] -=
                  counts[add_segment][r][b - bins_per_window];
            }
          }
          const uint64_t max_tumor{max(running_total[1][0],
                                       running_total[1][1])};
          const uint64_t min_normal{min(running_total[0][0],
                                        running_total[0][1])};
          const bool fully_in_segment{b >= all_in_segment_start &&
                b < all_in_segment_stop};
          const double max_ratio{1.0 * max_tumor / min_normal};
          if (fully_in_segment ||
              b < overlap_segment_start || b >= overlap_segment_stop) {
            min_max_ratio[fully_in_segment] =
                min(max_ratio, min_max_ratio[fully_in_segment]);
#if 0
            ++level_info[fully_in_segment][level_to_bin(max_ratio)];
#endif
          }
          if (do_basic_page)
            ratio_series->add_point(bin.abspos_start(), max_ratio);
        }
      }

      for (const bool fully_in_segment : {false, true}) {
        min_max_ratios.emplace_back(min_max_ratio[fully_in_segment],
                                    fully_in_segment);
      }

#if DO_HIST
      if (ratio >= 1 - edge && ratio < 1 + edge) {
        const unsigned int value_bin{static_cast<unsigned int>(
            hist_bins * (ratio - 1 + edge) / (2 * edge))};
        ++hist[lengthi][cpbi][value_bin];
      }
#endif
    }

    title_string << ", " << n_trials << " trials";
#if DO_HIST
    unique_lock<mutex> hist_graph_lock(graph_mutex);
    graphs.push_back(make_unique<PSGraph>(
        ps, title_string.str(), Bounds{1.0 - edge, 1.0 + edge}));
    graphs.back()->log_y(true);
    series.push_back(make_unique<PSXYSeries>(
        *graphs.back(), dark_circle_marker));
    PSXYSeries & hist_series{*series.back()};
    hist_graph_lock.unlock();
    for (unsigned int i{0}; i != hist[cpbi].size(); ++i) {
      const double ratio{1 - edge + i * 2 * edge / hist[lengthi][cpbi].size()};
      hist_series.add_point(ratio, hist[lengthi][cpbi][i]);
    }
#endif

    sort(min_max_ratios.begin(), min_max_ratios.end(),
         [](const pair<double, bool> & lhs, const pair<double, bool> & rhs) {
           return lhs.first < rhs.first;
         });
    ostringstream data_name;
    data_name << "sim_data." << length << "." << cpb << "."
    << out_title << ".txt";
    ofstream data_file{data_name.str().c_str()};
    data_file << setprecision(8);
    for (const pair<double, bool> val : min_max_ratios) {
      data_file << val.second << " " << val.first << "\n";
    }
  };

  // Run the simulation in parallel
  cerr << "Simulating autosome profiles" << endl;
  for (unsigned int lengthi{0}; lengthi != lengths.size(); ++lengthi) {
    for (unsigned int cpbi{0}; cpbi != cpbs.size(); ++cpbi) {
      pool.run(results, run_fun, lengthi, cpbi);
    }
  }
  while (results.size()) results.get();

  return 0;
} catch (Error & e) {
  cerr << "paa::Error:" << endl;
  cerr << e.what() << endl;
  return 1;
} catch (exception & e) {
  cerr << "std::exception" << endl;
  cerr << e.what() << endl;
  return 1;
} catch (...) {
  cerr << "unknown exception was caught" << endl;
  return 1;
}
```
