## Supplementary material for "G-Graph: An interactive genomic graph viewer": MUMdex Source Code for G-Graph: simulate_events2.cpp

```
//
// simulate_events
//
// simulate various event types to test mumdex
//
// Copyright 2016 Peter Andrews @ CSHL
//

#include <algorithm>
#include <array>
#include <exception>
#include <fstream>
#include <functional>
#include <iomanip>
#include <iostream>
#include <random>
#include <set>
#include <sstream>
#include <string>
#include <vector>

#include "error.h"
#include "fasta.h"
#include "mumdex.h"
#include "utility.h"

using std::array;
using std::cout;
using std::cerr;
using std::endl;
using std::exception;
using std::flush;
using std::function;
using std::max;
using std::min;
using std::mt19937_64;
using std::ofstream;
using std::ostringstream;
using std::random_device;
using std::set;
using std::setprecision;
using std::setw;
using std::string;
using std::uniform_int_distribution;
using std::uniform_real_distribution;
using std::vector;

using paa::Error;
using paa::Mappability;
using paa::MUM;
using paa::MUMdex;
using paa::MUMindex;
using paa::Pair;
using paa::Progress;
using paa::Reference;
using paa::SpaceOut;
using paa::reverse_complement;
using paa::sout;
using paa::tout;

struct ReadInfo {
  ReadInfo(const string & sequence_, const unsigned int pair_id_,
           const bool read_2_) :
      sequence{sequence_}, pair_id{pair_id_}, read_2{read_2_} { }

  string sequence;
  unsigned int pair_id;
  bool read_2;
  bool found{false};

  bool operator<(const ReadInfo & rhs) const {
    return sequence < rhs.sequence;
  }
  bool operator<(const string & rhs) const {
    return sequence < rhs;
  }
};

int main(int argc, char * argv[]) try {
  paa::exit_on_pipe_close();

  if (--argc != 2) throw Error("usage: simulate_events ref_fasta n_pairs");

  const Reference ref{argv[1]};
  const unsigned int n_pairs{static_cast<unsigned int>(atoi(argv[2]))};

  // Paramaters
  const double snp_prob{0.001};
  const double break_prob{0.001};
  const unsigned int read_length{151};
  const unsigned int min_mum{20};

  // Randomization
  random_device rd;
  mt19937_64 mersenne{rd()};
  using dist_real = uniform_real_distribution<double>;
  using dist_int_32 = uniform_int_distribution<uint32_t>;
  function<double()> realGen{bind(dist_real(0, 1), mersenne)};
  function<unsigned int()> baseGen{bind(dist_int_32(0, 3), mersenne)};
  function<unsigned int()> posGen{bind(dist_int_32(
      0, static_cast<unsigned int>(ref.size() - 1 - read_length)), mersenne)};

  // Create modified reference sequence
  vector<string> reference_segments;
  const string bases{"ACGT"};
  unsigned int n_snps{0};
  unsigned int n_breaks{0};
  Progress mod_progress(ref.size(), 0.1, "Modify reference");
  for (unsigned int c{0}; c != ref.n_chromosomes(); ++c) {
    for (unsigned int p{0}; p != ref.size(c); ++p) {
      mod_progress();
      // Break segments occasionally
      if (reference_segments.empty() ||
          (reference_segments.back().size() &&
           realGen() < break_prob)) {
        reference_segments.push_back("");
        if (reference_segments.size()) ++n_breaks;
      }
      char base{static_cast<char>(toupper(ref[c][p]))};
      // Mutate bases occasionally
      if (realGen() < snp_prob) {
        ++n_snps;
        char new_base;
        while ((new_base = bases[baseGen()]) == base) { }
        base = new_base;
      }
      reference_segments.back() += base;
    }
  }
  cout << "Introduced " << n_snps << " snps and " << n_breaks << " breaks\n";
  cout << "Shuffle and compose reference" << endl;
  shuffle(reference_segments.begin(), reference_segments.end(), mersenne);
  string modified_reference;
  modified_reference.reserve(ref.size());
  for (const string & segment : reference_segments) {
    modified_reference += segment;
  }
  reference_segments.clear();

  // Extract reads from modified reference
  vector<ReadInfo> reads;
  ofstream queries{"queries.sam"};
  ofstream sequences{"sequences.txt"};
  Progress reads_progress(n_pairs, 0.1, "Extract reads");
  set<string> unique_reads;
  for (unsigned int p{0}; p != n_pairs; ++p) {
    reads_progress();
    for (const bool read_2 : {false, true}) {
      // Read sequence
      string read;
      while (unique_reads.count(
                 read = modified_reference.substr(posGen(), read_length))) {}
      unique_reads.insert(read);
      reads.emplace_back(read, p, read_2);

      // Query output
      ostringstream out;
      SpaceOut<ostringstream> tsout{out, '\t'};
      tsout << p << (read_2 ? 141 : 77) << '*' << 0 << 0
            << '*' << '*' << 0 << 0
            << read << '*' << '\n';
      queries << out.str();
      sequences << read << endl;
    }
  }
  modified_reference.clear();
  unique_reads.clear();
  queries.close();
  sequences.close();

  cout << "Mapping queries" << endl;
  ostringstream mummer;
  mummer << "echo 'rm -Rf ./mumdex && "
         << "~/mumdex/mummer -verbose -samin -threads 12 -l "
         << min_mum << " ~/analysis/mums/hg19/chrAll.fa "
         << "queries.sam && echo Merging MUMdex && "
         << "~/mumdex/merge_mumdex mumdex 10 10 2> /dev/null' | bash";
  const time_t start_time{time(nullptr)};
  if (system(mummer.str().c_str()) == -1) {
    cerr << "Problem running mummer" << endl;
  }
  const time_t stop_time{time(nullptr)};
  const uint64_t elapsed = stop_time - start_time;
  sout << "MUMdex processing time was" << elapsed << "seconds or"
       << 60.0 * reads.size() / elapsed / 1000000
       << "million reads per minute" << endl;
  if (system("ms=$(du -bs mumdex | awk '{print $1}'); "
             "ss=$(du -bs sequences.txt | awk '{print $1}'); "
             "zs=$(cat sequences.txt | gzip -c | wc -c); "
             "echo MUMdex size is $ms bytes, sequence size is $ss bytes, "
             "sequence compression is $(perl -e 'print '$ss'/'$ms) or "
             "$(perl -e 'print '$zs'/'$ms) relative to zipped sequence") ==
      -1) {
    cerr << "Problem measuring compression" << endl;
  }
  // Check output for correctness
  cout << "Check correctness" << endl;
  sort(reads.begin(), reads.end());
  const MUMdex mumdex{"mumdex"};
  for (uint64_t p{0}; p != mumdex.n_pairs(); ++p) {
    uint64_t last_pair_id{0};
    const array<string, 2> pair_sequences = mumdex.sequences(p);
    for (const bool read_2 : {false, true}) {
      const string & sequence{pair_sequences[read_2]};
      auto found = lower_bound(reads.begin(), reads.end(), sequence);
      if (found == reads.end() || found->sequence != sequence) {
        throw Error("Sequence not found in lookup");
      }
      found->found = true;
      if (read_2 && last_pair_id != found->pair_id)
        throw Error("Pair Id Mismatch");
      last_pair_id = found->pair_id;;
    }
  }
  for (const ReadInfo & info : reads) {
    if (!info.found) {
        throw Error("Sequence not found in check");
    }
  }
  cout << "All reads found" << endl;

  return 0;
} catch (Error & e) {
  cerr << "paa::Error:" << endl;
  cerr << e.what() << endl;
  return 1;
} catch (exception & e) {
  cerr << "std::exception" << endl;
  cerr << e.what() << endl;
  return 1;
} catch (...) {
  cerr << "unknown exception was caught" << endl;
  return 1;
}
```
