## Supplementary material for "G-Graph: An interactive genomic graph viewer": MUMdex Source Code for G-Graph: simulate_events.cpp

```
//
// simulate_events
//
// simulate various event types to test mumdex
//
// Copyright 2016 Peter Andrews @ CSHL
//

#include <algorithm>
#include <array>
#include <exception>
#include <fstream>
#include <functional>
#include <iomanip>
#include <iostream>
#include <random>
#include <set>
#include <sstream>
#include <string>
#include <vector>

#include "error.h"
#include "fasta.h"
#include "mumdex.h"
#include "utility.h"

using std::array;
using std::cout;
using std::cerr;
using std::endl;
using std::exception;
using std::flush;
using std::function;
using std::max;
using std::min;
using std::mt19937_64;
using std::ofstream;
using std::ostringstream;
using std::random_device;
using std::set;
using std::setprecision;
using std::setw;
using std::string;
using std::uniform_int_distribution;
using std::uniform_real_distribution;
using std::vector;

using paa::Error;
using paa::Mappability;
using paa::MUM;
using paa::MUMdex;
using paa::MUMindex;
using paa::Pair;
using paa::Progress;
using paa::Reference;
using paa::SpaceOut;
using paa::reverse_complement;
using paa::sout;
using paa::tout;

const float snp_prob{1.0};
const unsigned int min_mum{75};
#if 0
const unsigned int min_length{900};
const unsigned int max_length{1023};
const unsigned int max_mum{800};
const unsigned int max_gap{1};
const unsigned int max_snp_length{1};
#else
const unsigned int min_length{151};
const unsigned int max_length{151};
const unsigned int max_mum{76};
const unsigned int max_gap{0};
const unsigned int max_snp_length{1};
#endif

class Event {
 public:
  string seq() const { return seq_; }
  string err() const { return err_; }
  string tag() const { return tag_; }

  mutable unsigned int chr[2]{0, 0};
  mutable unsigned int pos[2]{0, 0};
  mutable unsigned int len[2]{0, 0};
  mutable unsigned int edge[2]{0, 0};
  mutable unsigned int offset[2]{0, 0};
  mutable unsigned int flipped[2]{0, 0};
  mutable unsigned int gap{0};
  mutable string seq_{"seq"};
  mutable string err_{"*"};
  mutable string tag_{"tag"};
  mutable bool is_snp{false};
};

class EventMaker : public Event {
 public:
  using dist_int_32 = uniform_int_distribution<uint32_t>;
  using dist_real_32 = uniform_real_distribution<float>;
  EventMaker(const Reference & ref_arg, const Mappability & map_arg) :
      ref_{ref_arg}, map_{map_arg},
    posGen{bind(dist_int_32(0, static_cast<unsigned int>(ref_.size() - 1)),
                mersenne)},
    boolGen{bind(dist_int_32(0, 1), mersenne)},
    lenGen{bind(dist_int_32(min_mum, max_mum), mersenne)},
    gapGen{bind(dist_int_32(0, max_gap), mersenne)},
    edgeGen{bind(dist_int_32(0, max_gap), mersenne)},
    snpGen{bind(dist_int_32(1, max_snp_length), mersenne)},
    realGen{bind(dist_real_32(0, 1), mersenne)},
    baseGen{bind(dist_int_32(0, 3), mersenne)} { }

  string randseq(const unsigned int length) const {
    string result;
    for (unsigned int b{0}; b != length; ++b) {
      result += bases[baseGen()];
    }
    return result;
  }

  Event operator()() const {
    // Try to build bridge events randomly till a valid one is constructed
    bool good[2]{false, false};
    unsigned int n_tries{0};
    while (!good[0] || !good[1]) {
      while (!good[0] || !good[1]) {
        ++n_tries;
        ostringstream tagstream;
        unsigned int total_length{0};
        is_snp = realGen() < snp_prob;
        if (is_snp) {
          gap = snpGen();
        } else {
          gap = gapGen();
        }
        tagstream << "s." << is_snp << ".g." << gap << ".";
        for (const bool a : {false, true}) {
          if (a && !good[0]) break;
          good[a] = false;
          len[a] = lenGen();
          if (a && is_snp) {
            chr[1] = chr[0];
            flipped[1] = flipped[0];
            if (flipped[0]) {
              if (pos[0] < len[1] + gap) break;
              pos[1] = pos[0] - len[1] - gap;
            } else {
              pos[1] = pos[0] + len[0] + gap;
            }
            const unsigned int abspos{ref_.abspos(chr[1], pos[1])};
            if (map_.low(abspos) > 253) break;
            if (len[a] < map_.low(abspos)) break;
          } else {
            const unsigned int abspos{posGen()};
            const Reference::ChrPos chrpos{ref_.chrpos(abspos)};
            const unsigned int abspos2{ref_.abspos(chrpos)};
            if (abspos != abspos2) {
              throw Error("abspos problem") << abspos << abspos2;
            }
            chr[a] = chrpos.first;
            pos[a] = chrpos.second;
            flipped[a] = boolGen();
            if (map_.low(abspos) > 253) break;
            if (len[a] < map_.low(abspos)) break;
          }
          if (pos[a] + len[a] > ref_.size(chr[a])) break;
          edge[a] = edgeGen();
          if (a == 0) total_length += edge[a];
          offset[a] = total_length;
          total_length += len[a];
          if (a == 0) {
            total_length += gap;
          } else {
            total_length += edge[a];
          }
          tagstream << "c." << chr[a] << ".p." << pos[a] << ".l."
                    << len[a] << ".f." << flipped[a] << ".e."
                    << edge[a] << ".";
          if (a && total_length < min_length) break;
          if (total_length > max_length) break;
          good[a] = true;
        }
        tagstream << "l." << total_length << ".t." << n_tries;
        tag_ = tagstream.str();
      }
      seq_ = "";
      for (const bool a : {false, true}) {
        if (a == 0) {
          seq_ += randseq(edge[a]);
        } else {
          string gap_seq = randseq(gap);
          if (is_snp) {
            // Make sure is actually snp at edges of sequence
            const unsigned int low_a{flipped[0] ? 1U : 0U};
            while (toupper(gap_seq[0]) ==
                   ref_[chr[low_a]][pos[low_a] + len[low_a]] ||
                   toupper(gap_seq.back()) ==
                   ref_[chr[low_a]][pos[low_a] + len[low_a] + gap - 1]) {
              gap_seq = randseq(gap);
            }
            if (flipped[0]) {
              reverse_complement(&gap_seq);
            }
          }
          seq_ += gap_seq;
        }
        string mum{ref_.subseq(chr[a], pos[a], pos[a] + len[a])};
        if (flipped[a]) reverse_complement(&mum);
        seq_ += mum;
        if (a) {
          seq_ += randseq(edge[a]);
        }
      }
      if (seq_.find('N') != string::npos) good[0] = false;
    }
    // err_ = string(seq_.size(), 'B');
    return *this;
  }

 private:
  const Reference & ref_;
  const Mappability & map_;
  random_device rd{};
  const string bases{"ACGT"};
  mt19937_64 mersenne{rd()};
  function<uint32_t()> posGen;
  function<uint32_t()> boolGen;
  function<uint32_t()> lenGen;
  function<uint32_t()> gapGen;
  function<uint32_t()> edgeGen;
  function<uint32_t()> snpGen;
  function<float()> realGen;
  function<uint32_t()> baseGen;
};

int main(int argc, char * argv[]) try {
  paa::exit_on_pipe_close();

  if (--argc != 2) throw Error("usage: simulate_events ref_fasta n_events");

  const string fasta_name{argv[1]};
  const unsigned int n_events{static_cast<unsigned int>(atoi(argv[2]))};

  const Mappability map{fasta_name, true};
  const Reference ref{fasta_name};

  const EventMaker maker{ref, map};

  vector<Event> events;
  ofstream queries{"queries.sam"};
  Progress generate_progress(n_events, 0.01, "Generate queries loop");
  for (unsigned int n{0}; n != n_events; ++n) {
    generate_progress();
    ostringstream out;
    SpaceOut<ostringstream> tsout{out, '\t'};
    const Event event{maker()};
    events.push_back(event);
    tsout << n / 2 << (n % 2 ? 141 : 77) << '*' << 0 << 0
          << '*' << '*' << 0 << 0
          << event.seq() << event.err()
        // << event.tag()
          << '\n';
    // tout << n << out.str() << event.tag() << flush;
    queries << out.str();
  }
  queries.close();
  ostringstream mummer;
  mummer << "echo 'rm -Rf mumdex && ~/code/mumdex/mummer "
         << "-verbose -samin -qthreads 24 -l "
         << 2 << " ~/analysis/mums/hg19/chrAll.fa queries.sam "
         << "&& echo Merging MUMdex && "
         << "~/code/mumdex/merge_mumdex mumdex 10 10 2> /dev/null' | bash";
  cout << "Mapping queries" << endl;
  const time_t start_time{time(nullptr)};
  if (system(mummer.str().c_str()) == -1) {
    cerr << "Problem running mummer" << endl;
  }
  const time_t stop_time{time(nullptr)};
  const uint64_t elapsed = stop_time - start_time;
  sout << "MUMdex processing time was" << elapsed << "seconds or"
       << 60.0 * events.size() / elapsed / 1000000
       << "million reads per minute" << endl;
  if (system(("ms=$(du -bs mumdex | awk '{print $1}'); "
              "ss=" + std::to_string(n_events * (max_length + 1)) + "; "
              "echo MUMdex size is $ms bytes, sequence size is $ss bytes, "
              "sequence compression is $(perl -e 'print '$ss'/'$ms)").c_str())
      == -1) {
    cerr << "Problem measuring compression" << endl;
  }
  const MUMdex mumdex{"mumdex"};
  // Make sure each expected MUM is found, and record expected mum indices
  set<uint64_t> expected_mums;
  unsigned int n_seen{0};
  Progress check_progress(events.size(), 0.01, "Check MUMs loop");
  unsigned int n_mismatches{0};
  for (const Event & event : events) {
    check_progress();
    int last_read_pos{0};
    for (const bool a : {false, true}) {
      bool found{false};
      const string & seq{event.seq()};
      const unsigned int read_length{static_cast<unsigned int>(seq.size())};
      const unsigned int lowest_search_pos{read_length < event.pos[a] ?
            event.pos[a] - read_length : 0};
      for (const MUMindex pm : mumdex.region(
               event.chr[a], lowest_search_pos,
               event.chr[a], event.pos[a] + 1)) {
        const MUM mum{mumdex.mum(pm)};
        const int expected_position{mum.flipped() ?
              static_cast<int>(event.pos[a]) +
              static_cast<int>(event.len[a]) +
              static_cast<int>(event.offset[a]) - 1 :
              static_cast<int>(event.pos[a]) -
              static_cast<int>(event.offset[a])};
        if (mum.flipped() == event.flipped[a] &&
            mum.length() >= event.len[a] &&
            mum.read_start_position() == expected_position &&
            mum.offset() <= event.offset[a] &&
            mum.offset() + mum.length() >= event.offset[a] + event.len[a]) {
          const uint64_t pair_index{pm.pair_index()};
          const Pair pair{mumdex.pair(pm)};
          const unsigned int mum_in_pair{pm.mum_in_pair_index()};
          const uint64_t mum_index{pair.mums_start() + mum_in_pair};
          const array<string, 2> seqs = mumdex.sequences(pair_index);
          bool mismatch{false};
          for (unsigned int b{0}; b != seq.size(); ++b) {
            if (seqs[mum.read_2()][b] != toupper(seq[b])) {
              ++n_mismatches;
              mismatch = true;
              break;
            }
          }
          if (mismatch) continue;
          expected_mums.insert(mum_index);
          ++n_seen;
          if (event.is_snp && a && expected_position != last_read_pos)
            throw Error("SNP read positions disagree");
          found = true;
          last_read_pos = expected_position;
        }
      }
      if (!found) throw Error("missing event") << event.seq() << event.tag();
    }
  }
  const uint64_t n_expected{events.size() * 2};
  if (0)
    sout << "saw" << n_mismatches << "of approximately"
         << 1.0 * n_expected * (n_expected - 1) / 2 / ref.size() / 2
         << "expected mum collisions by birthday chance" << endl;
  sout << n_seen << "of" << n_expected << "expected mums found" << endl;
  const uint64_t n_spurious{mumdex.n_mums() - n_expected};
  sout << n_spurious << "spurious mums seen,"
       << 1.0 * n_spurious / n_expected
       << "spurious mums per expected mum" << endl;
  vector<unsigned int> length_counts(max_length);
  vector<unsigned int> excess_counts(max_length);
  unsigned int max_seen{0};
  Progress spurious_progress(n_spurious, 0.01, "Spurious check loop");
  const uint64_t min_size{10};
  const uint64_t max_size{40};
  const uint64_t n_elem_size{max_size - min_size};
  const uint64_t min_excess{0};
  const uint64_t max_excess{20};
  const uint64_t n_elem_excess{max_excess - min_excess};
  vector<vector<uint64_t>> combined_counts(n_elem_size,
                                           vector<uint64_t>(n_elem_excess));
  for (uint64_t m{0}; m != mumdex.n_mums(); ++m) {
    if (expected_mums.count(m)) continue;
    spurious_progress();
    const MUM mum{mumdex.mum(m)};
    ++length_counts[mum.length()];
    const unsigned int min_map{map.min(ref.abspos(mum.chromosome(),
                                                  mum.position0()),
                                       mum.length())};
    const unsigned int excess{mum.length() - min_map};
    ++excess_counts[excess];
    max_seen = max(max_seen, max(excess, mum.length()));
    for (uint64_t s{min_size}; s != max_size; ++s) {
      for (uint64_t e{min_excess}; e != max_excess; ++e) {
        if (mum.length() < s || excess < e) {
          ++combined_counts[s - min_size][e - min_excess];
        }
      }
    }
  }

  cout << setw(5) << "size"
       << setw(16) << "p_mum"
       << setw(16) << "mum_cdf"
       << setw(16) << "p_excess"
       << setw(16) << "excess_cdf" << endl;
  uint64_t n_length{0};
  uint64_t n_excess{0};
  for (unsigned int l{0}; l <= max_seen; ++l) {
    n_length += length_counts[l];
    n_excess += excess_counts[l];
    cout << setw(5) << l
         << setprecision(9) << setw(16) << 1.0 * length_counts[l] / n_spurious
         << setprecision(9) << setw(16) << 1.0 * n_length / n_spurious
         << setprecision(9) << setw(16) << 1.0 * excess_counts[l] / n_spurious
         << setprecision(9) << setw(16) << 1.0 * n_excess / n_spurious
         << endl;
  }

  cout << endl;
  cout << "x";
  for (uint64_t e{min_excess}; e != max_excess; ++e) {
    cout << "\t" << e;
  }
  cout << endl;
  cout.precision(4);
  // cout.setf(std::ios::scientific, std::ios::floatfield);
  for (uint64_t s{min_size}; s != max_size; ++s) {
    cout << s;
    for (uint64_t e{min_excess}; e != max_excess; ++e) {
      cout << "\t"
           << 1.0 * combined_counts[s - min_size][e - min_excess] / n_spurious;
    }
    cout << endl;
  }

  return 0;
} catch (Error & e) {
  cerr << "paa::Error:" << endl;
  cerr << e.what() << endl;
  return 1;
} catch (exception & e) {
  cerr << "std::exception" << endl;
  cerr << e.what() << endl;
  return 1;
} catch (...) {
  cerr << "unknown exception was caught" << endl;
  return 1;
}
```
