## Supplementary material for "G-Graph: An interactive genomic graph viewer": MUMdex Source Code for G-Graph: smooth_data.cpp

```
//
// smooth_data.cpp
//
// Perform lowess smoothing of x y dataset
//
// Copyright 2017 Peter Andrews @ CSHL
//

#include <fstream>
#include <iomanip>
#include <iostream>
#include <sstream>
#include <string>
#include <vector>
#include <climits>

#include "error.h"
#include "lowess.h"
#include "utility.h"

using std::cerr;
using std::cin;
using std::cout;
using std::endl;
using std::exception;
using std::ifstream;
using std::istringstream;
using std::ostringstream;
using std::setprecision;
using std::string;
using std::vector;

using paa::Error;

int main(int argc, char ** argv) {
  try {
    if (--argc != 1) throw Error("usage: smooth_data frac");

    const double frac{atof(argv[1])};

    vector<double> x;
    vector<double> y;

    double xv;
    double yv;
    while (cin >> xv >> yv) {
      x.push_back(xv);
      y.push_back(yv);
    }
    cerr << "Read " << x.size() << " data points" << endl;

    vector<double> smoothed(x.size());
    vector<double> resid(x.size());
    vector<double> weights(x.size());

    CppLowess::TemplatedLowess<std::vector<double>, double> Lowess;
    Lowess.lowess(x, y, frac, 10, 0.01, smoothed, resid, weights);

    cout << setprecision(12);
    for (unsigned int t{0}; t != x.size(); ++t) {
      cout << x[t] << " " << smoothed[t] << " " << y[t]
           << " " << y[t] / smoothed[t] << endl;
    }

    return 0;
  }
  catch(exception & e) {
    cerr << e.what() << endl;
    return 1;
  }
  catch(...) {
    cerr << "Some exception was caught." << endl;
    return 1;
  }
  return 0;
}
```
