## Supplementary material for "G-Graph: An interactive genomic graph viewer": MUMdex Source Code for G-Graph: snp_counts.cpp

```
//
// snp_counts.cpp
//
// Get base counts at a list of positions
//
// Copyright 2019 Peter Andrews @ CSHL
//

#include <algorithm>
#include <array>
#include <iostream>
#include <fstream>
#include <set>
#include <sstream>
#include <string>
#include <vector>

#include "bridges.h"
#include "error.h"
#include "mumdex.h"
#include "utility.h"

using std::array;
using std::cout;
using std::cerr;
using std::endl;
using std::ifstream;
using std::istringstream;
using std::max;
using std::max_element;
using std::set;
using std::sort;
using std::string;
using std::to_string;
using std::vector;

using paa::complement;
using paa::reverse_complement;
using paa::Bridge;
using paa::BridgeCounts;
using paa::Bridges;
using paa::ChromosomeIndexLookup;
using paa::Error;
using paa::Mappability;
using paa::MUM;
using paa::MUMindex;
// using paa::MUMdex;
using paa::OneBridgeInfo;
using paa::Pair;
using paa::Reference;

using MUMdex = paa::PreMappedMUMdex;
using MUMRegion = MUMdex::MUMRegion;

unsigned int base_to_int(const char b) {
  switch (b) {
    case 'A':
      return 0;
    case 'C':
      return 1;
    case 'G':
      return 2;
    case 'T':
      return 3;
    case 'N':
      return 4;
    default:
      throw Error("Unknown base to int") << b;
  }
}

char int_to_base(const unsigned int i) {
  switch (i) {
    case 0:
      return 'A';
    case 1:
      return 'C';
    case 2:
      return 'G';
    case 3:
      return 'T';
    case 4:
      return 'N';
    default:
      throw Error("Unknown int to base") << i;
  }
}

class CandInfo {
 public:
  class ParseError {};
  explicit CandInfo(const string & line) {
    istringstream input{line.c_str()};
    input >> chr >> pos;
    // pos += 1000;
    --pos;
    if (!input) throw ParseError{};
  }

  string chr{};
  unsigned int pos{};
};

int main(int argc, char ** argv)  try {
  // Check arguments
  if (--argc != 2) throw Error("usage: snp_counts position_list mumdex");

  // Process Arguments
  const string pos_file_name{argv[1]};
  ifstream pos_file{pos_file_name.c_str()};
  if (!pos_file) throw Error("Could not open file") << pos_file_name;

  const MUMdex mumdex{argv[2]};
  const Reference & ref{mumdex.reference()};
  const ChromosomeIndexLookup lookup{ref};
  const Mappability mappability{ref};

  // Read Input file into candidate objects
  string line;
  vector<CandInfo> info;
  // getline(pos_file, line);
  while (pos_file) {
    getline(pos_file, line);
    if (line.size()) {
      try {
        info.emplace_back(line);
      } catch (CandInfo::ParseError & e) {
        if (line[0] != 'c') {
          cerr << "parse error for line: " << line << endl;
        }
      }
    }
  }
  cerr << "Loaded " << info.size() << " positions" << endl;

  // Look for good reads to check for snp bridges
  const unsigned int read_length{151};
  for (const CandInfo & snp : info) {
    const unsigned int chr{lookup[snp.chr]};
    const unsigned int pos{snp.pos};
    const unsigned int start_pos{pos > read_length ? pos - read_length : 0};
    const unsigned int stop_pos{pos + 1};
    const MUMRegion region{mumdex.region(chr, start_pos, chr, stop_pos)};
    set<uint64_t> seen_pairs;
    array<unsigned int, 5> base_counts{{0, 0, 0, 0, 0}};
    for (const MUMindex index : region) {
      const uint64_t pair_index{index.pair_index()};
      const Pair pair{mumdex.pair(pair_index)};
      if (pair.dupe()) continue;
      const uint64_t mum_index{mumdex.mum_index(index)};
      const MUM mum{mumdex.mum(mum_index)};
      const int read_pos{mum.position0_to_read(pos)};
      if (read_pos < 0 ||
          read_pos >= static_cast<int>(pair.length(mum.read_2()))) continue;
      auto found = seen_pairs.find(pair_index);
      if (found != seen_pairs.end()) continue;
      uint64_t nearby_bases{0};
      array<unsigned int, 5> pair_base_counts{{0, 0, 0, 0, 0}};
      const array<std::string, 2> sequences(mumdex.sequences(pair_index));
      for (uint64_t m{mumdex.mums_start(pair_index)};
           m != mumdex.mums_stop(pair_index); ++m) {
        const MUM omum{mumdex.mum(m)};
        if (omum.chromosome() != chr) continue;
        if (omum.position0() > pos + read_length ||
            omum.position0() + omum.length() + read_length < pos) continue;
        nearby_bases += omum.length();
        const int oread_pos{mum.position0_to_read(pos)};
        if (oread_pos < 0 ||
            oread_pos >= static_cast<int>(pair.length(omum.read_2()))) continue;
        const char raw_base{sequences[omum.read_2()][oread_pos]};
        const char base{omum.flipped() ? complement(raw_base) : raw_base};
        ++pair_base_counts[base_to_int(base)];
      }
      if (nearby_bases < read_length) continue;
      auto max_iter = max_element(pair_base_counts.begin(),
                                   pair_base_counts.end());
      auto max_index = max_iter - pair_base_counts.begin();
      bool good{true};
      for (int64_t i{0}; i != pair_base_counts.size(); ++i) {
        if (i == max_index) continue;
        if (pair_base_counts[i] == pair_base_counts[max_index]) {
          good = false;
          break;
        }
      }
      if (!good) continue;
      seen_pairs.insert(pair_index);
      ++base_counts[max_index];
    }
    cout << snp.chr
         << '\t' << pos + 1;
    for (uint64_t b{0}; b != base_counts.size(); ++b) {
      cout << '\t' << base_counts[b];
    }
    sort(base_counts.begin(), base_counts.begin() + 4);
    const uint64_t total{base_counts[2] + base_counts[3]};
    const double proportion{total ? 1.0 * base_counts[2] / total : 1.0};
    cout << '\t' << total << '\t' << proportion << endl;
  }

  cerr << "done" << endl;

  return 0;
} catch (Error & e) {
  cerr << "paa::Error:" << endl;
  cerr << e.what() << endl;
  return 1;
} catch(std::exception & e) {
  cerr << e.what() << '\n';
  return 1;
} catch(...) {
  cerr << "Some exception was caught.\n";
  return 1;
}
```
