## Supplementary material for "G-Graph: An interactive genomic graph viewer": MUMdex Source Code for G-Graph: snp_hists.cpp

```
//
// snp_hists
//
// read in snp_counts tables, plot hists
//
// Copyright 2019 Peter Andrews @ CSHL
//

#include <algorithm>
#include <array>
#include <exception>
#include <iostream>
#include <fstream>
#include <sstream>
#include <string>
#include <utility>
#include <vector>

#include "error.h"
#include "psplot.h"

using std::array;
using std::cerr;
using std::cin;
using std::cout;
using std::endl;
using std::exception;
using std::ifstream;
using std::istringstream;
using std::pair;
using std::sort;
using std::string;
using std::to_string;
using std::vector;

using paa::Bounds;
using paa::Error;
using paa::PSDoc;
using paa::PSHSeries;
using paa::PSPage;

int main(int argc, char* argv[])  try {
  if (--argc < 3) throw Error("snp_hists out_name in_pos input ...");
  const string out_name{argv[1]};
  const string pos_name{argv[2]};
  argc -= 2;
  argv += 2;

  using Counts = array<unsigned int, 5>;
  using PosCounts = vector<Counts>;
  using AllCounts = vector<PosCounts>;
  string chr;
  unsigned int pos;
  unsigned int a;
  unsigned int c;
  unsigned int g;
  unsigned int t;
  unsigned int n;
  unsigned int tot;
  double r;
  string ref_allele;

  // Read input positions and determine minor allele
  vector<unsigned int> minor_alleles;
  ifstream pos_file{pos_name.c_str()};
  if (!pos_file) throw Error("Problem opening input pos") << pos_name;
  while (pos_file >> chr >> pos >> ref_allele >> a >> c >> g >> t) {
    vector<pair<unsigned int, unsigned int>> alleles;
    alleles.emplace_back(a, 0);
    alleles.emplace_back(c, 1);
    alleles.emplace_back(g, 2);
    alleles.emplace_back(t, 3);
    sort(alleles.begin(), alleles.end());
    minor_alleles.push_back(alleles[2].second);
  }

  // Read input files
  vector<string> names;
  AllCounts counts;
  for (int arg{0}; arg != argc; ++arg) {
    const string in_name{argv++[1]};
    ifstream in_file{in_name.c_str()};
    if (!in_file) throw Error("Problem opening input") << in_name;
    uint64_t p{0};
    while (in_file >> chr >> pos >> a >> c >> g >> t >> n >> tot >> r) {
      if (arg == 0) {
        counts.emplace_back(0);
        names.emplace_back(chr + " " + to_string(pos));
      }
      counts[p++].emplace_back(Counts{{a, c, g, t, a + c + g + t}});
    }
  }
  if (names.size() != minor_alleles.size())
    throw Error("names and minor size mismatch")
        << names.size() << minor_alleles.size();

  // Get totals
  for (uint64_t p{0}; p != counts.size(); ++p) {
    const string name{names[p]};
    const PosCounts & pos_counts{counts[p]};
    Counts totals{{0, 0, 0, 0, 0}};
    for (const Counts & count : pos_counts) {
      for (unsigned int i{0}; i != 5; ++i) {
        totals[i] += count[i];
      }
    }
    cout << name;
    for (unsigned int i{0}; i != 5; ++i) cout << '\t' << totals[i];
    cout << endl;
  }

  // Make hists
  PSDoc plots{out_name};
  plots.pdf(true);

  using Hist = PSHSeries<double, unsigned int>;
  Hist & all_hist = *Hist::create(plots, out_name + ";Minor / Total;N",
                                  Bounds{0.0, 1.1}, 110u);
  for (uint64_t p{0}; p != counts.size(); ++p) {
    const string & name{names[p]};
    PosCounts & pos_counts{counts[p]};
    Hist & hist = *Hist::create(plots,
                                out_name + " " + name + ";Minor / Total;N",
                                Bounds{0.0, 1.1}, 110u);
    const unsigned int minor{minor_alleles[p]};
    for (Counts & cts : pos_counts) {
      if (cts[4] > 2) {
        hist.add_point(1.0 * cts[minor] / cts[4]);
        all_hist.add_point(1.0 * cts[minor] / cts[4]);
      }
    }
  }

  return 0;
} catch (Error & e) {
  cerr << "paa::Error:" << endl;
  cerr << e.what() << endl;
  return 1;
} catch (exception & e) {
  cerr << "std::exception" << endl;
  cerr << e.what() << endl;
  return 1;
} catch (...) {
  cerr << "unknown exception was caught" << endl;
  return 1;
}
```
