## Supplementary material for "G-Graph: An interactive genomic graph viewer": MUMdex Source Code for G-Graph: snp_report.cpp

```
//
// snp_report.cpp
//
// Copyright 2017 Peter Andrews @ CSHL
//

#include <algorithm>
#include <iostream>
#include <fstream>
#include <map>
#include <set>
#include <sstream>
#include <string>
#include <vector>

#include "bridges.h"
#include "error.h"
#include "mumdex.h"
#include "utility.h"

using std::cout;
using std::cerr;
using std::endl;
using std::ifstream;
using std::istringstream;
using std::map;
using std::max;
using std::set;
using std::string;
using std::to_string;
using std::vector;

using paa::complement;
using paa::reverse_complement;
using paa::sout;
using paa::Bridge;
using paa::BridgeCounts;
using paa::Bridges;
using paa::ChromosomeIndexLookup;
using paa::Error;
using paa::Mappability;
using paa::MUM;
using paa::MUMdex;
using paa::Reference;

// using MUMdex = paa::PreMappedMUMdex;

unsigned int base_to_int(const char b) {
  switch (b) {
    case 'A':
      return 0;
    case 'C':
      return 1;
    case 'G':
      return 2;
    case 'T':
      return 3;
    default:
      throw Error("Unknown base to int") << b;
  }
}

char int_to_base(const unsigned int i) {
  switch (i) {
    case 0:
      return 'A';
    case 1:
      return 'C';
    case 2:
      return 'G';
    case 3:
      return 'T';
    default:
      throw Error("Unknown int to base") << i;
  }
}

void parse_counts(const string & in_string, unsigned int vals[4]) {
  istringstream in{in_string.c_str()};
  for (const unsigned int i : {0, 1, 2, 3}) {
    in >> vals[i];
    in.get();
  }
}

class CandInfo {
 public:
  class ParseError {};
  explicit CandInfo(const string & line_) : line{line_} {
    istringstream input{line.c_str()};
    input >> chrA >> posA >> highA >> chrB >> posB >> highB
          >> invariant >> offset;
    if (!input) {
      throw ParseError{};
    }
  }

  string chrA{};
  unsigned int posA{};
  bool highA{};
  string chrB{};
  unsigned int posB{};
  bool highB{};
  int invariant{};
  int offset{};
  string line{};
};


int main(int argc, char ** argv)  try {
  // Check arguments
  if (--argc < 3)
    throw Error("usage: snp_report count_dupes bridge_list mumdexes...");

  // Process Arguments
  const bool count_dupes{static_cast<bool>(atoi(argv[1]))};
  const string cand_file_name{argv[2]};
  ifstream cand_file{cand_file_name.c_str()};
  if (!cand_file) throw Error("Could not open file") << cand_file_name;

  // Read Input file into candidate objects
  string line;
  vector<CandInfo> info;
  while (cand_file) {
    getline(cand_file, line);
    if (line.size()) {
      try {
        const CandInfo i{line};
        info.push_back(i);
      } catch (CandInfo::ParseError & e) {
        if (line[0] != 'c') {
          cerr << "parse error for line: " << line << endl;
        }
      }
    }
  }
  cerr << "Loaded " << info.size() << " events" << endl;

  argc -= 2;
  argv += 2;

  // Load mumdexes
  vector<MUMdex> mumdexes;
  mumdexes.reserve(argc);
  while (argc) {
    mumdexes.emplace_back(argv[1]);
    --argc;
    ++argv;
  }
  const Reference & ref{mumdexes.front().reference()};
  const ChromosomeIndexLookup lookup{ref};
  const Mappability mappability{ref};

  const vector<string> bases{"A", "C", "G", "T"};

  const vector<string> mumdex_labels{[&mumdexes]() {
      vector<string> result;
      for (unsigned int i{0}; i != mumdexes.size(); ++i) {
        result.push_back(to_string(i + 1));
      }
      return result;
    }()};

  if (0) {
    // Header line
    cout << "chrA\tposA\thighA\tchrB\tposB\thighB\tinv\toff";
    const vector<string> counts{"cc", "ac", "rc", "bc", "br"};
    const string anchor{"AB"};
    const vector<string> members{"141", "149", "267", "268"};
    for (const bool ab : {false, true}) {
      for (const unsigned int c : {0, 1, 2, 3, 4}) {
        for (unsigned int m{0}; m != mumdexes.size(); ++m) {
          if (ab || c < 3) {
            cout << "\t" << members[m] << counts[c];
            if (c < 3) cout << anchor[ab];
          }
        }
      }
    }
    cout << endl;
  }

  cout << "chrA\tposA\thighA\tchrB\tposB\thighB\tinv\toff\tref\talt\trt\trcrt";
  for (unsigned int b{0}; b != bases.size(); ++b) {
    for (unsigned int m{0}; m != mumdexes.size(); ++m) {
      cout << "\tbc" << bases[b] << "-" << mumdex_labels[m];
    }
  }
  for (unsigned int m{0}; m != mumdexes.size(); ++m) {
    cout << "\tar" << "-" << mumdex_labels[m];
  }
  for (unsigned int b{0}; b != bases.size(); ++b) {
    for (unsigned int m{0}; m != mumdexes.size(); ++m) {
      cout << "\tcc" << bases[b] << "-" << mumdex_labels[m];
    }
  }
  for (unsigned int m{0}; m != mumdexes.size(); ++m) {
    cout << "\taac" << "-" << mumdex_labels[m];
  }
  for (unsigned int b{0}; b != bases.size(); ++b) {
    for (unsigned int m{0}; m != mumdexes.size(); ++m) {
      cout << "\tmac" << bases[b] << "-" << mumdex_labels[m];
    }
  }
  cout << endl;


  for (const CandInfo & i : info) {
    // Check that candidate is a snp
    if (i.highA != true || i.highB != false ||
        i.invariant != 0 || i.offset != 2)
      throw Error("Candidate is not a simple SNP:") << i.line;

    const string chrs[2]{i.chrA, i.chrB};
    const unsigned int chr[2]{lookup[i.chrA], lookup[i.chrB]};
    const unsigned int pos[2]{i.posA, i.posB};
    const unsigned int high[2]{i.highA, i.highB};

    const vector<BridgeCounts> new_bridge_counts{
      [&mumdexes, &ref, &mappability, &i, chr, pos, high, count_dupes]() {
        vector<BridgeCounts> result;
        for (unsigned int m{0}; m != mumdexes.size(); ++m) {
          const MUMdex & mumdex{mumdexes[m]};
          result.emplace_back(mumdex, ref, mappability,
                              chr, pos, high, i.offset, count_dupes);
        }
        return result;
      }()};

    for (const bool b : {false, true}) {
      cout << chrs[b] << "\t" << pos[b] << "\t" << high[b] << "\t";
    }
    const char ref_base{ref[chr[0]][pos[0] + 1]};
    string ref_string;
    ref_string += ref_base;
    cout << i.invariant << "\t" << i.offset << "\t" << ref_string;
    vector<unsigned int> total_val(mumdexes.size());
    vector<unsigned int> total_allele(bases.size());
    vector<vector<unsigned int>> values(
        mumdexes.size(), vector<unsigned int>(bases.size()));
    for (unsigned int b{0}; b != bases.size(); ++b) {
      const string this_base{bases[b]};
      for (unsigned int m{0}; m != mumdexes.size(); ++m) {
        unsigned int val{0};
        if (!new_bridge_counts[m].bridge.allele_count.count(this_base)) {
          if (this_base == ref_string) {
            val = new_bridge_counts[m].bridge.ref_count;
          }
        } else {
          if (this_base == ref_string)
            throw Error("Unexpected reference count");
          val = new_bridge_counts[m].bridge.allele_count.at(this_base);
        }
        total_allele[base_to_int(this_base[0])] += val;
        total_val[m] += val;
        values[m][b] = val;
      }
    }

    unsigned int alt_base{3 - base_to_int(ref_base)};
    for (unsigned int b{0}; b != bases.size(); ++b) {
      const string this_base{bases[b]};
      if (this_base != ref_string &&
          total_allele[b] >= total_allele[alt_base]) {
        alt_base = b;
      }
    }

    const string ref_triplet{ref.subseq(chr[0], pos[0], pos[0] + 3)};
    cout << "\t" << int_to_base(alt_base)
         << "\t" << ref_triplet
         << "\t" << reverse_complement(ref_triplet);

    for (unsigned int b{0}; b != bases.size(); ++b) {
      for (unsigned int m{0}; m != mumdexes.size(); ++m) {
        cout << "\t" << values[m][b];
      }
    }

    for (unsigned int m{0}; m != mumdexes.size(); ++m) {
      cout << "\t"
           << (total_val[m] ? 1.0 * values[m][alt_base] / total_val[m] : 0);
    }
    if (0) {
      for (unsigned int m{0}; m != mumdexes.size(); ++m) {
        const string alt_string{bases[alt_base]};
        if (new_bridge_counts[m].any[0].allele_count.count(alt_string)) {
          cout << "\t"
               << new_bridge_counts[m].any[0].allele_count.at(alt_string);
        } else {
          cout << "\t0";
        }
      }
    }
    for (unsigned int b{0}; b != bases.size(); ++b) {
      const string this_base{bases[b]};
      for (unsigned int m{0}; m != mumdexes.size(); ++m) {
        if (new_bridge_counts[m].any[0].allele_count.count(this_base) !=
            new_bridge_counts[m].any[1].allele_count.count(this_base)) {
          throw Error("Inconsistent any count 0");
        }
        if (new_bridge_counts[m].any[0].allele_count.count(this_base) &&
            new_bridge_counts[m].any[1].allele_count.count(this_base) &&
            new_bridge_counts[m].any[0].allele_count.at(this_base) !=
            new_bridge_counts[m].any[1].allele_count.at(this_base)) {
          throw Error("Inconsistent any count");
        }

        const unsigned int total_count{
          (new_bridge_counts[m].any[0].allele_count.count(this_base) ?
           new_bridge_counts[m].any[0].allele_count.at(this_base) : 0) +
          (new_bridge_counts[m].any[1].allele_count.count(this_base) ?
           new_bridge_counts[m].any[1].allele_count.at(this_base) : 0)};
        cout << "\t" << total_count / 2;
      }
    }
    for (unsigned int m{0}; m != mumdexes.size(); ++m) {
      const string alt_string{bases[alt_base]};
      cout << "\t" << std::max(
          new_bridge_counts[m].anchor[0].get_allele_count(alt_string),
          new_bridge_counts[m].anchor[1].get_allele_count(alt_string));
    }
    for (unsigned int b{0}; b != bases.size(); ++b) {
      const string this_base{bases[b]};
      for (unsigned int m{0}; m != mumdexes.size(); ++m) {
        cout << "\t" << std::max(
            new_bridge_counts[m].anchor[0].get_allele_count(this_base),
            new_bridge_counts[m].anchor[1].get_allele_count(this_base));
      }
    }
    cout << endl;
  }

  cerr << "done" << endl;

  return 0;
} catch (Error & e) {
  cerr << "paa::Error:" << endl;
  cerr << e.what() << endl;
  return 1;
} catch(std::exception & e) {
  cerr << e.what() << '\n';
  return 1;
} catch(...) {
  cerr << "Some exception was caught.\n";
  return 1;
}
```
