## Supplementary material for "G-Graph: An interactive genomic graph viewer": MUMdex Source Code for G-Graph: split_bed.pl

```
#! /usr/bin/perl -w

# Copyright 2015 Peter Andrews @ CSHL

use strict;

my $n_lines = $ARGV[0];

while (<STDIN>) {
    my ($chr, $start, $stop) = split;
    my $length = $stop - $start;
    if ($length <= $n_lines) {
        print "$chr\t$start\t$stop\n";
    } else {
        for (my $current = $start; $current < $stop; $current += $n_lines) {
            if ($current + 2 * $n_lines > $stop) {
                print "$chr\t$current\t$stop\n";
                last;
            } else {
                my $next = $current + $n_lines;
                print "$chr\t$current\t$next\n";                
            }
        }
    }
}
```
