## Supplementary material for "G-Graph: An interactive genomic graph viewer": MUMdex Source Code for G-Graph: strings.cpp

```
//
// strings.cpp
//
// useful string functions
//
// Copyright 2015 Peter Andrews @ CSHL
//

#include "strings.h"

#include <algorithm>
#include <string>

namespace paa {

std::string to_lower(const std::string & up) {
  std::string lower(up.size(), 'X');
  std::transform(up.begin(), up.end(), lower.begin(), ::tolower);
  return lower;
}
std::string to_upper(const std::string & low) {
  std::string upper(low.size(), 'X');
  std::transform(low.begin(), low.end(), upper.begin(), ::toupper);
  return upper;
}

void remove(std::string & input, const std::string & search) {
  const size_t pos = input.find(search);
  if (pos != std::string::npos) input.erase(pos, search.size());
}

}  // namespace paa
```
