## Supplementary material for "G-Graph: An interactive genomic graph viewer": MUMdex Source Code for G-Graph: submit_denovos.sh

```
#! /bin/bash

n_threads="$1"
if [ "$n_threads" = "" ] ; then
    n_threads=5
fi

bed=~/hg38/hg38.1M.bed
if [ "$2" != "" ] ; then
    bed="$2"
fi

out=out
if [ "$3" != "" ] ; then
    out="$3"
fi

n_jobs_max=2000 &&
n_jobs=$(jobids.sh C) &&
tac $bed |
while read chr start stop ; do
    dir=$out/$chr/$start-$stop
    name=$chr.$start
    if [ ! -e $dir ] ; then
        n_jobs=$((n_jobs + 1))
        if [ $n_jobs -le $n_jobs_max ] ; then
            mkdir -p $dir
            (
                cd $dir
                echo submitting job $n_jobs of $n_jobs_max in $dir &&
                qsub -N $name -l h_vmem=40G -pe smp $n_threads \
                    -cwd -o out.txt -e err.txt \
                    ~/mumdex/unstable/run_denovos.sh ~/hg38/hg38.fa \
                    ~/ssc_hg38/{families_2380.txt,bridges,samples} \
                    $chr $start $stop $n_threads > /dev/null ||
                error qsub failed
            ) || exit 1
        else
            echo max queue length has been reached
            break
        fi
    fi
done
```
