## Supplementary material for "G-Graph: An interactive genomic graph viewer": MUMdex Source Code for G-Graph: submit_samples.sh

```
#! /bin/bash

# setup shell environment
. ~/mumdex/utility/shell.sh

#
# process command line arguments
#

# lists of bam files to process
bam_list=(${1:-bams.txt})
for list in "${bam_list[@]}" ; do
    [ -e "$list" ] || error bam list $list not found
done

# number of threads to use
n_threads=${2:-10}
[ "$n_threads" -gt 0 ] || error bad n_threads $n_threads

# prefix name of the run - must start with a letter for sge
run_name=${3:-m}

# maximum number of jobs to put in queue
n_jobs_max=${4:-2000}
[ "$n_jobs_max" -gt 0 ] || error bad n_jobs_max $n_jobs_max

# Run jobs
n_jobs=$(jobids.sh C) &&
cat "${bam_list[@]}" | while read bam ; do
    size=$( # unused right now
        ls -l $bam |
        awk '{
size=int($5 * 1.2/1024/1024/1024);
printf "%d", (size > 130 ? size : 130);
}'
    )
    size=150 # usually good
    sample=$(basename $bam)
    sample=${sample%%.*}
    outdir=$PWD/samples/$sample
    if [ ! -e $outdir ] ; then 
        n_jobs=$((n_jobs + 1))
        if [ $n_jobs -le $n_jobs_max ] ; then
            (
                mkdir -p $outdir &&
                cd $outdir &&
                echo submitting job $n_jobs of $n_jobs_max for sample $sample &&
                qsub -N $run_name$sample -l h_vmem=${size}G -pe smp $n_threads \
                    -cwd -o out.txt -e err.txt ~/mumdex/unstable/run_sample.sh \
                    $bam $outdir $n_threads > /dev/null ||
                error qsub failed
            ) || exit 1
        else
            echo max queue length has been reached
            break
        fi
    fi
done
```
