## Supplementary material for "G-Graph: An interactive genomic graph viewer": MUMdex Source Code for G-Graph: subsample.cpp

```
//
// subsample
//
// subsample input
//
// Copyright 2014 Peter Andrews @ CSHL
//

#include <algorithm>
#include <exception>
#include <iostream>
#include <fstream>
#include <functional>
#include <random>
#include <string>
#include <vector>

#include "error.h"

using std::bind;
using std::cout;
using std::cerr;
using std::endl;
using std::exception;
using std::ifstream;
using std::mt19937_64;
using std::shuffle;
using std::string;
using std::vector;
using std::uniform_int_distribution;
using std::uniform_real_distribution;

using paa::Error;

int main(int argc, char* argv[], char * []) try {
  if (--argc < 2) throw Error("usage: subsample N input_file ...");

  vector<string> reservoir;
  const uint64_t N = atol((++argv)[0]);

  reservoir.reserve(N);
  uint64_t n = 0;

  std::random_device rd;
  auto mersenne = mt19937_64(rd());
  auto realGen = bind(uniform_real_distribution<double>(0.0, 1.0), mersenne);
  auto intGen = bind(uniform_int_distribution<uint64_t>(0, N - 1), mersenne);

  while (--argc) {
    const string input_file_name((++argv)[0]);
    ifstream input_file(input_file_name.c_str());
    if (!input_file) throw Error("Could not open file for input:")
                         << input_file_name;
    string line;
    while (getline(input_file, line) && input_file) {
      ++n;
      if (N == 0) {
        cout << line << endl;
      } else {
        if (reservoir.size() < N) {
          reservoir.push_back(line);
        } else {
          if (realGen() < 1.0 * N / n) {
            reservoir[intGen()] = line;
          }
        }
        if (n == N) shuffle(reservoir.begin(), reservoir.end(), mersenne);
      }
    }
  }
  if (N == 0) {
    cerr << "Output all input since N was 0" << endl;
    return 0;
  }
  if (n < N) {
    shuffle(reservoir.begin(), reservoir.end(), mersenne);
    cerr << "Not enough lines in subsample: only "
         << n << " of " << N << endl;
  }
  for (const string & line : reservoir) {
    cout << line << endl;
  }
  // cerr << "done" << endl;
  return 0;
} catch (Error & e) {
  cerr << "paa::Error:" << endl;
  cerr << e.what() << endl;
  return 1;
} catch (exception & e) {
  cerr << "std::exception" << endl;
  cerr << e.what() << endl;
  return 1;
} catch (...) {
  cerr << "unknown exception was caught" << endl;
  return 1;
}
```
