## Supplementary material for "G-Graph: An interactive genomic graph viewer": MUMdex Source Code for G-Graph: table_stats.cpp

```
//
// table_stats
//
// calculate averages and other stats over named columns of data
//
// Copyright 2018 Peter Andrews @ CSHL
//

#include <algorithm>
#include <exception>
#include <iostream>
#include <sstream>
#include <string>
#include <vector>

#include "error.h"
#include "stats.h"
#include "utility.h"

using std::cerr;
using std::cin;
using std::cout;
using std::endl;
using std::exception;
using std::istringstream;
using std::string;
using std::vector;

using paa::dne;
using paa::Error;
using paa::MAD;
using paa::NormalParams;

int main(int argc, char**)  try {
  --argc;
  if (argc != 0) throw Error("data_file | table_stats");

  string line;
  getline(cin, line);
  istringstream header_stream{line.c_str()};
  vector<string> header;
  while (header_stream >> line) header.push_back(line);
  vector<vector<double>> data(header.size());
  vector<unsigned int> has_zero(header.size());
  vector<unsigned int> has_neg(header.size());
  // vector<unsigned int> is_numeric(header.size());

  while (getline(cin, line)) {
    istringstream line_stream{line.c_str()};
    for (uint64_t c{0}; c != data.size(); ++c) {
      vector<double> & vals{data[c]};
      double input;
      line_stream >> input;
      if (!line_stream) throw Error("Bad input from line") << line;
      if (!(input > 0 || input < 0)) has_zero[c] = true;
      if (input < 0) has_neg[c] = true;
      vals.push_back(input);
    }
    if (!line_stream) throw Error("Parse Error");
  }

  auto report = [](const string & name, const vector<double> & vec) {
    const NormalParams norm{vec};
    const MAD mad{vec};
    cout << name
         << '\t' << vec.size()
         << '\t' << vec.front()
         << '\t' << vec.back()
         << '\t' << norm.mean
         << '\t' << norm.stdev
         << '\t' << norm.skew
         << '\t' << mad.median()
         << '\t' << mad.mad()
         << '\n';
  };

  cout << "column_name\tn\tmin\tmax\taverage\tstdev\tskew\tmedian\tmad\n";
  for (uint64_t c{0}; c != data.size(); ++c) {
    vector<double> vals{data[c]};
    sort(vals.begin(), vals.end());
    report(header[c], vals);
    if (has_zero[c]) {
      vector<double> nonzeros;
      for (const double val : vals)
        if (val > 0 || val < 0) nonzeros.push_back(val);
      if (nonzeros.size() && dne(nonzeros.front(), nonzeros.back()))
        report(header[c] + "_nonzeros", nonzeros);
    }
    if (has_neg[c]) {
      vector<string> tags{"_negatives", "_positives"};
      for (const bool pos : {false, true}) {
        vector<double> posneg;
        for (const double val : vals)
          if ((pos && val > 0) || (!pos && val < 0)) posneg.push_back(val);
        if (posneg.size() && dne(posneg.front(), posneg.back()))
          report(header[c] + tags[pos], posneg);
      }
    }
  }

  return 0;
} catch (Error & e) {
  cerr << "paa::Error:" << endl;
  cerr << e.what() << endl;
  return 1;
} catch (exception & e) {
  cerr << "std::exception" << endl;
  cerr << e.what() << endl;
  return 1;
} catch (...) {
  cerr << "unknown exception was caught" << endl;
  return 1;
}
```
