## Supplementary material for "G-Graph: An interactive genomic graph viewer": MUMdex Source Code for G-Graph: test_hmm.cpp

```
//
// test_hmm
//
// Test the hmm module
//
// Copyright 2018 Peter Andrews @ CSHL
//

#include <exception>
#include <iostream>
#include <functional>
#include <limits>
#include <random>
#include <string>
#include <vector>

#include "error.h"
#include "hmm.h"

using std::bind;
using std::cout;
using std::cerr;
using std::endl;
using std::exception;
using std::function;
using std::mt19937_64;
using std::random_device;
using std::string;
using std::uniform_real_distribution;
using std::vector;

using paa::Error;
using paa::HMM;
using paa::HMMparams;

int main(int argc, char * argv[]) try {
  if (0) std::cerr << argc << " " << argv[0] << std::endl;

  const double p_tails_1{0.95};
  const double p_tails_2{0.45};
  const vector<unsigned int> observations{[p_tails_1, p_tails_2]() {
      const unsigned int factor{100};
      const unsigned int n_coin_1{factor * 1000};
      const unsigned int n_coin_2{factor * 500};
      const unsigned int n_coin_1_again{factor * 500};
      random_device rd{};
      mt19937_64 mersenne{rd()};
      function<double()> gen{bind(uniform_real_distribution<double>(0, 1),
                                  mersenne)};
      vector<unsigned int> result;
      for (unsigned int t{0}; t != n_coin_1; ++t) {
        result.push_back(gen() < p_tails_1);
      }
      for (unsigned int t{0}; t != n_coin_2; ++t) {
        result.push_back(gen() < p_tails_2);
      }
      for (unsigned int t{0}; t != n_coin_1_again; ++t) {
        result.push_back(gen() < p_tails_1);
      }
      return result;
    }()};

  if (false) {
    for (unsigned int t{0}; t != observations.size(); ++t) {
      cout << " " << observations[t];
    }
    cout << endl;
  }

  HMMparams trial_params{2, 2};
  trial_params.initial_probs()[1] = 0.0000001;
  trial_params.initial_probs()[0] = 1 - trial_params.initial_probs()[1];

  trial_params.transitions()[0][1] = 0.0000001;
  trial_params.transitions()[0][0] = 1 - trial_params.transitions()[0][1];
  trial_params.transitions()[1][1] = trial_params.transitions()[0][0];
  trial_params.transitions()[1][0] = trial_params.transitions()[0][1];

  trial_params.emissions()[0][1] = p_tails_1;
  trial_params.emissions()[0][0] = 1 - trial_params.emissions()[0][1];
  trial_params.emissions()[1][1] = p_tails_2;
  trial_params.emissions()[1][0] = 1 - trial_params.emissions()[1][1];

  HMM hmm{trial_params, observations};
  hmm.baum_welch();

  return 0;
} catch (Error & e) {
  cerr << "paa::Error:" << endl;
  cerr << e.what() << endl;
  return 1;
} catch (exception & e) {
  cerr << "std::exception" << endl;
  cerr << e.what() << endl;
  return 1;
} catch (...) {
  cerr << "unknown exception was caught" << endl;
  return 1;
}
```
