## Supplementary material for "G-Graph: An interactive genomic graph viewer": MUMdex Source Code for G-Graph: test_mumdex.cpp

```
//
// test_mumdex
//
// test the MUMdex format
//
// Copyright Peter Andrews 2015 @ CSHL
//

#include <array>
#include <exception>
#include <functional>
#include <iostream>
#include <random>
#include <string>
#include <vector>

#include "error.h"
#include "mumdex.h"
#include "utility.h"

using std::array;
using std::bind;
using std::cout;
using std::cerr;
using std::endl;
using std::exception;
using std::mt19937_64;
using std::string;
using std::uniform_int_distribution;
using std::vector;

using paa::ClassInfo;
using paa::Pair;
using paa::MUM;
using paa::MUMindex;
using paa::SplitBases;
using paa::Error;
using paa::TExtraBases;
using paa::TAllBases;
using paa::MUMdex;
using paa::OldMappedVector;
using paa::GrowingVector;
using paa::tout;
using paa::terr;

std::random_device rd;
auto eng = mt19937_64(rd());
auto gen = bind(uniform_int_distribution<uint64_t>(0, 10), eng);
auto baseGen = bind(uniform_int_distribution<uint64_t>(1, 5), eng);

string random_bases(const unsigned int n) {
  string result;
  for (unsigned int i = 0; i != n; ++i) {
    result += paa::base_chars[baseGen()];
  }
  return result;
}

int main(int argc, char * argv[]) try {
  if (0) {
    ClassInfo<Pair>();
    ClassInfo<MUM>();
    ClassInfo<SplitBases>();
    ClassInfo<MUMindex>();
    ClassInfo<TExtraBases<GrowingVector>>();
    ClassInfo<TAllBases<GrowingVector>>();
    return 0;
  }

  if (--argc != 1) throw Error("usage: test_mumdex mumdex_name");
  const string mumdex_name{argv[1]};

  std::cerr << "loading reconstruction" << std::endl;
  const MUMdex mumdex(mumdex_name);
  std::cerr << "decoding final mumdex" << std::endl;
  for (unsigned int i = 0; i != mumdex.n_pairs(); ++i) {
    const std::array<std::string, 2> sequences(mumdex.sequences(i));
    for (const auto & sequence : sequences) {
      std::cout << sequence << std::endl;
    }
  }
  return 0;
  const auto & ref = mumdex.reference();
  if (1) {
    terr << "chr" << "position" << "readpos" << "flipped"
         << "offset" << "length" << "last" << endl;
    for (const auto index : mumdex.index()) {
      const auto pair = mumdex.pair(index.pair_index());
      const auto mum = mumdex.mum(index);
      terr << ref.name(mum.chromosome()) << mum.position0()
           << mum.read_position0(pair.length(mum.read_2()))
           << mum.flipped() << mum.offset() << mum.length()
           << mum.last_hit() << endl;
    }
  }

  if (0) {
    terr << "pair" << "read" << "dupe" << "bad" << "length"
         << "chr" << "position" << "readpos" << "flipped"
         << "offset" << "length" << "last" << "seq" << endl;
    for (uint64_t p = 0; p != mumdex.n_pairs(); ++p) {
      const auto pair = mumdex.pair(p);
      const auto seqs = mumdex.sequences(p);
      auto mum = mumdex.mums_begin(p);
      bool output_mum[2]{false, false};
      for (const bool r : {false, true}) {
        const auto seq = seqs[r];
        while (mum <= mumdex.mums_end(p)) {
          if ((mum == mumdex.mums_end(p) || !mum->read(r)) && output_mum[r])
            break;
          terr << p << r << pair.dupe() << pair.bad(r) << pair.length(r);
          if (mum == mumdex.mums_end(p) || !mum->read(r)) {
            terr << seq << endl;
            break;
          }
          terr << ref.name(mum->chromosome()) << mum->position0()
               << mum->read_position0(pair.length(r))
               << mum->flipped() << mum->offset() << mum->length()
               << mum->last_hit() << seq << endl;
          ++mum;
          output_mum[r] = true;
        }
      }
    }
  }

#if 0
  if (1) {
    for (auto mum = mumdexs[0].mums_begin(); mum != mumdexs[0].mums_end();
         ++mum) {
      cerr << ref.name(mum->chromosome()) << '\t' << mum->position() << '\t'
           << mum->read_2() << endl;
    }
  } else {
    const auto & mumdex = mumdexs[0];
    for (uint64_t m = 0; m != mumdex.mums_size(); ++m) {
      const auto mum = mumdex.sorted_mum(m);
      cerr << ref.name(mum.chromosome()) << '\t' << mum.position() << '\t'
           << mum.read_2() << endl;
    }
  }

  return 0;

  const string vec_name{"test.txt"};
  {
    OldMappedVector<int> test;
    test.push_back(1);
    test.push_back(2);
    test.emplace_back(3);
    test.push_back(4);
    test.push_back(5);
    test.emplace_back(6);
    test.push_back(7);
    test.emplace_back(8);
    for (unsigned int i = 0; i != 10000; ++i) {
      test.push_back(i);
    }
    test.save(vec_name);
  }
  OldMappedVector<int> test(vec_name);
  for (unsigned int i = 0; i != test.size(); ++i) {
    if (i < 10) cout << i << " " << test[i] << endl;
  }

  const unsigned int n_test_sequences = 10000;
  vector<string> originals;
  {
    TAllBases<GrowingVector> all;
    for (unsigned int i = 0; i != n_test_sequences; ++i) {
      string bases = random_bases(100 * gen());
      originals.push_back(bases);
      all.push_back(bases);
    }
    cerr << "stored info" << endl;
    all.save("testb.bases.bin", "testb.bases.extra.bin");
    cerr << "saved info" << endl;
  }

  TAllBases<OldMappedVector> all("testb");
  for (unsigned int i = 0; i != n_test_sequences; ++i) {
    const string retrieved = all.bases(i);
    if (retrieved != originals[i]) {
      cerr << "bases " << i << " do not agree" << endl;
      cerr << "  " << originals[i] << endl;
      cerr << "  " << retrieved << endl;
    }
  }
  cerr << "retrieved and verified info" << endl;
#endif

  return 0;
} catch (Error & e) {
  cerr << "paa::Error:" << endl;
  cerr << e.what() << endl;
  return 1;
} catch (exception & e) {
  cerr << "std::exception" << endl;
  cerr << e.what() << endl;
  return 1;
} catch (...) {
  cerr << "unknown exception was caught" << endl;
  return 1;
}
```
