## Supplementary material for "G-Graph: An interactive genomic graph viewer": MUMdex Source Code for G-Graph: test_mumdex.sh

```
#! /bin/bash

PATH=/data/software/samtools/samtools-0.1.16:$PATH

test=$(which count_anchors)
if [ "$test" = "" ] ; then
    echo The mumdex directory must be compiled and be in your path 1>&2    
    exit 1
fi

ref=$1
if [ "$ref" = "" ] ; then
    echo You must pass a reference fasta as the first argument 1>&2
    exit 1
fi
if [ ! -e "$ref" ] ; then
    echo The reference fasta $ref was not found 1>&2
    exit 1
fi
if [ -e "$ref.bin/" ] && [ ! -e $ref.bin/ref.seq.bin ] ; then
    echo Looks like an older format $ref.bin/ directory was found 1>&2
    echo Please remove it or rename it before proceeding 1>&2
    exit 1
fi

bed=$2
if [ "$bed" = "" ] ; then
    echo You must pass a bed file as the second argument 1>&2
    exit 1
fi
if [ ! -e "$bed" ] ; then
    echo The bed file $bed was not found 1>&2
    exit 1
fi

bam=$3
fasta1=$3
fasta2=$4

if [ "$bam" = "" ] ; then
    echo You must pass a paired end bam file as the third argument 1>&2
    echo or two paired end fasta or fast2 files as arguments 3 and 4 1>&2
    exit 1
fi

if [ "$fasta2" = "" ] ; then
    if [ ! -e "$bam" ] ; then
        echo The bam file $bam was not found 1>&2
        exit 1
    fi
    if [ -e mumdex ] ; then
        echo skipping mummer because mumdex/ exists
    else
        echo running mummer
        mummer -l 1 -samin -qthreads 1 -rcref -verbose $ref \
            <(namepair <(samtools view $bam | head -n 100000))
        echo merging mumdex parts and indexing mums
        merge_mumdex mumdex
    fi
else
    if [ ! -e "$fasta1" ] ; then
        echo Could not find fasta or fastq file $fasta1 1>&2
        exit 1
    fi
    if [ ! -e "$fasta2" ] ; then
        echo Could not find fasta or fastq file $fasta2 1>&2
        exit 1
    fi
    if [ -e mumdex ] ; then
        echo skipping mummer because mumdex/ exists
    else
        echo running mummer
        mummer -normalmem -l 1 -samin -qthreads 12 -rcref -verbose $ref \
            <(fastqs_to_sam $fasta1 $fasta2)
        echo merging mumdex parts and indexing mums
        merge_mumdex mumdex
    fi
fi

pair_view mumdex > pairs.txt

exit 0

echo
echo --------------------------------------
echo counting anchors over the bed file $bed
count_anchors $bed mumdex

echo
echo --------------------------------------
echo converting from binary to show a few nonzero count output lines
(echo chr pos base abspos inr ina outr outa inm outm insup outsup ;
    show_counts_special $bed mumdex ) | grep -v "0 0 0 0" | head -n 20

echo
echo --------------------------------------
echo demonstrating example pair and mum access in mumdex
mumdex_examples mumdex
```
