## Supplementary material for "G-Graph: An interactive genomic graph viewer": MUMdex Source Code for G-Graph: test_numerical.cpp

```
//
// test_numerical
//
// test the numerical.h header
//
// Copyright Peter Andrews 2017 @ CSHL
//

#include <algorithm>
#include <cmath>
#include <exception>
#include <functional>
#include <future>
#include <iomanip>
#include <iostream>
#include <limits>
#include <random>
#include <utility>
#include <vector>

#include "error.h"
#include "numerical.h"
#include "threads.h"

using std::cout;
using std::cerr;
using std::endl;
using std::exception;
using std::fabs;
using std::future;
using std::mt19937_64;
using std::numeric_limits;
using std::random_device;
using std::setprecision;
using std::sqrt;
using std::uniform_int_distribution;
using std::vector;

using paa::Error;
using paa::GoldenMinimizer;
using paa::MultiDimMinimizer;
using paa::ThreadPool;

double test_fun(const double x) {
  return (x - 5.000000001) * (x - 5.000000001) + 10;
}

inline double complicated_fun(const double x,
                              const double offset,
                              const double scale1,
                              const double period1,
                              const double scale2,
                              const double period2,
                              const double phase) {
  return offset + scale1 * cos(period1 * x) + scale2 * cos(period2 * x + phase);
}

inline double adapted_fun(const double x, const vector<double> & parameters) {
  return complicated_fun(x,
                         parameters[0],
                         parameters[1],
                         parameters[2],
                         parameters[3],
                         parameters[4],
                         parameters[5]);
}

class DataTester {
 public:
  DataTester(const vector<double> & x_vals_,
             const vector<double> & y_vals_) :
      x_vals{x_vals_}, y_vals{y_vals_} { }
  double evaluate_block(const vector<double> & parameters,
                        const unsigned int start,
                        const unsigned int stop) const {
    double residual{0.0};
    for (unsigned int xi{start}; xi != stop; ++xi) {
      // residual += fabs(adapted_fun(x_vals[xi], parameters) - y_vals[xi]);
      const double diff{adapted_fun(x_vals[xi], parameters) - y_vals[xi]};
      residual += diff * diff;
    }
    return residual;
  }
  double operator()(const vector<double> & parameters) const {
    const unsigned int n_blocks{32};
    const unsigned int block_size{static_cast<unsigned int>(
        std::max(1UL, x_vals.size() / n_blocks))};
    for (unsigned int start{0} ; start < x_vals.size() ; start += block_size) {
      const unsigned int stop{
        std::min(static_cast<unsigned int>(x_vals.size()), start + block_size)};
      futures.push_back(pool.run([this, &parameters, start, stop]() noexcept {
            return evaluate_block(parameters, start, stop);
          }));
    }
    double result{0};
    for (future<double> & fut : futures) {
      result += fut.get();
    }
    futures.clear();
    return result;
  }
  const vector<double> & x_vals;
  const vector<double> & y_vals;
  mutable vector<future<double>> futures{};
  mutable ThreadPool pool{32};
};

int main(int argc, char * argv[]) try {
  if (--argc != 0) throw Error("usage: test_numerical");
  if (0) cout << argv[0] << endl;
  const GoldenMinimizer minimizer(test_fun, -1000, 1000);
  cerr << "Minimum value is at " << setprecision(20) << minimizer.min() << endl;

  // Randomizer
  random_device rd;
  auto mersenne = mt19937_64(rd());
  uniform_int_distribution<uint64_t> udist{
    0, numeric_limits<uint64_t>::max()};

  // Set up data for multidimensional minimization
  const unsigned int n_points{10000};
  const double x_min{0};
  const double x_max{10};
  const double step{(x_max - x_min) / n_points};
  vector<double> x_vals(n_points);
  vector<double> y_vals(n_points);
  const vector<double> test_params{100, 22.1, 15, 15.2, 3, 1};
  const vector<double> start_params{200, 23, 15, 6.3, 3.5, 2};
  // const vector<double> start_params{120, 2.5, 15.3, 3.0, 3.3, 2.05};
  // const vector<double> start_params{150, 30, 16, 5.0, 1, 1.1};
  for (unsigned int xi{0}; xi != n_points; ++xi) {
    const double x{x_min + xi * step};
    const double y{adapted_fun(x, test_params)};
    x_vals[xi] = x;
    y_vals[xi] = y + 10.0 * udist(mersenne) / numeric_limits<uint64_t>::max();
  }

  // Fit model to data
  const DataTester tester{x_vals, y_vals};

  cerr << setprecision(6);
  MultiDimMinimizer<DataTester, GoldenMinimizer> multi_minimizer{tester};
  const vector<double> best_params{multi_minimizer.minimize(start_params)};

  // Output data + fit values
  cout << setprecision(20);
  cout << "x y initial best" << endl;
  for (unsigned int xi{0}; xi != n_points; ++xi) {
    const double x{x_vals[xi]};
    cout << x << " " << y_vals[xi]
         << " " << adapted_fun(x, start_params)
         << " " << adapted_fun(x, best_params)
         << endl;
  }

  cerr << "Residual is " << tester(best_params) << endl;

  return 0;
} catch (Error & e) {
  cerr << "paa::Error:" << endl;
  cerr << e.what() << endl;
  return 1;
} catch (exception & e) {
  cerr << "std::exception" << endl;
  cerr << e.what() << endl;
  return 1;
} catch (...) {
  cerr << "unknown exception was caught" << endl;
  return 1;
}
```
