## Supplementary material for "G-Graph: An interactive genomic graph viewer": MUMdex Source Code for G-Graph: test_psplot.cpp

```
//
// test_psplot.cpp
//
// test psplot.h and tsv.h header
//
// Copyright 2016 Peter Andrews @ CSHL
//

#include <exception>
#include <functional>
#include <iostream>
#include <memory>
#include <fstream>
#include <random>
#include <vector>

#include "error.h"
#include "psplot.h"
#include "tsv.h"

using std::bind;
using std::cin;
using std::cout;
using std::cerr;
using std::endl;
using std::exception;
using std::function;
using std::mt19937_64;
using std::normal_distribution;
using std::ofstream;
using std::random_device;
using std::vector;

using paa::triangle;
using paa::Bounds;
using paa::Error;
using paa::Marker;
using paa::MarkerGraph;
using paa::PSDoc;
using paa::PSGraph;
using paa::PSHSeries;
using paa::PSPage;
using paa::PSXYSeries;
using paa::TSV;

int main(int argc, char*[]) try {
  if (--argc != 0) throw Error("usage: test_psplot");

  //
  // psplot - plotting where you can control details
  //

  // The main document (can have several per program)
  PSDoc ps{"psplot", "psplot test"};

  // Demo of marker types and colors - a small subset of possibilities
  MarkerGraph marker_graph{ps};

  // Test multiple layouts per page
  // Plots the same graph on the page several times for simplicity
  PSPage layout_page{ps, "Layout Test",
        "1 2 (0.3) =0 1 2 (0.6) 1 +1 0 2 2 -0 0 2 1- -1 1 2 2 |0 0 3 3|-+="};
  PSGraph layout_graph{layout_page, "Layout;X;Y"};
  PSXYSeries layout_series{layout_graph};
  for (unsigned int s{0}; s != 17; ++s) {
    layout_series.add_point(s, s * s);
    layout_page.add(&layout_graph);
  }

  // Test histogram
  random_device rd;
  mt19937_64 mersenne{rd()};
  using normal = normal_distribution<double>;
  function<double()> Gen1{bind(normal(30, 10), mersenne)};
  function<double()> Gen2{bind(normal(55, 15), mersenne)};
  PSPage hpage{ps};
  PSGraph hgraph{hpage, "Test Hist;X;frequency", Bounds{0, 100}};
  PSHSeries<int, uint64_t> hseries{hgraph, 100, "1 0 0", true, "Red !"};
  for (unsigned int n{0}; n != 1000000; ++n) {
    hseries.add_point(Gen1());
  }
  PSHSeries<int, uint64_t> hseries2{hgraph, 100, "0 0 1", true, "Blue !"};
  for (unsigned int n{0}; n != 10000; ++n) {
    hseries2.add_point(Gen2());
  }

  // Raw series - skips the page and graph, just attaches to doc
  const Marker filled_marker{triangle(), 1.0, "0 0 0", 1, true};
  PSXYSeries raw_series{ps, "Raw Series;X;Y", filled_marker};
  const vector<unsigned int> data{1, 2, 3, 4, 5, 6, 7, 8, 9, 10};
  for (const int i : data) {
    raw_series.add_point(i, sqrt(i));
  }

  // Raw graph - skips the page, just attaches to doc
  PSGraph raw_graph{ps, "Raw Graph with Arbitrary Postscript;X;Y"};
  raw_series.add(&raw_graph);  // Add existing series to another graph
  layout_series.add(&raw_graph);  // Again

  // Arbitrary postscript on page test - add to last graph
  raw_graph
      .ps("0 0 0 c np 10 10 m 200 200 l sp")  // points on page
      .ps("0 0 1 c np 2 150 gc m 8 200 gc l sp")  // graph coords
      .ps("0 1 0 c np 0.5 0 gfc m 1 0.5 gfc l sp")  // fractional coords
      .ps("1 0 0 c np 0 1 pfc m 1 0 pfc l sp");  // page fractional coords

  // Also make an eps version of a particular graph
  raw_graph.eps("arbitrary");

  // Nodoc series - goes to a standalone file
  PSXYSeries nodoc_series{"nodoc_series", "nodoc series"};
  for (const int i : data) {
    nodoc_series.add_point(i, sqrt(i));
  }

  // Nodoc graph - goes to standalone file
  PSGraph nodoc_graph{"nodoc_graph", "nodoc graph"};
  PSXYSeries nodoc_graph_series{nodoc_graph};
  for (const int i : data) {
    nodoc_graph_series.add_point(i, sin(i));
  }

  // Nodoc page - goes to standalone file
  PSPage nodoc_page{"nodoc_page", "nodoc graph", ""};
  PSGraph nodoc_page_graph{nodoc_page};
  PSXYSeries nodoc_page_series{nodoc_page_graph};
  for (const int i : data) {
    nodoc_page_series.add_point(i, log(i + 1));
  }

  //
  // TSV - simpler plotting mostly from text file input
  //

  // Create data file for TSV to read
  ofstream tsv_out{"tsv.txt"};
  tsv_out << "x y z c" << endl;
  for (unsigned int i{0}; i != 1000; ++i) {
    const char c{static_cast<char>('A' + (i % 26))};
    tsv_out << i << " " << sin(i / 10.0) << " " << sqrt(i) << " "
            << c << endl;
  }
  tsv_out.close();

  // Load TSV
  cerr << "TSV:" << endl;
  TSV tsv{"tsv.txt"};

  // Plot all vs all - 3 graphs
  tsv.plot();

  // Add a new series w, based on two existing series x and y
  tsv.add_data("w",
               [](const unsigned int c1,
                  const double c2) noexcept {
                 return fabs(c1 + 10 * c2);
               },
               "x", "y");

  // Make better axis label
  tsv("w").label("abs(x + 10 * y)");

  // Plot all vs one - 3 graphs
  tsv.plot("w");

  // Select based upon column c
  tsv.select("c == B");

  // Set limits for y, set log for x
  tsv("y").range(-1.1, 1.1);
  tsv("x").log(true);

  // Plot one vs anoher, with above selection - one plot
  tsv.plot("x", "y");

  // Remove selection
  tsv.select("");

  // Decide to not plot some automatically from now on
  tsv.do_not_plot("y", "z");

  // Plot remaining - only one plot
  tsv.plot();

  return 0;
} catch (Error & e) {
  cerr << "Error:" << endl;
  cerr << e.what() << endl;
  return 1;
} catch (exception & e) {
  cerr << "exception" << endl;
  cerr << e.what() << endl;
  return 1;
} catch (...) {
  cerr << "unknown exception was caught" << endl;
  return 1;
}
```
