## Supplementary material for "G-Graph: An interactive genomic graph viewer": MUMdex Source Code for G-Graph: test_psshade.cpp

```
//
// test_psshade
//
// test fast plotting
//
// Copyright 2018 Peter Andrews @ CSHL
//

#include <exception>
#include <iostream>
#include <fstream>
#include <functional>
#include <random>
#include <string>

#include "error.h"
#include "psplot.h"
#include "utility.h"

using std::cout;
using std::cerr;
using std::endl;
using std::exception;
using std::function;
using std::ifstream;
using std::mt19937_64;
using std::random_device;
using std::string;
using std::normal_distribution;

using paa::commas;
using paa::Error;
using paa::Marker;
using paa::Progress;
using paa::PSDoc;
using paa::PSGraph;
using paa::PSShade;
using paa::PSXYSeries;

int main(int argc, char * argv[]) try {
  if (--argc < 1) throw Error("test_psshade n_points [in_file...]");

  const uint64_t n_points{static_cast<uint64_t>(atol(argv[1]))};

  random_device rd;
  mt19937_64 mersenne(rd());
  normal_distribution<double> dist{0, 0.4};
  function<double()> gen{bind(dist, std::ref(mersenne))};

  // plot
  PSDoc plots{"shade"};

  PSGraph shade_graph{plots,
        "Shade Test - " + commas(n_points) + " Points;X;Y"};
  PSShade shade{shade_graph, "1 0 1"};
  const Marker marker{paa::circle(), 0.2, "0 0 1", 0.1, true};
  PSXYSeries other{shade_graph, marker};
  PSShade shade2{plots, "Shade Test Doc - " + commas(n_points) + " Points;X;Y"};
  PSShade shade3{plots, "Shade Test - " + commas(n_points) + " Points;X;Y"};

  Progress progress{n_points, 0.01, "Filling four series"};
  for (uint64_t p{0}; p != n_points; ++p) {
    const double g1{gen()};
    const double g2{gen()};
    const double x{20.0 * p / n_points};
    const double y{g1 + x * sin(x + g2)};
    shade.add_point(x, y);
    shade2.add_point(x, -y);
    shade3.add_point(2 * g1, g2);
    progress();
  }
  for (uint64_t p{0}; p != 10000; ++p) {
    const double x{20.0 * p / 10000};
    const double y{-20 + 2 * x + gen()};
    other.add_point(x, y);
  }

  if (argc > 1) {
    --argc;
    ++argv;
    while (argc--) {
      const string file_name{argv++[1]};
      cerr << "Loading data from " << file_name
           << " and filling three series" << endl;
      ifstream in_file{file_name.c_str()};
      if (!in_file) throw Error("Problem opening input file") << file_name;
      double x;
      double y;
      PSShade & user_shade{*PSShade::create(plots, file_name + ";x;y")};
      PSShade & user_shade2{*PSShade::create(plots, file_name + ";x;y")};
      user_shade2.histeq(false);
      PSShade & user_shade3{*PSShade::create(plots, file_name + ";x;y")};
      while (in_file >> x >> y) {
        user_shade.add_point(x, y);
        user_shade2.add_point(x, y);
        if (fabs(x - 1) < 0.5 && fabs(y - 1) < 0.5) user_shade3.add_point(x, y);
      }
    }
  }
  cerr << "Rendering graphs as a PS text document" << endl;

  return 0;
} catch (Error & e) {
  cerr << "paa::Error:" << endl;
  cerr << e.what() << endl;
  return 1;
} catch (exception & e) {
  cerr << "std::exception" << endl;
  cerr << e.what() << endl;
  return 1;
} catch (...) {
  cerr << "unknown exception was caught" << endl;
  return 1;
}
```
