## Supplementary material for "G-Graph: An interactive genomic graph viewer": MUMdex Source Code for G-Graph: test_talk.sh

```
#! /bin/bash

function run() {
    [ "$@" = true ] ||
    (
        echo
        80cols.sh -C - 
        "$@"
        80cols.sh -C -
    )
}

(rsync -a --exclude talk2018 --exclude 13023 --exclude 12029 --exclude 'neighbors.*' --exclude 'names.*' ~/web/talk/:/data/web/paa/talk &&
rsync -a --exclude="*.o" --exclude=talk2018 ~/mumdex/ mumdex.com:/data/web/paa/mumdex &&
ssh mumdex.com 'cd /data/web/paa/mumdex && . ~/.bash_profile && make -j 8 talk2018 && cp talk2018 ../talk/') > ~/remote.txt 2>&1 &

mmake talk2018 lint &&
run ~/mumdex/talk2018 &&
cp ~/mumdex/talk2018 ~/web/talk/talk2018
# ssh andmac open http://wigclust19.cshl.edu/talk/ || (echo problem! ; exit 1) 
cstatus=$?

echo
cat ~/remote.txt

wait || exit 1
[ $? = 0 ] && [ $cstatus = 0 ] &&
ssh andmac open -g http://mumdex.com/talk/?page=22 || (echo problem! ; exit 1) 
```
