## Supplementary material for "G-Graph: An interactive genomic graph viewer": MUMdex Source Code for G-Graph: test_threads.cpp

```
//
// test_threads
//
// Make sure thread pool class behaves as expected
//
// Copyright 2017 Peter Andrews @ CSHL
//

#include <exception>
#include <iostream>
#include <string>

#include "error.h"
#include "threads.h"

using std::cout;
using std::cerr;
using std::endl;
using std::exception;
using std::string;

using paa::Error;
using paa::ThreadPool;

int main(int argc, char * argv[]) try {
  if (0) cout << argc << argv[1] << endl;

  paa::test_thread_pool();

  return 0;
} catch (Error & e) {
  cerr << "paa::Error:" << endl;
  cerr << e.what() << endl;
  return 1;
} catch (exception & e) {
  cerr << "std::exception" << endl;
  cerr << e.what() << endl;
  return 1;
} catch (...) {
  cerr << "unknown exception was caught" << endl;
  return 1;
}
```
