## Supplementary material for "G-Graph: An interactive genomic graph viewer": MUMdex Source Code for G-Graph: test_x11.cpp

```
//
// test_x11.cpp
//
// test x11
//
// Copyright 2016 Peter Andrews @ CSHL
//

#include <algorithm>
#include <exception>
#include <iostream>
#include <string>
#include <utility>
#include <vector>

#include "error.h"
#include "x11plot.h"

using std::cerr;
using std::cout;
using std::endl;
using std::exception;
using std::max;
using std::string;
using std::vector;

namespace paa {

class X11Circles : public X11Win {
 public:
  using X11Win::X11Win;

  explicit X11Circles(X11App & app__) : X11Win(app__) {
    XSelectInput(display(), window, StructureNotifyMask | ExposureMask |
                 KeyPressMask | ButtonMotionMask | ButtonPressMask);
  }

  virtual void button_press(const XButtonEvent & event) {
    skip = event.x + 1;
    factor = 1.0 * event.y / height();
    draw();
  }
  virtual void motion(const XMotionEvent & event) {
    skip = event.x + 1;
    factor = 1.0 * event.y / height();
    draw();
  }

  virtual void draw() {
    std::vector<XArc> arcs;
    XArc circle;
    circle.width = width() * factor;
    circle.height = height() * factor;
    circle.angle1 = 0;
    circle.angle2 = 64 * 360;
    XFillRectangle(display(), window, fill_gc, 0, 0, width(), height());

    for (int x{0}; x <= width() + 10 * circle.width; x += skip) {
      for (int y{0}; y <= height() + 10 * circle.height; y += skip) {
        circle.x = x;
        circle.y = y;
        circle.x -= 5.5 * circle.width;
        circle.y -= 5.5 * circle.height;
        arcs.emplace_back(circle);
        if (arcs.size() == max_request / 3) {
          XDrawArcs(display(), window, gc, &arcs[0],
                    static_cast<unsigned int>(arcs.size()));
          arcs.clear();
        }
      }
    }

    if (arcs.size()) XDrawArcs(display(), window, gc, &arcs[0],
                               static_cast<unsigned int>(arcs.size()));
    XFlush(display());
  }

  virtual ~X11Circles() { }

 private:
  unsigned int skip{30};
  double factor{0.05};
};

}  // namespace paa

using paa::Error;
using paa::X11App;
using paa::X11Circles;
using paa::X11Graph;
using paa::X11TextGrid;
using paa::X11Win;

int main(int argc, char*[]) try {
  if (--argc != 0) throw Error("usage: test_x11");

  const unsigned int ns{3};
  const unsigned int np{10000};
  vector<vector<double>> xs(ns);
  vector<vector<double>> ys(ns);
  for (unsigned int s{0}; s != ns; ++s) {
    for (unsigned int x{0}; x != np; ++x) {
      xs[s].push_back(x);
      ys[s].push_back(max(-2.5, (s + 1) * sin(10.0 * x / np + s)));
    }
  }

  X11App app;
  app.create<X11Circles>();
  app.create<X11Graph>(xs[0], ys[0], xs[1], ys[1], xs[2], ys[2]);
  vector<vector<string>> text{
    {"hello", "there", "my", "friend"},
    {"World", "there", "mistake", "me"}, {"1", "2", "", "4"}};
  using Vec = std::vector<unsigned int>;
  using Cells = std::vector<std::pair<unsigned int, unsigned int>>;
  app.create<X11TextGrid>(text, Cells{}, Cells{},
                          Vec{0}, Vec{0}, Vec{1}, Vec{},
                          nullptr, nullptr, nullptr);
  app.run();

  return 0;
} catch (Error & e) {
  cerr << "Error:" << endl;
  cerr << e.what() << endl;
  return 1;
} catch (exception & e) {
  cerr << "exception" << endl;
  cerr << e.what() << endl;
  return 1;
} catch (...) {
  cerr << "unknown exception was caught" << endl;
  return 1;
}
```
