## Supplementary material for "G-Graph: An interactive genomic graph viewer": MUMdex Source Code for G-Graph: transmission.cpp

```
//
// transmission
//
// get some counts for transmission study
//
// Copyright 2018 Peter Andrews @ CSHL
//

#include <algorithm>
#include <array>
#include <chrono>
#include <exception>
#include <functional>
#include <future>
#include <iostream>
#include <mutex>
#include <sstream>
#include <string>
#include <utility>
#include <vector>

#include "block_reader.h"
#include "bridges.h"
#include "error.h"
#include "files.h"
#include "mumdex.h"
#include "population.h"
#include "sequence.h"
#include "threads.h"
#include "utility.h"

using std::array;
using std::cerr;
using std::cout;
using std::defer_lock;
using std::endl;
using std::exception;
using std::flush;
using std::future;
using std::lock_guard;
using std::lower_bound;
using std::mutex;
using std::ostream;
using std::ostringstream;
using std::pair;
using std::string;
using std::unique_lock;
using std::vector;

using paa::file_size;
using paa::get_block_size;
using paa::serr;
using paa::sout;
using paa::BridgeInfo;
using BlockMerger = paa::BlockMerger<BridgeInfo>;
using BlockReader = paa::BlockReader<BridgeInfo>;
using paa::ChromosomeIndexLookup;
using paa::ConsensusSequence;
using paa::Error;
using paa::Family;
using paa::FileVector;
using paa::Mappability;
using paa::MUM;
using paa::MUMindex;
using paa::Pair;
using paa::Population;
using paa::Reference;
using paa::Sample;
using paa::Similarity;
using paa::SpaceOut;
using paa::ThreadPool;

#define USE_FILES 1

#if USE_FILES
using MUMdex = paa::FileMUMdex;
using Bridges = paa::FileBridges;
#else
using MUMdex = paa::MUMdex;
using Bridges = paa::Bridges;
#endif

using Anchors = MUMdex::Anchors;
using Bridge = Bridges::Bridge;

const bool verbose{false};

const bool exclude_snps{true};
const unsigned int min_support_length{20};
const unsigned int min_mate_count{1};
const unsigned int min_mate_support{20};
const unsigned int min_excess_mappability{0};

const unsigned int t1{10};
const unsigned int t2{2};
const unsigned int t3{1};
const unsigned int T2{50};
const unsigned int T3{0};

uint64_t n_no_cut{0};
uint64_t n_not_snp{0};
uint64_t n_one_family{0};
uint64_t n_children{0};
uint64_t n_children_one_family{0};
uint64_t n_big_count{0};
uint64_t n_good_support{0};
uint64_t n_good_mate_support{0};
uint64_t n_good_excess{0};
uint64_t n_adjacent_cut{0};
uint64_t n_parent_coverage_cut{0};

void bridge_out(ostream & out, const BridgeInfo & bridge,
                const bool end_line = false) {
  out << " " << bridge.chr1()
      << " " << bridge.pos1()
      << " " << bridge.high1()
      << " " << bridge.chr2()
      << " " << bridge.pos2()
      << " " << bridge.high2()
      << " " << bridge.invariant()
      << " " << bridge.offset()
      << " " << bridge.bridge_count();
  if (end_line) out << endl;
}

int main(int argc, char* argv[])  try {
  if (--argc != 7)
    throw Error("usage: transmission ref family_file bridges_dir "
                "n_threads chromosome start stop");

  // Other command line arguments
  const string reference_file{argv[1]};
  const Reference ref{reference_file};
  const ChromosomeIndexLookup chr_lookup{ref};
  const Mappability map{reference_file, true};
  const string family_file{argv[2]};
  const Population pop{family_file};
  const string bridges_dir{argv[3]};
  const unsigned int n_threads{static_cast<unsigned int>(atoi(argv[4]))};
  const string chromosome_name{argv[5]};
  const unsigned int chromosome{chr_lookup[chromosome_name]};
  const unsigned int chromosome_offset{ref.offset(chromosome)};
  const unsigned int start{static_cast<unsigned int>(atoi(argv[6]))};
  const unsigned int stop{static_cast<unsigned int>(atoi(argv[7]))};

  ThreadPool pool{n_threads};

  // Set up bridge files to start position, in parallel to save time
  vector<future<BlockReader>> futures;
  futures.reserve(pop.n_samples());
  uint64_t block_size{4096};
  for (const auto s : pop.samples()) {
    ostringstream bridges_name;
    bridges_name << bridges_dir << "/" << pop.sample(s) << "/"
                 << get_bridge_file_name(ref, chromosome);
    if (!s) {
      block_size = std::max(block_size, get_block_size(bridges_name.str()));
      cerr << "Block size is " << block_size << endl;
    }
    futures.push_back(pool.run([start, stop](const string name){
          return BlockReader{name, start, stop};
        }, bridges_name.str()));
  }
  vector<BlockReader> readers;
  unsigned int n_samples_loaded{0};
  for (const auto s : pop.samples()) {
    if (s != n_samples_loaded) throw Error("Sample sanity check");
    readers.push_back(futures[n_samples_loaded].get());
    serr << pop.family(pop.family(s))
         << pop.sample(s)
         << pop.member(s)
         << ++n_samples_loaded
         << endl;
  }

  ThreadPool load_pool{n_threads};
  BlockMerger merger{load_pool, block_size = 32768, readers};

  // Document the run
  serr << "Run parameters:" << endl
       << "  n_threads" << n_threads << endl
       << "  family_file" << family_file << endl
       << "  bridges_dir" << bridges_dir << endl
       << "  chromosome" << chromosome_name << endl
       << "  start" << start << endl
       << "  stop" << stop << endl
       << "  n_families" << pop.n_families() << endl
       << "  n_samples" << pop.n_samples() << endl
       << "  exclude_snps" << exclude_snps << endl
       << "  min_support_length" << min_support_length << endl
       << "  min_mate_count" << min_mate_count << endl
       << "  min_mate_support" << min_mate_support << endl
       << "  min_excess_mappability" << min_excess_mappability << endl;

  sout << "family"
       << "members"
       << "samples"
       << "sex"
       << "n_in_family"

       << "bridges"
       << "n_t1"
       << "n_t2"
       << "n_t3"

       << "chr"
       << "chrA" << "posA" << "highA"
       << "chrB" << "posB" << "highB"
       << "invariant" << "offset"
       << "type" << "orient"

       << "supA" << "supB"
       << "emapA" << "emapB"
       << "mapA" << "mapB"
       << "mCA" << "mCB"
       << "mSA" << "mSB"

       << "max_bridge_count"

       << "pop_n_families"
       << "pop_n_samples"
       << "pop_n_parents"
       << "pop_n_children"
       << "pop_total"
       << "pop_parent_total"
       << "pop_child_total"
       << "pop_parent_counts"
       << "pop_child_counts"
       << "pop_info"
       << endl;

  // Loop, adding one bridge at a time to all_bridges if the same event
  vector<BridgeInfo> all_bridges;
  vector<Sample> all_samples;
  vector<unsigned int> parent_counts;
  vector<unsigned int> children_counts;
  unsigned int total_bridge_count{0};
  unsigned int parent_bridge_count{0};
  unsigned int parent_seen{0};
  unsigned int child_bridge_count{0};
  unsigned int child_seen{0};
  uint64_t n_bridges{0};
  const uint64_t notify_interval{10000000};
  bool reset{true};
  BridgeInfo last_bridge{};
  while (true) {
    // New bridge expected - start fresh
    if (reset) {
      all_bridges.clear();
      all_samples.clear();
      parent_counts.clear();
      children_counts.clear();
      total_bridge_count = 0;
      parent_bridge_count = 0;
      parent_seen = 0;
      child_bridge_count = 0;
      child_seen = 0;
      reset = false;
    }

    // Add a bridge to the list if the same event
    bool bridge_done{true};
    if (merger.available()) {
      const BlockMerger::Item & item{merger.next()};
      const BridgeInfo current{item.first};
      // if (all_bridges.size() && current < all_bridges.back()) {
      if (current < last_bridge) {
        cerr << "Bridge misordering" << endl;
        bridge_out(cerr, last_bridge);
        cerr << endl;
        bridge_out(cerr, current);
        const Sample sample{item.second};
        cerr << " " << pop.sample(sample);
        cerr << endl;
        exit(1);
      }
      last_bridge = current;
      if (all_bridges.empty() || !(all_bridges.back() < current)) {
        ++n_no_cut;
        bridge_done = false;
        all_bridges.push_back(current);
        total_bridge_count += current.bridge_count();
        const Sample sample{item.second};
        all_samples.emplace_back(sample);
        if (pop.is_parent(sample)) {
          parent_bridge_count += current.bridge_count();
          ++parent_seen;
        } else {
          child_bridge_count += current.bridge_count();
          ++child_seen;
        }
        merger.advance();
      }
    } else {
      if (all_samples.empty()) {
        if (verbose) cerr << "bridge queue empty" << endl;
        break;
      }
    }

    // No new bridges added, so bridge list is ready to process
    if (bridge_done) {
      reset = true;

      // The first bridge, the exemplar since all are the same signature
      const BridgeInfo & bridge{all_bridges[0]};

      if ((++n_bridges % notify_interval) == 0)
        cerr << "progress: " << n_bridges << " " << bridge.pos1() << " "
             << 1.0 * (bridge.pos1() - start) / (stop - start) << endl;

      // SNP cut
      const bool is_snp{bridge.chr1() == bridge.chr2() &&
            bridge.invariant() == 0 && bridge.high1() != bridge.high2()};
      if (is_snp) {
        if (exclude_snps) continue;
        throw Error("Cannot deal with snps here");
      }

      // Count families seen
      const unsigned int n_families{[&all_samples, &pop] {
          unsigned int n{0};
          Family last_family{100000};
          for (const Sample sample : all_samples) {
            const Family family{pop.family(sample)};
            if (family != last_family) {
              ++n;
              last_family = family;
            }
          }
          return n;
        }()};

      // Seen at big counts in parents
      const unsigned int n_parents_t1{[&all_samples, &all_bridges, &pop] {
          unsigned int n{0};
          for (unsigned int b{0}; b != all_bridges.size(); ++b)
            if (pop.is_parent(all_samples[b]) &&
                all_bridges[b].bridge_count() >= t1) ++n;
          return n;
        }()};

      // Initial T1 cut
      if (n_parents_t1 == 0) continue;

      // Count parents seen at over t2
      const unsigned int n_parents_t2{[&all_samples, &all_bridges, &pop] {
          unsigned int n{0};
          for (unsigned int b{0}; b != all_bridges.size(); ++b)
            if (pop.is_parent(all_samples[b]) &&
                all_bridges[b].bridge_count() > t2) ++n;
          return n;
        }()};

      // Initial T2 cut
      if (n_parents_t2 > T2 + 1) continue;

      // Seen at small counts in parents
      const unsigned int n_parents_t3{[&all_samples, &all_bridges, &pop] {
          unsigned int n{0};
          for (unsigned int b{0}; b != all_bridges.size(); ++b)
            if (pop.is_parent(all_samples[b]) &&
                all_bridges[b].bridge_count() > 0 &&
                all_bridges[b].bridge_count() <= t3) ++n;
          return n;
        }()};

      // Initial T3 cut - plus one for one family with surprise
      if (n_parents_t3 > T3 + 1) continue;

      // Get mappability information
      const unsigned int mum1_abspos{chromosome_offset + bridge.pos1()};
      const unsigned int mum2_abspos{ref.offset(bridge.chr2()) + bridge.pos2()};
      const unsigned int mum1_map{map.low_high(bridge.high1(), mum1_abspos)};
      const unsigned int mum2_map{map.low_high(bridge.high2(), mum2_abspos)};

      for (unsigned int s{0}; s != all_samples.size(); ++s)
        if (pop.is_parent(all_samples[s])) {
          parent_counts.push_back(all_bridges[s].bridge_count());
        } else {
          children_counts.push_back(all_bridges[s].bridge_count());
        }

      ostringstream pop_info;
      Family last_family{100000000};
      for (unsigned int s{0}; s != all_samples.size(); ++s) {
        if (s) pop_info << ",";
        const Sample sample{all_samples[s]};
        const Family family{pop.family(sample)};
        if (family != last_family) {
          pop_info << pop.family(family) << ":";
          last_family = family;
        }
        pop_info << pop.member(sample)[0] << all_bridges[s].bridge_count();
      }

      // Look at each family separately
      const unsigned int n_samples{static_cast<unsigned int>(
          all_samples.size())};
      while (all_samples.size()) {
        // Gather samples and bridges for family
        static vector<Sample> samples;
        static vector<BridgeInfo> bridges;
        samples.clear();
        bridges.clear();
        while (all_samples.size() &&
               (samples.empty() ||
                pop.family(samples.back()) ==
                pop.family(all_samples.back()))) {
          samples.push_back(all_samples.back());
          all_samples.pop_back();
          bridges.push_back(all_bridges.back());
          all_bridges.pop_back();
        }

        vector<unsigned int> family_parent_counts;
        for (unsigned int b{0}; b != bridges.size(); ++b) {
          const Sample sample{samples[b]};
          if (pop.is_parent(sample))
            family_parent_counts.push_back(bridges[b].bridge_count());
        }
        sort(family_parent_counts.begin(), family_parent_counts.end());

        if (family_parent_counts.empty()) continue;
        if (family_parent_counts.back() < t1) continue;
        if (family_parent_counts.size() == 2) {
          if (family_parent_counts.front() > t2) continue;
        } else {
          if (n_parents_t3 > T3) continue;
        }

        // Gather max support counts and lengths
        unsigned int max_bridge_count{0};
        unsigned int max_anchor1_support{0};
        unsigned int max_anchor2_support{0};
        unsigned int max_mate_anchor1_count{0};
        unsigned int max_mate_anchor1_support{0};
        unsigned int max_mate_anchor2_count{0};
        unsigned int max_mate_anchor2_support{0};
        for (const BridgeInfo & sample_bridge : bridges) {
          if (max_bridge_count < sample_bridge.bridge_count())
            max_bridge_count = sample_bridge.bridge_count();
          if (max_anchor1_support < sample_bridge.anchor1_length())
            max_anchor1_support = sample_bridge.anchor1_length();
          if (max_anchor2_support < sample_bridge.anchor2_length())
            max_anchor2_support = sample_bridge.anchor2_length();
          if (max_mate_anchor1_count < sample_bridge.mate_anchor1_count())
            max_mate_anchor1_count = sample_bridge.mate_anchor1_count();
          if (max_mate_anchor2_count < sample_bridge.mate_anchor2_count())
            max_mate_anchor2_count = sample_bridge.mate_anchor2_count();
          if (max_mate_anchor1_support < sample_bridge.mate_anchor1_length())
            max_mate_anchor1_support = sample_bridge.mate_anchor1_length();
          if (max_mate_anchor2_support < sample_bridge.mate_anchor2_length())
            max_mate_anchor2_support = sample_bridge.mate_anchor2_length();
        }

        // Low bridge count cut
        if (max_bridge_count < t1) continue;
        ++n_big_count;

        // Short anchor support cut
        if (max_anchor1_support < min_support_length ||
            max_anchor2_support < min_support_length) continue;
        ++n_good_support;

        // Short mate support cut
        if (max_mate_anchor1_count < min_mate_count ||
            max_mate_anchor2_count < min_mate_count ||
            max_mate_anchor1_support < min_mate_support ||
            max_mate_anchor2_support < min_mate_support) continue;
        ++n_good_mate_support;

        // Short excess mappability cut
        if (max_anchor1_support < mum1_map) {
          cerr << "Mappability error" << endl;
          exit(1);
        }
        if (max_anchor1_support < min_excess_mappability + mum1_map)
          continue;
        if (max_anchor2_support < min_excess_mappability +  mum2_map)
          continue;
        ++n_good_excess;

        ostringstream out;

        // individuals info
        const Family family{pop.family(samples.front())};
        const vector<Sample> & family_members{pop.samples(family)};
        out << pop.family(family);
        for (unsigned int s{0}; s != samples.size(); ++s)
          out << (s ? "," : " ") << pop.member(samples[s]);
        for (unsigned int s{0}; s != samples.size(); ++s)
          out << (s ? "," : " ") << pop.sample(samples[s]);
        for (unsigned int s{0}; s != samples.size(); ++s)
          out << (s ? "," : " ") << pop.sex(samples[s]);
        out << " " << samples.size();

        for (unsigned int m{0}; m != family_members.size(); ++m) {
          out << (m ? "," : " ");
          const Sample sample{family_members[m]};
          const vector<Sample>::const_iterator found{
            find(samples.begin(), samples.end(), sample)};
          if (found == samples.end()) {
            out << 0;
          } else {
            const unsigned int count{
              bridges[found - samples.begin()].bridge_count()};
            out << count;
          }
        }
        out << " " << n_parents_t1
            << " " << n_parents_t2
            << " " << n_parents_t3;

        out << " " << bridge.chr1()
            << " " << ref.name(bridge.chr1())
            << " " << bridge.pos1()
            << " " << bridge.high1()
            << " " << ref.name(bridge.chr2())
            << " " << bridge.pos2()
            << " " << bridge.high2()
            << " " << bridge.invariant()
            << " " << bridge.offset()
            << " " << bridge.description()
            << " " << bridge.orientation_char();

        // anchor info
        out << " " << max_anchor1_support
            << " " << max_anchor2_support
            << " " << max_anchor1_support - mum1_map
            << " " << max_anchor2_support - mum2_map
            << " " << mum1_map
            << " " << mum2_map
            << " " << max_mate_anchor1_count
            << " " << max_mate_anchor2_count
            << " " << max_mate_anchor1_support
            << " " << max_mate_anchor2_support;

        // bridge counts
        out << " " << max_bridge_count;

        // population bridge counts
        out << " " << n_families
            << " " << n_samples
            << " " << parent_seen
            << " " << child_seen
            << " " << total_bridge_count
            << " " << parent_bridge_count
            << " " << child_bridge_count;

        // parent counts
        if (parent_counts.size()) {
          for (unsigned int p{0}; p != parent_counts.size(); ++p)
            out << (p ? ',' : ' ') << parent_counts[p];
        } else {
          out << " 0";
        }
        // children counts
        if (children_counts.size()) {
          for (unsigned int c{0}; c != children_counts.size(); ++c)
            out << (c ? ',' : ' ') << children_counts[c];
        } else {
          out << " 0";
        }

        // detailed population info
        out << " " << pop_info.str();

        cout << out.str() << endl;
      }
    }
  }

  cerr << "finished bridge loop" << endl;

  return 0;
} catch (Error & e) {
  cerr << "paa::Error:" << endl;
  cerr << e.what() << endl;
  return 1;
} catch (exception & e) {
  cerr << "std::exception" << endl;
  cerr << e.what() << endl;
  return 1;
} catch (...) {
  cerr << "unknown exception was caught" << endl;
  return 1;
}
```
