## Supplementary material for "G-Graph: An interactive genomic graph viewer": MUMdex Source Code for G-Graph: unequal_bridges.cpp

```
//
// unequal_bridges
//
// find bridges with different counts in different samples
//
// Copyright 2016 Peter Andrews @ CSHL
//

#include <algorithm>
#include <deque>
#include <exception>
#include <iostream>
#include <queue>
#include <sstream>
#include <string>
#include <utility>
#include <vector>

#include "bridges.h"
#include "error.h"
#include "files.h"
#include "mumdex.h"
#include "poisson.h"
#include "population.h"
#include "utility.h"

using std::cerr;
using std::cout;
using std::deque;
using std::endl;
using std::exception;
using std::lower_bound;
using std::max;
using std::min;
using std::move;
using std::ostringstream;
using std::priority_queue;
using std::string;
using std::vector;

using paa::ref_ptr;
using paa::serr;
using paa::sout;
using paa::BridgeInfo;
using paa::ChromosomeIndexLookup;
using paa::Error;
using paa::FileVector;
using paa::Mappability;
using paa::MergeHelper;
using paa::MergeHelperCompare;
using paa::Poisson;
using paa::Reference;
using paa::Sample;
using paa::SpaceOut;

#if 0
using MUMdex = paa::FileMUMdex;
using Bridges = paa::FileBridges;
#else
using MUMdex = paa::MUMdex;
using Bridges = paa::Bridges;
#endif

using Anchors = MUMdex::Anchors;
using Bridge = Bridges::Bridge;

const bool verbose{false};

const bool output_snps{false};
const unsigned int min_support_length{20};
const unsigned int min_mate_support_length{20};

uint64_t n_no_cut{0};
uint64_t n_not_snp{0};
uint64_t n_two_or_less{0};
uint64_t n_one_family{0};
uint64_t n_children{0};
uint64_t n_children_one_family{0};
uint64_t n_no_rare_inherited{0};
uint64_t n_one_parent{0};
uint64_t n_big_count{0};
uint64_t n_good_support{0};
uint64_t n_good_mate_support{0};
uint64_t n_good_excess{0};
uint64_t n_good_pop{0};
uint64_t n_adjacent_cut{0};
uint64_t n_parent_coverage_cut{0};

const Reference * paa::ref_ptr;

int main(int argc, char* argv[])  try {
  --argc;
  if (argc < 6) {
    throw Error("usage: unequal_bridges ref chromosome start stop sample ...");
  }

  // Command line arguments
  const string reference_file{argv[1]};
  const Reference ref{reference_file};
  ref_ptr = & ref;
  const ChromosomeIndexLookup chr_lookup{ref};
  const Mappability map{reference_file, true};
  const string chromosome_name{argv[2]};
  const unsigned int chromosome{chr_lookup[chromosome_name]};
  const unsigned int chromosome_offset{ref.offset(chromosome)};
  const unsigned int start{static_cast<unsigned int>(atoi(argv[3]))};
  const unsigned int stop{static_cast<unsigned int>(atoi(argv[4]))};

  argc -= 4;
  argv += 4;

  // Set up queue of bridge files
  deque<MergeHelper> helpers;
  using PQueue = priority_queue<MergeHelper*, vector<MergeHelper *>,
      MergeHelperCompare>;
  PQueue queue{MergeHelperCompare()};
  unsigned int n_samples_skipped{0};
  vector<string> samples;
  vector<double> n_pairs;
  unsigned int si{0};
  while (argc--) {
    const string sample_name{argv[1]};
    samples.push_back(sample_name);
    const MUMdex mumdex{sample_name + "/mumdex"};
    n_pairs.push_back(mumdex.n_pairs());
    ++argv;
    ostringstream bridges_name;
    bridges_name << sample_name <<  "/bridges_snp/"
                 << get_bridge_file_name(ref, chromosome);
    MergeHelper helper{bridges_name.str(), Sample{si++}, start, stop};
    if (helper.valid_start()) {
      helpers.push_back(move(helper));
      queue.push(&helpers.back());
    } else {
      ++n_samples_skipped;
    }
  }
  const double mean_n_pairs{1.0 * accumulate(n_pairs.begin(), n_pairs.end(),
                                             0UL) / samples.size()};

  cout << "chrA posA hA chrB posB hB inv off o sA sB mcA mcB msA msB";
  for (unsigned int s{0}; s != samples.size(); ++s) {
    cout << " " << samples[s];
  }
  if (0) {
    for (unsigned int s{0}; s != samples.size(); ++s) {
      cout << " n_" << samples[s];
    }
    cout << " avg";
    for (unsigned int s{0}; s != samples.size(); ++s) {
      cout << " p_" << samples[s];
    }
  }
  cout << " mapA mapB p lp" << endl;

  // Loop, adding one bridge at a time to all_bridges if the same event
  vector<BridgeInfo> all_bridges;
  vector<unsigned int> all_samples;
  vector<unsigned int> counts;
  vector<double> ncounts;
  unsigned int total_bridge_count{0};
  bool reset{true};
  while (true) {
    // New bridge expected - start fresh
    if (reset) {
      all_bridges.clear();
      all_samples.clear();
      total_bridge_count = 0;
      reset = false;
    }

    // Add a bridge to the list if the same event
    bool bridge_done{true};
    if (queue.size()) {
      MergeHelper * top{queue.top()};
      const BridgeInfo current{top->current()};
      if (all_bridges.empty() || !(all_bridges.back() < current)) {
        ++n_no_cut;
        bridge_done = false;
        all_bridges.push_back(current);
        total_bridge_count += static_cast<uint64_t>(current.bridge_count());
        const unsigned int sample{top->sample()};
        all_samples.emplace_back(sample);
        queue.pop();
        if (top->advance()) {
          queue.push(top);
        }
      }
      if (queue.empty()) bridge_done = true;
      if (bridge_done) reset = true;
    } else {
      if (all_samples.empty()) {
        if (verbose) cerr << "bridge queue empty" << endl;
        break;
      }
    }

    // No new bridges added, so bridge list is ready to process
    if (bridge_done) {
      if (total_bridge_count < 5) continue;
      // The first bridge, the exemplar since all are the same
      const BridgeInfo & bridge{all_bridges[0]};
      if (bridge.invariant() == 0 && !output_snps) continue;
      const unsigned int max_anchor1_length{[&all_bridges]() {
          unsigned int m{0};
          for (const BridgeInfo & b : all_bridges) {
            m = max(m, b.anchor1_length());
          }
          return m;
        }()};
      if (max_anchor1_length < min_support_length) continue;
      const unsigned int max_anchor2_length{[&all_bridges]() {
          unsigned int m{0};
          for (const BridgeInfo & b : all_bridges) {
            m = max(m, b.anchor2_length());
          }
          return m;
        }()};
      if (max_anchor2_length < min_support_length) continue;
      const unsigned int mate_anchor1_count{[&all_bridges]() {
          unsigned int m{0};
          for (const BridgeInfo & b : all_bridges) {
            m += b.mate_anchor1_count();
          }
          return m;
        }()};
      const unsigned int mate_anchor2_count{[&all_bridges]() {
          unsigned int m{0};
          for (const BridgeInfo & b : all_bridges) {
            m += b.mate_anchor2_count();
          }
          return m;
        }()};
      const unsigned int max_mate_anchor1_length{[&all_bridges]() {
          unsigned int m{0};
          for (const BridgeInfo & b : all_bridges) {
            m = max(m, b.mate_anchor1_length());
          }
          return m;
        }()};
      if (max_mate_anchor1_length < min_mate_support_length) continue;
      const unsigned int max_mate_anchor2_length{[&all_bridges]() {
          unsigned int m{0};
          for (const BridgeInfo & b : all_bridges) {
            m = max(m, b.mate_anchor2_length());
          }
          return m;
        }()};
      if (max_mate_anchor2_length < min_mate_support_length) continue;
      ostringstream out;
      SpaceOut<ostringstream> ssout(out);
      ssout << ref.name(bridge.chr1()) << bridge.pos1() << bridge.high1()
           << ref.name(bridge.chr2()) << bridge.pos2() << bridge.high2()
           << bridge.invariant() << bridge.offset() << bridge.orientation_char()
           << max_anchor1_length << max_anchor2_length
           << mate_anchor1_count << mate_anchor2_count
           << max_mate_anchor1_length << max_mate_anchor2_length;
      counts.clear();
      counts.resize(samples.size());
      ncounts.clear();
      ncounts.resize(samples.size());
      unsigned int avail_sample{0};
      unsigned int min_count(-1);
      unsigned int max_count{0};
      double max_ncount{0};
      double total_ncount{0};
      for (unsigned int sample{0}; sample != samples.size(); ++sample) {
        if (avail_sample < all_samples.size() &&
            sample == all_samples[avail_sample]) {
          counts[sample] = all_bridges[avail_sample].bridge_count();
          ++avail_sample;
        } else {
          counts[sample] = 0;
        }
        ssout << counts[sample];
        ncounts[sample] = counts[sample] / n_pairs[sample];
        total_ncount += ncounts[sample];
        min_count = min(min_count, counts[sample]);
        max_count = max(max_count, counts[sample]);
        max_ncount = max(max_ncount, ncounts[sample]);
      }
      if (min_count * 10 > max_count) continue;  // Need ?
      const double mean_count{1.0 * total_ncount / samples.size()};
      if (0) {
        for (unsigned int s{0}; s != samples.size(); ++s) {
          ssout << ncounts[s] * mean_n_pairs;
        }
        ssout << mean_count * mean_n_pairs;
      }
      double p{1};
      for (unsigned int s{0}; s != samples.size(); ++s) {
        const unsigned int count{counts[s]};
        const double sample_mean{mean_count * n_pairs[s]};
        const Poisson poisson{sample_mean, count};
        const double cdf{min(poisson.cdf(count), poisson.ucdf(count))};
        // const double sp{max(min_prob, min(1.0, 2 * cdf))};
        const double sp{min(1.0, 2 * cdf)};
        p *= sp;
        // ssout << cdf;
      }

      // Get mappability information
      const unsigned int mum1_abspos{chromosome_offset + bridge.pos1()};
      const unsigned int mum2_abspos{ref.offset(bridge.chr2()) + bridge.pos2()};
      const unsigned int mum1_map{map.low_high(bridge.high1(), mum1_abspos)};
      const unsigned int mum2_map{map.low_high(bridge.high2(), mum2_abspos)};
      ssout << mum1_map << mum2_map;
      ssout << p << log10(p);
      if (p < 0.001) cout << out.str() << endl;
    }
  }

  cerr << "finished bridge loop" << endl;

  return 0;
} catch (Error & e) {
  cerr << "paa::Error:" << endl;
  cerr << e.what() << endl;
  return 1;
} catch (exception & e) {
  cerr << "std::exception" << endl;
  cerr << e.what() << endl;
  return 1;
} catch (...) {
  cerr << "unknown exception was caught" << endl;
  return 1;
}
```
