## Supplementary material for "G-Graph: An interactive genomic graph viewer": MUMdex Source Code for G-Graph: validate_mumdex.cpp

```
//
// validate_mumdex.cpp
//
// Copyright 2017 Peter Andrews @ CSHL
//

#include <algorithm>
#include <iostream>
#include <fstream>
#include <map>
#include <sstream>
#include <string>
#include <vector>

#include "bridges.h"
#include "error.h"
#include "mumdex.h"
#include "utility.h"

using std::cout;
using std::cerr;
using std::endl;
using std::ifstream;
using std::istringstream;
using std::map;
using std::max;
using std::string;
using std::vector;

using paa::sout;
using paa::Bridge;
using paa::Bridges;
using paa::ChromosomeIndexLookup;
using paa::Error;
using paa::MUMdex;
using paa::Reference;

void parse_counts(const string & in_string, unsigned int vals[4]) {
  istringstream in{in_string.c_str()};
  for (const unsigned int i : {0, 1, 2, 3}) {
    in >> vals[i];
    in.get();
  }
}

class CandInfo {
 public:
  class ParseError {};
  explicit CandInfo(const string & line) {
    istringstream input{line.c_str()};
    string bridges_;
    string anchorsA_;
    string anchorsB_;
    string coveragesA_;
    string coveragesB_;
    input >> family >> parent >> kids >> nFam >> nSam >> nInFam
          >> chr >> chrA >> posA >> highA >> chrB >> posB >> highB
          >> type >> ori >> invariant >> offset >> bridge
          >> supA >> supB >> mCA >> mSA >> mCB >> mSB >> mapA >> mapB
          >> minParCov;
    const bool full_info{false};
    if (full_info) {
      input >> bridges_ >> anchorsA_ >> anchorsB_
            >> coveragesA_ >> coveragesB_
            >> pNP >> pNB >> pMed >> pMax >> tBC >> nBC >> nS;
      parse_counts(bridges_, bridges);
      parse_counts(anchorsA_, anchorsA);
      parse_counts(anchorsB_, anchorsB);
      parse_counts(coveragesA_, coveragesA);
      parse_counts(coveragesB_, coveragesB);
    }
    if (!input) {
      throw ParseError{};
    }
  }
  string family{};
  string parent{};
  string kids{};
  unsigned int nFam{};
  unsigned int nSam{};
  unsigned int nInFam{};
  unsigned int chr{};
  string chrA{};
  unsigned int posA{};
  bool highA{};
  string chrB{};
  unsigned int posB{};
  bool highB{};
  string type{};
  char ori{};
  int invariant{};
  int offset{};
  unsigned int bridge{};
  unsigned int supA{};
  unsigned int supB{};
  unsigned int mCA{};
  unsigned int mSA{};
  unsigned int mCB{};
  unsigned int mSB{};
  unsigned int mapA{};
  unsigned int mapB{};
  unsigned int minParCov{};
  unsigned int bridges[4]{0, 0, 0, 0};
  unsigned int anchorsA[4]{0, 0, 0, 0};
  unsigned int anchorsB[4]{0, 0, 0, 0};
  unsigned int coveragesA[4]{0, 0, 0, 0};
  unsigned int coveragesB[4]{0, 0, 0, 0};
  unsigned int pNP{};
  unsigned int pNB{};
  unsigned int pMed{};
  unsigned int pMax{};
  unsigned int tBC{};
  unsigned int nBC{};
  unsigned int nS{};
};

int main(int argc, char ** argv)  try {
  // Check arguments
  if (--argc < 3)
    throw Error("usage: validate_mumdex cand missing mumdexes...");

  // Input file
  const string cand_file_name{argv[1]};
  ifstream cand_file{cand_file_name.c_str()};
  if (!cand_file) throw Error("Could not open file") << cand_file_name;

  // Read Input file into candidate objects
  string line;
  vector<CandInfo> info;
  while (cand_file) {
    getline(cand_file, line);
    if (line.size()) {
      try {
        CandInfo i{line};
        info.push_back(i);
      } catch (CandInfo::ParseError & e) {
        if (line[0] != 'f' || line[1] != 'a') {
          cerr << "parse error for line: " << line << endl;
        }
      }
    }
  }

  // Load missing sample information
  map<string, string> missing_text{[argv]() {
      ifstream input{argv[2]};
      if (!input) throw Error("Problem opening missing samples file");
      string family;
      string message;
      map<string, string> result;
      while (input >> family) {
        getline(input, message);
        result[family] = message;
      }
      return result;
    }()};

  argc -= 2;
  argv += 2;

  // map<string, vector<unsigned char>> missing_samples{[&missing_text](){ }()};

  // Load mumdexes for family members
  vector<MUMdex> mumdexes;
  mumdexes.reserve(argc);
  while (argc) {
    mumdexes.emplace_back(argv[1]);
    --argc;
    ++argv;
  }
  const Reference & ref{mumdexes.front().reference()};
  const ChromosomeIndexLookup lookup{ref};

  const unsigned int early{152};
  unsigned int n_success{0};
  unsigned int n_good_success{0};
  unsigned int n_good_primers{0};
  unsigned int n_bad_primer{0};

  // Header line
  cout << "family\tkid\tchrA\tposA\thighA\tchrB\tposB\thighB\tinv\toff";
  const vector<string> counts{"cc", "ac", "bc"};
  const string anchor{"AB"};
  const string members{"FMPS"};
  for (const bool ab : {false, true}) {
    for (const unsigned int c : {0, 1, 2}) {
      for (unsigned int m{0}; m != mumdexes.size(); ++m) {
        if (ab || c < 2)
          cout << "\t" << members[m] << counts[c] << anchor[ab];
      }
    }
  }
  cout << "\tbad\tkidok\tfactor\tsuccess\tsuccess_or_bad\tmissing" << endl;;

  for (const CandInfo & i : info) {
    unsigned int max_base_seen{0};
    cout << i.family << "\t" << i.kids;
    const string chrs[2]{i.chrA, i.chrB};
    const unsigned int chr[2]{lookup[i.chrA], lookup[i.chrB]};
    const unsigned int pos[2]{i.posA, i.posB};
    const unsigned int high[2]{i.highA, i.highB};

    for (const bool b : {false, true}) {
      cout << "\t" << chrs[b] << "\t" << pos[b] << "\t" << high[b];
    }
    cout << "\t" << i.invariant << "\t" << i.offset;

    vector<unsigned int> bridge_counts(mumdexes.size());

    // How many mums contain the anchor base
    for (const bool b : {false, true}) {
      // Find counts
      vector<unsigned int> base_counts(mumdexes.size());
      vector<unsigned int> anchor_counts(mumdexes.size());
      for (unsigned int m{0}; m != mumdexes.size(); ++m) {
        const MUMdex & mumdex{mumdexes[m]};

        // Count bridges
        if (b) {
          for (const Bridge & bridge : Bridges{
              chr[0], pos[0], static_cast<bool>(high[0]),
                  chr[1], pos[1], static_cast<bool>(high[1]),
                  i.invariant, mumdex}) {
            if (0) cout << bridge.pair_index();
            ++bridge_counts[m];
          }
        }

        const auto begin_index = mumdex.lower_bound(
            chr[b], pos[b] > early ? pos[b] - early : 0);
        const auto end_index = mumdex.lower_bound(
            chr[b], pos[b] + 1);
        for (auto index = begin_index; index != end_index; ++index) {
          // const auto pair = mumdex.pair(*index);
          const auto mum = mumdex.mum(*index);
          if (pos[b] >= mum.position0() &&
              pos[b] < mum.position0() + mum.length()) {
            ++base_counts[m];
            if ((pos[b] == mum.position0() && !high[b] &&
                 (mum.flipped() ? !mum.touches_end() : mum.offset())) ||
                (pos[b] == mum.position0() + mum.length() - 1 && high[b] &&
                 (mum.flipped() ? mum.offset() : !mum.touches_end()))) {
              ++anchor_counts[m];
            }
          }
        }
      }

      // Output counts
      for (unsigned int m{0}; m != mumdexes.size(); ++m) {
        max_base_seen = max(base_counts[m], max_base_seen);
        cout << "\t" << base_counts[m];
      }
      for (unsigned int m{0}; m != mumdexes.size(); ++m) {
        cout << "\t" << anchor_counts[m];
      }
    }
    const unsigned int max_kids{max(bridge_counts[2], bridge_counts[3])};
    const unsigned int max_parents{max(bridge_counts[0], bridge_counts[1])};
    // const unsigned int max_family{max(max_kids, max_parents)};
    const double factor{max_kids / (max_parents + 1.0)};
    bool correct_kid{false};
    for (unsigned int m{0}; m != mumdexes.size(); ++m) {
      cout << "\t";
      if ((m == 2 && i.kids != "sibling") ||
          (m == 3 && i.kids != "proband")) {
        if (bridge_counts[m] > 10 * max_parents) correct_kid = true;
        // cout << "+";
      }
      cout << bridge_counts[m];
    }
    const bool bad_primer{max_base_seen < 10};
    if (max_base_seen < 10) ++n_bad_primer;
    const bool success{correct_kid && factor > 10};
    if (success) ++n_success;
    bool good_or_bad_success{false};
    if (missing_text[i.family].empty()) {
      ++n_good_primers;
      if (success) {
        ++n_good_success;
        good_or_bad_success = true;
      }
    } else {
      good_or_bad_success = true;
    }

    cout << "\t" << bad_primer << "\t" << correct_kid
         << "\t" << static_cast<unsigned int>(factor)
         << "\t" << success
         << "\t" << good_or_bad_success
         << "\t" << missing_text[i.family] << endl;
  }

  std::cerr << "good success rate " << 1.0 * n_good_success /
      n_good_primers << " " << n_good_success << " " << n_good_primers << endl;
  std::cerr << "success rate " << 1.0 * n_success / info.size() << endl;
  std::cerr << "bad primer rate " << 1.0 * n_bad_primer / info.size() << endl;

  return 0;
} catch (Error & e) {
  cerr << "paa::Error:" << endl;
  cerr << e.what() << endl;
  return 1;
} catch(std::exception & e) {
  cerr << e.what() << '\n';
  return 1;
} catch(...) {
  cerr << "Some exception was caught.\n";
  return 1;
}
```
