## Supplementary material for "G-Graph: An interactive genomic graph viewer": MUMdex Source Code for G-Graph: venn.cpp

```
//
// venn.cpp
//
//
// Copyright 2018 Peter Andrews @ CSHL
//

#include <algorithm>
#include <iostream>
#include <string>
#include <vector>
#include <climits>

#include "error.h"
#include "numerical.h"
#include "utility.h"

using std::cerr;
using std::cin;
using std::cout;
using std::endl;
using std::exception;
using std::string;
using std::vector;

using paa::Error;
using paa::GoldenMinimizer;

class Circle {
 public:
  Circle(
    const double x__,
    const double y__,
    const double r__) :
  x_{x__},
  y_{y__},
  r_{r__} { }

  double x() const { return x_; }
  Circle & x(const double x__) {
    x_ = x__;
    return *this;
  }
  double y() const { return y_; }
  Circle & y(const double y__) {
    y_ = y__;
    return *this;
  }
  double r() const { return r_; }
  Circle & r(const double r__) {
    r_ = r__;
    return *this;
  }

 private:
  double x_;
  double y_;
  double r_;
};

class AreaObjFun {
 public:
  AreaObjFun(
      const double & area_,
      const double & r_,
      const double & R_) :
      area{area_},
    r{r_},
    R{R_} {
      using std::swap;
      if (R < r) swap(r, R);
    }
#if 1
  double operator()(const double d) const {
    const double diff{area -
          (r * r * acos((d * d + r * r - R * R) / (2 * d * r)) +
           R * R * acos((d * d + R * R - r * r) / (2 * d * R)) -
           sqrt((-d + r + R) * (d + r - R) * (d - r + R) * (d + r + R)) / 2)};
    return diff * diff;
  }
#else
  double operator()(const double d) const {
    const double d1{(r * r - R * R + d * d) / (2 * d)};
    const double d2{(R * R - r * r + d * d) / (2 * d)};
    return diff * diff;
  }
#endif

 private:
  double area;
  double r;
  double R;
};

const double pi{atan2(0, -1)};

double area(const double radius) {
  return pi * radius * radius;
}

class Venn {
 public:
  Venn(const double width_, const double height_, const double padding_,
       const double n1_, const double n2_, const double n12_) :
      width{width_},
    height{height_},
    padding{padding_},
    n1{n1_},
    n2{n2_},
    n12{n12_} {
      if (width < height) throw Error("Expect width greater than height");
      if (n1 <= 0 || n2 <= 0 || n12 <= 0)
        throw Error("Venn input numbers may not be zero");
      if (n1 < n2) throw Error("n1 cannot be less than n2");
      const double r1{std::min(height / 2 - padding, width / 4 - padding)};
      const double r2{sqrt(r1 * r1 * n2 / n1)};
      const AreaObjFun obj_fun{pi * r1 * r1 * n12 / n1, r1, r2};
      GoldenMinimizer minimizer(obj_fun, 0, r1 + r2);
      const double d{minimizer.min()};
      const double x_pad{(width - r1 - r2 - d) / 2};
      std::cerr << "pi " << pi
                << " r1 " << r1
                << " r2 " << r2
                << " d " << d
                << " a1 " << area(r1)
                << " a2 " << area(r2)
                << " a12 " << pi * r1 * r1 * n12 / n1
                << std::endl;
      c1 = Circle{x_pad + r1, height / 2, r1};
      c2 = Circle{x_pad + r1 + d, height / 2, r2};
    }

  string ps() const {
    std::ostringstream out;
    out << "%!PS-Adobe-3.0 EPSF-3.0\n"
        << "%%BoundingBox: 0 0 " << width << " " << height << "\n"
        << "2 setlinewidth\n"
        << "newpath " << c1.x() << " " << c1.y() << " " << c1.r()
        << " 0 360 arc gsave 1 0 0 setrgbcolor fill grestore "
        << "gsave stroke grestore gsave clip \n"
        << "newpath " << c2.x() << " " << c2.y() << " " << c2.r()
        << " 0 360 arc gsave 1 0 1 setrgbcolor fill grestore stroke\n";
    return out.str();
  }

 private:
  double width;
  double height;
  double padding;
  double n1;
  double n2;
  double n12;
  Circle c1{0, 0, 0};
  Circle c2{0, 0, 0};
};

int main(int argc, char ** argv) try {
  if (--argc != 3) throw Error("usage: venn n1 n2 n12");

  const double n1{atof(argv[1])};
  const double n2{atof(argv[2])};
  const double n12{atof(argv[3])};

  const Venn venn{792, 612, 20, n1, n2, n12};
  cout << venn.ps() << endl;

  return 0;
} catch (Error & e) {
  cerr << "paa::Error:" << endl;
  cerr << e.what() << endl;
  return 1;
} catch(exception & e) {
  cerr << e.what() << endl;
  return 1;
} catch(...) {
  cerr << "Some exception was caught." << endl;
  return 1;
}
```
