## Supplementary material for "G-Graph: An interactive genomic graph viewer": MUMdex Source Code for G-Graph: wg_smash_comparison.cpp

```
//
// wg_smash_comparison.cpp
//
// Use WG CN as ground truth to assess SMASH segment calls
//
// Copyright 2017 Peter Andrews @ CSHL
//

#include <algorithm>
#include <exception>
#include <fstream>
#include <iostream>
#include <map>
#include <sstream>
#include <string>
#include <utility>
#include <vector>

#include "cn.h"
#include "error.h"
#include "mumdex.h"
#include "psplot.h"
#include "stats.h"
#include "strings.h"

using std::cin;
using std::cout;
using std::cerr;
using std::endl;
using std::exception;
using std::ifstream;
using std::istringstream;
using std::map;
using std::ofstream;
using std::ostringstream;
using std::pair;
using std::string;
using std::vector;

using paa::Bin;
using paa::Bounds;
using paa::ChromosomeIndexLookup;
using paa::CN_abspos;
using paa::CN_Bin;
using paa::CN_Bins;
using paa::Error;
using paa::MAD;
using paa::Marker;
using paa::PSDoc;
using paa::PSGraph;
using paa::PSHSeries;
using paa::PSXYSeries;
using paa::Reference;
using paa::Segment;

int main(int argc, char* argv[]) try {
  if (--argc != 4)
    throw Error("usage: wg_smash_comparison ref bin_file smash_dir wg_calls ");

  // Process arguments
  const Reference ref{argv[1]};
  const ChromosomeIndexLookup lookup{ref};

  const string bins_name{argv[2]};
  const vector<Bin> all_bins{load_bins(bins_name, ref)};
  // List of previously determined good bins
  const vector<Bin> bins{[&all_bins]() {
      vector<Bin> result;
      for (const Bin & bin : all_bins) {
        if (!bin.bad()) {
          result.push_back(bin);
        }
      }
      return result;
    }()};

  const string smash_dir{argv[3]};
  map<string, vector<Segment>> all_segments;

  const string wg_calls_name{argv[4]};
  ifstream wg_calls_file{wg_calls_name.c_str()};
  if (!wg_calls_file)
    throw Error("Problem opening wg calls file") << wg_calls_name;
  string line;
  string chr_name;
  uint64_t start;
  uint64_t stop;
  unsigned int size;
  int state;
  string family;
  string sample;
  string origin;
  string destination;
  unsigned int ngenes;
  string span;
  unsigned int plot_index;
  unsigned int n{0};
  getline(wg_calls_file, line);
  while (getline(wg_calls_file, line)) {
    istringstream line_stream{line.c_str()};
    line_stream >> chr_name >> start >> stop >> size >> state
                >> family >> sample >> origin >> destination
                >> ngenes >> span >> plot_index;
    start /= 10;
    stop /= 10;
    size /= 10;
    const unsigned int chr{lookup[string("chr") + chr_name]};
    if (!line_stream) throw Error("Problem parsing line") << line;
    ++n;

    // Which family members are the event found in
    vector<string> members;
    if (destination == "child")
      members.push_back(family + "-D");
    if (origin == "both" || origin == "father")
      members.push_back(family + "F-D");
    if (origin == "both" || origin == "mother")
      members.push_back(family + "M-D");

    const string expected_state{state > 0 ? "gain" : "loss"};

    if (0)
      cerr << chr_name
           << " " << start
           << " " << stop
           << " " << size
           << " " << expected_state
           << " " << family
           << " " << origin
           << " " << destination
           << " " << members.size() << endl;

    for (const string member : members) {
      // Load segment data if necessary
      if (!all_segments.count(member)) {
        ostringstream segments_name;
        segments_name << smash_dir << "/" << member << "/" << member
                      << "_" << all_bins.size() << "_bins_segments.txt";
        all_segments[member] = Segment::load(segments_name.str(), lookup);
      }

      // See if SMASH segment almost matches up with WG call...
      const vector<Segment> & segments{all_segments[member]};
      unsigned int bases_called_correctly{0};
      for (const Segment & segment : segments) {
        if (segment.chr != chr) continue;
        if (segment.type != expected_state) continue;
        if (0)
          cerr << "Here " << segments.size() << " " << segment.type
               << " " << expected_state
               << " " << chr << " " << segment.chr
               << " " << start << " " << stop
               << " " << segment.startpos << " " << segment.stoppos
               << endl;
        if (segment.startpos > stop) continue;
        if (segment.stoppos <= start) continue;
        if (segment.startpos > start &&
            segment.stoppos <= stop) {
          bases_called_correctly += segment.stoppos - segment.startpos;
          continue;
        }
        if (segment.startpos < start &&
            segment.stoppos >= stop) {
          bases_called_correctly += stop - start;
          continue;
        }
        if (segment.startpos < start && segment.stoppos < stop) {
          bases_called_correctly += segment.stoppos - start;
          continue;
        }
        if (segment.startpos > start && segment.stoppos > stop) {
          bases_called_correctly += stop - segment.startpos;
          continue;
        }
        throw Error("Unexpected segment overlap error");
      }
      cout << segments.size()
           << " " << bases_called_correctly
           << " " << size
           << " " << 1.0 * bases_called_correctly / size
           << " " << member
           << " " << chr_name
           << " " << start
           << " " << stop
           << " " << expected_state
           << " " << plot_index
           << endl;
    }
  }
  cerr << "Read " << n << " lines from " << wg_calls_name << endl;

  return 0;
} catch (Error & e) {
  cerr << "paa::Error:" << endl;
  cerr << e.what() << endl;
  return 1;
} catch (exception & e) {
  cerr << "std::exception" << endl;
  cerr << e.what() << endl;
  return 1;
} catch (...) {
  cerr << "unknown exception was caught" << endl;
  return 1;
}
```
