## Supplementary material for "G-Graph: An interactive genomic graph viewer": MUMdex Source Code for G-Graph: x11plot.cpp

```
//
// x11plot
//
// data exploration
//
// Copyright 2016 Peter Andrews @ CSHL
//

#include <algorithm>
#include <exception>
#include <iostream>
#include <fstream>

#include "error.h"
#include "threads.h"
#include "tsv.h"
#include "x11plot.h"

using std::cout;
using std::cerr;
using std::endl;
using std::exception;
using std::ofstream;

using paa::Error;
using paa::ThreadPool;
using paa::TSV;
using paa::X11Plotter;

int main(int argc, char** argv) try {
  if (--argc != 1) throw Error("usage: x11plot input_file");

  // TSV reads in a tabular text file
  TSV tsv{argv[1]};

  // Thread pool
#ifdef __CYGWIN__
unsigned int n_threads{1};
#else
unsigned int n_threads{std::max(std::thread::hardware_concurrency(), 1U)};
#endif
  ThreadPool pool{n_threads};

  // Create an X11 plotter from TSV
  X11Plotter plotter{tsv, &pool};

  return 0;
} catch (Error & e) {
  cerr << "Error:" << endl;
  cerr << e.what() << endl;
  return 1;
} catch (exception & e) {
  cerr << "exception" << endl;
  cerr << e.what() << endl;
  return 1;
} catch (...) {
  cerr << "unknown exception was caught" << endl;
  return 1;
}
```
